# Identification of Direct-acting nsP2 Helicase Inhibitors with Antialphaviral Activity

**DOI:** 10.1101/2025.03.04.641060

**Authors:** Muthu Ramalingam Bose, John D. Sears, Kacey M. Talbot, Yuan-Wei Norman Su, Scott Houliston, Mohammed Anwar Hossain, Zachary W. Davis-Gilbert, Chunqing Zhao, Hans J. Oh, Peter J. Brown, Marcia K. Sanders, Stella R. Moorman, Durbadal Ojha, Jane E. Burdick, Isabella Law, Noah L. Morales, Sabian A. Martinez, Peter Loppnau, Julia Garcia Perez, Adam M. Drobish, Thomas E. Morrison, Zachary J. Streblow, Daniel N. Streblow, Cheryl H. Arrowsmith, Ava Vargason, Rafael M. Couñago, Levon Halabelian, Jamie J. Arnold, Craig E. Cameron, Nathaniel J. Moorman, Mark T. Heise, Timothy M. Willson

## Abstract

Alphaviruses are mosquito-borne RNA viruses that pose a significant public health threat, with no FDA-approved antiviral therapeutics available. The non-structural protein 2 helicase (nsP2hel) is an enzyme involved in unwinding dsRNA essential for alphavirus replication. This study reports the discovery and optimization of first-in-class oxaspiropiperidine inhibitors targeting nsP2hel. Structure-activity relationship (SAR) studies identified potent cyclic sulfonamide analogs with nanomolar antiviral activity against chikungunya virus (CHIKV). Biochemical analyses of nsP2hel ATPase and RNA unwindase activities showed these compounds act by a non-competitive mode suggesting that they are allosteric inhibitors. Viral resistance mutations mapped to nsP2hel and a fluorine-labeled analog exhibited direct binding to the protein by ^19^F NMR. The lead inhibitor, **2o**, demonstrated broad-spectrum antialphaviral activity, reducing titers of CHIKV, Mayaro virus (MAYV), and Venezuelan equine encephalitis virus (VEEV). These findings support nsP2hel as a viable target for development of broad-spectrum direct-acting antialphaviral drugs.

**Graphical Abstract:** 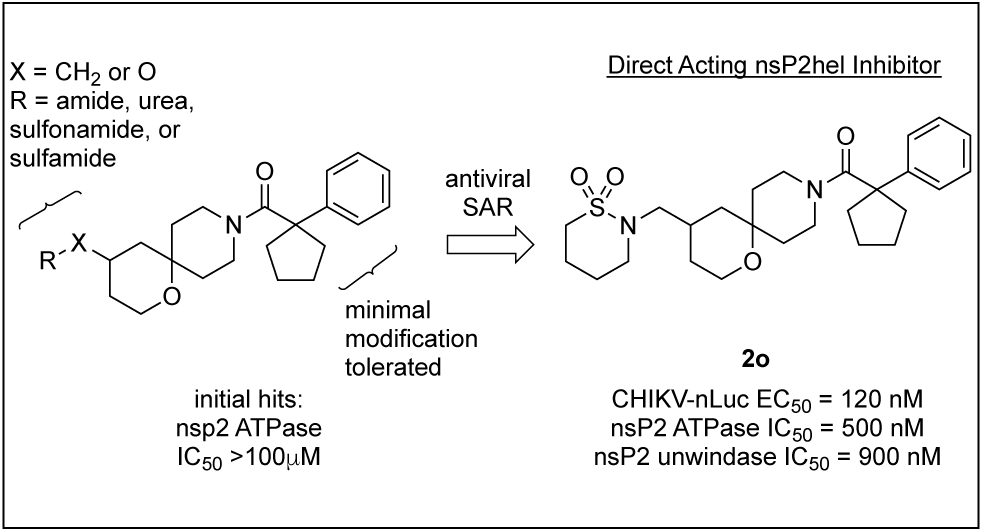

## Introduction

Alphaviruses are mosquito-borne RNA viruses that pose a significant public health risk both within the United States and globally.^1^ Alphaviruses such as chikungunya virus (CHIKV), Ross River virus (RRV), and O’nyong-nyong virus (ONNV) have caused large scale epidemics of debilitating acute and chronic arthralgia. Likewise, Venezuelan (VEEV), Western (WEEV) and Eastern (EEEV) Equine Encephalitis viruses trigger neurologic disease with high rates of morbidity and mortality^2^ with a recent outbreak of EEEV in New England resulting in issuance of multiple public health alerts.^3^ The threat posed by these viruses is likely to increase as global warming expands the footprint of their mosquito vectors. Moreover, viruses such as VEEV and EEEV are also considered potential bioterror threats due to their capacity for weaponization. Despite the public health danger posed by these viruses, there are currently no FDA-approved drugs for any alphavirus-caused disease. Broadly acting antiviral therapeutics are badly needed for protection against both existing and future alphavirus threats.

The highly conserved alphavirus non-structural protein 2 (nsP2) is a promising target for direct-acting antiviral drugs as it mediates a variety of processes that are essential for viral RNA synthesis and virulence.^4^ One of the main functions of nsP2 is RNA unwindase activity that is primarily mediated by an N-terminal helicase domain (nsP2hel) and is an essential step in viral RNA synthesis.^4^ nsP2hel is classified as an SF1 helicase by sequence homology of the conserved Walker A/B Motifs and RNA binding domains.^5^ ATP is a cofactor for the nsP2hel RNA unwindase with energy for separation of the bases derived from hydrolysis to ADP. As a result, nsP2hel also functions as an ATPase.^6^ However, a clear understanding of the biochemistry of nsp2hel as a helicase motor enzyme and its precise functions in alphavirus replication are incomplete at a molecular and cellular level. Although viral RNA helicases are compelling targets for development of direct-acting antiviral drugs,^7^ the dynamic nature of these motor enzymes has made it difficult to identify, characterize, and optimize small molecule inhibitors using conventional target-based approaches to drug discovery.^8^ Herpes simplex virus (HSV) helicase inhibitors have been successfully optimized into clinical candidates using cell-based antiviral assays as the primary drivers for development of structure-activity relationships,^9, 10^ but remarkably there have been no biophysical or structural studies that reveal the details of the molecular interactions of these inhibitors with the helicase protein. Analysis of the X-ray structures of mammalian helicase inhibitors has revealed that the majority of them function by allosteric mechanisms, although there are some rare examples of DNA/RNA competitive or ATP competitive inhibitors.^8^ The dearth of direct-acting helicase inhibitors has stifled research on the molecular mechanism of action of these motor enzymes and their role in human diseases and viral replication.

As part of the program to develop new direct-acting antialphaviral drugs 1-oxa-9-azaspiro[5,5]undecanes (oxaspiropiperidines) **1a** and **2a** (Figure 1) were recently identified as weak inhibitors of the ATPase activity of CHIKV nsP2.^11^ In this report we describe the optimization of these oxaspiropiperidines into a series of potent nsP2 ATPase inhibitors with antialphaviral activity and demonstrate that they are direct-acting allosteric inhibitors of the nsP2hel motor enzyme.

**Figure 1.**
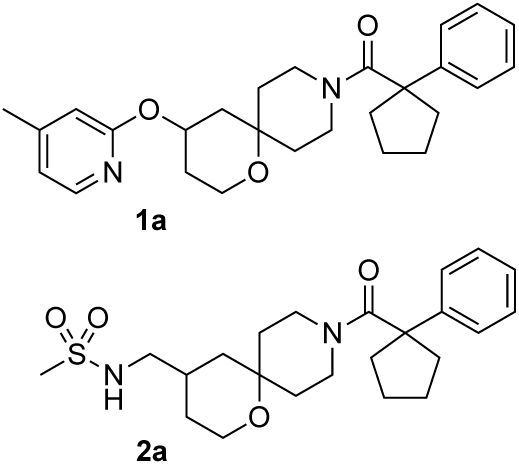
Oxaspiropiperidine nsP2 ATPase inhibitors

## Results and Discussion

### Scheme 1. Synthesis of 1a and analogs (1b–d)*^a^*

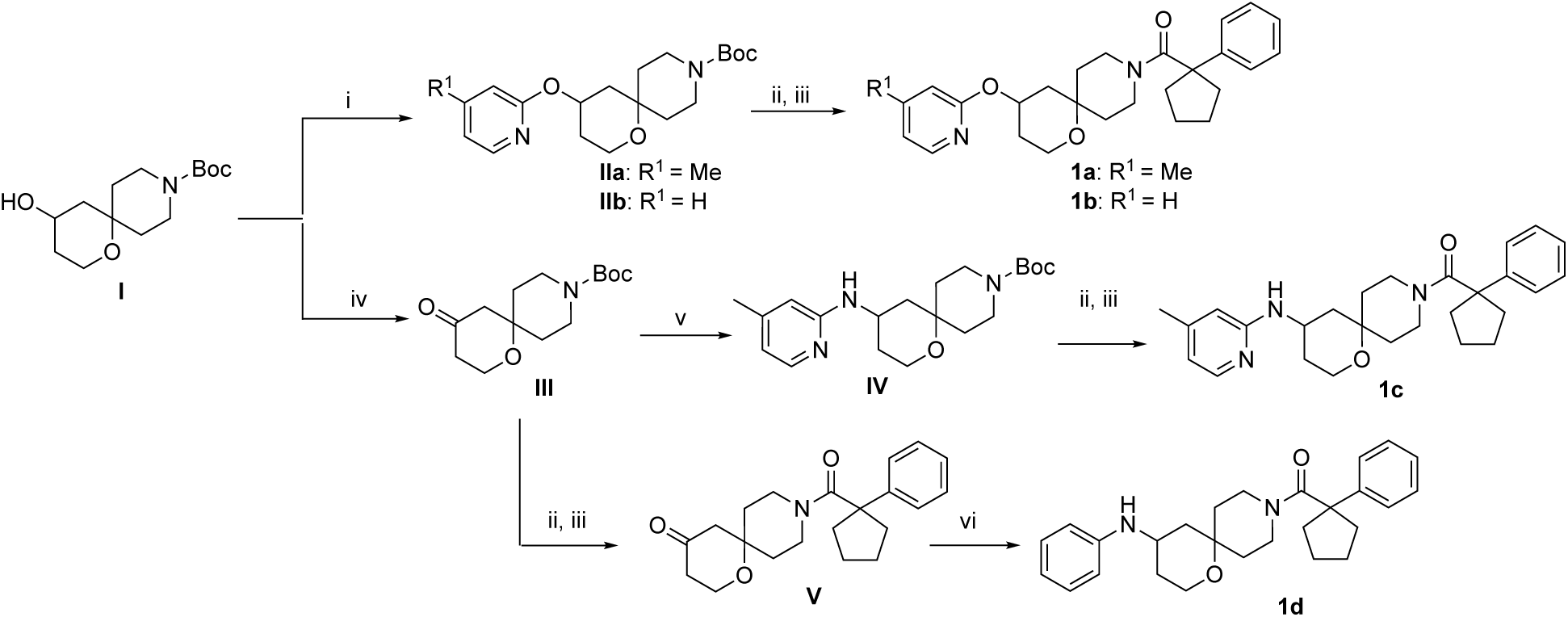

*^a^*Reagents and conditions: (i) KO*^t^*Bu, DMF, 50 °C, 1 h; then 4-methyl-2-chloropyridine or 2-chloropyridine, 85 °C, 4 h; (ii) TFA, DCM; (iii) 1-phenyl-1-cyclopentanecarboxylic acid, HATU, DIPEA, DMF; (iv) PDC, DCM; (v) 2-aminopyridine, Ti(O^i^Pr)_4_, THF, 60 °C; then NaBH_4_, rt; (vi) aniline, NaBH(OAc)_3_, acetic acid, toluene, 140 °C, microwave.

### Synthetic Chemistry

Intermediate **I** containing the oxaspiropiperidine core found in **1a**–**d** was synthesized following literature procedure.^12^ Deprotonation of **I** with KO*^t^*Bu and S_N_Ar reaction with a 2-chloropyridine furnished analogs **IIa** and **IIb** (Scheme 1). Removal of the Boc group, followed by an amide coupling reaction with 1-phenyl-1-cyclopentanecarboxylic acid yielded 2-alkoxypyridines **1a** and **1b**. Oxidation of **I** by PDC furnished ketone **III** which underwent reductive amination with 2-aminopyridine to yield intermediate **IV**. Boc deprotection followed by amide coupling with 1-phenyl-1-cyclopentanecarboxylic acid yielded 2-aminopyridine **1c**. Boc deprotection of **III** followed by amide coupling with 1-phenyl-1-cyclopentanecarboxylic acid furnished intermediate **V** which upon reductive amination under microwave conditions yielded aniline **1d**.

#### Scheme 2. Synthesis of 2a and sulfonamide (2b–h, 5a–g, 6a–m, 7a-c), amide (3a–l), and urea (3p–u) analogs*^a^*

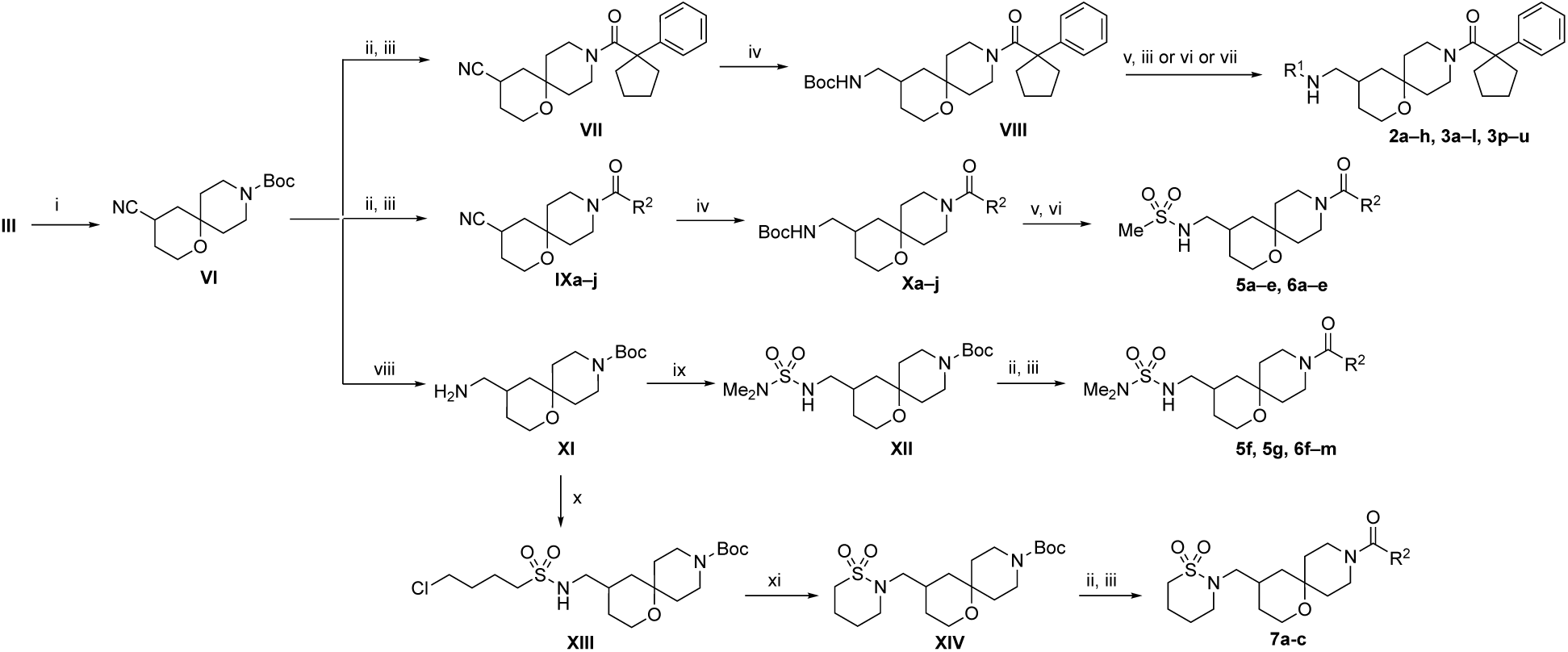

*^a^*Reagents and conditions: (i) TOSMIC, KO*^t^*Bu, DME; (ii) TFA, DCM; (iii) R^2^-COOH, HATU, DIPEA, DMF or TBTU, pyridine; (iv) NiCl_2_, NaBH_4_, Boc_2_O, MeOH; (v) HCl in dioxane; (vi) sulfonyl chloride, Et_3_N, DCM; (vii) isocyanate, Et_3_N, DCM; (viii) MeOH, aq. NH_3_, Ra-Ni, 60 °C (ix) *N*,*N*-dimethylsulfomyl chloride, Et_3_N, DCM; (xi) 4-chloro-1-butylsulfonyl chloride, Et_3_N, DCM; (xi) Cs_2_CO_3_, DMF, 50 °C.

Synthesis of aminomethyl analogs of the oxaspiropiperidine utilized a Van Leusen^13^ reaction of intermediate **III** with TosMIC to generate nitrile intermediate **VI** (Scheme 2). Boc deprotection of **VI** followed by amide coupling with 1-phenyl-1-cyclopentanecarboxylic acid provided intermediate **VII**. Reduction of the nitrile using nickel boride in the presence of Boc anhydride furnished the Boc-protected amine **VIII**. Boc deprotection followed by sulfonylation, amidation, or treatment with an isocyanate yielded the sulfonamide analogs **2a**–**h**, amide analogs **3a**–**l**, and urea analogs **3p**–**u**, respectively. Deprotection of intermediate **VI** followed by amide coupling with a range of carboxylic acids provided intermediates **IXa**– **j**, which underwent nitrile reduction using nickel boride in the presence of Boc anhydride to afford intermediates **Xa**–**j**. Boc deprotection followed by reaction with methanesulfonyl chloride furnished sulfonamide analogs **5a**–**e** and **6a**–**e**. Reduction of the nitrile in **VI** to a primary amine using Raney nickel furnished intermediate **XI** which was treated with *N*,*N*-dimethylsulfonyl chloride, Boc deprotected, and coupled with a range of carboxylic acids to yield *N*,*N*-dimethylsulfamate analogs **5f–g** and **6f**–**m**. Alternatively, reaction of primary amine **XI** with 4-chlorobutylsulfonyl chloride furnished intermediate **XIII** which underwent intramolecular cyclization in the presence of Cs_2_CO_3_ to generate intermediate **XIV**. Boc deprotection of **XIV** followed by amide coupling with three fluorophenyl-substituted carboxylic acids yielded **7a**–**c**.

#### Scheme 3. Synthesis of tertiary sulfonamides (2i–m) and amide (3m)*^a^*

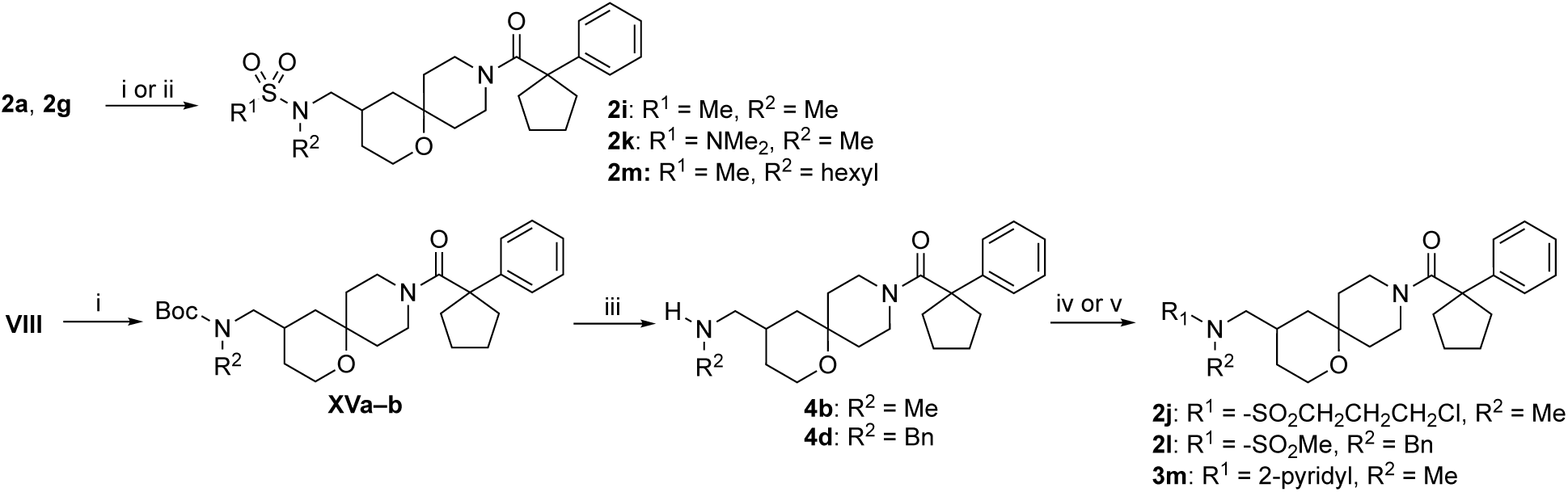

*^a^*Reagents and conditions: (i) NaH, MeI or BnBr, DMF; (ii) K_2_CO_3_, 1-iodohexane, DMF, 85 °C; (iii) 4M HCl in dioxane; (iv) 3-chloropropylsulfonyl chloride or MeSO_2_Cl, Et_3_N, DCM; (v) pyridine-2-carboxylic acid, HATU, DIPEA, DMF.

*N*-Alkylation of sulfonamides **2a** and **2g** with iodomethane or iodohexane afforded the tertiary sulfonamide **2i**, **2k**, and **2m** (Scheme 3). *N*-Methylation of the Boc-protected intermediate **VIII** followed by Boc deprotection afforded secondary amine analogs **4b** and **4d**. Coupling of **4b** with 3-chloropropylsulfonyl chloride, methane sulfonyl chloride, or pyridine-2-carboxylic acid yielded the tertiary sulfonamides **2j** and **2l** and tertiary amide **3m**.

#### Scheme 4. Synthesis of chloroalkyl sulfonamides and amide (2c, 2d, 3e), cyclic sulfonamides (2n–2o), cyclic amides (3n–o), and secondary amine (4c)*^a^*

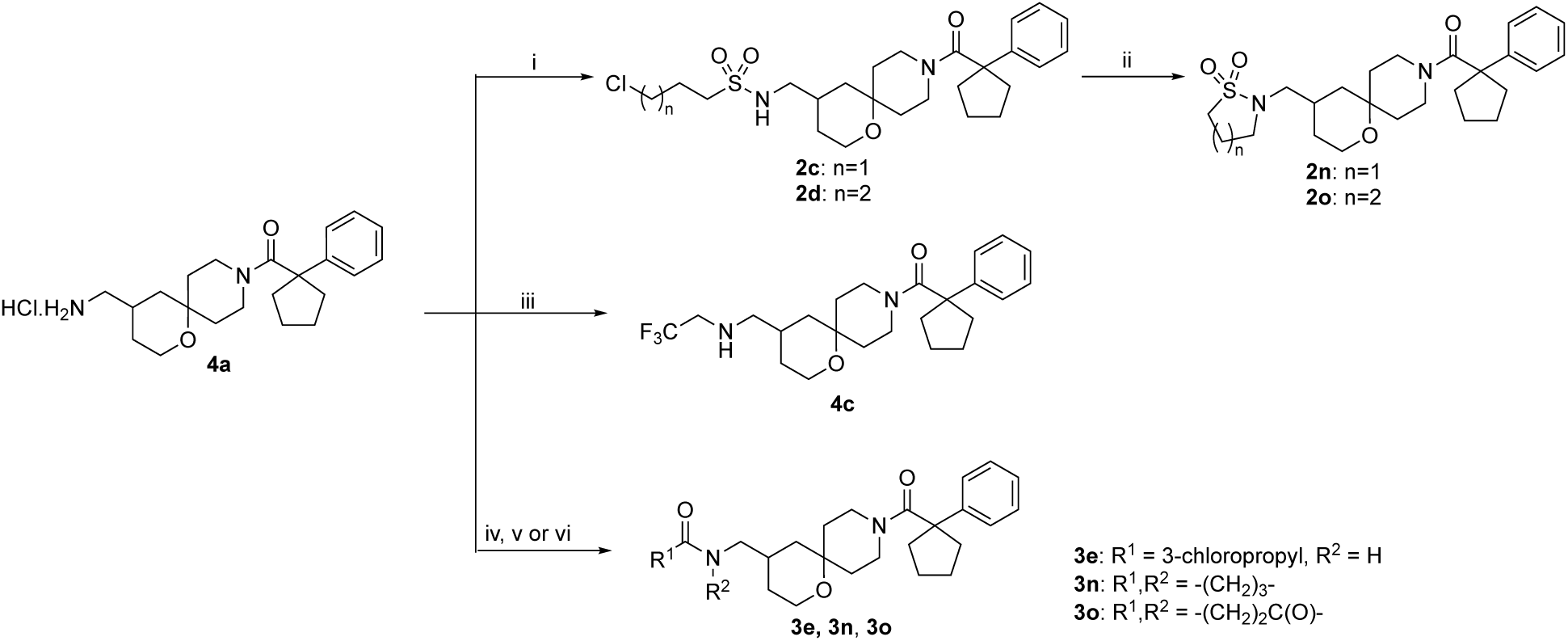

*^a^*Reagents and conditions: (i) sulfonyl chloride, Et_3_N, DCM; (ii) Cs_2_CO_3_, DMF, 50 °C; (iii) 2,2,2-trifluoroethyl trifluoromethylsulfonate, DIPEA, 50 °C, dioxane; (iv) 3-chloropropanoyl chloride, Et_3_N, DCM; (v) NaH, THF, rt; (vi) Succinimide, DMAP, acetic acid, 140 °C.

The cyclic sulfonamides **2n** and **2o**, secondary amine **4c**, secondary amide **3e**, and cyclic amides **3n** and **3o** were synthesized from the HCl salt of primary amine **4a (**Scheme 4). Reaction of **4a** with *n*-chloroalkylsulfonyl chlorides produced **2c** and **2d**. Intramolecular cyclization of **2c** and **2d** under basic conditions furnished the respective cyclic sulfonamides **2n** and **2o**. Alkylation of **4a** with 2,2,2-trifluoroethyl trifluoromethylsulfonate afforded secondary amine **4c**. Reaction of **4a** with 3-chloropropanoyl chloride produced **3e** which underwent intramolecular cyclization under basic conditions to generate the cyclic amide **3n**. Reaction of **4a** with succinic anhydride in acetic acid under reflux furnished the succinimide analog **3o**.

#### Scheme 5. Synthesis of tertiary alcohol 4e*^a^*

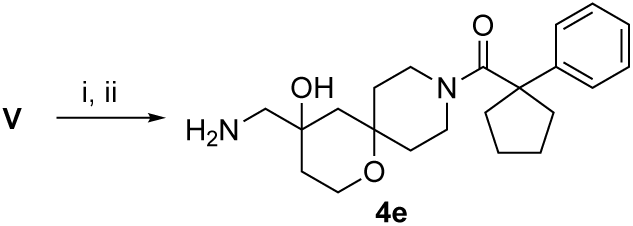

*^a^*Reagents and conditions: (i) TMSCN, Cu(OTf)_2,_ ACN ; (ii) LiAlH_4_, THF

Synthesis of analog **4e** containing a tertiary alcohol on the spirocyclic tetrahydropyran ring used a copper-catalyzed addition of trimethylsilyl cyanide to ketone **V** followed by chemoselective reduction of the nitrile using lithium aluminum hydride at 0 °C for 1h to generate aminoalcohol **4e**.

Spirocyclic compounds containing 1,1-disubstituted cyclopentyl amides displayed complex ^13^C NMR spectra. Many resonances corresponding to the spiropiperidine and the cyclopentane carbons either were not observed, were very broad in CDCl₃, DMSO-d_6_, CD₃OD, and acetone-d_6_, or were partially hidden by the solvent (Figure S1). Several of these aliphatic ^13^C NMR signals were not obscured by the solvent in pyridine-d_5_ and CD₃CN (Figure S2); however, the spiropiperidine carbon signals remained broad and were difficult to assign. The line broadening can be attributed to molecular rotation and transverse relaxation (T₂) on the NMR time scale.^14^ To fully assign the ^13^C spectra, variable temperature NMR was utilized. For a representative case, comparative spectra of **2n** at 0, 25, and 60 °C in pyridine-d_5_ is shown in Figure 2. At 25 °C, many of the resonances were broad and those corresponding to the spiropiperidine carbons (P) could not be assigned. At 60 °C, several of the carbon resonances were sharper, including those corresponding to C4 on the spirocycle, C7 on the cyclic sulfonamide, and C13/14 on the cyclopentane. However, signals for the spiropiperidine carbons in the 38–44 ppm region were notably absent. At 0 °C, multiple signals were observed in the 38–44 ppm region including resonances for all spiropiperidine carbons (P) of **2n**. However, several of the carbon atoms exhibited multiple resonances, further supporting the existence of multiple conformations on the NMR time scale. Additional assignment of carbon resonances utilized ^1^H-^13^C two-dimensional NMR studies that were conducted at 60 °C (HSQC Figure S3 and HMBC Figure S4) and at 0 °C (HSQC Figure S5) in pyridine-d_5_. Based on these 2D data, the cyclopentane carbons C_12_/C_15_ (Figure 2) were assigned as the broad singlets between 39–40 ppm, which were more clearly resolved at 0 °C in CD₃CN (Figure S6). Using this combination of variable temperature ^13^C and 2D NMR in multiple solvents all 21 carbon resonances of **2n** could be observed, although specific assignment of the individual spiropiperidine carbons (P) was not possible (Figure 2).

**Figure 2.**
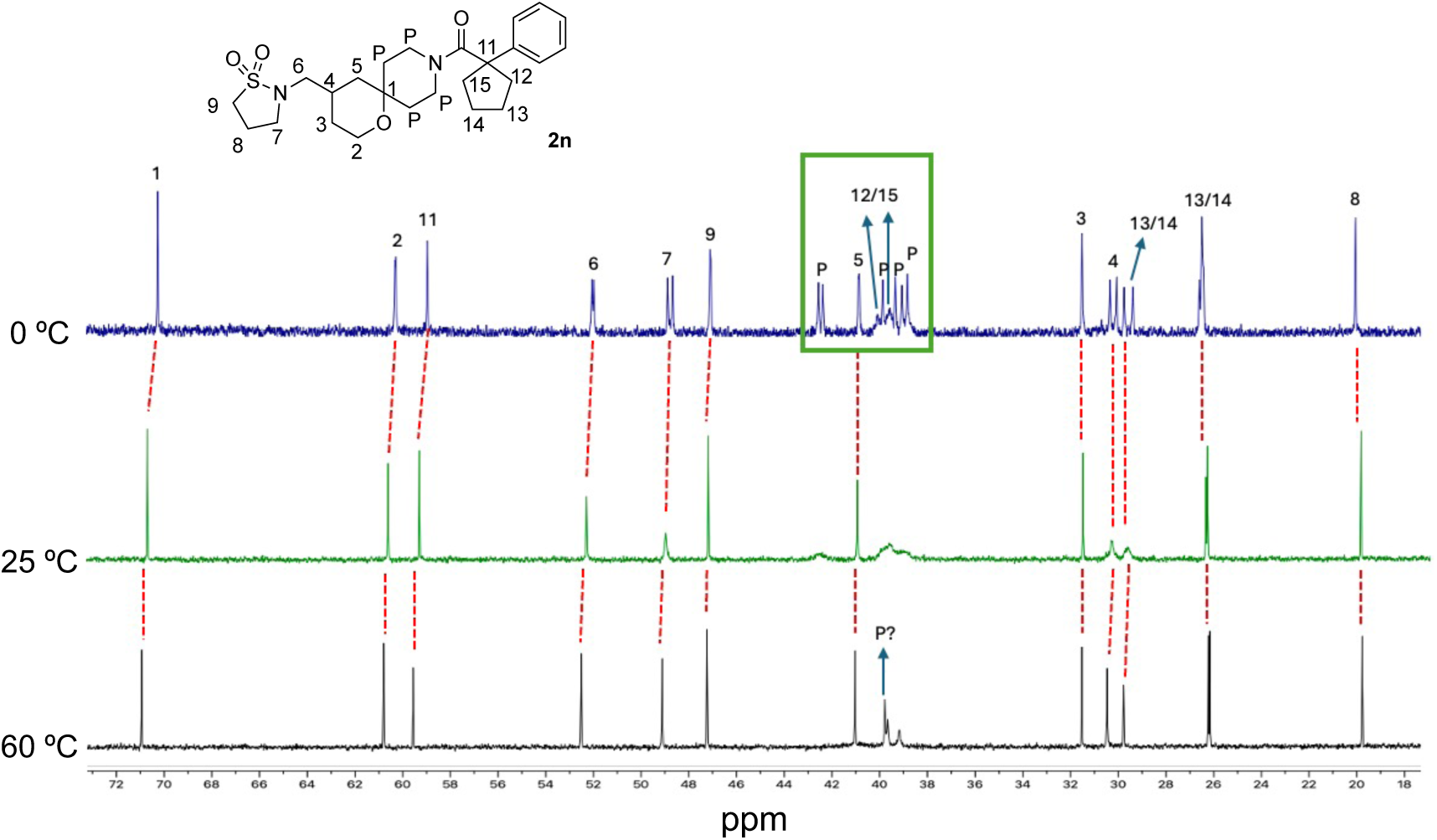
Variable temperature ^13^C NMR of **2n** in pyridine-d_5_ showing the resonances of the sp^3^ carbons from 18–72 ppm. Numbers above the resonances in the spectra correspond to the carbons labelled in the chemical structure. Individual assignment of the piperidine carbons (P) was not possible. Dashed red lines indicate the respective resonances at the variable temperatures.

### Biological Evaluation

Measurement of antiviral activity employed a CHIKV-nLuc reporter assay using an attenuated CHIKV vaccine strain (181/25)^15^ with an nLuc reporter inserted downstream of the capsid gene.^16^ Assays were performed in human fibroblast MRC5 cells at 6 h post inoculation with the virus and luciferase activity used as a measure of CHIKV replication and polyprotein expression. In the CHIKV-nLuc assay the known alphavirus polymerase inhibitor NHC^17^ had an EC_50_ = 1.0 μM.^16^ Secondary screening for breadth was performed using a VEEV-nLuc reporter. nsP2 ATPase inhibition was determined with a multistep ADP-glo assay as previously described for the initial hit characterization.^11^

Initial hits **1a** and **2a** demonstrated antiviral activity with micromolar potency in the CHIKV-nLuc assay (Table 1), but showed weaker ATPase inhibition (IC_50_ >100 μM) than was recorded in the original report.^11^ **1a** and **2a** were 10-fold less active in the VEEV-nLuc assay (Table S1). Given the weak ATPase activity of **1a** and **2a** and concerns that characterization of mammalian helicase inhibitors using biochemical assays was challenging due to assay artifacts caused by compound aggregation or protein interference,^18,19^ we opted to use data from the CHIKV viral replication assay to drive the initial optimization of the screening hits. However, activity of new analogs was also collected in the CHIKV nsP2 ATPase assay in parallel to allow cross-correlations to be made between the antiviral and biochemical measures of helicase inhibition as potency of the inhibitors improved. This approach of using an antiviral assay to develop the primary SAR and drive a medicinal chemistry campaign had been successful in the development of HSV helicase inhibitors.^9,10^

**Table 1.**
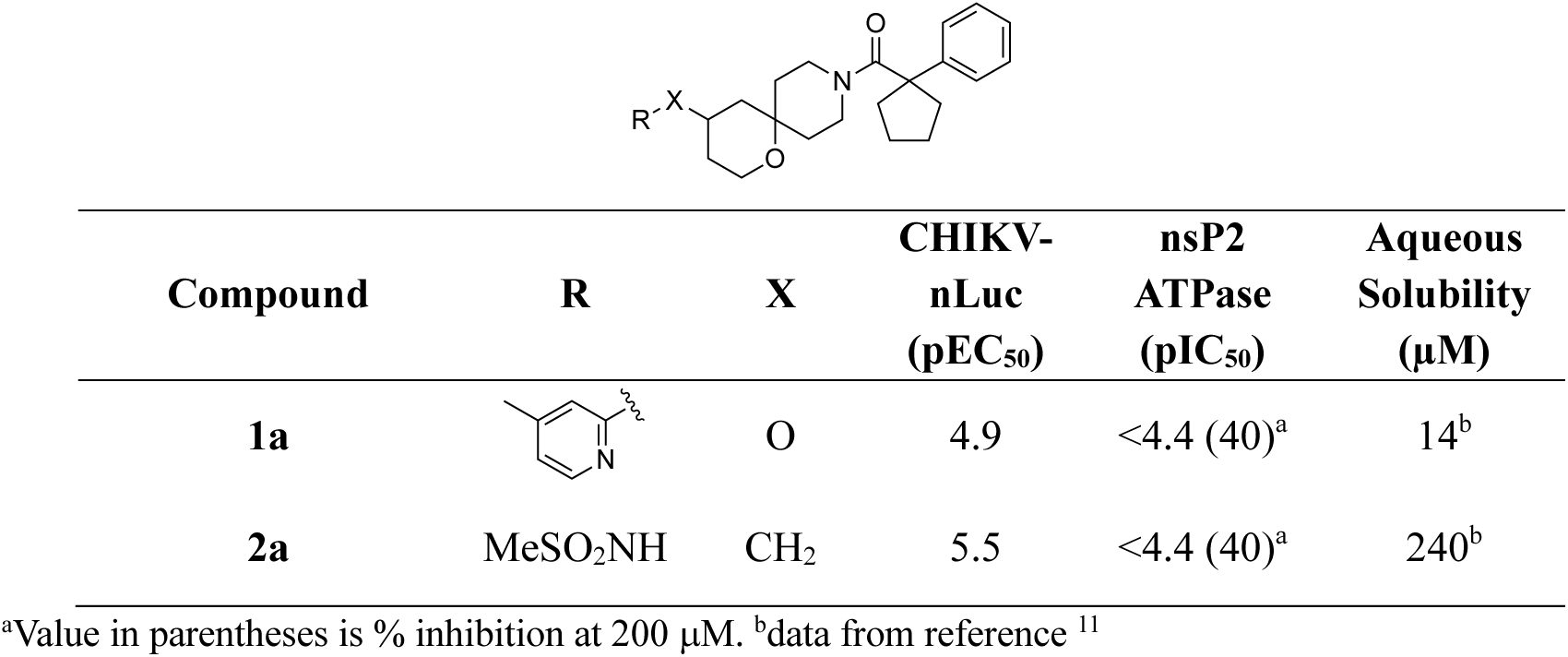
nsP2 ATPase Inhibitors from High Throughput Screening.

| Compound | R | X | CHIKV-nLuc<br>(pEC<sub>50</sub>) | nsP2<br>ATPase<br>(pIC<sub>50</sub>) | Aqueous<br>Solubility<br>(μM) |
| --- | --- | --- | --- | --- | --- |
| **1a** |  | O | 4.9 | <4.4 (40)<sup>a</sup> | 14<sup>b</sup> |
| **2a** | MeSO<sub>2</sub>NH | CH<sub>2</sub> | 5.5 | <4.4 (40)<sup>a</sup> | 240<sup>b</sup> |

<sup>a</sup>Value in parentheses is % inhibition at 200 μM. <sup>b</sup>data from reference <sup>11</sup>

**Table 2.**
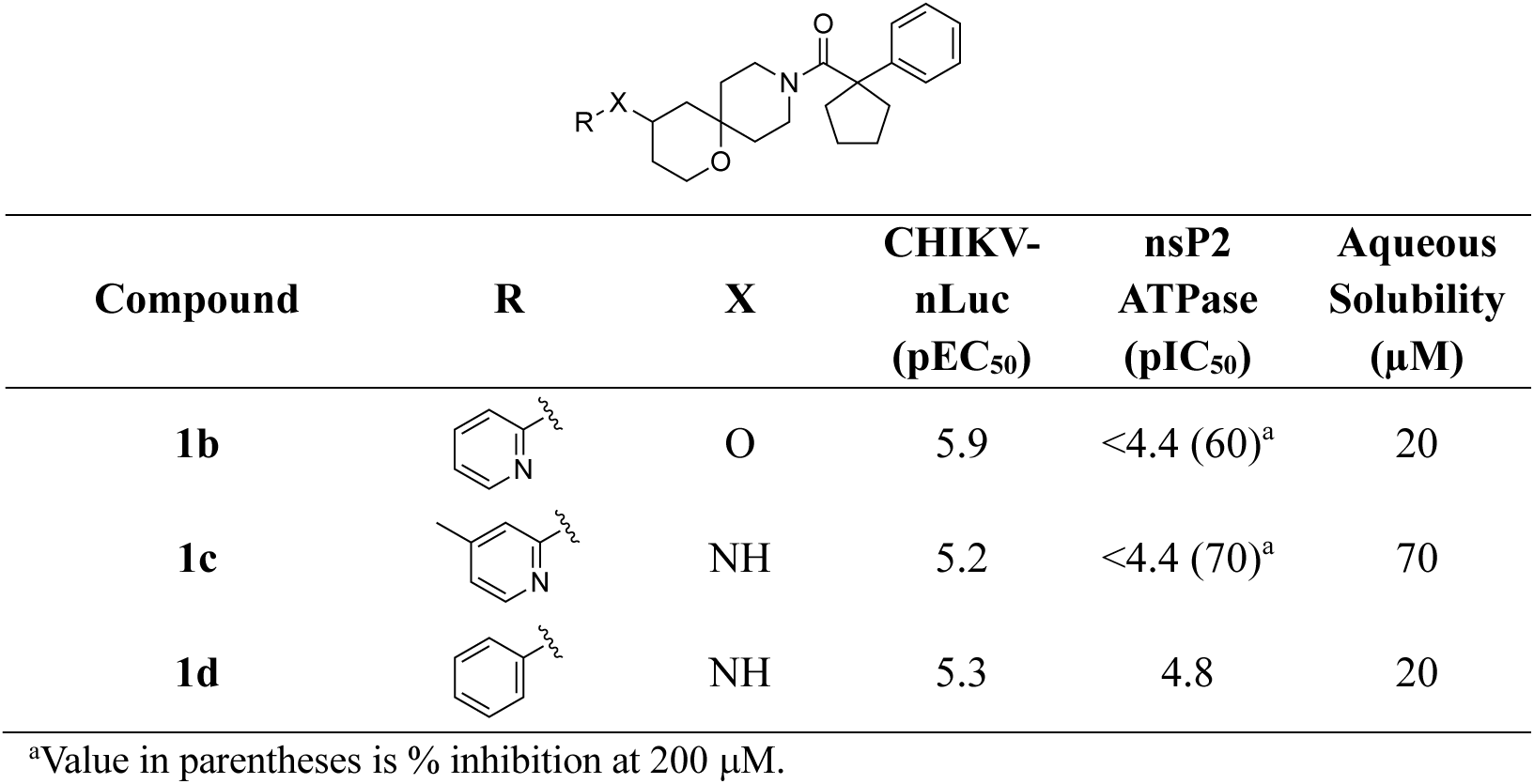
Antialphaviral Activity and Solubility of 1b–d.

| Compound | R | X | CHIKV-nLuc<br>(pEC<sub>50</sub>) | nsP2<br>ATPase<br>(pIC<sub>50</sub>) | Aqueous<br>Solubility<br>(μM) |
| --- | --- | --- | --- | --- | --- |
| **1b** |  | O | 5.9 | <4.4 (60)<sup>a</sup> | 20 |
| **1c** |  | NH | 5.2 | <4.4 (70)<sup>a</sup> | 70 |
| **1d** |  | NH | 5.3 | 4.8 | 20 |

<sup>a</sup>Value in parentheses is % inhibition at 200 μM.

The SAR campaign began with the exploration of the contribution of the pyridyl ether group to the activity of **1a**. Three analogs were synthesized with modifications to either the ether linkage or the pyridine ring. The analog **1b**, lacking the 4-methyl substitution on the 2-pyridyl ring showed improved antiviral activity against CHIKV with EC_50_ ∼ 1 μM. The aminopyridine analog (**1c**) and the simple aniline analog (**1d**) showed slightly lower activity against CHIKV. In contrast to their antiviral activity, analogs **1b–d** only showed robust ATPase inhibition at concentrations >30 μM. Since spirocycles **1a-d** had only low to moderate aqueous solubility, our attention turned to the methyl sulfonamide analog (**2a**) which was more active in the CHIKV antiviral assay and had a better solubility profile.

To explore SAR of **2a**, a series of secondary and tertiary sulfonamides were synthesized (Table 3). Within the secondary alkyl sulfonamides, the ethyl analog (**2b**) had antiviral activity equivalent to **2a**, but the chloropropyl (**2c**) and chlorobutyl (**2d**) analogs were more active with CHIKV EC_50_ ∼ 1 μM. The cyclopropyl (**2e**) and benzene (**2f**) sulfonamides had similar antiviral activity to **2a**, but the benzene sulfonamide (**2f**) demonstrated low aqueous solubility. The secondary sulfamide analogs **2g** and **2h** showed similar antiviral activity to methyl sulfonamide (**2a**). Switching to the corresponding *N*-methyl tertiary sulfonamides (**2i–j**) and sulfamide (**2k**) led to a small increase in CHIKV inhibitory activity. Although the *N*-methyl chloropropyl analog (**2j**) exhibited an 8-fold increase in antiviral activity compared to **2a** it had only moderate aqueous solubility. The *N*-benzyl tertiary sulfonamide (**2l**) was less active against CHIKV and had very poor aqueous solubility. The *n*-hexyl tertiary sulfonamide (**2m**), although demonstrating equivalent antiviral activity to the secondary sulfonamide **2a**, also had very poor aqueous solubility. Intriguingly, the five-membered cyclic tertiary sulfonamide (**2n**) exhibited an increase in CHIKV inhibitory activity compared to the methyl sulfonamide (**2a**). The six-membered cyclic sulfonamide (**2n**) demonstrated an even greater improvement with EC_50_ = 130 nM, a 25-fold increase in antiviral activity compared to **2a**. Notably the analogs **2j** and **2o** with the best antiviral activity also showed strong ATPase inhibition with IC_50_ ≤ 1 μM.

**Table 3.**
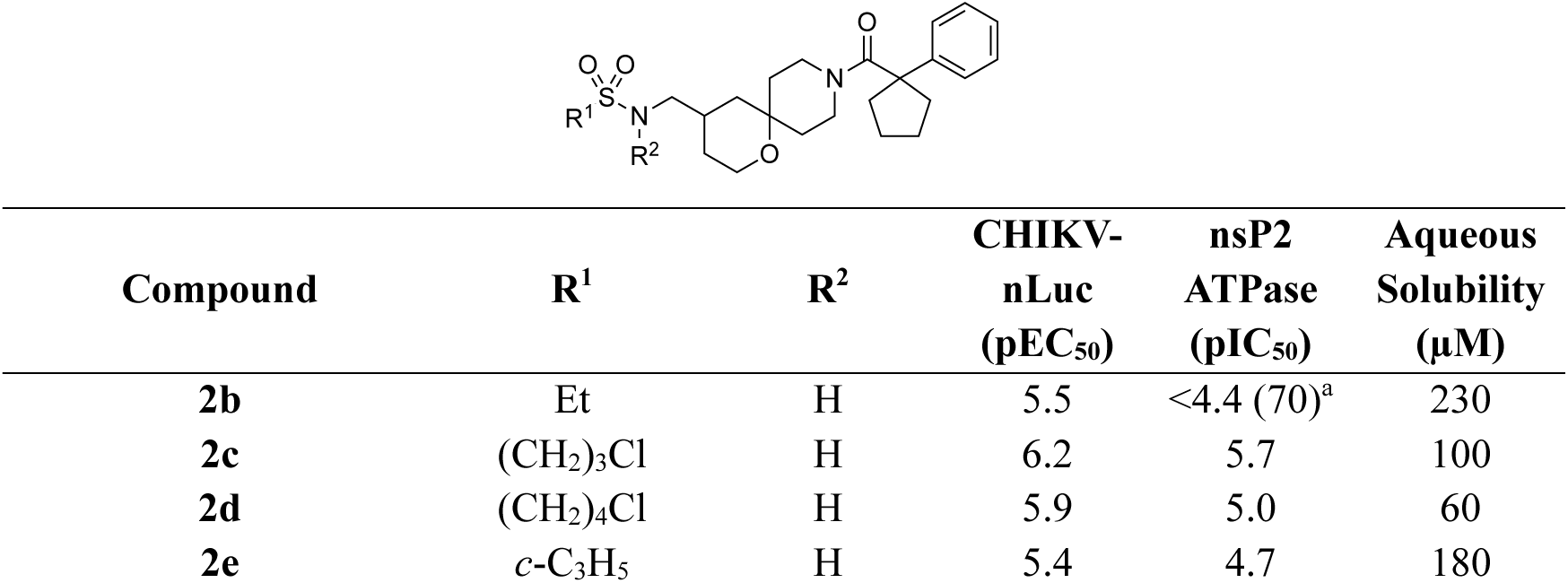
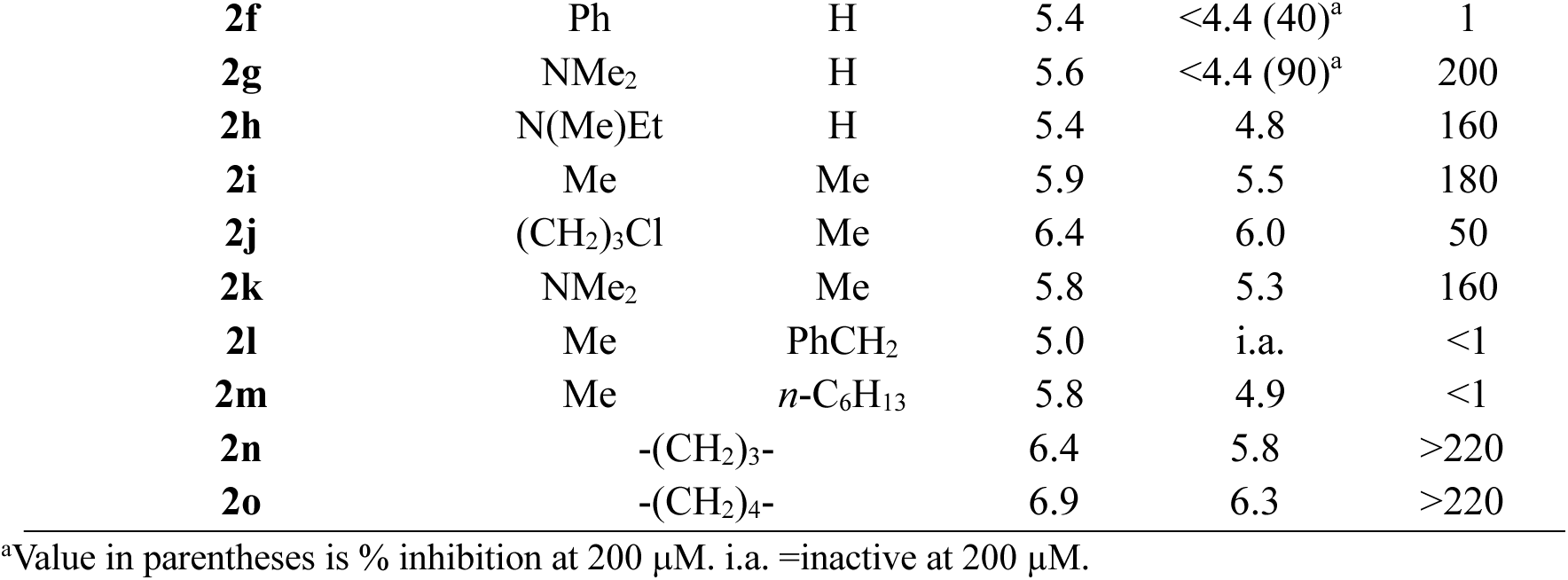
Antialphaviral Activity and Solubility of Sulfonamide and Sulfamides 2b–n.

| Compound | R<sup>1</sup> | R<sup>2</sup> | CHIKV-nLuc<br>(pEC<sub>50</sub>) | nsP2<br>ATPase<br>(pIC<sub>50</sub>) | Aqueous<br>Solubility<br>(μM) |
| --- | --- | --- | --- | --- | --- |
| **2b** | Et | H | 5.5 | <4.4 (70)<sup>a</sup> | 230 |
| **2c** | (CH<sub>2</sub>)<sub>3</sub>Cl | H | 6.2 | 5.7 | 100 |
| **2d** | (CH<sub>2</sub>)<sub>4</sub>Cl | H | 5.9 | 5.0 | 60 |
| **2e** | *c*-C<sub>3</sub>H<sub>5</sub> | H | 5.4 | 4.7 | 180 |
| **2f** | Ph | H | 5.4 | <4.4 (40)<sup>a</sup> | 1 |
| **2g** | NMe<sub>2</sub> | H | 5.6 | <4.4 (90)<sup>a</sup> | 200 |
| **2h** | N(Me)Et | H | 5.4 | 4.8 | 160 |
| **2i** | Me | Me | 5.9 | 5.5 | 180 |
| **2j** | (CH<sub>2</sub>)<sub>3</sub>Cl | Me | 6.4 | 6.0 | 50 |
| **2k** | NMe<sub>2</sub> | Me | 5.8 | 5.3 | 160 |
| **2l** | Me | PhCH<sub>2</sub> | 5.0 | i.a. | <1 |
| **2m** | Me | *n*-C<sub>6</sub>H<sub>13</sub> | 5.8 | 4.9 | <1 |
| **2n** | -(CH<sub>2</sub>)<sub>3</sub>- |  | 6.4 | 5.8 | >220 |
| **2o** | -(CH<sub>2</sub>)<sub>4</sub>- |  | 6.9 | 6.3 | >220 |

<sup>a</sup>Value in parentheses is % inhibition at 200 μM. i.a. =inactive at 200 μM.

These results demonstrated that introduction of an *N*-methyl group onto the initial hit **2a** enhanced antiviral potency against CHIKV. However, further increasing the size of the tertiary nitrogen substituent (**2l** and **2m**) was detrimental to antiviral activity and aqueous solubility. In addition, the linear chloropropyl analogs **2c** and **2j** sulfonamides had improved antiviral activity. Gratifyingly, the cyclic tertiary sulfonamides (**2n–o**) exhibited improved potency compared to their linear counterparts and maintained good solubility profiles.

Structure-activity across a series of amide and urea analogs was also explored (Table 4). The isopropyl (**3a**) amide maintained antiviral activity against CHIKV, but the *n*-pentyl amide (**3b**) showed improved activity. The cyclopropyl (**3c**) and cyclohexyl (**3d**) amides also retained antiviral activity. Switching to the chloropropyl amide (**3e**) resulted in an increase in CHIKV inhibitory activity, similar to the effect that was seen in the sulfonamide series (Table 3). The benzamide (**3f**) had equivalent activity to the cyclohexyl amide (**3d**), showing that aromaticity conferred no advantage over a hydrophobic alkyl substituent. Addition of methoxy substituents to the benzamide (**3g**–**h**) did not affect antiviral activity but a larger *p*-alkoxy substituted benzamide (**3i**) was inactive at concentrations <30 μM. Three isomeric pyridyl amides (**3j**–**l**) were tested. All three were active against CHIKV, but the 2-pyridyl analog (**3j**) had the best combination of antiviral activity and aqueous solubility. The 3-pyridyl analog (**3k**) showed lower solubility, while the 4-pyridyl analog (**3l**) had weaker antiviral activity. Overall, secondary amides showed comparable antiviral activity to their corresponding sulfonamides suggesting that they functioned as isosteric replacements. In two examples (**3e** and **3f**) the amides were more soluble than the corresponding sulfonamide analogs (**2c** and **2f**). To explore whether tertiary substitution in the amide series would also improve activity the *N*-methylated 2-pyridyl analog (**3m**) was synthesized. However, unlike the sulfonamide series, the *N*-methylated amide (**3m**) was less active the secondary amide (**3j**). Likewise, the cyclic lactam (**3n**) was less active than the sulfonamide analog (**2m**), further demonstrating that among the tertiary substituted analogs sulfonamides were preferred over amides. The urea analogs (**3p**–**u**) showed comparable activity to the amides. The unsubstituted urea (**3p**) and the isopropyl (**3q**) or *t*-butyl (**3r**) substituted ureas showed antiviral EC_50_ ∼ 10 μM. The aromatic ureas (**3s**–**3u)** were more active, but had much lower aqueous solubility as was seen in the sulfonamide series (Table 3). In general, ATPase inhibition was weaker in the amide series with only succinimide **3o** demonstrating an IC_50_ ∼ 1 μM.

**Table 4.** Antialphaviral Activity and Solubility of Amides and Ureas 3a–u.

| Compound | R<sup>1</sup> | R<sup>2</sup> | CHIKV-nLuc<br>(pEC<sub>50</sub>) | nsP2<br>ATPase<br>(pIC<sub>50</sub>) | Aqueous<br>Solubility<br>(μM) |
| --- | --- | --- | --- | --- | --- |
| **3a** | *i*-Pr | H | 5.4 | <4.4 (80)<sup>a</sup> | 170 |
| **3b** | *n*-C<sub>5</sub>H<sub>11</sub> | H | 5.7 | 5.2 | 60 |
| **3c** | *c*-C<sub>3</sub>H<sub>5</sub> | H | 5.4 | <4.4 (80)<sup>a</sup> | 160 |
| **3d** | *c*-C<sub>6</sub>H<sub>11</sub> | H | 5.6 | 4.5 | 130 |
| **3e** | (CH<sub>2</sub>)<sub>3</sub>Cl | H | 6.5 | 5.7 | 170 |
| **3f** | Ph | H | 5.6 | 4.8 | 70 |
| **3g** | 4-MeO-Ph | H | 5.4 | 4.7 | 35 |
| **3h** | 3-MeO-Ph | H | 5.3 | <4.4 (70)<sup>a</sup> | 19 |
| **3i** | 4-(CH<sub>2</sub>)<sub>6</sub>O-Ph | H | <4.4 (70)<sup>a</sup> | i.a. | <1 |
| **3j** | 2-pyridyl | H | 5.8 | <4.4 (90)<sup>a</sup> | 190 |
| **3k** | 3-pyridyl | H | 5.6 | 4.9 | 90 |
| **3l** | 4-pyridyl | H | 5.0 | <4.4 (70)<sup>a</sup> | 170 |
| **3m** | 2-pyridyl | Me | 5.3 | <4.4 (80)<sup>a</sup> | 180 |
| **3n** | -(CH<sub>2</sub>)<sub>3</sub>- |  | 5.6 | <4.4 (90)<sup>a</sup> | 200 |
| **3o** | -(CH<sub>2</sub>)<sub>2</sub>C(O)- |  | 6.0 | 5.9 | 160 |
| **3p** | NH<sub>2</sub> | H | 4.9 | <4.4 (50)<sup>a</sup> | 190 |
| **3q** | *i*-PrNH | H | 4.8 | <4.4 (50)<sup>a</sup> | 180 |
| **3r** | *t*-BuNH | H | 5.0 | <4.4 (80)<sup>a</sup> | 170 |
| **3s** | PhNH | H | 5.9 | 4.8 | 10 |
| **3t** | 4-MeO(C<sub>6</sub>H<sub>4</sub>)NH | H | 5.4 | 4.7 | 10 |
| **3u** | 2-MeO(C<sub>6</sub>H<sub>4</sub>)NH | H | 5.4 | i.a. | <10 |

<sup>a</sup>Value in parentheses is % inhibition at 40 μM (CHIKV-nLuc) or 200 μM (ATPase). i.a. =inactive at 200 μM. NT = not tested.

To potentially improve aqueous solubility a series of amine analogs was synthesized (Table 5). Although the amine analogs showed good aqueous solubility, it was no better than many of the sulfonamides and amides and the primary amine (**4a**) and methylamine (**4b**) exhibited poor antiviral activity. The trifluoroethylamine (**4c**) and benzyl amine (**4d**) had improved CHIKV inhibitory activity, but ATPase inhibition remained low. Introduction of a hydroxyl group on the spirocyclic core in analog **4e** did not improve antiviral activity or ATPase inhibition. In general, the amine analogs (**4a**–**e)** demonstrated inferior antiviral activity compared to the amides and sulfonamides demonstrating the preference for at least one H-bond acceptor at this end of the inhibitor.

**Table 5.** Antialphaviral Activity and Solubility of Amines 4a–e.

| Compound | R | X | CHIKV-nLuc<br>(pEC<sub>50</sub>) | nsP2<br>ATPase<br>(pIC<sub>50</sub>) | Aqueous<br>Solubility<br>(μM) |
| --- | --- | --- | --- | --- | --- |
| **4a** | H | H | <4.4 (60)<sup>a</sup> | i.a. | 160 |
| **4b** | Me | H | <4.4 (60)<sup>a</sup> | i.a. | 170 |
| **4c** | CF<sub>3</sub>CH<sub>2</sub> | H | 4.9 | <4.4 (60)<sup>a</sup> | 170 |
| **4d** | PhCH<sub>2</sub> | H | 5.0 | <4.4 (80)<sup>a</sup> | 180 |
| **4e** | H | OH | <4.4 (50)<sup>a</sup> | i.a. | 170 |

<sup>a</sup>Value in parentheses is % inhibition at 40 μM (CHIKV-nLuc) or 200 μM (ATPase). i.a. =inactive at 200 μM.

Switching attention to the amidopiperidine side of the spirocyclic core, the role of the substituents on the quaternary carbon of the amide was explored (Table 6). The methylene analog (**5a**) lacking the cyclopentyl group was inactive. Likewise, the gem-dimethyl analog (**5b**) was inactive. Even small changes to the size of the ring were poorly tolerated, with the cyclobutyl analog (**5c**) being inactive and the cyclohexyl analog (**5d**) showing only weak antiviral activity. Incorporation of an oxygen atom as a pyran ring (**5e**) further diminished antiviral activity. In the sulfamide series the tetrahydropyran (**5f**) and keto (**5g**) analogs were also inactive. Loss in antiviral activity was accompanied by low or no measurable ATPase inhibition. We concluded that the cyclopentyl substituent adjacent to the amide carbonyl was essential for antialphaviral activity and the inability to tolerate even small modifications suggested that it played a critical role in nsP2hel inhibition.

**Table 6.**
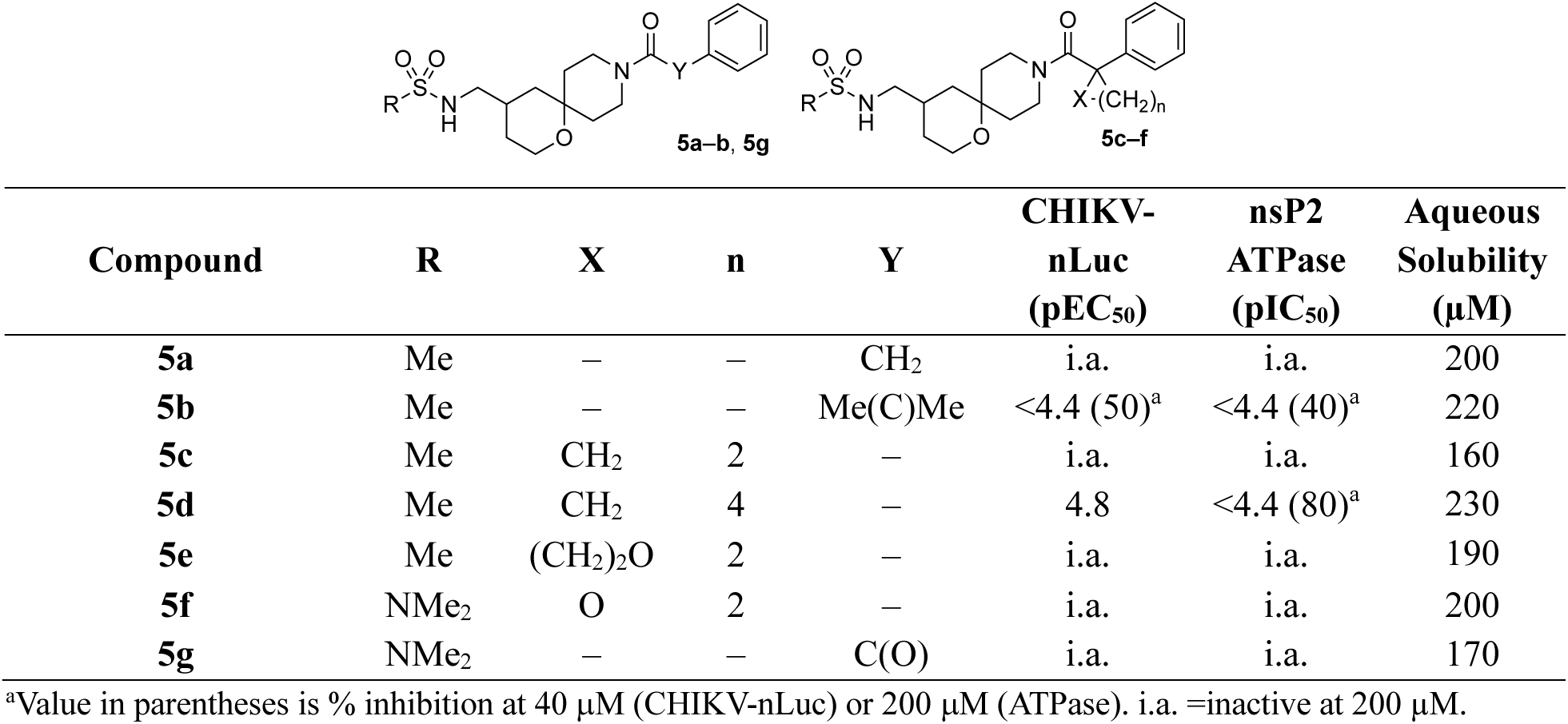
Antialphaviral Activity and Solubility of Sulfonamides and Sulfamides 5a–g.

The contribution of the terminal phenyl group to the antiviral activity of the inhibitors was probed in a series of analogs (Table 7). Substituting the phenyl group for a methyl group gave analog **6a** which lacked antiviral activity. Turning to more subtle modifications, we found that *para*-substituents on the phenyl ring in analogs **6b**–**d** were also not tolerated. In contrast, the *meta*-chlorophenyl analog (**6e**) mirrored the activity of the parent phenyl analog (**2a**). In the sulfamide series *meta*-substituted phenyl analogs were also active, with the 3-methoxyphenyl (**6f**) and *meta*-tolyl (**6g**) analogs demonstrating similar activity and only the 3-trifluromethyl analog (**6h**) showing lower activity than the parent **2g**. The *ortho*-chloro (**6i**) and *ortho-*fluoro (**6j**) substituted analogs also showed good activity against CHIKV. Replacement of the phenyl group with pyridine resulted in analogs **6k**–**m** with weak or no antiviral activity. Overall, the terminal phenyl substituent was shown to be essential for antiviral activity, with little tolerance for modification. Only the *ortho*-chloro (**6i**) and *ortho-*fluoro (**6j**) analogs maintained antiviral activity and potent ATPase inhibition. It appeared that the piperidine amide with both cyclopentyl and phenyl substituents was a critical pharmacophore for antiviral activity suggesting that it adopted a conformation that fit into a key hydrophobic pocket in the nsP2hel protein.

**Table 7.** Antialphaviral Activity and Solubility of Sulfonamides and Sulfamides 6a–m.

| Compound | R<sub>1</sub> | R<sub>2</sub> | CHIKV-nLuc<br>(pEC<sub>50</sub>) | nsP2<br>ATPase<br>(pIC<sub>50</sub>) | Aqueous<br>Solubility<br>(μM) |
| --- | --- | --- | --- | --- | --- |
| **6a** | Me | Me | <4.4 (40)<sup>a</sup> | i.a. | 110 |
| **6b** | Me | 4-MeO-Ph | i.a. | i.a. | 210 |
| **6c** | Me | 4-Cl-Ph | <4.4 (70)<sup>a</sup> | i.a. | 220 |
| **6d** | Me | 4-Me-Ph | i.a. | i.a. | 160 |
| **6e** | Me | 3-Cl-Ph | 5.6 | 4.7 | >220 |
| **6f** | NMe<sub>2</sub> | 3-MeO-Ph | 5.2 | <4.4 (80)<sup>a</sup> | 170 |
| **6g** | NMe<sub>2</sub> | 3-Me-Ph | 5.4 | <4.4 (90)<sup>a</sup> | 140 |
| **6h** | NMe<sub>2</sub> | 3-F<sub>3</sub>C-Ph | 4.7 | <4.4 (40)<sup>a</sup> | 90 |
| **6i** | NMe<sub>2</sub> | 2-Cl-Ph | 5.9 | 5.1 | 240 |
| **6j** | NMe<sub>2</sub> | 2-F-Ph | 5.3 | 4.5 | 230 |
| **6k** | NMe<sub>2</sub> | 2-pyridyl | <4.4 (70)<sup>a</sup> | <4.4 (40)<sup>a</sup> | 170 |
| **6l** | NMe<sub>2</sub> | 3-pyridyl | <4.4 (50)<sup>a</sup> | i.a. | 160 |
| **6m** | NMe<sub>2</sub> | 4-pyridyl | i.a. | i.a. | 180 |

<sup>a</sup>Value in parentheses is % inhibition at 40 μM (CHIKV-nLuc) or 200 μM (ATPase). i.a. =inactive at 200 μM.

Cyclic sulfonamide **2o** emerged as the most potent analog in the CHIKV-nLuc viral replication assays. To confirm that this activity was seen on the wild type virus and to explore the breadth of antialphaviral activity, **2o** was tested for its ability to reduce viral titer against clinical isolates of three alphaviruses: CHIKV, VEEV, and MAYV (Figure 3). Sulfonamide **2o** produced a dose dependent effect with a 10^3^–10^8^ reduction in viral titer on the three alphaviruses. It was most potent against the athritogenic alphaviruses MAYV and CHIKV with EC_50_ 1–3 μM but only showed activity at ∼10 μM against the encephalitic alphavirus VEEV.^20^ The weaker antiviral activity against VEEV matched the trend observed in the VEEV-nLuc reporter assay across the full series of oxapiropiperidines (Table S1 and Figure S9).

**Figure 3.**
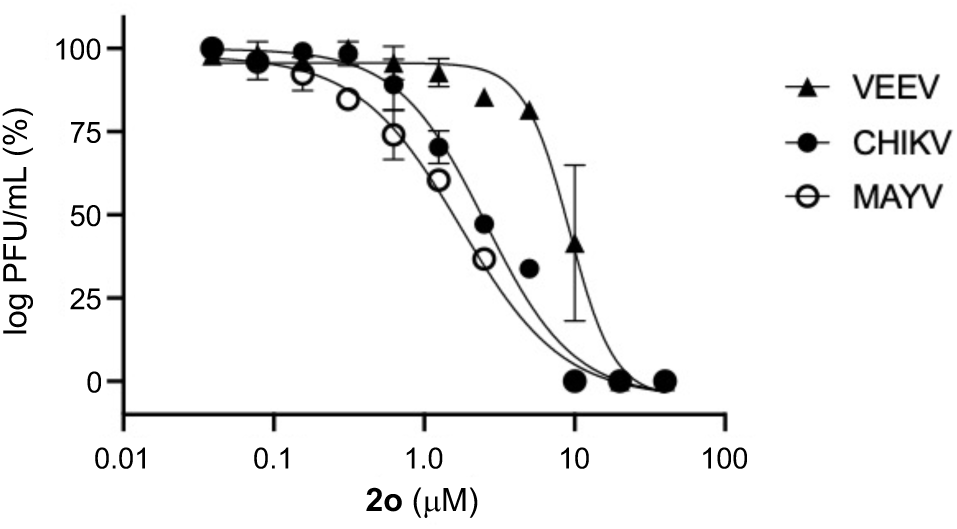
Antialphaviral activity of **2o**. Titer reduction in cells infected with the respective alphaviruses measured by plaque assay, n =3 ± SD.

To demonstrate that the cellular effects on viral replication were likely to be the result of nsp2hel inhibition a series of additional biophysical and biochemical assays were performed. First, the correlation between the EC_50_ for CHIKV antiviral assay and the IC_50_ for nsP2hel ATPase inhibition was compared using 26 analogs for which a complete dose response was determined in the respective assays. A robust positive correlation was observed (Figure 4) with an *R*^2^ = 0.85, which is highly supportive of helicase inhibition as the mechanism of action of the oxaspiropiperidines.

**Figure 4.**
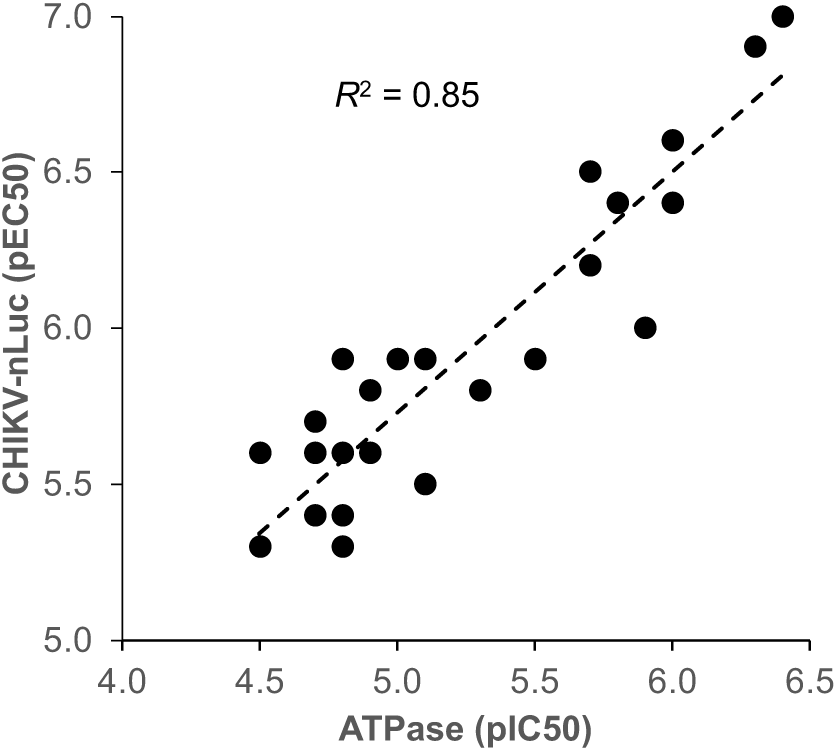
Correlation of CHIKV antiviral activity with nsP2 ATPase inhibition.

**Figure 5.**
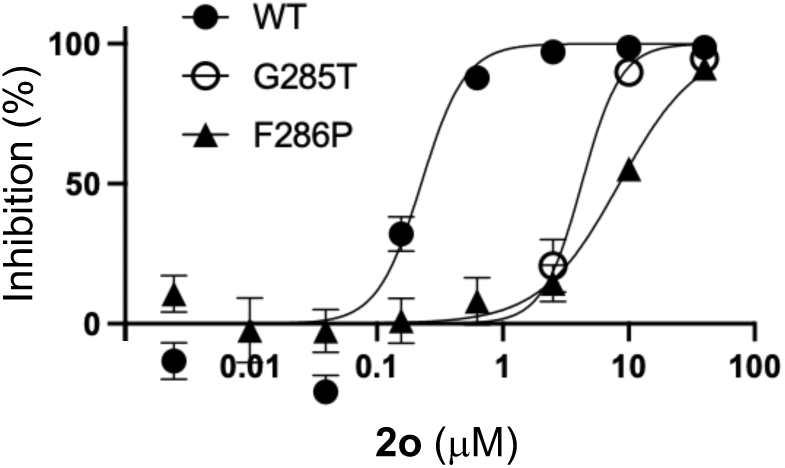
Activity of **2o** on point mutants of CHIKV-nLuc located in the RecA1 domain of nsP2hel.

Next, sulfonamide **2o** was tested against two synthetic point mutants of CHIKV-nLuc that had been generated to probe the function of conserved amino acids across diverse alphaviruses. The point mutants G285T and F286P are located in the RecA1 core of the nsP2hel domain (Figure S7),^21^ but do not affect the antiviral activity of NHC, an inhibitor of the viral polymerase (Figure S8). When tested on the G285T and F286P mutants, there was a >10-fold right shift in the EC_50_ for inhibition of viral replication by **2o**, clearly indicating that the antiviral potency of the sulfonamide was decreased by specific point mutations within the nsp2hel domain.

Although the correlation between biochemical ATPase inhibition and antiviral activity and the viral resistance from point mutants in the nsP2hel domain provided strong evidence of direct targeting of the viral helicase, we sought additional biophysical evidence of direct interaction of the spiropiperidines with the nsP2 protein. SAR in the sulfonamide and sulfamide series (Table 7) suggested that the phenyl substituent on the tertiary amide may reside in a tight lipophilic pocket, since only small halogen substituents on the ring were tolerated. To develop a potential fluorine-substituted probe for use in NMR experiments three analogs of the potent cyclic sulfonamide **2o** were prepared (Scheme 2). Fluorine substitution was paced at the *ortho* (**7a**), *meta* (**7b**), and *para* (**7c**) positions on the phenyl ring. Testing in the CHIKV-nLuc viral replication and ATPase inhibition assays showed that the 3,5-difluorophenyl derivative (**7c**) was the most potent of the analogs with activity equivalent to the parent cyclic sulfonamide **2o** (Table 8).

**Table 8.**
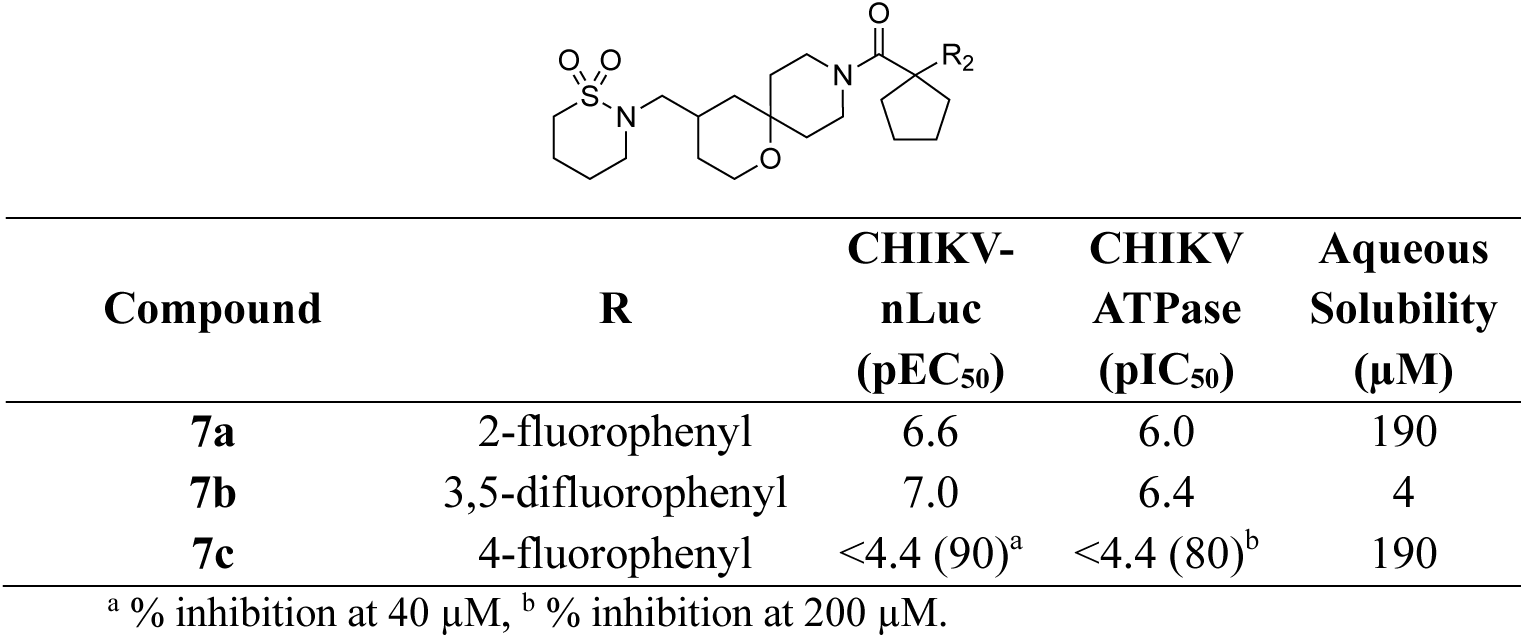
Antialphaviral Activity and Solubility of Fluorophenyl Analogs 7a–c.

| Compound | R | CHIKV-nLuc<br>(pEC<sub>50</sub>) | CHIKV<br>ATPase<br>(pIC<sub>50</sub>) | Aqueous<br>Solubility<br>(μM) |
| --- | --- | --- | --- | --- |
| **7a** | 2-fluorophenyl | 6.6 | 6.0 | 190 |
| **7b** | 3,5-difluorophenyl | 7.0 | 6.4 | 4 |
| **7c** | 4-fluorophenyl | <4.4 (90)<sup>a</sup> | <4.4 (80)<sup>b</sup> | 190 |

<sup>a</sup> % inhibition at 40 μM, <sup>b</sup> % inhibition at 200 μM.

Despite the relatively poor aqueous solubility, **7b** was selected as an ^19^F NMR probe. The compound alone showed a strong signal at −109.78 ppm from the two fluorine atoms in the ^19^F NMR (Figure 6). Upon addition of purified protein representing the nsP2hel domain alone a dose dependent downfield shift and line broadening of the of the ^19^F signal was observed in the NMR spectra. These data demonstrate a biophysical interaction between **7b** and the nsP2hel domain protein and support the conclusion that the antiviral oxaspiropiperidine sulfonamides are direct nsP2hel inhibitors.

**Figure 6.**
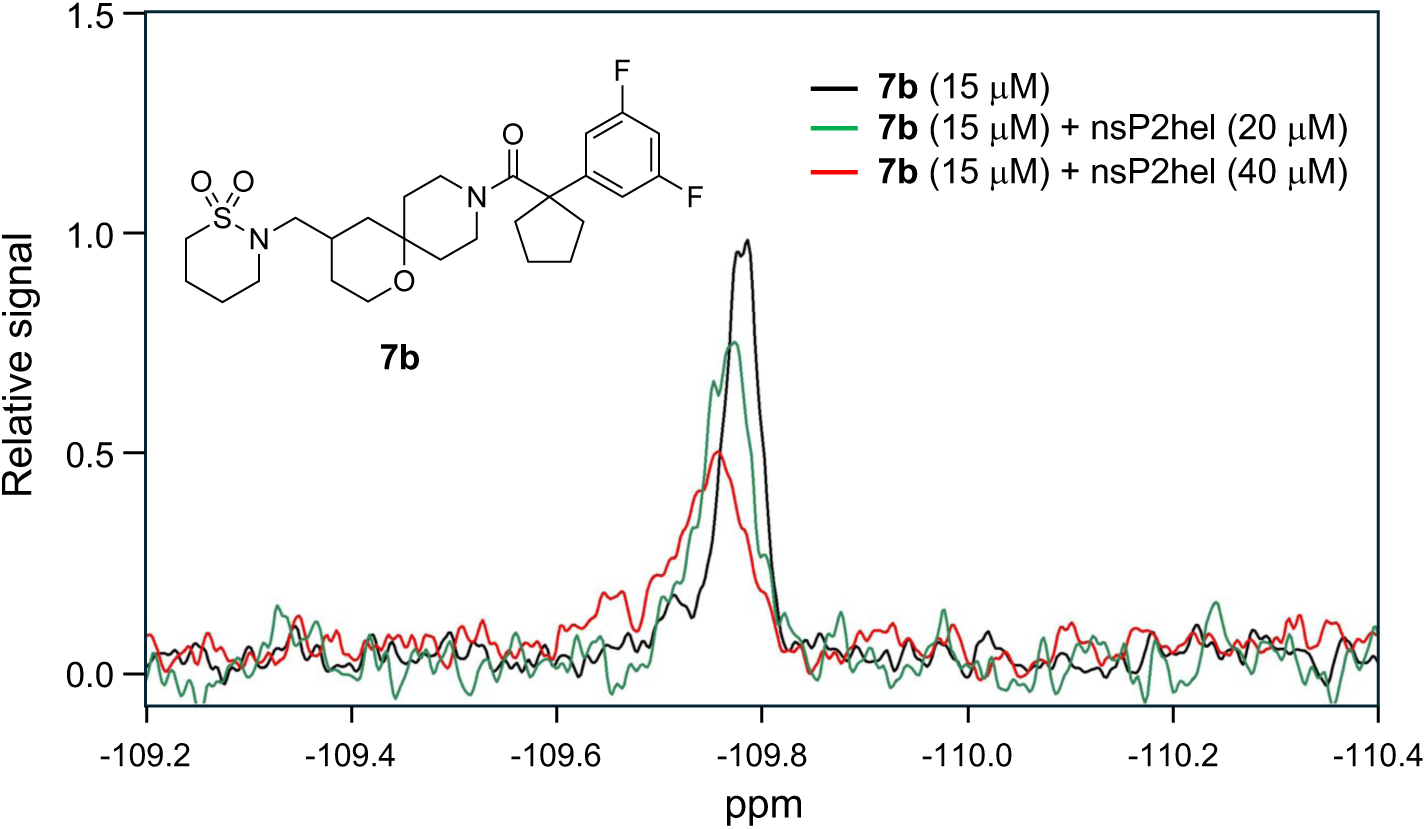
^19^F NMR spectra of **7b** in the absence or presence of nsP2hel protein (aa 1–464). Relative concentrations of the ligand and protein are indicated.

To further characterize the mechanism of nsP2hel inhibition by **2o**, we evaluated the impact of the compound on the ATPase and RNA helicase activities (Figure 7). Using an assay that directly monitored the conversion of ^32^P-ATP to ^32^P-ADP by nsP2, the effect of **2o** was measured at two time points. As had been seen in the multistep ADP-glo assay, the cyclic sulfonamide **2o** was a potent inhibitor of nsP2 ATPase activity (Figure 7A). Notably, the IC_50_ of **2o** for inhibition of ATPase activity was the same at both the 5 min and 10 min time points (Figure 7B). The ATPase activity of nsP2 was also measured across a range of ATP concentrations to determine maximum rate of catalysis (*V*_max_) and the substrate concentration at which the reaction rate was at its half maximum value (*K*_m_). In the presence of **2o** (1.0 or 2.5 μM) the *V*_max_ was decreased, but the *K*_m_ was unchanged (Figure 7C). The effect of **2o** on the RNA helicase activity of nsP2 was studied using an assay that employed a forked RNA duplex composed of a 39mer loading strand and a 19mer displaced strand. Production of the displaced ssRNA strand was inhibited by **2o** dose dependently with an IC_50_ = 0.9 μM (Figure 7D). However, **2o** did not change the binding affinity of fluorescein-labelled dsRNA to nsP2 as measured by fluorescence polarization (Figure 7E), suggesting that the inhibitor did not directly displace the RNA substrate from the enzyme. Thus, kinetic analysis of nsP2 ATPase inhibition by **2o** at multiple time points and multiple substrate concentrations was consistent with a non-competitive mechanism where the inhibitor was bound to both the enzyme and the enzyme•ATP complex. Likewise, inhibition of nsP2 helicase activity by **2o** did not involve direct displacement of the RNA substrate. Together, these data are consistent with a mechanism of helicase inhibition by **2o** consistent with an allosteric mechanism of action that may interfere with conformational dynamics of the enzyme required for ATP hydrolysis both in the absence and presence of RNA.

**Figure 7.**
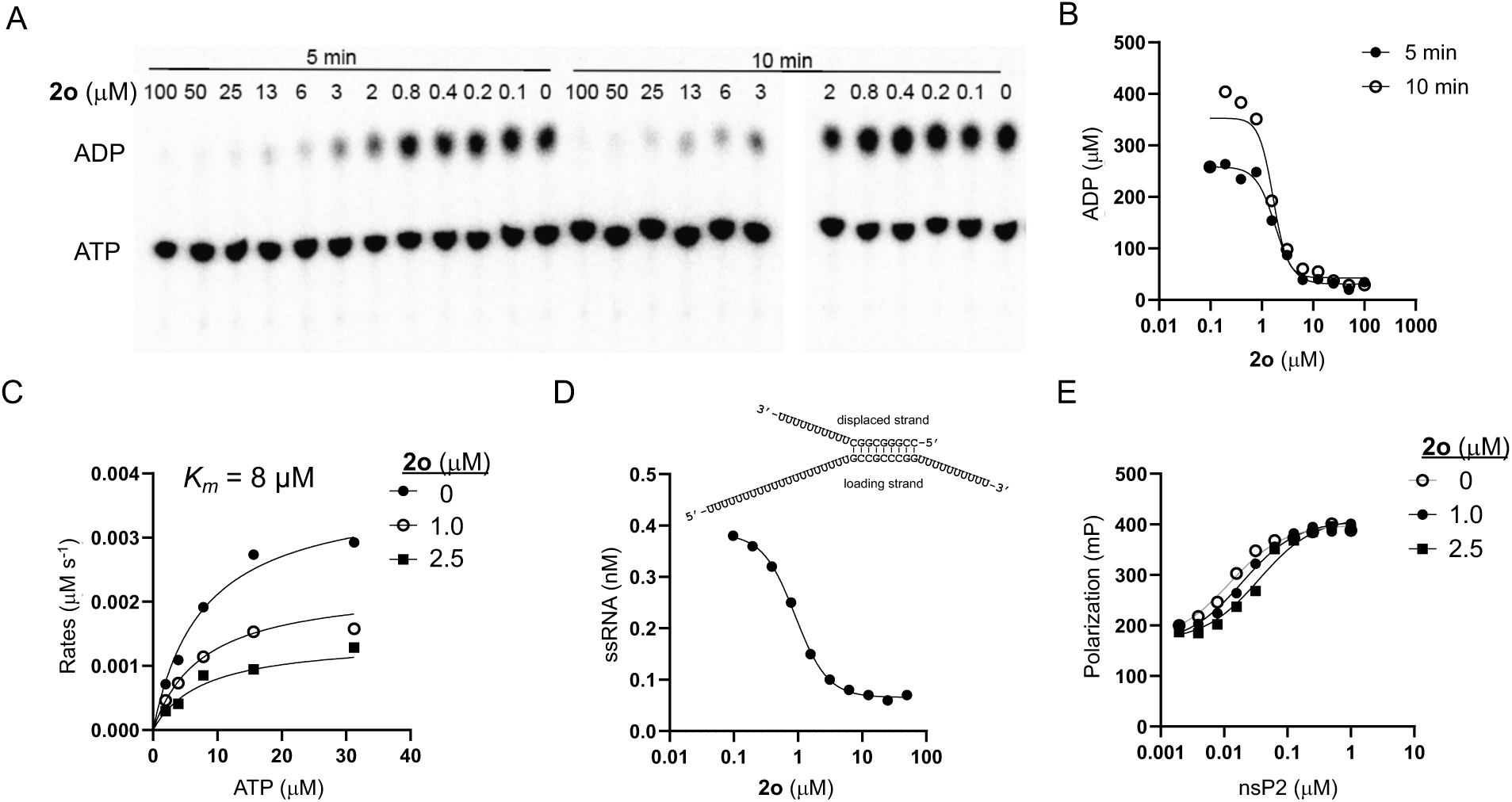
Mechanism of nsP2hel inhibition by **2o**. (A) and (B) Conversion of ^32^P-ATP to ^32^P-ADP by nsP2 was monitored by TLC at two time points. The IC_50_ of **2o** for inhibition of ADP formation did not change. (C) Rate of conversion of ^32^P-ATP to ^32^P-ADP by nsP2 at different substrate concentrations. The enzyme *K*_m_ was not affected by **2o**. (D) **2o** inhibited conversion of dsRNA to ssRNA by nsP2. Structure of the forked dsRNA with 5’ and 3’ overhangs is shown. (E) **2o** did not preclude binding of fluorescein-labeled dsRNA to increasing concentrations of nsP2.

## Conclusions

We used a plate-base viral replication assay containing an nLuc reporter to explore the SAR of a series of oxaspiropiperidines resulting in the identification of first-in-class CHIKV nsP2hel inhibitors with direct-acting antiviral activity. Similar to the optimization of HSV helicase inhibitors,⁹ the success of the medicinal chemistry campaign relied on robust high-throughput antiviral assays, as developing early SAR trends in the ATPase assay was challenging due to the initial screening hits exhibiting biochemical activity only in the high micromolar range. The optimized inhibitors contain an oxaspiropiperidine core that was capped with an unusual amide containing both a cyclopentane ring and a phenyl group. Although this highly substituted amide resulted in complex NMR spectra due to restricted rotation in relation to oxaspiropiperidine core, both the cyclopentane ring and phenyl ring were critical for potent helicase inhibition. All modifications to the cycloalkyl group resulted in a dramatic decrease in antiviral activity and only halogens were accommodated on the phenyl ring suggesting that both amide substituents were bound into a tight lipophilic pocket in the helicase enzyme. The other end of the oxaspiropiperidine core was more tolerant of modifications. Incorporation of a cyclic sulfonamide led to the identification of **2o** which was a potent inhibitor of the helicase ATPase and RNA unwindase that also demonstrated antiviral activity in the 100 nM range. Multiple lines of evidence demonstrated that these oxaspiropiperidines were direct inhibitors of nsP2hel: ATPase inhibition correlated with antiviral activity; point mutants in the helicase domain of nsP2 resulted in viral resistance; and a difluoro-substituted analog showed direct interaction with the protein by ^19^F NMR. Kinetic studies demonstrated that **2o** was a non-competitive inhibitor of enzyme activity, which indicated that it was likely to be a Type 1 allosteric inhibitor of the helicase.^8^ Although it was tempting to build models in which these nsP2hel inhibitors were docked into the region defined by the resistance mutants (G285/F286), there is as yet no direct evidence that the molecules bind to the protein at this precise location. In fact, there are multiple mechanisms by which point mutations can perturb conformational changes between the helicase RecA domains to inhibit procession of the motor enzyme along its nucleotide substrate,^22^ which would in turn lead indirectly to resistance to an allosteric inhibitor.^8^ Ongoing research with these inhibitors using a combination of biophysics, chemoproteomics, viral resistance induction, and structural studies will provide a more complete picture of the molecular basis of small molecule nsP2hel inhibition. In the interim, **2o** will be a valuable tool to study the role of nsP2hel in viral replication and serve as a lead for the development of drugs effective against SF1 helicases across the full spectrum of pathogenic alphaviruses.

## Experimental Section

### General Information

All reactions were conducted in oven-dried glassware under a dry nitrogen atmosphere unless otherwise specified. All reagents and solvents were obtained from commercial sources and used without further purification. No unexpected safety hazards were encountered during the syntheses. Analytical thin layer chromatography (TLC) was performed on pre-coated silica gel plates (200 μm, F_254_ indicator), visualized under UV light or by staining with iodine and KMnO₄. Column chromatography utilized pre-loaded silica gel cartridges on a Biotage automated purification system. ¹H and ¹³C NMR spectra were recorded in DMSO-*d*₆, CD_3_CN, CDCl_3_, Acetone-*d*_6_, Pyridine-*d*_5_ and CD_3_OD at 400/500/700 and 101/126/176 MHz, respectively, on Bruker spectrometers. Chemical shifts (*δ*) are reported in parts per million (ppm) downfield from tetramethylsilane for ¹H NMR, with major peaks designated as s (singlet), d (doublet), t (triplet), q (quartet), and m (multiplet), dd (doublet of doublets), td (triplet of doublets), qd (quartet of doublets), tt (triplet of triplets) and ddd (doublet of doublet of doublets). High-resolution mass spectrometry (HRMS) analyses were performed at the UNC Department of Chemistry Mass Spectrometry Core Laboratory using a Q Exactive HF-X mass spectrometer. Liquid chromatography-mass spectrometry (LC-MS) was conducted on an Agilent 1290 Infinity II LC System with an Agilent Infinity Lab PoroShell 120 EC-C18 column (30 °C, 2.7 μm, 2.1 × 50 mm), employing a 5−95% CH₃CN in water eluent, with 0.2% (v/v) formic acid as the modifier and a flow rate of 1 mL/min. Preparative high-performance liquid chromatography (HPLC) was executed using an Agilent 1260 Infinity II LC System equipped with a Phenomenex C18 column (PhenylHexyl, 30 °C, 5 μm, 75 × 30 mm), with a 5−95% CH₃CN in water eluent and 0.05% (v/v) trifluoroacetic acid as the modifier, at a flow rate of 30 mL/min. Analytical HPLC data were recorded on a Waters Alliance HPLC with a PDA detector or an Agilent 1260 Infinity II series with a PDA detector (EC-C18, 100 mm × 4.6 mm, 3.5 μm), using a 10−90% CH₃CN in water eluent at a flow rate of 0.5 mL/min. Final compounds were isolated as non-hygroscopic foams. In a few cases where the compound was isolated as a gum, addition of (Et)_2_O and evaporation yielded a foam as the final form. The final compounds were confirmed to be >95% pure by HPLC analysis. The low intensity or absence of carbon signals in spiropiperidines is attributed to molecular rotations occurring on the NMR time scale. *Tert*-butyl 4-hydroxy-1-oxa-9-azaspiro[5.5]undecane-9-carboxylate (**I**), and all carboxylic acids and sulfonyl chlorides were obtained from commercial sources.

### General Methods

#### General Procedure-I

To a stirred solution of the Boc-protected intermediate (1.0 eq), 4 M HCl in dioxane (5 volumes) was added, or the intermediate was dissolved in DCM (10 volumes) followed by the addition of trifluoroacetic acid (5 volumes). The mixture was stirred at 0–25 °C for 1–4 h. Upon completion (based on TLC and LCMS), the reaction mixture was concentrated, and the resulting crude product was either used without further purification or purified using preparative HPLC, if required, to afford the desired compound.

#### General procedure-II

To a stirred solution of carboxylic acid (1.1 eq) in DMF, HATU (1.5 eq) and DIPEA (4.0 eq), were added at 0 °C. The reaction mixture was stirred at 0 °C for 10 min after which the amine salt was added at the same temperature. Reaction mixture was stirred at rt for 16 h. On completion of the reaction (based on TLC and LCMS), the reaction mixture was diluted with water and extracted with EtOAc. The organic layers were combined and washed with brine, dried over Na_2_SO_4_, filtered and concentrated to give crude compound. The resulting crude product was purified by combiflash or preparative HPLC to afford the desired product.

#### General procedure-III

To a stirred solution of acid (1.0 eq) and 2-(1*H*-benzotriazole-1-yl)-1,1,3,3-tetramethylaminium tetrafluoroborate (TBTU) (1.5 eq) in pyridine was added corresponding amine substituent (1.2 eq) and the reaction was stirred at 25 °C for 6 h. On completion of the reaction (based on TLC and LCMS analysis), the reaction was poured into water, extracted with EtOAc, and the combined organic layers dried over anhydrous Na_2_SO_4_, filtered, and concentrated. The crude product was purified by combiflash or preparative HPLC to afford the desired product.

#### General procedure-IV

To a stirred solution of carbonitrile (1.0 eq) in MeOH (10 volumes) at 0 °C, (Boc)_2_O (2.5 eq) and NiCl_2_·6H_2_O (0.2 eq) were added. After stirring for 15 minutes, NaBH_4_ (10.0 eq) was added portion-wise over 20 minutes. The reaction was then stirred at 25 °C for 6 h. On completion of the reaction (based on TLC and LCMS analysis), diethylenetriamine (2.0 eq) was added and stirred for 30 minutes. The solvent was evaporated and the purple residue was dissolved in EtOAc, washed with saturated NaHCO_3_, dried over Na_2_SO_4_, filtered, and concentrated. The crude product was purified by combiflash to afford the desired compound.

#### General procedure-V

To a stirred solution of amine salt (1.0 eq) and Et_3_N (4 eq) in DCM (5 volumes) at 0°C was added sulfonyl chlorides (1.5 eq) or isocyanates (1.5 eq) or acid chloride (1.2 eq) and stirred at rt for 2–16 h. Reaction progress was monitored by TLC and LCMS. After completion, the solvent was removed in the vacuo and the residue was diluted with water. The aqueous layer was extracted with EtOAc, and the combined organic layers were washed with brine, dried over Na_2_SO_4_, filtered, and concentrated. The crude product was purified by combiflash or preparative HPLC to afford the desired product.

#### General procedure-VI

A solution of chloroalkylsulfonamide (1 eq) and Cs₂CO₃ (2 eq) in DMF was stirred at 60 °C for 2 h. Alternatively, chloroalkylamide (1 eq) was added to NaH (2 eq) in THF at 0 °C and stirred at 25 °C for 2 h. After completion (based on LCMS and TLC), the reaction mixture was diluted with water and extracted with EtOAc. The combined organic layers were washed with brine, dried over Na_2_SO_4_, filtered, and concentrated to obtain the crude compound. The resulting crude product was purified by combiflash or preparative HPLC to afford the desired product.

#### General procedure-VII

A solution of tert-butyl 4-hydroxy-1-oxa-9-azaspiro[5.5]undecane-9-carboxylate (1 eq) in DMF was added KO*^t^*Bu (2 eq) under argon. The resulting mixture was stirred at 50 °C for 2 h, after which a solution of the respective aryl halide (1.5 eq) in DMF was added and stirred at 85 °C for 4 h. After completion (based on LCMS and TLC), the reaction mixture was diluted with water and extracted with EtOAc. The combined organic layers were washed with brine, dried over Na_2_SO_4_, filtered, and concentrated to obtain the crude compound. The resulting crude product was purified by combiflash to afford the desired product.

#### General procedure-VIII

To a suspension of NaH in DMF at 0 °C, secondary sulfonamides or carbamates were added and stirred at room temperature for 1 h. The reaction mixture was then cooled again to 0 °C, followed by the addition of MeI or BnBr, and stirred at room temperature for 12 h. Alternatively, a solution of secondary sulfonamides in DMF was added K_2_CO_3_ and 1-iodohexane and stirred at 85 °C for 12 h. On completion of the reaction (based on TLC and LCMS analysis), the reaction mixture was quenched with cold water and extracted with EtOAc. The combined organic layers were washed with brine, dried over Na_2_SO_4_, filtered, and concentrated to obtain the crude compound. The resulting crude product was purified by combiflash or preparative HPLC to afford the desired product.

#### Synthesis of intermediates IIa, IIb and III–IV

*O*-alkylation of *tert*-butyl 4-hydroxy-1-oxa-9-azaspiro[5.5]undecane-9-carboxylate **I** with 2-chloropyridines following General Procedure-VII afforded **IIa** and **IIb**.

#### Tert-butyl 4-((4-methylpyridin-2-yl)oxy)-1-oxa-9-azaspiro[5.5]undecane-9-carboxylate (IIa)

White foam (46% yield). ^1^H NMR (700 MHz, CDCl_3_): δ 8.14–8.04 (m, 1H), 6.85 (s, 1H), 6.67–6.62 (m, 1H), 5.55 (br s, 1H), 3.96–3.92 (m, 1H), 3.88–3.70 (m, 3H), 3.24–3.13 (m, 2H), 2.40–2.36 (m, 3H), 2.19 (d, *J* = 12.5 Hz, 1H), 2.17–2.08 (m, 1H), 2.06–1.97 (m, 1H), 1.84 (d, *J* = 13.6 Hz, 1H), 1.77–1.70 (m, 1H), 1.69–1.63 (m, 1H), 1.63–1.56 (m, 2H), 1.50 (s, 9H). MS (ESI) *m*/*z*: 363.2 [M + H]^+^.

#### Tert-butyl 4-((pyridin-2-yl)oxy)-1-oxa-9-azaspiro[5.5]undecane-9-carboxylate (IIb)

White foam (75% yield). ^1^H NMR (400 MHz, CDCl_3_): δ 8.11–8.09 (m, 1H), 7.57–7.53 (m, 1H), 6.87–6.75 (m, 1H), 6.67 (d, *J* = 8.4 Hz, 1H), 5.38 (dt, *J* = 9.4, 4.9 Hz, 1H), 3.89 (dt, *J* = 12.3, 4.4 Hz, 1H), 3.79–3.63 (m, 3H), 3.23–3.05 (m, 2H), 2.13–1.99 (m, 2H), 1.80 (d, *J* = 13.8 Hz, 1H), 1.74–1.65 (m, 1H), 1.63–1.47 (m, 4H), 1.45–1.42 (m, 9H). MS (ESI) *m*/*z*: 349.2 [M + H]^+^.

#### Tert-butyl 4-oxo-1-oxa-9-azaspiro[5.5]undecane-9-carboxylate (III)

To a stirred solution of tert-butyl 4-hydroxy-1-oxa-9-azaspiro[5.5]undecane-9-carboxylate **I** (35.0 g, 129.15 mmol, 1.0 eq) in DCM (350 mL) was added PDC (72.84 g, 193.72 mmol, 1.5 eq) was added portion-wise at 25 °C. The reaction mixture was stirred at the same temperature for 16 h. Reaction progress was monitored by TLC and LCMS, after completion, the mixture was filtered through high flow bed of celite and the filtrate was washed with 5% HCl solution (100 mL) and the desired compound extracted in DCM (500 mL x 2). The organic layer was dried over Na_2_SO_4_ and concentrated to obtained crude which was purified through combiflash using 30% EtOAc in n-hexane as elution system to afford *tert*-butyl 4-oxo-1-oxa-9-azaspiro[5.5]undecane-9-carboxylate **III** as a white solid (23 g, 66% yield). MS (ESI) *m*/*z*: 270.2 [M + H]^+^.

#### Tert-butyl 4-((4-methylpyridin-2-yl)amino)-1-oxa-9-azaspiro[5.5]undecane-9-carboxylate (IV)

To a solution of *tert*-butyl 4-oxo-1-oxa-9-azaspiro[5.5]undecane-9-carboxylate **III** (30 mg, 0.111 mmol, 1 eq) in THF (0.6 mL) was added 2-amino-4-methylpyridine (21.7 mg, 0.2 mmol, 1.8 eq) and titanium isoproxide (0.06 mL, 0.228 mmol, 2 eq). The resulting mixture was stirred at 60 °C for 12 h. Removal of THF under vacuum afforded a yellow crude solid. The yellow solid was then dissolved in methanol and cooled to 0 °C. NaBH_4_ was added portion-wise, and the mixture was stirred at 25 °C for 6 h. Methanol was removed under vacuum, and the residue was diluted with DCM. The solid was filtered, and the filtrate was concentrated under vacuum, followed by purification using combiflash with 100% EtOAc as eluent to furnish *tert*-butyl 4-((4-methylpyridin-2-yl)amino)-1-oxa-9-azaspiro[5.5]undecane-9-carboxylate (**IV**) as a colorless oil (24 mg, 60%). MS (ESI) *m*/*z*: 362.2 [M + H]^+^.

#### Synthesis of 1a–c and 1d

Boc deprotection of **IIa**, **IIb** and **IV** using TFA/DCM followed by amide coupling using the General Procedures I and II afforded **1a**–**c**.

#### (4-((4-Methylpyridin-2-yl)oxy)-1-oxa-9-azaspiro[5.5]undecan-9-yl)(1-phenylcyclopentyl)methanone (1a)

White foam (60% yield, over two steps). ^1^H NMR (700 MHz, CDCl_3_): δ 8.01–7.93 (m, 1H), 7.27 (dd, *J* = 14.0, 6.6 Hz, 2H), 7.22–7.12 (m, 3H), 6.71 (s, 1H), 6.53–6.48 (m, 1H), 5.31–5.29 (m, 1H), 4.30–4.21 (m, 1H), 3.81–3.79 (m, 1H), 3.72–3.52 (m, 1H), 3.23–3.19 (m, 1H), 3.12 – 2.99 (m, 1H), 2.88–2.80 (m, 1H), 2.48–2.35 (m, 2H), 2.29 (s, 3H), 2.04 (br s, 1H), 2.01–1.88 (m, 2H), 1.83 –1.82 (m, 1H), 1.77–1.69 (m, 4H), 1.64–1.30 (m, 5H), 0.94–0.78 (m, 1H). ^13^C NMR (176 MHz, CDCl_3_) δ 174.6, 146.0, 128.8, 126.2, 125.2, 118.5, 112.0, 71.7, 58.9, 58.6, 42.0, 41.7, 41.1, 39.3, 38.5, 38.3, 37.9, 37.1, 36.8, 31.9, 25.4, 21.2, 14.3. HRMS (ESI) *m/z*: [M + H]^+^ calculated for C_27_H_35_N_2_O_3_: 435.2647, found 435.2639.

#### (1-Phenylcyclopentyl)(4-((pyridin-2-yl)oxy)-1-oxa-9-azaspiro[5.5]undecan-9-yl)methanone (1b)

White foam (61% yield, over two steps). ^1^H NMR (500 MHz, DMSO-*d*_6_): δ 8.16–8.06 (m, 1H), 7.72–7.60 (m, 1H), 7.32 (t, *J* = 7.7 Hz, 2H), 7.24–7.13 (m, 3H), 6.94 (ddd, *J* = 7.2, 5.0, 1.0 Hz, 1H), 6.71 (d, *J* = 8.3 Hz, 1H), 5.20 (s, 1H), 4.05 (br s, 1H), 3.70 (dt, *J* = 12.1, 4.3 Hz, 1H), 3.57–3.50 (m, 1H), 3.17–3.12 (m, 1H), 3.02–2.62 (m, 3H), 2.30–2.29 (m, 2H), 1.92 1.91 (m, 3H), 1.84 −1.80 (m, 1H), 1.63–1.61 (m, 4H), 1.51 –1.46 (m, 1H), 1.34–1.28 (m, 2H), 1.20–1.15 (m, 1H), 0.83–0.81 (m, 1H). ^13^C NMR (126 MHz, DMSO-*d*_6_) δ 173.0, 162.3, 146.8, 145.6, 139.3, 128.7, 126.0, 124.8, 116.9, 111.2, 71.1, 67.2, 58.1, 57.9, 42.1, 40.5, 40.4, 31.7, 25.0, 24.9. HRMS (ESI) *m/z*: [M + H]^+^ calculated for C_26_H_33_N_2_O_3_: 421.2491, found 421.2485.

#### (4-((4-Methylpyridin-2-yl)amino)-1-oxa-9-azaspiro[5.5]undecan-9-yl)(1-phenylcyclopentyl)methanone (1c)

White foam (20% yield, over two steps). ^1^H NMR (400 MHz, CDCl_3_): δ 8.92 (s, 1H), 7.69 (d, *J* = 6.4 Hz, 1H), 7.30 (t, *J* = 7.6 Hz, 2H), 7.24–7.15 (m, 3H), 6.57 (d, *J* = 6.2 Hz, 1H), 6.48–6.43 (m, 1H), 4.48–4.37 (m, 4H), 3.81 (dd, *J* = 12.2, 4.7 Hz, 1H), 3.72–3.50 (m, 2H), 3.31–3.24 (m, 1H), 3.11–2.82 (m, 2H), 2.43 (s, 3H), 2.30–2.10 (m, 2H), 1.90 (d, *J* = 13.2 Hz, 2H), 1.73–1.69 (m, 3H), 1.66–1.58 (m, 2H), 1.51–1.11 (m, 3H), 0.83–0.65 (m, 1H). ^13^C NMR (100 MHz, CDCl_3_) δ152.6, 136.3, 128.9, 126.6, 125.2, 117.1, 114.2, 113.9, 108.5, 71.1, 59.5, 58.7, 45.9, 41.4, 38.5, 37.9, 37.3, 32.0, 29.6, 25.2, 22.7. HRMS (ESI) *m/z*: [M + H]^+^ calculated for C_27_H_36_N_3_O_2_: 434.2807, found 434.2802.

#### 4-(Phenylamino)-1-oxa-9-azaspiro[5.5]undecan-9-yl)(1-phenylcyclopentyl)methanone (1d)

Boc deprotection of Tert-butyl 4-oxo-1-oxa-9-azaspiro[5.5]undecane-9-carboxylate **III** using TFA/DCM followed by amide coupling using the General Procedures I and II afforded 9-(1-phenylcyclopentane-1-carbonyl)-1-oxa-9-azaspiro[5.5]undecan-4-one **V** as a white foam (64% yield). MS (ESI) *m*/*z*: 342.2 [M + H]^+^. Step-2: to a solution of 9-(1-phenylcyclopentane-1-carbonyl)-1-oxa-9-azaspiro[5.5]undecan-4-one **V** (100 mg, 0.293 mmol, 1 eq) and aniline (0.03 mL, 0.293 mmol, 1 eq) in toluene (2 mL) were added acetic acid (42 mL, 0.733 mmol, 2.5 eq) and sodium triacetoxyborohydride (99 mg, 0.47 mmol, 1.6 eq) and heated at 140 °C in microwave for 20 min. After completion (based on TLC and LCMS), the reaction was quenched with water, extracted with EtOAc, washed with brine, dried over anhydrous Na_2_SO_4_, filtered, and concentrated. The residue was purified by preparative HPLC afforded (4-(phenylamino)-1-oxa-9-azaspiro[5.5]undecan-9-yl)(1-phenylcyclopentyl)methanone **1d** as a white foam (62% yield). ^1^H NMR (500 MHz, CD_3_CN): δ 7.40–7.32 (m, 4H), 7.27–7.20 (m, 3H), 7.17 (t, *J* = 7.4 Hz, 1H), 7.11 (d, *J* = 7.9 Hz, 2H), 4.17–4.08 (m, 1H), 3.71–3.75 (m, 3H), 3.21 (br s, 1H), 2.99–2.82 (m, 3H), 2.38–2.25 (m, 2H), 2.14–2.04 (m, 1H), 1.89–1.75 (m, 3H), 1.72–1.70 (s, 4H), 1.49 (tt, *J* = 13.0, 6.5 Hz, 2H), 1.30–1.23 (m, 2H), 1.09– 0.58 (m, 2H). ^13^C NMR (126 MHz, CD_3_CN) δ 174.7, 147.0, 140.0, 130.7, 129.6, 127.0, 126.0, 125.2, 120.1, 72.1, 66.2, 59.9, 59.3, 52.4, 40.7, 31.5, 26.0, 15.6. HRMS (ESI) *m/z*: [M + H]^+^ calculated for C_27_H_35_N_2_O_2_: 419.2698, found 419.2691.

#### Synthesis of intermediates VI–VIII, IXa–j and Xa–j

To a stirred solution of *tert*-butyl 4-oxo-1-oxa-9-azaspiro[5.5]undecane-9-carboxylate **III** (18 g, 66.91 mmol, 1.0 equiv.) in DME (180 mL), EtOH (6 mL) was added, and the mixture was cooled to 0°C. TosMIC (17.61 g, 90.32 mmol, 1.35 equiv.) was then added, followed by the portion-wise addition of KO*^t^*Bu (18.73 g, 167.27 mmol, 2.5 equiv.) at 0 °C. Reaction mixture was stirred at 0 °C for 0.5 h, then allowed to stir at room temperature for another 0.5 h, and subsequently stirred at 40 °C for 0.5 h. Reaction progress was monitored by TLC, after completion, the mixture was directly filtered through celite, the pad washed with DME and filtrate concentrated under reduced pressure to obtain crude material. The resulting crude was purified through combiflash chromatography using 20% EtOAc in n-Hexane as elution system afforded *tert*-butyl 4-cyano-1-oxa-9-azaspiro[5.5]undecane-9-carboxylate (**VI**) as a white solid. (9 g, 48% yield). ^1^H NMR (400 MHz, CDCl_3_): δ 3.86–3.71 (m, 3H), 3.61–3.55 (m, 1H), 3.17–3.10 (m, 1H), 3.03–2.96 (m, 1H), 2.87 (tt, *J* = 11.0, 4.2 Hz, 1H), 2.02–1.90 (m, 2H), 1.89–1.77 (m, 2H), 1.77–1.65 (m, 2H), 1.58–1.48 (m, 1H), 1.45 (s, 9H), 1.36–1.26 (m, 1H). MS (ESI) *m*/*z*: 281.2 [M+H]^+^. Step-2: Boc deprotection of **VI** using TFA/DCM followed by amide coupling with different carboxylic acids using the General Procedures I and II afforded **VII** and **IXa**–**j**. Step-3: Reduction of carbonitriles (**VII** and **IXa**–**j**) with NiCl_2_.6H_2_O/NaBH_4_ in presence of (Boc)_2_O following the general procedure IV afforded **VIII** and **Xa**–**j**.

#### 9-(1-Phenylcyclopentane-1-carbonyl)-1-oxa-9-azaspiro[5.5]undecane-4-carbonitrile (VII)

White foam (78% yield, over two steps). ^1^H NMR (500 MHz, CDCl_3_): δ 7.32–7.27 (m, 2H), 7.23–7.15 (m, 3H), 4.35 (br s, 1H), 3.73–3.69 (m, 1H), 3.63–3.14 (m, 2H), 2.94 –2.64 (m, 3H), 2.47–2.07 (m, 3H), 1.88–1.85 (m, 2H), 1.80–1.60 (m, 7H), 1.57 (s, 2H), 1.28–1.13 (m, 1H), 0.91–0.45 (m, 1H). MS (ESI) *m*/*z*: 353.2 [M + H]^+^.

#### 9-(2-Phenylacetyl)-1-oxa-9-azaspiro[5.5]undecane-4-carbonitrile (IXa)

Colorless gum (87% yield, over two steps). ^1^H NMR (500 MHz, CDCl_3_): δ 7.33–7.28 (m, 2H), 7.25–7.20 (m, 3H), 4.36–4.32 (m, 1H), 3.80–3.75 (m, 1H), 3.73–3.69 (m, 2H), 3.63–3.46 (m, 2H), 3.36–2.84 (m, 2H), 2.80–2.74 (m, 2H), 2.17– 1.92 (m, 1H), 1.91–1.78 (m, 2H), 1.76–1.67 (m, 1H), 1.65–1.59 (m, 1H), 1.54–0.91 (m, 2H). MS (ESI) *m*/*z*: 299.2 [M + H]^+^.

#### 9-(2-Methyl-2-phenylpropanoyl)-1-oxa-9-azaspiro[5.5]undecane-4-carbonitrile (IXb)

White foam (80% yield, over two steps). ^1^H NMR (500 MHz, CDCl_3_): δ 7.44–7.30 (m, 2H), 7.25–7.21 (m, 3H), 4.38 (br s, 1H), 3.75–3.69 (m, 1H), 3.49–3.17 (m, 2H), 2.98–2.82 (m, 3H), 2.03–1.95 (m, 1H), 1.83–1.64 (m, 3H), 1.63–1.32 (m, 9H), 1.01–0.63 (m, 1H). MS (ESI) *m*/*z*: 327.2 [M + H]^+^.

#### 9-(1-Phenylcyclobutane-1-carbonyl)-1-oxa-9-azaspiro[5.5]undecane-4-carbonitrile (IXc)

White foam (86% yield, over two steps). ^1^H NMR (500 MHz, CD_3_OD): δ 7.42–7.31 (m, 4H), 7.29–7.19 (m, 1H), 4.89–4.77 (m, 1H), 4.27–4.12 (m, 1H), 3.71–3.63 (m, 1H), 3.63–3.42 (m, 1H), 3.18–3.14 (m, 1H), 3.09– 2.79 (m, 4H), 2.77–2.68 (m, 1H), 2.59–2.42 (m, 1H), 2.34–2.27 (m, 1H), 2.14–1.81 (m, 3H), 1.74–1.58 (m, 3H), 1.57–1.40 (m, 1H), 1.29–1.18 (m, 1H), 0.88–0.59 (m, 1H). MS (ESI) *m*/*z*: 339.2 [M + H]^+^.

#### 9-(4-Phenyltetrahydro-2H-pyran-4-carbonyl)-1-oxa-9-azaspiro[5.5]undecane-4-carbonitrile (IXe)

White foam (68% yield, over two steps). ^1^H NMR (500 MHz, DMSO-*d*_6_): δ 7.39–7.36 (m, 2H), 7.28–7.22 (m, 3H), 3.78–3.73 (m, 2H), 3.62–3.50 (m, 3H), 3.48–3.37 (m, 1H), 3.32–3.20 (m, 2H), 3.16–3.06 (m, 1H), 2.82–2.74 (m, 1H), 2.73–2.69 (m, 3H), 2.11–2.09 (m, 2H), 1.90–1.70 (m, 4H), 1.60–1.49 (m, 1H), 1.38 (*t*, *J* = 12.4 Hz, 1H), 0.95–0.85 (m, 2H). MS (ESI) *m*/*z*: 368.2 [M + H]^+^.

#### 9-(1-Methylcyclopentane-1-carbonyl)-1-oxa-9-azaspiro[5.5]undecane-4-carbonitrile (IXf)

White solid (90% yield, over two steps). ^1^H NMR (500 MHz, DMSO-*d*_6_): δ 3.82 (br s, 1H), 3.69–3.65 (m, 1H), 3.55 (td, *J* = 11.8, 2.6 Hz, 1H), 3.17 (tt, *J* = 11.6, 3.9 Hz, 1H), 2.97–2.69 (m, 4H), 1.93–1.82 (m, 2H), 1.67– 1.59 (m, 1H), 1.58–1.42 (m, 8H), 1.32–1.16 (m, 1H), 1.18 (s, 3H). MS (ESI) *m*/*z*: 291.2 [M + H]^+^.

#### 9-(1-(4-Chlorophenyl)cyclopentane-1-carbonyl)-1-oxa-9-azaspiro[5.5]undecane-4-carbonitrile (IXh)

White foam (55% yield, over two steps). ^1^H NMR (500 MHz, CDCl_3_): δ 7.22–7.20 (m, 2H), 7.08– 7.06 (m, 2H), 4.27 (br s, 1H), 3.66 (dt, *J* = 12.5, 4.0 Hz, 1H), 3.49–3.15 (m, 2H), 2.87–2.75 (m, 3H), 2.43– 2.33 (m, 2H), 1.93–1.78 (m, 3H), 1.74–1.61 (m, 7H), 1.57–1.46 (m, 1H), 1.44–1.23 (m, 1H), 1.15–1.07 (m, 1H), 0.93–0.57 (m, 1H). MS (ESI) *m*/*z*: 387.2 [M + H]^+^.

#### 9-(1-(p-Tolyl)cyclopentane-1-carbonyl)-1-oxa-9-azaspiro[5.5]undecane-4-carbonitrile (IXi)

White foam (65% yield, over two steps). ^1^H NMR (500 MHz, CDCl_3_): δ 7.15–7.03 (m, 4H), 4.34 (br s, 1H), 3.74– 3.70 (m, 1H), 3.59–3.12 (m, 2H), 2.93–2.82 (m, 3H), 2.50–2.36 (m, 1H), 2.32 (s, 3H), 2.13–1.86 (m, 3H), 1.83–1.39 (m, 9H), 1.37–0.43 (m, 3H). MS (ESI) *m*/*z*: 367.3 [M + H]^+^.

#### 9-(1-(3-chlorophenyl)cyclopentane-1-carbonyl)-1-oxa-9-azaspiro[5.5]undecane-4-carbonitrile (IXj)

White foam (46% yield, over two steps). ^1^H NMR (500 MHz, CDCl_3_): δ 7.33–7.29 (m, 1H), 7.26– 7.24 (m, 2H), 7.14 (dt, *J* = 7.6, 1.5 Hz, 1H), 4.43 (br s, 1H), 3.80 (dt, *J* = 12.5, 4.0 Hz, 1H), 3.59–3.51 (m, 1H), 3.28–3.03 (m, 2H), 2.90–2.81 (m, 2H), 2.62–2.34 (m, 2H), 2.09–1.89 (m, 4H), 1.87–1.71 (m, 7H), 1.67–1.37 (m, 2H), 1.02–0.61 (m, 1H). MS (ESI) *m*/*z*: 387.2 [M + H]^+^.

#### Tert-butyl ((9-(1-phenylcyclopentane-1-carbonyl)-1-oxa-9-azaspiro[5.5]undecan-4-yl)methyl)carbamate VIII

Reduction of carbonitrile **VII** using the General Procedure IV afforded **VIII** as a white foam (83% yield). ^1^H NMR (700 MHz, CDCl_3_): δ 7.29 (t, *J* = 7.5 Hz, 2H), 7.21–7.18 (m, 3H), 4.55 (br s, 1H), 4.35–4.24 (m, 1H), 3.68 (dd, *J* = 12.6, 5.1 Hz, 1H), 3.53–3.34 (m, 1H), 3.21–3.09 (m, 1H), 2.97– 2.80 (m, 3H), 2.53–2.45 (m, 1H), 2.39–2.26 (m, 1H), 2.13–2.03 (m, 1H), 1.93–1.85 (m, 1H), 1.79 – 1.67 (m, 6H), 1.57–1.49 (m, 2H), 1.42 (s, 9H), 1.36–1.34 (m, 1H), 1.17–1.09 (m, 2H), 1.07–0.92 (m, 1H), 0.91– 0.59 (m, 2H). MS (ESI) *m*/*z*: 457.3 [M + H]^+^.

#### Tert-butyl ((9-(2-phenylacetyl)-1-oxa-9-azaspiro[5.5]undecan-4-yl)methyl)carbamate (Xa)

Reduction of carbonitrile **IXa** using the General Procedure-IV afforded **Xa** as a white foam (69% yield). ^1^H NMR (500 MHz, CDCl_3_): δ 7.26–7.20 (m, 2H), 7.18–7.13 (m, 3H), 4.67 (br s, 1H), 4.31–4.12 (m, 1H), 3.71–3.62 (m, 3H), 3.55–3.27 (m, 3H), 3.20–2.96 (m, 1H), 2.94–2.78 (m, 3H), 2.09 (d, *J* = 15.2 Hz, 1H), 1.75 (s, 1H), 1.55–1.47 (m, 1H), 1.45–1.39 (m, 1H), 1.36 (s, 9H), 1.33–1.32 (m, 1H), 1.10 (td, *J* = 12.6, 5.1 Hz, 2H), 0.92 (s, 1H). MS (ESI) *m*/*z*: 403.3 [M + H]^+^.

#### Tert-butyl ((9-(2-methyl-2-phenylpropanoyl)-1-oxa-9-azaspiro[5.5]undecan-4-yl)methyl)carbamate (Xb)

Reduction of carbonitrile **IXb** using the General Procedure-IV afforded **Xb** as a white foam (50% yield). ^1^H NMR (500 MHz, CDCl_3_): δ 7.34–7.30 (m, 2H), 7.23–7.19 (m, 3H), 4.55–4.33 (m, 2H), 3.73– 3.61 (m, 1H), 3.49–3.34 (m, 1H), 3.10–2.82 (m, 5H), 1.90–1.60 (m, 5H), 1.55–1.52 (m, 6H), 1.43 (s, 9H), 1.40–1.36 (m, 1H), 1.16–1.07 (m, 1H), 0.96–0.86 (m, 1H). MS (ESI) *m*/*z*: 431.3 [M + H]^+^.

#### Tert-butyl ((9-(1-phenylcyclobutane-1-carbonyl)-1-oxa-9-azaspiro[5.5]undecan-4-yl)methyl)carbamate (Xc)

Reduction of carbonitrile **IXc** using the General Procedure-IV afforded **Xc** as a white foam (58% yield). ^1^H NMR (500 MHz, CDCl_3_): δ 7.39–7.29 (m, 4H), 7.25–7.17 (m, 1H), 4.54 (br s, 1H), 4.27 (d, *J* = 13.2 Hz, 1H), 3.69 (d, *J* = 11.5 Hz, 1H), 3.53–3.35 (m, 1H), 3.10–2.97 (m, 2H), 2.91–2.78 (m, 3H), 2.83–2.69 (m, 1H), 2.55–2.36 (m, 1H), 2.31–2.18 (m, 1H), 2.15–2.05 (m, 1H), 1.99 (q, *J* = 9.2 Hz, 1H), 1.99–1.85 (m, 1H), 1.82–1.64 (m, 2H), 1.59–1.47 (m, 2H), 1.42 (s, 9H), 1.34–1.30 (m, 1H), 1.16–1.06 (m, 2H), 1.00–0.82 (m, 1H), 0.81–0.48 (m, 1H). MS (ESI) *m*/*z*: 443.3 [M + H]^+^.

#### Tert-butyl ((9-(1-phenylcyclohexane-1-carbonyl)-1-oxa-9-azaspiro[5.5]undecan-4-yl)methyl)carbamate (Xd)

Boc deprotection of **VI** using TFA/DCM followed by amide coupling with 1-phenylcyclohexane-1-carboxylic acid using the General Procedures-I and -II afforded carbonitrile **IXd** (MS (ESI) *m*/*z*: 367.2 [M + H]^+^) which was 80% pure and used directly in the next step. Reduction of **IXd** with NiCl_2_.6H_2_O/NaBH_4_ in presence of (Boc)_2_O following the general procedure-IV afforded **Xd** as a white solid (35% yield, over three steps). ^1^H NMR (500 MHz, CDCl_3_): δ 7.33–7.26 (m, 3H), 7.25–7.15 (m, 1H), 7.21–7.18 (m, 1H), 4.54 (br s, 1H), 4.24–4.11 (m, 2H), 3.67 (dd, *J* = 12.0, 4.9 Hz, 1H), 3.44–3.40 (m, 1H), 3.00–2.83 (m, 4H), 2.33–2.24 (m, 2H), 1.82–1.61 (m, 8H), 1.60–1.50 (m, 3H), 1.43–1.34 (m, 10H), 1.31– 1.23 (m, 2H), 1.11 (qd, *J* = 12.6, 5.0 Hz, 2H), 0.94–0.86 (m, 1H). MS (ESI) *m*/*z*: 471.4 [M + H]^+^.

#### Tert-butyl (9-(4-phenyltetrahydro-2H-pyran-4-carbonyl)-1-oxa-9-azaspiro[5.5]undecan-4-yl)methyl)carbamate (Xe)

Reduction of carbonitrile **IXe** using the General Procedure IV afforded **Xe** as a white foam (68% yield). ^1^H NMR (500 MHz, CDCl_3_): δ 7.37–7.30 (m, 2H), 7.28–7.19 (m, 3H), 4.60 (s, 1H), 4.28–4.26 (m, 1H), 3.95–3.71 (m, 4H), 3.70–3.61 (m, 1H), 3.49–3.42 (m, 1H), 3.30–2.86 (m, 4H), 2.81 (s, 2H), 2.24–2.17 (m, 2H), 2.03 –1.74 (m, 2H), 1.53 (d, *J* = 13.6 Hz, 1H), 1.42 (s, 9H), 1.36–1.22 (m, 2H), 1.10 (qd, *J* = 12.7, 5.1 Hz, 1H), 0.95–0.81 (m, 1H), 0.73–0.54 (m, 1H). MS (ESI) *m*/*z*: 473.3 [M + H]^+^.

#### Tert-butyl ((9-(1-methylcyclopentane-1-carbonyl)-1-oxa-9-azaspiro[5.5]undecan-4-yl)methyl)carbamate (Xf)

Reduction of carbonitrile **IXf** using the General Procedure-IV afforded **Xf** as a white solid (54% yield). ^1^H NMR (500 MHz, CDCl_3_): δ 4.61 (br s, 1H), 4.13–3.97 (m, 1H), 3.80–3.76 (m, 1H), 3.57 (td, *J* = 12.3, 2.4 Hz, 1H), 3.28–2.96 (m, 4H), 2.20–2.14 (m, 3H), 1.94–1.80 (m, 1H), 1.66–1.51 (m, 10H), 1.50–1.48 (m, 1H), 1.43 (s, 9H), 1.28–1.25 (m, 4H), 1.23–1.16 (m, 1H), 1.05 (t, *J* = 12.8 Hz, 1H). MS (ESI) *m*/*z*: 395.3 [M + H]^+^.

#### Tert-butyl ((9-(1-(4-methoxyphenyl)cyclopentane-1-carbonyl)-1-oxa-9-azaspiro[5.5]undecan-4-yl)methyl)carbamate (Xg)

Boc deprotection of **VI** using TFA/DCM followed by amide coupling with 1-(3-methoxyphenyl)cyclopentane-1-carboxylic acid using the General Procedures-I and -II afforded carbonitrile **IXg** (MS (ESI) *m*/*z*: 383.3 [M + H]^+^)which was 85% pure and used directly in the next step. Reduction of **IXg** with NiCl_2_.6H_2_O/NaBH_4_ in presence of (Boc)_2_O following the general procedure-IV afforded **Xg** as a white foam (21% yield, over three steps). ^1^H NMR (500 MHz, CDCl_3_): δ 7.12–7.10 (m, 2H), 6.83 (d, *J* = 8.7 Hz, 2H), 4.56 (br s, 1H), 4.30–4.27 (s, 1H), 3.79 (s, 3H), 3.69 (dd, *J* = 12.1, 4.8 Hz, 1H), 3.49–3.39 (s, 1H), 3.25–3.08 (m, 2H), 2.93–2.84 (m, 2H), 2.42–2.30 (m, 2H), 2.07–2.00 (m, 1H), 1.85–1.70 (m, 2H), 1.70 (s, 4H), 1.56–1.50 (m, 1H), 1.42 (s, 9H), 1.39–1.35 (m, 1H), 1.25 (s, 2H), 1.12 (qd, *J* = 12.7, 5.1 Hz, 2H), 0.92–0.65 (m, 3H). MS (ESI) *m*/*z*: 487.3 [M + H]^+^.

#### Tert-butyl ((9-(1-(4-chlorophenyl)cyclopentane-1-carbonyl)-1-oxa-9-azaspiro[5.5]undecan-4-yl)methyl)carbamate (Xh)

Reduction of carbonitrile **IXh** using the General Procedure-IV afforded **Xh** as a white foam (55% yield). ^1^H NMR (500 MHz, CDCl_3_): δ 7.29–7.23 (m, 2H), 7.13 (d, *J* = 8.5 Hz, 2H), 4.57–4.28 (m, 2H), 3.69 (dd, *J* = 11.8, 4.9 Hz, 1H), 3.57–3.36 (m, 1H), 3.16–3.07 (m, 1H), 2.92–2.82 (m, 3H), 2.41 (s, 2H), 2.17–1.79 (m, 4H), 1.76–1.65 (m, 5H), 1.63–1.49 (m, 2H), 1.42 (s, 9H), 1.41–1.35 (m, 1H), 1.22–1.07 (m, 2H), 0.92–0.71 (m, 2H). MS (ESI) *m*/*z*: 491.2 [M + H]^+^.

#### Tert-butyl ((9-(1-(p-tolyl)cyclopentane-1-carbonyl)-1-oxa-9-azaspiro[5.5]undecan-4-yl)methyl)carbamate (Xi)

Reduction of carbonitrile **IXi** using the General Procedure-IV afforded **Xi** as a white foam (60% yield). ^1^H NMR (500 MHz, CDCl_3_): δ 7.10–7.06 (m, 4H), 4.59–4.27 (m, 2H), 3.68 (dd, *J* = 12.3, 5.2 Hz, 1H), 3.55–3.32 (m, 1H), 3.22 (s, 1H), 3.14–2.74 (m, 4H), 2.43 (s, 1H), 2.30 (s, 4H), 2.18– 1.94 (m, 2H), 1.87 (s, 2H), 1.76–1.63 (m, 5H), 1.53 (d, *J* = 12.8 Hz, 1H), 1.42 (s, 9H), 1.39–1.35 (m, 1H), 1.31–1.07 (m, 2H), 1.01–0.56 (m, 2H). MS (ESI) *m*/*z*: 471.3 [M + H]^+^.

#### Tert-butyl ((9-(1-(3-chlorophenyl)cyclopentane-1-carbonyl)-1-oxa-9-azaspiro[5.5]undecan-4-yl)methyl)carbamate (Xj)

Reduction of carbonitrile **IXj** using the General Procedure-IV afforded **Xj** as a white foam (52% yield). ^1^H NMR (500 MHz, CDCl_3_): δ 7.26–7.15 (m, 3H), 7.0–7.07 (m, 1H), 4.56–4.31 (m, 2H), 3.73–3.51–3.39 (m, 1H), 3.28–2.73 (m, 5H), 2.53–2.30 (m, 2H), 2.20–1.97 (m, 2H), 1.85–1.83 (m, 1H), 1.77–1.64 (m, 6H), 1.55 (d, *J* = 7.4 Hz, 1H), 1.43 (s, 10H), 1.13 (qd, *J* = 12.7, 5.1 Hz, 2H), 1.01– 0.65 (m, 2H). MS (ESI) *m*/*z*: 491.2[M + H]^+^.

#### Synthesis of intermediates XI–XIV

To a stirred solution of *tert*-butyl 4-cyano-1-oxa-9-azaspiro[5.5]undecane-9-carboxylate **VI** (10 g, 35.714 mmol, 1.0 eq.) in MeOH (10 volumes) at 25 °C was added Raney Ni (approx. 2.5 g) and then added Aq. Ammonia (0.5 volume). Reaction mixture was stirred at 70 °C for 15 hours. Reaction progress was monitored by TLC, after completion, the mixture was filtered through high flow celite and washed with MeOH. The filtrate was concentrated to obtain crude. The resulting crude was purified through reverse phase purification using C18 column and Acetonitrile in 0.2% NH_3_ in H_2_O as eluent. The pure fraction was collected and lyophilized to afford *tert*-butyl 4-(aminomethyl)-1-oxa-9-azaspiro[5.5]undecane-9-carboxylate **XI** as pale yellow sticky liquid (2.3 g, 23% yield). ^1^H NMR (400 MHz, CDCl_3_): δ 3.86–3.50 (m, 4H), 3.20 (br s, 1H), 3.02 (br s, 1H), 2.55 (d, J = 6.3 Hz, 2H), 2.14 (d, J = 14.2 Hz, 1H), 1.65 (d, J = 14.0 Hz, 2H), 1.61–1.49 (m, 5H), 1.45 (s, 9H), 1.32–1.24 (m, 1H), 1.23–1.11 (m, 1H), 1.03 (t, J = 12.7 Hz, 1H). MS (ESI) *m*/*z*: 185.3 [M + H-100]^+^. Step-2: Sulfonylation of **XI** with *N,N*-Dimethylsulfamoyl chloride using the General Procedure-V furnished *Tert*-butyl 4-(((*N*,*N*-dimethylsulfamoyl)amino)methyl)-1-oxa-9-azaspiro[5.5]undecane-9-carboxylate **XII** as a colorless gum (80% yield). MS (ESI) *m*/*z*: 292.0 [M + H]^+^.

#### Tert-butyl4-((1,1-dioxido-1,2-thiazinan-2-yl)methyl)-1-oxa-9-azaspiro[5.5]undecane-9-carboxylate (XIV)

Sulfonylation of **XI** with 4-chloro-1-butylsulfonylchloride using the General Procedure-V to afford *Tert*-butyl 4-(((4-chlorobutyl)sulfonamido)methyl)-1-oxa-9-azaspiro[5.5]undecane-9-carboxylate **XIII** as a colorless gum (88% yield). ^1^H NMR (500 MHz, CDCl_3_): δ 3.77–3.67 (m, 3H), 3.52 (td, *J* = 12.3, 2.4 Hz, 1H), 3.37–3.31 (m, 2H), 3.20–3.08 (m, 1H), 3.05–2.77 (m, 6H), 2.21–2.13 (m, 2H), 2.09 (dd, *J* = 14.1, 3.7 Hz, 1H), 1.93–1.89 (m, 1H), 1.69–1.52 (m, 4H), 1.48–1.46 (m, 2H), 1.42 (s, 9H), 1.29–1.10 (m, 2H), 1.00 (t, *J* = 12.8 Hz, 1H). MS (ESI) *m*/*z*: 439.2 [M + H]^+^. Step-2: Intramolecular cyclization of intermediate **XIII** using the General Procedure-VI with Cs_2_CO_3_ afforded *Tert*-butyl 4-((1,1-dioxido-1,2-thiazinan-2-yl)methyl)-1-oxa-9-azaspiro[5.5]undecane-9-carboxylate **XIV** as a white solid (83% yield). ^1^H NMR (500 MHz, CDCl_3_): δ 3.81–3.68 (m, 3H), 3.55 (tq, *J* = 12.4, 2.3 Hz, 1H), 3.36 (d, *J* = 5.3 Hz, 2H), 3.21–3.14 (m, 1H), 3.05–2.84 (m, 5H), 2.22–2.14 (m, 2H), 2.12–2.09 (m, 1H), 1.95–1.89 (m, 1H), 1.70–1.54 (m, 4H), 1.52 –1.48 (m, 2H), 1.44 –1.43(m, 9H), 1.32–1.12 (m, 2H), 1.05–0.99 (m, 1H). MS (ESI) *m*/*z*: 403.2 [M + H]^+^.

#### Synthesis of 2a–h, 5a–e and 6a–e

Sulfonylation of intermediate (**VIII**) following the General Procedure-V afforded sulfonamides (**2a***–***h**)

#### N-((9-(1-Phenylcyclopentane-1-carbonyl)-1-oxa-9-azaspiro[5.5]undecan-4-yl)methyl)methanesulfonamide (2a)

White foam (71% yield). ^1^H NMR (500 MHz, CD_3_CN): δ 7.36–7.29 (m, 2H), 7.23–7.19 (m, 3H), 5.11 (d, *J* = 8.4 Hz, 1H), 4.14 (br s, 1H), 3.61 (dd, *J* = 11.9, 5.0 Hz, 1H), 3.46–3.45 (m, 1H), 3.18–2.97 (m, 2H), 2.83 (s, 3H), 2.82–2.74 (m, 2H), 2.38–2.26 (m, 2H), 2.14–2.04 (m, 3H), 1.74–1.51 (m, 6H), 1.47–1.43 (m, 2H), 1.27–1.18 (m, 1H), 1.05 (qd, *J* = 12.6, 5.2 Hz, 1H), 0.88–0.86 (m, 3H). ^13^C NMR (126 MHz, CD_3_CN): δ 174.6, 147.2, 129.6, 127.0, 126.1, 71.0, 60.8, 59.3, 49.8, 40.5, 39.9, 31.7 31.0, 26.1, 26.1. HRMS (ESI) *m/z*: [M + H]^+^ calculated for C_23_H_35_N_2_O_4_S: 435.2317, found 435.2311.

#### N-((9-(1-Phenylcyclopentane-1-carbonyl)-1-oxa-9-azaspiro[5.5]undecan-4-yl)methyl)ethanesulfonamide (2b)

White foam (71% yield). ^1^H NMR (400 MHz, CD_3_CN): δ 7.34–7.30 (m, 2H), 7.23–7.19 (m, 3H), 5.14 (t, *J* = 6.5 Hz, 1H), 4.14 (br s, 1H), 3.60 (dd, *J* = 12.0, 5.0 Hz, 1H), 3.53–3.33 (m, 1H), 3.25 −3.13 (m, 1H), 2.84 –2.74(m, 3H), 2.84–2.74 (m, 3H), 2.58–2.22 (m, 4H), 2.12–2.01 (m, 1H), 1.70–1.66 (m, 5H), 1.56 (d, *J* = 13.3 Hz, 1H), 1.49–1.35 (m, 2H), 1.22 (t, *J* = 7.4 Hz, 3H), 1.16–0.97 (m, 2H), 0.88–0.73 (m, 2H). ^13^C NMR (126 MHz, CD_3_CN): δ 174.6, 147.1, 129.5, 126.9, 126.0, 71.0, 60.7, 59.3, 49.6, 46.6, 40.5, 30.9, 26.0, 8.5. HRMS (ESI) *m/z*: [M + H]^+^ calculated for C_24_H_37_N_2_O_4_S: 449.2474, found 449.2466.

#### 3- Chloro-N-((9-(1-phenylcyclopentane-1-carbonyl)-1-oxa-9-azaspiro[5.5]undecan-4-yl)methyl)propane-1-sulfonamide (2c)

White foam (64% yield). ^1^H NMR (500 MHz, CD_3_CN): δ 7.34– 7.31 (m, 2H), 7.24–7.18 (m, 3H), 5.32–5.13 (m, 1H), 4.14 (br s, 1H), 3.68 (t, *J* = 6.4 Hz, 2H), 3.61 (dd, *J* = 12.0, 5.0 Hz, 1H), 3.56–3.37 (m, 1H), 3.21–3.18 (m, 1H), 3.11–3.07 (m, 2H), 3.04–2.90 (m, 1H), 2.88– 2.74 (m, 3H), 2.40–2.30 (m, 2H), 2.18–2.08 (m, 3H), 2.07–2.-5 (m, 1H), 1.88–1.78 (m, 1H), 1.72–1.64 (m, 4H), 1.56 (d, *J* = 13.0 Hz, 1H), 1.47–1.43 (m, 2H), 1.31–1.11 (m, 1H), 1.04 (qd, *J* = 12.6, 5.1 Hz, 1H), 0.99–0.65 (m, 3H). ^13^C NMR (126 MHz, CD_3_CN): δ 174.6, 147.2, 129.6, 126.9, 126.0, 71.0, 60.7, 59.3, 49.6, 44.1, 40.5, 31.8, 31.0, 27.9, 26.1, 26.1. HRMS (ESI) *m/z*: [M + H]^+^ calculated for C_25_H_38_ClN_2_O_4_S: 497.2241, found 497.2235.

#### 4- Chloro-N-((9-(1-phenylcyclopentane-1-carbonyl)-1-oxa-9-azaspiro[5.5]undecan-4-yl)methyl)butane-1-sulfonamide (2d)

White foam (60% yield). ^1^H NMR (500 MHz, CD_3_CN): δ 7.36–7.28 (m, 2H), 7.26–7.16 (m, 3H), 5.18 (d, *J* = 6.8 Hz, 1H), 4.14–4.05 (m, 1H), 3.61 (t, *J* = 6.1 Hz, 3H), 3.47–3.42 (m, 1H), 3.18–3.14 (m, 1H), 2.98 (dd, *J* = 8.3, 6.4 Hz, 3H), 2.88–2.70 (m, 3H), 2.38–2.30 (m, 2H), 2.13–1.98 (m, 1H), 1.90–1.81 (m, 5H), 1.75–1.62 (m, 5H), 1.56 (d, *J* = 12.9 Hz, 1H), 1.47–1.44 (m, 2H), 1.27–1.13 (m, 1 H), 1.04 (qd, *J* = 12.6, 5.2 Hz, 1H), 0.89–0.72 (m, 3H). ^13^C NMR (126 MHz, CD_3_CN): δ 174.6, 147.2, 129.6, 126.9, 126.0, 71.0, 60.7, 59.3, 51.4, 45.5, 40.5, 31.6, 31.0, 26.1, 26.0, 21.9. HRMS (ESI) *m/z*: [M + H]^+^ calculated for C_26_H_40_ClN_2_O_4_: 511.2397, found 511.2404.

#### N-((9-(1-Phenylcyclopentane-1-carbonyl)-1-oxa-9-azaspiro[5.5]undecan-4-yl)methyl)cyclopropanesulfonamide (2e)

White foam (59% yield). ^1^H NMR (400 MHz, CDCl_3_): δ 7.32–7.30 (m, 2H), 7.23–7.15 (m, 3H), 4.39–4.17 (m, 2H), 3.71 (dd, *J* = 12.0, 4.8 Hz, 1H), 3.58–3.33 (m, 1H), 3.24–3.22 (m, 1H), 3.12–2.75 (m, 4H), 2.57–2.28 (m, 3H), 2.09–2.04 (m, 1H), 1.89–1.59 (m, 11H), 1.46– 1.41 (m, 1H), 1.21–1.11 (m, 3H), 1.04–0.94 (m, 2H), 0.83–0.62 (m, 1H). ^13^C NMR (100 MHz, CDCl_3_): δ 174.7, 145.9, 128.8, 126.3, 125.2, 70.4, 60.3, 58.7, 49.4, 40.1, 31.5, 30.2, 25.4, 5.5 (2). HRMS (ESI) *m/z*: [M + H]^+^ calculated for C_25_H_37_N_2_O_4_S: 461.2474, found 461.2468.

#### N-((9-(1-Phenylcyclopentane-1-carbonyl)-1-oxa-9-azaspiro[5.5]undecan-4-yl)methyl)benzenesulfonamide (2f)

White foam (43% yield). ^1^H NMR (400 MHz, CD_3_CN): δ 7.81–7.70 (m, 2H), 7.61–7.46 (m, 3H), 7.30 (t, *J* = 7.6 Hz, 2H), 7.23–7.12 (m, 3H), 5.48 (t, *J* = 6.5 Hz, 1H), 4.12 (br s, 1H), 3.56 (dd, *J* = 11.6, 4.9 Hz, 1H), 3.39–3.30 (m, 1H), 3.22–2.69 (m, 3H), 2.68–2.57 (m, 2H), 2.31– 2.11 (m, 5H), 1.93–1.77 (m, 1H), 1.68–1.67 (m, 4H), 1.49–1.26 (m, 3H), 1.11–0.54 (m, 4H). ^13^C NMR (126 MHz, CDCl_3_): δ 174.7, 140.1, 132.9, 129.3, 128.8, 126.3, 125.2, 70.3, 60.2, 58.7, 49.2, 40.0, 38.5, 31.1, 30.2, 25.4. HRMS (ESI) *m/z*: [M + H]^+^ calculated for C_28_H_37_N_2_O_4_S: 497.2474, found 497.2472.

#### {4-[(Dimethylaminosulfonylamino)methyl]-1-oxa-9-aza-9-spiro[5.5]undecyl}(1-phenylcyclopentyl)methanone (2g)

White foam (72% yield). ^1^H NMR (400 MHz, CD_3_CN): δ 7.37–7.33 (m, 2H), 7.27– 7.20 (m, 3H), 5.27–5.14 (m, 1H), 4.17 (br s, 1H), 3.63 (dd, *J* = 12.0, 4.8 Hz, 1H), 3.47–3.42 (m, 1H), 3.21–3.10 (m, 2H), 2.86–2.74 (m, 3H), 2.72 (s, 6H), 2.54–2.32 (m, 5H), 2.14–2.05 (m, 1H), 1.78–1.70 (m, 5H), 1.60 –1.56 (m, 1H), 1.47–1.46 (m, 1H), 1.13–0.87 (m, 4H). ^13^C NMR (126 MHz, CD_3_CN): δ 174.6, 147.1, 129.6, 127.0, 126.0, 71.0, 60.8, 59.3, 50.2, 40.6, 38.4, 31.6, 31.0, 26.1. HRMS (ESI) *m/z*: [M + H]^+^ calculated for C_24_H_38_N_3_O_4_S: 464.2583, found 464.2579.

#### {4-[(N-Ethyl-N-methylaminosulfonylamino)methyl]-1-oxa-9-aza-9-spiro[5.5]undecyl}(1-phenylcyclopentyl)methanone (2h)

White foam (48% yield). ^1^H NMR (500 MHz, CDCl_3_): δ 7.30 (t, *J* = 7.6 Hz, 2H), 7.23–7.15 (m, 3H), 4.31 (br s, 1H), 3.70 (dd, *J* = 12.0, 4.9 Hz, 1H), 3.52–3.49 (m, 1H), 3.39–3.36 (m, 1H), 3.20 (q, *J* = 7.2 Hz, 3H), 2.97–2.96 (m, 1H), 2.84–2.79 (m, 2H), 2.77 (s, 3H), 2.56–2.26 (m, 2H), 2.17–2.08 (m, 1H), 1.91–1.89 (m, 2H), 1.79–1.69 (m, 4H), 1.69–1.58 (m, 3H), 1.45–1.38 (m, 1H), 1.23–1.09 (m, 5H), 1.00–0.77 (m, 2H), 0.71–0.64 (s, 1H). ^13^C NMR (126 MHz, CDCl_3_): δ 174.6, 146.0, 128.8, 126.3, 125.2, 70.4, 60.5, 60.3, 58.7, 49.4, 45.4, 40.2, 34.3, 30.9, 30.4, 25.4, 21.2, 14.3, 13.2. HRMS (ESI) *m/z*: [M + H]^+^ calculated for C_25_H_40_N_3_O_4_S: 478.2739, found 478.2733. Boc deprotection of intermediates (**Xa–j**) followed by sulfonylation with various sulfonyl chlorides following the General Procedures I and V afforded sulfonamides (**5a–e** and **6a–e**)

#### N-((9-(Phenylacetyl)-1-oxa-9-azaspiro[5.5]undecan-4-yl)methyl)methanesulfonamide (5a)

The compound **5a** was prepared from intermediate **Xa**. White foam (65% yield). ^1^H NMR (500 MHz, CD_3_CN): δ 7.33–7.30 (m, 2H), 7.27–7.20 (m, 3H), 5.28–5.25 (m, 1H), 4.14–4.09 (m, 1H), 3.73–3.65 (m, 3H), 3.64– 3.57 (m, 1H), 3.56–3.50 (m, 1H), 3.39–2.99 (m, 2H), 2.91–2.85 (m, 4H), 2.83–2.79 (m, 1H), 2.21–2.04 (m, 1H), 1.92–1.77 (m, 1H), 1.67–1.51 (m, 2H), 1.49–1.31 (m, 2H), 1.26–1.05 (m, 2H), 0.99–0.92 (m, 1H). ^13^C NMR (126 MHz, CD_3_CN): δ 170.4, 137.0, 129.8, 129.4, 127.4, 70.9, 60.9, 60.8, 49.8, 42.7, 42.6, 40.8, 40.6, 40.5, 40.3, 39.8, 39.7, 38.4, 38.3, 31.7, 31.6, 31.0, 30.2, 29.6. (additional carbon signals observed due to rotamers). HRMS (ESI) *m/z*: [M + H]^+^ calculated for C_19_H_29_N_2_O_4_S: 381.1848, found 381.1841.

#### N-((9-(2-Methyl-2-phenylpropanoyl)-1-oxa-9-azaspiro[5.5]undecan-4-yl)methyl)methanesulfonamide (5b)

The compound **5b** was prepared from intermediate **Xb**. White foam (71% yield). ^1^H NMR (500 MHz, CD_3_CN): δ 7.36–7.33 (m, 2H), 7.26–7.21 (m, 3H), 5.21–5.08 (m, 1H), 4.16 (br s, 1H), 3.65–3.56 (m, 1H), 3.54–3.40 (m, 1H), 3.19–2.89 (m, 2H), 2.83 (s, 3H), 2.82–2.76 (m, 2H), 2.51–2.33 (m, 2H), 1.86–1.67 (m, 1H), 1.50–1.45 (m, 1H), 1.49 (dd, *J* = 3.7, 2.0 Hz, 8H), 1.10–0.70 (m, 4H). ^13^C NMR (126 MHz, CD_3_CN): δ 174.8, 147.8, 129.7, 127.1, 125.8, 70.9, 60.7, 49.8, 47.6, 40.5, 39.8, 31.7, 31.0, 29.5. HRMS (ESI) *m/z*: [M + H]^+^ calculated for C_21_H_33_N_2_O_4_S: 409.2161, found 409.2152.

#### N-((9-(1-Phenylcyclobutane-1-carbonyl)-1-oxa-9-azaspiro[5.5]undecan-4-yl)methyl)methanesulfonamide (5c)

The compound **5c** was prepared from intermediate **Xc**. White foam (60% yield). ^1^H NMR (500 MHz, CD_3_CN): δ 7.39–7.35 (m, 4H), 7.28–7.21 (m, 1H), 5.12 (br s, 1H), 4.12– 4.07 (m, 1H), 3.64–3.54 (m, 1H), 3.53–3.38 (m, 1H), 3.05–2.96 (m, 2H), 2.88–2.80 (m, 5H), 2.78–2.66 (m, 2H), 2.42–2.22 (m, 1H), 2.29–2.15 (m, 1H), 2.12–1.95 (m, 2H), 1.89–1.65 (m, 3H), 1.56 (t, *J* = 14.9 Hz, 1H), 1.44– 1.38 (m, 2H), 1.12 (t, *J* = 7.0 Hz, 1H), 1.07–0.98 (m, 1H), 0.92–0.62 (m, 2H). ^13^C NMR (126 MHz, CD_3_CN): δ 174.3, 145.0, 129.7, 127.2, 125.8, 70.9, 60.7, 53.1, 49.7, 41.9, 41.8, 40.5, 40.4, 39.8, 39.6, 39.4, 38.6, 38.5, 33.6, 33.5, 32.6, 32.4, 31.6, 31.6, 30.9, 29.5, 29.3. HRMS (ESI) *m/z*: [M + H]^+^ calculated for C_22_H_33_N_2_O_4_S: 421.2161, found 421.2150.

#### N-((9-(1-Phenylcyclohexane-1-carbonyl)-1-oxa-9-azaspiro[5.5]undecan-4-yl)methyl)methanesulfonamide (5d)

The compound **5d** was prepared from intermediate **Xd**. White foam (65% yield). ^1^H NMR (500 MHz, CD_3_CN): δ 7.36–7.31 (m, 2H), 7.29–7.24 (m, 2H), 7.24–7.19 (m, 1H), 5.13–5.10 (m, 1H), 4.26–3.65 (m, 1H), 3.63–3.44 (m, 2H), 3.01–2.90 (m, 1H), 2.88–2.69 (m, 6H), 2.26– 2.04 (m, 5H), 1.76 –1.51(m, 9H), 1.47–1.44 (m, 1H), 1.30–1.25 (m, 2H), 1.04 (qd, *J* = 12.6, 5.1 Hz, 2H), 0.85 (t, *J* = 12.8 Hz, 1H). ^13^C NMR (126 MHz, CD_3_CN) :δ 173.7, 147.6, 129.7, 127.1, 126.1, 70.9, 60.7, 51.9, 49.7, 40.4, 39.8, 39.7, 31.6, 30.9, 29.6, 26.5, 24.4 (2C). HRMS (ESI) *m/z*: [M + H]^+^ calculated for C_24_H_37_N_2_O_4_S: 449.2474.

#### N-((9-(4-Phenyltetrahydro-2H-pyran-4-carbonyl)-1-oxa-9-azaspiro[5.5]undecan-4-yl)methyl)methanesulfonamide (5e)

The compound **5e** was prepared from intermediate **Xe**. White foam (63% yield). ^1^H NMR (500 MHz, DMSO-*d*_6_): δ 7.39–7.36 (m, 2H), 7.31–7.20 (m, 3H), 6.92 (t, *J* = 6.1 Hz, 1H), 4.11–4.01 (m, 1H), 3.78–3.74 (m, 2H), 3.65–3.53 (m, 3H), 3.38 (br s, 1H), 3.30–3.16 (m, 2 H), 2.97– 2.93 (m, 1H), 2.84 (s, 3H), 2.75–2.65 (m, 3H), 2.17–2.06 (m, 2H), 1.99–1.62 (m, 4H), 1.54–1.40 (m, 2H), 1.02–0.55 (m, 4H). ^13^C NMR (126 MHz, DMSO-*d*_6_): δ 171.4, 145.0, 129.0, 126.6, 124.9, 69.7, 64.6, 64.6, 59.5, 48.6, 48.4, 48.3, 30.3, 30.0, 28.5. HRMS (ESI) *m/z*: [M + H]^+^ calculated for C_23_H_35_N_2_O_5_S: 451.2266, found 451.2256.

#### N-((9-(1-Methylcyclopentane-1-carbonyl)-1-oxa-9-azaspiro[5.5]undecan-4-yl)methyl)methanesulfonamide (6a)

The compound 6a was prepared from intermediate **Xf**. White solid (65% yield). 1H NMR (500 MHz, CD_3_CN) δ 5.19–5.16 (m, 1H), 3.97–3.89 (m, 2H), 3.70 (ddd, J = 11.9, 5.2, 1.6 Hz, 1H), 3.59 (td, J = 12.3, 2.4 Hz, 1H), 3.20–2.82 (m, 7H), 2.19–2.04 (m, 3H), 1.91–1.84 (m, 1H), 1.66 –1.46 (m, 10H), 1.35–1.25 (m, 1H), 1.23 (s, 3H), 1.19–1.07 (m, 1H), 0.99 (t, J = 12.9 Hz, 1H). ^13^C NMR (126 MHz, CD_3_CN) δ 176.7, 71.2, 60.8, 49.9, 49.8, 40.7, 39.8, 39.2, 31.8, 31.1, 30.2, 26.6, 26.0. HRMS (ESI) *m/z*: [M + H]^+^ calculated for C_18_H_32_N_2_O_4_S: 373.2161, found 373.2153.

#### N-((9-(1-(4-Methoxyphenyl)cyclopentane-1-carbonyl)-1-oxa-9-azaspiro[5.5]undecan-4-yl)methyl)methanesulfonamide (6b)

The compound **6b** was prepared from intermediate **Xg**. White foam (53% yield). ^1^H NMR (500 MHz, CD_3_CN): δ 7.16–7.09 (m, 2H), 6.90–6.83 (m, 2H), 5.12 (br s, 1H), 4.13 (br s, 1H), 3.76 (s, 3H), 3.63–3.60 (m, 1H), 3.47–3.40 (m, 1H), 3.28–3.22 (m, 1H), 3.04–2.92 (m, 1H), 2.86–2.76 (m, 5H), 2.33–2.22 (m, 7H), 2.11–2.01 (m, 2H), 1.85–1.78 (m, 2H), 1.66 (s, 3H), 1.56 (d, *J* = 13.0 Hz, 1H), 1.48–1.45(m, 1H), 1.09–1.01 (m, 1H), 0.87–0.69 (m, 1H). ^13^C NMR (126 MHz, CD_3_CN): δ 174.7, 158.8, 139.1, 127.2, 114.8, 71.0, 60.7, 58.6, 55.8, 49.8, 40.6, 31.7, 31.0, 25.9. HRMS (ESI) *m/z*: [M + H]^+^ calculated for C_24_H_37_N_2_O_5_S: 465.2423 found 465.2413.

#### N-((9-(1-(4-Chlorophenyl)cyclopentane-1-carbonyl)-1-oxa-9-azaspiro[5.5]undecan-4-yl)methyl)methanesulfonamide (6c)

The compound **6c** was prepared from intermediate **Xh**. White foam (67% yield). ^1^H NMR (500 MHz, CD_3_CN): δ 7.36–7.29 (m, 2H), 7.23–7.16 (m, 2H), 5.14 (br s, 1H), 4.14 (br s, 1H), 3.66–3.57 (m, 1H), 3.57–3.41 (m, 1H), 3.14–2.97 (m, 2H), 2.97 (s, 1H), 2.86–2.76 (m, 5H), 2.46–2.27(m, 5H), 2.13–2.03 (m, 1H), 1.91–1.75 (m, 2H), 1.73–1.63 (m, 4H), 1.57 (d, *J* = 13.1 Hz, 1H), 1.51–1.35 (m, 2H), 1.06 (qd, *J* = 12.6, 5.2 Hz, 1H), 0.88–0.74 (m, 2H). ^13^C NMR (126 MHz, CD_3_CN): δ 174.1, 146.1, 132.1, 129.5, 127.9, 70.9, 60.8, 58.9, 49.8, 40.5, 39.8, 31.7, 31.0, 26.0, 26.0. HRMS (ESI) *m/z*: [M + H]^+^ calculated for C_23_H_34_ClN_2_O_4_S: 469.1928, found 469.1921.

#### N-((9-(1-(p-tolyl)cyclopentane-1-carbonyl)-1-oxa-9-azaspiro[5.5]undecan-4- yl)methyl)methanesulfonamide (6d)

The compound **6d** was prepared from intermediate **Xi**. White foam (60% yield). ^1^H NMR (500 MHz, CD_3_CN): δ 7.13 (d, *J* = 8.1 Hz, 2H), 7.10–7.08 (m, 2H), 5.14–5.13 (m, 1H), 4.13 (br s, 1H), 3.63–3.55 (m, 1H), 3.44–3.43 (m, 1H), 3.19 –2.92 (m, 2H), 2.83 (s, 3H), 2.82–2.73 (m, 2H), 2.36–2.29 (m, 5H), 2.19–1.99 (m, 2H), 1.90–1.75 (m, 2H), 1.69–1.63 (m, 5H), 1.56 (d, *J* = 13.2 Hz, 1H), 1.49–1.45 (m, 3H), 1.05 (tt, *J* = 12.6, 6.3 Hz, 1H), 0.87–0.75 (m, 2H). ^13^C NMR (126 MHz, CD_3_CN): δ 174.8, 144.1, 136.4, 130.1, 125.9, 71.0, 60.7, 58.9, 49.7, 40.4, 39.8, 31.6, 30.9, 26.0, 25.9, 20.8. HRMS (ESI) *m/z*: [M + H]^+^ calculated for C_24_H_37_N_2_O_4_: 449.2474, found 449.2465.

#### *N*-((9-(1-(3-chlorophenyl)cyclopentane-1-carbonyl)-1-oxa-9-azaspiro[5.5]undecan-4-yl)methyl)methanesulfonamide (*6e*)

The compound **6e** was prepared from intermediate **Xj.** White foam (62% yield). ^1^H NMR (500 MHz, CD_3_CN): δ 7.36–7.33 (m, 1H), 7.26–7.19 (m, 2H), 7.14 (m, 1H), 5.31–5.28 (m, 1H), 4.17–4.12 (m, 1H), 3.75–3.68 (m, 1H), 3.43–3.42 (m, 1H), 3.14–2.97 (m, 3H), 2.84–2.79 (m, 5H), 2.38–2.15 (m, 6H), 2.08–2.03 (m, 2H), 1.68–1.62 (m, 3H), 1.57 (d, *J* = 12.9 Hz, 1H), 1.48–1.45 (m, 1H), 1.10–1.01 (m, 1H), 0.88–0.73 (m, 3H). ^13^C NMR (126 MHz, CD_3_CN): δ 174.7, 158.8, 139.1, 127.2, 114.8, 71.0, 60.7, 58.6, 55.8, 49.8, 40.6, 39.8, 31.7, 31.0, 25.9. HRMS (ESI) *m/z*: [M + H]^+^ calculated for C_23_H_34_ClN_2_O_4_S: 469.1928, found 469.1925. HRMS (ESI) *m/z*: [M + H]^+^ calculated for C_23_H_34_ClN_2_O_4_S: 469.1928, found 469.1925.

#### Synthesis of compound 3a–l and 3p–u

Boc deprotection of intermediate **VIII** using General Procedure-I with 4M HCl/dioxane, followed by amide coupling with various carboxylic acids using General Procedure-II, afforded compounds **3a***–***l**. Alternatively, reaction with isocyanates following General Procedure-V yielded compounds **3p***–***u.**

#### N-((9-(1-Phenylcyclopentane-1-carbonyl)-1-oxa-9-azaspiro[5.5]undecan-4-yl)methyl)isobutyramide (3a)

White foam (70% yield, over two steps). ^1^H NMR (500 MHz, CD_3_CN): δ 7.34–7.29 (m, 2H), 7.24–7.18 (m, 3H), 6.27 (br s, 1H), 4.13–4.07 (m, 1H), 3.59 (dd, *J* = 11.8, 5.0 Hz, 1H), 3.53–3.28 (m, 2H), 3.17–3.10 (m, 1H), 2.95–2.79 (m, 5H), 2.41–2.40 (m, 1H), 2.33–2.27 (m,2H), 2.14–1.99 (m, 1H), 1.68–1.66 (m, 5H),1.53 –1.33 (m, 3H), 1.07–0.98 (m, 7H), 0.92–0.52 (m, 3H). ^13^C NMR (126 MHz, CD_3_CN) δ 177.7, 174.6, 147.1, 129.6, 127.0, 126.1, 71.0, 60.9, 59.3, 45.6, 40.8, 35.9, 31.5, 26.1, 26.0, 20.0 (2C). HRMS (ESI) *m/z*: [M + H]^+^ calculated for C_26_H_39_N_2_O_3_: 427.2960, found 427.2954.

#### N-((9-(1-Phenylcyclopentane-1-carbonyl)-1-oxa-9-azaspiro[5.5]undecan-4-yl)methyl)hexanamide (3b)

White foam (48% yield, over two steps). ^1^H NMR (500 MHz, CD_3_CN): δ 7.35–7.29 (m, 2H), 7.24– 7.17 (m, 3H), 6.32–6.29 (m, 1H), 4.13 (br s, 1H), 3.59 (dd, *J* = 11.7, 4.9 Hz, 1H), 3.53–3.17 (m, 2H), 3.09– 2.54 (m, 5H), 2.38–2.30 (m, 2H), 2.15–2.05 (m, 4H), 1.75–1.62 (m, 5H), 1.57–1.45 (m, 3H), 1.40–1.11 (m, 7H), 1.02 (qd, *J* = 12.6, 5.1 Hz, 1H), 0.89 (t, *J* = 7.2 Hz, 3H), 0.83–0.58 (m, 2H). ^13^C NMR (126 MHz, CD_3_CN): δ 174.5, 173.7, 147.2, 129.6, 127.0, 126.0, 71.0, 60.9, 59.3, 45.7, 40.8, 36.8, 32.1, 31.5, 26.3, 26.1, 26.0, 23.1, 14.3. HRMS (ESI) *m/z*: [M + H]^+^ calculated for C_28_H_43_N_2_O_3_: 455.3273, found 455.3268.

#### *N*-((9-(1-Phenylcyclopentane-1-carbonyl)-1-oxa-9-azaspiro[5.5]undecan-4-yl)methyl)cyclopropanecarboxamide (3c)

White foam (59% yield, over two steps). ^1^H NMR (400 MHz, CDCl_3_): δ 7.32–7.28 (m, 2H), 7.23–7.16 (m, 3H), 5.78 (br s, 1H), 4.30–3.95 (m, 5H), 3.69 (d, *J* = 11.3 Hz, 1H), 3.49–2.74 (m, 6H), 2.44–2.32 (m, 2H), 2.14–1.91 (m, 3H), 1.54–1.46 (m, 3H), 1.37–1.32 (m, 2H), 1.22–1.07 (m, 2H), 0.99–0.95 (m, 3H), 0.79–0.63 (m, 3H). ^13^C NMR (100 MHz, CDCl_3_): δ 175.2, 174.6, 128.9, 126.4, 125.2, 70.3, 60.4, 58.7, 45.9, 40.2, 30.8, 30.3, 25.3, 15.0, 7.5. HRMS (ESI) *m/z*: [M + H]^+^ calculated for C_26_H_37_N_2_O_3_: 425.2804, found 425.2797.

#### N-((9-(1-Phenylcyclopentane-1-carbonyl)-1-oxa-9-azaspiro[5.5]undecan-4-yl)methyl)cyclohexanecarboxamide (3d)

White foam (62% yield, over two steps). ^1^H NMR (500 MHz, CD_3_CN): δ 7.33–7.30 (m, 2H), 7.24–7.18 (m, 3H), 6.28–6.23 (m, 1H), 4.13 (br s, 1H), 3.58 (dd, *J* = 12.2, 4.7 Hz, 1H), 3.49–3.31 (m, 1H), 3.17–2.76 (m, 6H), 2.42–2.31 (m, 2H), 2.08–2.00 (m, 2H), 1.87–1.58 (m, 11H), 1.49–1.42 (m, 1H), 1.41–1.06 (m, 8H), 1.06–0.95 (m, 1H), 0.90–0.61 (m, 2H). ^13^C NMR (126 MHz, CD_3_CN): δ 176.8, 174.6, 147.1, 129.5, 126.9, 126.1, 71.0, 60.9, 59.3, 45.9, 45.5, 40.8, 31.5, 31.2, 30.5, 30.5, 26.5, 26.1. HRMS (ESI) *m/z*: [M + H]^+^ calculated for C_29_H_43_N_2_O_3_: 467.3273, found 467.3268.

#### 4-Chloro-N-((9-(1-phenylcyclopentane-1-carbonyl)-1-oxa-9-azaspiro[5.5]undecan-4-yl)methyl)butanamide (3e)

The compound **3e** prepared from the acylation of **4a** using the General Procedure-IV. Colorless gum (81% yield, over two steps). ^1^H NMR (500 MHz, CDCl_3_): δ 7.29 (t, *J* = 7.7 Hz, 2H), 7.23–7.16 (m, 3H), 5.56 (br s, 1H), 4.35–4.30 (m, 1H), 3.74–3.56 (m, 3H), 3.35–3.33 (m, 1H), 3.22 (m, 1H), 3.08–3.05 (m, 2H), 2.98–2.80 (m, 2H), 2.50–2.47 (m, 1H), 2.35 (t, *J* = 7.1 Hz, 3H), 2.14– 2.04 (m, 3H), 1.88–1.73 (m, 6H), 1.56–1.34 (m, 3H), 1.18–1.10 (m, 1H), 1.02–0.61 (m, 4H). ^13^C NMR (126 MHz, CDCl_3_): δ 174.6, 171.9, 146.0, 128.8, 128.8, 126.3, 125.2, 125.2, 70.4, 60.4, 58.7, 45.6, 44.6, 40.3, 33.3 (2C), 30.9, 30.5, 30.4, 28.1, 25.5, 25.4, 23.9, 21.3. HRMS (ESI) *m/z*: [M + H]^+^ calculated for C_26_H_38_ClN_2_O_3_: 461.2571, found 461.2562.

#### N-((9-(1-Phenylcyclopentane-1-carbonyl)-1-oxa-9-azaspiro[5.5]undecan-4-yl)methyl)benzamide (3f)

White foam (43% yield, over two steps). ^1^H NMR (500 MHz, CD_3_CN): δ 7.80–7.73 (m, 2H), 7.55– 7.48 (m, 1H), 7.47–7.40 (m, 2H), 7.33–7.25 (m, 2H), 7.23–7.16 (m, 3H), 7.09 (t, *J* = 6.1 Hz, 1H), 4.15 (br s, 1H), 3.60 (dd, *J* = 11.8, 5.0 Hz, 1H), 3.54–3.32 (m, 1H), 3.15–3.12 (m, 3H), 3.01–2.95 (m, 1H), 2.86– 2.67 (m, 1H), 2.37–2.22 (m, 2H), 2.16–1.96 (m, 2H), 1.86–1.80 (m, 1H), 1.73–1.60 (m, 5H), 1.56 (d, *J* = 12.7 Hz, 1H), 1.48–1.35 (m, 2H), 1.16–0.56 (m, 4H). ^13^C NMR (126 MHz, CD_3_CN): δ 174.6, 167.9, 147.1, 135.9, 132.1, 129.6, 129.3, 127.9, 126.9, 126.0, 71.0, 60.9, 59.3, 46.4, 41.0, 31.5, 31.4, 26.1, 26.0. HRMS (ESI) *m/z*: [M + H]^+^ calculated for C_29_H_37_N_2_O_3_: 461.2804, found 461.2796.

#### 4-Methoxy-N-((9-(1-phenylcyclopentane-1-carbonyl)-1-oxa-9-azaspiro[5.5]undecan-4-yl)methyl)benzamide (3g)

White foam (63% yield, over two steps). ^1^H NMR (500 MHz, CD_3_CN): δ 7.76– 7.69 (m, 2H), 7.34–7.27 (m, 2H), 7.23–7.15 (m, 3H), 7.00–6.87 (m, 3H), 4.26–4.04 (m, 1H), 3.83 (s, 3H), 3.60 (dd, *J* = 12.4, 4.7 Hz, 1H), 3.44–3.39 (m, 1H), 3.28–3.20 (m, 1H), 3.11 (t, *J* = 6.4 Hz, 2H), 3.06–2.91 (m, 1H), 2.77 _ 2.75 (m, 1H), 2.49–2.24 (m, 2H), 2.20–2.01 (m, 2H), 1.90–1.80 (m, 1H), 1.69–1.66 (m, 4H), 1.58–1.50 (m, 1H), 1.47–1.36 (m, 2H), 1.31–1.04 (m, 2H), 1.00–0.60 (m, 3H). ^13^C NMR (126 MHz, CD_3_CN): δ 174.6, 167.5, 162.9, 147.1, 129.7, 129.5, 128.0, 126.9, 126.0, 114.4, 71.0, 60.9, 59.3, 56.1, 46.3, 40.9, 31.5, 31.3, 26.0 (2C) HRMS (ESI) *m/z*: [M + H]^+^ calculated for C_30_H_39_N_2_O_4_: 491.2910, found 491.2903.

#### 3-Methoxy-N-((9-(1-phenylcyclopentane-1-carbonyl)-1-oxa-9-azaspiro[5.5]undecan-4-yl)methyl)benzamide (3h)

White foam (62% yield, over two steps). ^1^H NMR (500 MHz, CD_3_CN): δ 7.41–7.26 (m, 5H), 7.23–7.16 (m, 3H), 7.07–7.05 (m, 1H), 7.04–6.96 (m, 1H), 4.15 (br s, 1H), 3.83 (s, 3H), 3.62– 3.47 (m, 1H), 3.22–3.18 (m, 3H), 2.98–2.73 (m, 2H), 2.40–2.32 (m, 2H), 2.18–2.00 (m, 3H), 1.86–1.80 (m, 1H), 1.67–1.55 (m, 5H), 1.48–1.37 (m, 2H), 1.20–1.04 (m, 2H), 1.01–0.59 (m, 3H). ^13^C NMR (126 MHz, CD_3_CN): δ 174.5, 167.6, 160.7, 147.2, 137.4, 130.5, 129.6, 126.9, 126.0, 120.1, 117.9, 113.2, 71.0, 60.9, 59.3, 56.0, 46.4, 31.5, 31.4, 26.1. HRMS (ESI) *m/z*: [M + H]^+^ calculated for C_30_H_39_N_2_O_4_: 491.2910, found 491.2903.

#### 4-(Hexyloxy)-N-((9-(1-phenylcyclopentane-1-carbonyl)-1-oxa-9-azaspiro[5.5]undecan-4-yl)methyl)benzamide (3i)

White foam (59% yield, over two steps). ^1^H NMR (500 MHz, CDCl_3_): δ 7.69 (d, *J* = 8.7 Hz, 2H), 7.31–7.26 (m, 2H), 7.22–7.13 (m, 3H), 6.91–6.89 (dt, *J* = 8.8, 1.9 Hz, 2H), 6.17 (br s, 1H), 4.29 (br s, 1H), 4.00–3.97 (m, 2H), 3.74–3.66 (m, 1H), 3.56–3.35 (m, 1H), 3.32–3.17 (m, 3H), 3.07–2.77 (m, 2H), 2.46–2.33 (m, 2H), 2.12–2.05 (m, 4H), 1.83–1.76 (m, 2H), 1.75–1.66 (m, 4H), 1.59–1.39 (m, 5H), 1.37–1.30 (m, 4H), 1.25–1.16 (m, 2H), 1.08–0.96 (m, 1H), 0.92–0.86 (m, 3H), 0.83–0.61 (m, 1H). ^13^C NMR (126 MHz, CDCl_3_): δ 174.6, 167.4, 162.0, 146.0, 128.8, 128.7, 126.5, 126.3, 125.1, 114.4, 70.4, 68.4, 60.4, 58.7, 46.0, 41.8, 40.4, 39.4, 38.5, 37.8, 31.7, 31.0, 30.5, 29.2, 25.8, 25.4, 22.7, 14.2. HRMS (ESI) *m/z*: [M + H]^+^ calculated for C_35_H_49_N_2_O_4_: 561.3692, found 561.3685.

#### N-((9-(1-Phenylcyclopentane-1-carbonyl)-1-oxa-9-azaspiro[5.5]undecan-4-yl)methyl)picolinamide (3j)

White foam (59% yield, over two steps). ^1^H NMR (400 MHz, CD_3_CN): δ 8.58–8.56 (m, 1H), 8.17–8.05 (m, 2H), 7.91 (td, *J* = 7.7, 1.7 Hz, 1H), 7.52–7.49 (m, 1H), 7.34–7.26 (m, 2H), 7.23–7.14 (m, 3H), 4.15 (br s, 1H), 3.60 (dd, *J* = 11.9, 5.0 Hz, 1H), 3.45–3.39 (m, 1H), 3.24–3.14 (m, 3H), 2.95–2.89 (m, 1H), 2.83–2.51 (m, 3H), 2.36–2.22 (m, 2H), 1.91–1.77 (m, 2H), 1.66–1.52 (m, 5H), 1.47–1.36 (m, 2H), 1.24– 1.04 (m, 2H), 0.97–0.58 (m, 2H). ^13^C NMR (126 MHz, CD_3_CN): δ 174.6, 165.1, 151.1, 149.3, 147.1, 138.6, 129.6, 127.3, 126.9, 126.0, 122.7, 71.0, 60.8, 59.3, 45.8, 40.9, 31.7, 31.3, 26.1, 26.0. HRMS (ESI) *m/z*: [M + H]^+^ calculated for C_28_H_36_N_3_O_3_: 462.2756, found 462.2746.

#### N-((9-(1-Phenylcyclopentane-1-carbonyl)-1-oxa-9-azaspiro[5.5]undecan-4-yl)methyl)nicotinamide (3k)

White foam (40% yield, over two steps). ^1^H NMR (500 MHz, CD_3_CN): δ 8.95 (s, 1H), 8.71–8.70 (m, 1H), 8.18–8.14 (m, 1H), 7.55–7.44 (m, 1H), 7.36–7.26 (m, 2H), 7.22–7.17 (m, 3H), 4.84 (br s, 1H), 4.15 (br s, 1H), 3.55–3.40 (m, 1H), 3.25–3.06 (m, 3H), 2.96–2.79 (m, 3H), 2.57–1.97 (m, 4H), 1.92–1.73 (m, 2H), 1.72–1.62 (m, 4H), 1.58 (d, *J* = 12.8 Hz, 1H), 1.48–1.40 (m, 1H), 1.40–1.20 (m, 1H), 1.15–1.06 (m, 1H), 0.93–0.67 (m, 3H). ^13^C NMR (126 MHz, CD_3_CN): δ 174.6, 165.9, 151.6, 148.2, 147.1, 137.0, 131.9, 129.6, 126.9, 126.0, 124.8, 71.0, 60.8, 59.3, 46.4, 40.8, 31.4, 31.3, 26.1, 26.0. HRMS (ESI) *m/z*: [M + H]^+^ calculated for C_28_H_36_N_3_O_3_: 462.2756, found 462.2747.

#### N-((9-(1-Phenylcyclopentane-1-carbonyl)-1-oxa-9-azaspiro[5.5]undecan-4-yl)methyl)isonicotinamide (3l)

White foam (48% yield, over two steps). ^1^H NMR (500 MHz, CD_3_CN): δ 8.75–8.70 (m, 2H), 7.75–7.73 (m, 2H), 7.32–7.29 (m, 3H), 7.23–7.15 (m, 3H), 4.88–4.35 (m, 2H), 4.15 (br s, 1H), 3.66–3.58 (m, 1H), 3.53–3.51 (m, 2H), 3.15 (t, *J* = 6.4 Hz, 3H), 3.05–2.70 (m, 2H), 2.37–2.08 (m, 3H), 2.10–2.03 (m, 1H), 1.87–1.82 (m, 1H), 1.69–1.55(m, 4H), 1.57 (d, *J* = 12.9 Hz, 1H), 1.47–1.43 (m, 1H), 1.18–1.05 (m, 2H), 0.98–0.93 (m, 3H). ^13^C NMR (126 MHz, CD_3_CN): δ 174.6, 165.8, 149.7, 147.1, 144.4, 126.9, 126.0, 122.8, 71.0, 60.8, 59.3, 46.5, 40.8, 31.3, 26.1, 26.0. HRMS (ESI) *m/z*: [M + H]^+^ calculated for C_28_H_36_N_3_O_3_: 462.2756, found 462.2746.

#### (4-(Aminomethyl)-1-oxa-9-azaspiro[5.5]undecan-9-yl)(1-phenylcyclopentyl)methanone (3p)

White foam (79% yield, over two steps). ^1^H NMR (400 MHz, CD_3_CN): δ 7.39–7.30 (m, 2H), 7.28–7.20 (m, 3H), 5.86–5.12 (m, 4H), 4.16 (br s, 1H), 3.62 (dd, *J* = 11.6, 5.0 Hz, 1H), 3.46–3.21 (m, 2H), 3.00–2.82 (m, 4H), 2.39–2.07 (m, 3H), 1.94–1.62 (m, 6H), 1.53–1.51 (m, 1H), 1.43–1.38 (m, 2H), 1.19–1.00 (m, 2H), 0.87– 0.75 (m, 2H). ^13^C NMR (126 MHz, CD_3_CN): δ 174.7, 147.1, 129.6, 126.9, 126.0, 70.9, 60.8, 59.3, 46.9, 40.5, 31.6, 31.0, 26.0, 26.0. HRMS (ESI) *m/z*: [M + H]^+^ calculated for C_23_H_34_N_3_O_3_: 400.2600, found 400.2592.

#### 1-Isopropyl-3-((9-(1-phenylcyclopentane-1-carbonyl)-1-oxa-9-azaspiro[5.5]undecan-4-yl)methyl)urea (3q)

White foam (89% yield, over two steps). ^1^H NMR (500 MHz, CD_3_CN): δ 7.35–7.27 (m, 2H), 7.25–7.17 (m, 3H), 4.70–4.13 (m, 4H), 3.74–3.69 (m, 1H), 3.59 (dd, *J* = 12.0, 4.9 Hz, 1H), 3.47 – 3.18 (m, 2H), 3,00–2.76 (m, 4H), 2.37–2.22 (m, 2H), 2.13–1.99 (m, 1H), 1.91–1.73 (m, 1H), 1.74–1.60 (m, 5H), 1.56–1.39 (m, 2H), 1.40–1.34 (m, 1H), 1.27–1.05 (m, 7H), 1.03–0.93 (m, 1H), 0.83–0.83 (m, 2H). ^13^C NMR (126 MHz, CD_3_CN): δ 174.7, 158.9, 147.1, 129.5, 126.9, 126.0, 70.9, 60.9, 59.3, 46.7, 42.7, 40.7, 31.9, 31.1, 26.0, 26.0, 23.4. HRMS (ESI) *m/z*: [M + H]^+^ calculated for C_26_H_40_N_3_O_3_: 442.3069, found 442.3061.

#### 1-(Tert-butyl)-3-((9-(1-phenylcyclopentane-1-carbonyl)-1-oxa-9-azaspiro[5.5]undecan-4-yl)methyl)urea (3r)

White foam (61% yield, over two steps). ^1^H NMR (500 MHz, CD_3_CN): δ 7.34–7.30 (m, 2H), 7.23–7.19 (m, 3H), 4.13 (br s, 1H), 3.63–3.54 (m, 1H), 3.44–3.40 (m, 2H), 3.17–2.95 (m, 2H), 2.83–2.78 (m, 3H), 2.41–2.34 (m, 3H), 2.10–1.97 (m, 2H), 1.91–1.79 (m, 1H), 1.69–1.67 (m, 5H), 1.47 (d, *J* = 12.8 Hz, 1H), 1.39–1.33 (m, 1H), 1.25–1.24 (m, 9H), 1.14–1.10 (m, 1H), 1.00 (tt, *J* = 12.6, 6.5 Hz, 1H), 0.81–0.63 (m, 3H). ^13^C NMR (126 MHz, CD_3_CN): δ 174.6, 158.8, 147.1, 129.6, 127.0, 126.0, 70.9, 60.9, 59.3, 50.6, 46.4, 40.7, 31.2, 29.6, 26.1. HRMS (ESI) *m/z*: [M + H]^+^ calculated for C_27_H_42_N_3_O_3_: 456.3226, found 456.3217.

#### 1-Phenyl-3-((9-(1-phenylcyclopentane-1-carbonyl)-1-oxa-9-azaspiro[5.5]undecan-4-yl)methyl)urea (3s)

White foam (34% yield, over two steps). ^1^H NMR (400 MHz, CD_3_CN): δ 7.40–7.34 (m, 2H), 7.34–7.28 (m, 2H), 7.27–7.16 (m, 5H), 7.13 (br s, 1H), 6.97–6.93 (m, 1H), 5.25 (t, *J* = 6.0 Hz, 1H), 4.14 (br s, 1H), 3.60 (dd, *J* = 12.1, 4.9 Hz, 1H), 3.45–3.39 (m, 1H), 3.25–3.11 (m, 1H), 2.97–2.93 (m, 3H), 2.79–2.78 (m, 1H), 2.36 –2.28 (m, 2H), 2.15–2.04 (m, 3H), 2.09–2.01 (m, 1H), 1.88–1.77 (m, 1H), 1.71 –1.63 (m, 4H), 1.52 (d, *J* = 12.9 Hz, 1H), 1.42–1.37 (m, 1H), 1.10–0.68 (m, 4H). ^13^C NMR (126 MHz, CD_3_CN): δ 174.6, 156.4, 147.1, 141.2, 129.6, 129.5, 126.9, 126.0, 122.7, 119.4, 70.9, 60.9, 59.3, 46.4, 40.7, 31.9, 31.1, 26.0 (2C). HRMS (ESI) *m/z*: [M + H]^+^ calculated for C_29_H_38_N_3_O_3_:476.2913, found 476.2905.

#### 1-(4-Methoxyphenyl)-3-((9-(1-phenylcyclopentane-1-carbonyl)-1-oxa-9-azaspiro[5.5]undecan-4-yl)methyl)urea (3t)

White foam (64% yield, over two steps). ^1^H NMR (500 MHz, CD_3_CN): δ 7.34–7.27 (m, 2H), 7.26–7.12 (m, 6H), 6.83–6.78 (m, 2H), 5.47–5.02 (m, 1H), 4.13 (br s, 1H), 3.73 (s, 3H), 3.63–3.54 (m, 1H), 3.45–3.36 (m, 1H), 3.24–3.08 (m, 1H), 3.07–2.96 (m, 1H), 2.92 (d, *J* = 6.5 Hz, 2H), 2.79–2.36 (m, 5H), 2.13–1.99 (m, 1H), 1.76–1.61 (m, 5H), 1.51–1.48 (m, 1H), 1.41–1.35 (m, 2H), 1.03 (qd, *J* = 12.6, 5.1 Hz, 1H), 0.84–0.65 (m, 3H). ^13^C NMR (126 MHz, CD_3_CN): δ 174.6, 156.8, 156.1, 147.1, 134.1, 129.6, 126.9, 126.0, 121.9, 114.8, 70.9, 60.9, 59.3, 56.0, 46.5, 40.7, 32.0, 31.2, 26.1, 26.0. HRMS (ESI) *m/z*: [M + H]^+^ calculated for C_30_H_40_N_3_O_4_: 506.3019, found 506.3010.

#### 1-(2-Methoxyphenyl)-3-((9-(1-phenylcyclopentane-1-carbonyl)-1-oxa-9-azaspiro[5.5]undecan-4-yl)methyl)urea (3u)

White foam (67% yield, over two steps). ^1^H NMR (500 MHz, CD_3_CN): δ 8.10–8.03 (m, 1H), 7.33–7.28 (m, 3H), 7.24–7.16 (m, 3H), 6.93–6.88 (m, 2H), 6.87–6.84 (m, 1H), 5.82–5.61 (m, 1H), 4.21–4.12 (m, 1H), 3.82 (s, 3H), 3.62–3.55 (m, 1H), 3.46–3.41 (m, 1H), 3.16–2.73 (m, 5H), 2.36–1.97 (m, 4H), 1.90–1.77 (m, 1H), 1.69–1.66 (m, 5H), 1.51 (d, *J* = 13.1 Hz, 1H), 1.41–1.38 (m, 2H), 1.05 (tt, *J* = 12.6, 6.3 Hz, 1H), 1.00–0.50 (m, 3H). ^13^C NMR (126 MHz, CD_3_CN): δ 174.6, 156.4, 148.7, 147.1, 130.5, 129.6, 126.9, 126.0, 122.4, 121.6, 119.3, 111.3, 70.9, 60.9, 59.3, 56.4, 46.4, 40.7, 31.9, 31.2, 26.1, 26.0. HRMS (ESI) *m/z*: [M + H]^+^ calculated for C_30_H_40_N_3_O_4_: 506.3019, found 506.3011.

#### Synthesis of 5f, 5g, 6f–m and 7a–c

Boc deprotection of **XII** and **XIV** using TFA/DCM followed by amide coupling with carboxylic acids using the General Procedures I and II afforded (**5f**, **5g**, **6f***–***m**) and **7a***–***c** respectively.

#### {4-[(Dimethylaminosulfonylamino)methyl]-1-oxa-9-aza-9-spiro[5.5]undecyl}(2-phenyltetrahydro-2-furyl)methanone (5f)

White foam (77% yield, over two steps). ^1^H NMR (400 MHz, DMSO-*d*_6_, 80 °C): δ 7.38–7.24 (m, 5H), 6.82 (br s, 1H), 4.13 (q, *J* = 7.2 Hz, 1H), 4.00–3.68 (m, 3H), 3.66–3.52 (m, 1H), 3.44–3.43 (m, 1H), 3.07–3.03 (m, 2H), 2.75–2.70 (m, 2H), 2.66–2.61 (m, 6H), 1.87–1.83 (m, 2H), 1.80–1.60 (m, 3H), 1.55 (d, *J* = 13.2 Hz, 1H), 1.35–1.09 (m, 3H), 1.06–0.96 (m, 1H), 0.83–0.79 (m, 2H). ^13^C NMR (101 MHz, DMSO-*d*_6_): δ 169.1, 143.6, 128.4, 127.1, 123.5, 89.1, 88.9, 69.8, 69.6, 59.5, 48.8, 37.6, 30.3, 30.1, 30.0, 29.0, 28.5, 25.0, 24.3. HRMS (ESI) *m/z*: [M + H]^+^ calculated for C_23_H_35_N_2_O_5_S: 451.2266, found 451.2256.

#### 1-{4-[(Dimethylaminosulfonylamino)methyl]-1-oxa-9-aza-9-spiro[5.5]undecyl}-2-phenyl-1,2-ethanedione (5g)

Pale yellow thick oil (84% yield, over two steps). ^1^H NMR (400 MHz, CDCl_3_, 45 °C): δ 8.02–7.87 (m, 2H), 7.71–7.59 (m, 1H), 7.57–7.43 (m, 2H), 4.40 (br s, 1H), 4.06–3.81 (m, 2H), 3.65–3.46 (m, 2H), 3.32–3.26 (m, 2H), 3.11–2.92 (m, 2H), 2.82–2.80 (m, 6H), 2.37–2.16 (m, 1H), 1.97–1.89 (m, 1H),1.72 –1.58 (m, 3H), 1.42–1.33 (m, 1H), 1.30–1.19 (m, 1H), 1.11–0.87 (m, 2H). ^13^C NMR (126 MHz, DMSO-*d_6_*): δ 169.1 (3C), 143.7, 143.6, 143.4, 143.3, 128.5, 128.4, 127.1, 123.5 (2C), 89.1, 88.9, 69.8, 69.6, 69.3, 69.2, 48.9, 48.8, 41.2, 41.1, 41.0, 38.6, 38.4, 38.2, 38.1, 37.6 (2C), 30.3, 30.2, 30.1, 30.0, 29.9, 29.0, 28.5, 25.0 (3C), 24.9, 24.3. (additional carbon signals observed due to rotamers). HRMS (ESI) *m/z*: [M + H]^+^ calculated for C_20_H_30_N_3_O_5_S: 424.1906, found 424.1897.

#### {4-[(Dimethylaminosulfonylamino)methyl]-1-oxa-9-aza-9-spiro[5.5]undecyl}[1-(m-methoxyphenyl)cyclopentyl]methanone (6f)

White foam (41% yield, over two steps). ^1^H NMR (500 MHz, CD_3_CN): δ 7.24 (t, *J* = 7.9 Hz, 1H), 6.80–6.75 (m, 2H), 6.75–6.72 (m, 1H), 5.16 (br s, 1H), 4.13 (br s, 1H), 3.76 (s, 3H), 3.61 (dd, *J* = 12.0, 5.0 Hz, 1H), 3.51–3.21 (m, 2H), 2.98 –2.69 (m, 9H), 2.41–2.31 (m, 2H), 2.14–2.03 (m, 3H), 1.84–1.80 (m, 1H), 1.75–1.53 (m, 6H), 1.49–1.37 (m, 2H), 1.30–1.12 (m, 1H), 1.04 (qd, *J* = 12.6, 5.2 Hz, 1H), 0.93–0.69 (m, 2H). ^13^C NMR (126 MHz, CD_3_CN): δ 174.5, 160.9, 148.8, 130.6, 112.1, 111.8, 71.0, 60.8, 59.3, 55.8, 50.2, 40.7, 38.4, 31.6, 31.1, 26.1. HRMS (ESI) *m/z*: [M + H]^+^ calculated for C_25_H_40_N_3_O_5_S: 494.2688, found 494.2682.

#### {4-[(Dimethylaminosulfonylamino)methyl]-1-oxa-9-aza-9-spiro[5.5]undecyl}[1-(m-tolyl)cyclopentyl]methanone (6g)

White foam (13% yield, over two steps). ^1^H NMR (400 MHz, DMSO-*d_6_*, 80 °C) δ 7.20 (t, *J* = 6.4 Hz, 1H), 6.99–7.02 (m, 3H), 6.81 (s, 1H), 3.58–3.60 (m, 2H), 3.42 - 3.45 (m, 2H), 2.99 - 3.02 (m, 2H), 2.79 - 2.85 (m, 2H), 2.70–2.71 (m, 2H), 2.65 (s, 6H), 2.29 (m, 5H), 1.91 (s, 2H), 1.73 (s, 2H), 1.64 (s, 4H), 1.48 - 1.51 (m, 2H), 1.17 - 1.18 (m, 1H), 0.98 - 1.02 (m, 1H), 0.81 - 0.84 (m, 1H). ^13^C NMR (101 MHz, DMSO-d_6_) 173.5, 146.2, 138.1, 129,0, 127.1, 125.9, 122.3, 70.2, 60.0, 58.3, 49.3, 38.7, 38.1, 30.7, 30.5, 25.5, 21.7. C_25_H_40_N_3_O_4_S: 478.2739, found 478.2731.

#### {4-[(Dimethylaminosulfonylamino)methyl]-1-oxa-9-aza-9-spiro[5.5]undecyl}{1-[m-(trifluoromethyl)phenyl]cyclopentyl}methanone (6h)

White foam (24% yield, over two steps). ^1^H NMR (400 MHz, DMSO-*d_6_*, 80 °C) δ 7.60 –7.54 (m, 3H), 7.40 (s, 1H), 6.80 (s, 1H), 3.41–3.61 (m, 4H), 3.00 – 3.06 (m, 2H), 2.66–2.83 (m, 10H), 2.42–2.49 (m, 2H), 1.95–1.99 (m, 2H), 1.67–1.92 (m, 6H), 1.47–1.56 (m, 2H), 1.26 (m, 1H), 1.01–1.05 (m, 1H), 0.81–0.87 (m, 1H). ^13^C NMR (101 MHz, DMSO-*d*_6_): δ 172.8, 147.5, 130.5, 129.5, 123.4, 121.8, 70.1, 60.0, 58.3, 49.2, 40.6, 39.4, 38.1, 30.7, 30.4, 25.5. HRMS (ESI) *m/z*: [M + H]^+^ calculated for C_25_H_37_F_3_N_3_O4S: 532.2457, found 532.2451.

#### [1-(o-Chlorophenyl)cyclopentyl]{4-[(dimethylaminosulfonylamino)methyl]-1-oxa-9-aza-9-spiro[5.5]undecyl}methanone (6i)

White foam (18% yield, over two steps). ^1^H NMR (500 MHz, CD_3_CN): δ 7.48 (dd, *J* = 7.9, 1.6 Hz, 1H), 7.39–7.37 (m, 1H), 7.34–7.31 (m, 1H), 7.24–7.21 (m, 1H), 5.16 (br s, 1H), 4.09 (br s, 1H), 3.63–3.33 (m, 2H), 3.16 (d, *J* = 13.5 Hz, 1H), 3.02–2.91 (m, 1H), 2.81–2.75 (m, 2H), 2.69 (s, 6H), 2.36–2.22 (m, 3H), 2.13–2.00 (m, 1H), 1.89–1.51 (m, 7H), 1.51–1.39 (m, 2H), 1.27–1.24 (m, 1H), 1.03 (qd, *J* = 12.6, 5.2 Hz, 1H), 0.94–0.51 (m, 3H). ^13^C NMR (126 MHz, CD_3_CN): δ 173.5, 144.2, 133.9, 132.0, 128.7, 128.3, 128.1, 71.1, 60.8, 58.7, 50.2, 38.4, 31.6, 31.1, 25.6. HRMS (ESI) *m/z*: [M + H]^+^ calculated for C_24_H_37_ClN_3_O_4_S: 498.2193, found 498.2185.

#### {4-[(Dimethylaminosulfonylamino)methyl]-1-oxa-9-aza-9-spiro[5.5]undecyl}[1-(o-fluorophenyl)cyclopentyl]methanone (6j)

White foam (45% yield, over two steps). ^1^H NMR (500 MHz, CD_3_CN): δ 7.36 (td, *J* = 8.1, 1.8 Hz, 1H), 7.29–7.23 (m, 1H), 7.17 (td, *J* = 7.6, 1.4 Hz, 1H), 7.06 (ddd, *J* = 11.9, 8.1, 1.3 Hz, 1H), 5.18 (br s, 1H), 4.11 (br s, 1H), 3.60 (dd, *J* = 12.1, 4.9 Hz, 1H), 3.57–3.35 (m, 1H), 3.22–2.98 (m, 2H), 2.83–2.69 (m, 3H), 2.69 (s, 6H), 2.47–2.36 (m, 3H), 2.14–2.02 (m, 1H), 1.90–1.81 (m, 1H), 1.77–1.61 (m, 5H), 1.61–1.51 (m, 1H), 1.49–1.48 (m, 2H), 1.24–0.98 (m, 2H), 0.85–0.68 (m, 2H). ^13^C NMR (126 MHz, CD_3_CN): δ 173.6, 161.2 (d, *J* = 247 Hz), 134.0 (d, *J* = 12.6 Hz), 129.1 (d, *J* = 8.2 Hz), 128.0 (d, *J* = 5 Hz), 125.4 (d, *J* = 2.5 Hz), 117.1 (d, *J* = 23 Hz), 71.0, 60.8, 55.9, 50.1, 40.6, 38.4, 31.6, 31.1, 25.7. HRMS (ESI) *m/z*: [M + H]^+^ calculated for C_24_H_37_FN_3_O_4_S: 482.2489, found 482.2480.

#### {4-[(Dimethylaminosulfonylamino)methyl]-1-oxa-9-aza-9-spiro[5.5]undecyl}[1-(2-pyridyl)cyclopentyl]methanone (6k)

White foam (86% yield, over two steps). ^1^H NMR (400 MHz, DMSO-*d*_6_, 80 °C): δ 8.51 (d, *J* = 4.8 Hz, 1H), 7.75 (t, *J* = 15.6 Hz, 2H), 7.22 (t, *J* = 11.6 Hz, 1H), 7.16 (d, *J* = 7.6 Hz, 1H), 6.81 (s, 1H), 3.59 (t, *J* = 12 Hz, 2H), 3.44 (t, *J* = 22.4 Hz, 2H), 2.96 (s, 1H), 2.82 - 2.65 (m, 8H), 2.33 - 2.20 (m, 4H), 1.74 (m, 2H), 1.62 (s, 3H), 1.56 (t, *J* = 30.8 Hz, 2H), 1.02 - 0.98 (m, 5H), 0.83 (t, *J* = 25.6 Hz, 2H). ^13^C NMR (101 MHz, DMSO-*d*_6_) δ 173.0, 164.3, 149.0, 137.6, 121.9, 120.9, 70.2, 60.7, 60.0, 49.3, 30.7, 30.5, 26.0. HRMS (ESI) *m/z*: [M + H]^+^ calculated for C_23_H_37_N_4_O_4_S: 465.2535, found 465.2525.

#### {4-[(dimethylaminosulfonylamino)methyl]-1-oxa-9-aza-9-spiro[5.5]undecyl}[1-(3-pyridyl)cyclopentyl]methanone (6l)

White foam (2% yield, over two steps). ^1^H NMR (400 MHz, DMSO-*d*_6_, 80 °C): δ 8.42 (s, 2H), 7.60 (d, *J* = 8.0 Hz, 1H), 7.36 - 7.33 (m, 1H), 6.82 (br s, 1H), 3.64–3.52 (m, 2H), 3.45–3.39 (m, 1H), 2.87–2.84 (m, 1H), 2.72–2.69 (m, 2H), 2.65 (s, 6H), 2.36 (br s, 3H), 1.96 (br s, 2H), 1.81 (br s, 3H), 1.67 (br s, 4H), 1.56–1.50 (m, 2H), 1.26–1.20 (m, 1H), 1.15–0.98 (m, 3H), 0.85–0.78 (m, 1H). ^13^C NMR (400 MHz, DMSO-*d_6_*, 80 °C) 173.0, 147.8, 146.9, 141.5, 133.1, 124.2, 70.2, 60.0, 56.7, 49.2, 38.1, 30.7, 30.4, 25.4. HRMS (ESI) *m/z*: [M + H]^+^ calculated for C_23_H_37_N_4_O_4_S: 465.2535, found 465.2525.

#### {4-[(Dimethylaminosulfonylamino)methyl]-1-oxa-9-aza-9-spiro[5.5]undecyl}[1-(4-pyridyl)cyclopentyl]methanone (6m)

White foam (16% yield, over two steps). ^1^H NMR (400 MHz, DMSO-*d*_6_, 80 °C): δ 8.51 (d, *J* = 6.0 Hz, 2H), 7.18 (d, *J* = 6.0 Hz, 2H), 6.81 (s, 1H), 3.61 - 3.57 (m, 2H), 3.45 - 3.39 (m, 2H), 2.99 (t, *J* = 12.0 Hz, 1H), 2.82 (t, *J* = 11.2 Hz, 1H), 2.76 - 2.70 (m, 2H), 2.65 (s, 6H), 2.38 - 2.34 (m, 2H), 1.95 - 1.90 (m, 2H), 1.82 (br s, 1H), 1.74 (br s, 1H), 1.66 (br s, 4H), 1.53 (t, *J* = 12.8 Hz, 2H), 1.20 (br s, 1H), 1.05 (br s, 1H), 1.03 - 0.96 (m, 2H), 0.82 (t, *J* = 12.8 Hz, 1H). ^13^C NMR (101 MHz, DMSO-*d*_6_): δ 172.1, 155.0, 150.5, 120.7, 70.2, 60.0, 58.0, 49.3, 38.1, 30.7, 30.5, 28.9, 25.7. HRMS (ESI) *m/z*: [M + H]^+^ calculated for C_23_H_37_N_4_O_4_S: 465.2535, found 465.2526.

#### (4-((1,1-Dioxido-1,2-thiazinan-2-yl)methyl)-1-oxa-9-azaspiro[5.5]undecan-9-yl)(1-(2-fluorophenyl)cyclopentyl)methanone (7a)

White foam (39% yield, over two steps). ^1^H NMR (500 MHz, CD_3_CN): δ 7.38–7.34 (m, 1H), 7.28–7.24 (m, 1H), 7.17 –7.15 (m, 1H), 7.08–7.04 (m, 1H), 4.11 (br s, 1H), 3.61 (dd, *J* = 12.0, 5.0 Hz, 1H), 3.50–3.40 (m, 1H), 3.34–3.14 (m, 3H), 3.07–2.91 (m, 3H), 2.88–2.81 (m, 3H), 2.53–2.23 (m, 2H), 2.13 (d, *J* = 3.3 Hz, 1H), 2.10–2.02 (m, 3H), 1.89–1.79 (m, 2H), 1.75–1.64 (m, 4H), 1.62–1.50 (m, 3H), 1.46–1.42 (m, 2H), 1.02 (qd, *J* = 12.6, 5.1 Hz, 2H), 0.84–0.66 (m, 2H). ^13^C NMR (126 MHz, CD_3_CN): δ 173.6, 161.2 (d, *J* = 247 Hz), 134.1 (d, *J* = 14 Hz), 129.1 (d, *J* = 8 Hz), 128.0 (d, *J* = 5 Hz), 125.4 (d, *J* = 5 Hz), 117.0 (d, *J* = 23 Hz), 71.0, 60.9, 55.6, 54.10, 51.2, 47.98, 40.72, 38.7, 31.2, 25.8, 24.6, 20.9. HRMS (ESI) *m/z*: [M + H]^+^ calculated for C_26_H_38_FN_2_O_4_S: 493.2536, found 493.2529.

#### (1-(3,5-Difluorophenyl)cyclopentyl)(4-((1,1-dioxido-1,2-thiazinan-2-yl)methyl)-1-oxa-9-azaspiro[5.5]undecan-9-yl)methanone (7b)

White foam (42% yield, over two steps). ^1^H NMR (500 MHz, DMSO-*d*_6_): δ 7.12–7.08 (m, 1H), 6.87–6.83 (m, 2H), 4.18–3.91 (m, 1H), 3.61–3.58 (m, 1H), 3.45–3.39 (m, 1H), 3.26–3.25 (m, 2H), 3.09–2.98 (m, 4H), 2.86–2.58 (m, 3H), 2.36–2.31 (m, 2H), 2.10–1.97 (m, 3H), 1.83–1.78 (m, 3H), 1.71–1.48 (m, 7H), 1.47–1.27 (m, 2H), 1.19–1.03 (m, 1H), 1.03–0.65 (m, 3H). ^13^C NMR (126 MHz, DMSO-*d*_6_): δ 171.90, 162.6 (dd, *J* = 246.4, 13.6 Hz), 150.4, 108.3 (d, *J* = 20 Hz), 108.2 (d, *J* = 20 Hz), 101.7 (t, *J* = 25 Hz), 69.7, 59.6, 58.0, 52.8, 50.0, 46.6, 30.1, 29.3, 28.4, 24.9 (2C), 23.5, 19.5. HRMS (ESI) *m/z*: [M + H]^+^ calculated for C_26_H_37_F_2_N_2_O_4_S: 511.2442, found 511.2436.

#### (4-((1,1-Dioxido-1,2-thiazinan-2-yl)methyl)-1-oxa-9-azaspiro[5.5]undecan-9-yl)(1-(4-fluorophenyl)cyclopentyl)methanone (7c)

White foam (55% yield, over two steps). ^1^H NMR (500 MHz, CD_3_CN): δ 7.27–7.16 (m, 2H), 7.10–6.98 (m, 2H), 4.15 (br s, 1H), 3.63–3.59 (m, 1H), 3.52–3.34 (m, 1H), 3.31–3.16 (m, 3H), 3.01–2.90 (m, 3H), 2.88 –2.80 (m, 3H), 2.37–2.21 (m, 3H), 2.12–1.98 (m, 3H), 1.84– 1.80 (m, 2H), 1.67 (dd, *J* = 8.1, 4.2 Hz, 4H), 1.57 –1.52 (m, 3H), 1.46–1.41 (m, 2H), 1.27–1.12 (m, 1 H), 1.02 (qd, *J* = 12.6, 5.1 Hz, 1H), 0.84–0.70 (m, 2H). ^13^C NMR (126 MHz, CD_3_CN): δ 174.30, 162.0 (d, *J* = 243 Hz) 143.2 (d, *J* = 2.5 Hz), 127.9 (d, *J* = 8.8 Hz), 116.1 (d, *J* = 21 Hz), 71.0, 60.9, 58.8, 54.1, 51.2, 48.0, 40.7, 38.8, 30.7, 26.0, 25.9 (2C), 24.6, 20.8. HRMS (ESI) *m/z*: [M + H]^+^ calculated for C_26_H_38_FN_2_O_4_S: 493.2536, found 493.2539.

#### Synthesis of 2i–m, 4b, 4d and 3m

*N*-alkylation of **2a** and **2g** using the General Procedure VIII afforded **2i**, **2k** and **2m**.

#### N-Methyl-N-((9-(1-phenylcyclopentane-1-carbonyl)-1-oxa-9-azaspiro[5.5]undecan-4-yl)methyl)methanesulfonamide (2i)

The compound **2i** was prepared from *N*-methylation of **2a**. white foam (52% yield). ^1^H NMR (500 MHz, CD_3_CN): δ 7.35–7.29 (m, 2H), 7.24–7.18 (m, 3H), 4.16 (br s, 1H), 3.61 (dd, *J* = 11.8, 5.0 Hz, 1H), 3.51 −3.44 (m, 1H), 3.19 (br s, 1H), 3.08–2.89 (m, 1H), 2.87–2.75 (m, 3H), 2.74 (s, 3H), 2.72 (s, 3H), 2.39–2.22 (m, 2H), 2.18–2.02 (m, 1H), 1.88–1.82 (m, 2H), 1.73–1.63 (m, 4H), 1.54 (d, *J* = 13.1 Hz, 1H), 1.44–1.39 (m, 2H), 1.34–1.14 (m, 1H), 1.03 (qd, *J* = 12.6, 5.1 Hz, 1H), 0.84–0.70 (m, 3H). ^13^C NMR (126 MHz, CD_3_CN): δ 174.6, 147.2, 129.6, 126.9, 126.0, 71.0, 60.8, 59.3, 56.9, 40.7, 35.9, 34.7, 31.2, 29.7, 26.1, 26.0. HRMS (ESI) *m/z*: [M + H]^+^ calculated for C_24_H_37_N_2_O_4_S: 449.2474, found 449.2466.

#### (4-{[(Dimethylaminosulfonyl)-N-methylamino]methyl}-1-oxa-9-aza-9-spiro[5.5]undecyl)(1-phenylcyclopentyl)methanone (2k)

The compound **2k** was prepared from *N*-methylation of **2g**. White foam (56% yield). ^1^H NMR (500 MHz, CD_3_CN): δ 7.34 –7.30 (m, 2H), 7.24–7.18 (m, 3H), 4.15 (br s, 1H), 3.61 (dd, *J* = 11.9, 5.0 Hz, 1H), 3.49–3.18 (m, 2H), 2.96–2.85 (m, 4H), 2.73–2.72 (m, 9H), 2.40–2.27 (m, 2H), 2.13–1.99 (m, 1H), 1.87–1.79 (m, 1H), 1.71–1.66 (m, 5H), 1.53 (d, *J* = 13.1 Hz, 1H), 1.42–1.38 (m, 2H), 1.30–1.11 (m, 1H), 1.03 (qd, *J* = 12.6, 5.1 Hz, 1H), 0.97–0.47 (m, 3H).^13^C NMR (126 MHz, CD_3_CN): δ 174.5, 147.2, 129.6, 126.9, 126.0, 71.0, 60.8, 59.3, 57.6, 40.7, 39.3, 38.5, 36.2, 31.2, 29.7, 26.1 (2C). HRMS (ESI) *m/z*: [M + H]^+^ calculated for C_25_H_40_N_3_O_4_S: 478.2739, found 478.2730.

#### N-Hexyl-N-((9-(1-phenylcyclopentane-1-carbonyl)-1-oxa-9-azaspiro[5.5]undecan-4-yl)methyl)methanesulfonamide (2m)

The compound **2m** was prepared from *N*-hexylation of **2a**. Colorless gum (48% yield). ^1^H NMR (500 MHz, CD_3_CN): δ 7.35–7.28 (m, 2H), 7.24–7.17 (m, 3H), 4.29–4.06 (m, 1H), 3.61 (dd, *J* = 11.8, 4.9 Hz, 1H), 3.53–3.29 (m, 1H), 3.26–2.93 (m, 4H), 2.89–2.77 (m, 3H), 2.75 (s, 3H), 2.41–2.30 (m, 2H), 2.21–2.00 (m, 2H), 1.84–1.82 (m, 2H), 1.73–1.63 (m, 4H), 1.59–1.37 (m, 5H), 1.33–1.27 (m, 7H), 1.02 (qd, *J* = 12.7, 5.2 Hz, 1H), 0.91–0.68 (m, 5H). ^13^C NMR (126 MHz, CD_3_CN): δ 174.5, 147.2, 129.6, 127.0, 126.0, 71.1, 60.9, 59.3, 55.5, 50.2, 40.9, 37.3, 32.1, 30.4, 29.7, 27.1, 26.1, 23.3, 14.3. HRMS (ESI) *m/z*: [M + H]^+^ calculated for C_29_H_47_N_2_O_4_S: 519.3256, found 519.3254.

*N*-Alkylation of carbamate **VIII** using General Procedure-VIII afforded intermediates **XVa***–***b**, followed by Boc deprotection using the General Procedure **I** with 4M HCl in dioxane afforded **4b** and **4d**, and subsequent reaction with sulfonyl chlorides or acid chloride using General Procedure-V furnished **2j**, **2l** and **3m**.

#### Tert-butyl methyl((9-(1-phenylcyclopentane-1-carbonyl)-1-oxa-9-azaspiro[5.5]undecan-4-yl)methyl)carbamate (XVa)

The compound **XVa** was prepared from *N*-methylation of intermediate **VIII**. White foam (73% yield). ^1^H NMR (500 MHz, CD_3_CN): δ 7.35–7.27 (m, 2H), 7.25–7.17 (m, 3H), 4.14–4.04 (m, 1H), 3.59 (dd, *J* = 12.1, 5.0 Hz, 1H), 3.52–3.33 (m, 1H), 3.18–2.92 (m, 4H), 2.84–2.76 (m, 4H), 2.42–2.25 (m, 2H), 2.11–2.00 (m, 1H), 1.89–1.78 (m, 1H), 1.76–1.57 (m, 5H), 1.44–1.37 (m, 11H), 1.31– 1.27 (m, 2H), 1.05 (qd, *J* = 12.6, 5.1 Hz, 1H), 0.96–0.56 (m, 3H). MS (ESI) *m*/*z*: 471.3 [M + H]^+^.

#### Tert-butyl benzyl((9-(1-phenylcyclopentane-1-carbonyl)-1-oxa-9-azaspiro[5.5]undecan-4-yl)methyl)carbamate (XVb)

The compound **XVb** was prepared from *N*-benzylation of intermediate **VIII**. white foam (67% yield). ^1^H NMR (500 MHz, CD_3_CN): δ 7.34–7.29 (m, 4H), 7.26–7.18 (m, 6H), 4.42–4.33 (m, 2H), 4.09–4.05 (m, 1H), 3.56 (dd, *J* = 11.9, 4.9 Hz, 1H), 3.45–3.23 (m, 1H), 3.23–3.07 (m, 1H), 2.97 (s, 3H), 2.78–2.71 (m, 1H), 2.47–2.21 (m, 2H), 2.12–1.98 (m, 1H), 1.91–1.74 (m, 2H), 1.70–1.66 (m, 4H), 1.48–1.35 (m, 11H), 1.32–1.21 (m, 2H), 1.07–0.98 (m, 1H), 0.92–0.48 (m, 3H). MS (ESI) *m*/*z*: 547.3 [M + H]^+^.

#### (4-((Methylamino)methyl)-1-oxa-9-azaspiro[5.5]undecan-9-yl)(1-phenylcyclopentyl)methanone (4b)

The compound **4b** was prepared from Boc deprotection of **XVa**. white foam (87% yield). ^1^H NMR (500 MHz, CD_3_OD): δ 7.36–7.30 (m, 2H), 7.26–7.18 (m, 3H), 4.29–4.20 (m, 1H), 3.76–3.57 (m, 3H), 3.53–3.35 (m, 1H), 3.09–3.01 (m, 1H), 2.93–2.73 (m, 3H), 2.68 (s, 3H), 2.45–1.90 (m, 6H), 1.80–1.67 (m, 4H), 1.62–1.60 (m, 1H), 1.54–1.44 (m, 2H), 1.24–1.16 (m, 1H), 1.10–0.58 (m, 3H). ^13^C NMR (126 MHz, CDCl_3_): δ 176.6, 146.7, 129.9, 127.5, 126.2, 73.6, 72.5, 71.2, 60.8, 62.2, 59.9, 55.9, 43.7, 43.1, 40.4, 34.1, 30.8, 29.4, 26.1. HRMS (ESI) *m/z*: [M + H]^+^ calculated for C_23_H_35_N_2_O_2_: 371.2698, found 371.2691.

#### (4-((Benzylamino)methyl)-1-oxa-9-azaspiro[5.5]undecan-9-yl)(1-phenylcyclopentyl)methanone (4d)

The compound **4d** was prepared from Boc deprotection of **XVb**. white foam (88% yield). ^1^H NMR (500 MHz, CD_3_CN): δ 8.62 (br s, 2H), 7.48–7.45 (m, 2H), 7.43–7.40 (m, 3H), 7.33–7.30 (m, 2H), 7.22–7.17 (m, 3H), 4.11 (s, 2H), 4.07–3.82 (m, 1H), 3.73–3.62 (m, 1H), 3.59 (dd, *J* = 11.7, 4.6 Hz, 1H), 3.53–3.44 (m, 1H), 3.14–2.89 (m, 3H), 2.82–2.74 (m, 3H), 2.35–2.22 (m, 2H), 2.13–1.99 (m, 2H), 1.90–1.77 (m, 1H), 1.72–1.61 (m, 4H), 1.59 (d, *J* = 13.0 Hz, 1H), 1.51–1.47 (m, 1H), 1.10–1.03 (m, 1H), 0.90–0.73 (d, *J* = 65.3 Hz, 3H). ^13^C NMR (126 MHz, CD_3_CN): δ 174.6, 147.1, 132.2, 131.1, 130.3, 129.8, 127.0, 126.0, 71.8, 70.9, 67.6, 66.2, 60.3, 59.3, 53.7, 52.3, 30.7, 39.9, 28.8, 26.1, 26.0, 15.6. HRMS (ESI) *m/z*: [M + H]^+^ calculated for C_29_H_39_N_2_O_2_: 447.3011, found 447.3000.

#### 3-Chloro-N-methyl-N-((9-(1-phenylcyclopentane-1-carbonyl)-1-oxa-9-azaspiro[5.5]undecan-4-yl)methyl)propane-1-sulfonamide (2j)

*N*-sulfonylation of **4b** with 3-chloropropanesulfonyl chloride using the General Procedure-V afforded 3-Chloro-N-methyl-N-((9-(1-phenylcyclopentane-1-carbonyl)-1-oxa-9-azaspiro[5.5]undecan-4-yl)methyl)propane-1-sulfonamide **2j** as white foam. ^1^H NMR (500 MHz, CD_3_CN): δ 7.36–7.29 (m, 2H), 7.23–7.19 (m, 3H), 4.16 (br s, 1H), 3.68 (t, *J* = 6.4 Hz, 2H), 3.61 (dd, *J* = 12.1, 4.8 Hz, 1H), 3.56–3.36 (m, 1H), 3.27–3.09 (m, 1H), 3.09–3.03 (m, 2H), 2.93–2.82 (m, 3H), 2.77 (s, 3H), 2.39– 2.30 (m, 2H), 2.20–2.12 (m, 3H), 2.1–2.00 (m, 1H), 1.9–1.63 (m, 6H), 1.54–1.51 (m, 1H), 1.45–1.34 (m, 2H), 1.29–1.13 (m, 1H), 1.04 (tt, *J* = 12.7, 6.4 Hz, 1H), 0.95–0.57 (m, 3H). ^13^C NMR (126 MHz, CD_3_CN): δ 174.5, 147.2, 129.6, 126.9, 126.0, 71.0, 60.8, 59.3, 56.7, 46.7, 44.2, 40.6, 35.7, 31.1, 29.6, 27.4, 26.1, 26.0. HRMS (ESI) *m/z*: [M + H]^+^ calculated for C_26_H_40_ClN_2_O_4_S: 511.2397, found 511.2394.

#### N-Benzyl-N-((9-(1-phenylcyclopentane-1-carbonyl)-1-oxa-9-azaspiro[5.5]undecan-4-yl)methyl)methanesulfonamide (2l)

*N*-sulfonylation of **4d** with methanesulfonyl chloride using the General Procedure-V afforded *N*-Benzyl-N-((9-(1-phenylcyclopentane-1-carbonyl)-1-oxa-9-azaspiro[5.5]undecan-4-yl)methyl)methanesulfonamide **2l** as a white foam. ^1^H NMR (500 MHz, CDCl_3_): δ 7.35–7.32 (m, 6H), 7.26–7.09 (m, 4H), 4.39 (d, *J* = 14.8 Hz, 1H), 4.29–4.11 (m, 2H), 3.61–3.58 (m, 1H), 3.39–2.98 (m, 3H), 2.95 (d, *J* = 7.2 Hz, 2H), 2.92–2.81 (m, 1H), 2.78 (s, 3H), 2.73 – 2.22 (m, 3H), 2.05–1.90 (m, 2H),1.79– 1.56 (m, 5H), 1.42–1.27 (m, 3H), 1.05 – 0.32 (m, 4H).^13^C NMR (126 MHz, CDCl_3_): δ 174.6, 136.4, 128.9, 128.8, 128.4, 126.3, 125.1, 70.4, 60.3, 58.7, 55.2, 53.6, 41.6, 40.0, 38.5, 37.7, 30.5, 29.8, 29.5, 28.8, 25.4. HRMS (ESI) *m/z*: [M + H]^+^ calculated for C_30_H_41_N_2_O_4_S: 525.2787, found 525.2779.

#### N-methyl-N-((9-(1-phenylcyclopentane-1-carbonyl)-1-oxa-9-azaspiro[5.5]undecan-4-yl)methyl)picolinamide (3m)

Amidation of **4b** with 2-pyridinecarboxylic acid by using the General Procedure-II afforded *N*-methyl-*N*-((9-(1-phenylcyclopentane-1-carbonyl)-1-oxa-9-azaspiro[5.5]undecan-4-yl)methyl)picolinamide **3m** as a white foam. ^1^H NMR (500 MHz, CD_3_CN): δ 8.57–8.54 (m, 1H), 7.99– 7.83 (m, 1H), 7.51–7.40 (m, 2H), 7.38–7.14 (m, 5H), 4.14 (br s, 1H), 3.72–3.61 (m, 1H), 3.58–3.49 (m, 1H), 3.37–3.33 (, 1H), 3.27–3.23 (m, 1H), 3.13–3.11 (m, 2H), 3.00 (s, 3H), 2.88–2.63 (m, 3H), 2.39–2.22 (m, 2H), 2.10–2.04 (m, 1H), 1.81–1.80 (m, 1H), 1.68–1.56 (m, 4H), 1.57 (d, J = 12.6 Hz, 1H), 1.46–1.37 (m, 1H), 1.26–1.08 (m, 2H), 0.97 (br s, 1H), 0.80 (qd, J = 12.5, 5.2 Hz, 1H), 0.61 (br s, 1H). ^13^C NMR (126 MHz, CD_3_CN): δ 174.6, 174.6, 169.3, 168.9, 155.2, 154.9, 148.6, 148.4, 147.1, 147.1, 139.6, 139.1, 129.6, 127.0, 126.0, 126.0, 125.7, 124.3, 124.0, 71.0, 70.9, 60.9, 60.7, 59.3, 57.1, 54.1, 40.8, 40.3, 38.0, 34.1, 31.3, 30.9, 30.1, 29.5, 26.1, 26.0. (additional carbon signals observed due to rotamers). HRMS (ESI) m/z: [M + H]^+^ calculated for C_29_H_38_N_3_O_3_: 476.2913, found 476.2903.

#### *Synthesis of* (4-(aminomethyl)-1-oxa-9-azaspiro[5.5]undecan-9-yl)(1-phenylcyclopentyl)methanone hydrochloride 4ª

Boc deprotection of **VIII** using General Procedure-I with 4M HCl/dioxane to afford (4-(aminomethyl)-1-oxa-9-azaspiro[5.5]undecan-9-yl)(1-phenylcyclopentyl)methanone hydrochloride **4a** as a white solid (86% yield). ^1^H NMR (400 MHz, CD_3_OD): δ 7.36–7.30 (m, 2H), 7.26–7.18 (m, 3H), 4.23 (br s, 1H), 3.80–3.32 (m, 5H), 3.12–2.89 (m, 1H), 2.89 (s, 1H), 2.77–2.67 (m, 2H), 2.59–1.84 (m, 6H), 1.73–1.62 (m, 6H), 1.55–1.40 (m, 2H), 1.25–0.59 (m, 4H). ^13^C NMR (100 MHz, CD_3_OD): δ 176.6, 146.7, 129.9, 127.5, 126.2, 73.6, 72.5, 71.3, 62.2, 60.9, 59.9, 46.0, 43.7, 43.2, 40.4, 39.5, 38.0, 30.8, 30.2, 26.0. HRMS (ESI) *m/z*: [M + H]^+^ calculated for C_22_H_33_N_2_O_2_: 357.2542, found 357.2532.

#### Synthesis of 2n, 2o and 3e

Intramolecular cyclization of **2c** and **2d** using the General Procedure-VI with Cs_2_CO_3_/DMF afforded **2n** and **2o** respectively. Alternatively, Intramolecular cyclization of **3e** using the General Procedure-VI with NaH/THF furnished **3n**

#### (4-((1,1-Dioxidoisothiazolidin-2-yl)methyl)-1-oxa-9-azaspiro[5.5]undecan-9-yl)(1-phenylcyclopentyl)methanone (2n)

White foam (65% yield). ^1^H NMR (500 MHz, CD_3_CN): δ 7.34–7.31 (m, 2H), 7.23–7.19 (m, 3H), 4.13 (br s, 1H), 3.61 (dd, *J* = 11.9, 5.0 Hz, 1H), 3.55–3.36 (m, 1H), 3.23–3.10 (m, 3H), 3.07–3.04 (m, 2H), 3.02–2.75 (m, 2H), 2.74–2.64 (m, 2H), 2.49–2.29 (m, 2H), 2.28–2.20 (m, 2H), 2.10–2.03 (m, 1H), 1.81–1.79 (m, 2H), 1.71–1.66 (m, 4H), 1.57 (d, *J* = 13.1 Hz, 1H), 1.48–1.44 (m, 3H), 1.03 (qd, *J* = 12.6, 5.1 Hz, 1H), 0.84–0.71 (m, 3H). ^13^C NMR (126 MHz, CD_3_CN): δ 174.6, 147.2, 129.6, 126.9, 126.0, 71.0, 60.8, 59.3, 51.8, 48.7, 47.2, 40.7, 31.3, 30.0, 26.1, 26.0, 19.6. HRMS (ESI) *m/z*: [M + H]^+^ calculated for C_25_H_37_N_2_O_4_S: 461.2474, found 461.2465.

#### (4-((1,1-dioxido-1,2-thiazinan-2-yl)methyl)-1-oxa-9-azaspiro[5.5]undecan-9-yl)(1-phenylcyclopentyl)methanone (2°

White foam (61% yield). ^1^H NMR (500 MHz, CD_3_CN): δ 7.36–7.29 (m, 2H), 7.25–7.18 (m, 3H), 4.15 (br s, 1H), 3.60 (dd, *J* = 11.9, 4.9 Hz, 1H), 3.47–3.38 (m, 1H), 3.31–3.18 (m, 3H), 2.99–2.97 (m, 1H), 2.95–2.90 (m, 2H), 2.88–2.77 (m, 3H), 2.46–2.25 (m, 2H), 2.15–1.99 (m, 3H),1.86–1.67 (m, 2H), 1.73–1.64 (m, 4H), 1.57–1.52 (m, 3H), 1.44–1.40 (m, 2H), 1.20–1.11 (m, 1H), 1.02 (tt, *J* = 12.6, 6.3 Hz, 1H), 0.83–0.69 (m, 3H). ^13^C NMR (126 MHz, CD_3_CN): δ 174.6, 147.2, 129.6, 126.9, 126.0, 71.0, 60.8, 59.3, 54.1, 51.2 48.0, 40.7, 31.2, 30.7, 26.1, 26.0, 24.6, 20.8. HRMS (ESI) *m/z*: [M + H]^+^ calculated for C_26_H_39_N_2_O_4_S: 475.2630, found 475.2622.

#### 1-((9-(1-Phenylcyclopentane-1-carbonyl)-1-oxa-9-azaspiro[5.5]undecan-4-yl)methyl)pyrrolidin-2-one (3n)

White foam (44% yield). ^1^H NMR (500 MHz, CD_3_CN): δ 7.34–7.31 (m, 2H), 7.24–7.18 (m, 3H), 4.14 (br s, 1H), 3.59 (dd, *J* = 11.9, 5.0 Hz, 1H), 3.49–3.40 (m, 1H), 3.35–3.23 (m, 2H), 3.18–2.92 (m, 4H), 2.84–2.81 (m, 1H), 2.37–2.33 (m, 2H), 2.21–1.97 (m, 3H), 1.89–1.77 (m, 1H), 1.76–1.53 (m, 5H), 1.43 (d, *J* = 13.3 Hz, 2H), 1.33–1.29 (m, 1H), 1.25–1.11 (m, 1H), 1.03 (qd, *J* = 12.6, 5.1 Hz, 1H), 0.99–0.84 (m, 3H). ^13^C NMR (126 MHz, CD_3_CN): δ 175.6, 174.5, 147.1, 129.6, 126.9, 126.0, 71.0, 60.8, 59.3, 49.1, 48.3, 40.8, 31.4, 31.3, 29.5, 26.1, 26.0, 18.7. HRMS (ESI) *m/z*: [M + H]^+^ calculated for C_26_H_37_N_2_O_3_: 425.2804, found 425.2792.

#### Synthesis of compound 4c and 3o

To a suspension of (4-(aminomethyl)-1-oxa-9-azaspiro[5.5]undecan-9-yl)(1-phenylcyclopentyl)methanone hydrochloride **4a** (20 mg, 0.05 mmol, 1 eq) in dioxane (0.4 mL) were added DIPEA (18 μL, 0.1 mmol, 2 eq) and 2,2,2-Trifluoroethyl trifluoromethylsulfonate (9 μL, 0.06 mmol, 1.2 eq) and the mixture was stirred at 25 °C for 30 min, then heated at 50 °C for 12 h. After completion (based on TLC and LCMS), the reaction mixture was concentrated, and the resulting crude product was purified using preparative HPLC to afford (1-Phenylcyclopentyl)(4-(((2,2,2-trifluoroethyl)amino)methyl)-1-oxa-9-azaspiro[5.5]undecan-9-yl)methanone **4c** as a white foam (14 mg, 63% yield). ^1^H NMR (500 MHz, CD_3_OD): δ 7.35–7.30 (m, 2H), 7.26–7.18 (m, 3H), 4.24 (br s, 1H), 3.91–3.51 (m, 4H), 3.41–3.33 (m, 1H), 3.09–3.03 (m, 1H), 2.95–2.87 (m, 3H), 2.46–2.17 (m, 4H), 2.08–1.56 (m, 8H), 1.51–1.47 (m, 2H), 1.24–0.93 (m, 3H), 0.82–0.67 (m, 1H).^13^C NMR (126 MHz, CD_3_OD): δ 176.6, 162.7, 162.4, 146.7, 129.9, 127.5, 126.2, 121.2 (q, *J* = 290 Hz), 71.3, 60.8, 59.9, 55.6, 43.2, 40.6, 31.0, 29.4, 26.1. HRMS (ESI) *m/z*: [M + H]^+^ calculated for C_24_H_34_F_3_N_2_O_2_: 439.2572, found 439.2563.

#### 1-((9-(1-Phenylcyclopentane-1-carbonyl)-1-oxa-9-azaspiro[5.5]undecan-4-yl)methyl)pyrrolidine-2,5-dione (3o)

To a stirred solution of amine salt (70 mg, 0.18 mmol, 1.0 eq) and succinic anhydride (20 mg, 0.20 mmol, 1.1 eq) in acetic acid (0.7 mL), a catalytic amount of DMAP was added. The solution was refluxed for 24 h. Upon completion of the reaction (based on TLC and LCMS), the solution was cooled to room temperature, and the solvent was removed under reduced pressure. The residue was dissolved in dichloromethane (50 mL), and the organic layer was washed with saturated NaHCO_3_ solution (10 mL) and 1 N HCl (10 mL) solution. The combined organic layers dried over anhydrous Na_2_SO_4_, filtered, and concentrated. The crude was purified by combi flash using eluent 80% EtOAc in hexane to afford 1-((9-(1-Phenylcyclopentane-1-carbonyl)-1-oxa-9-azaspiro[5.5]undecan-4-yl)methyl)pyrrolidine-2,5-dione **3o** as a white foam (30 mg, 37% yield). ^1^H NMR (500 MHz, CD_3_CN): δ 7.35–7.28 (m, 2H), 7.23–7.19 (m, 3H), 4.15 (br s, 1H), 3.57 (dd, *J* = 11.9, 5.0 Hz, 1H), 3.47–3.30 (m, 1H), 3.18 (h, *J* = 6.4 Hz, 3H), 2.93–2.70 (m, 2H), 2.58–2.36 (m, 6H), 2.16–2.08 (m, 2H), 1.81–1.65 (m, 5H), 1.43 (d, *J* = 13.3 Hz, 2H), 1.39–1.18 (m, 2H), 1.06 (qd, *J* = 12.7, 5.1 Hz, 1H), 0.89–0.65 (m, 3H). ^13^C NMR (126 MHz, CD_3_CN) δ 178.9, 174.5, 147.1, 129.6, 126.9, 126.0, 70.9, 60.7, 59.3, 44.9, 40.7, 30.2 31.2, 28.8, 26.1, 26.0. HRMS (ESI) *m/z*: [M + H]^+^ calculated for C_26_H_35_N_2_O_4_: 439.2597, found 439.2588.

#### Synthesis of 4-(Aminomethyl)-4-hydroxy-1-oxa-9-azaspiro[5.5]undecan-9-yl)(1-phenylcyclopentyl)methanone (4e)

A solution of 9-(1-Phenylcyclopentane-1-carbonyl)-1-oxa-9-azaspiro[5.5]undecan-4-one **V** (200 mg, 0.586 mmol, 1 eq) in acetonitrile (2 mL) was treated with TMSCN (87.2 mg, 879 mmol, 1.5 eq) in the presence of Cu(OTf)_2_ (10.6 mg, 0.0293 mmol, 0.05 eq) in dry acetonitrile (2 mL) at rt for 3 h. The reaction mixture was diluted with DCM (20 mL) and washed with sat. NaHCO_3_. The combined organic layers dried over anhydrous Na_2_SO_4_, filtered, and concentrated. The solvent removed in vacuo to obtain the cyanohydrin crude. The crude product was used in the next step without further purification. Next, to a stirred solution of cyanohydrin (250 mg, 0.567 mmol, 1eq) in dry THF (5 mL) at 0 °C was added LiAlH_4_ (53.8 mg, 1.42 mmol, 2.5 eq) and stirred at the same temperature for 1 h. Upon completion, add EtOAc followed by methanol and water to quench the excess LiAlH_4_. The reaction mixture was filtered through celite bed and filtrate concentrated under reduced pressure obtain crude product which was subjected to prep HPLC to afford tertiary alcohol (4-(aminomethyl)-4-hydroxy-1-oxa-9-azaspiro[5.5]undecan-9-yl)(1-phenylcyclopentyl)methanone **4e** (colorless gum, 84 mg, 41% yield over two steps). ^1^H NMR (500 MHz, CD_3_CN): δ 7.35–7.29 (m, 2H), 7.25–7.19 (m, 5H), 4.30–3.61 (m, 4H), 3.56–3.51 (m, 1H), 3.17–2.79 (m, 5H), 2.38–2.29 (m, 3H), 2.12–1.98 (m, 1H), 1.78–1.58 (m, 4H), 1.56–1.46 (m, 3H), 1.44–1.27 (m, 1H), 1.37–1.17 (s, 1H), 1.06–0.65 (m, 2H). ^13^C NMR (126 MHz, CD_3_CN) ^13^C NMR (126 MHz, CD_3_CN) δ 174.5, 147.2, 129.5, 126.9, 126.0, 70.7, 69.9, 59.3, 57.2, 55.3, 40.0, 35.3, 26.1. HRMS (ESI) *m/z*: [M + H]^+^ calculated for C_22_H_33_N_2_O_3_: 373.2491, found 373.2481.

### Biological Assays

#### CHIKV-nLuc Reporter Assay

MRC5 hTERT fibroblasts cultured in DMEM supplemented with 10% FBS and 1% penicillin-streptomycin were treated with an eight-point four-fold serial dilution of compound, NHC, (MedChemExpress, HY-125033) or DMSO one hour prior to infection with nLuc reporter virus for CHIKV (181/25-nLuc-Capsid)^16^ at an MOI of 0.1 or left uninfected. After 6 h, cells infected with virus were treated with Nano-Glo luciferase assay reagent to detect nLuc luminescence. To assess cell viability, uninfected cells were treated with CellTiterGlo 2.0 reagent (Promega). In each case, luminescence was measured on Promega GloMax plate reader. Percent inhibition and viability were calculated by normalization to DMSO control wells. EC_50_ values were calculated from compounds run in technical and biological triplicate using a three-parameter Hill equation.

#### CHIKV nsP2 ATPase Assay

CHIKV nsP2 ATPase activity was tested using the Promega ADP-glo assay kit in a 384-well format. CHIKV nsP2 sequence from an attenuated vaccine strain (181/25)^15^ was cloned into a pET28 vector with a N-terminal TwinStrep 6-His SUMO tag and expressed in Rosetta 2 E. coli (Novagen) for 48h at 15 °C. E.coli were lysed in a sodium phosphate buffer (50 mM NaHPO_4_ pH 7.8, 20% glycerol, 500 mM NaCl, 1 mM TCEP) by sonication, nucleic acids were precipitated using polyethylenamine, and the lysate was clarified by centrifugation at 70,000 g for 30 min. Protein from the clarified lysate was precipitated using ammonium sulfate to 40% saturation and the precipitated proteins including nsP2 were pelleted by centrifugation at 70,000 g for 30 min. The pellet was resuspended in sodium phosphate buffer and run over a Streptactin-XT column (Cyvita) and washed with 100 column volumes of buffer prior to elution by on column cleavage using sumo protease ULP1. The eluant was concentrated and dialyzed into a HEPES buffer (25 mM HEPES pH 7.4, 250 mM NaCl, 20% glycerol, 1 mM TCEP). 1 nM of purified CHIKV nsP2 was incubated with compound diluted in an 8-point 3-fold dose response starting at 250 µM for 30 min prior to addition of 30 µM ATP. After 1 h at room temperature 2 µL of ADP-Glo reagent (Promega) was added and the reaction was incubated for 40 min followed by the addition of Kinase detection reagent. After 30 min the assay plate was read for luminescence using a Tecan Spark. Percent inhibition of ATPase activity was determined by comparing compound treated samples to DMSO after subtracting background samples containing substrate and no enzyme. IC_50_ values were calculated from compounds run in technical and biological triplicate using a three-parameter Hill equation.

For kinetic experiments, reactions were performed at 30 °C. in a buffer containing 25 mM HEPES pH 7.5, 1 mM ATP, 1 mM Mg acetate, 1 mM TCEP, 50 mM K glutamate, 10% (vol/vol) DMSO, and 0.05 µCi/µL [α^32^P]-ATP. Reactions were initiated by the addition of 20 nM nsP2 protein. Reactions were quenched at various times by addition of EDTA to 250 mM. nsP2 was diluted immediately prior to use in 25 mM HEPES pH 7.5, 1 mM TCEP and 20% glycerol. The volume of enzyme added to any reaction was always less than or equal to one–tenth the total volume. 1 µL of the quenched reaction was spotted onto polyethyleneimine-cellulose TLC plates (EM Science). TLC plates were developed in 0.3 M potassium phosphate, pH 7.0, dried and exposed to a PhosphorImager screen. TLC plates were visualized by using a PhosphorImager and quantified by using the ImageQuant software (Molecular Dynamics) to determine the amount of ATP hydrolyzed to ADP. The amount of ADP was plotted as a function of time. For determination of K_m_ the ATPase assay was performed using 0.5 nM nsP2 mixed with increasing concentrations of ATP in the presence of 0, 1.0 or 2.0 μM of **2o**.

#### Alphavirus Titer Reduction Assay

Primary Normal Human Dermal Fibroblast (NHDF) cells were cultured and plated onto 48-well cell plates at a cell density of 40,000 cells/well and incubated at 37 °C and 5% CO_2_. Compound **2o** (10 mM in DMSO) was used to prepare an 11-point dose-response curve ranging from 40 µM to 0.039 µM by diluting 1:1 with DMEM supplemented with 5% fetal bovine serum (FBS) and 1x penicillin-streptomycin-glutamine (PSG). Media-only and DMSO (5 µL/mL) were included as negative controls. 1 h before infection, 200 µL of diluted **2o** and controls were added to the 48-well plates of confluent NDHF cells in triplicate. Cells were infected with a multiplicity of infection equal to 1 in a volume of 100 µL. At 2 hpi, infection media were removed, cells were washed two times with PBS, and fresh media containing **2o** dilution series was added to the cells. At 24 hpi, 20 µL of supernatants were transferred to 96-well plates and a limiting dilution plaque assay was performed on a confluent monolayer of Vero cells to determine levels of virus production. For the plaque assay, 10-fold serial dilutions were made in the 96-well plates and 100 µL of each dilution was transferred onto Vero cells of a 48-well plate and incubated with continuous rocking. At 2 hpi, the cells were overlaid with 250 µL of 0.3% high-/0.3% low-viscosity carboxymethyl cellulose prepared in DMEM supplemented with 5% FBS and 1x PSG. At 48–72 hpi depending upon the virus used in the assay, the plates were fixed with 3.7% formalin and stained with 0.4% methylene blue dye. Plaques were enumerated using a dissecting microscope. Data was transformed and normalized before curve fitting with GraphPad Prism v10 software.

#### CHIKV nsP2 helicase assay

dsRNA substrates were produced by annealing 100 nM RNA oligonucleotides (U20-ss9-U10: UUUUUUUUUUUUUUUUUUUUGCCGCCCGGUUUUUUUUUU and ss9-U10: CCGGGCGGCUUUUUUUUUU) in 25 mM HEPES pH 7.5 and 50 mM NaCl in a Progene Thermocycler (Techne) with a molar ratio of 1:1.4 for labeled:unlabeled strands. Annealing reaction mixtures were heated to 90 °C for 1 min and slowly cooled (5 °C/min) to 10 °C. Reactions were performed at 30 °C. 50 nM nsP2 protein was mixed with 1 nM [^32^P]-labeled unwinding substrate in 25 mM HEPES pH 7.5, 5 mM ATP, 5 mM Mg acetate, 1 mM TCEP, 20 mM KGlutamate, 1% (vol/vol) DMSO, and 0.5 µM RNA Trap. Reactions were initiated by the addition of 50 nM nsP2. The trapping strand was a 9-mer RNA (CCGGGCGGC) that was complementary to the ^32^P-labeled displaced strand. Reactions were quenched at various times by addition 2X Quench Buffer (200 mM EDTA, 0.66% SDS, 2 mM (UMP) poly(rU), 0.05% Bromophenol Blue, 0.05% Xylene Cyanol, and 10% Glycerol). nsP2 was diluted immediately prior to use in 25 mM HEPES pH 7.5, 1 mM TCEP and 20% glycerol. The volume of enzyme added to any reaction was always less than or equal to one–tenth the total volume. 10 µL of the quenched reaction mixtures was loaded on a 20% acrylamide, 0.53% bisacrylamide native polyacrylamide gel containing 1× TBE. Electrophoresis was performed in 1× TBE at 15 mA. Gels were visualized by using a phosphorimager and quantitated by using the ImageQuant software (Molecular Dynamics) to determine the amount of strand separation. The fraction of RNA that had been unwound by the helicase was corrected for the efficiency of the trapping strand in preventing reannealing of products. This correction factor was determined for each experiment by heating a sample of the RNA substrate at 95 °C for 1 min then cooled to 10 °C at a rate of 5 °C/min in the presence of the trapping strand. The quantity of unwinding substrate that was prevented from reannealing was typically ∼98%. The amount of RNA unwound was plotted as a function of time.

#### Fluorescence polarization assay

Increasing concentrations of nsP2 were mixed with 1 nM of fluorescein-labeled dsRNA in a binding buffer containing 25 mM HEPES pH 7.5, 1 mM Mg acetate, 1 mM TCEP, 100 mM KGlutamate, and 1% (vol/vol) DMSO. Reactions were incubated at room temperature for 10 min in a 384-well non-binding plate and then read with a dual wavelength 485/20 485/20 fluorescence polarization cube using a Biotek Synergy H1 plate reader. Data from protein titration experiments were fit to a hyperbola: 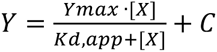 where X is the concentration of protein, Y is degree of polarization, K_d,app_ is the apparent dissociation constant, and Y_max_ is the maximum value of Y.

#### ^19^F NMR

CHIKV nsP2hel domain (UniProt A0A1U9VX00 aa 536-1000) was subcloned into a pET28-derived *E. coli* expression vector, pET28-MHL, generating a construct with an N-terminal His_6_-tag followed by a TEV cleavage site. The construct was transformed into *E. coli* BL21 (DE3) pRARE2 competent cells. An overnight starter culture was grown in LB media supplemented with kanamycin and chloramphenicol. This starter culture was then used to inoculate TB media supplemented with the same antibiotics. Cultures were grown at 37 °C until an OD_600_ of 0.8 was reached, at which point the temperature was reduced to 16 °C. Protein expression was induced with 0.5 mM IPTG using a LEX system, and cultures were grown overnight. Cells were harvested by centrifugation, and the resulting cell pellets were stored at −80 °C. For purification the supernatant was incubated with nickel affinity resin in an open column, and the protein was eluted with 250 mM imidazole. The eluted fractions were subjected to TEV protease cleavage overnight at 4 °C. The cleaved protein was separated by incubating the sample with a fresh nickel resin, collecting only the flowthrough (unbound fraction). The protein was further purified by size exclusion chromatography with a HiLoad 16/60 Superdex 200 gel filtration column on an ÄKTA Pure chromatography system (GE Healthcare) and eluted in a buffer containing 20 mM Hepes (pH 7.5), 200 mM NaCl, 5% glycerol, and 2 mM TCEP. Protein fractions containing pure nsP2hel protein, as confirmed by SDS-PAGE, were pooled together and concentrated up to 8.25 mg/mL using a 3 kDa cutoff protein spin concentrator, and flash-frozen in liquid nitrogen and stored at −80 °C. Spectra of the compound **7b** (15 μM) were acquired on a Bruker Avance III spectrometer operating at 600 MHz, equipped with a QCI probe at 298K, and collected with and without the presence of CHIKV nsP2hel protein (20 or 40 μM, buffered in 20 mM HEPES pH 7.5, 200 mM NaCl, 5% glycerol, 2 mM TCEP and 1% DMSO). TFA (100 μM) was added as an internal reference. 1.5k transients were acquired over a sweep width of 150 ppm; an exponential window function (LB=5 Hz) was applied prior to Fourier transformation.

### Kinetic Solubility Assay

Kinetic solubility of final compounds was determined in triplicate in pH 7.4 phosphate buffer using 10 mM stock solutions at 2.0 % DMSO and CLND detection at Analiza, Inc.

## Supporting Information

Table S1, VEEV-nLuc data. Figure S1–S5, Variable temperature 2D NMR of **2n**. Figure S6, Sequence and structure of CHIKV nsP2hel. Figure S7, Inhibition of CHIKV-nLuc replication by NHC. Figure S8, Correlation of CHIKV-nLuc with VEEV-nLuc (PDF).

## Author Contributions

Conceptualization, T.M.W., M.T.H., N.J.M., C.J.C.; investigation, M.B., J.D.S., K.M.T., Y.S., S.H., M.H., Z.W.D., C.Z., H.J.O., P.J.B., M.K.S., S.R.M., D.O., J.B., I.L., N.M., S.A.M., P.A.L., J.G.P., A.M.D., T.E.M., Z.J.S., D.N.S.; methodology, M.B., J.D.S., K.M.T., T.J.M., D.N.S., C.H.A., A.M.V., R.M.C., L.H., J.J.A., C.J.C. M.T.H., N.J.M., T.M.W.; writing—original draft, T.M.W., M.B.; writing—review and editing, M.T.H., N.J.M., C.J.C., Z.W.D., H.J.O., P.J.B., M.K.S., P.A.L., T.E.M., L.H., K.T.M., C.E.C.. All authors have read and agreed to the published version of the manuscript.

## Data Availability

The original contributions presented in the study are included in the article/Supplementary Material, further inquiries can be directed to the corresponding author.

## Acknowledgments

The authors acknowledge the support of Piramal Pharma Solutions for synthesis of chemical intermediates. We thank Alex Tropsha (UNC) and Konstantin Popov (UNC) for the original sequence analysis that led to nomination of the CHIKV-nLuc point mutants.

## Abbreviations

CHIKV: Chikungunya virus
MAYV: Mayaro virus
VEEV: Venezuelan equine encephalitis virus
DME: dimethoxyethane
nLuc: nanoluciferase
hpi: hours post infection

