## Supporting Information for "Identification of Direct-acting nsP2 Helicase Inhibitors with Antialphaviral Activity"

#### Table of Contents


| | | |
|---|---|---|
| Table S1 | VEEV-nLuc data | S2 |
| Figure S1 | $^{13}$C NMR of **2n** (126 MHz, Acetone-$d_6$, $CD_3OD$, DMSO-$d_6$ and $CDCl_3$) | S3 |
| Figure S2 | $^{13}$C NMR of **2n** (126 MHz, $CD_3CN$ and Pyridine-$d_5$) | S4 |
| Figure S3 | $^{1}$H-$^{13}$C HSQC of **2n** (500 MHz, 60 ºC, Pyridine-$d_5$) | S5 |
| Figure S4 | $^{1}$H-$^{13}$C HMQC of **2n** (500 MHz, 60 ºC, Pyridine-$d_5$) | S6 |
| Figure S5 | $^{1}$H-$^{13}$C HSQC of **2n** (500 MHz, 0 ºC, Pyridine-$d_5$) | S7 |
| Figure S6 | $^{13}$C NMR of **2n** (126 MHz, 0 ºC, Pyridine-$d_5$ and $CD_3CN$) | S8 |
| Figure S7 | Sequence and structure of CHIKV nsP2hel | S9 |
| Figure S8 | Inhibition of CHIKV-nLuc replication by NHC | S10 |
| Figure S9 | Correlation of CHIKV-nLuc with VEEV-nLuc | S11 |
| Figure S10 | Analytical spectra new compounds | S12–S162 |

**Table S1.** VEEV-nLuc data

| Compound | VEEV-nLuc ($pEC_{50}$) | Compound | VEEVnLuc ($pEC_{50}$) |
|---|---|---|---|
| **1a** | 4.7 | **3p** | 5.4 |
| **2a** | 5.1 | **3q** | <4.4 (60)[a] |
| **1b** | <4.4 (60)[a] | **3r** | <4.4 (50)[a] |
| **1c** | <4.4 (80)[a] | **3s** | <4.4 (70)[a] |
| **1d** | 4.8 | **3t** | <4.4 (80)[a] |
| **2b** | 4.9 | **3u** | <4.4 (40)[a] |
| **2c** | 5 | **4a** | <4.4 (40)[a] |
| **2d** | 5 | **4b** | 4.7 |
| **2e** | <4.4 (50)[a] | **4c** | <4.4 (60)[a] |
| **2f** | <4.4 (90)[a] | **4d** | 5 |
| **2g** | 5.1 | **4e** | <4.4 (70)[a] |
| **2h** | 4.5 | **5a** | <4.4 (20)[a] |
| **2i** | 4.9 | **5b** | <4.4 (40)[a] |
| **2j** | 5 | **5c** | <4.4 (20)[a] |
| **2k** | 4.9 | **5d** | <4.4 (60)[a] |
| **2l** | <4.4 (70)[a] | **5e** | <4.4 (20)[a] |
| **2m** | 5 | **5f** | <4.4 (<10)[a] |
| **2n** | 5.8 | **5g** | <4.4 (40)[a] |
| **2o** | 6.3 | **6a** | <4.4 (20)[a] |
| **3a** | 4.7 | **6b** | <4.4 (<10)[a] |
| **3b** | 4.9 | **6c** | <4.4 (30)[a] |
| **3c** | <4.4 (70)[a] | **6d** | <4.4 (<10)[a] |
| **3d** | 4.6 | **6e** | 5.0 |
| **3e** | 5.5 | **6f** | 4.7 |
| **3f** | 4.8 | **6g** | 4.9 |
| **3g** | 4.7 | **6h** | <4.4 (50)[a] |
| **3h** | 5.2 | **6i** | 5.7 |
| **3i** | <4.4 (60)[a] | **6j** | 5.1 |
| **3j** | 4.9 | **6k** | <4.4 (40)[a] |
| **3k** | 4.9 | **6l** | <4.4 (50)[a] |
| **3l** | <4.40 (40)[a] | **6m** | <4.4 (<10)[a] |
| **3m** | <4.4 (60)[a] | **7a** | 5.8 |
| **3n** | 5.4 | **7b** | 5.8 |
| **3o** | 6.3 | **7c** | <4.4 (80)[a] |

[a]Value in parentheses is % inhibition at 40 μM

**Figure S1.** $^{13}$C NMR of **2n** (126 MHz, acetone-$d_6$, $CD_3OD$, DMSO-$d_6$ and $CDCl_3$)

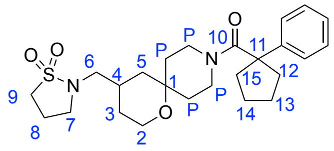

**2n**

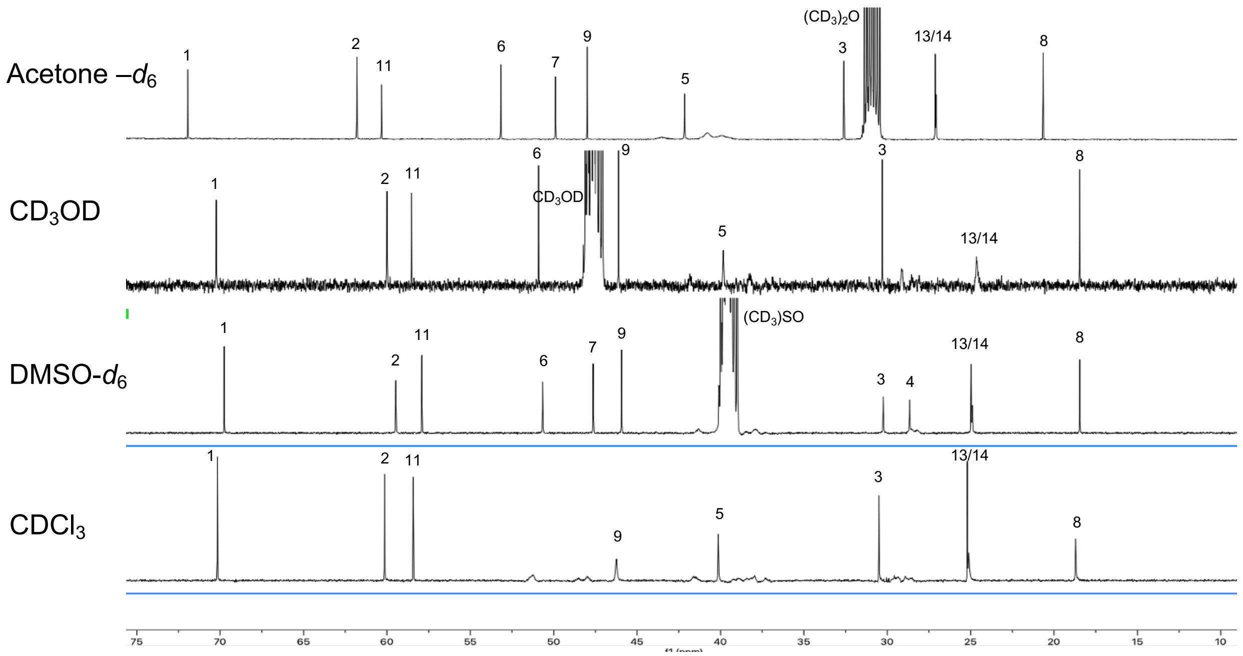

**Figure S2.** $^{13}$C NMR of **2n** (126 MHz, $CD_3CN$ and pyridine-$d_5$)

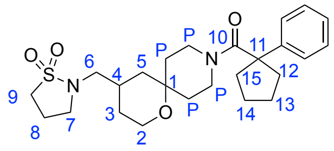

**2n**

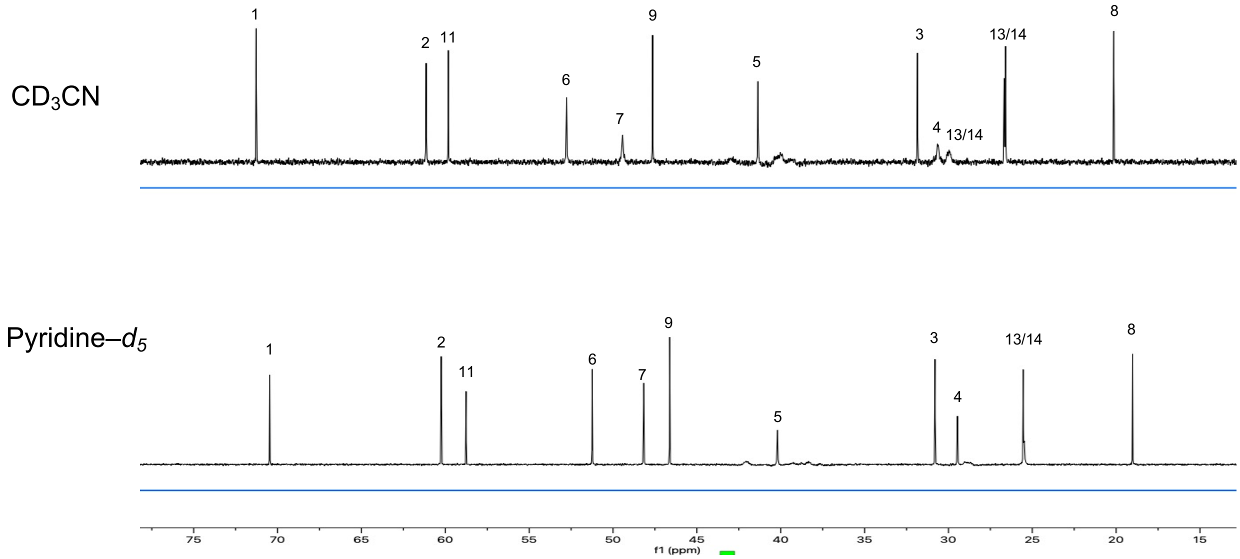

**Figure S3.** $^1$H-$^{13}$C HSQC of **2n** (500 MHz, 60 ºC, pyridine-$d_5$)

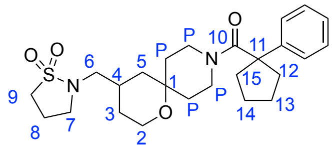

**2n**

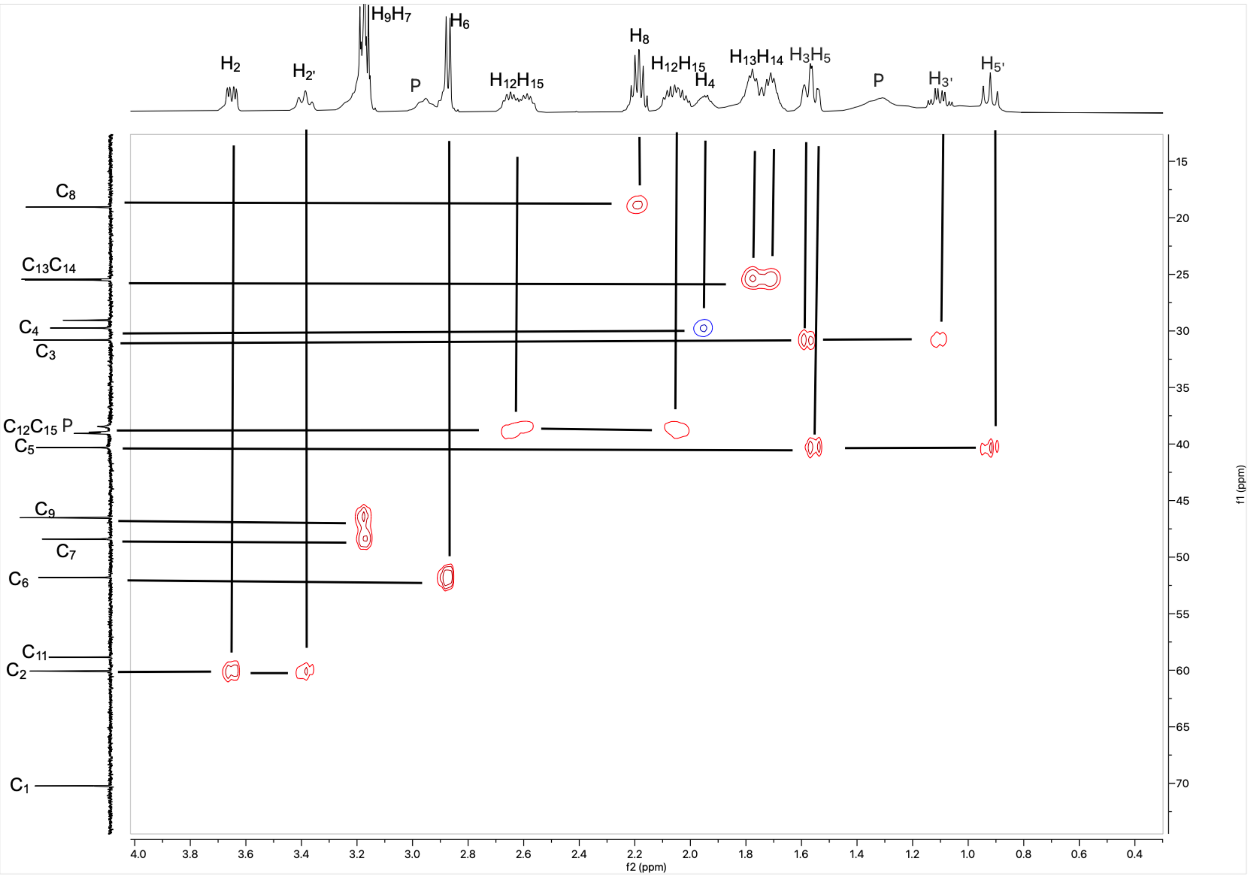

**Figure S4.** $^{1}H$-$^{13}C$ HMQC of **2n** (500 MHz, 60 ºC, pyridine-*d₅*)

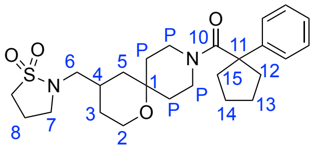


**2n**

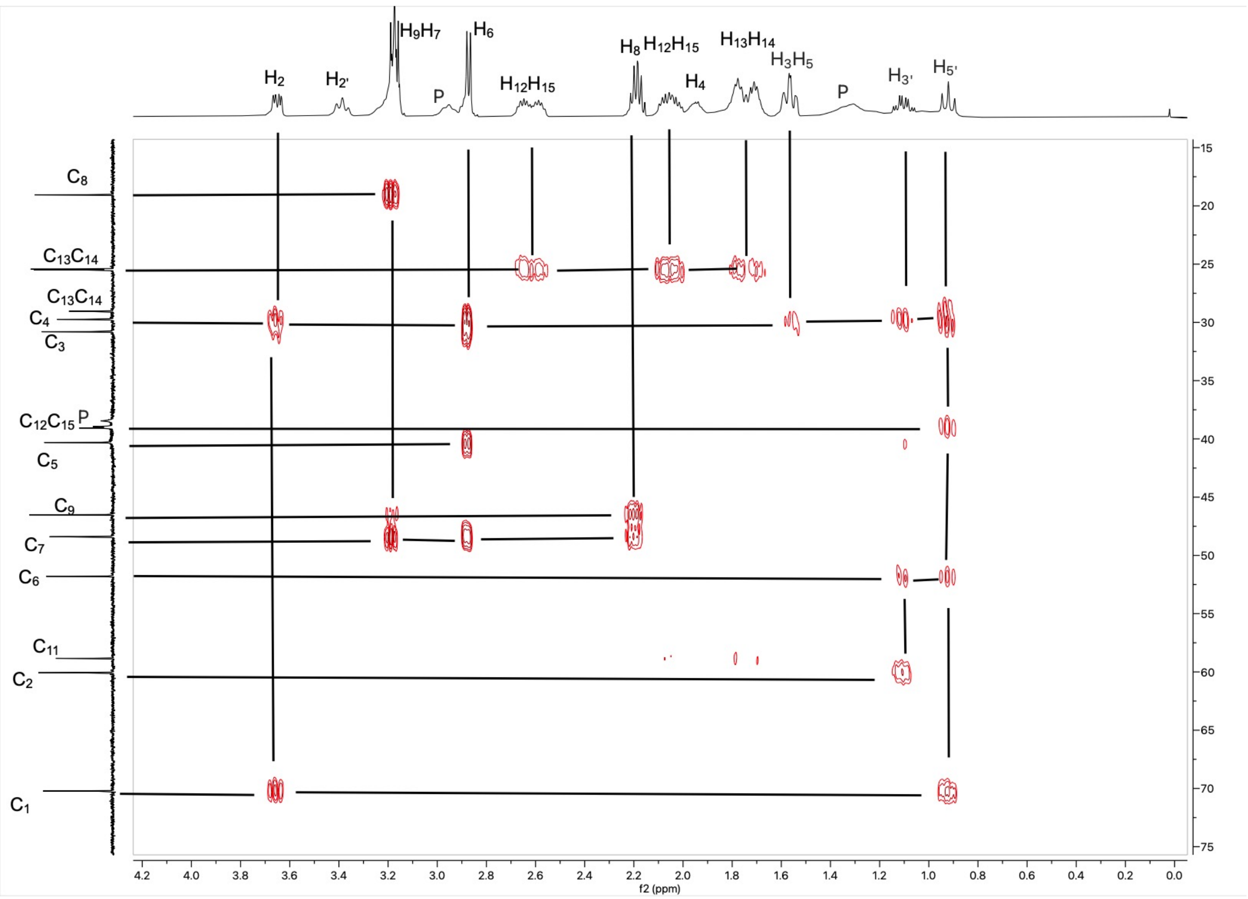

**Figure S5.** $^{1}H$-$^{13}C$ HSQC of **2n** (500 MHz, 0 ºC, pyridine-$d_5$)

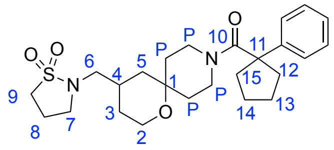


**2n**

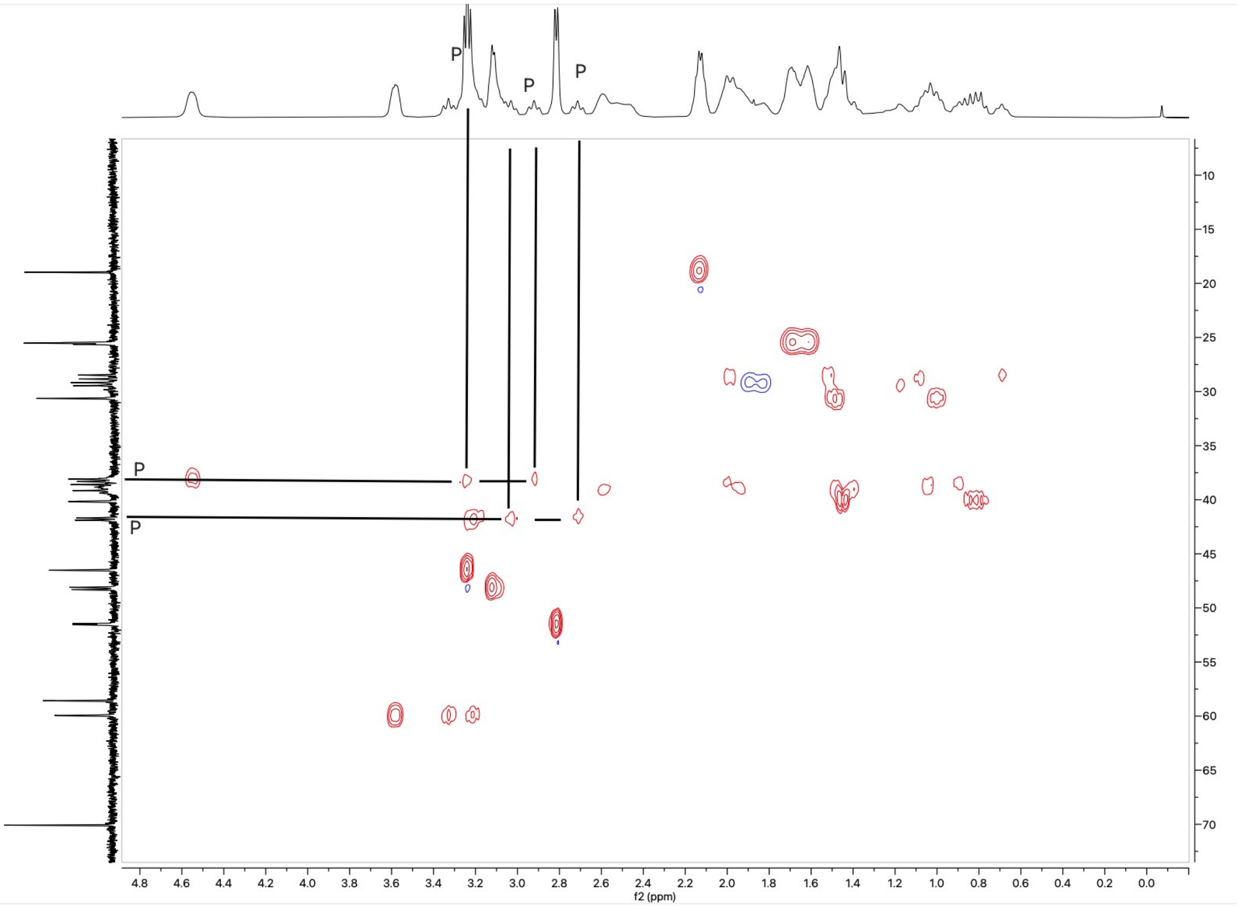

**Figure S6.** $^{13}$C NMR of **2n** (126 MHz, 0 ºC, pyridine-$d_5$ and $CD_3CN$)

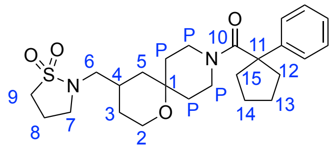

**2n**

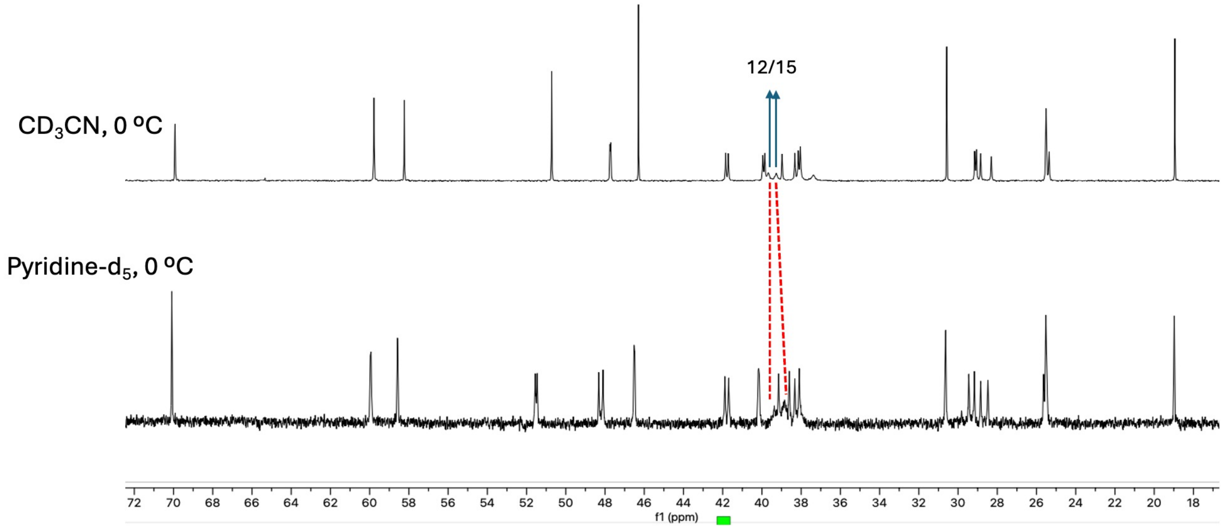

**Figure S7.** Sequence and structure of CHIKV nsP2hel

```
CHIKV nsP2 GIIETPRGAIKVTAQPTDHVVGEYLVLSPQTVLRSQKLSLIHALAEQVKTCTHNGRAGRY 60
CHIKV nsP2 AVEAYDGRVLVPSGYAISPEDFQSLSESATMVYNEREFVNRKLHHIAMHGPALNTDEESY 120
CHIKV nsP2 ELVRAERTEHEYVYDVDQRRCCKKEEAAGLVLVGDLTNPPYHEFAYEGLKIRPACPYKIA 180
CHIKV nsP2 VIGVFGVPGSGKSAIIKNLVTRQDLVTSGKKENCQEISTDVMRQRGLEITARTVDSLLLN 240
CHIKV nsP2 GCNRPVDVLYVDEAFACHSGTLLALIALVRPRQKVVLCGDPKQCGFFNMMQMKVNYNHNI 300
CHIKV nsP2 CTQVYHKSISRRCTLPVTAIVSSLHYEGKMRTTNEYNKPIVVDTTGSTKPDPGDLVLTCF 360
CHIKV nsP2 RGWVKQLQIDYRGYEVMTAAASQGLTRKGVYAVRQKVNENPLYASTSEHVNVLLTRTEGK 420
CHIKV nsP2 LVWKTLSGDPWIKTLQNPPKGNFKATIKEWEVEHASIMAGICSHQ    465
```

A. Amino acid sequence of CHIKV nsP2hel protein. The N-terminal helicase RecA1 and RecA2 domains are marked. G285 and F286 in the RecA1 domain are shown in red.

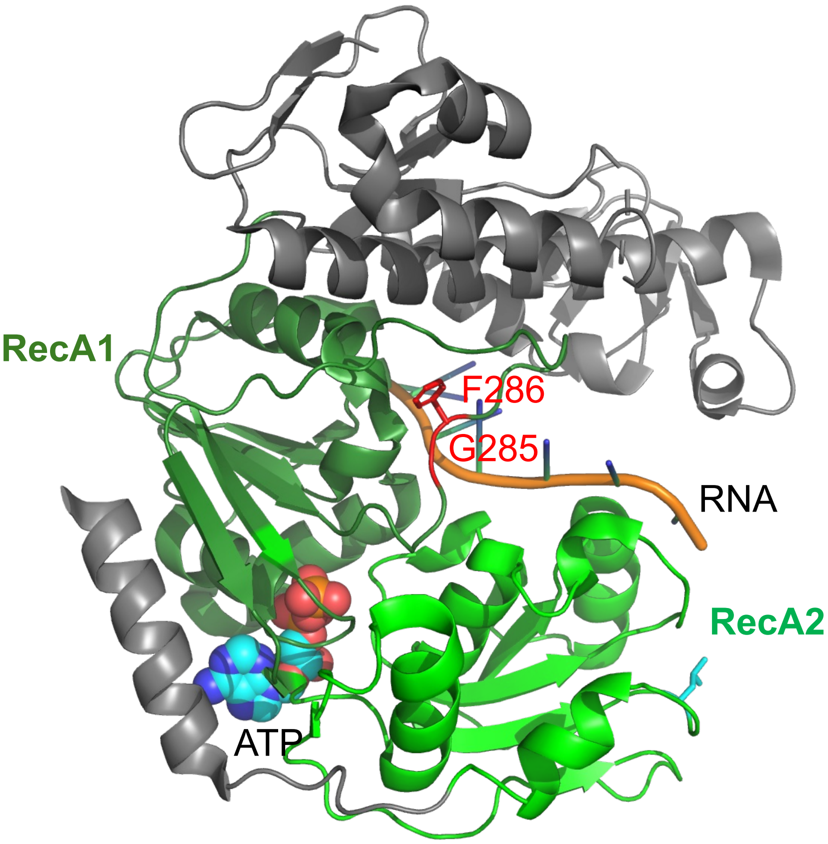


B. X-ray crystal structure of nsP2hel (PDB: 6JIM). RecA1 (light green) and RecA2 (dark green) domains are colored. G285 and F286 in the RecA1 domain are colored red. The ATP analog is shown as a space filling model colored by atom type (C, cyan; N, blue; O, red). ssRNA is shown in gold.

**Figure S8.** Inhibition of CHIKV-nLuc replication by NHC

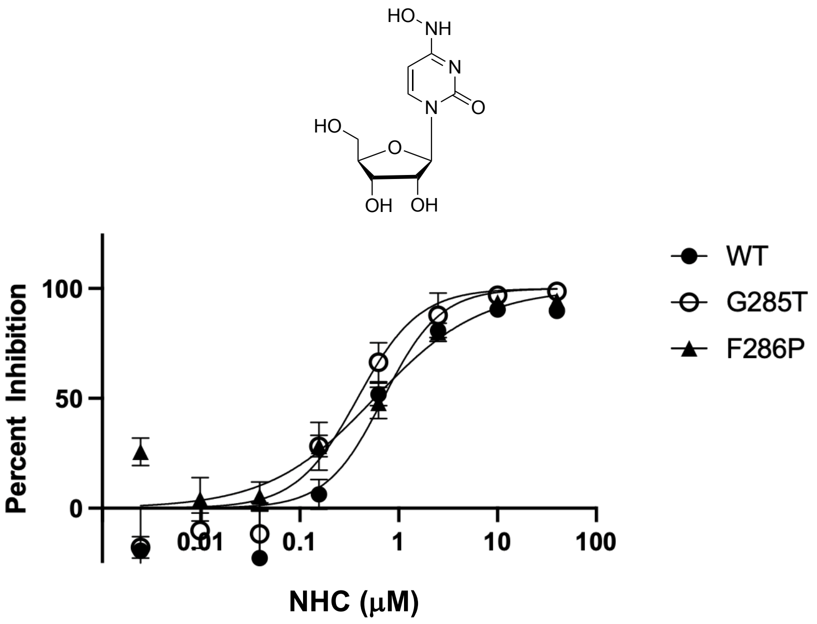

**Figure S9.** Correlation of CHIKV-nLuc with VEEV-nLuc

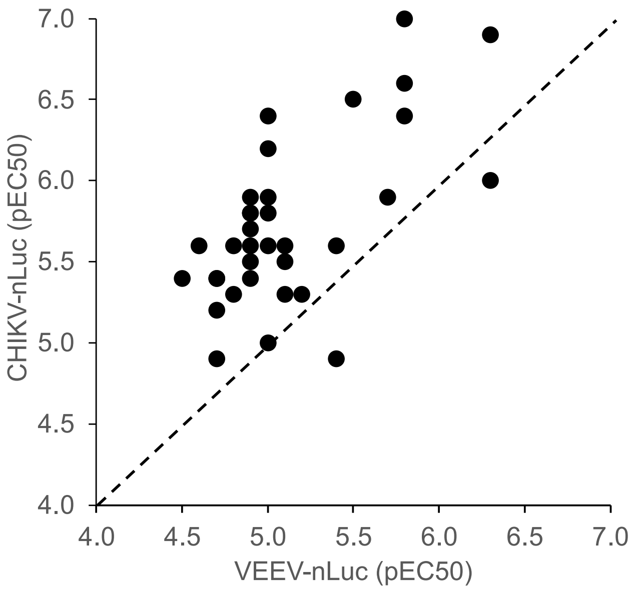


The correlation shows 34 oxaspiropiperidine analogs for which a full dose response curve was determined in both assays. The dashed line represents equivalent antiviral activity against CHIKV and VEEV. Most inhibitors reside on the left of the line indicating antiviral activity on CHIKV > VEEV.

**Figure S9.** Analytical spectra new compounds

$^1$H NMR (700 MHz, $CDCl_3$) for **1a**

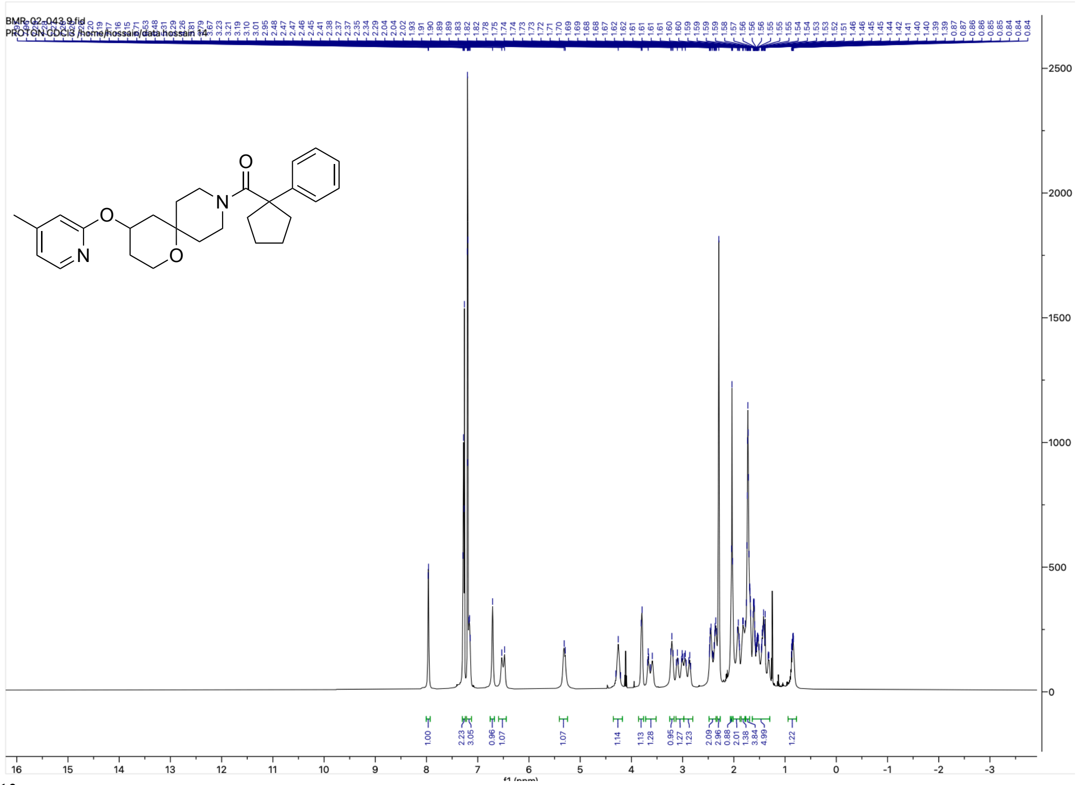


$^{13}$C NMR (176 MHz, $CDCl_3$) for **1a**

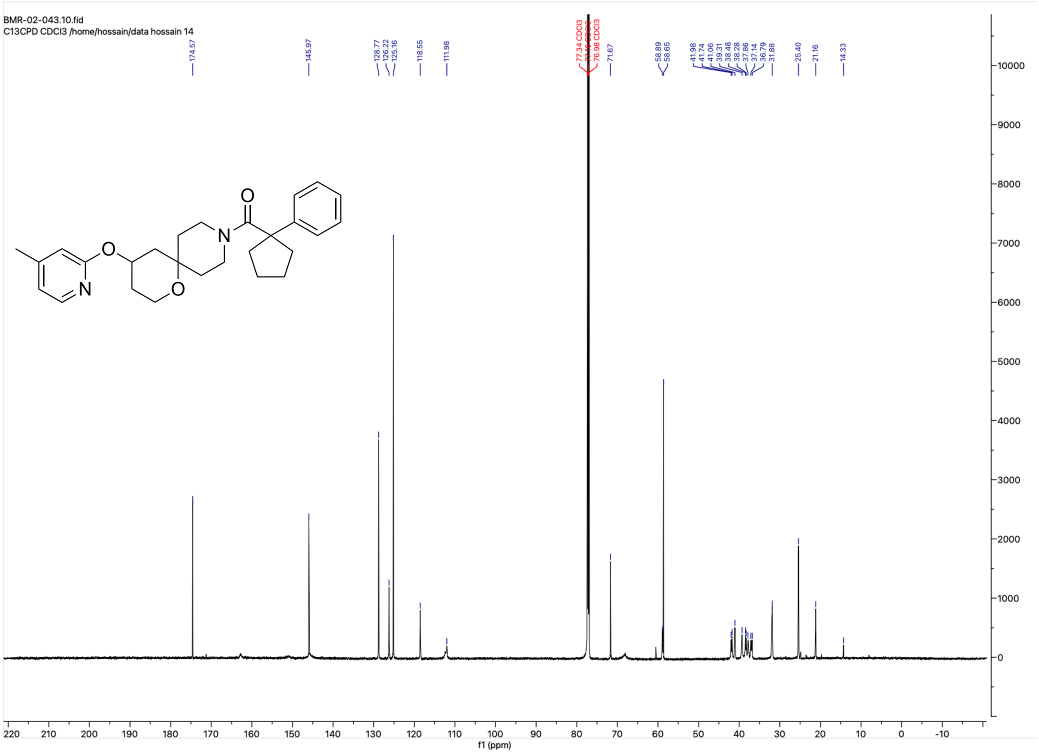

HRMS for **1a**

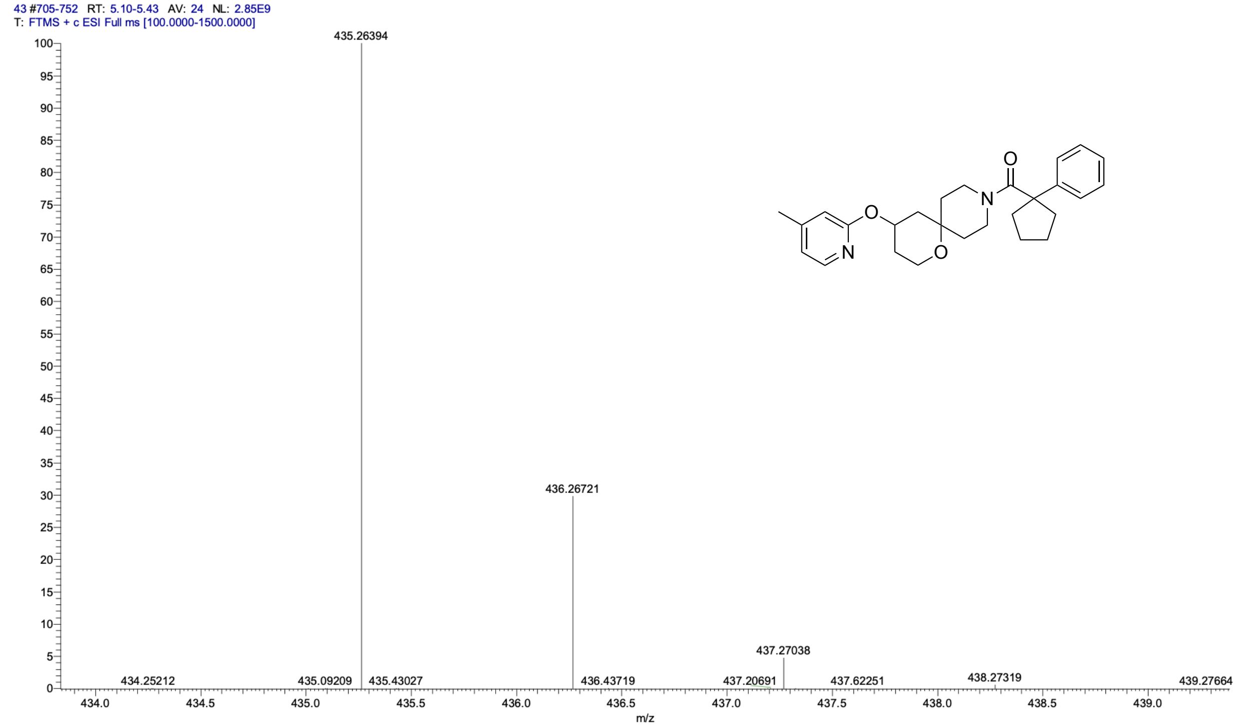


HPLC trace for **1a**

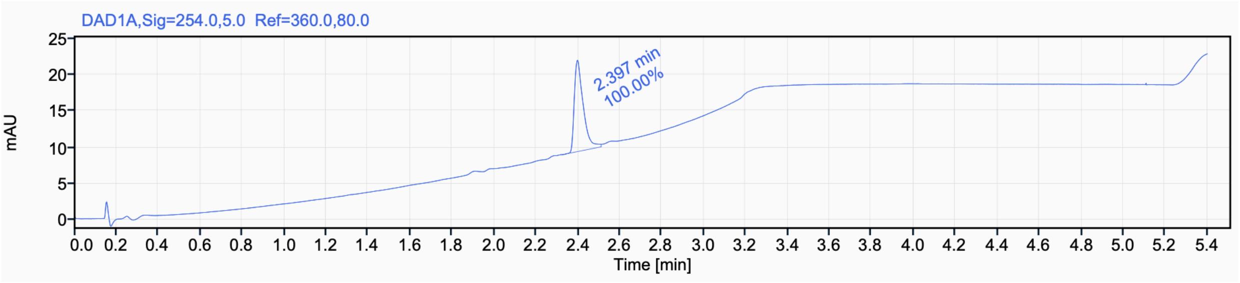


Signal Name DAD1A

| RT (min) | Signal description | Area | Area% | Peak Resolution USP |
|---|---|---|---|---|
| 2.397 | DAD1A,Sig=254.0,5.0 Ref=360.0,80.0 | 37.6 | 100.00 | |

$^{1}$H NMR (500 MHz, DMSO) for **1b**

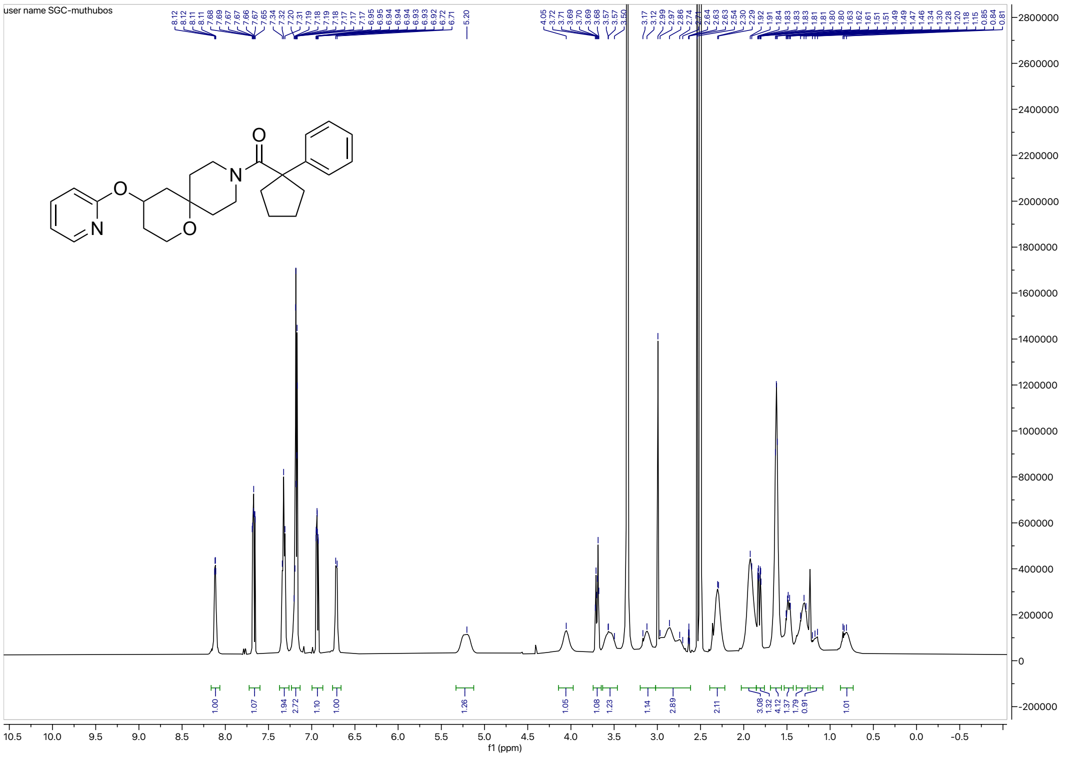

$^{13}$C NMR (126 MHz, DMSO) for **1b**

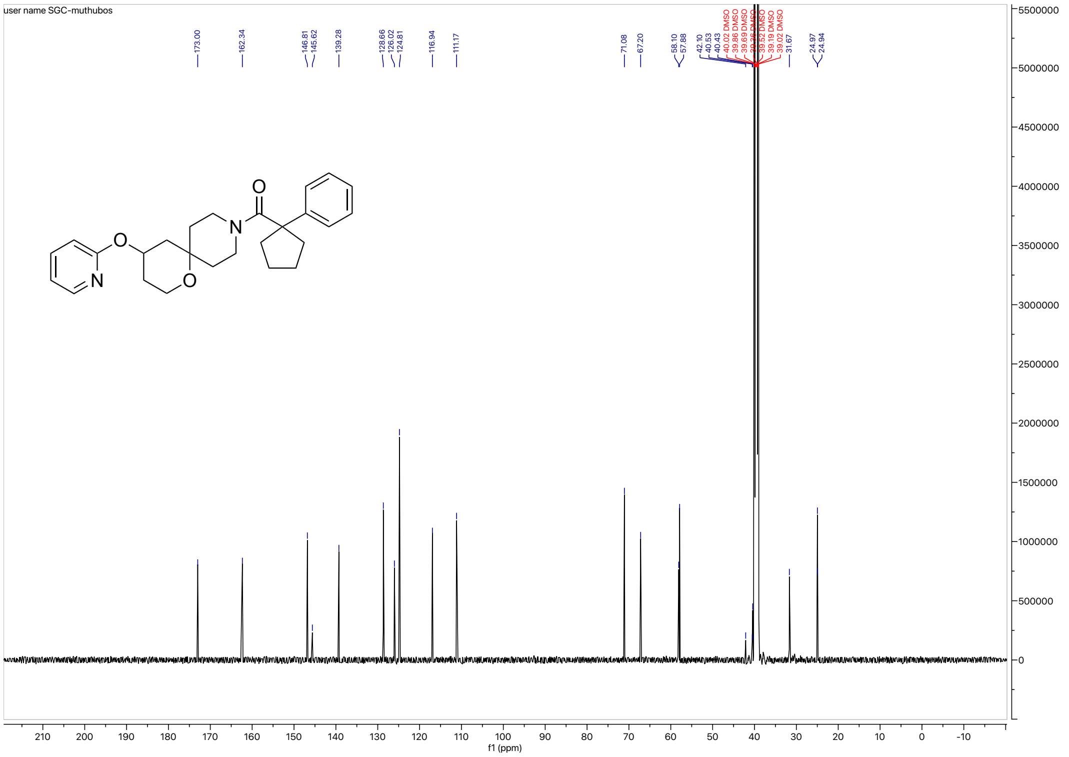

#### HRMS for **1b**

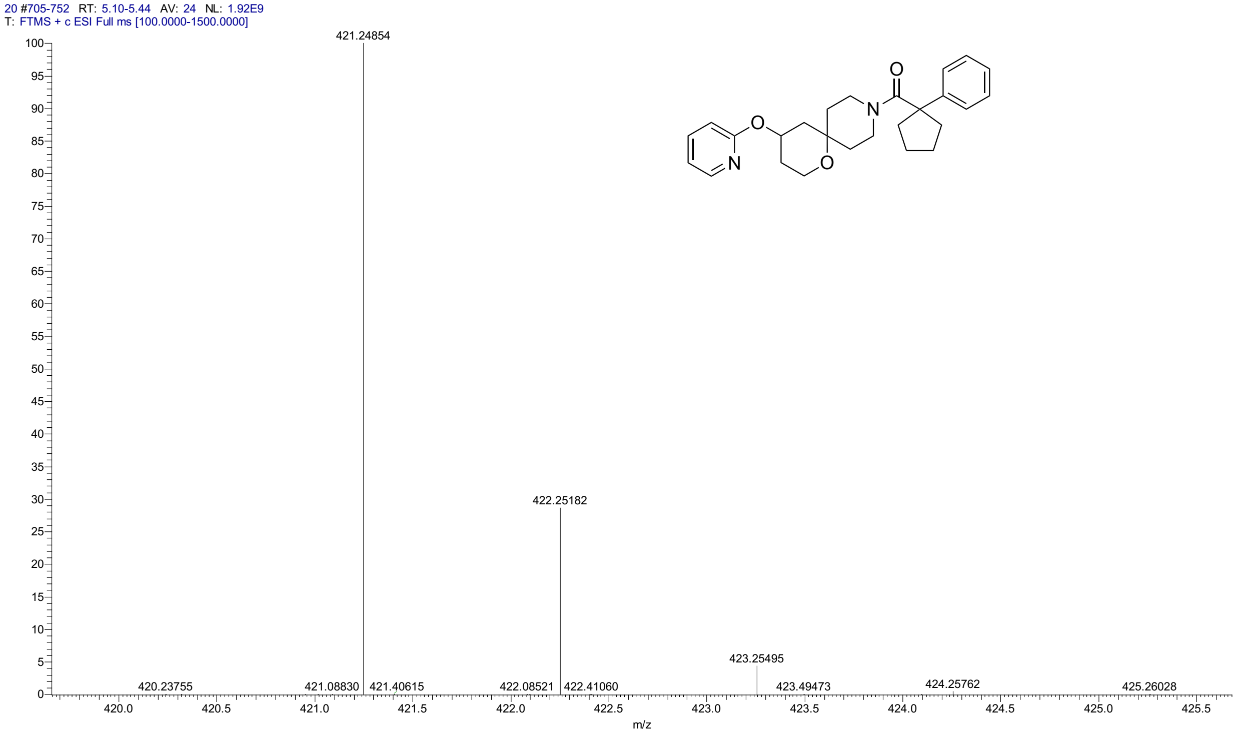


#### HPLC trace for **1b**

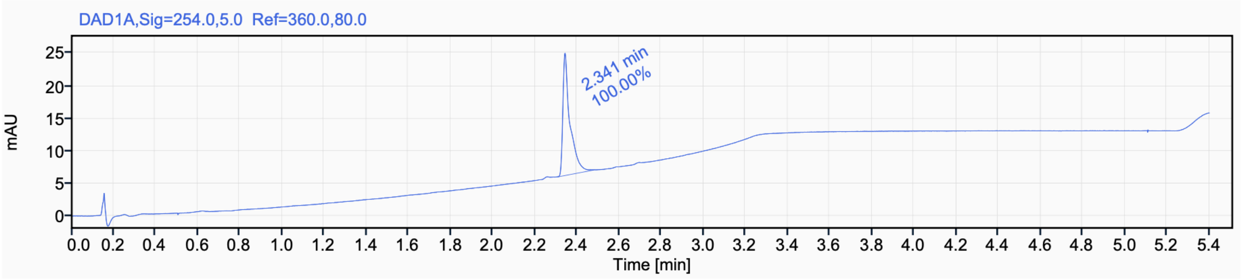


Signal Name DAD1A

| RT (min) | Signal description | Area | Area% | Peak Resolution USP |
|---|---|---|---|---|
| 2.341 | DAD1A,Sig=254.0,5.0 Ref=360.0,80.0 | 44.0 | 100.00 | |

[1]H NMR (400 MHz, $CDCl_3$) for **1c**

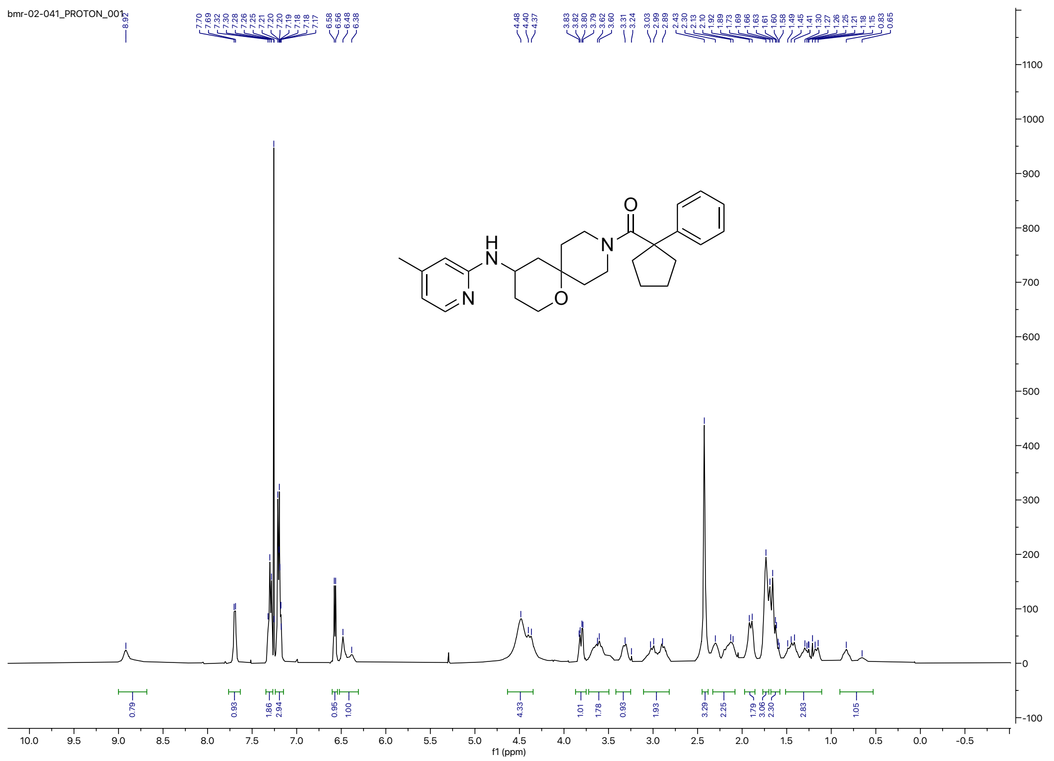


[13]C NMR (100 MHz, $CDCl_3$ for **1c**

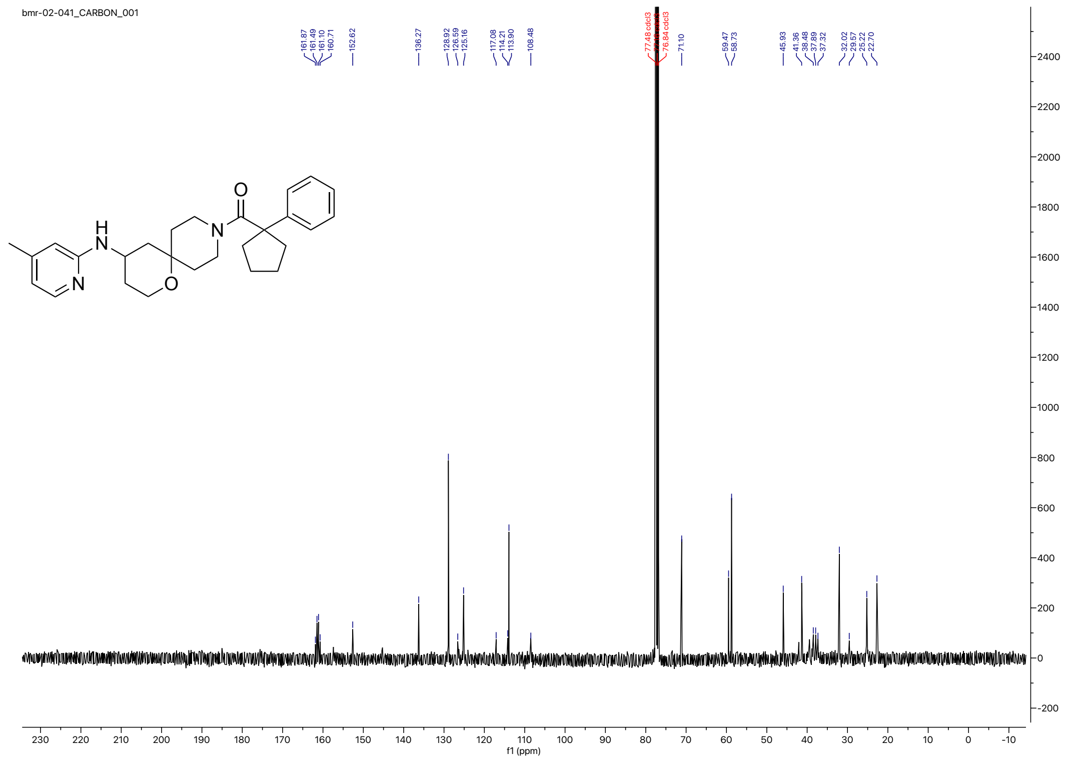

#### HRMS for **1c**

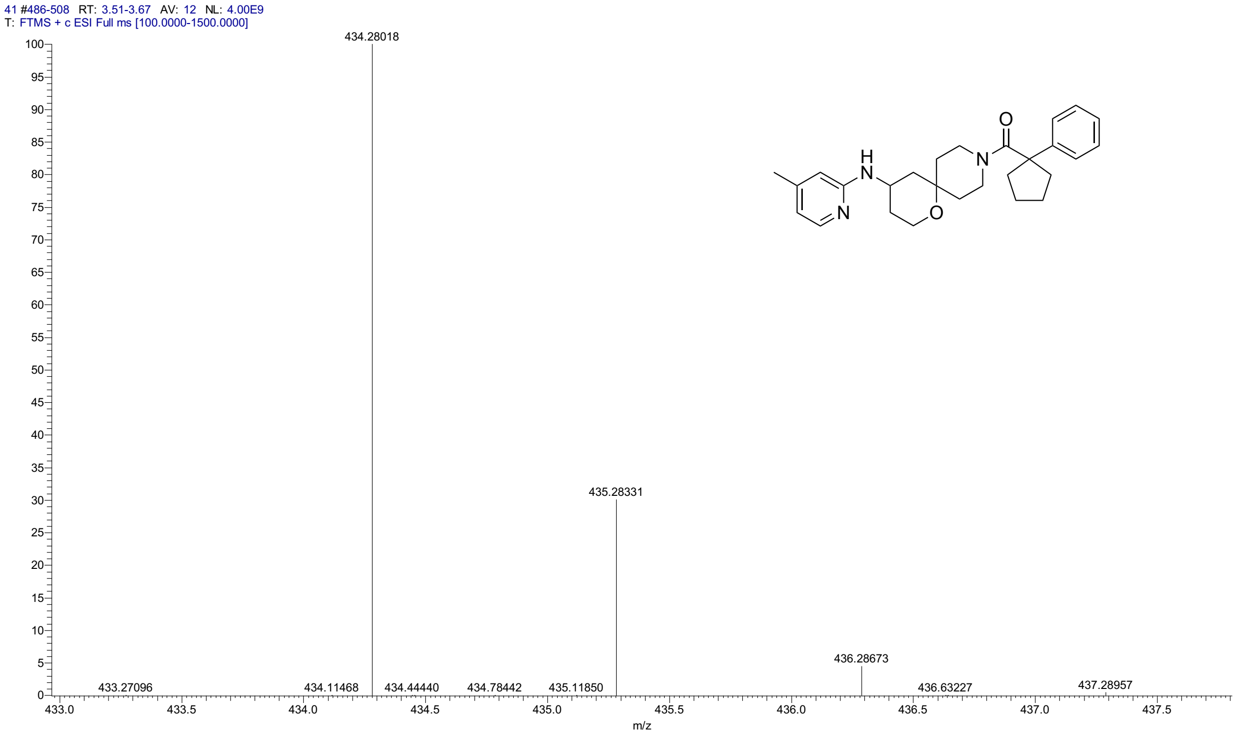


#### HPLC for **1c**

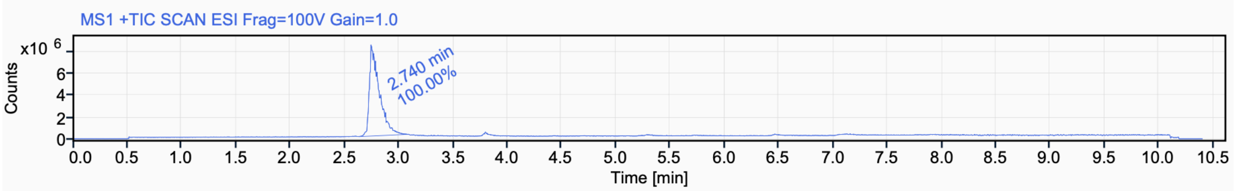


Signal Name MS1 +TIC SCAN ESI Frag=100V Gain=1.0

| RT (min) | Signal description | m/z | Purity UV(‰) / MS(%) | Area | Area% |
|---|---|---|---|---|---|
| 2.740 | MS1 +TIC SCAN ESI Frag=100V Gain=1.0 | | | 53985755.4 | 100.00 |

[1]H NMR (500 MHz, $CD_3CN$) for **1d**

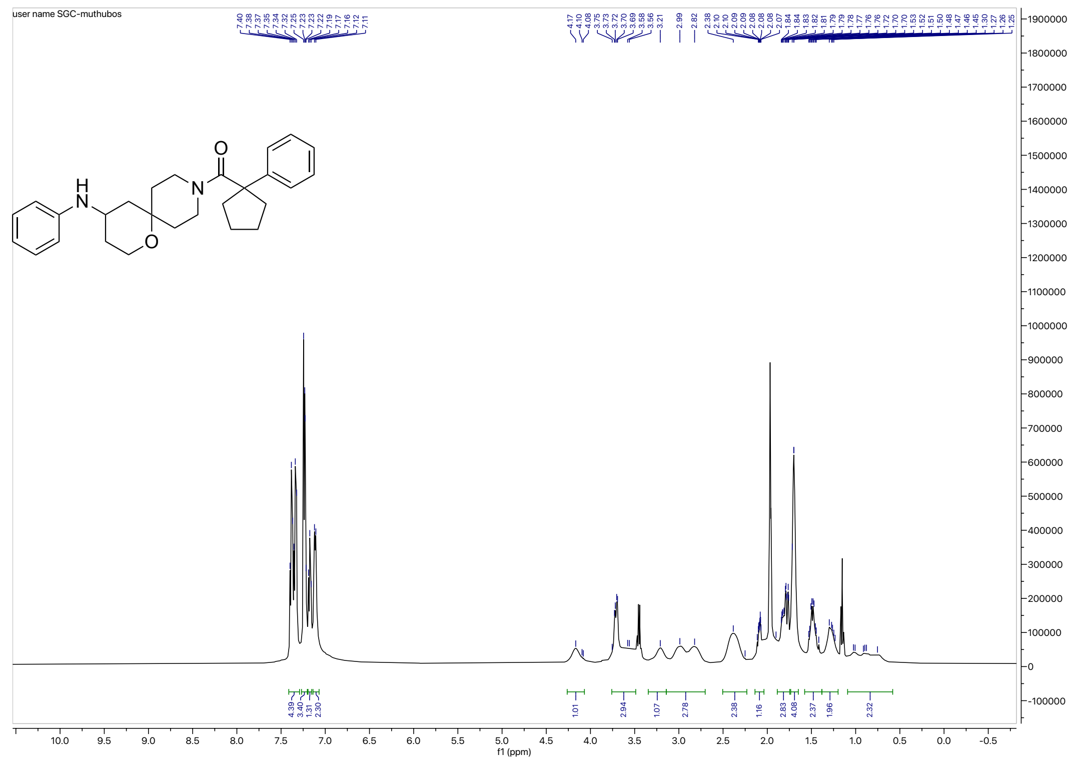

[13]C NMR (126 MHz, $CD_3CN$) for **1d**

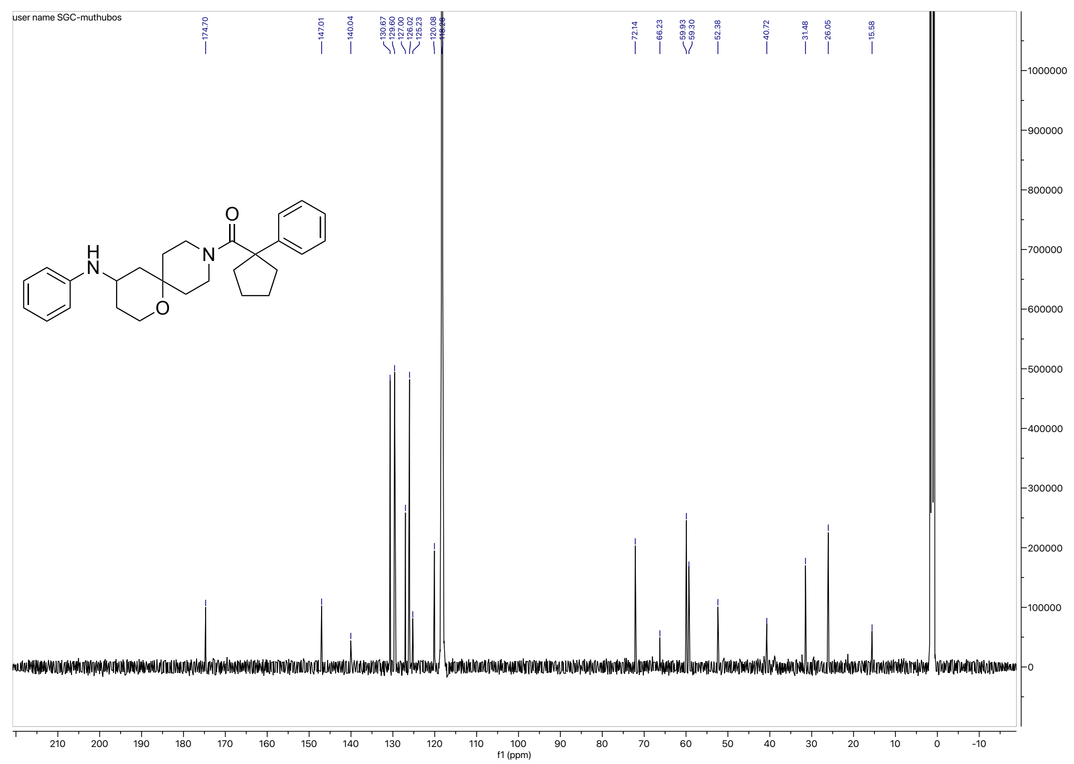

HRMS for **1d**

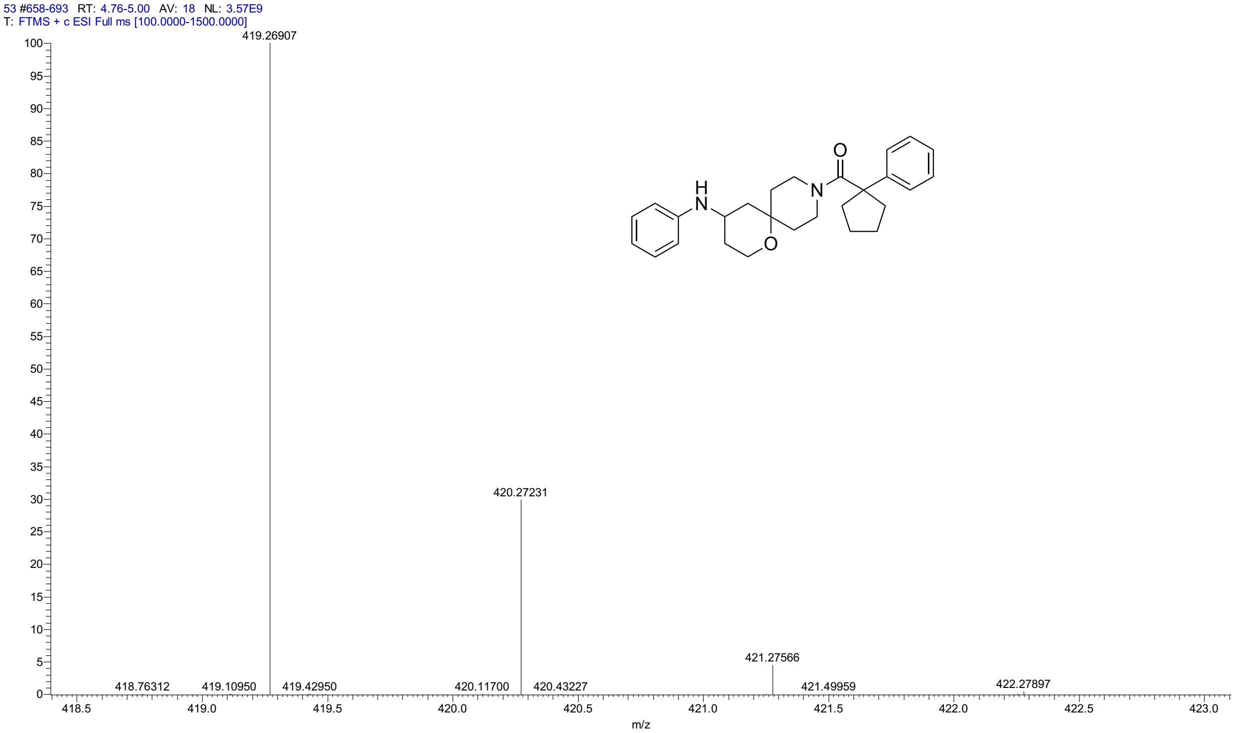


HPLC for **1d**

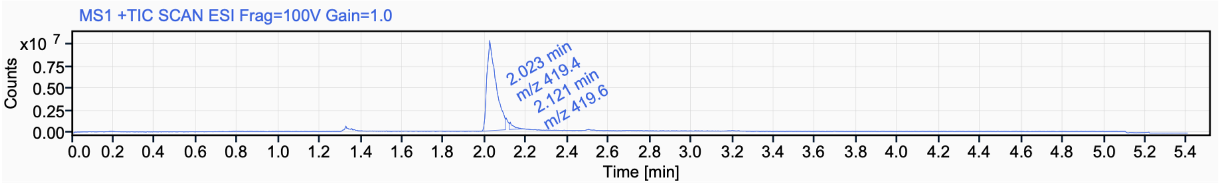


Signal Name MS1 +TIC SCAN ESI Frag=100V Gain=1.0

| RT (min) | Signal description | m/z | Purity UV(‰) / MS(%) | Area | Area% |
|---|---|---|---|---|---|
| 2.023 | MS1 +TIC SCAN ESI Frag=100V Gain=1.0 | 419.4 | | 30999637.1 | 96.68 |
| 2.121 | MS1 +TIC SCAN ESI Frag=100V Gain=1.0 | 419.6 | | 1064444.7 | 3.32 |

$^{1}$H NMR (500 MHz, $CD_3CN$) for **2a**

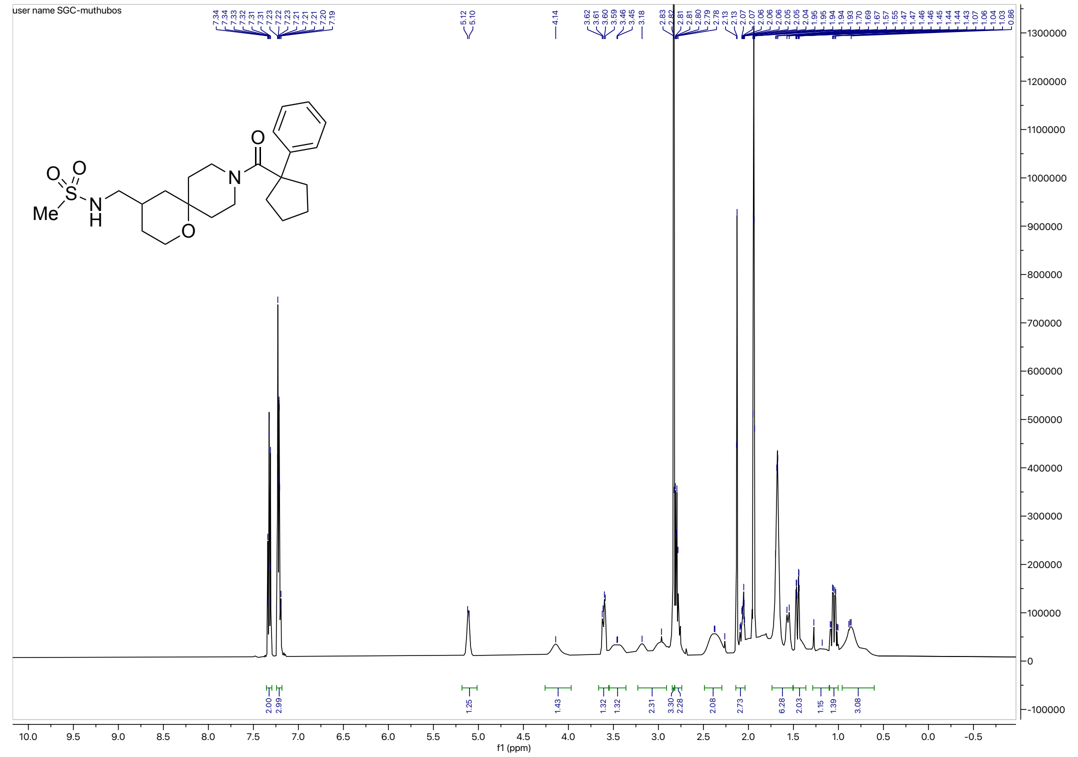

$^{13}$C NMR (126 MHz, $CD_3CN$) for **2a**

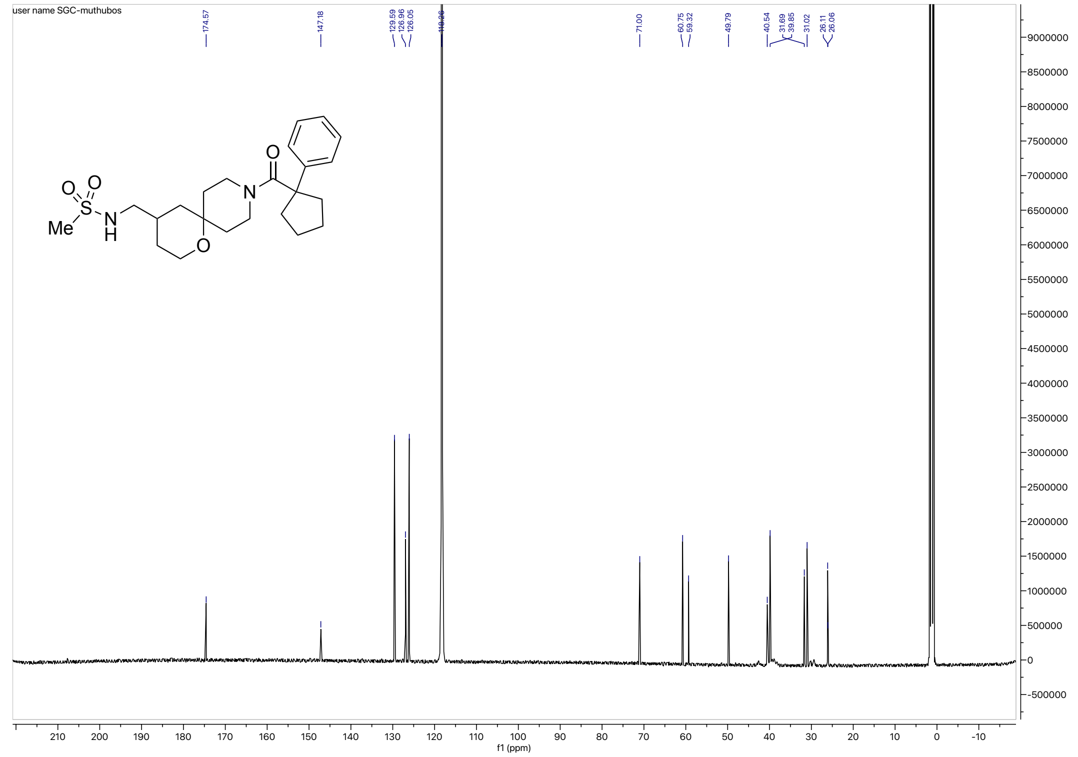

HRMS for **2a**

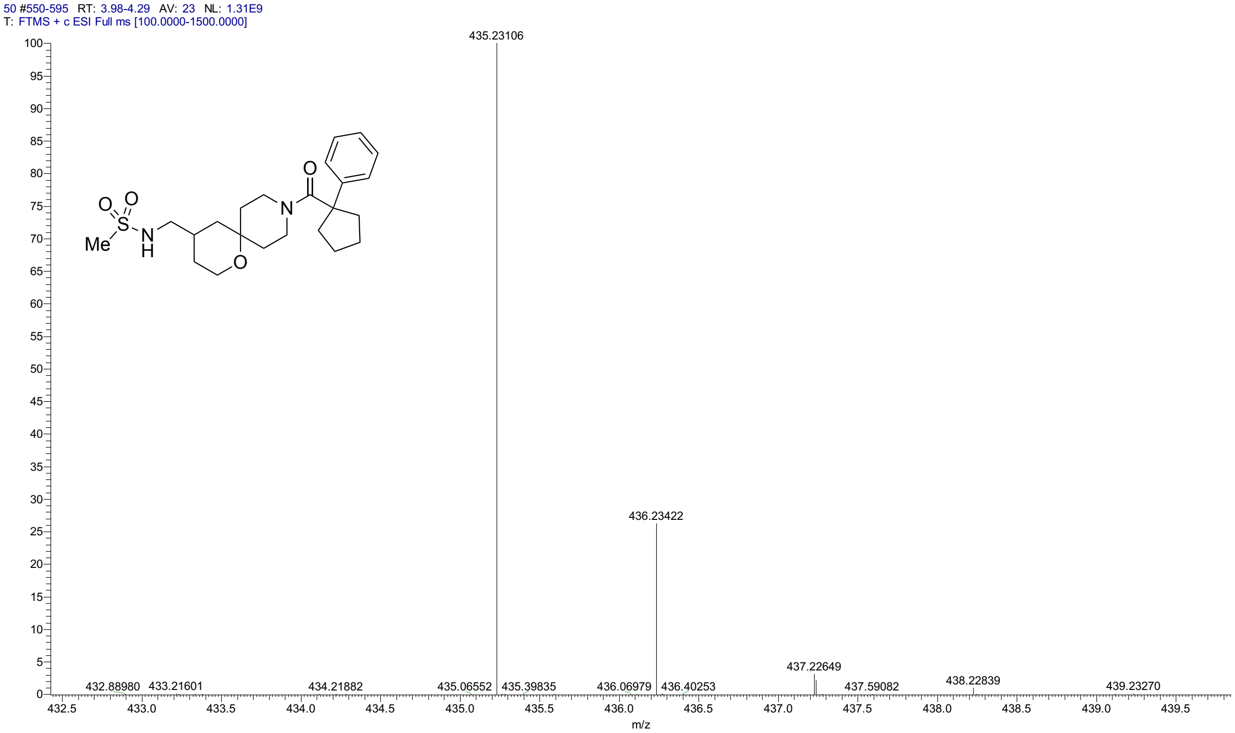


HPLC for **2a**

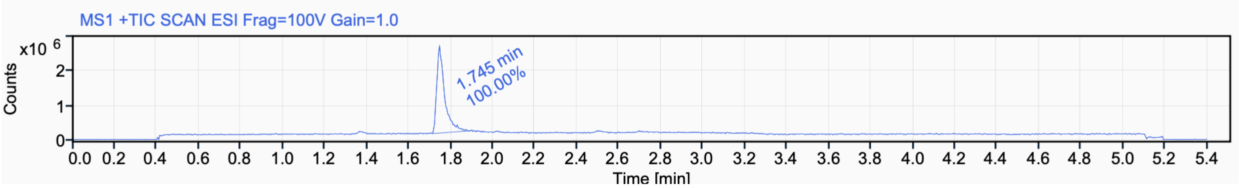


Signal Name MS1 +TIC SCAN ESI Frag=100V Gain=1.0

| RT (min) | Signal description | Area | Area% | Peak Resolution USP |
|---|---|---|---|---|
| 1.745 | MS1 +TIC SCAN ESI Frag=100V Gain=1.0 | 6645361.7 | 100.00 | |

#### $^{1}$H NMR (400 MHz, $CD_3CN$) for **2b**

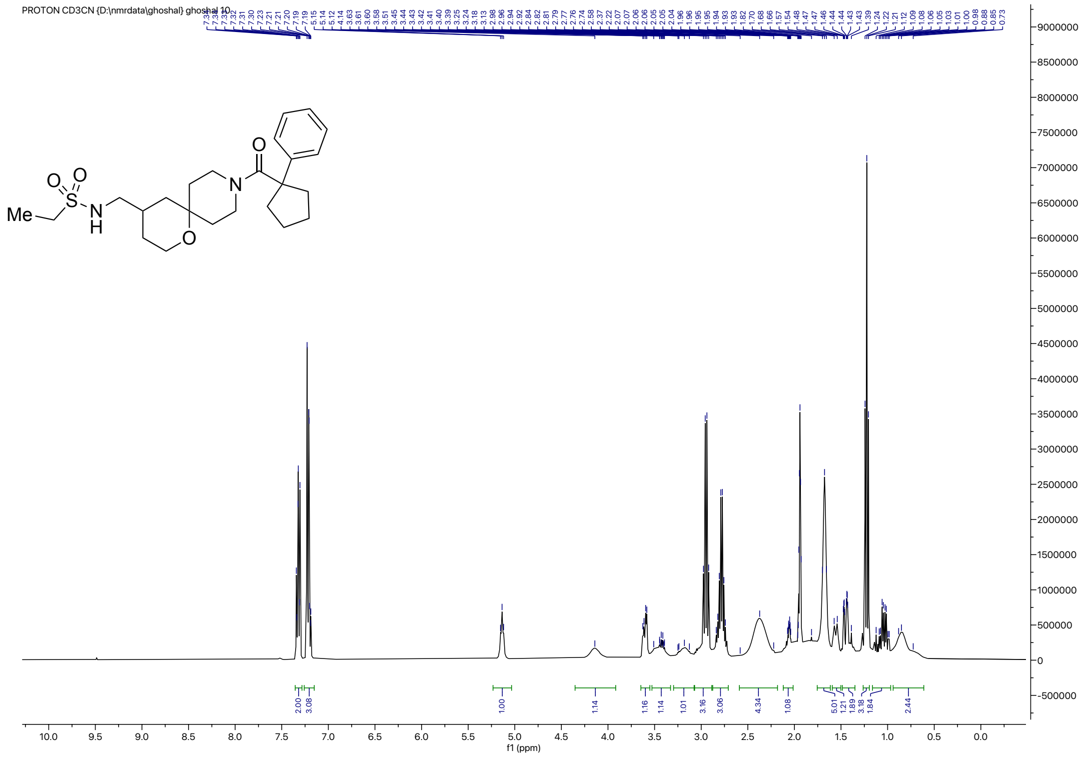


#### $^{13}$C NMR (126 MHz, $CD_3CN$) for **2b**

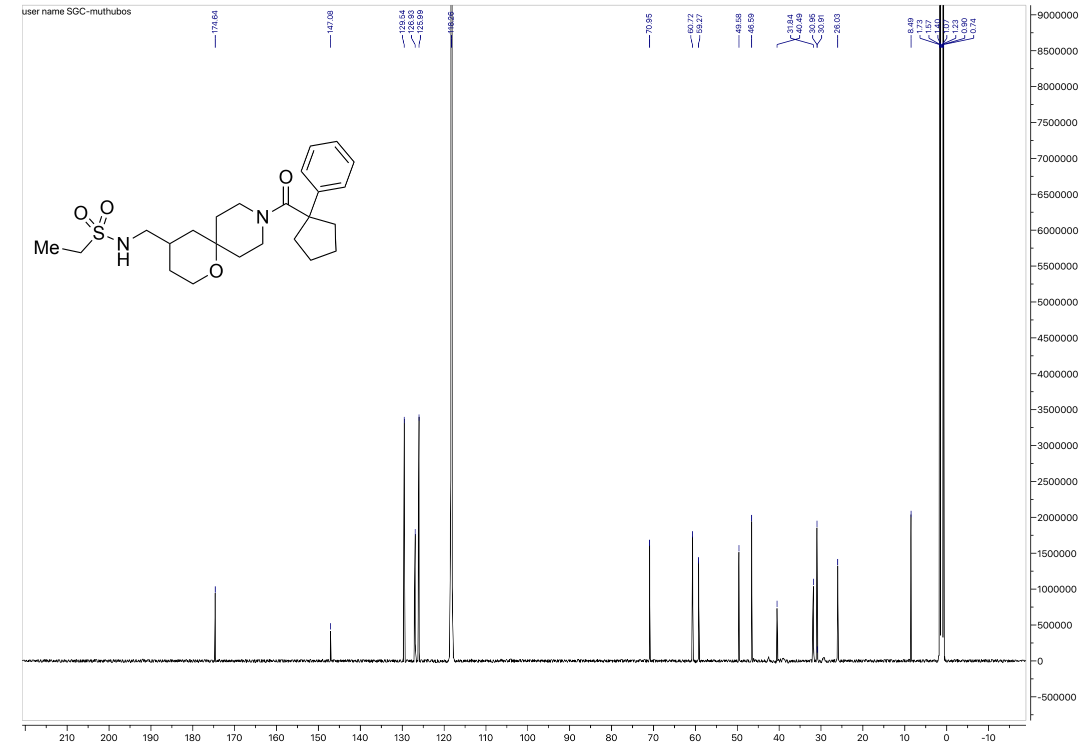

#### HRMS for **2b**

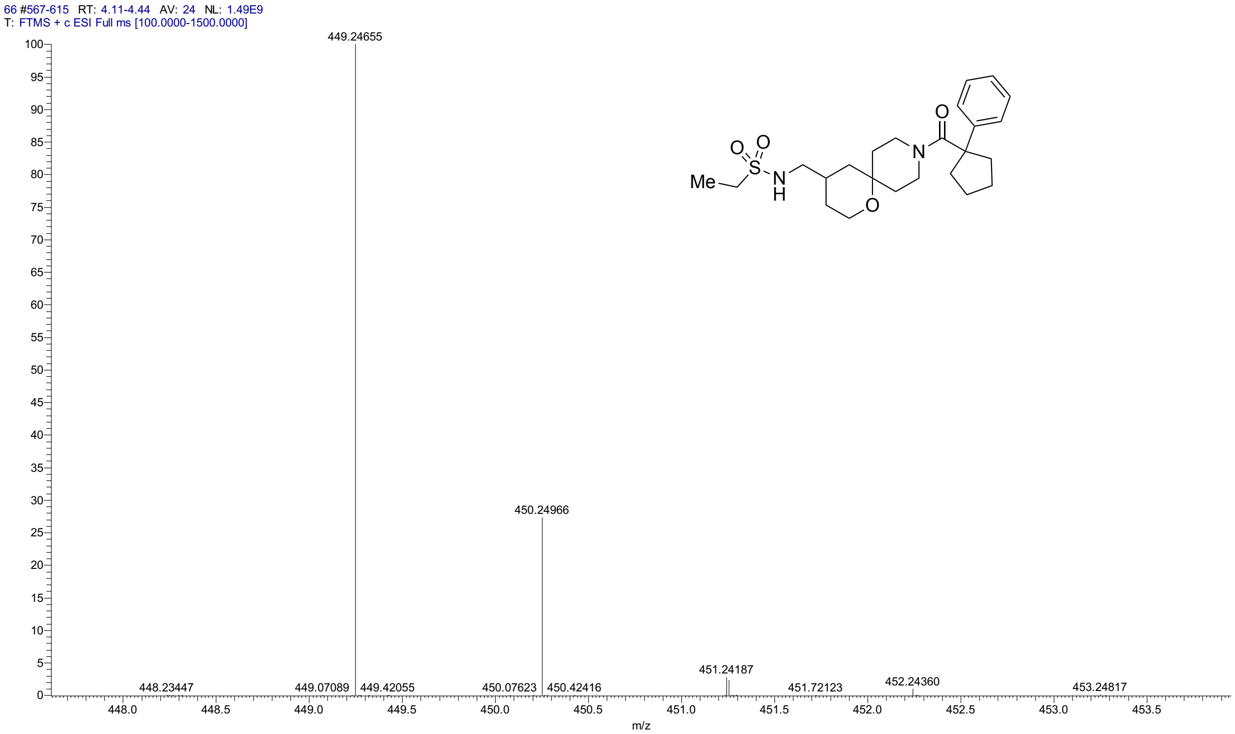


#### HPLC for **2b**

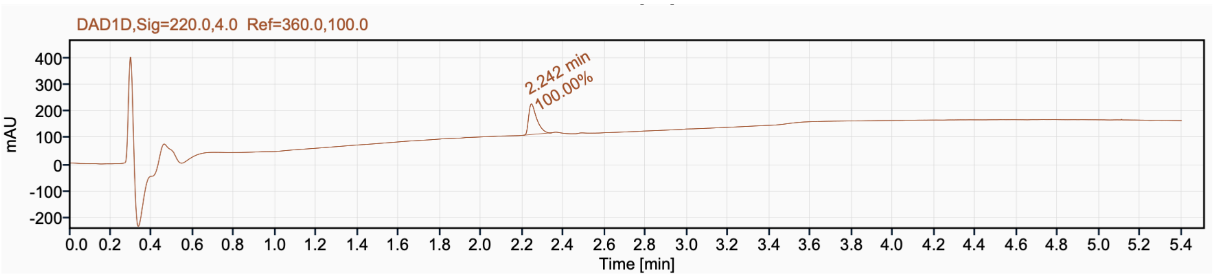


Signal Name DAD1D

| RT (min) | Signal description | Area | Area% | Peak Resolution USP |
|---|---|---|---|---|
| 2.242 | DAD1D,Sig=220.0,4.0 Ref=360.0,100.0 | 309.0 | 100.00 | |

[1]H NMR (500 MHz, $CD_3CN$) for **2c**

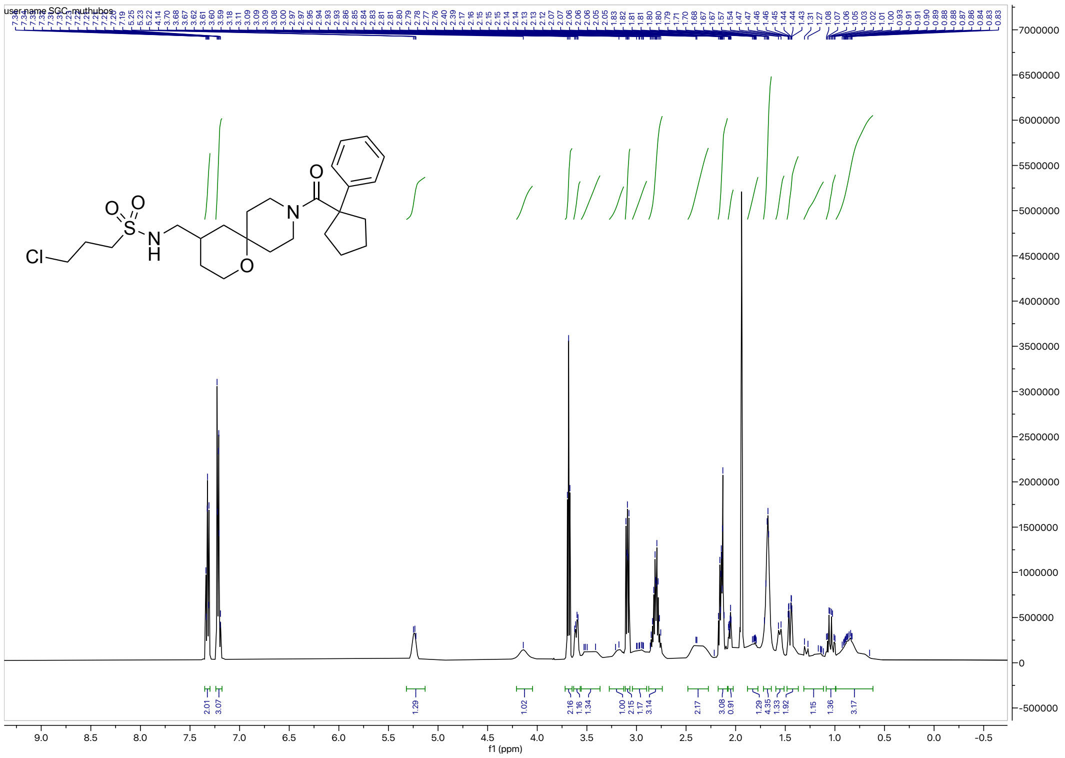

[13]C NMR (126 MHz, $CD_3CN$) for **2c**

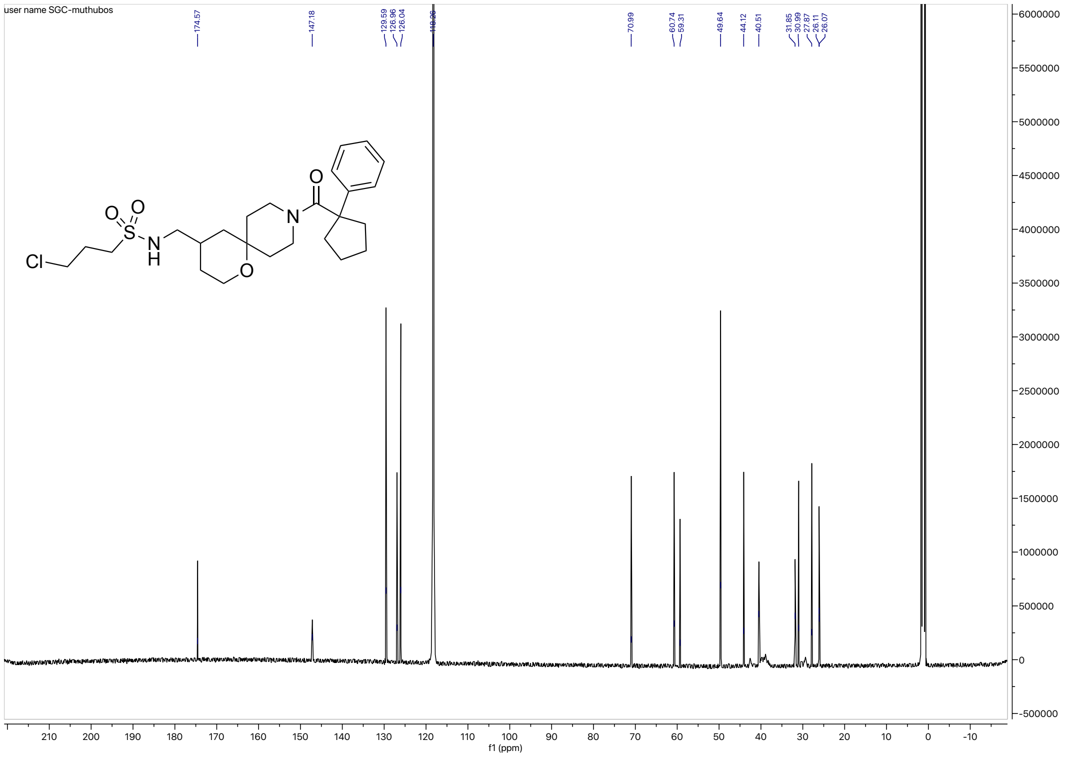

HRMS for **2C**

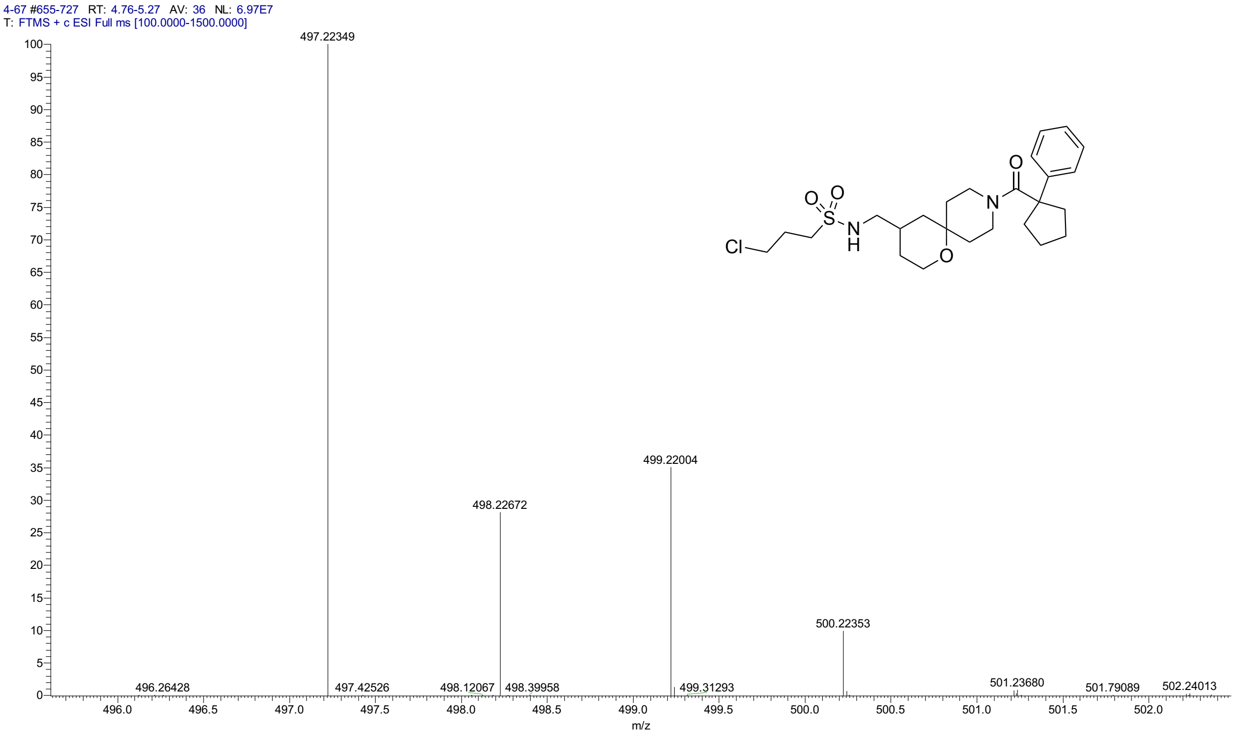


HPLC for **2C**

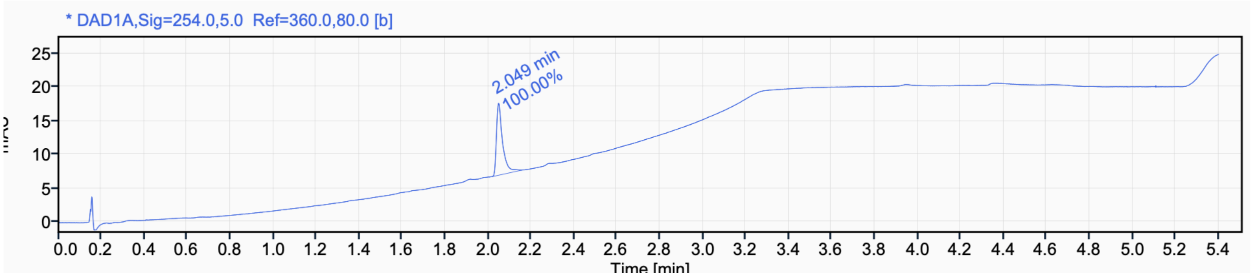


Signal Name DAD1A

| RT (min) | Signal description | m/z | Purity UV(‰) / MS(%) | Area | Area% |
|---|---|---|---|---|---|
| 2.049 | * DAD1A,Sig=254.0,5.0 Ref=360.0,80.0 [b] | | | 21.1 | 100.00 |

$^{1}H$ NMR (500 MHz, $CD_3CN$) for **2d**

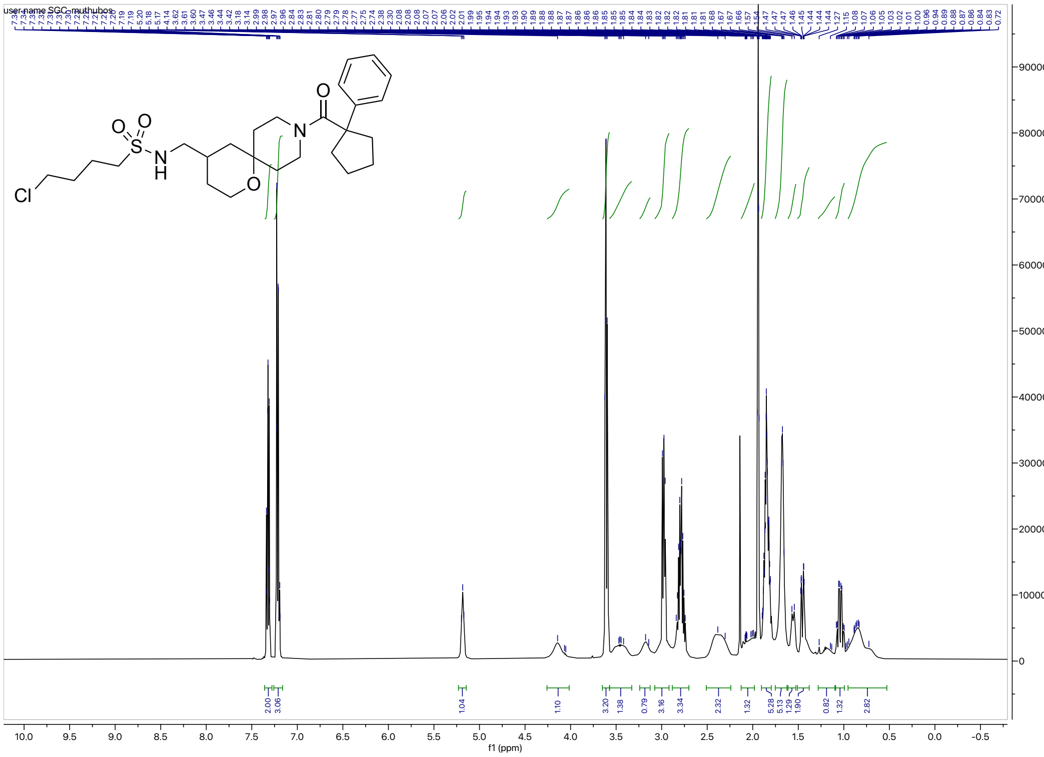


$^{13}C$ NMR (126 MHz, $CD_3CN$) for **2d**

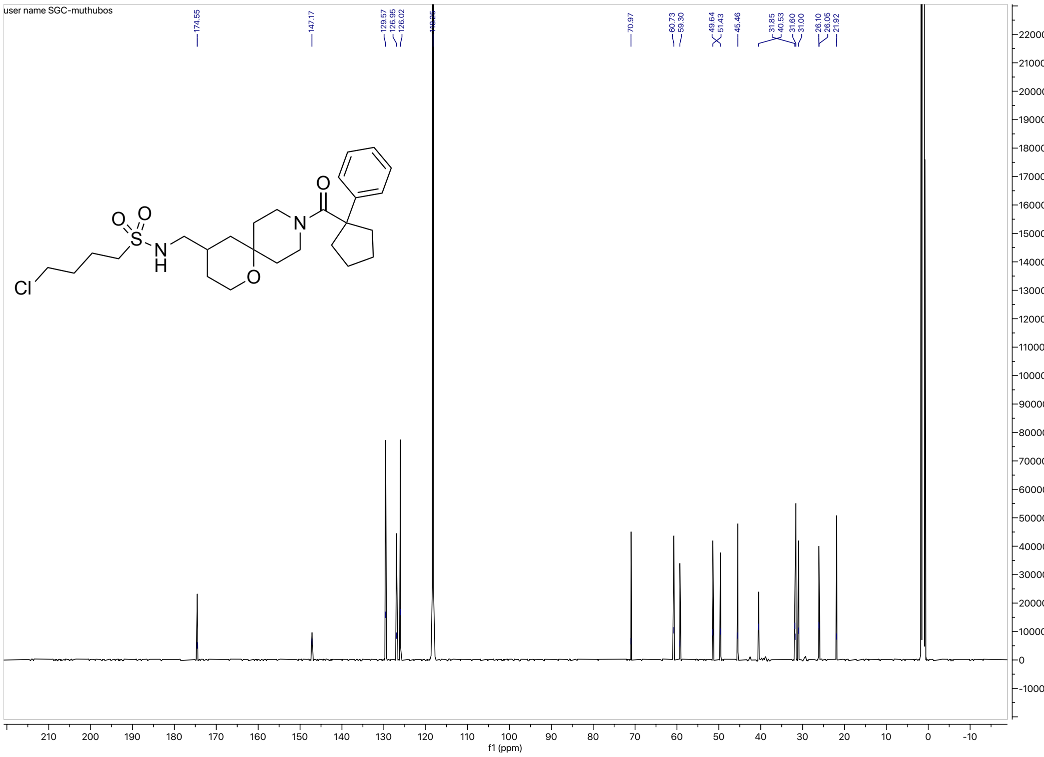

#### HRMS for **2d**

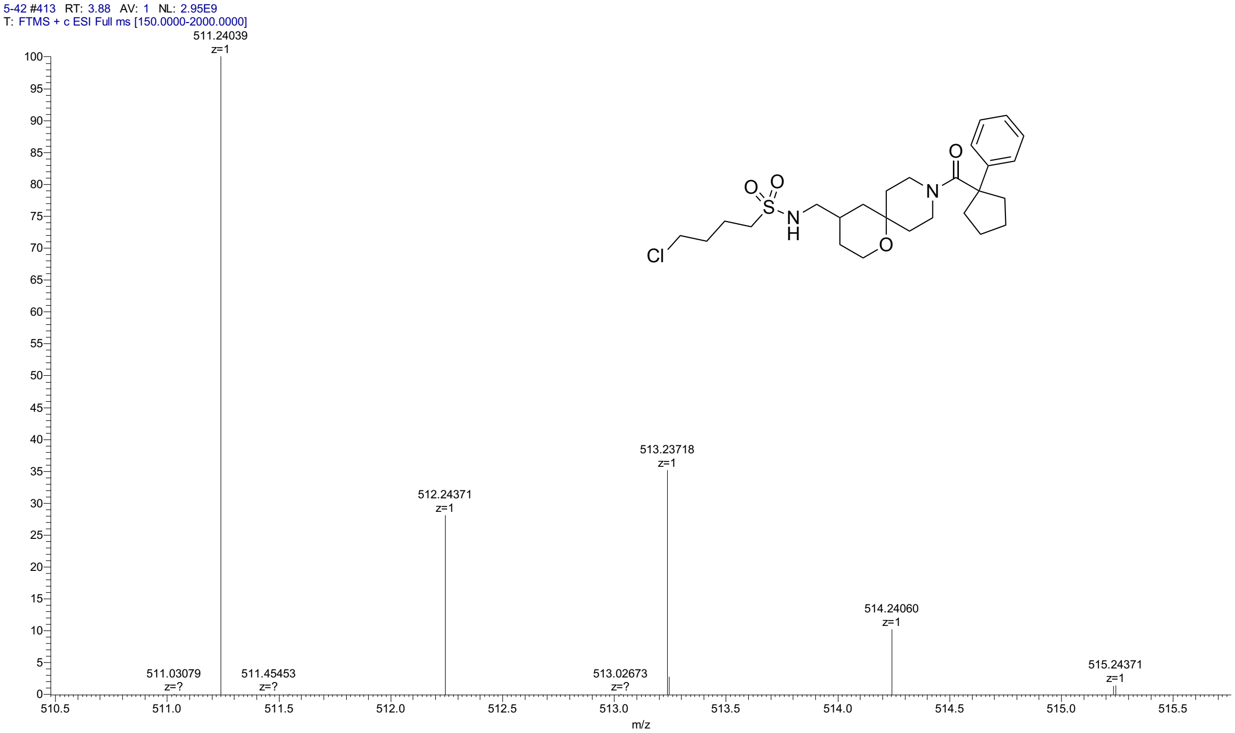


#### HPLC for **2d**

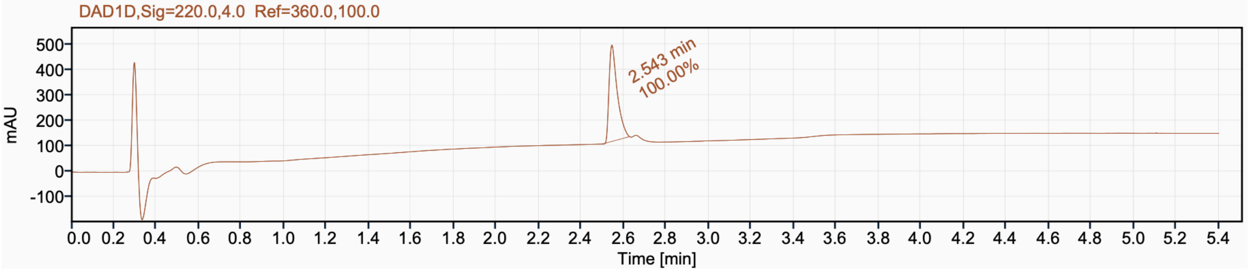


Signal Name DAD1D

| RT (min) | Signal description | Area | Area% | Peak Resolution USP |
|---|---|---|---|---|
| 2.543 | DAD1D,Sig=220.0,4.0 Ref=360.0,100.0 | 1011.0 | 100.00 | |

$^{1}H$ NMR (400 MHz, $CDCl_3$) for **2e**

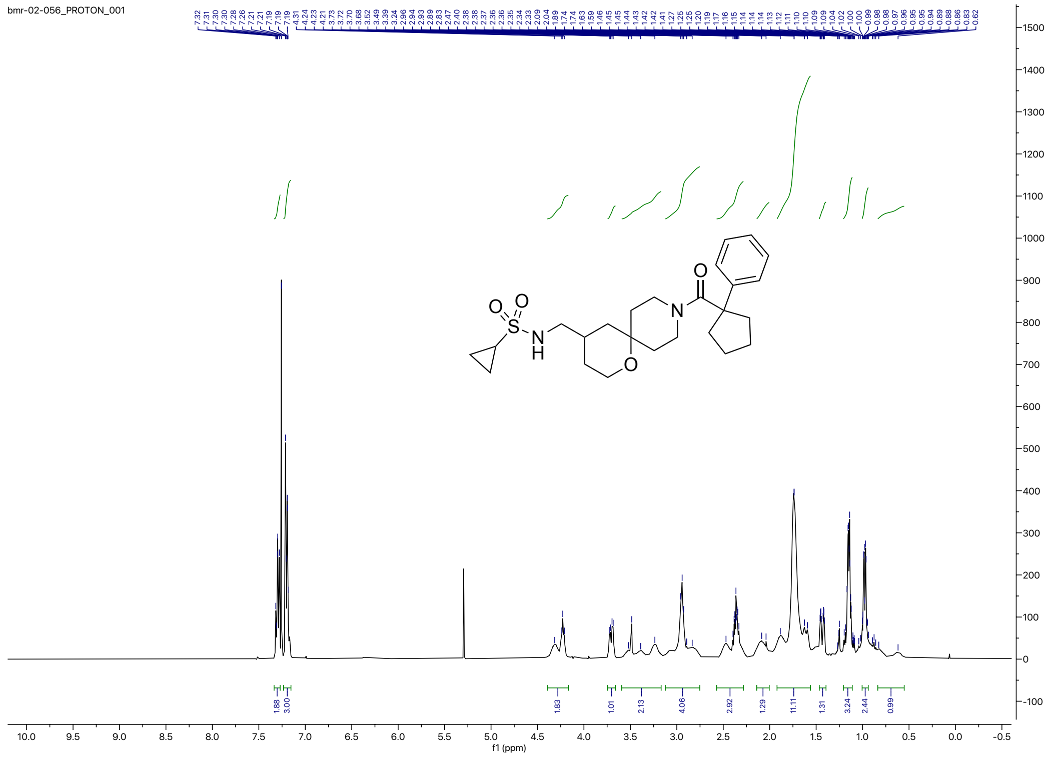

$^{13}C$ NMR (100 MHz, $CDCl_3$) for **2e**

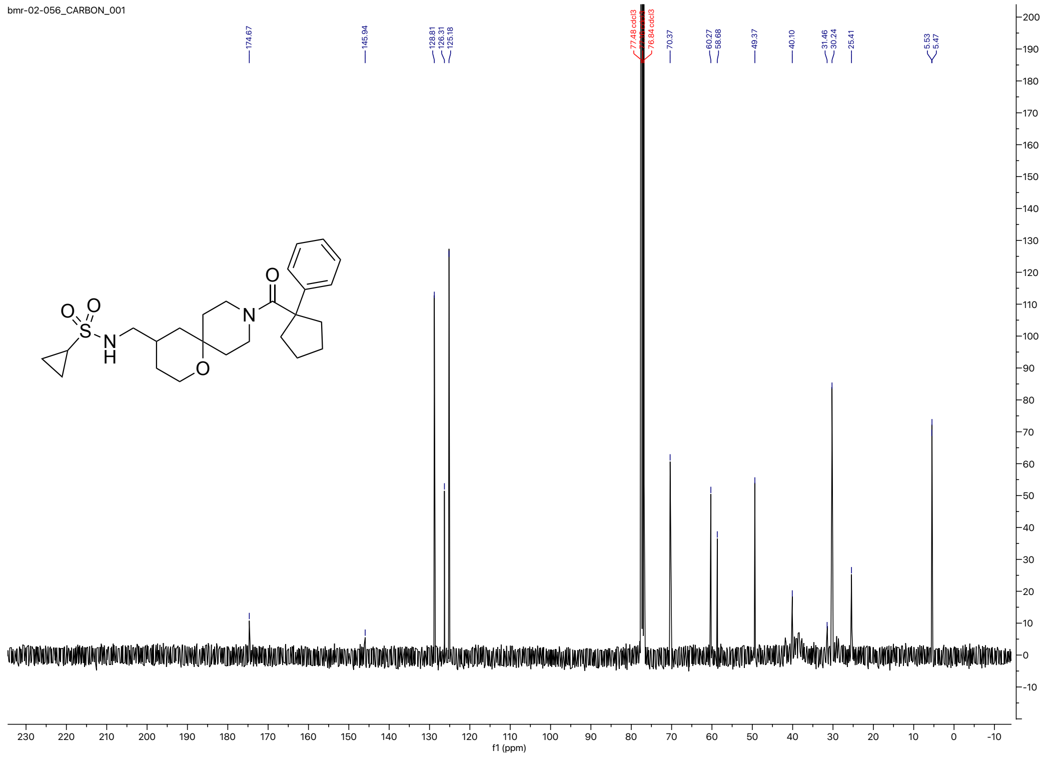

HRMS for **2e**

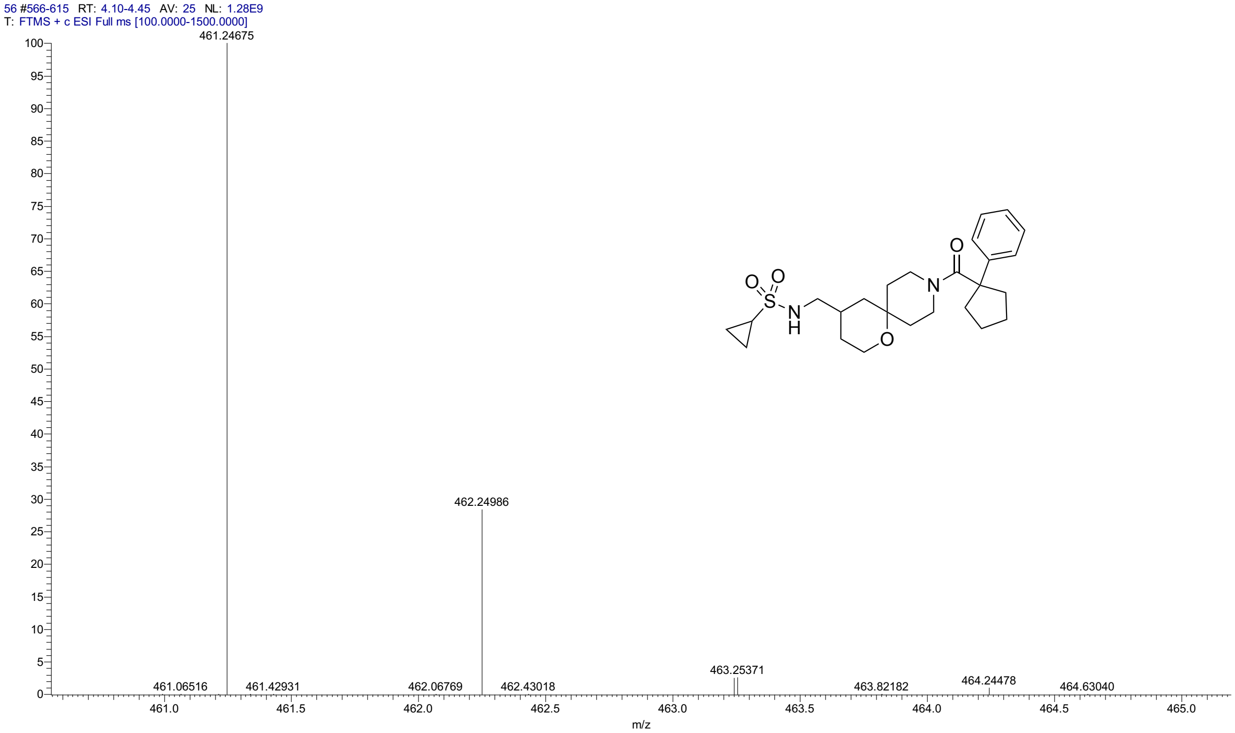


HPLC for **2e**

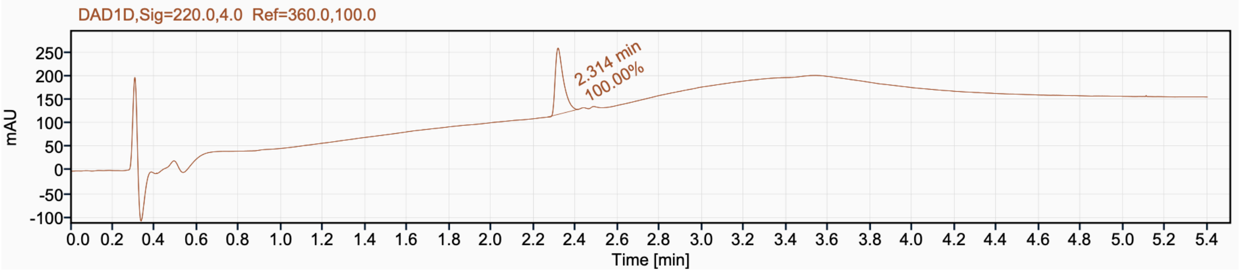


Signal Name DAD1D

| RT (min) | Signal description | Area | Area% | Peak Resolution USP |
|---|---|---|---|---|
| 2.314 | DAD1D,Sig=220.0,4.0 Ref=360.0,100.0 | 381.2 | 100.00 | |

[1]H NMR (400 MHz, $CD_3CN$) for **2f**

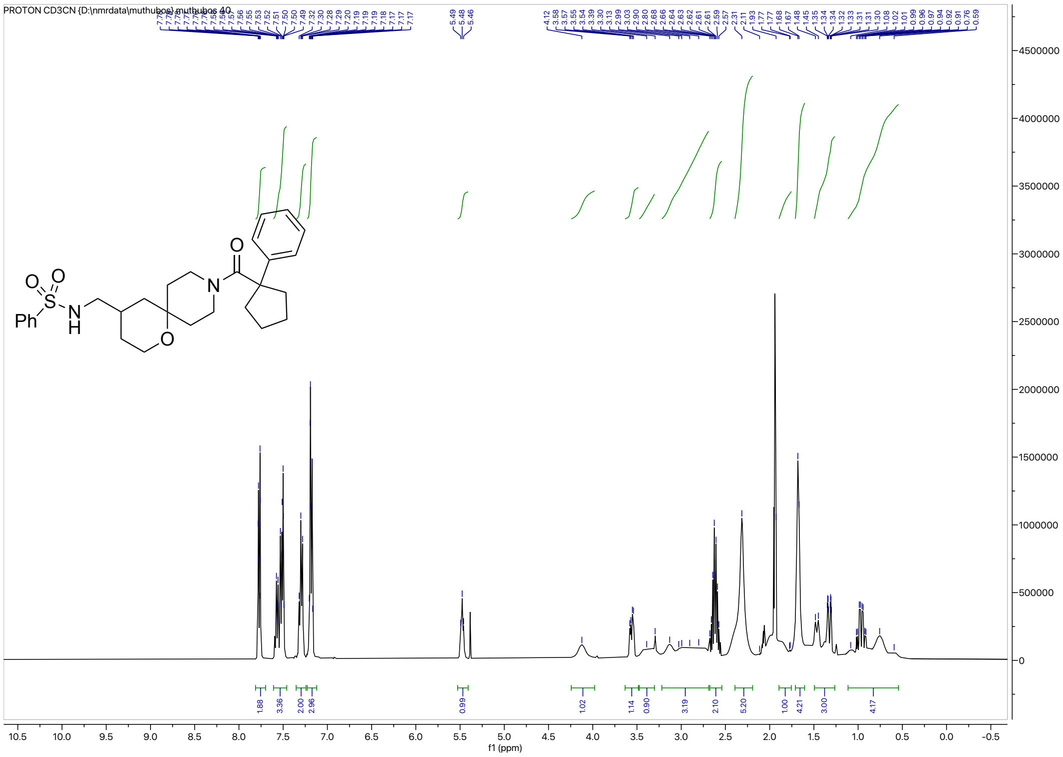

[13]C NMR (126 MHz, $CDCl_3$) for **2f**

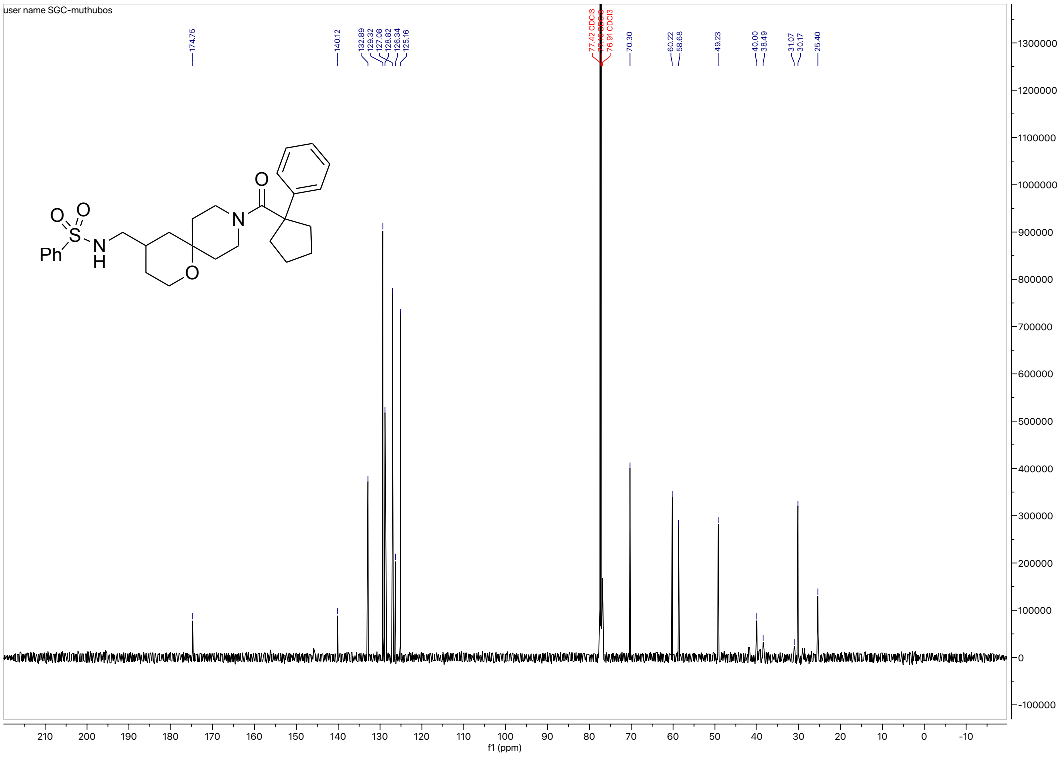

HRMS for **2f**

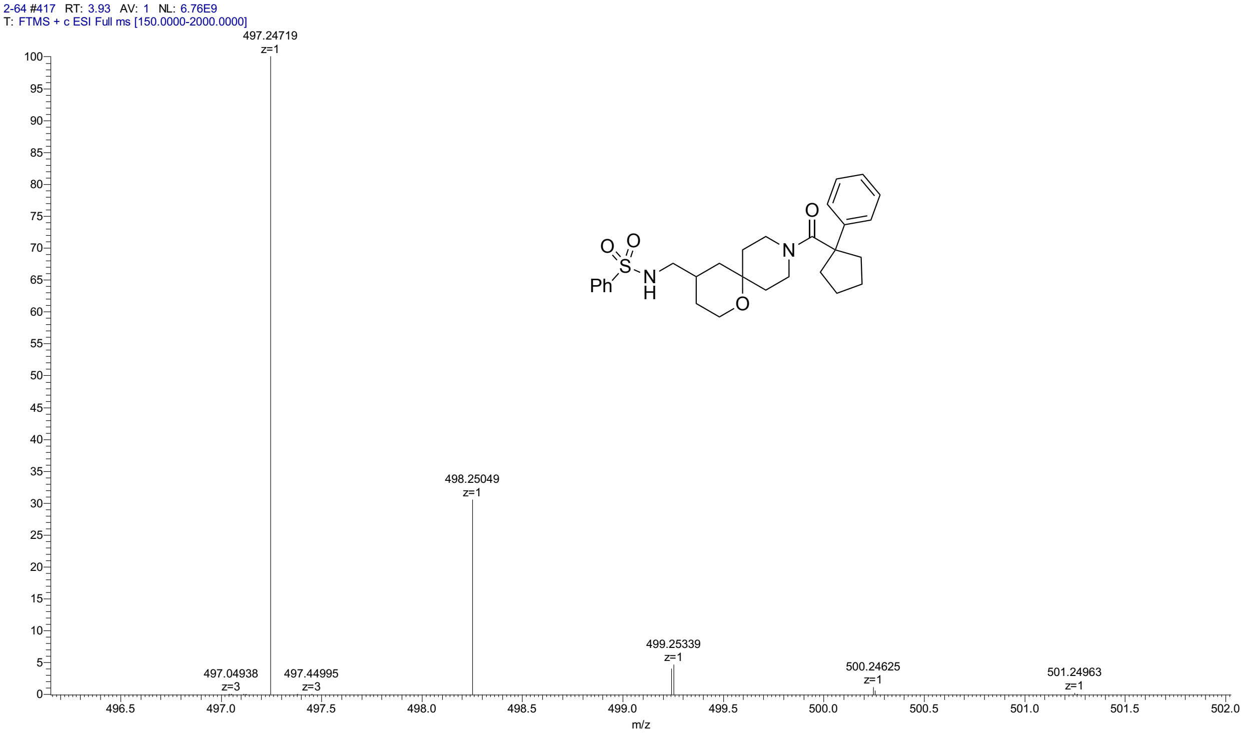


HPLC for **2f**

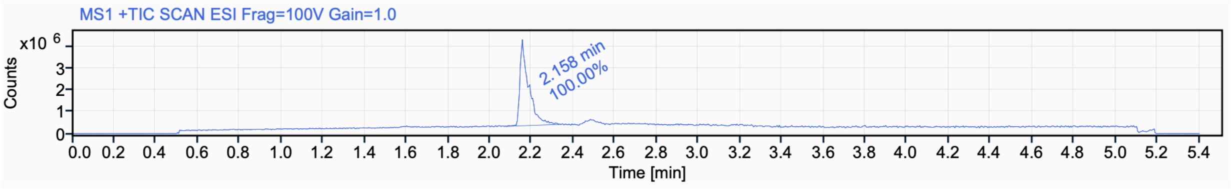


Signal Name MS1 +TIC SCAN ESI Frag=100V Gain=1.0

| RT (min) | Signal description | Area | Area% | Peak Resolution USP |
|---|---|---|---|---|
| 2.158 | MS1 +TIC SCAN ESI Frag=100V Gain=1.0 | 11563796.8 | 100.00 | |

$^{1}$H NMR (400 MHz, $CD_3CN$) for **2g**

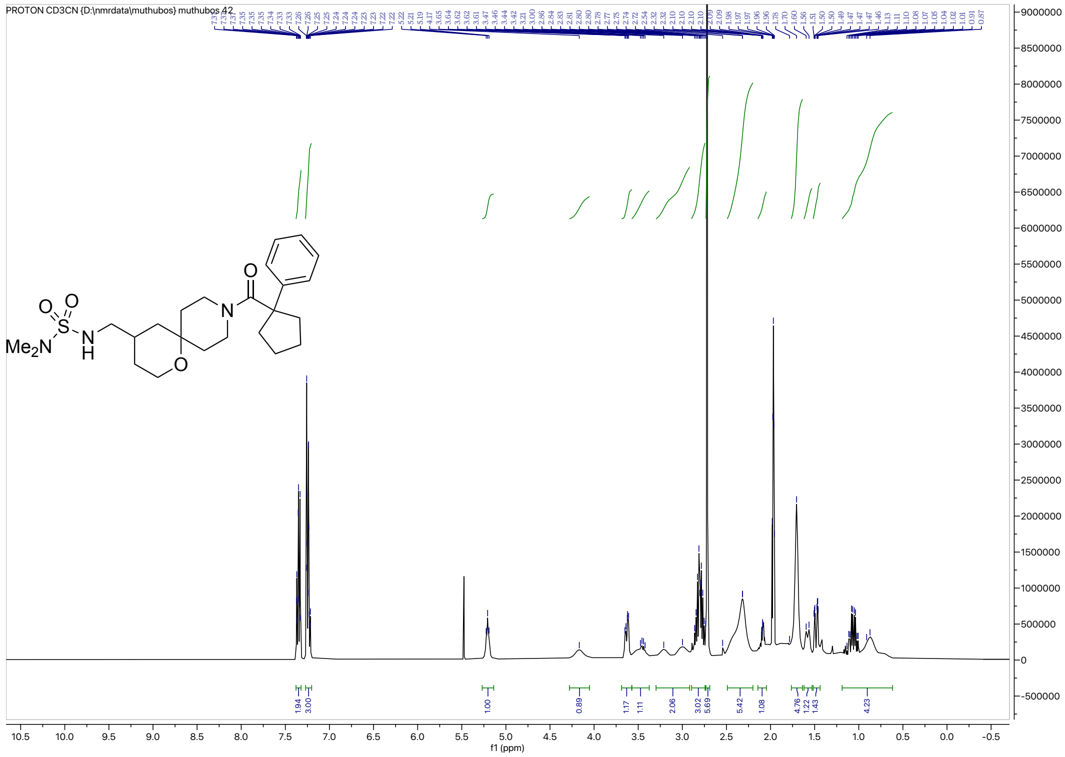

$^{13}$C NMR (126 MHz, $CD_3CN$) for **2g**

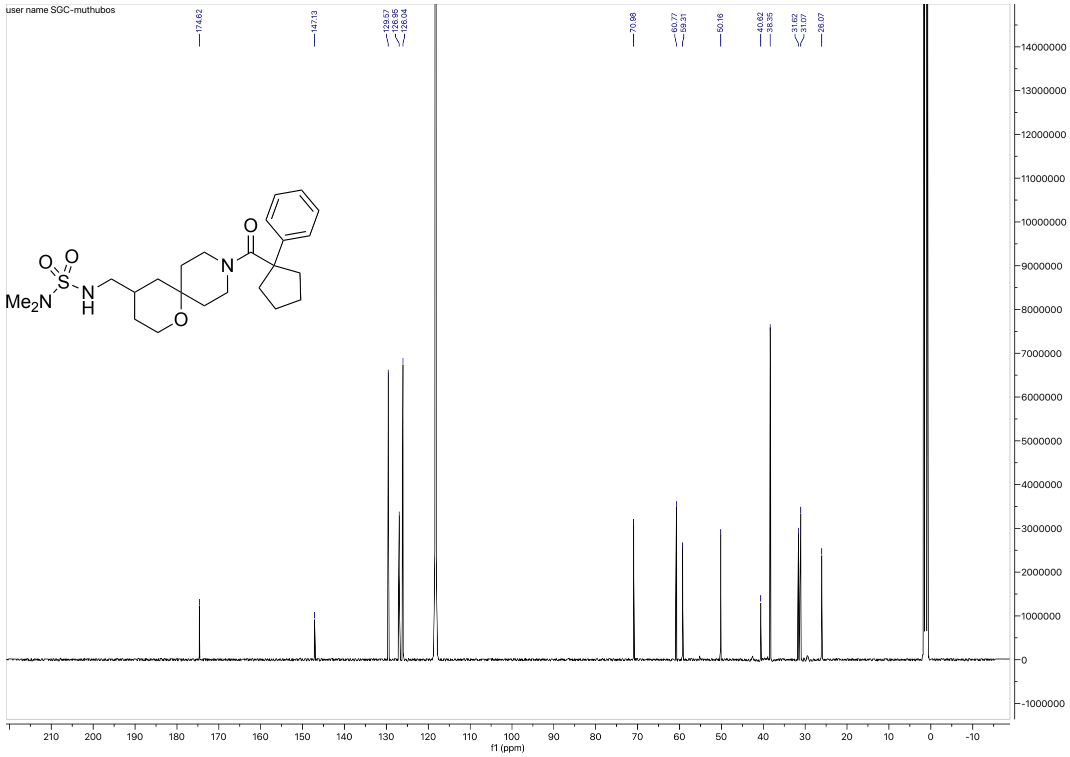

HRMS for **2g**

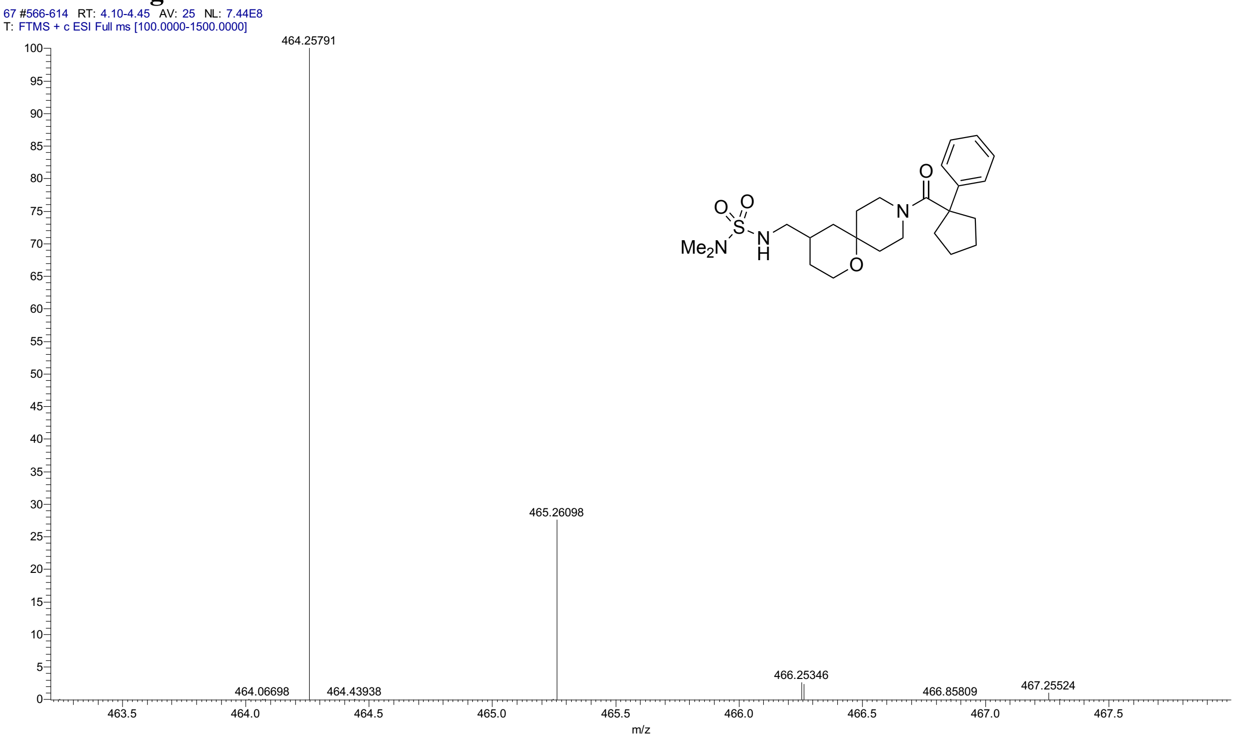


HPLC for **2g**

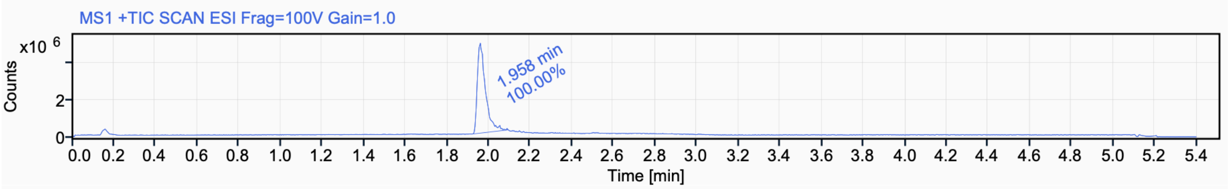


Signal Name MS1 +TIC SCAN ESI Frag=100V Gain=1.0

| RT (min) | Signal description | m/z | Purity UV(‰) / MS(%) | Area | Area% |
|---|---|---|---|---|---|
| 1.958 | MS1 +TIC SCAN ESI Frag=100V Gain=1.0 | | | 13186384.3 | 100.00 |

$^{1}$H NMR (500 MHz, $CDCl_3$) for **2h**

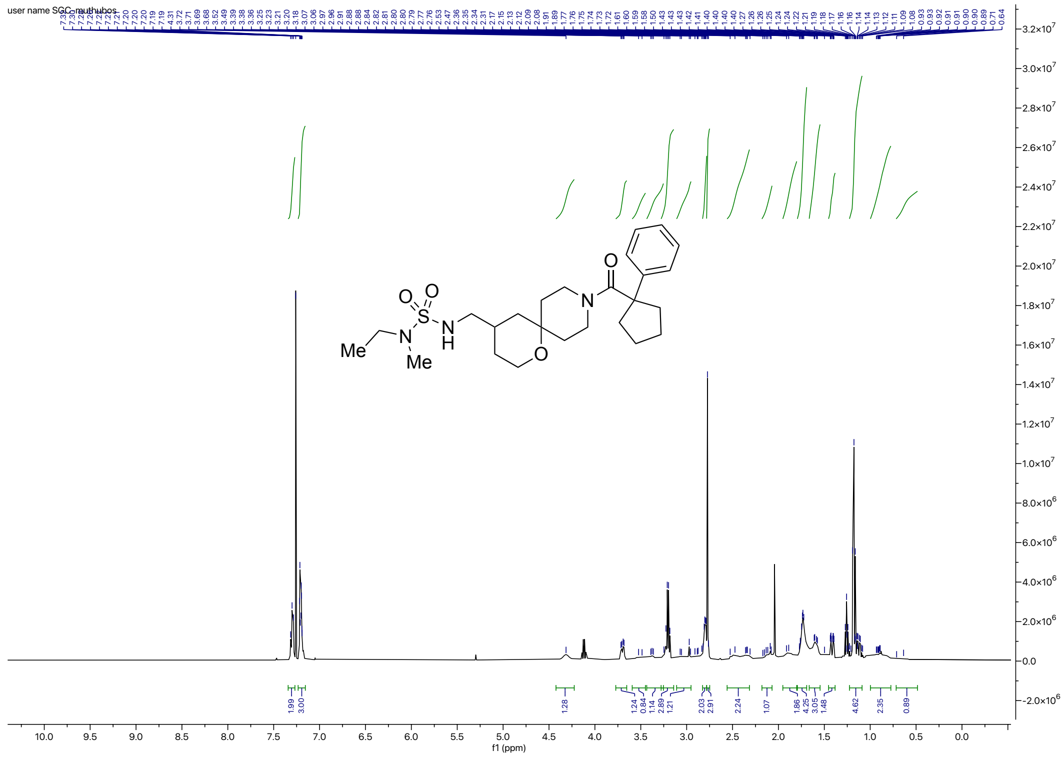

$^{13}$C NMR (126 MHz, $CDCl_3$) for **2h**

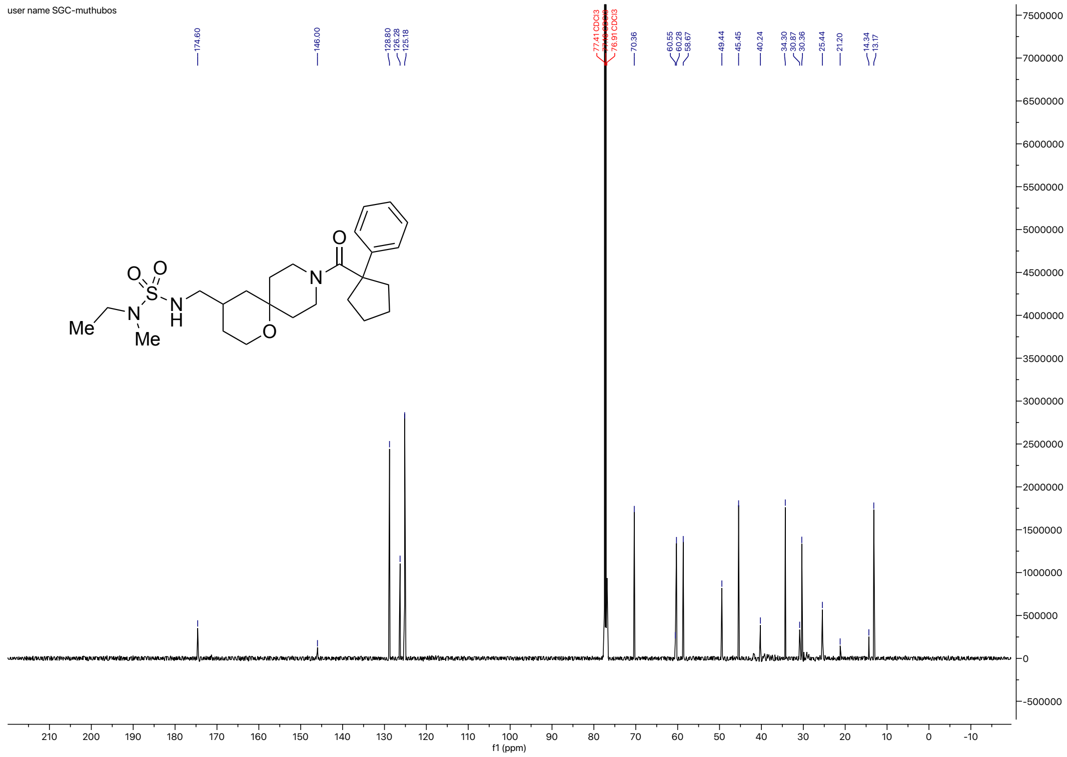

### HRMS for **2h**

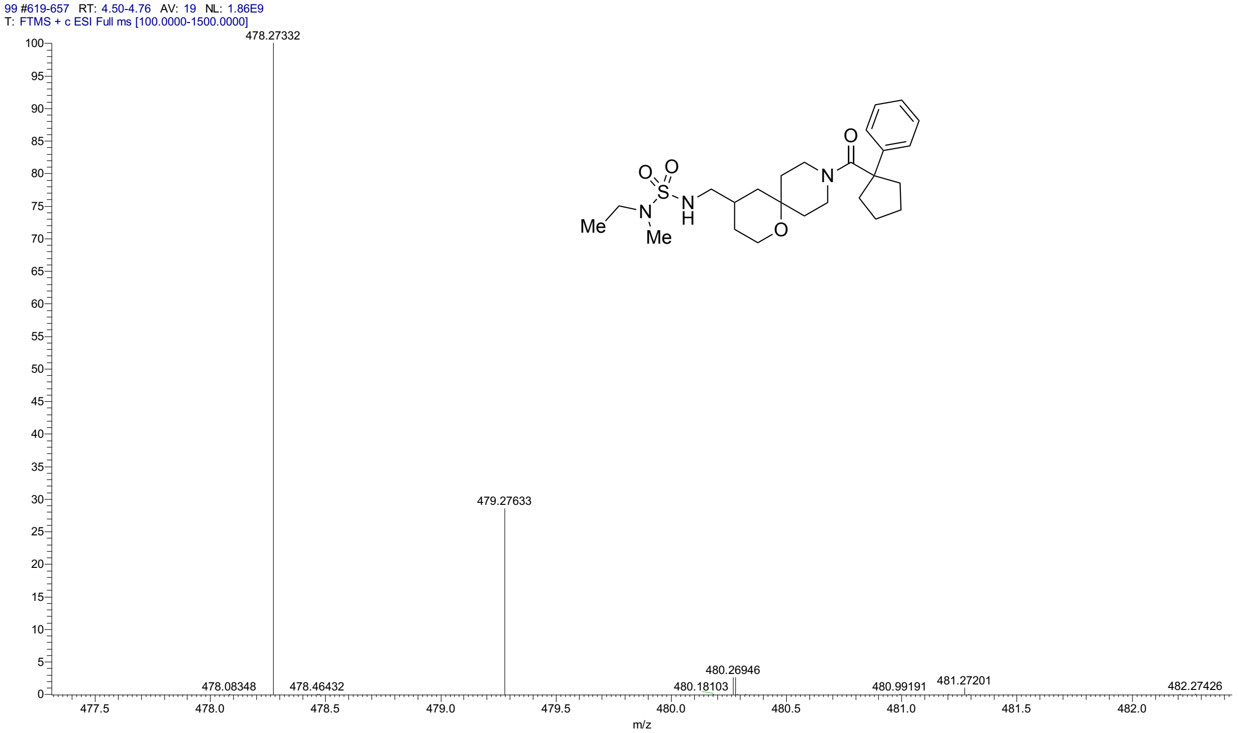


### HPLC for **2h**

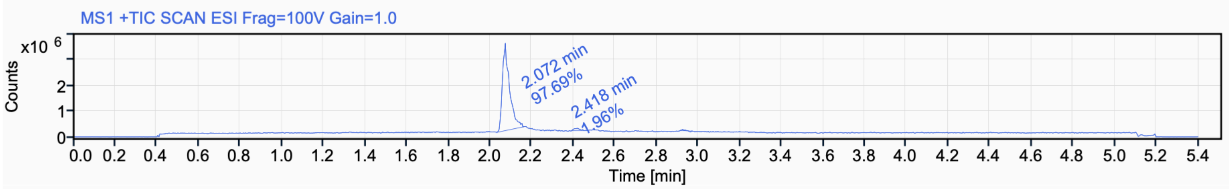


Signal Name MS1 +TIC SCAN ESI Frag=100V Gain=1.0

| RT (min) | Signal description | m/z | Purity UV(‰) / MS(%) | Area | Area% |
|---|---|---|---|---|---|
| 2.072 | MS1 +TIC SCAN ESI Frag=100V Gain=1.0 | | | 8104501.2 | 97.69 |
| 2.418 | MS1 +TIC SCAN ESI Frag=100V Gain=1.0 | | | 162322.4 | 1.96 |
| 2.924 | MS1 +TIC SCAN ESI Frag=100V Gain=1.0 | | | 29558.1 | 0.36 |

$^{1}$H NMR (500 MHz, $CD_3CN$) for **2i**

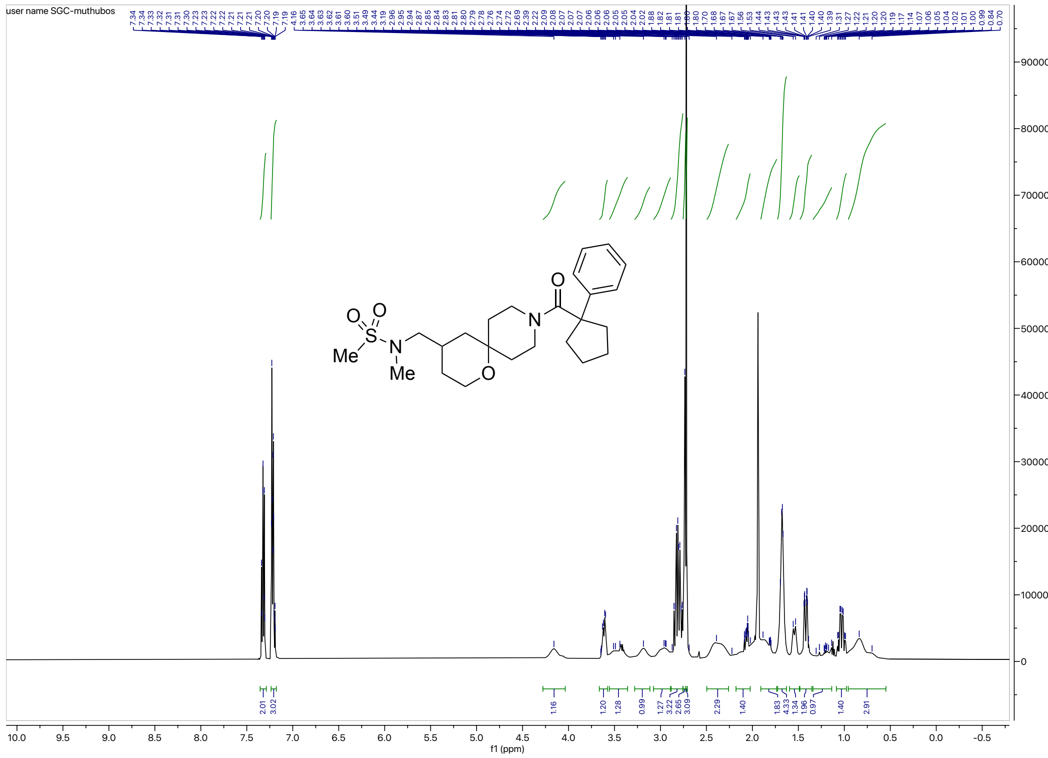


$^{13}$C NMR (126 MHz, $CD_3CN$) for **2i**

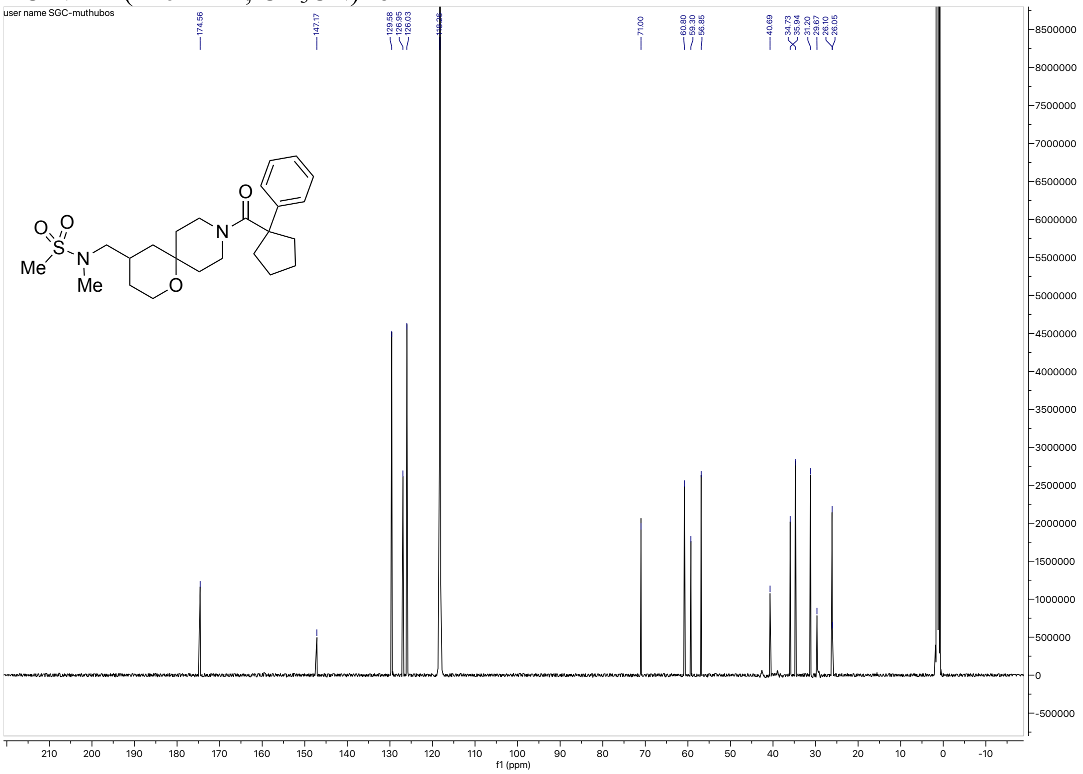

#### HRMS for **2i**

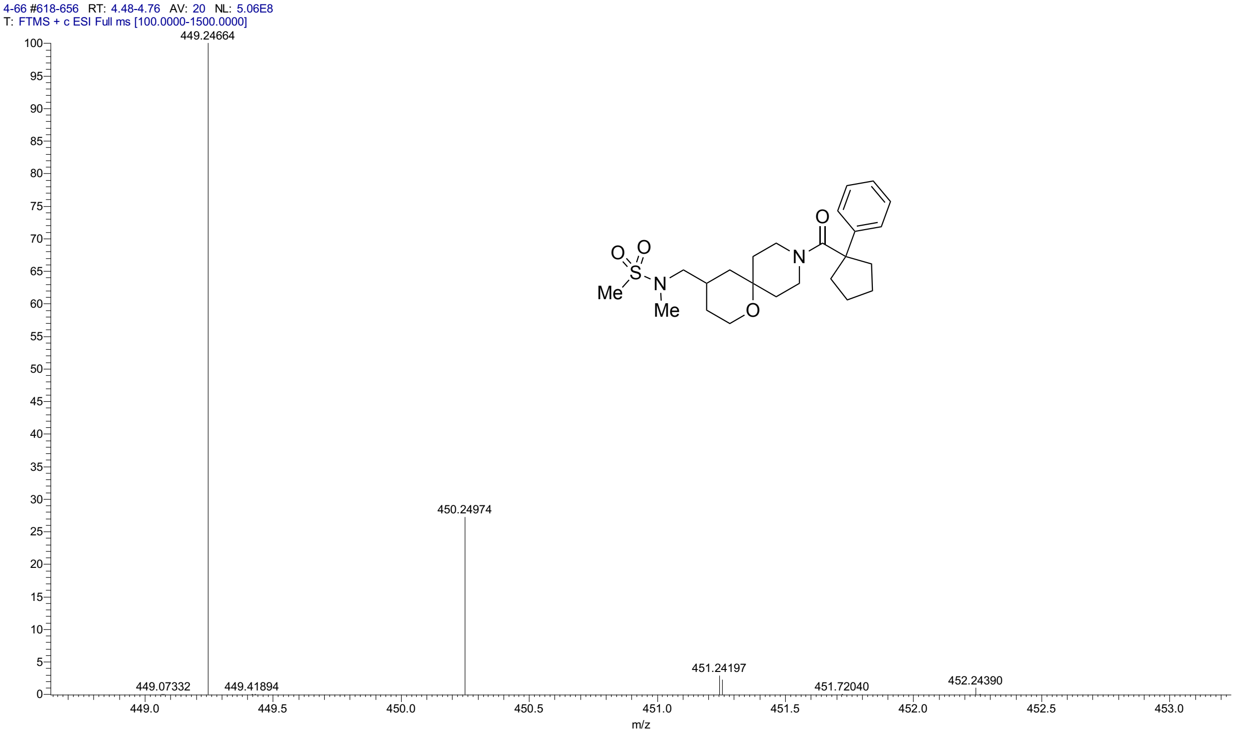


#### HPLC for **2i**

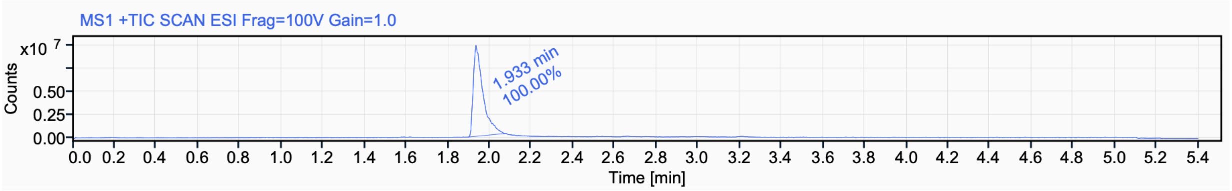


Signal Name MS1 +TIC SCAN ESI Frag=100V Gain=1.0

| RT (min) | Signal description | m/z | Purity UV(‰) / MS(%) | Area | Area% |
|---|---|---|---|---|---|
| 1.933 | MS1 +TIC SCAN ESI Frag=100V Gain=1.0 | | | 31401454.1 | 100.00 |

[1]H NMR (500 MHz, $CD_3CN$) for **2j**

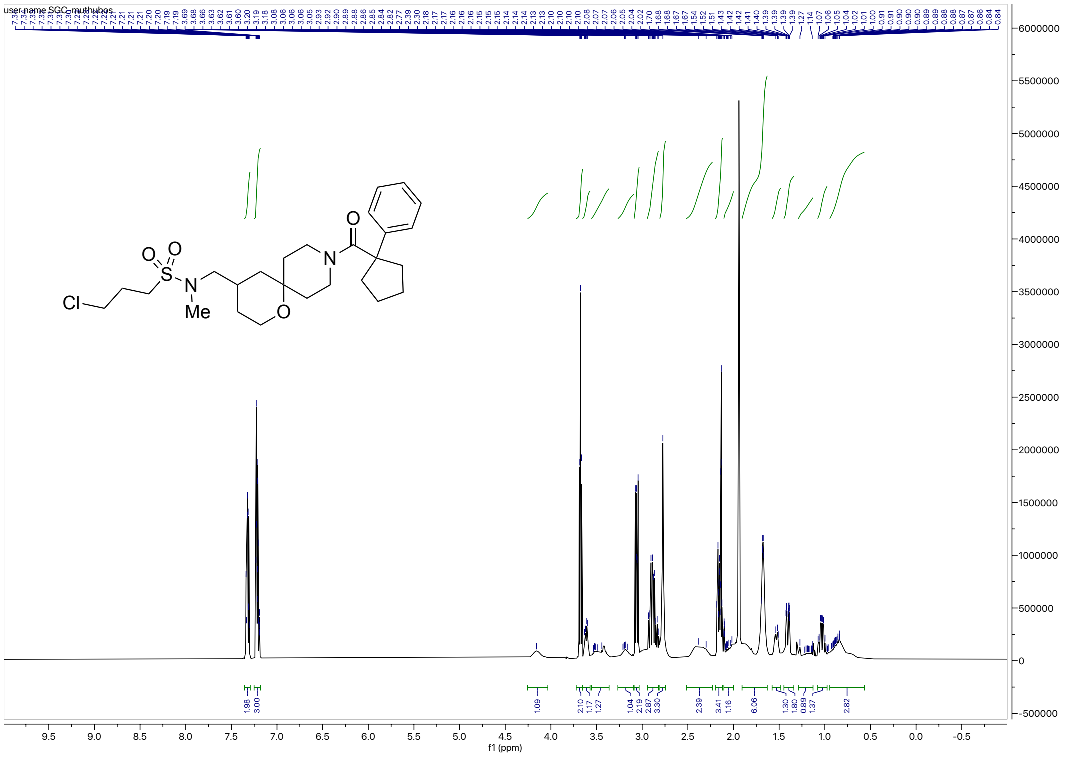

[13]C NMR (126 MHz, $CD_3CN$) for **2j**

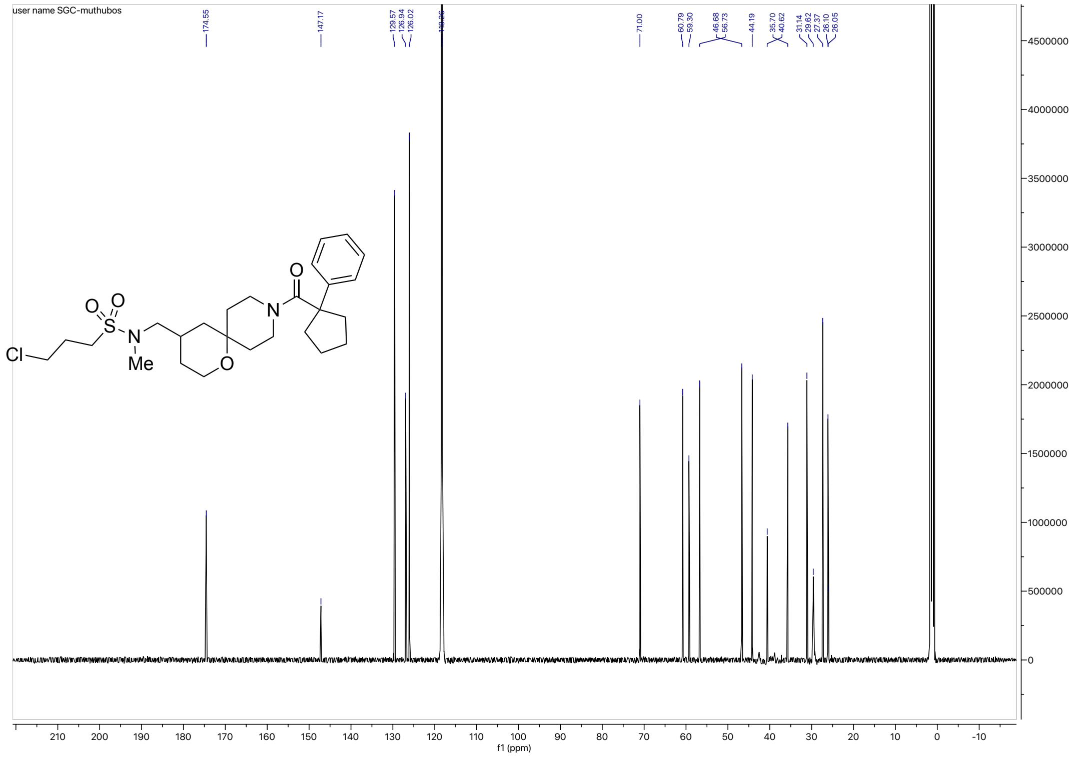

HRMS for **2j**

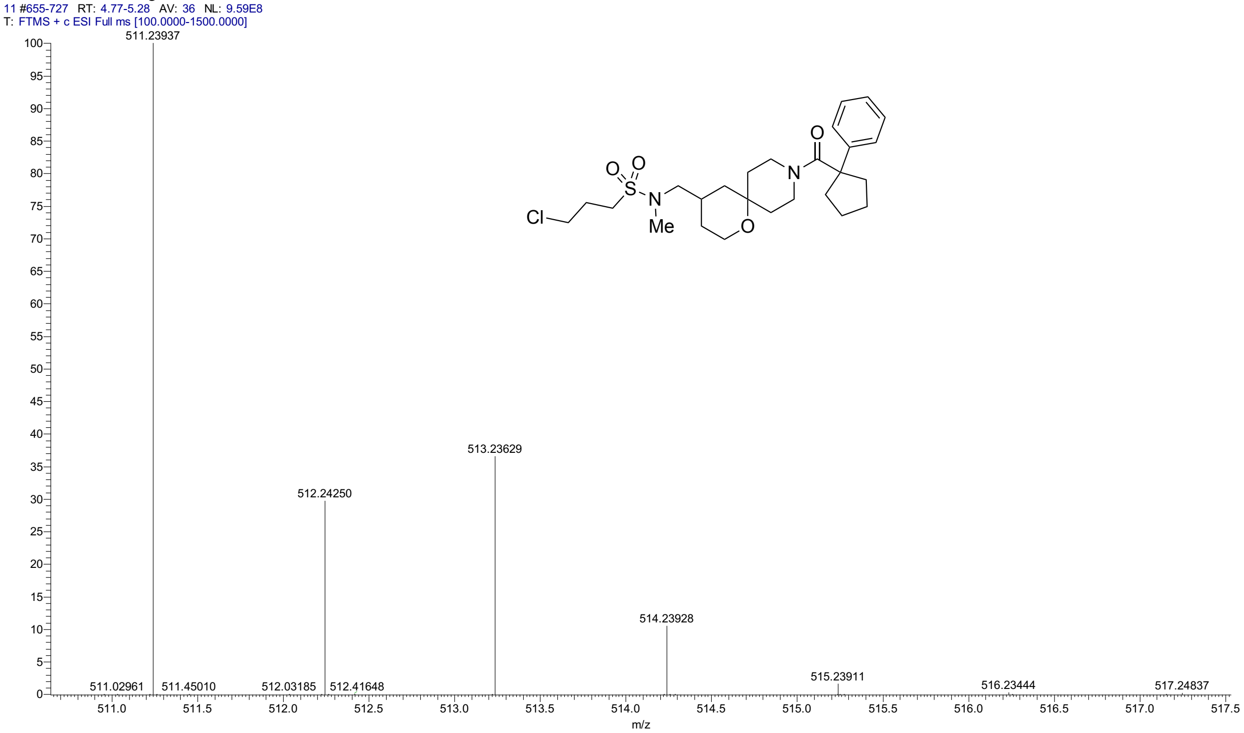


HPLC for **2j**

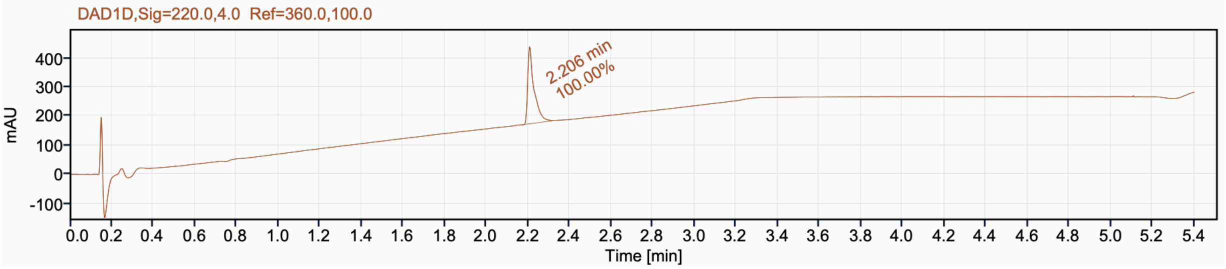


Signal Name DAD1D

| RT (min) | Signal description | Area | Area% | Peak Resolution USP |
|---|---|---|---|---|
| 2.206 | DAD1D,Sig=220.0,4.0 Ref=360.0,100.0 | 565.1 | 100.00 | |

$^{1}$H NMR (500 MHz, $CD_3CN$) for **2k**

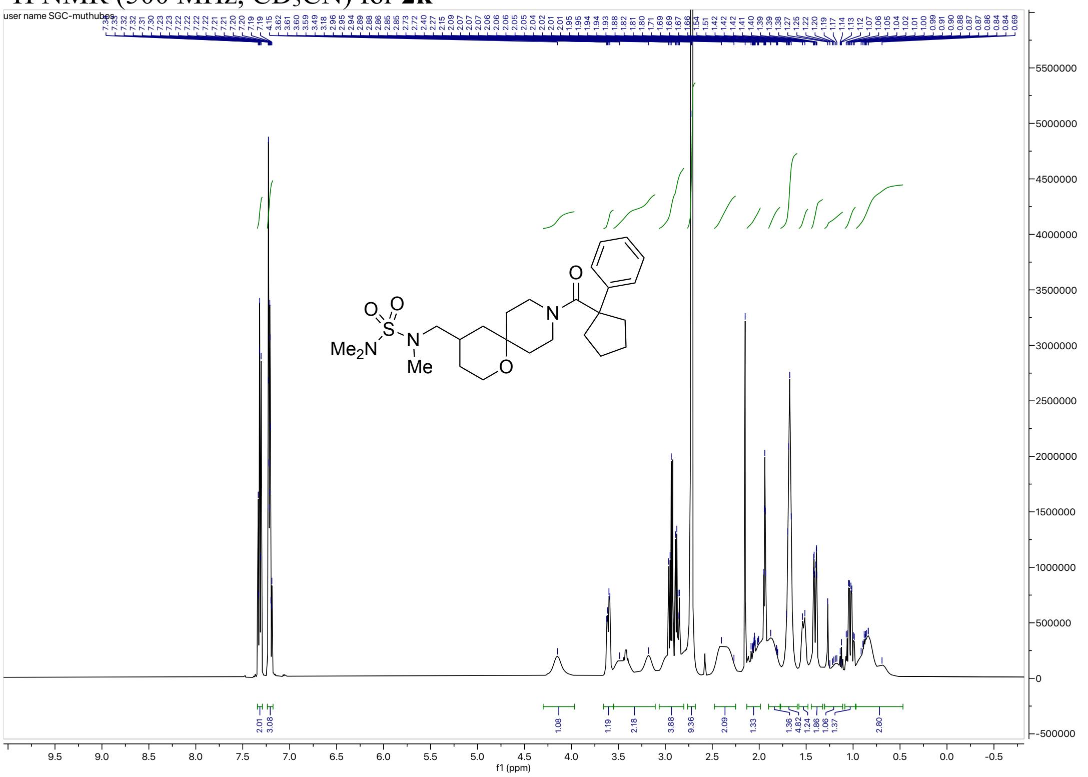

$^{13}$C NMR (126 MHz, $CD_3CN$) for **2k**

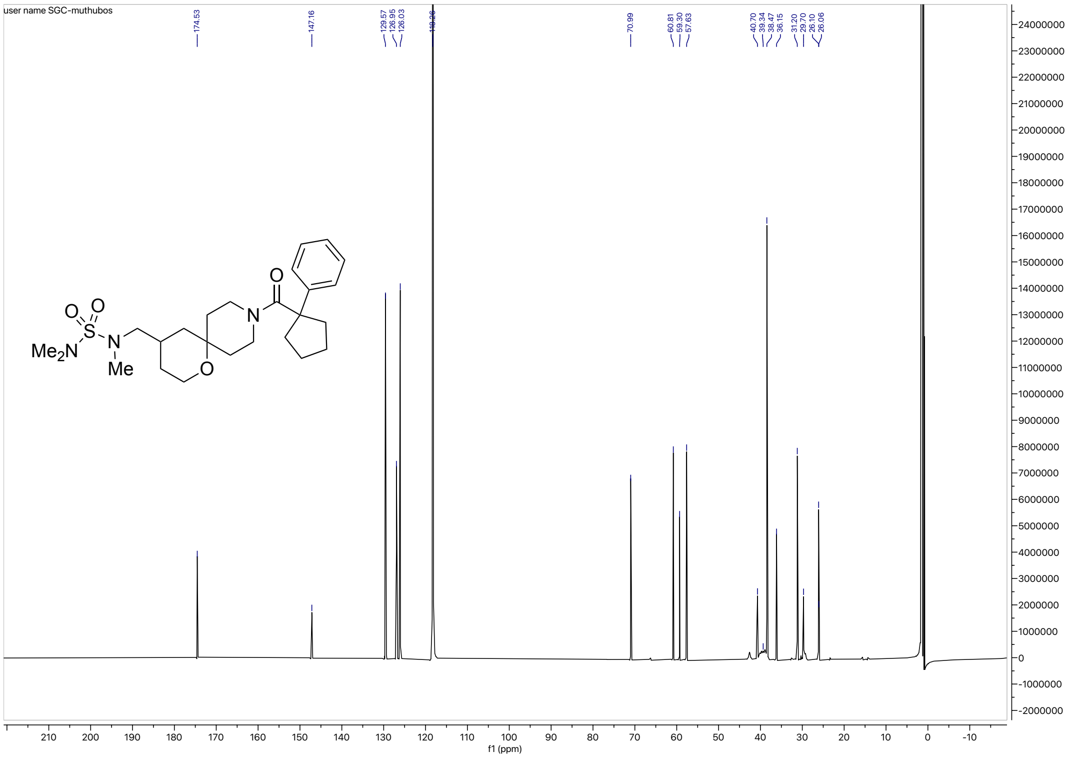

#### HRMS for **2k**

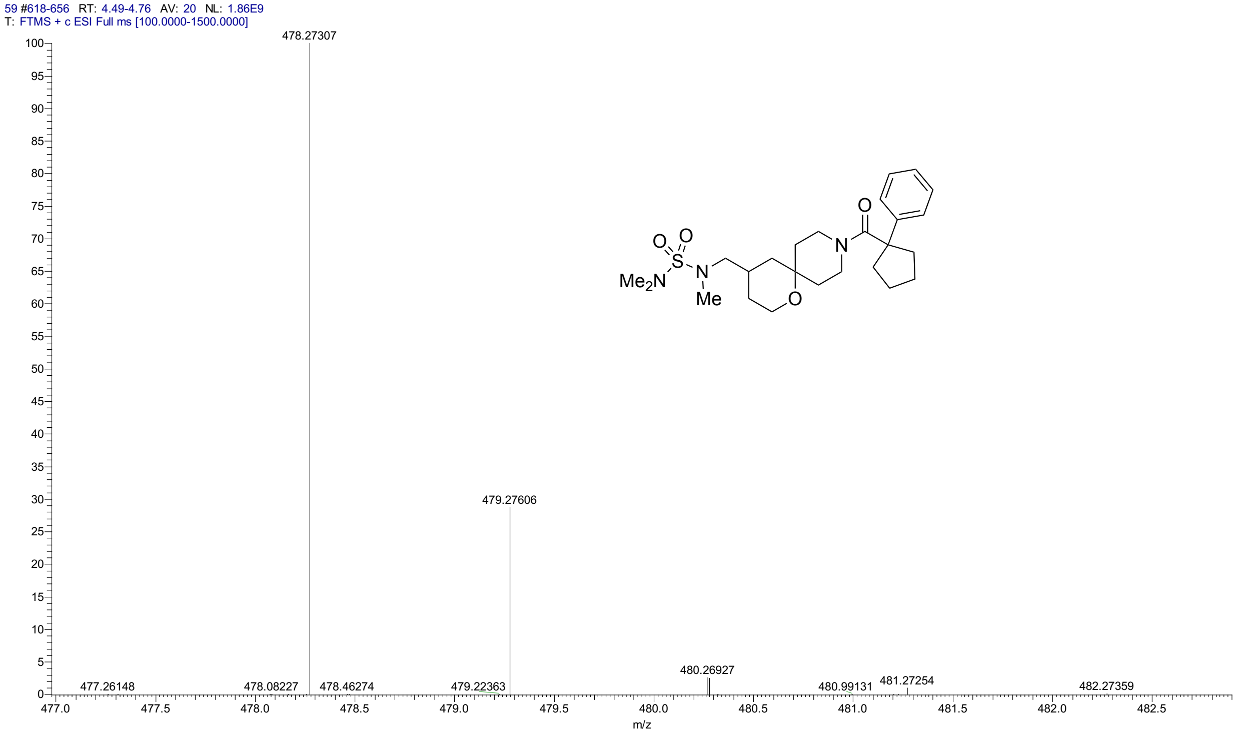


#### HPLC for **2k**

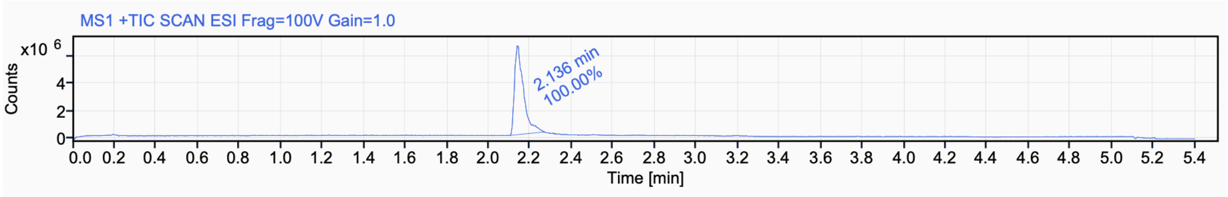


Signal Name MS1 +TIC SCAN ESI Frag=100V Gain=1.0

| RT (min) | Signal description | m/z | Purity UV(‰) / MS(%) | Area | Area% |
|---|---|---|---|---|---|
| 2.136 | MS1 +TIC SCAN ESI Frag=100V Gain=1.0 | | | 19491885.0 | 100.00 |

$^{1}H$ NMR (500 MHz, $CDCl_3$) for **2l**

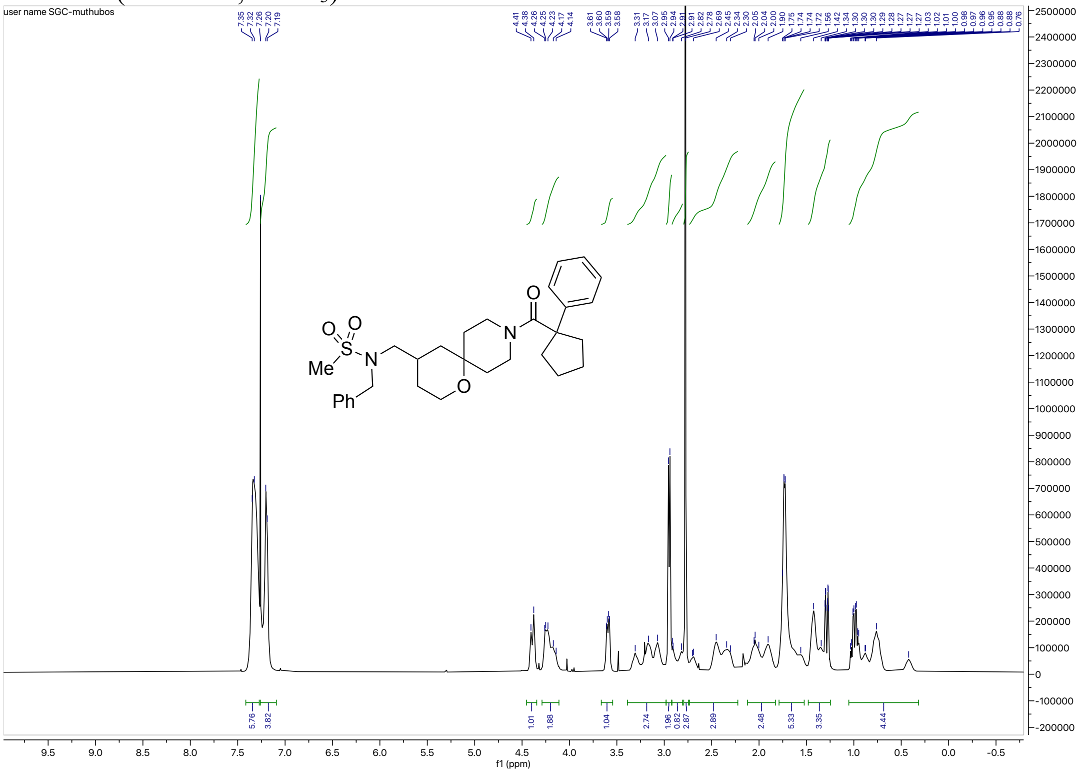

$^{13}C$ NMR (126 MHz, $CDCl_3$) for **2l**

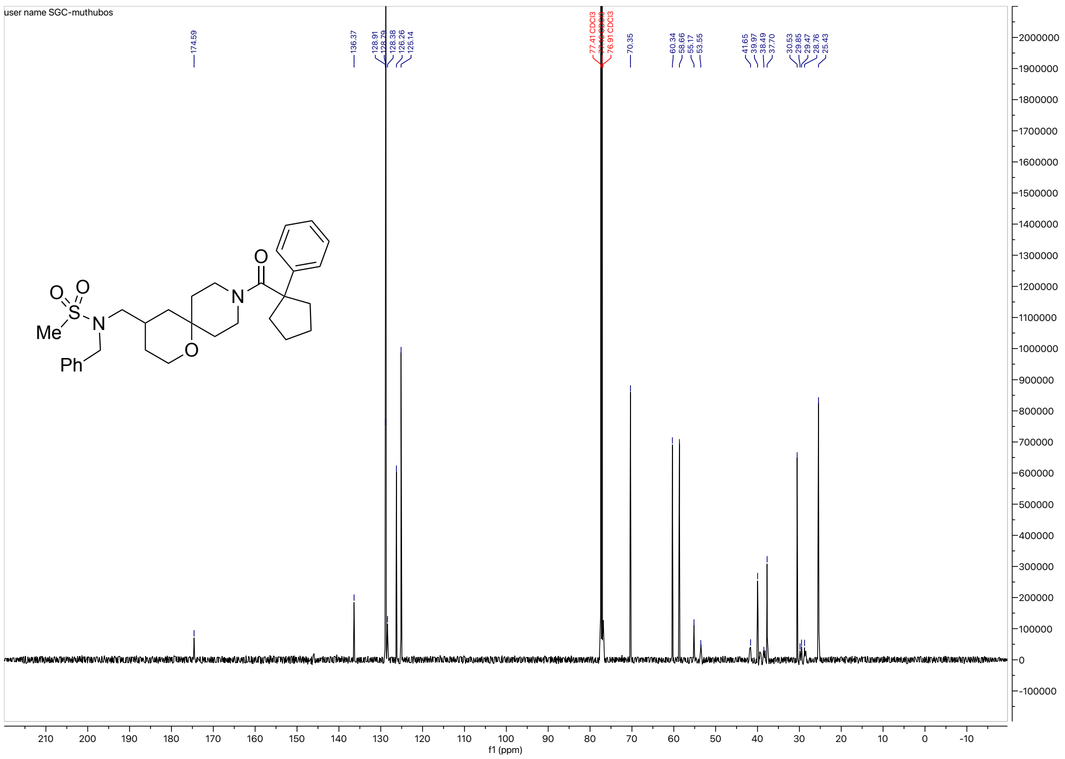

HRMS for **2l**

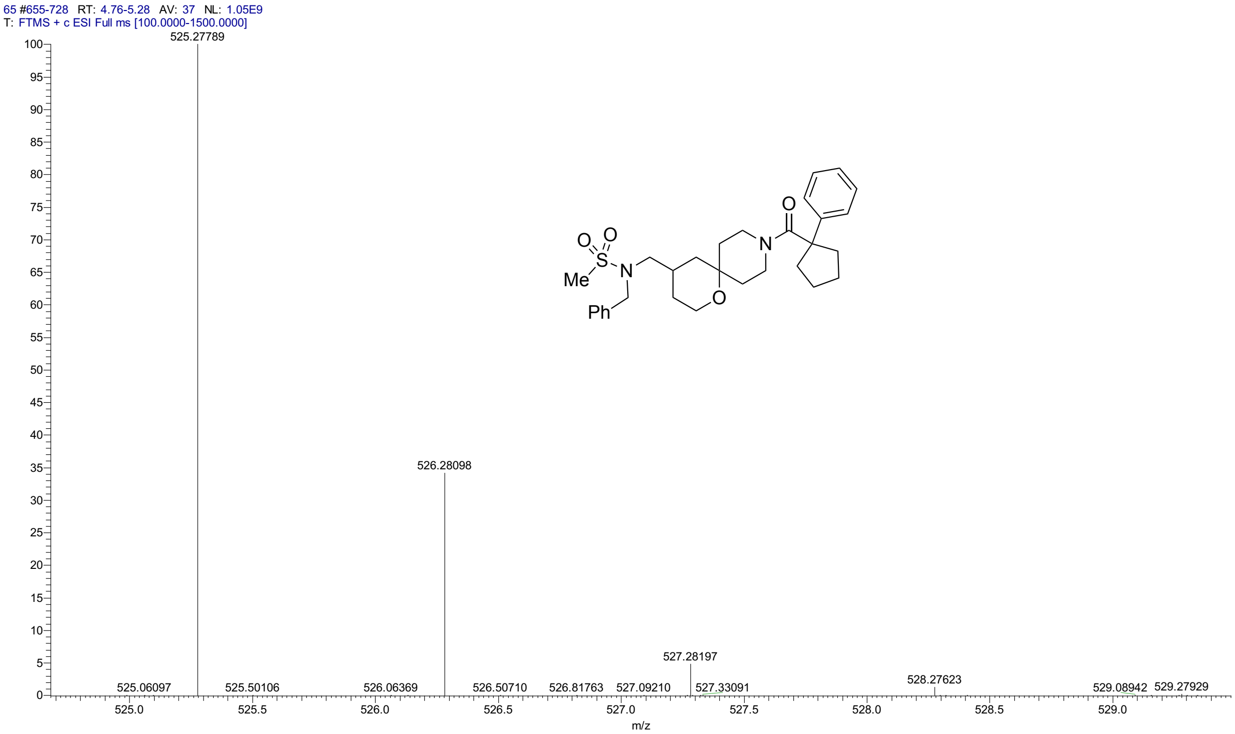


HPLC for **2l**

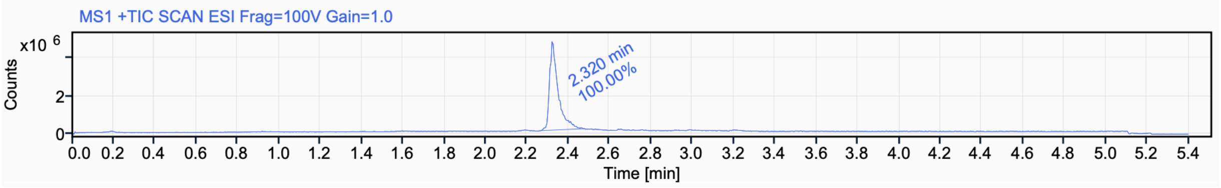


Signal Name MS1 +TIC SCAN ESI Frag=100V Gain=1.0

| RT (min) | Signal description | m/z | Purity UV(‰) / MS(%) | Area | Area% |
|---|---|---|---|---|---|
| 2.320 | MS1 +TIC SCAN ESI Frag=100V Gain=1.0 | | | 13585042.2 | 100.00 |

$^{1}$H NMR (500 MHz, $CD_3CN$) for **2m**

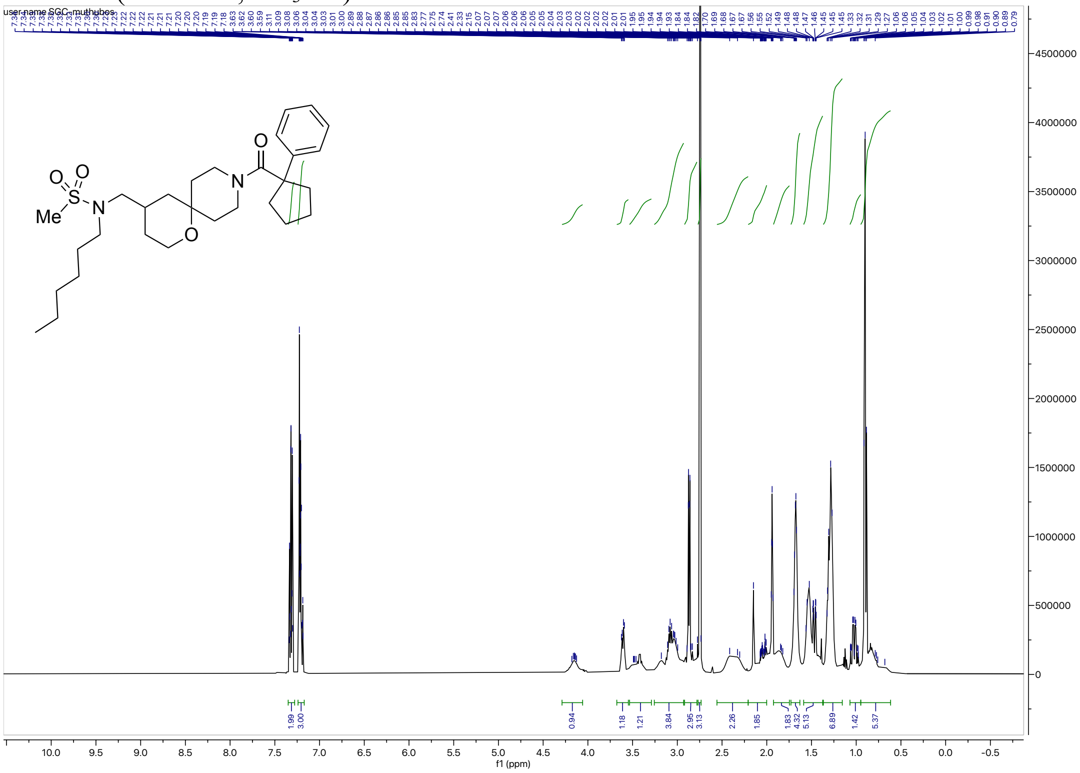


$^{13}$C NMR (126 MHz, $CD_3CN$) for **2m**

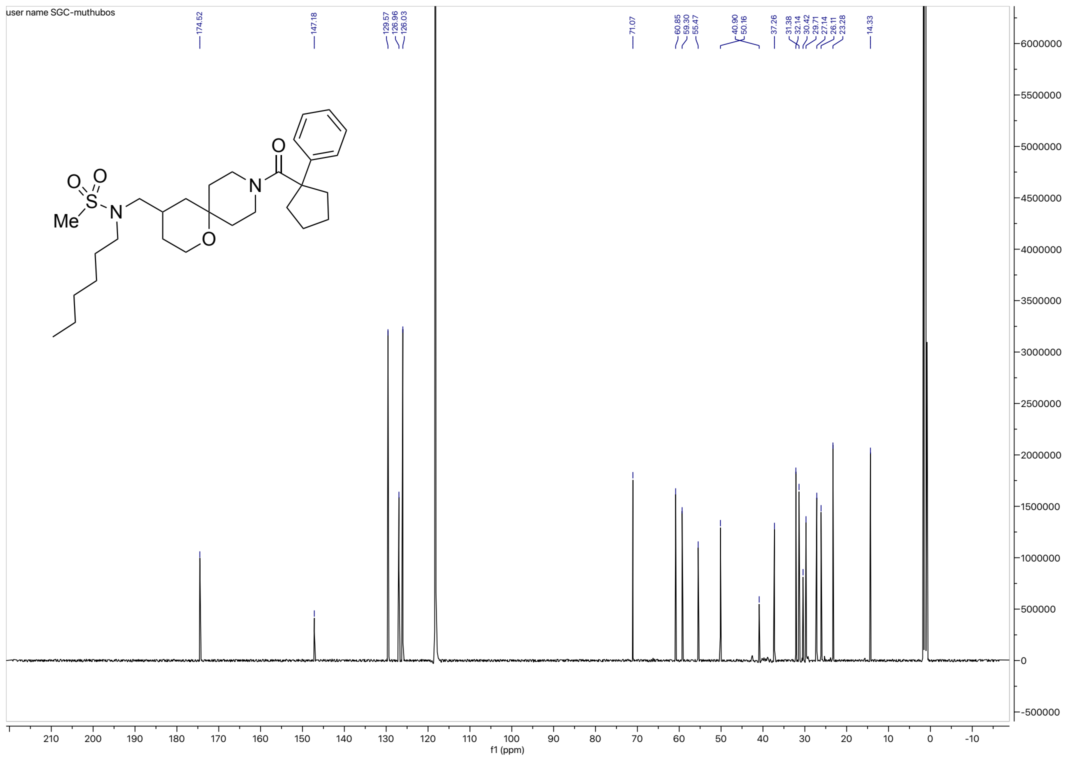

### HRMS for **2m**

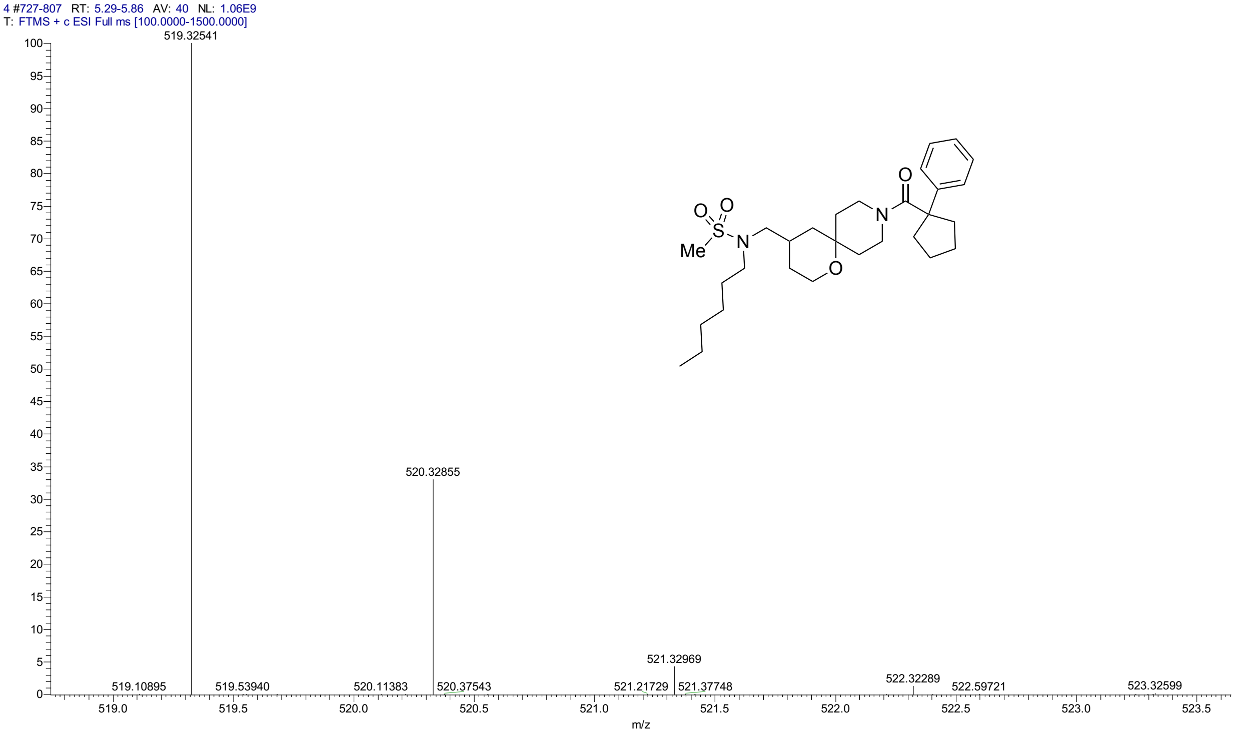


### HPLC for **2m**

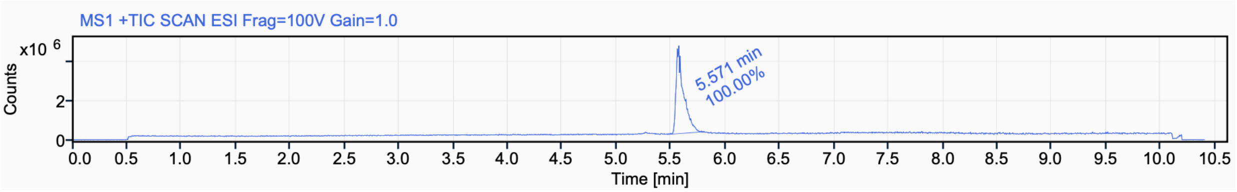


Signal Name MS1 +TIC SCAN ESI Frag=100V Gain=1.0

| RT (min) | Signal description | Area | Area% | Peak Resolution USP |
|---|---|---|---|---|
| 5.571 | MS1 +TIC SCAN ESI Frag=100V Gain=1.0 | 21390179.2 | 100.00 | |

[1]H NMR (500 MHz, $CD_3CN$) for **2n**

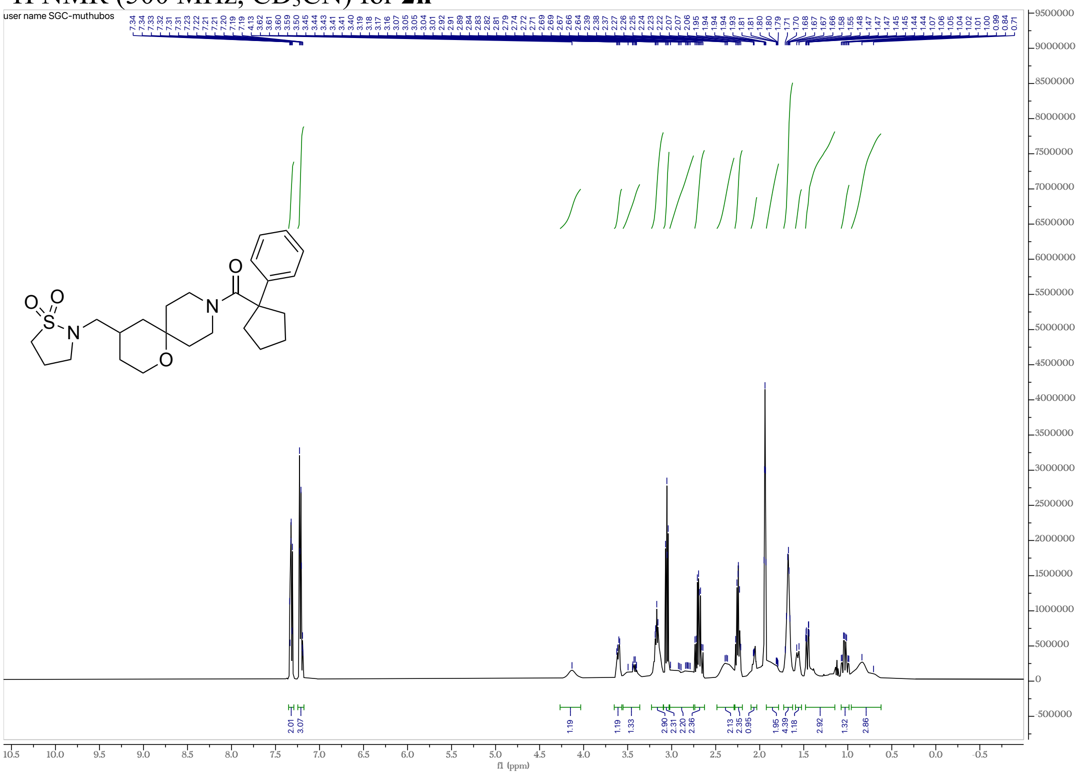


[13]C NMR (126 MHz, $CD_3CN$) for **2n**

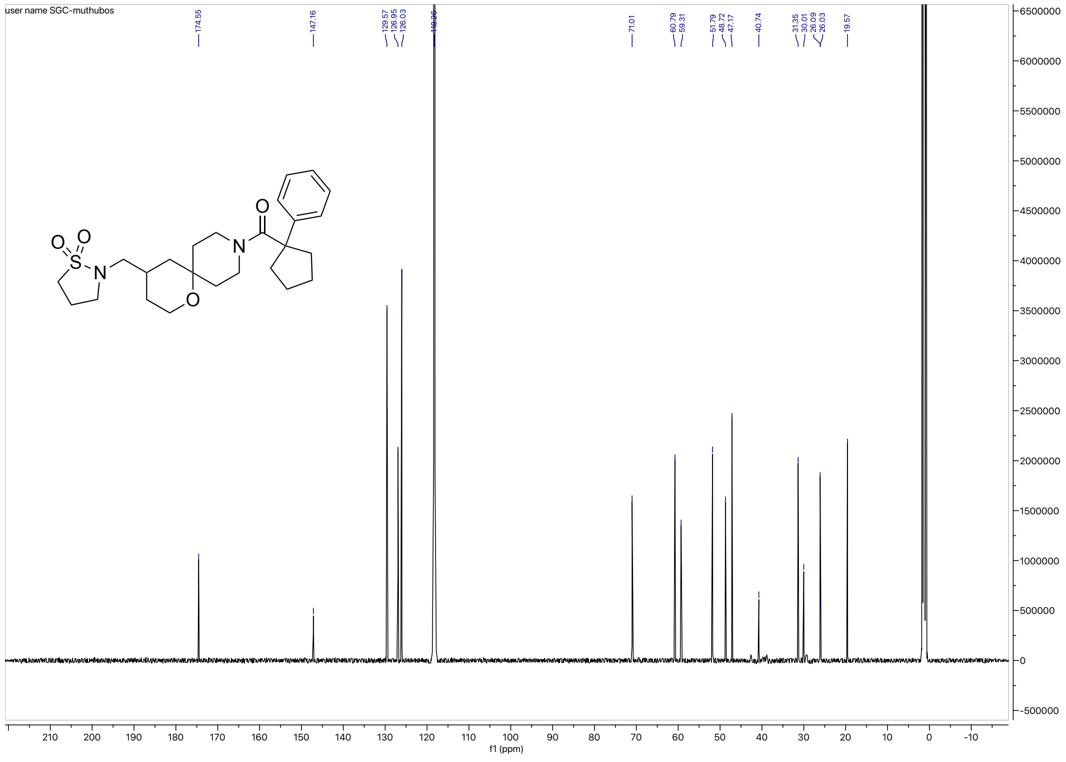

HRMS for **2n**

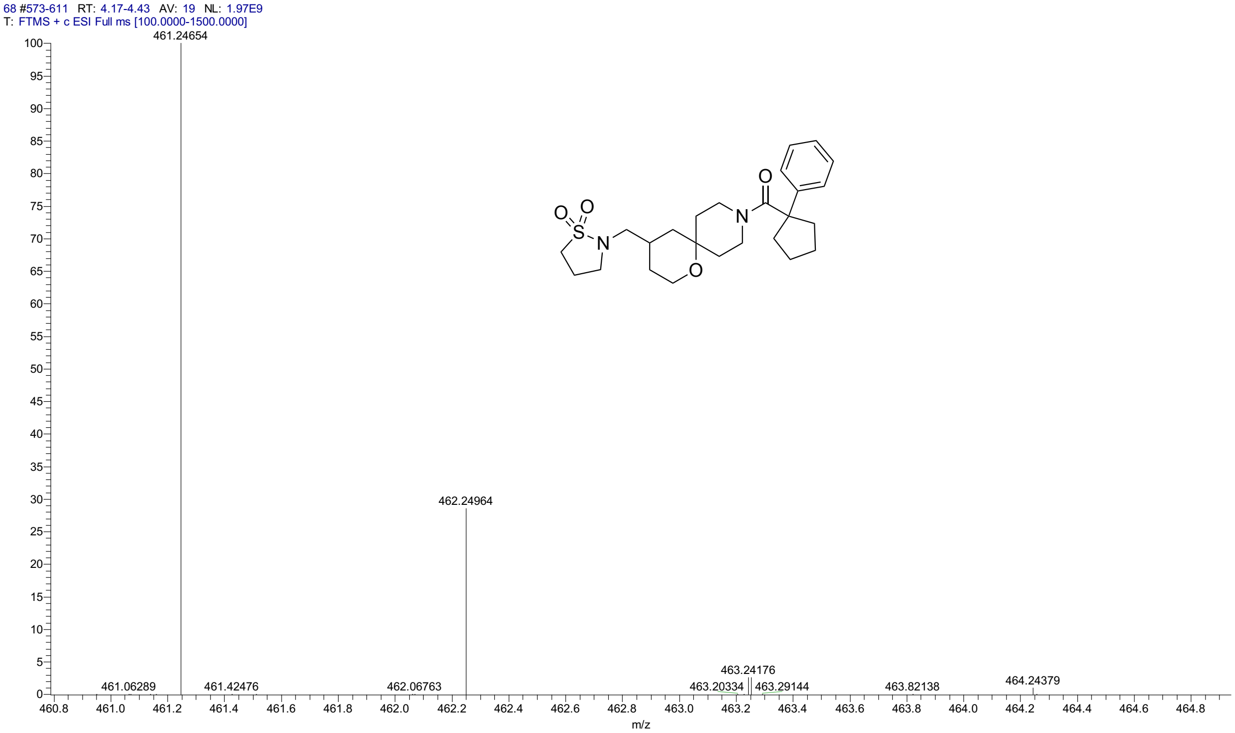


HPLC for **2n**

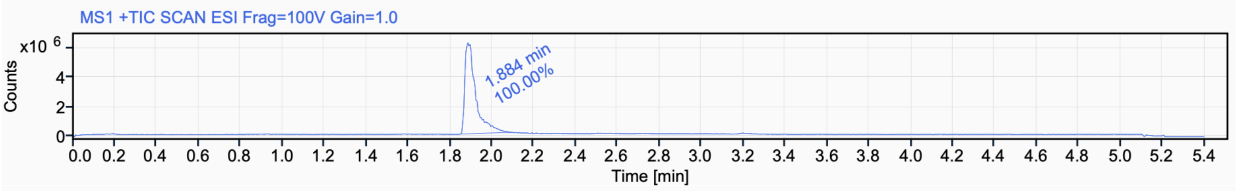


Signal Name MS1 +TIC SCAN ESI Frag=100V Gain=1.0

| RT (min) | Signal description | m/z | Purity UV(‰) / MS(%) | Area | Area% |
|---|---|---|---|---|---|
| 1.884 | MS1 +TIC SCAN ESI Frag=100V Gain=1.0 | | | 22275219.3 | 100.00 |

$^{1}$H NMR (500 MHz, $CD_3CN$) for **2o**

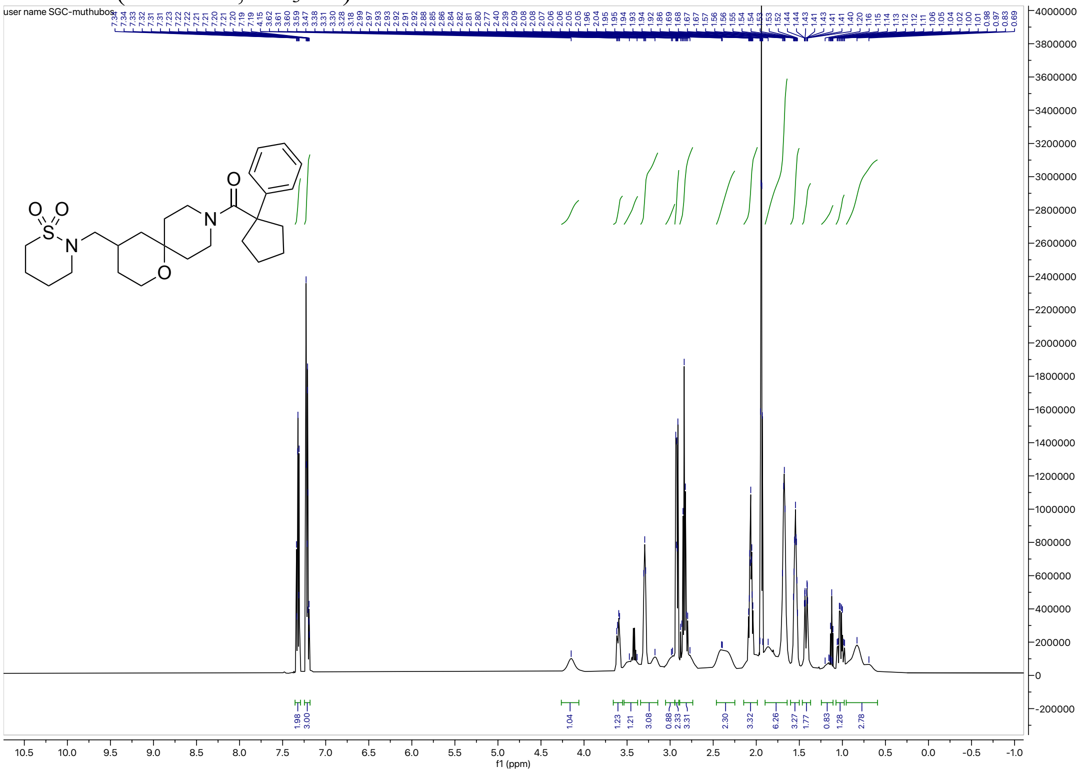

$^{13}$C NMR (126 MHz, $CD_3CN$) for **2o**

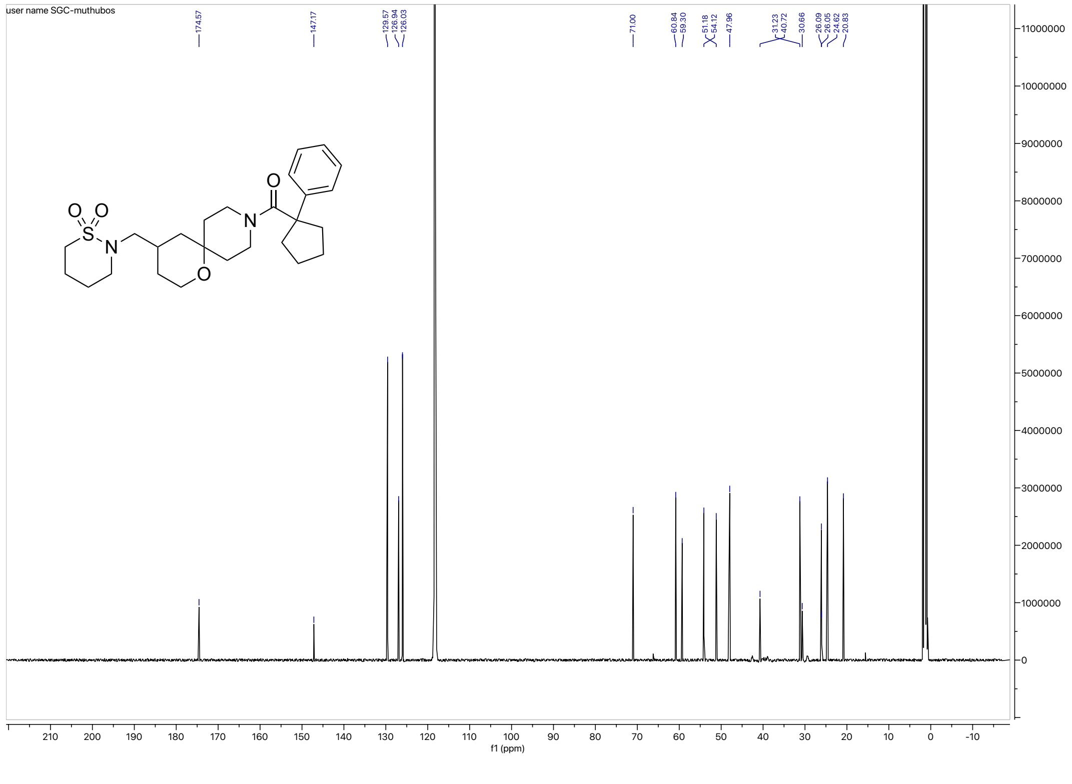

HRMS for **2o**

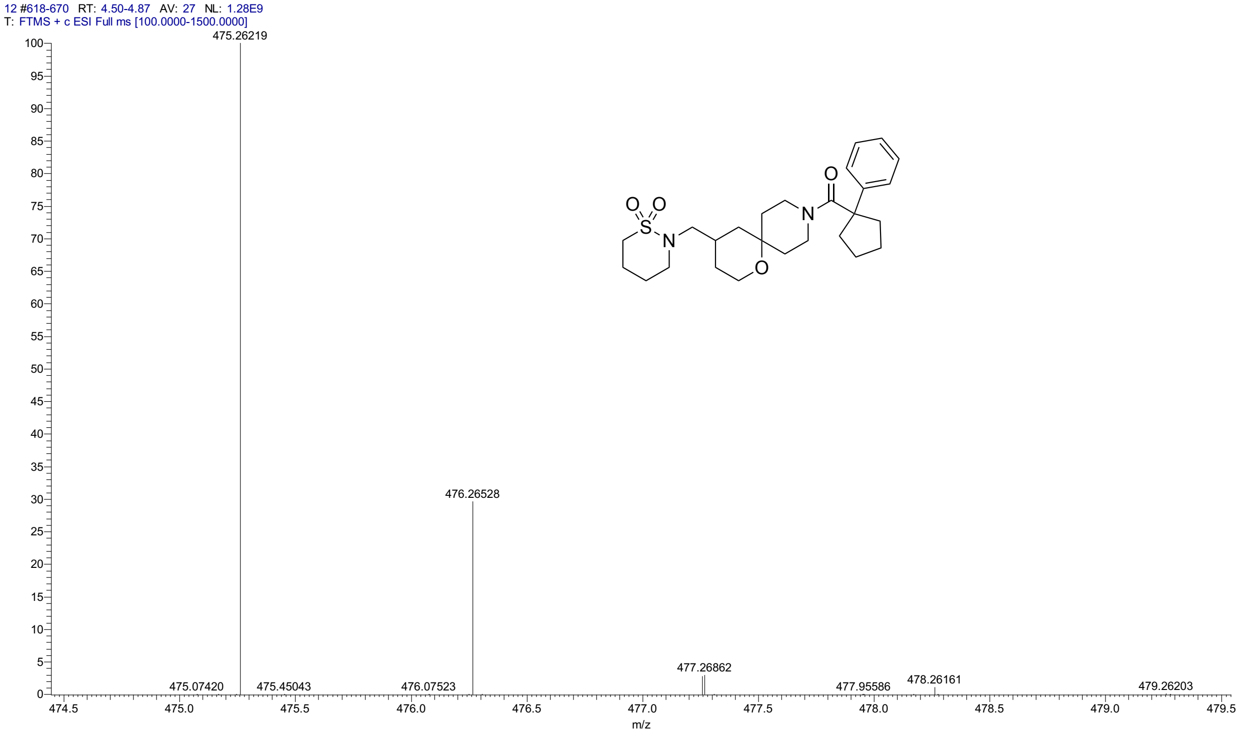


HPLC for **2o**

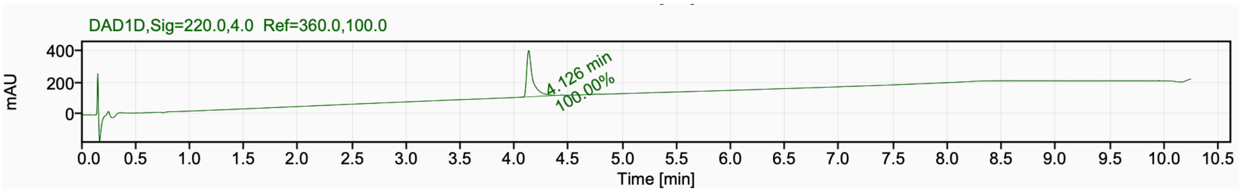


Signal Name DAD1D

| RT (min) | Signal description | Area | Area% | Peak Resolution USP |
|---|---|---|---|---|
| 4.126 | DAD1D,Sig=220.0,4.0 Ref=360.0,100.0 | 1127.4 | 100.00 | |

[1]H NMR (500 MHz, $CD_3CN$) for **3a**

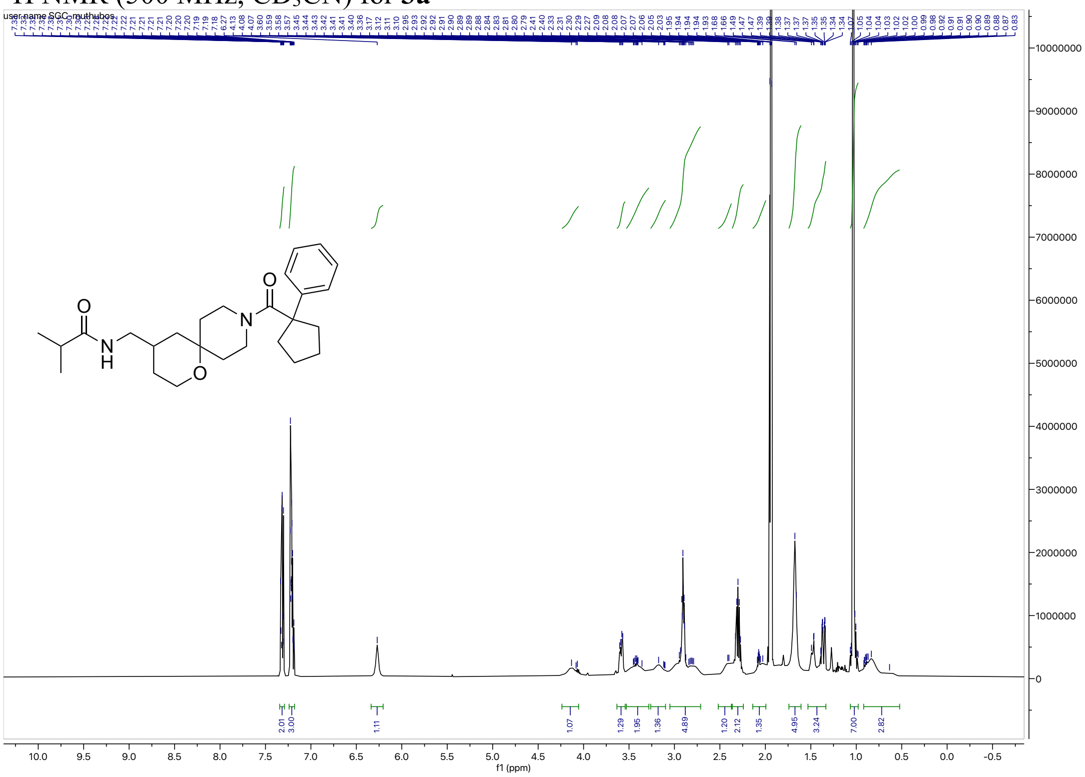


[13]C NMR (126 MHz, $CD_3CN$) for **3a**

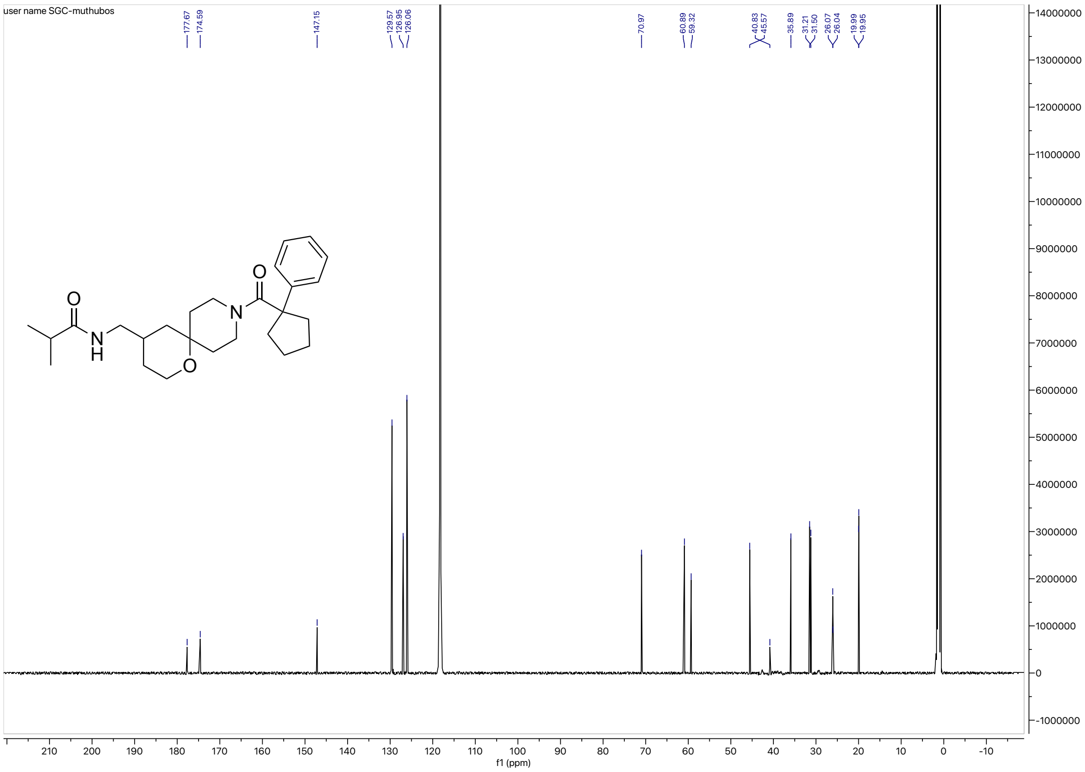

HRMS for **3a**

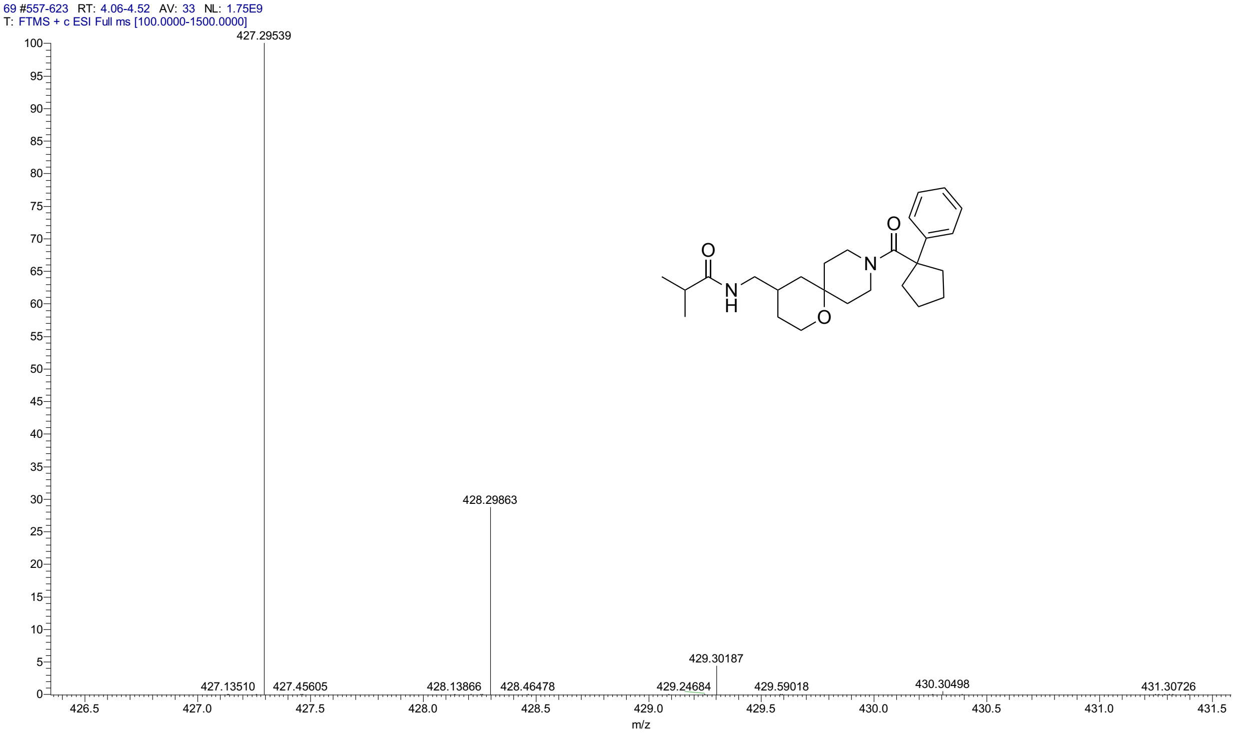


HPLC for **3a**

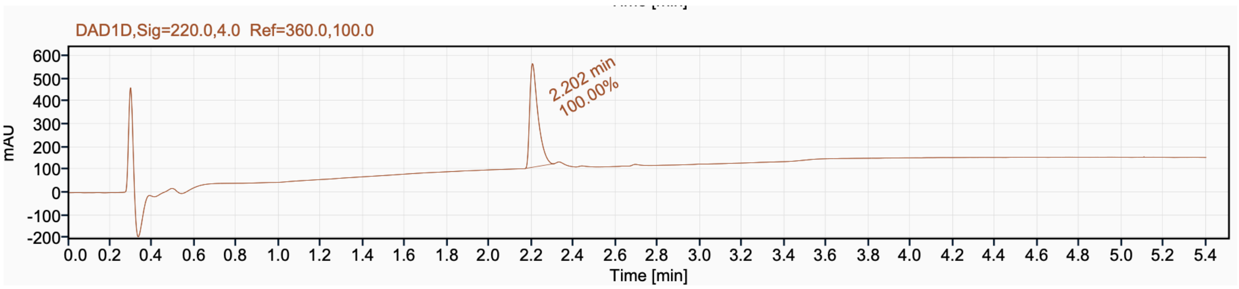


Signal Name DAD1D

| RT (min) | Signal description | Area | Area% | Peak Resolution USP |
|---|---|---|---|---|
| 2.202 | DAD1D,Sig=220.0,4.0 Ref=360.0,100.0 | 1249.8 | 100.00 | |

$^1$H NMR (500 MHz, $CD_3CN$) for **3b**

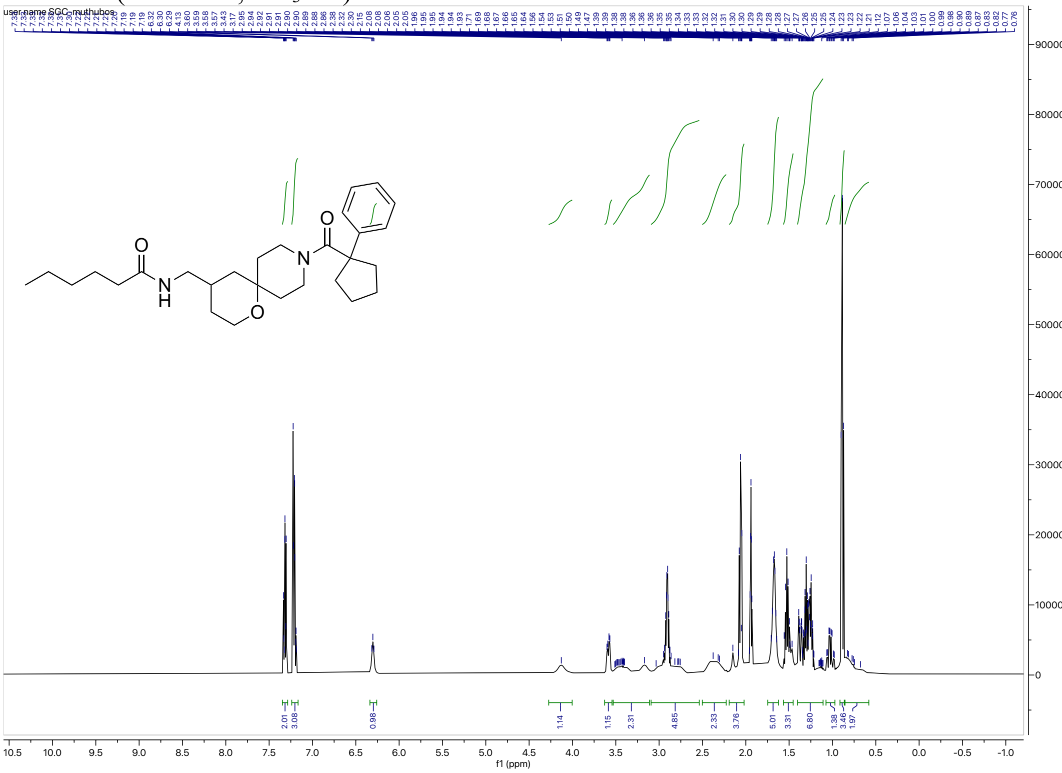

$^{13}$C NMR (126 MHz, $CD_3CN$) for **3b**

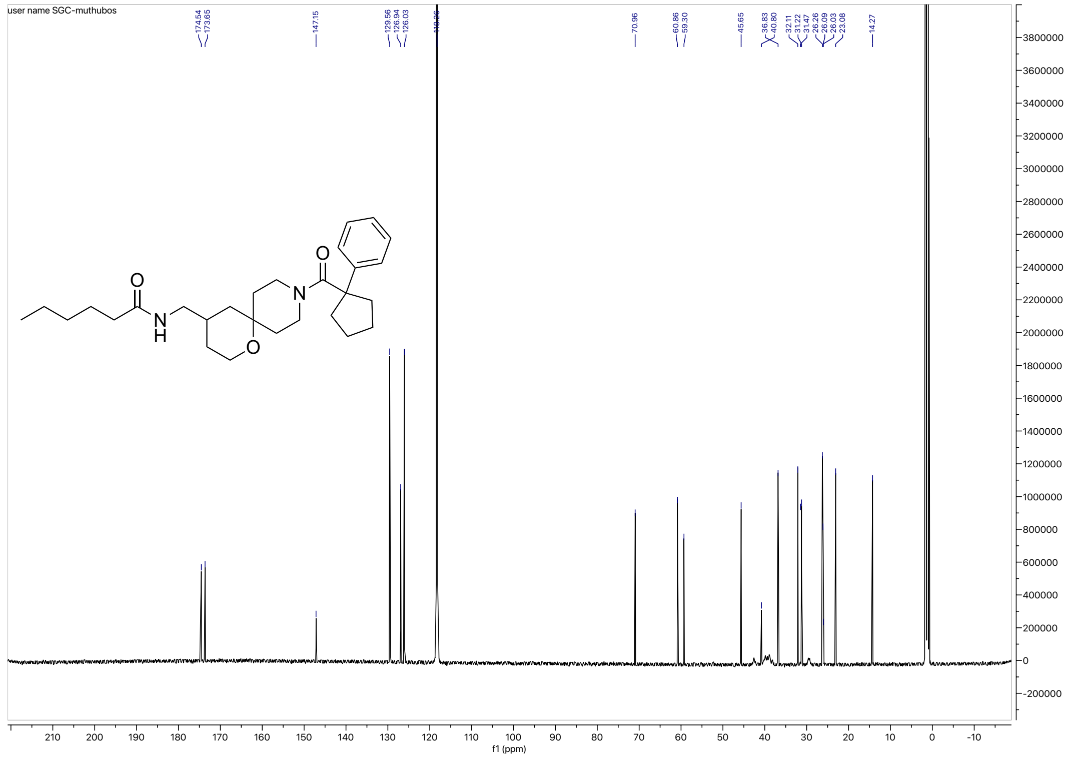

HRMS for **3b**

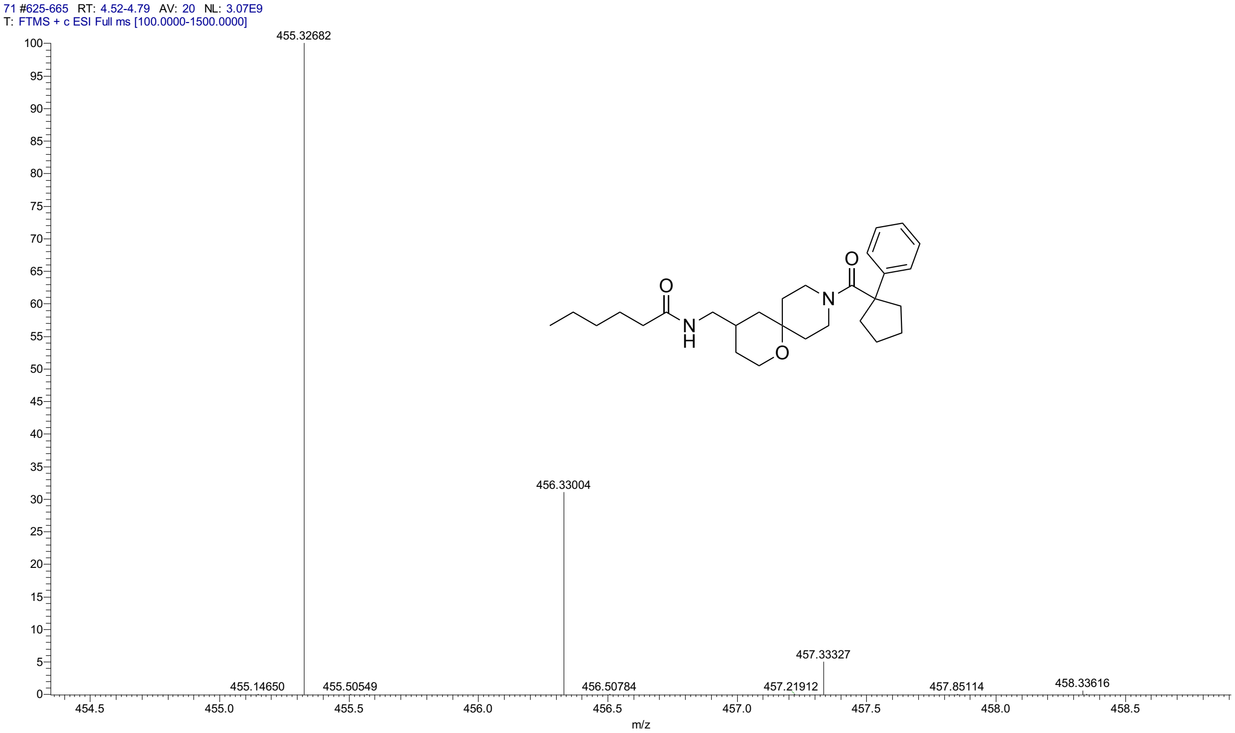


HPLC for **3b**

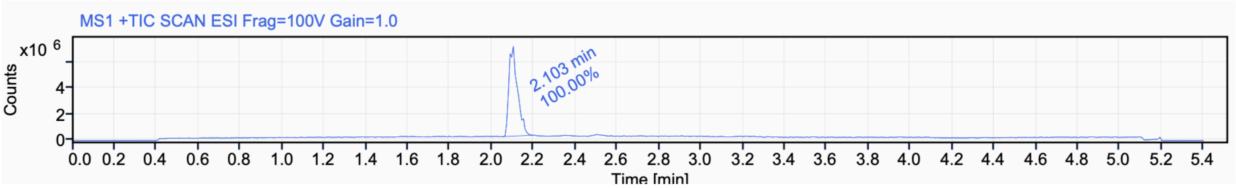


Signal Name MS1 +TIC SCAN ESI Frag=100V Gain=1.0

| RT (min) | Signal description | Area | Area% | Peak Resolution USP |
|---|---|---|---|---|
| 2.103 | MS1 +TIC SCAN ESI Frag=100V Gain=1.0 | 19793639.7 | 100.00 | |

#### $^{1}$H NMR (400 MHz, $CDCl_3$) for **3c**

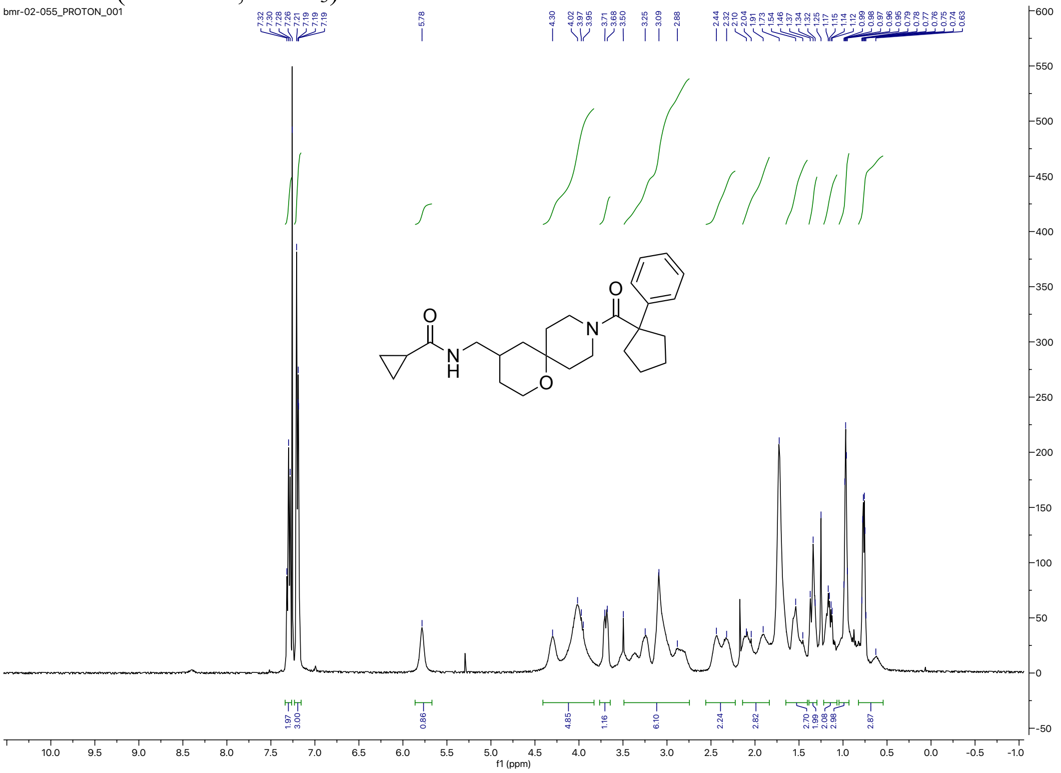


#### $^{13}$C NMR (126 MHz, $CDCl_3$) for **3c**

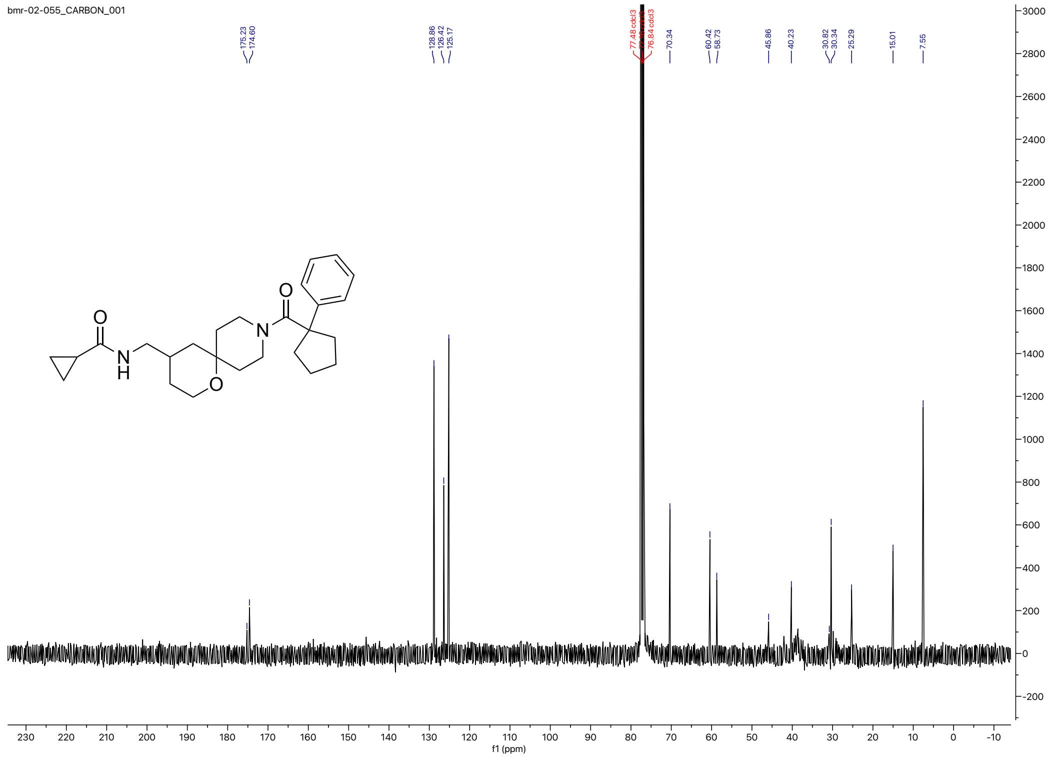

HRMS for **3c**

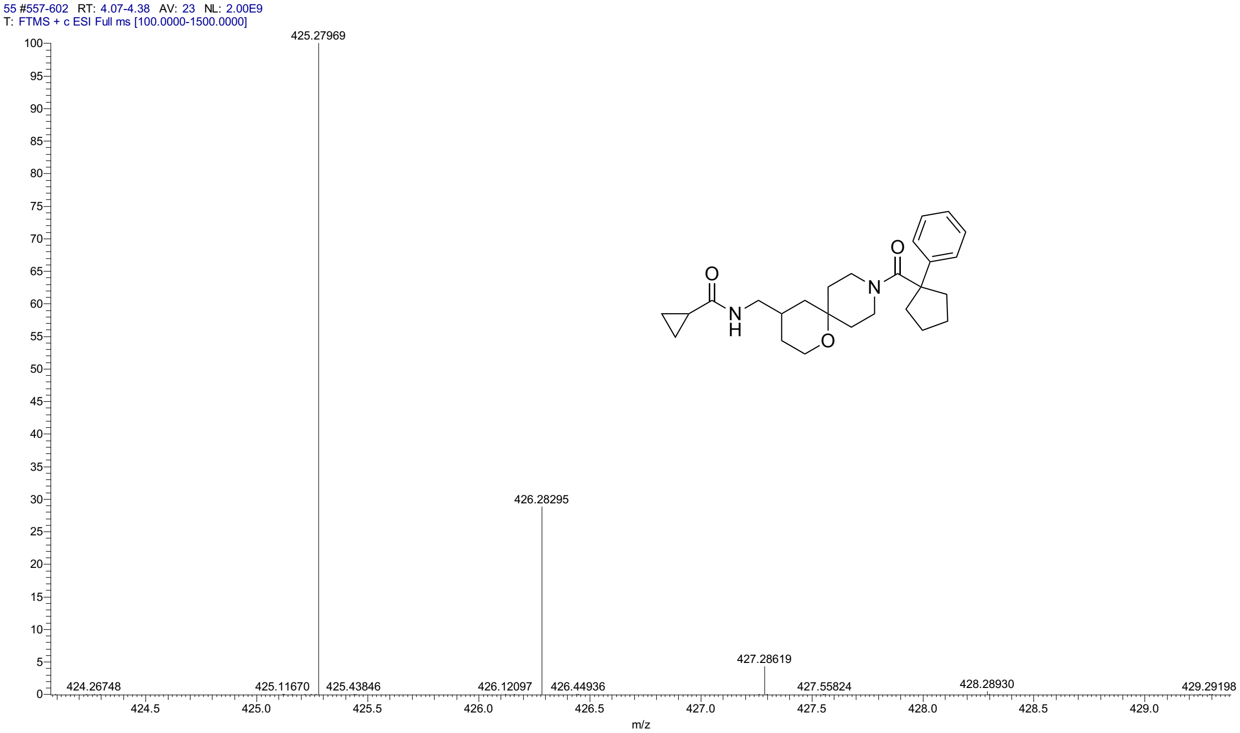


HPLC for **3c**

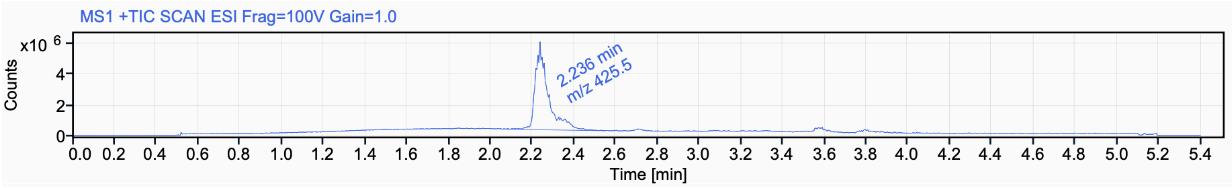


Signal Name MS1 +TIC SCAN ESI Frag=100V Gain=1.0

| RT (min) | Signal description | Area | Area% | Peak Resolution USP |
| --- | --- | --- | --- | --- |
| 2.236 | MS1 +TIC SCAN ESI Frag=100V Gain=1.0 | 24258027.9 | 100.00 | |

$^{1}H$ NMR (500 MHz, $CD_3CN$) for **3d**

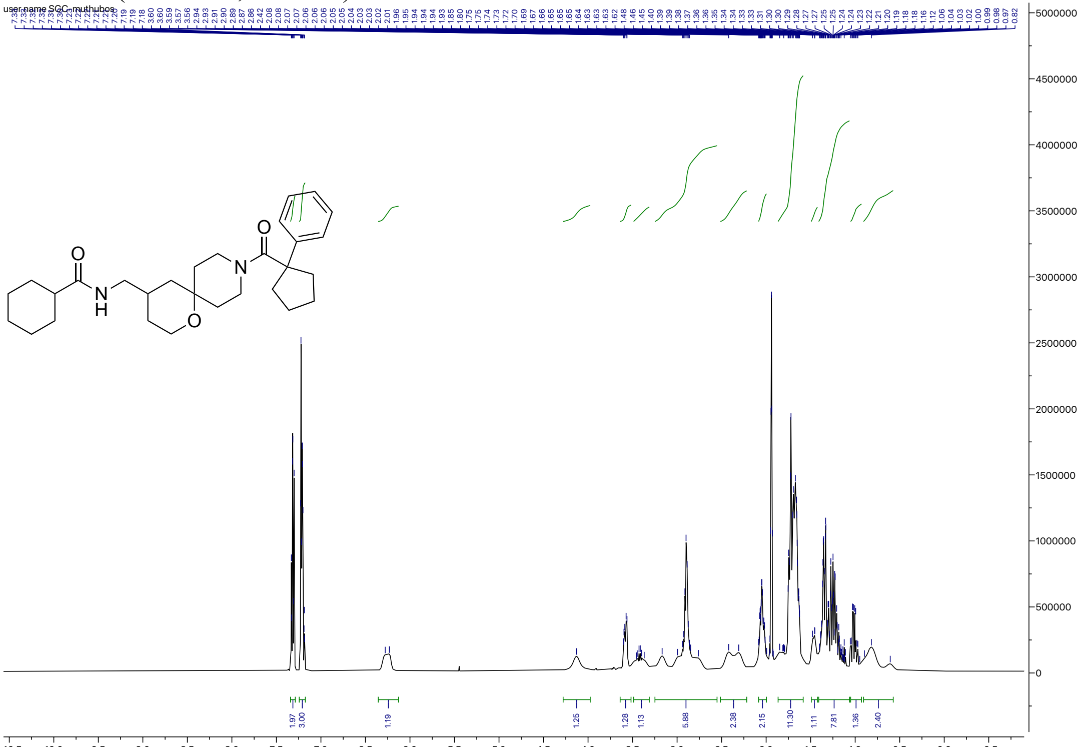

$^{13}C$ NMR (126 MHz, $CD_3CN$) for **3d**

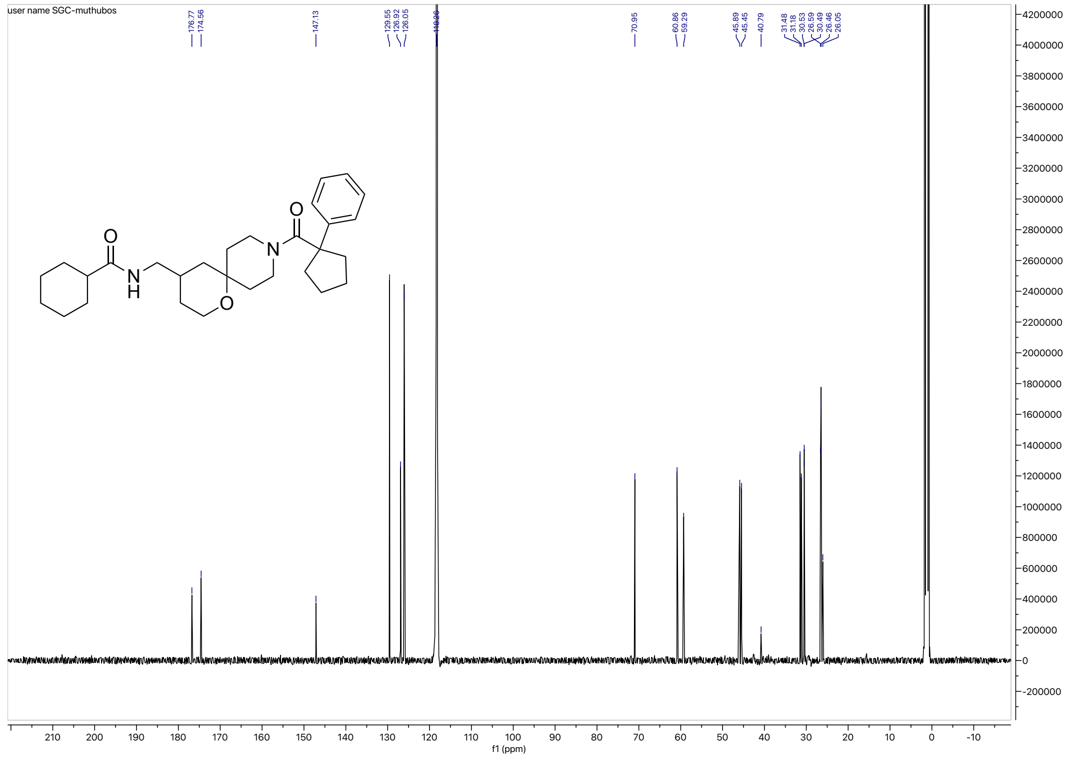

HRMS for **3d**

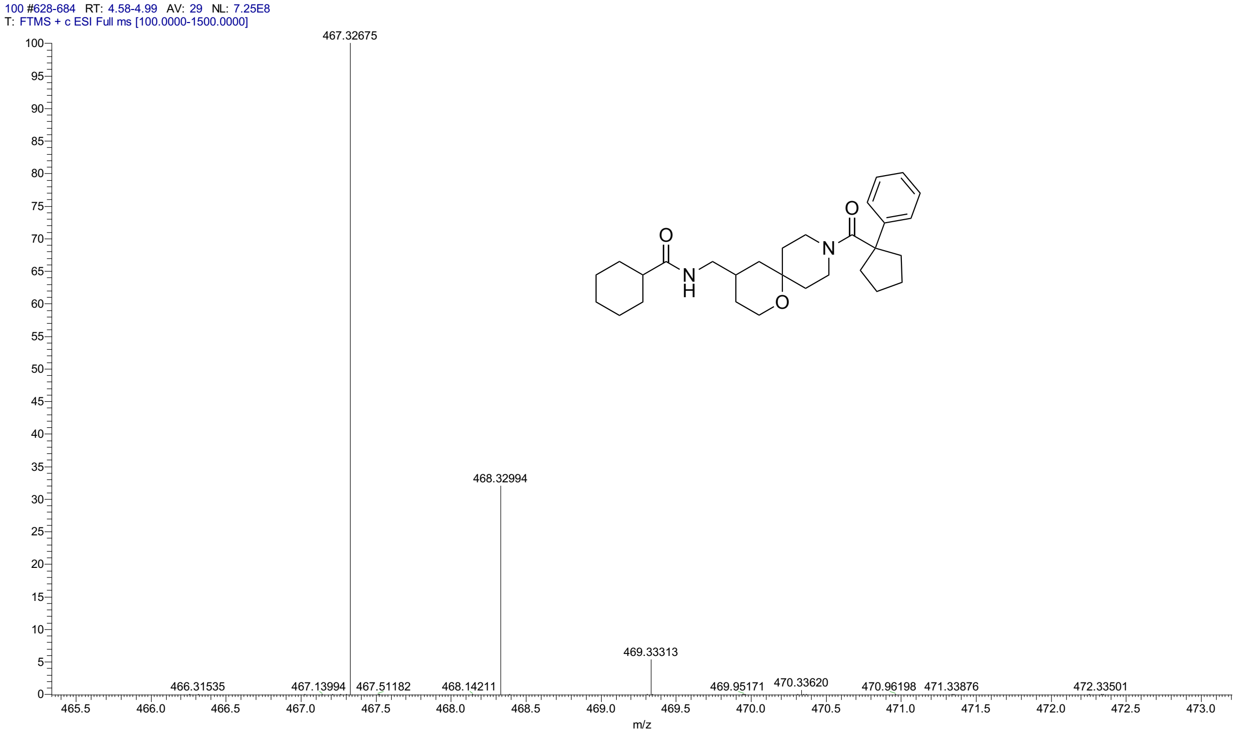


HPLC for **3d**

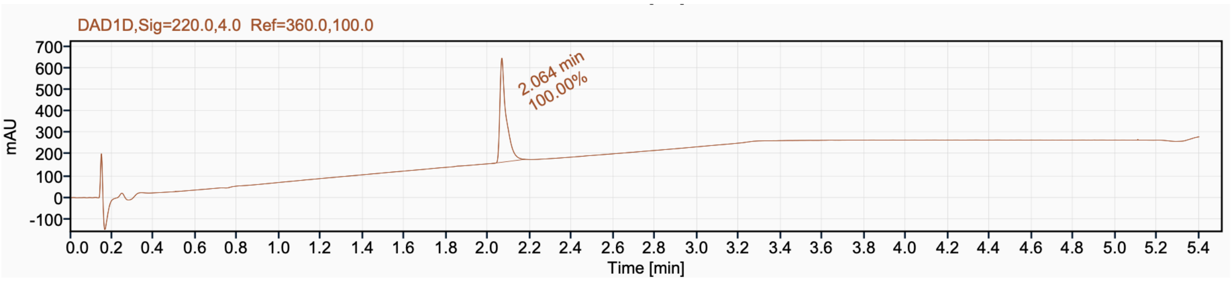

$^{1}H$ NMR (500 MHz, $CDCl_3$) for **3e**

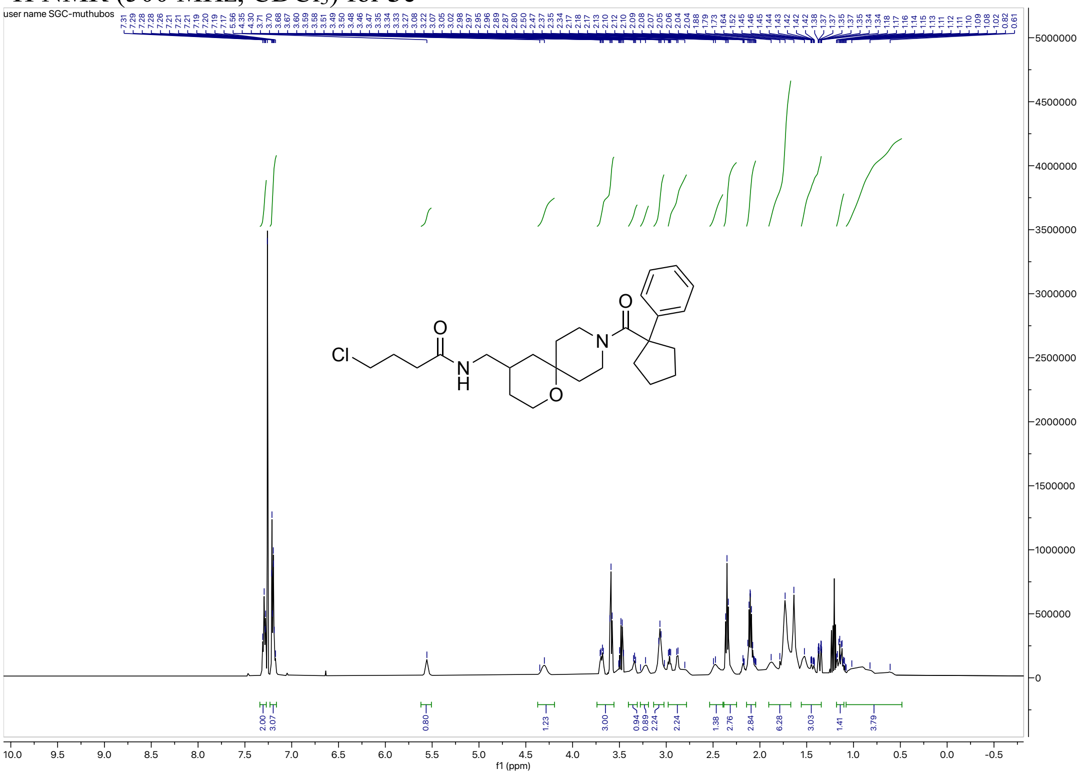


$^{13}C$ NMR (126 MHz, $CDCl_3$) for **3e**

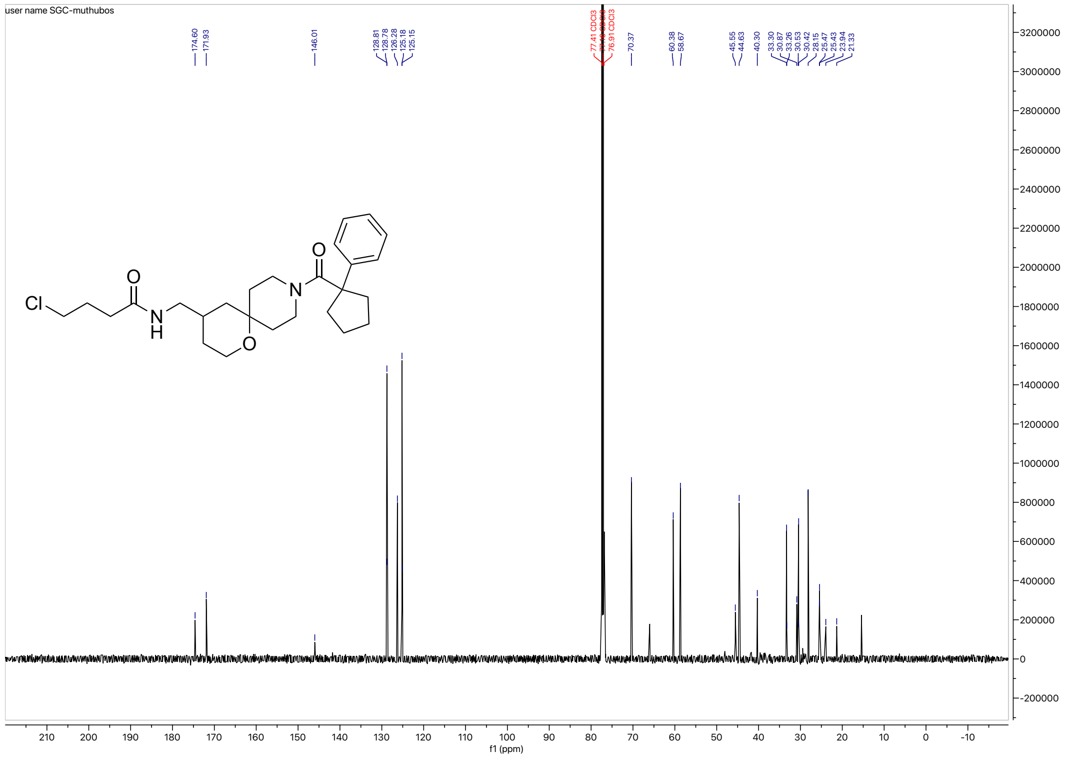

HRMS for **3e**

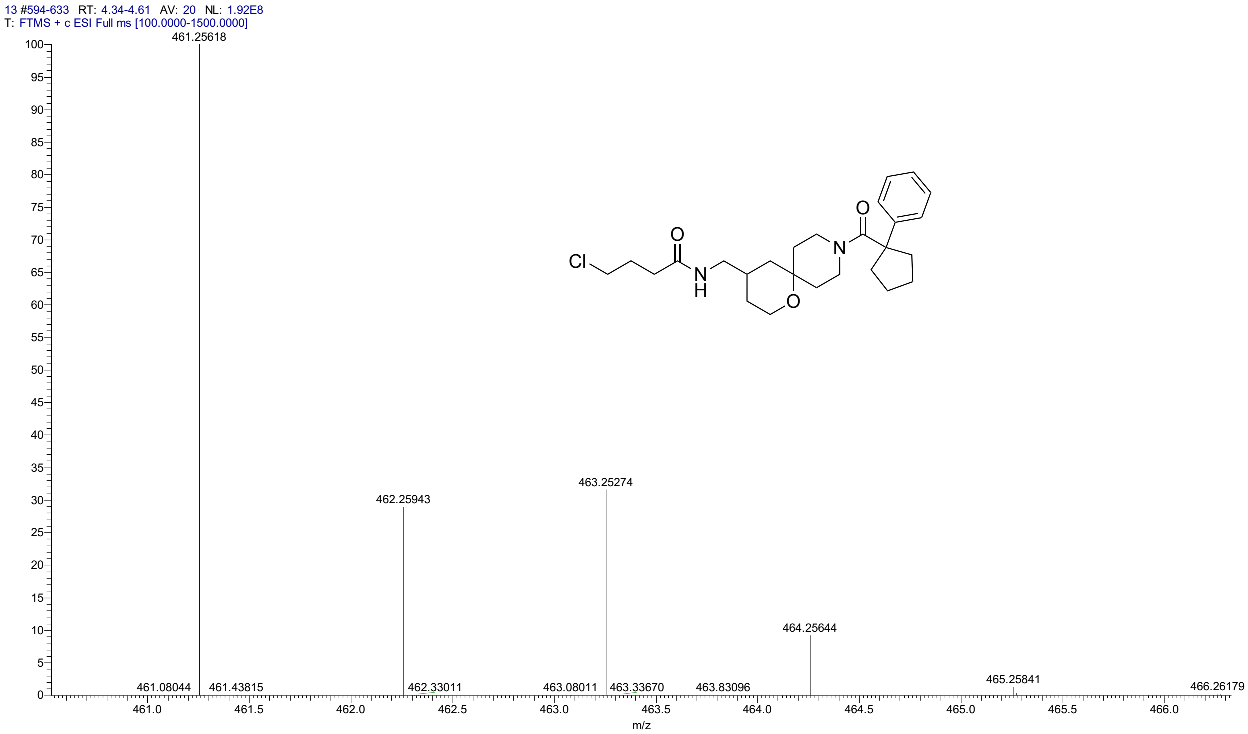


HPLC for **3e**

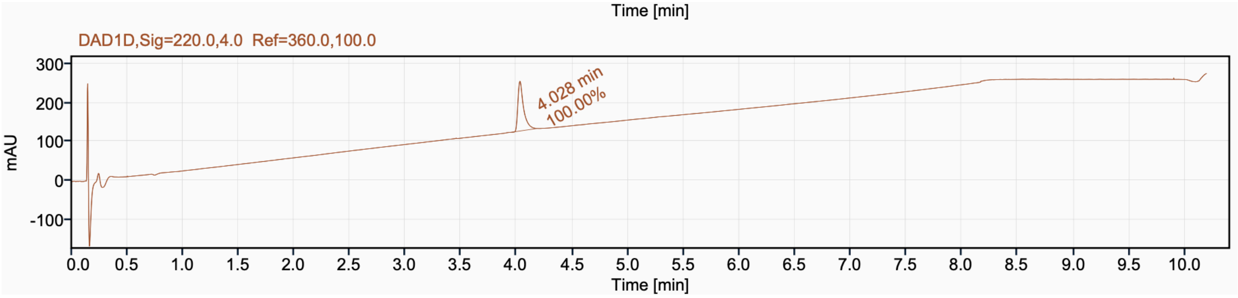


| Signal Name | DAD1D | | | |
|---|---|---|---|---|
| **RT (min)** | **Signal description** | **Area** | **Area%** | **Peak Resolution USP** |
| 4.028 | DAD1D,Sig=220.0,4.0 Ref=360.0,100.0 | 448.4 | 100.00 | |

[1]H NMR (500 MHz, $CD_3CN$) for **3f**

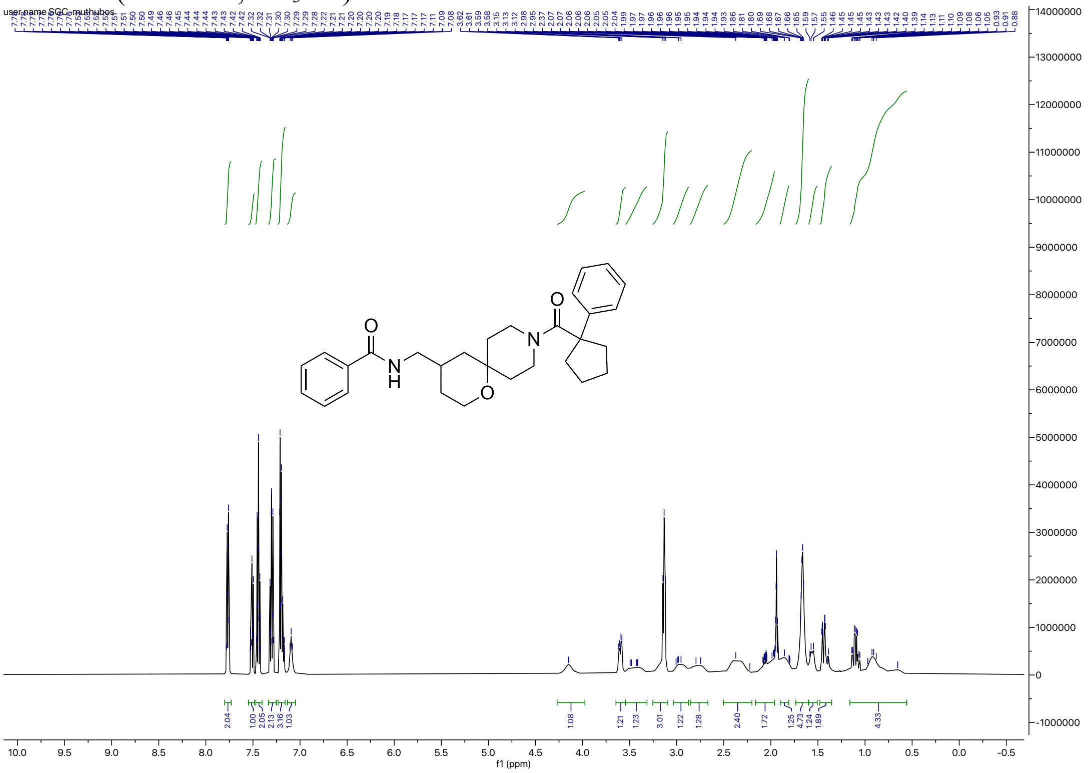

[13]C NMR (126 MHz, $CD_3CN$) for **3f**

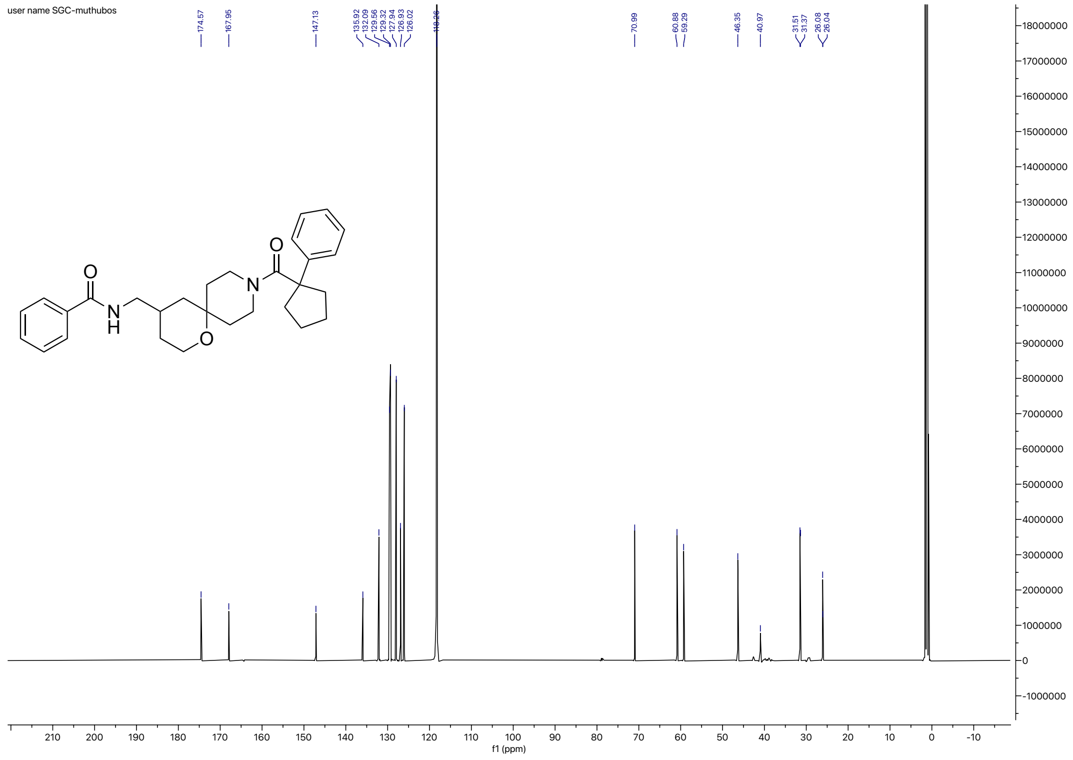

HRMS for **3f**

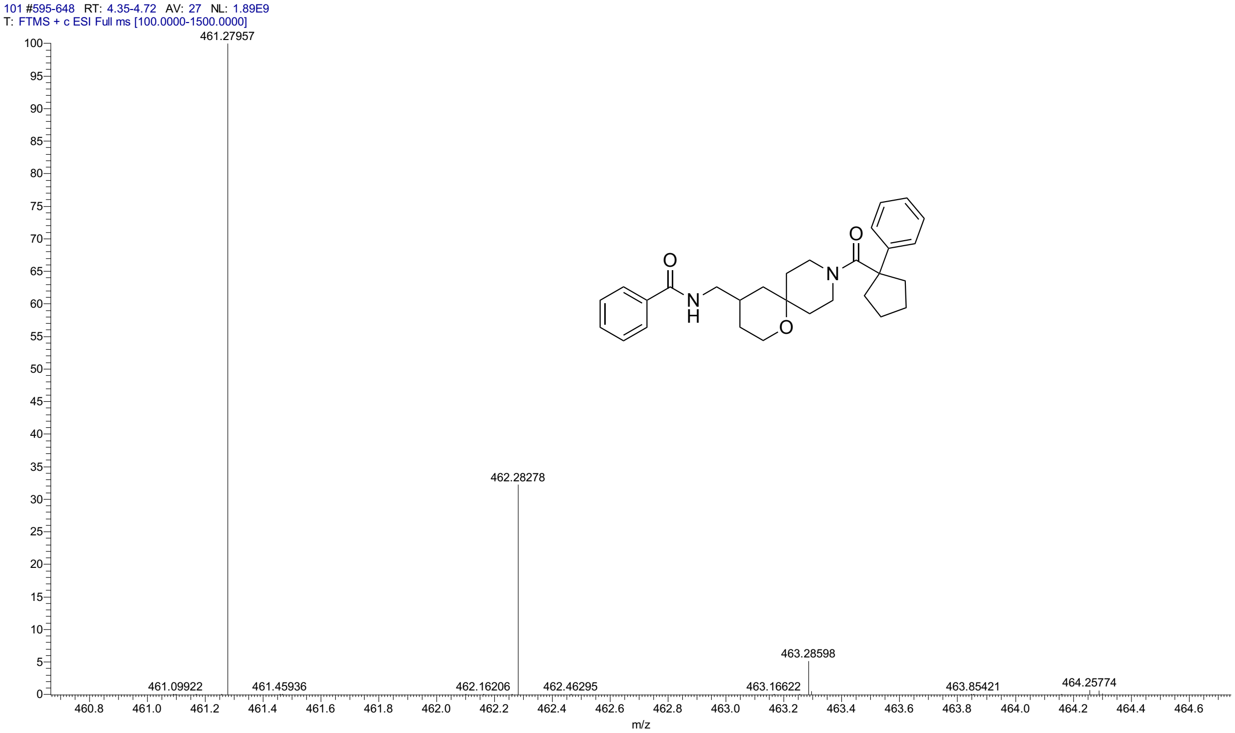


HPLC for **3f**

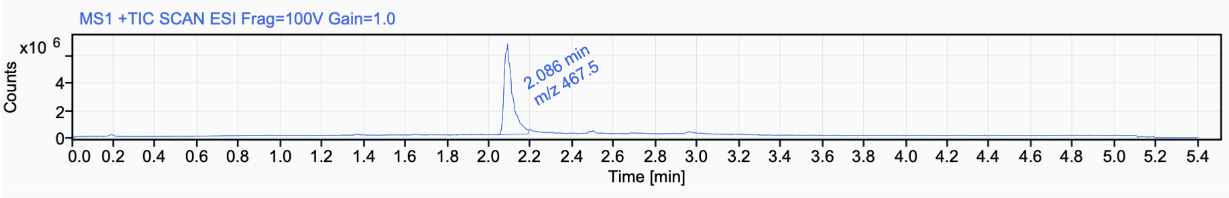


Signal Name MS1 +TIC SCAN ESI Frag=100V Gain=1.0

| RT (min) | Signal description | m/z | Purity UV(‰) / MS(%) | Area | Area% |
|---|---|---|---|---|---|
| 2.086 | MS1 +TIC SCAN ESI Frag=100V Gain=1.0 | 467.5 | | 18701243.4 | 100.00 |

$^{1}$H NMR (500 MHz, $CD_3CN$) for **3g**

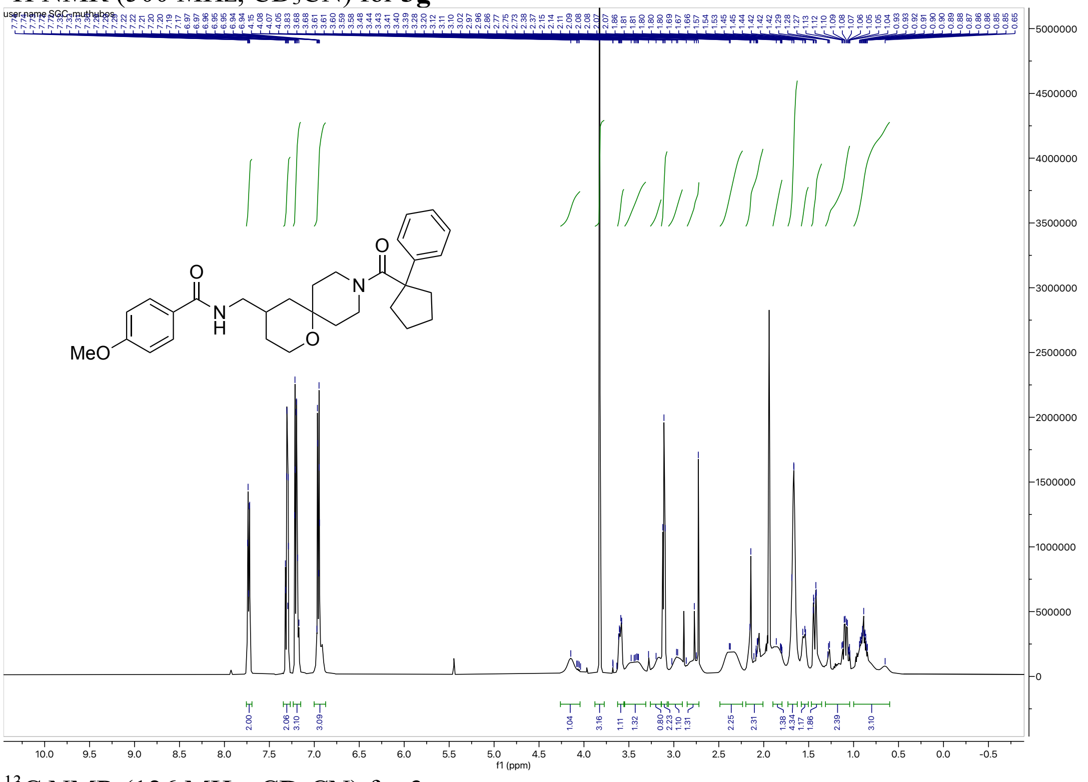

$^{13}$C NMR (126 MHz, $CD_3CN$) for **3g**

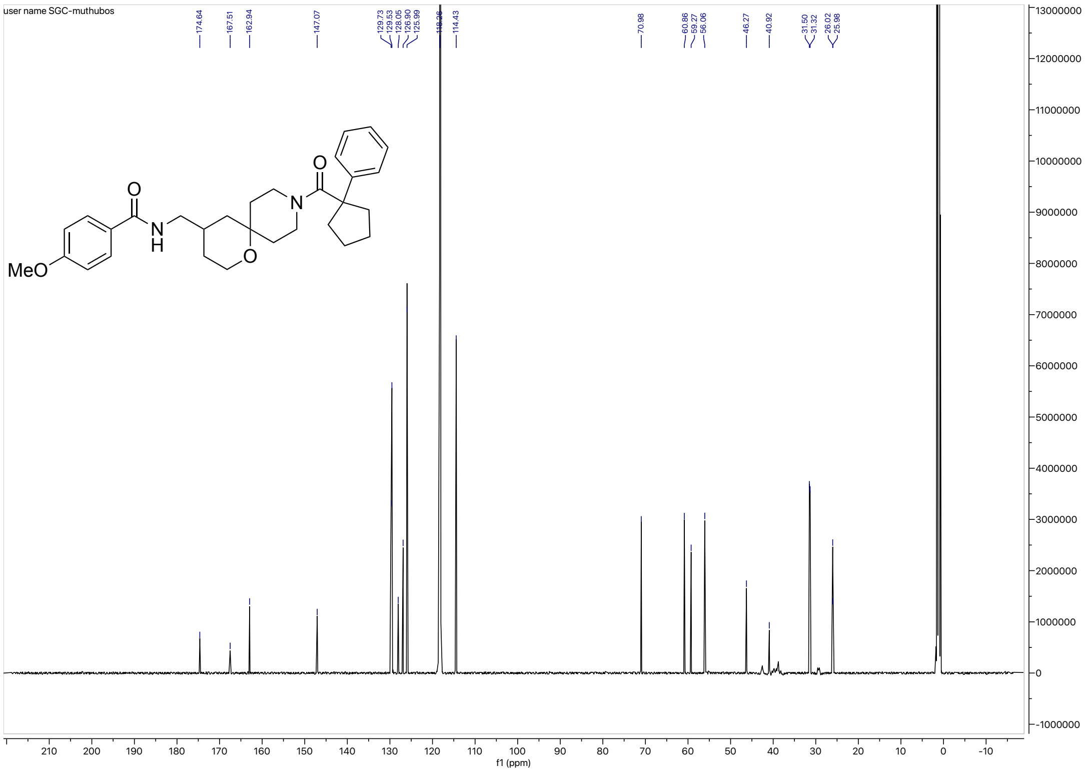

HRMS for **3g**

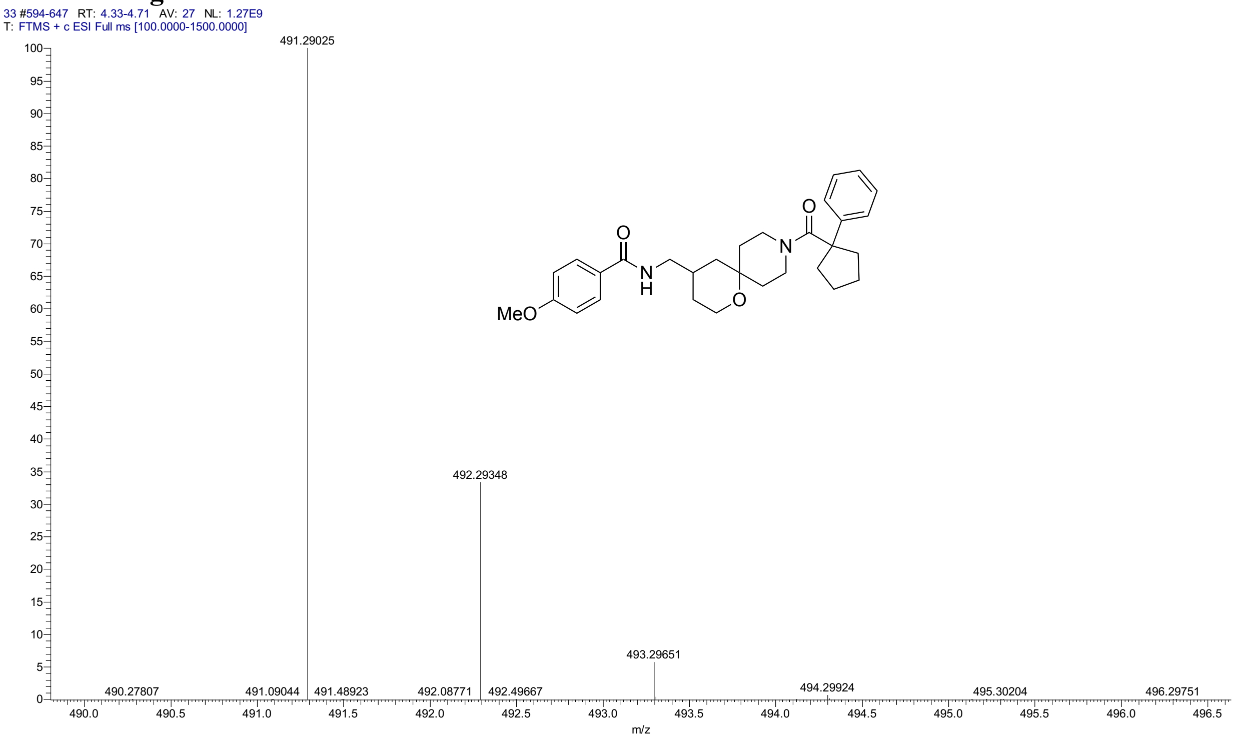


HPLC for **3g**

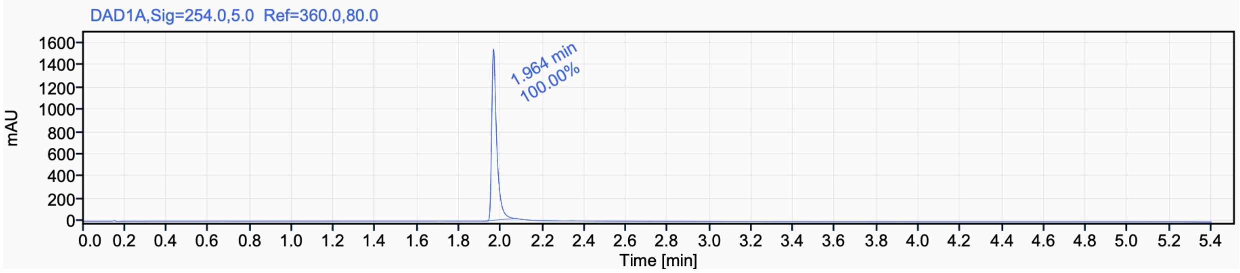


Signal Name DAD1A

| RT (min) | Signal description | Area | Area% | Peak Resolution USP |
|---|---|---|---|---|
| 1.964 | DAD1A,Sig=254.0,5.0 Ref=360.0,80.0 | 2572.7 | 100.00 | |

$^{1}$H NMR (500 MHz, $CD_3CN$) for **3h**

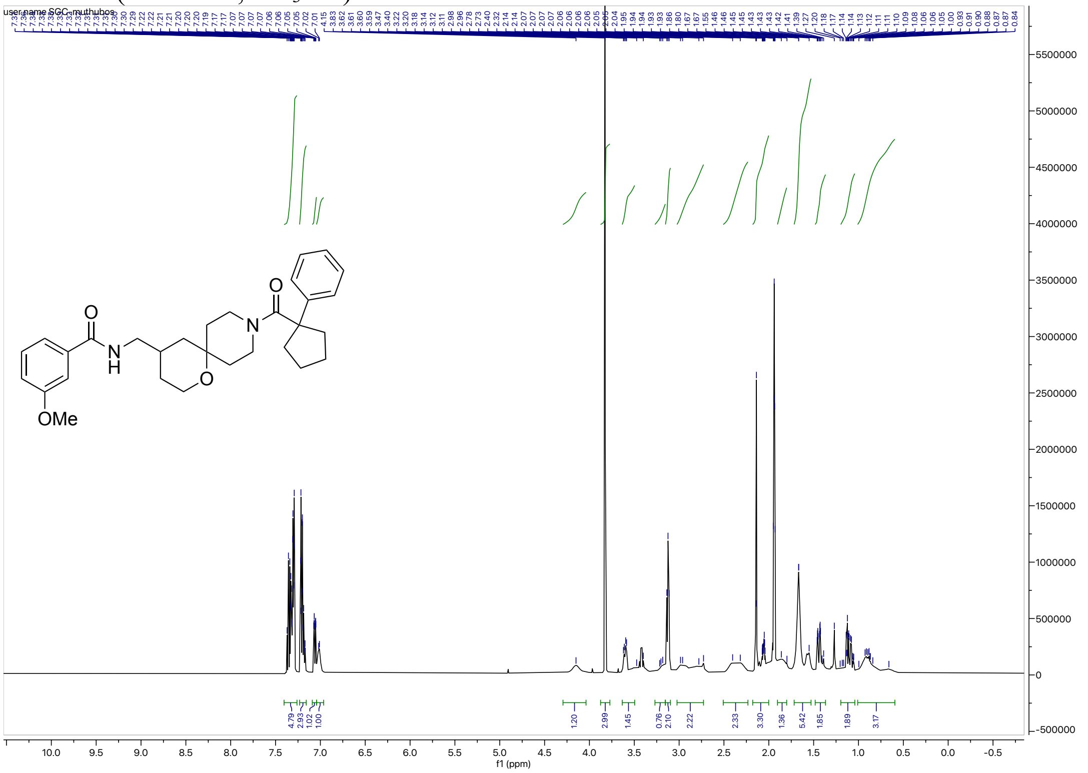

$^{13}$C NMR (126 MHz, $CD_3CN$) for **3h**

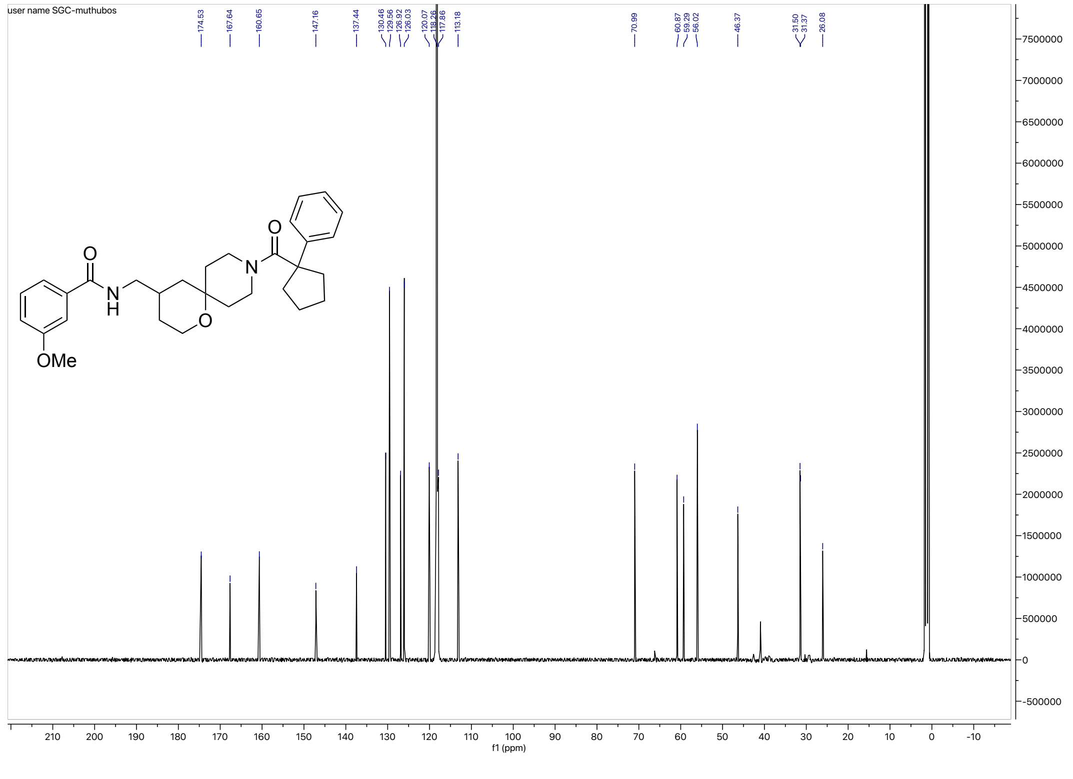

#### HRMS for **3h**

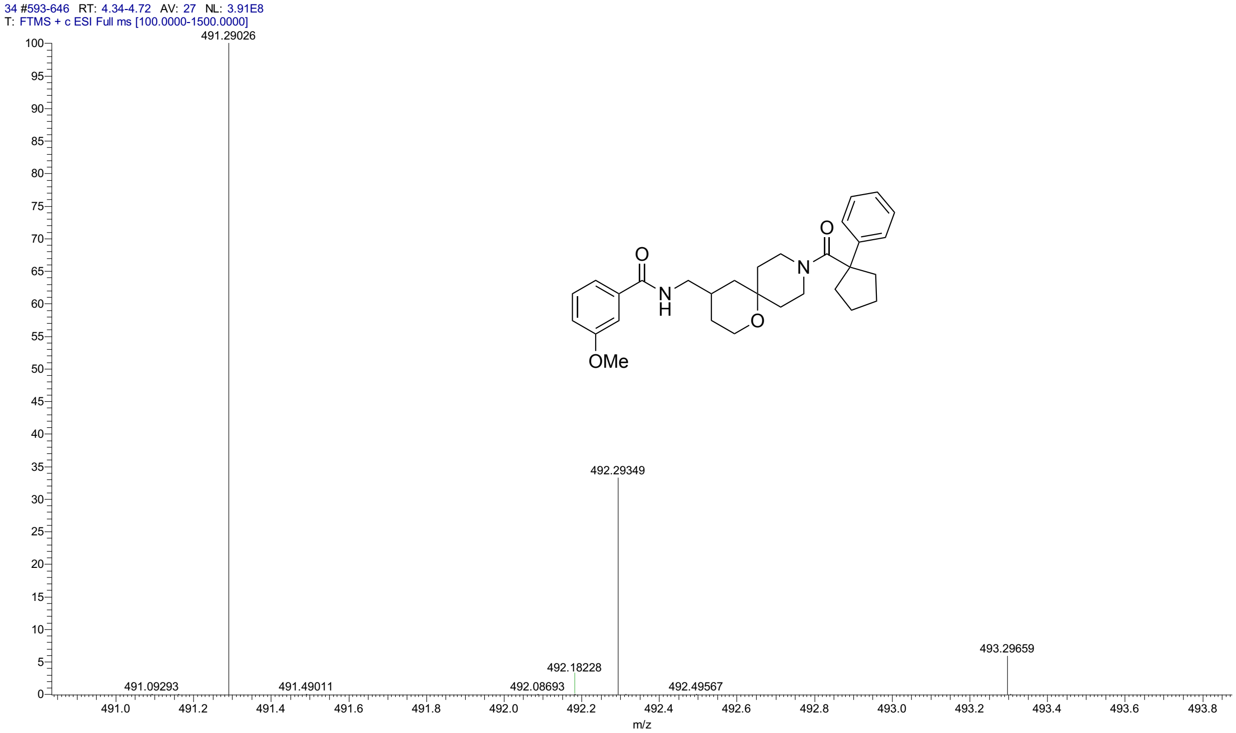


#### HPLC for **3h**

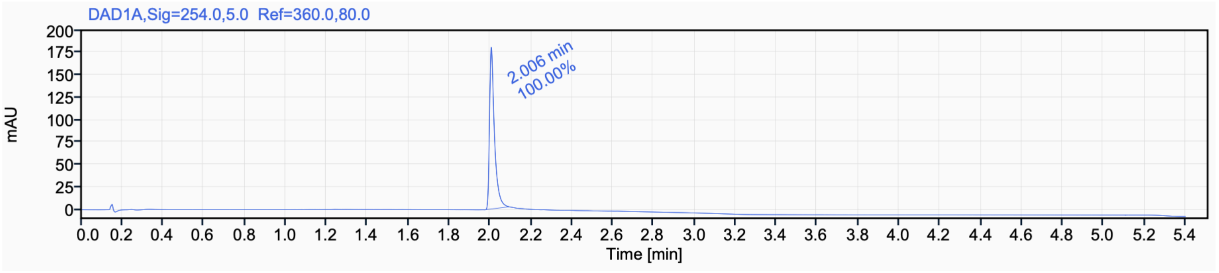


Signal Name DAD1A

| RT (min) | Signal description | Area | Area% | Peak Resolution USP |
|---|---|---|---|---|
| 2.006 | DAD1A,Sig=254.0,5.0 Ref=360.0,80.0 | 303.6 | 100.00 | |

$^{1}H$ NMR (500 MHz, $CDCl_3$) for **3i**

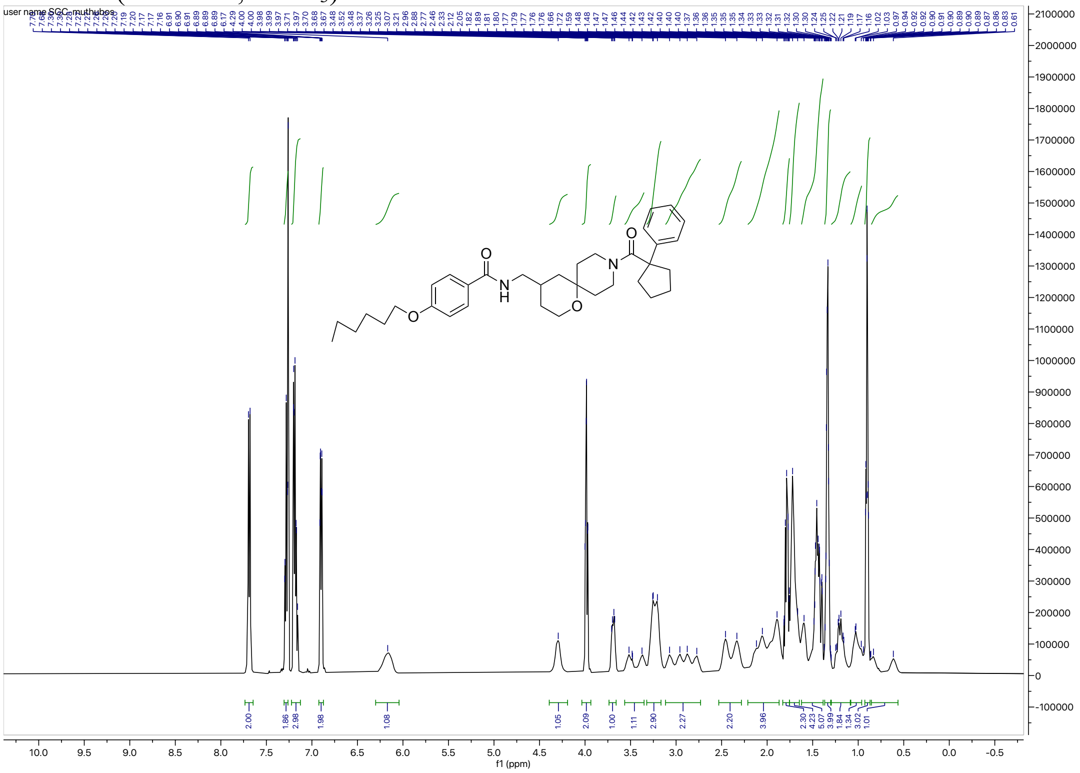


$^{13}C$ NMR (126 MHz, $CDCl_3$) for **3i**

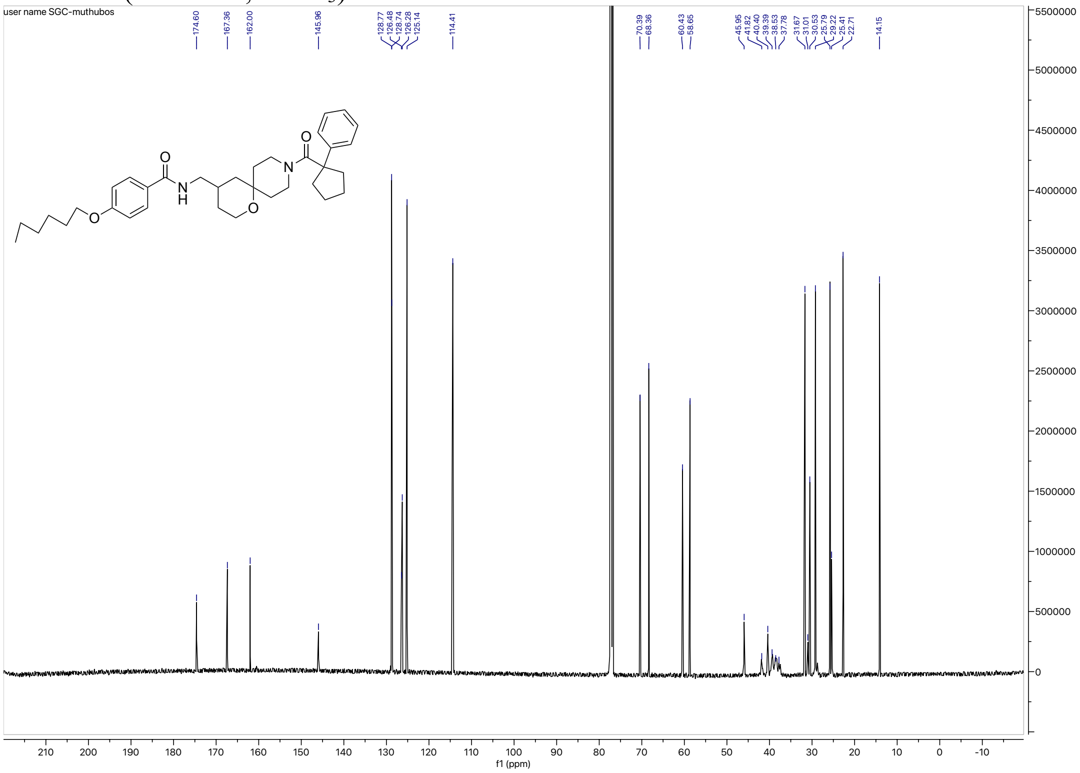

#### HRMS for **3i**

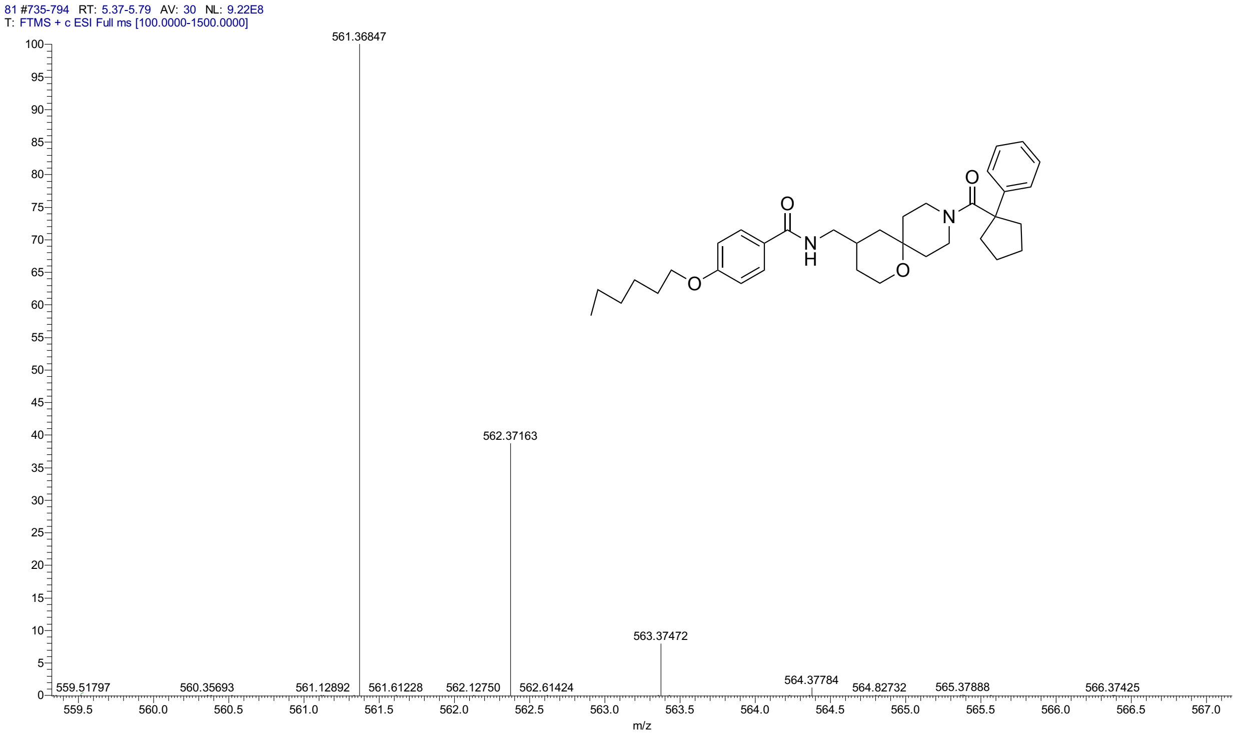


#### HPLC for **3i**

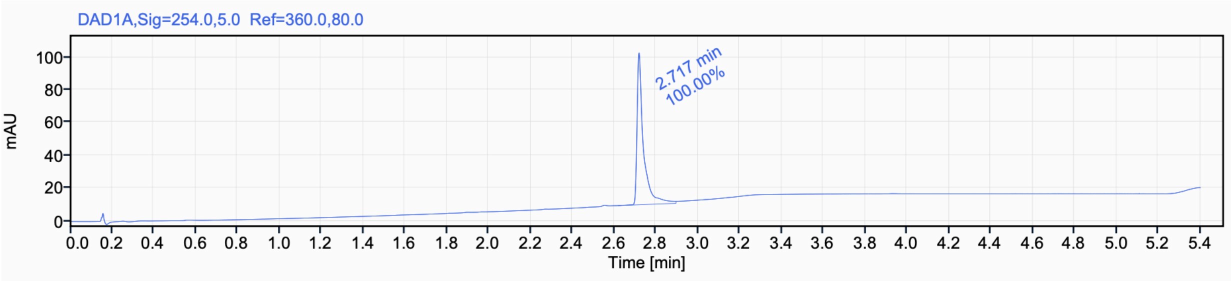


Signal Name DAD1A

| RT (min) | Signal description | m/z | Purity UV(‰) / MS(%) | Area | Area% |
|---|---|---|---|---|---|
| 2.717 | DAD1A,Sig=254.0,5.0 Ref=360.0,80.0 | | | 196.6 | 100.00 |

$^{1}$H NMR (400 MHz, CD$_3$CN) for **3j**

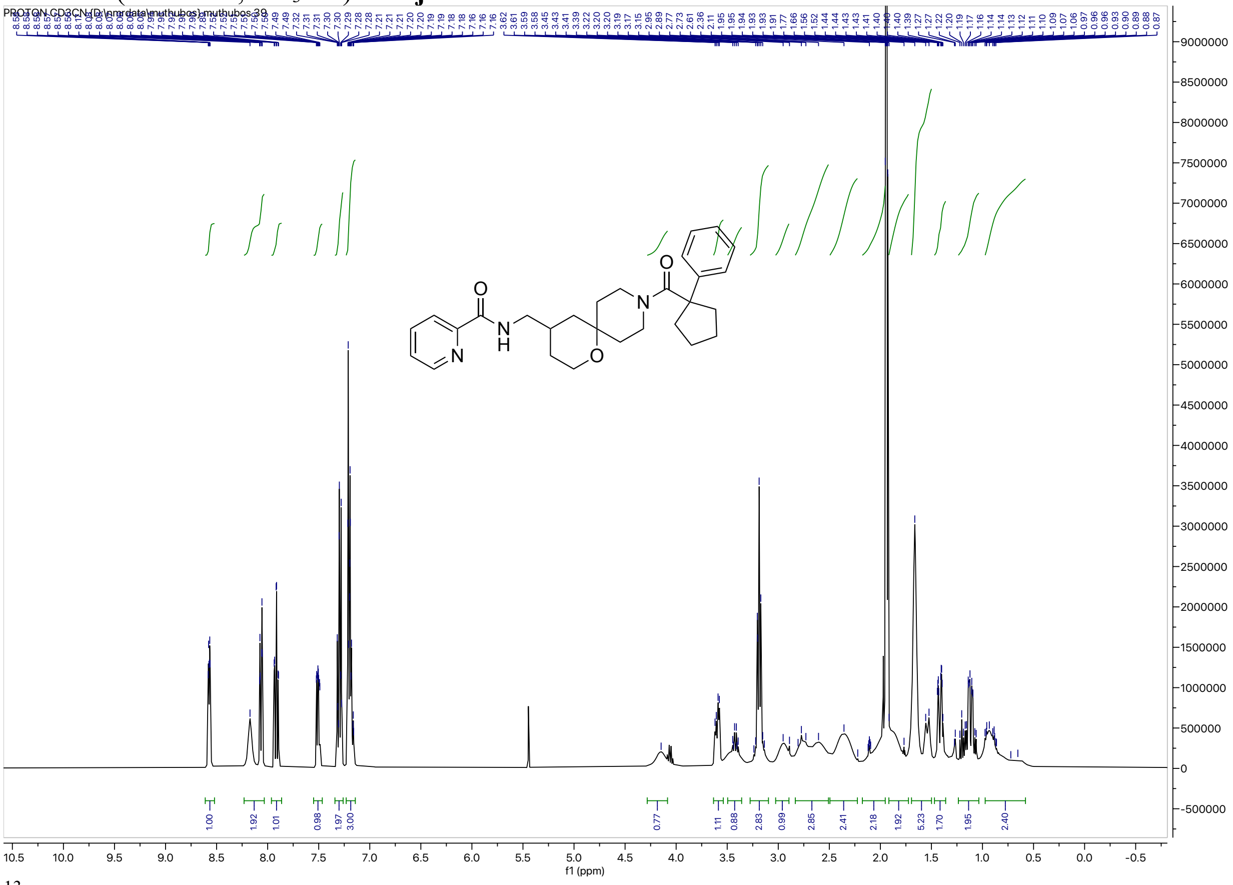

$^{13}$C NMR (126 MHz, CD$_3$CN) for **3j**

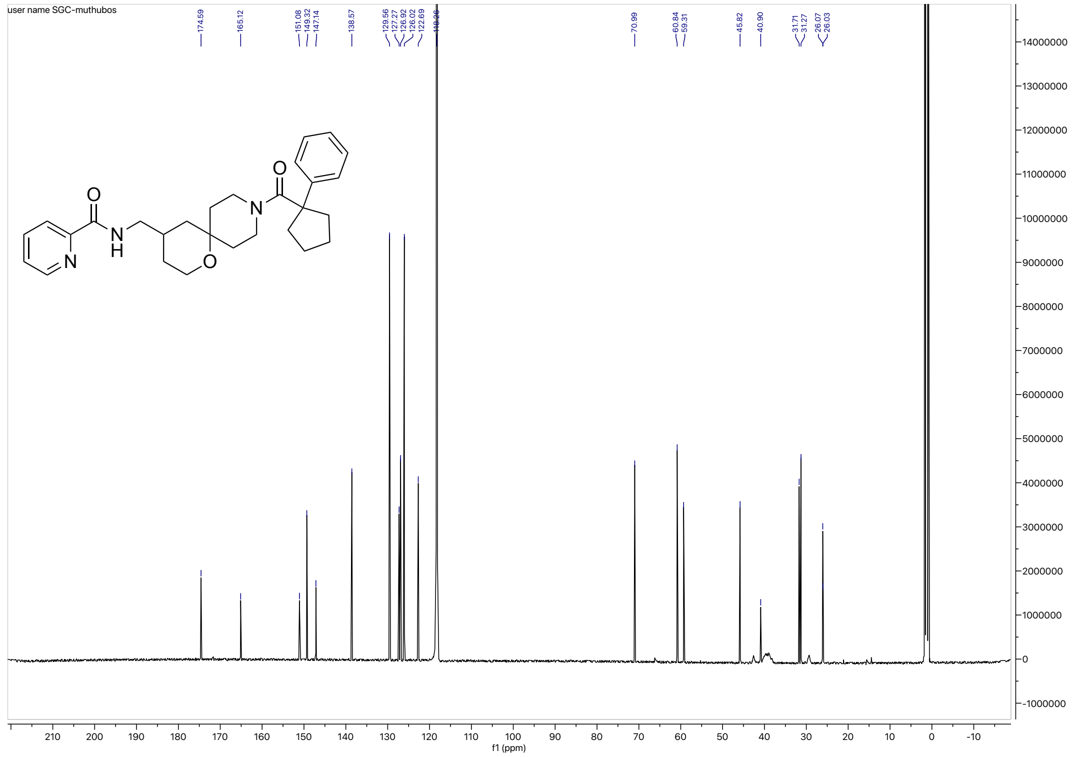

HRMS for **3j**

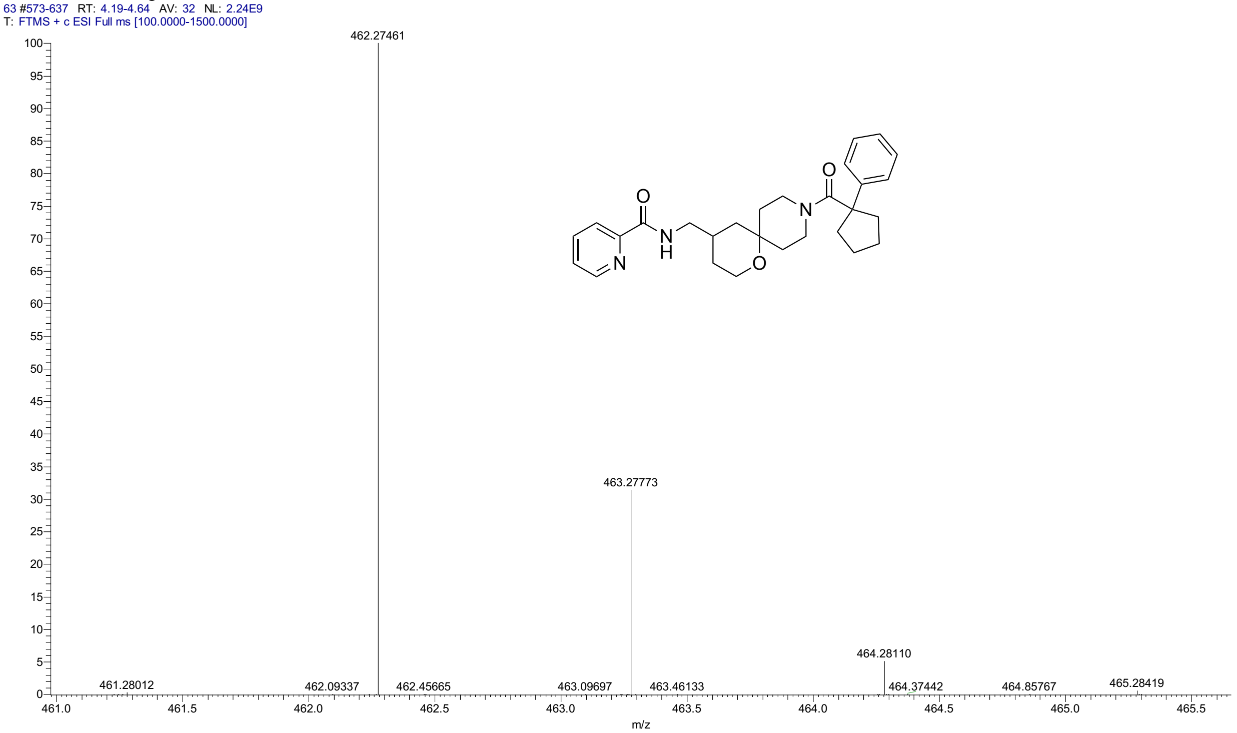


HPLC for **3j**

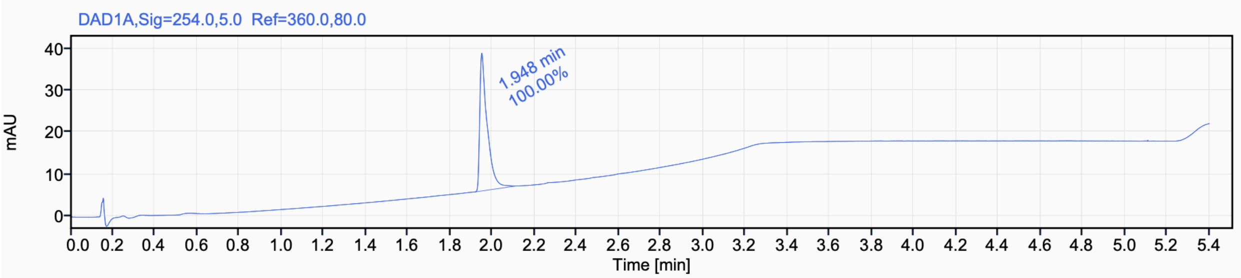


Signal Name DAD1A

| RT (min) | Signal description | m/z | Purity UV(‰) / MS(%) | Area | Area% |
|---|---|---|---|---|---|
| 1.948 | DAD1A,Sig=254.0,5.0 Ref=360.0,80.0 | | | 80.7 | 100.00 |

$^{1}$H NMR (500 MHz, $CD_3CN$) for **3k**

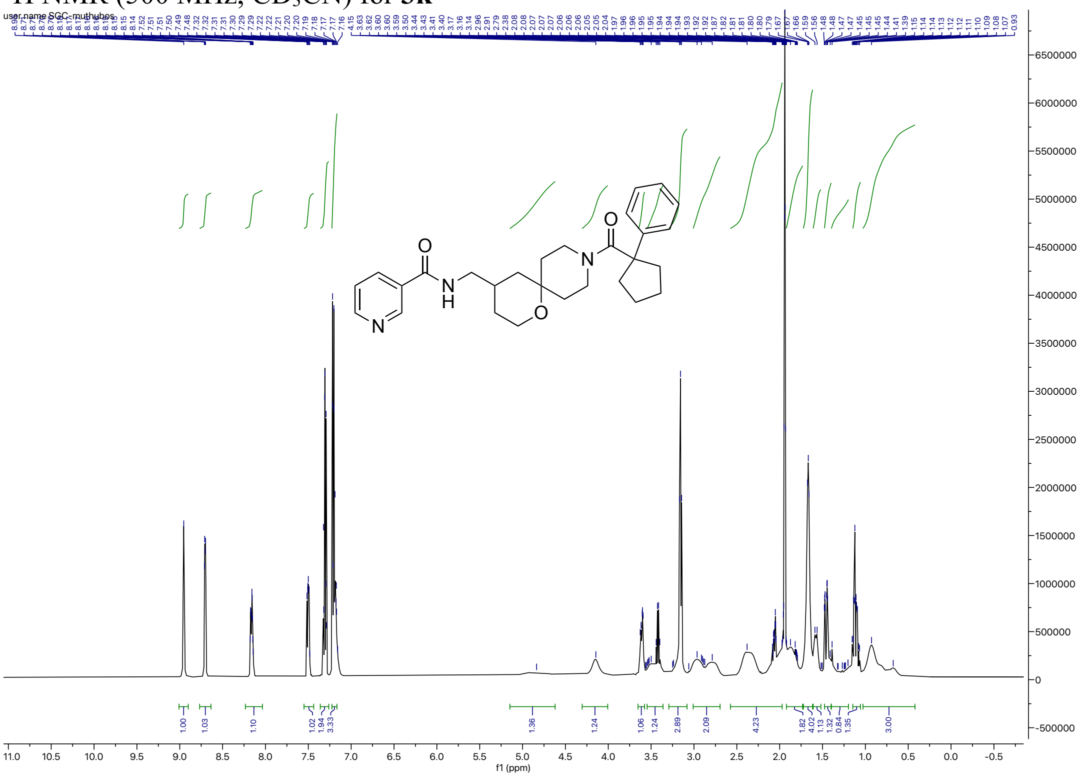


$^{13}$C NMR (126 MHz, $CD_3CN$) for **3k**

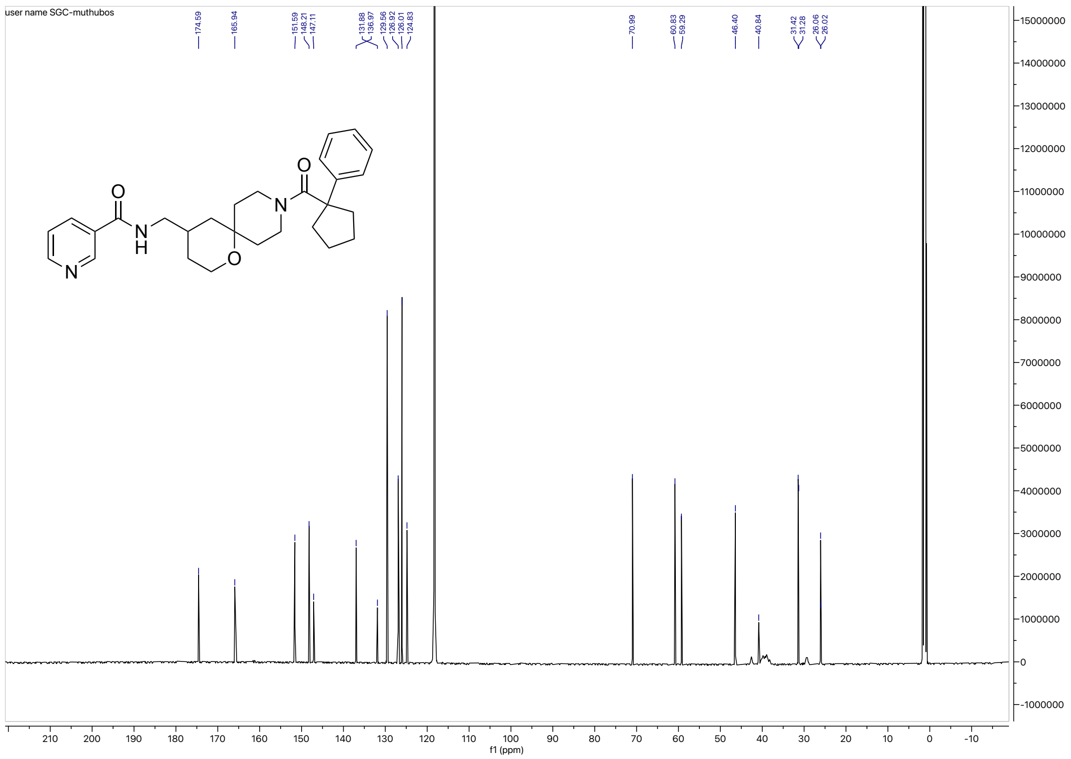

HRMS for **3k**

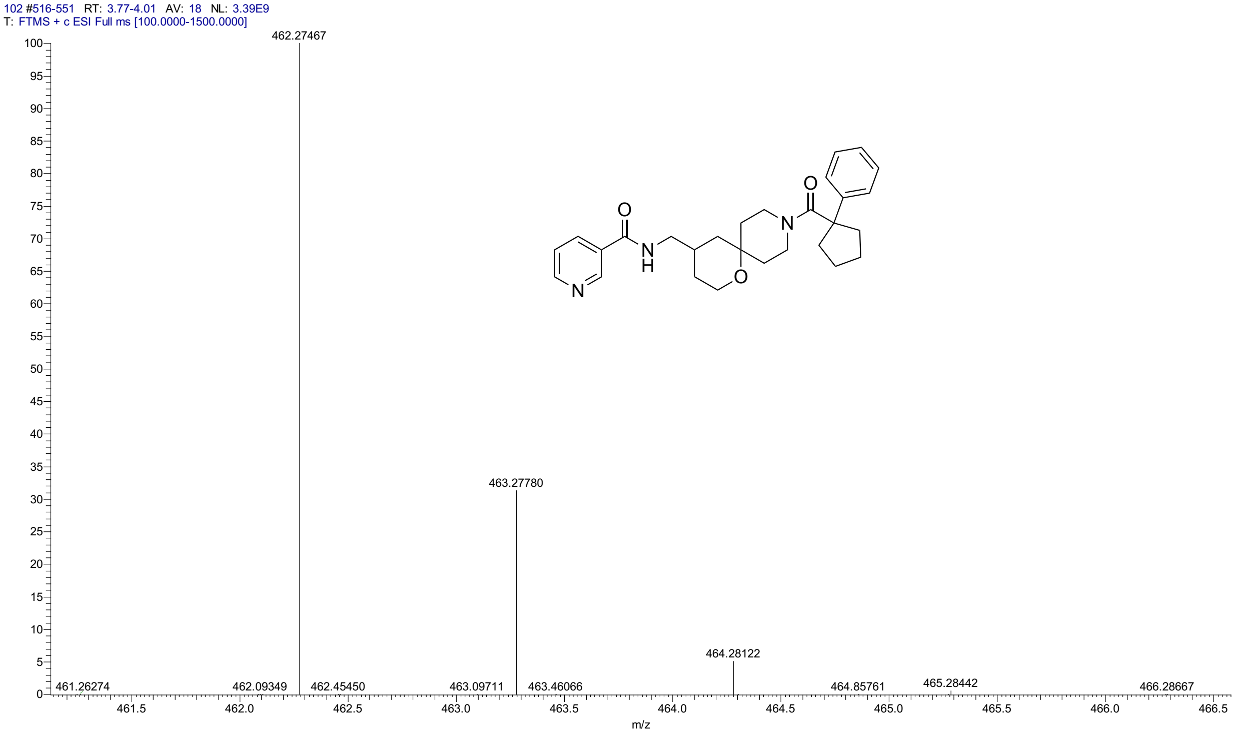


HPLC for **3k**

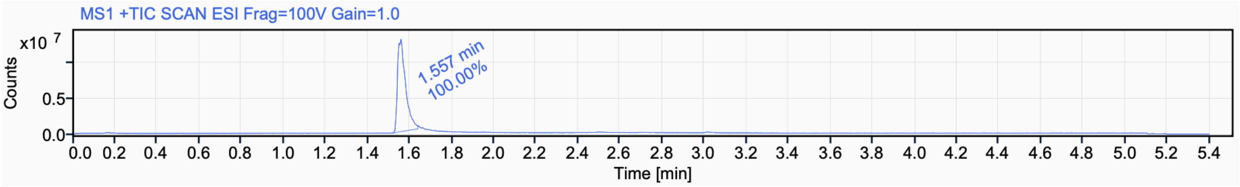

$^{1}$H NMR (500 MHz, $CD_3CN$) for **3l**

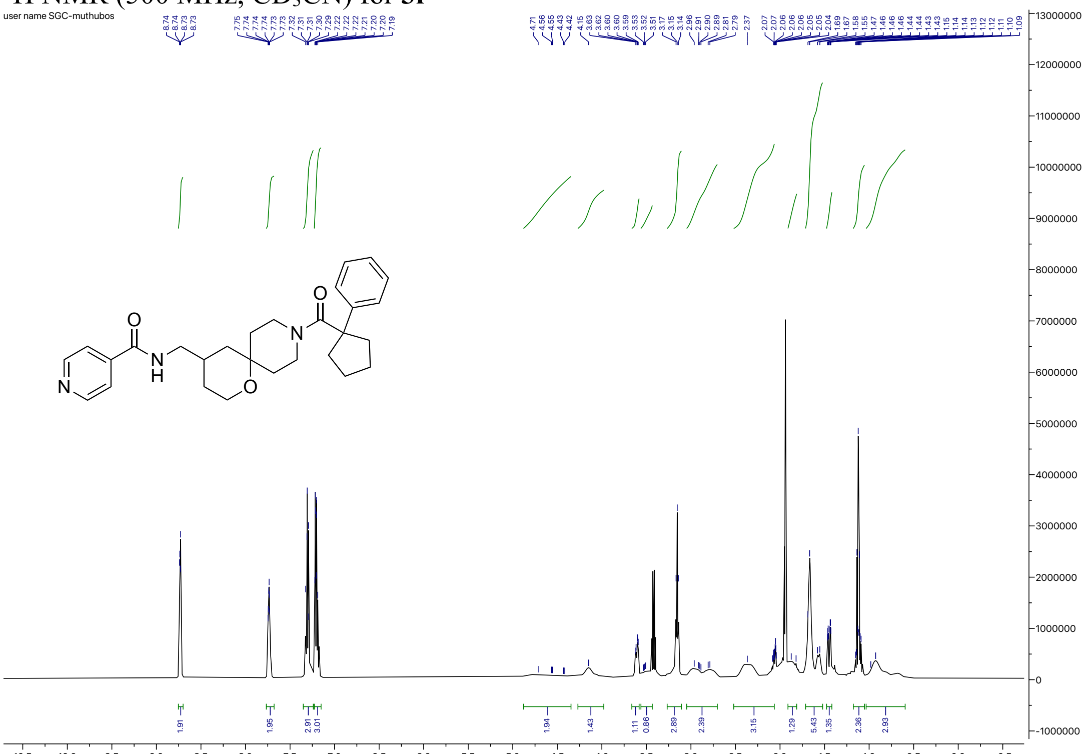


$^{13}$C NMR (126 MHz, $CD_3CN$) for **3l**

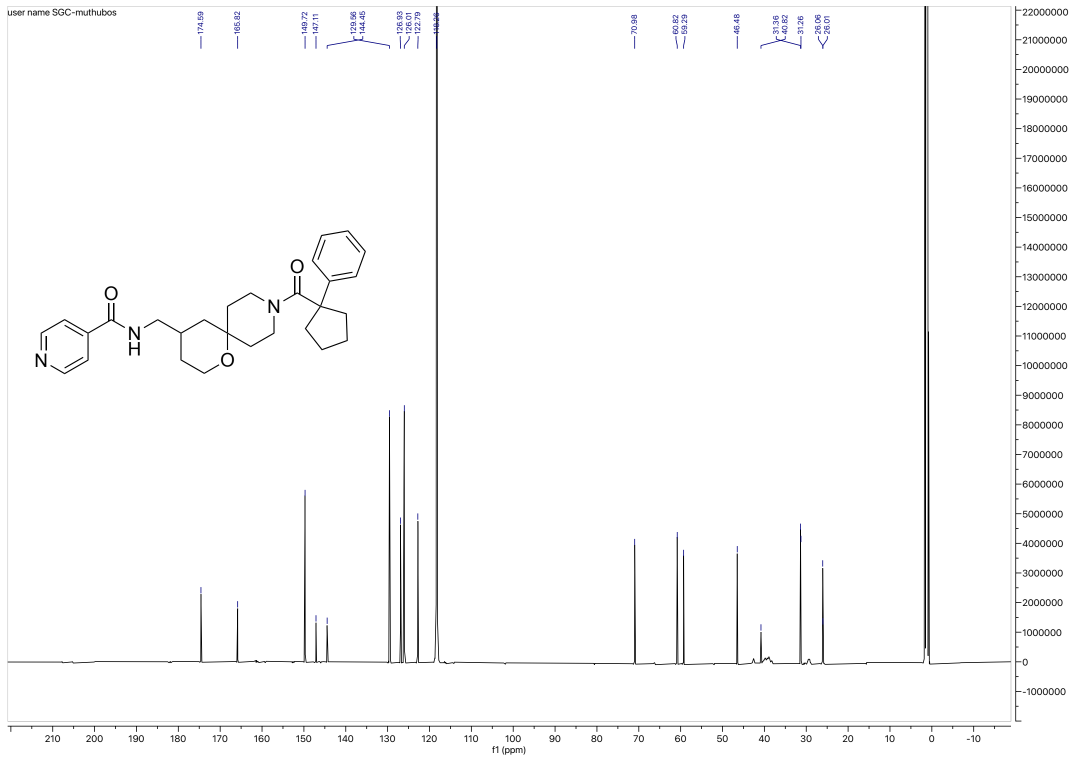

### HRMS for **3l**

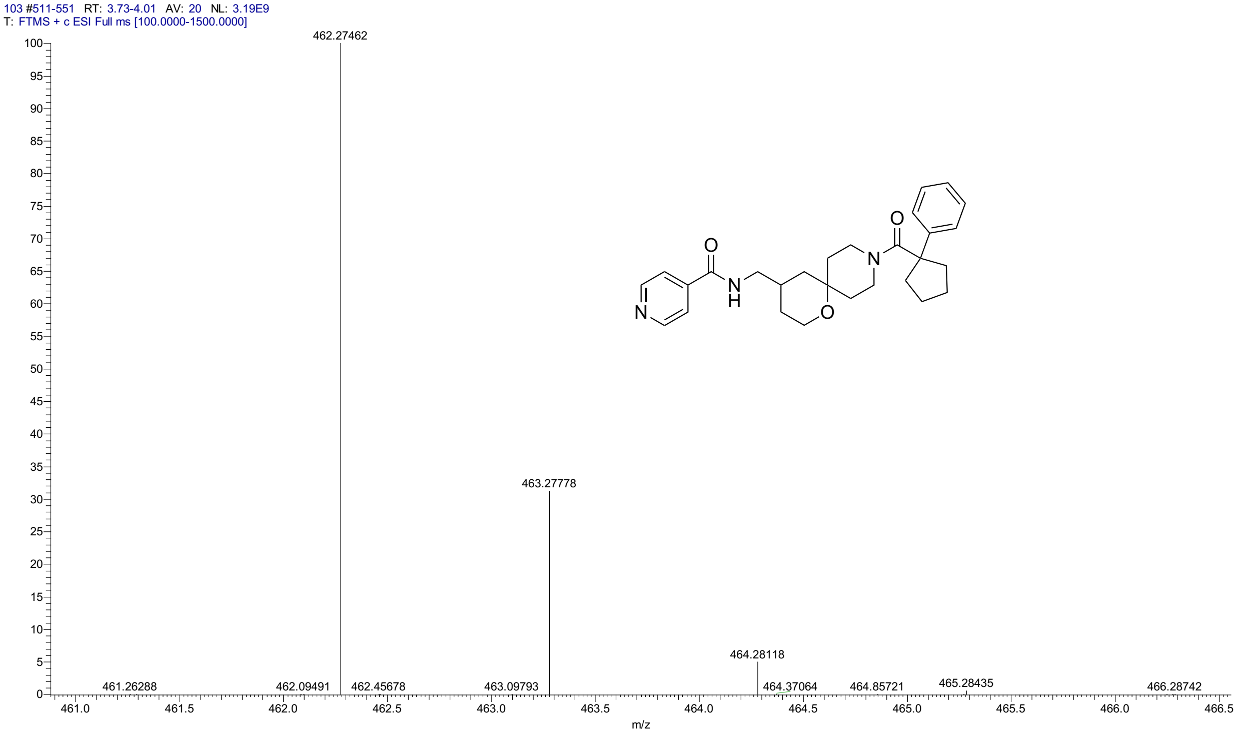


### HPLC for 3l

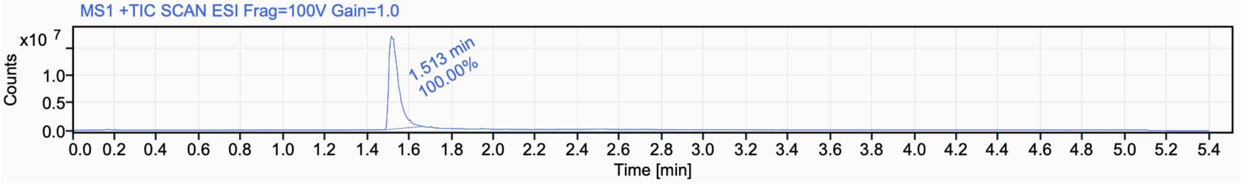


Signal Name MS1 +TIC SCAN ESI Frag=100V Gain=1.0

| RT (min) | Signal description | m/z | Purity UV(‰) / MS(%) | Area | Area% |
|---|---|---|---|---|---|
| 1.513 | MS1 +TIC SCAN ESI Frag=100V Gain=1.0 | | | 55503759.8 | 100.00 |

[1]H NMR (500 MHz, $CD_3CN$) for **3m**

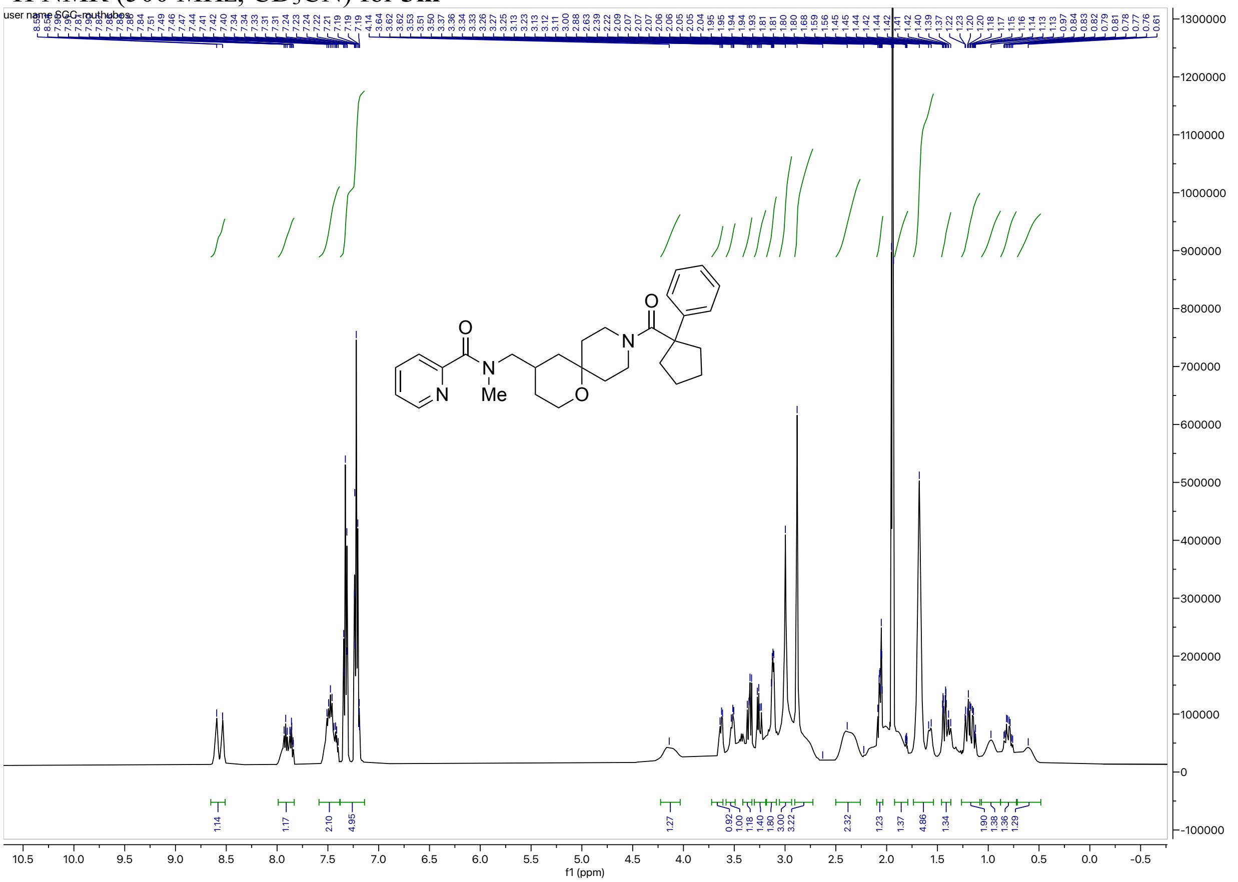


[13]C NMR (126 MHz, $CD_3CN$) for **3m**

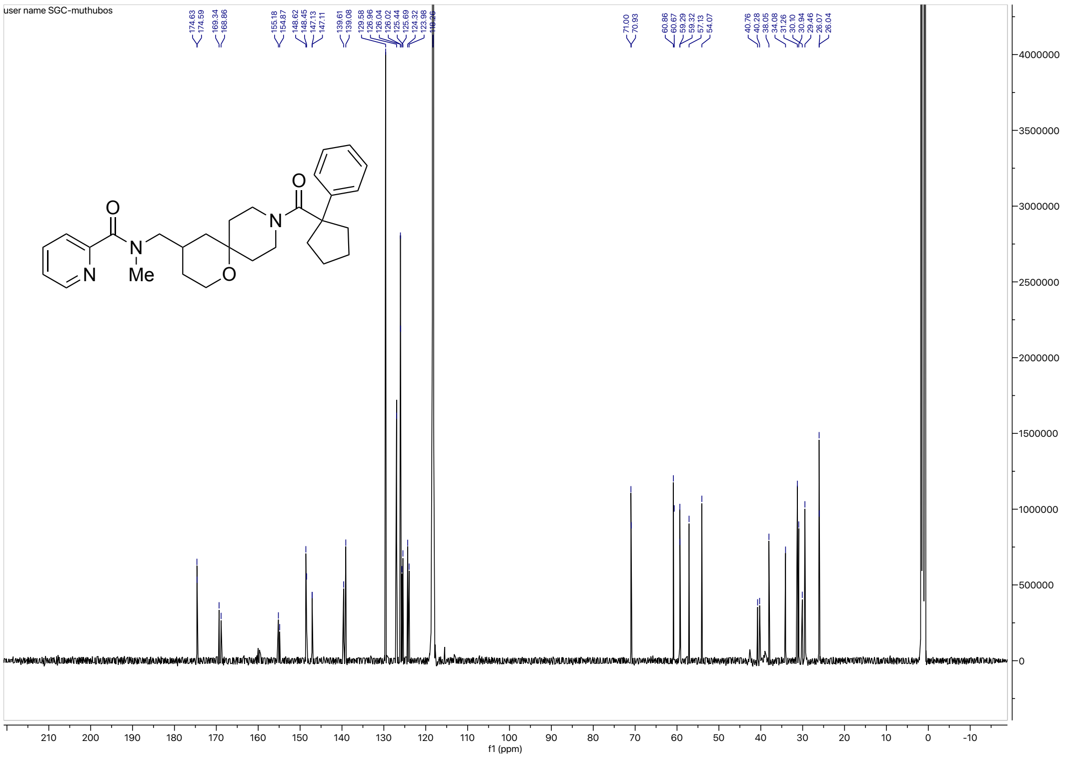

HRMS for **3m**

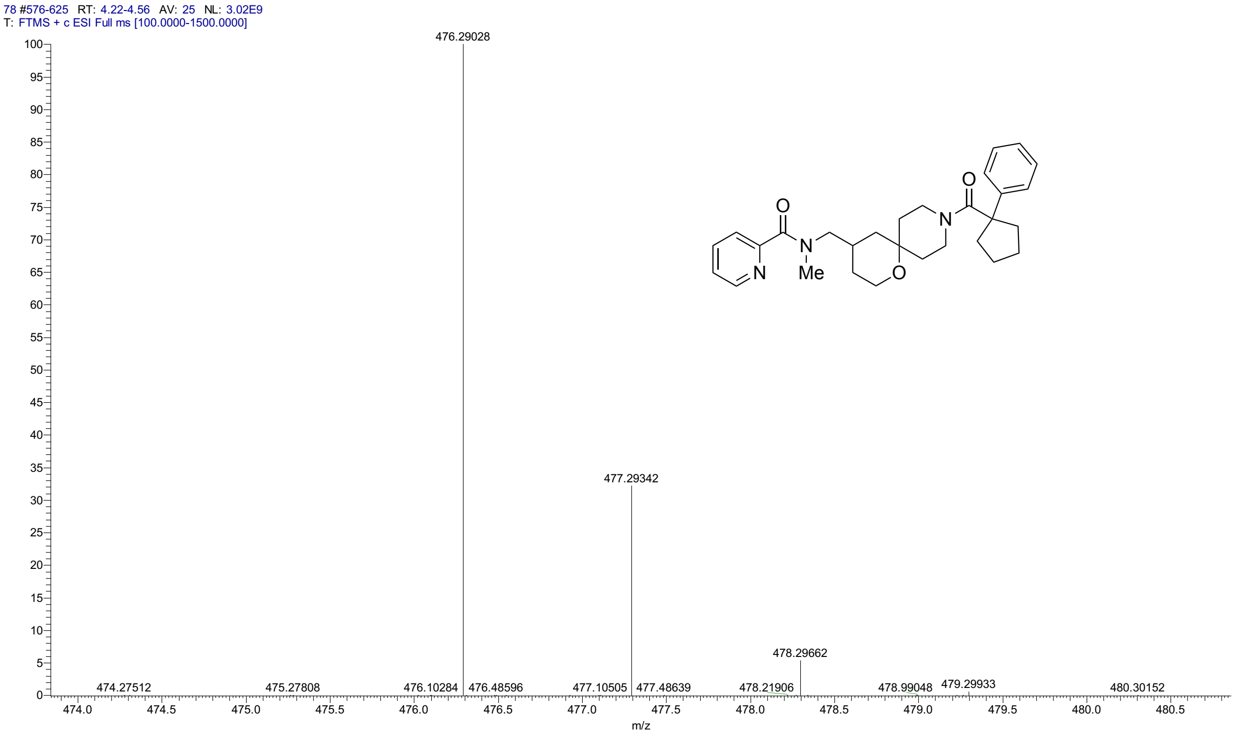


HPLC for **3m**

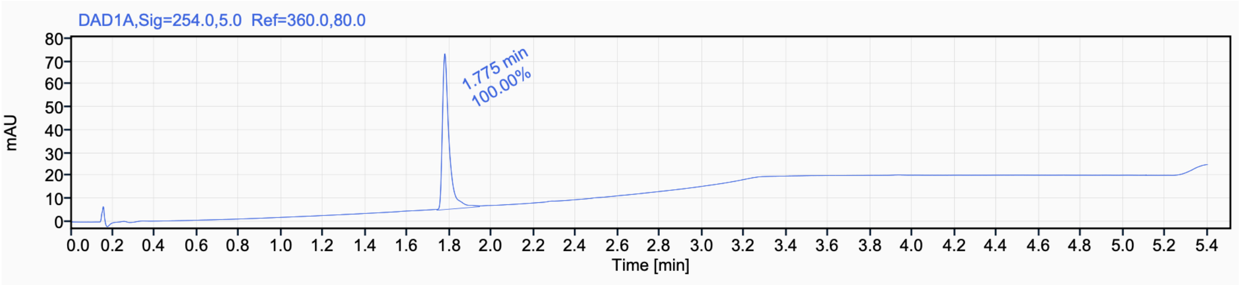


Signal Name DAD1A

| RT (min) | Signal description | m/z | Purity UV(‰) / MS(%) | Area | Area% |
|---|---|---|---|---|---|
| 1.775 | DAD1A,Sig=254.0,5.0 Ref=360.0,80.0 | | | 147.1 | 100.00 |

[1]H NMR (500 MHz, $CD_3CN$) for **3n**

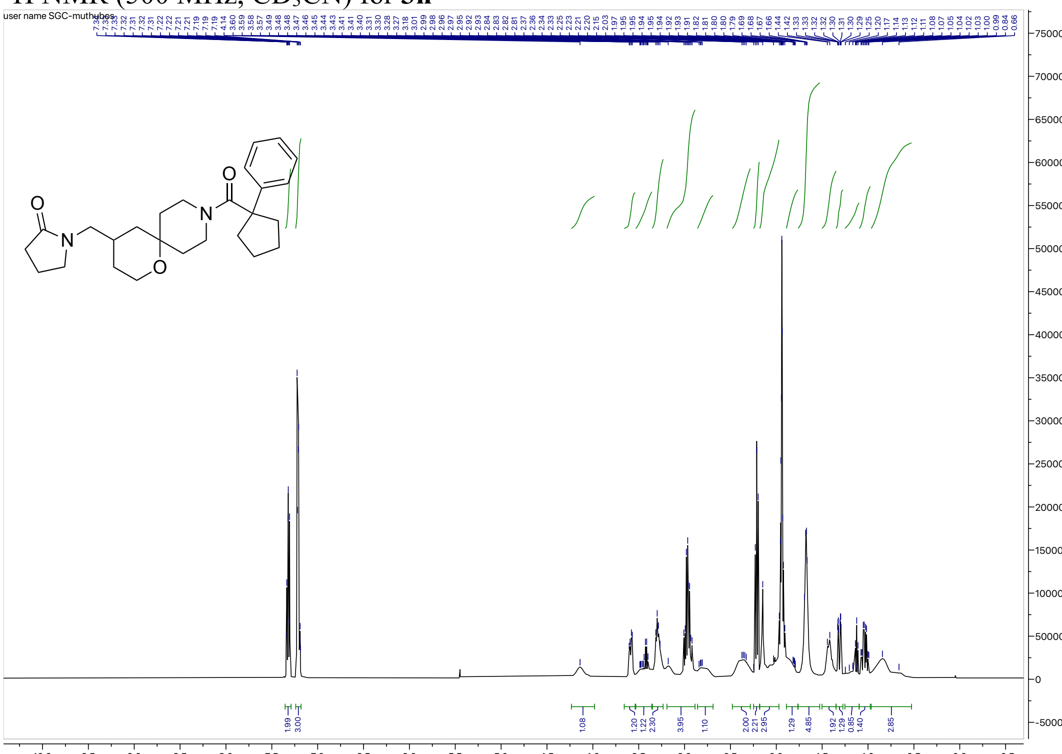


[13]C NMR (126 MHz, $CD_3CN$) for **3n**

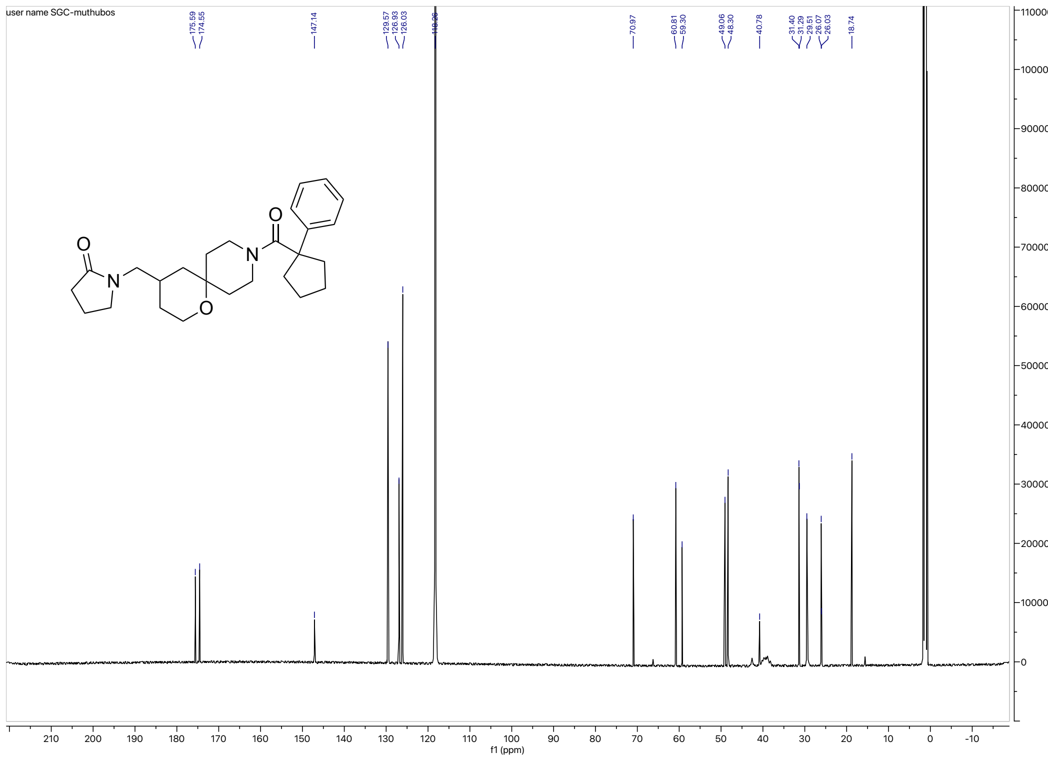

#### HRMS for **3n**

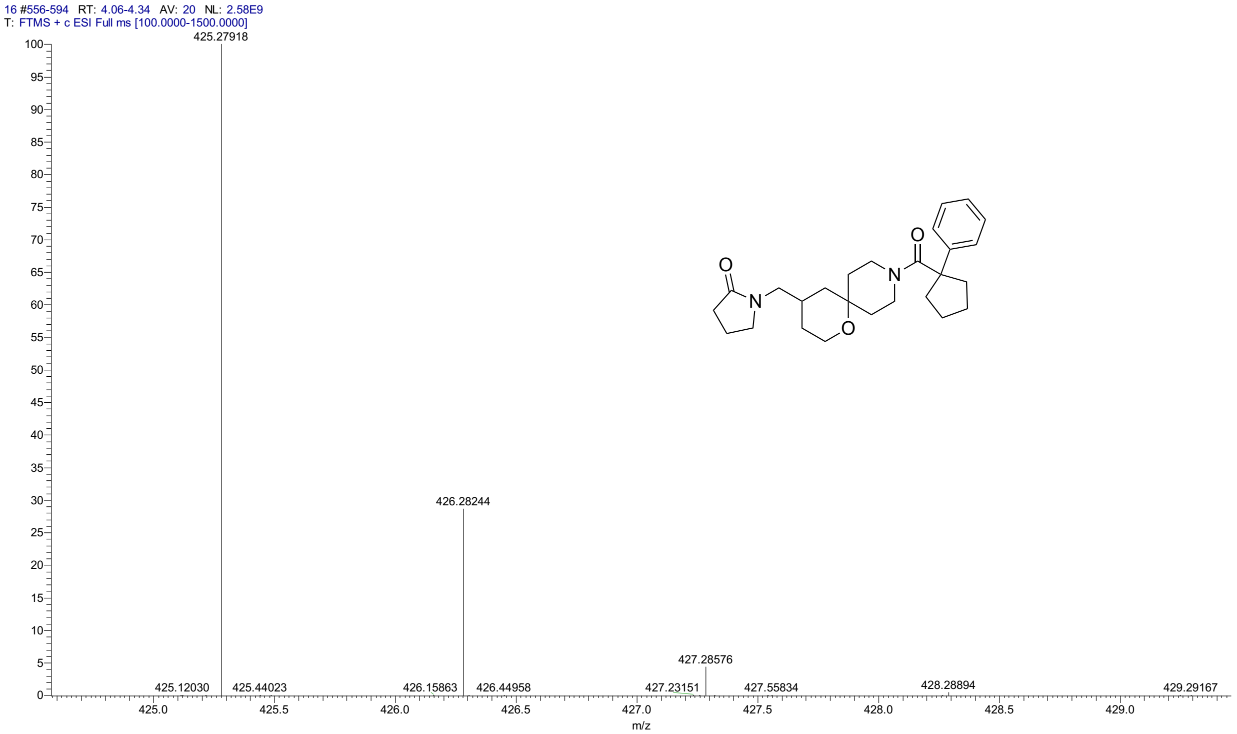


#### HPLC for **3n**

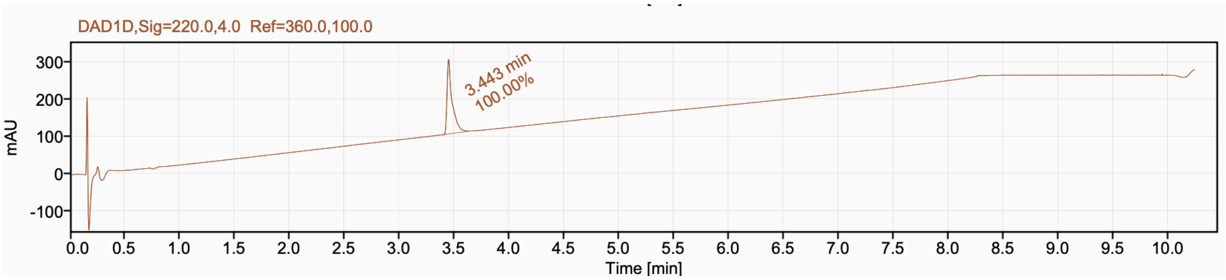


Signal Name DAD1D

| RT (min) | Signal description | Area | Area% | Peak Resolution USP |
|---|---|---|---|---|
| 3.443 | DAD1D,Sig=220.0,4.0 Ref=360.0,100.0 | 654.7 | 100.00 | |

#### $^{1}$H NMR (500 MHz, $CD_3CN$) for **3o**

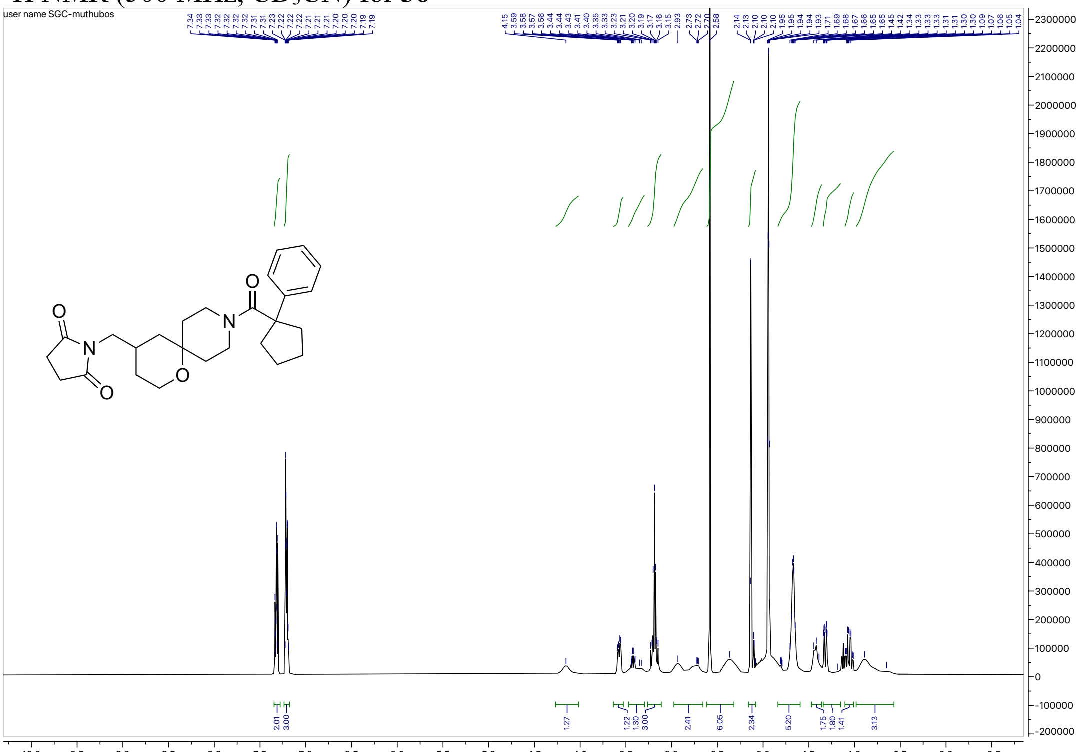


#### $^{13}$C NMR (126 MHz, $CD_3CN$) for **3o**

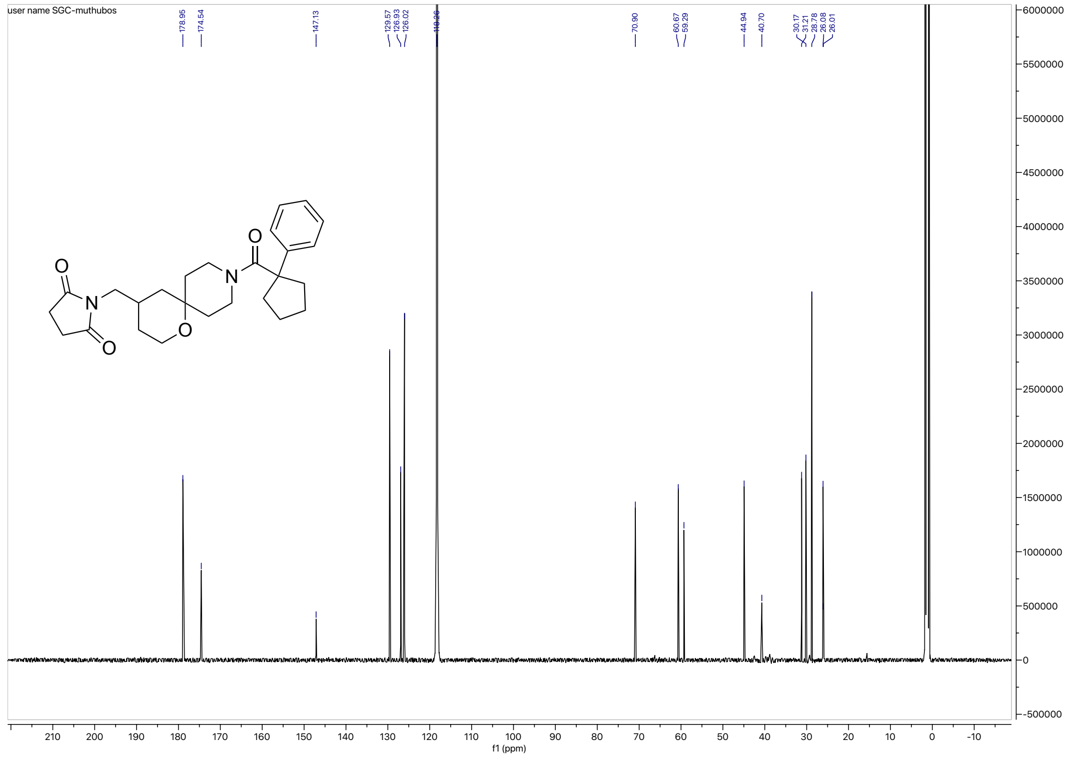

#### HRMS for **3o**

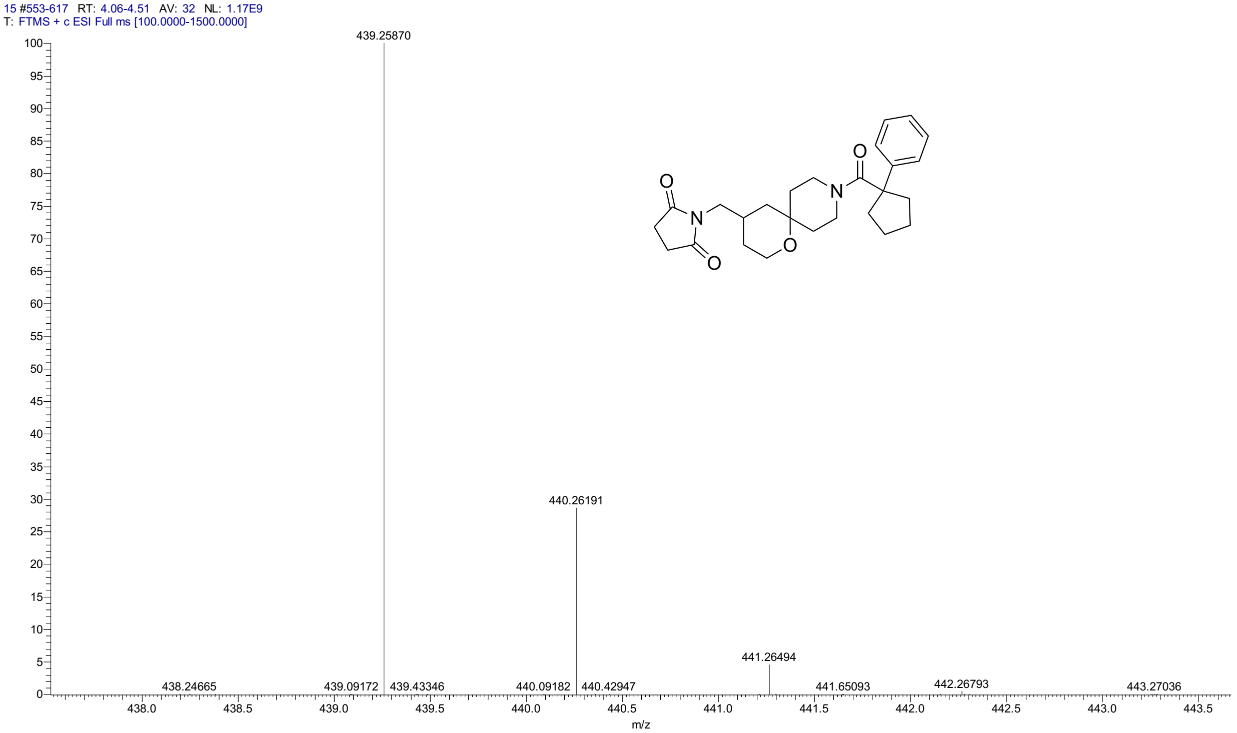


#### HPLC for **3o**

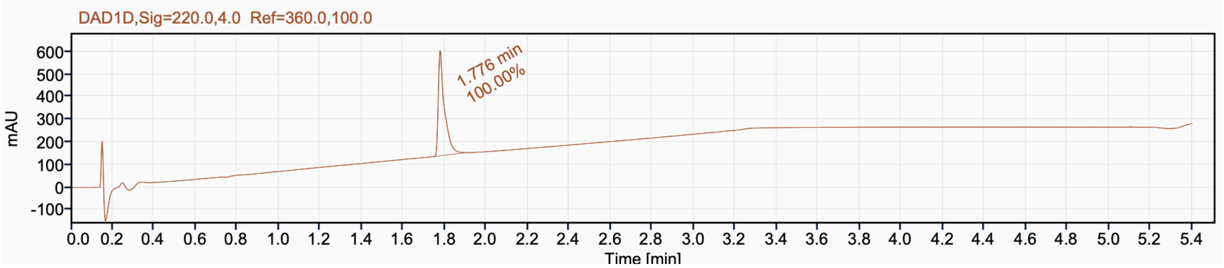


Signal Name DAD1D

| RT (min) | Signal description | Area | Area% | Peak Resolution USP |
|---|---|---|---|---|
| 1.776 | DAD1D,Sig=220.0,4.0 Ref=360.0,100.0 | 966.9 | 100.00 | |

$^{1}H$ NMR (400 MHz, $CD_3CN$) for **3p**

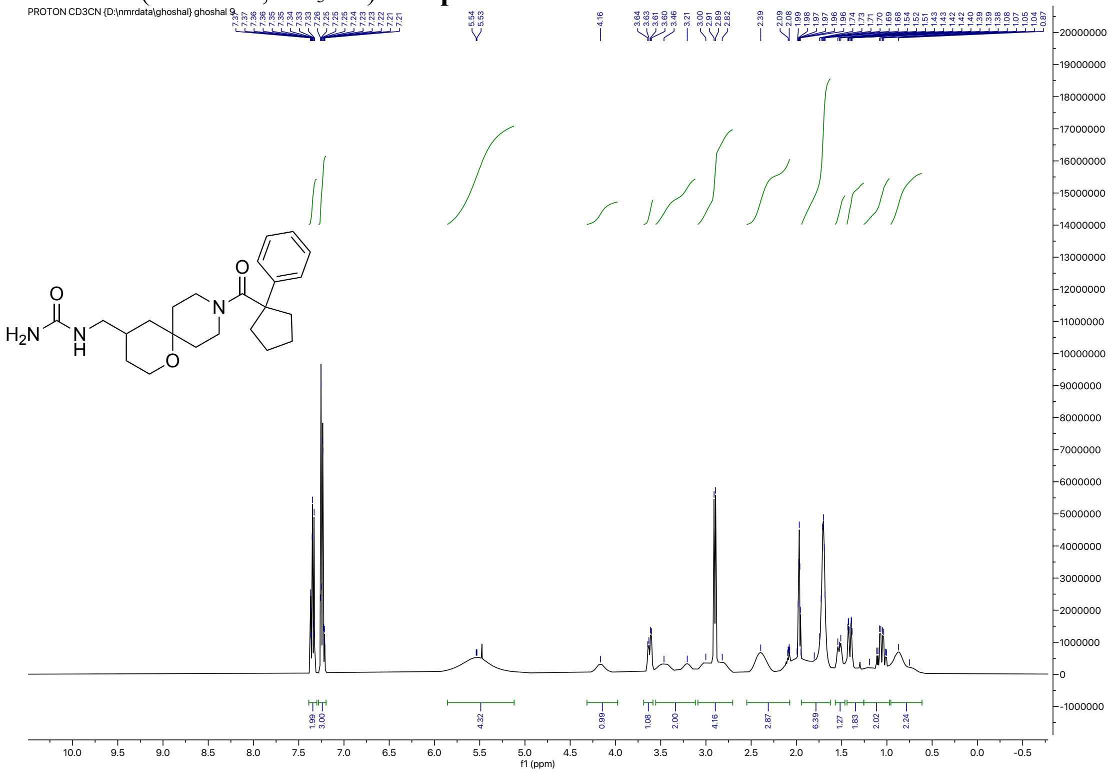

$^{13}C$ NMR (126 MHz, $CD_3CN$) for **3p**

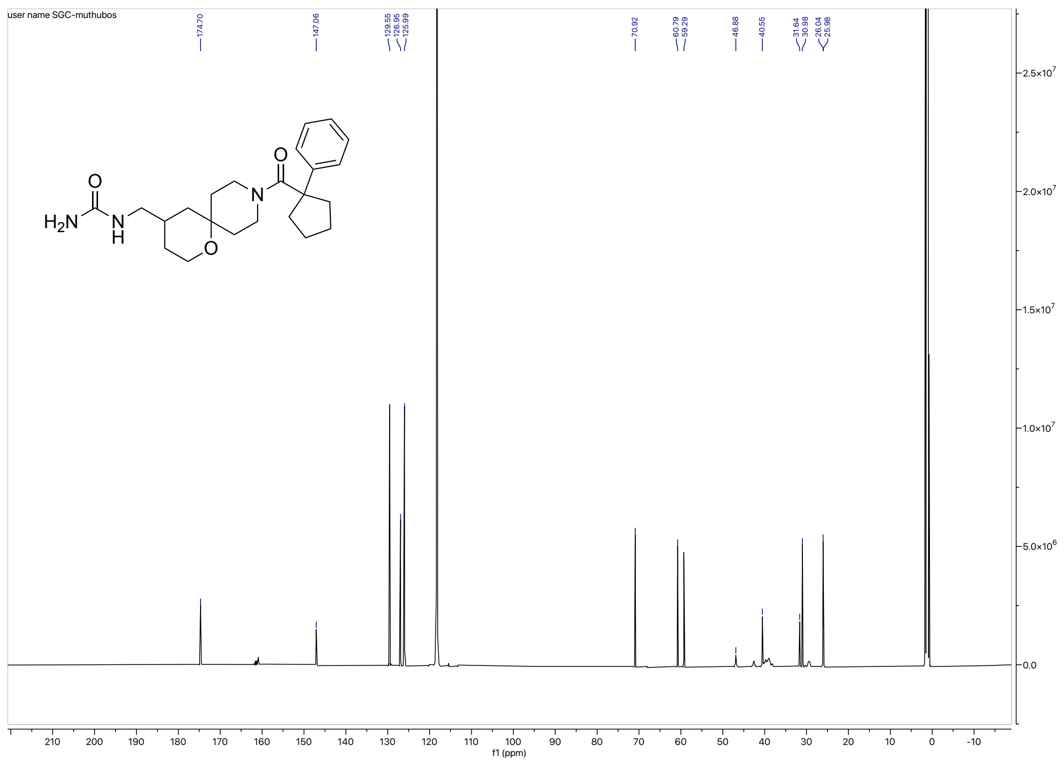

### HRMS for **3p**

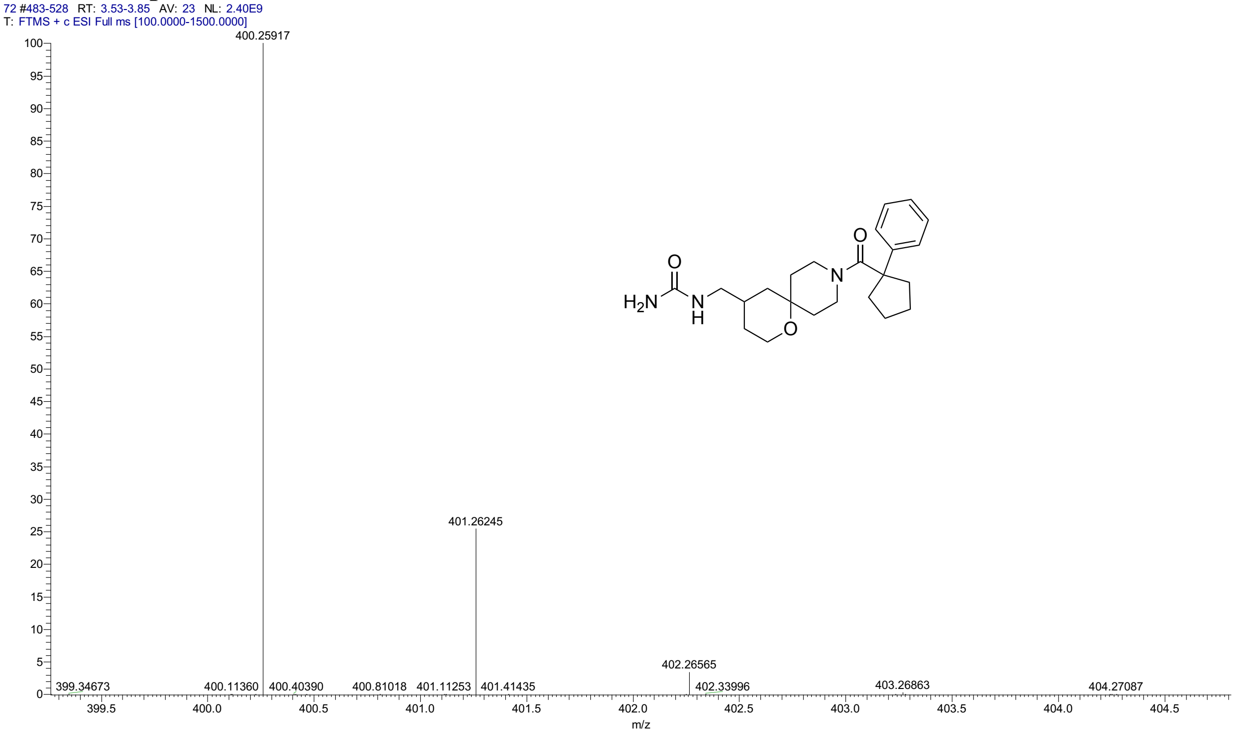


### HPLC for **3p**

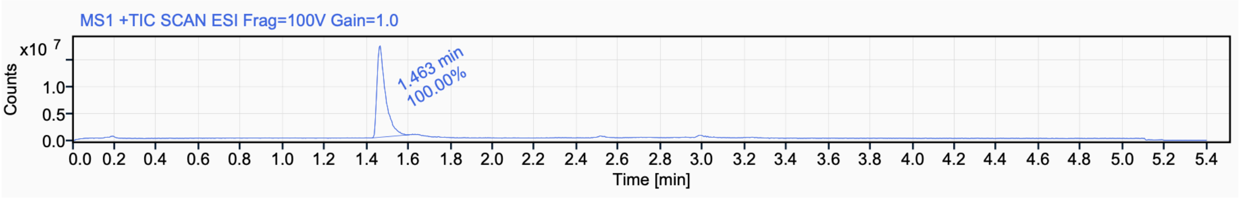


| Signal Name | MS1 +TIC SCAN ESI Frag=100V Gain=1.0 | | | |
|---|---|---|---|---|
| **RT (min)** | **Signal description** | **Area** | **Area%** | **Peak Resolution USP** |
| 1.463 | MS1 +TIC SCAN ESI Frag=100V Gain=1.0 | 49005591.2 | 100.00 | |

$^{1}$H NMR (500 MHz, $CD_3CN$) for **3q**

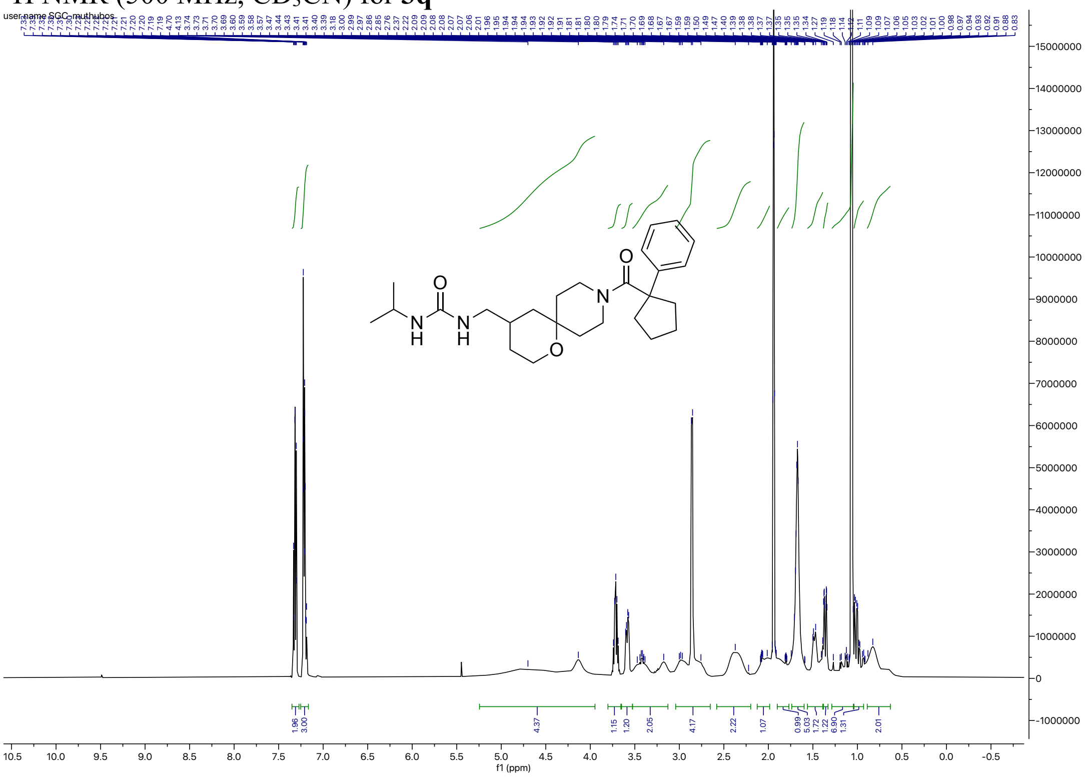


$^{13}$C NMR (126 MHz, $CD_3CN$) for **3q**

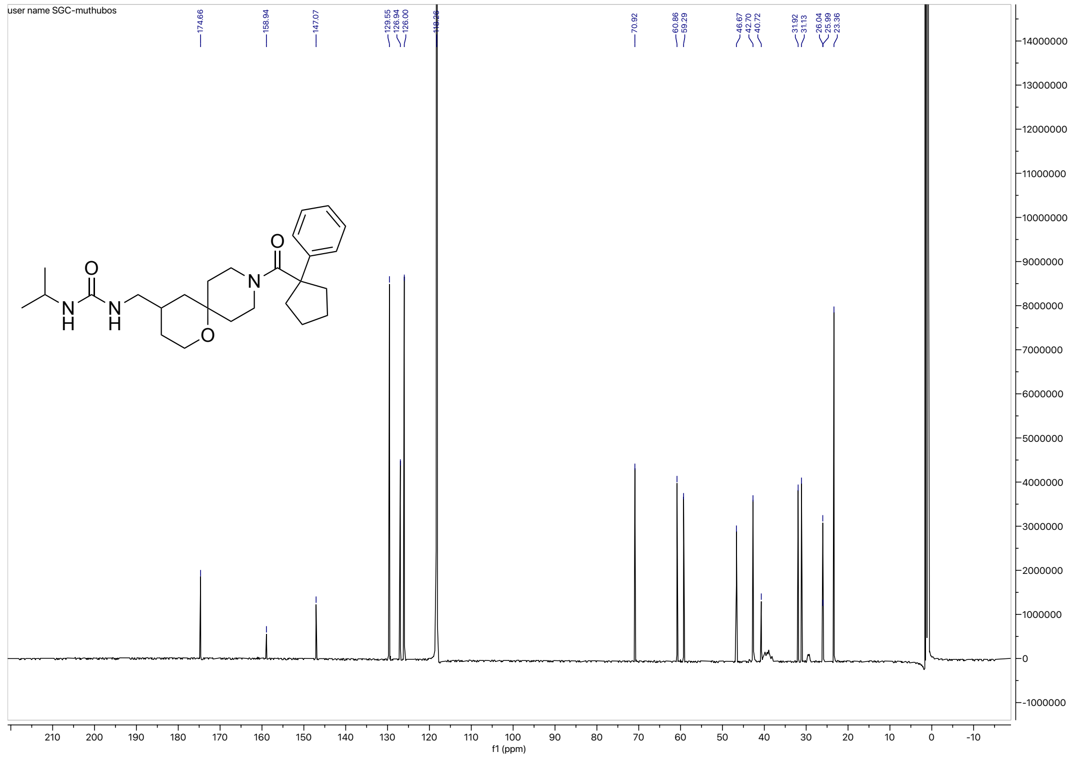

#### HRMS for **3q**

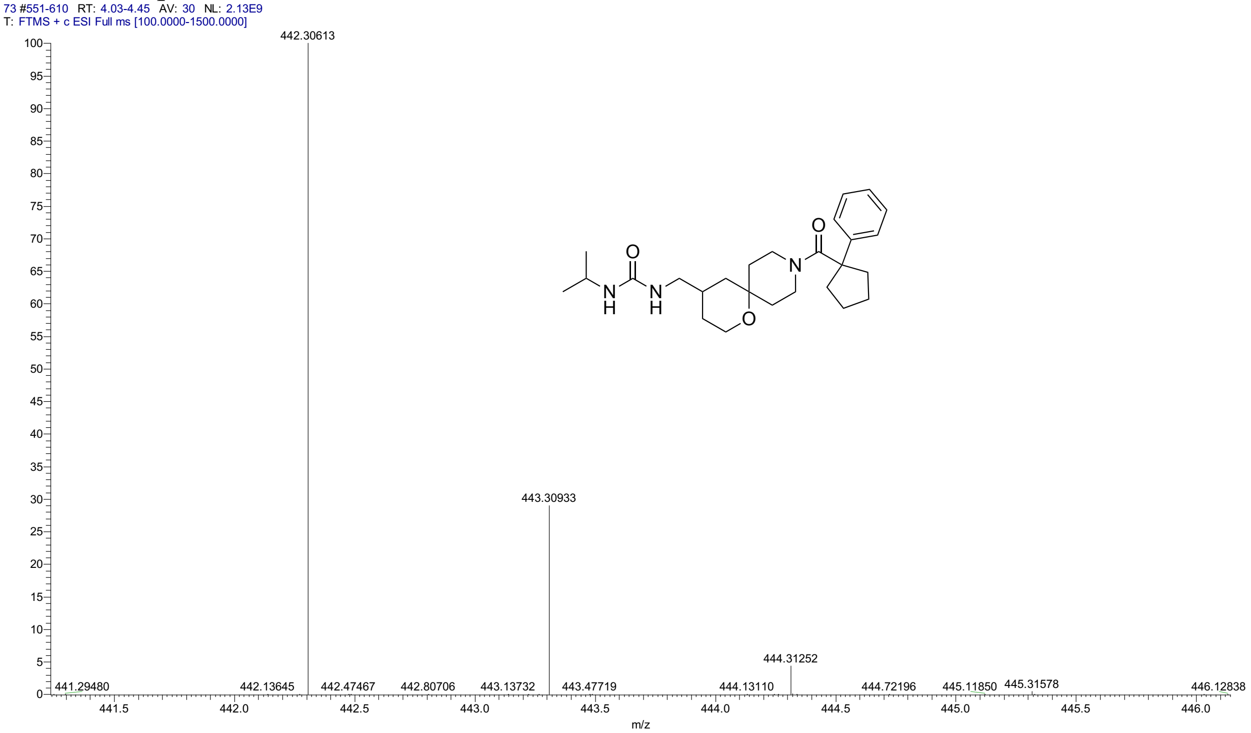


#### HPLC for **3q**

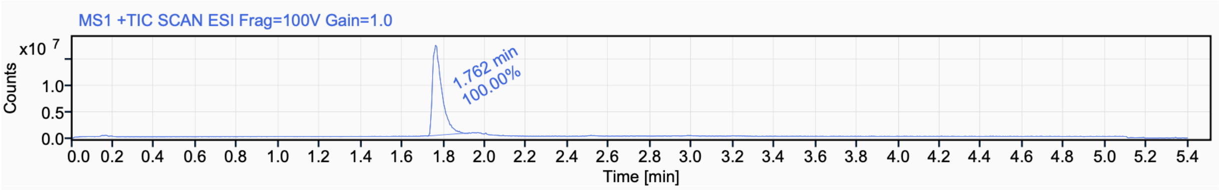


Signal Name MS1 +TIC SCAN ESI Frag=100V Gain=1.0

| RT (min) | Signal description | m/z | Purity UV(‰) / MS(%) | Area | Area% |
|---|---|---|---|---|---|
| 1.762 | MS1 +TIC SCAN ESI Frag=100V Gain=1.0 | | | 54618137.8 | 100.00 |

#### $^{1}$H NMR (500 MHz, $CD_3CN$) for **3r**

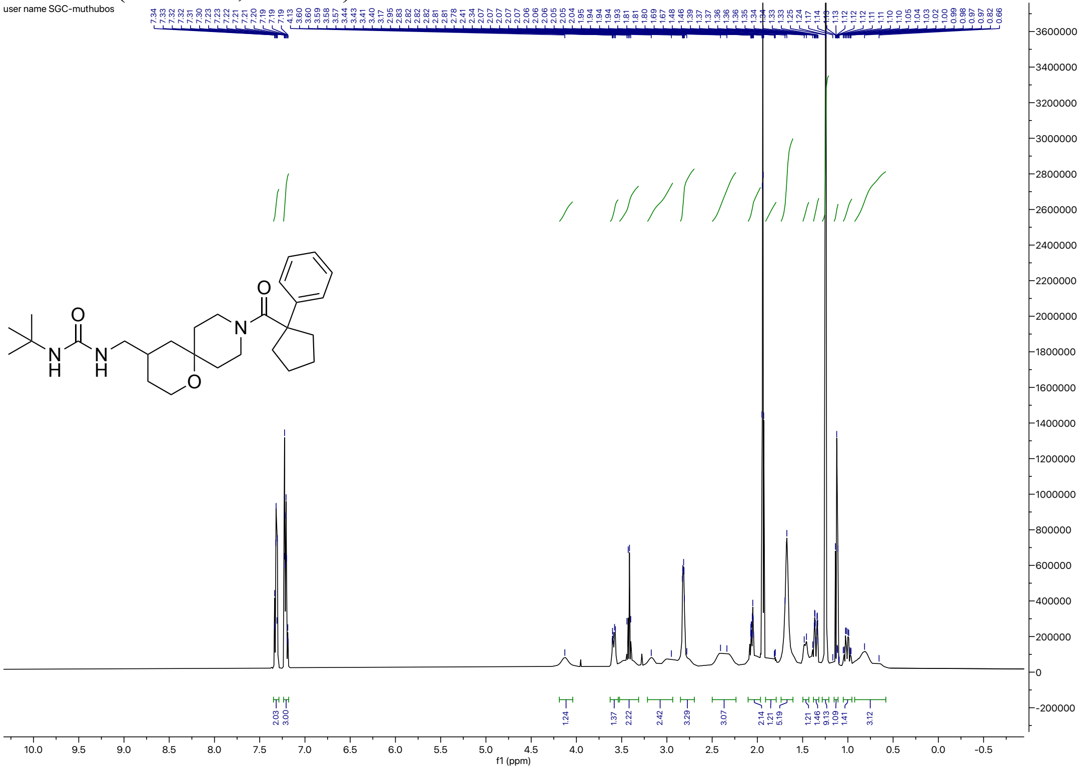


#### $^{13}$C NMR (126 MHz, $CD_3CN$) for **3r**

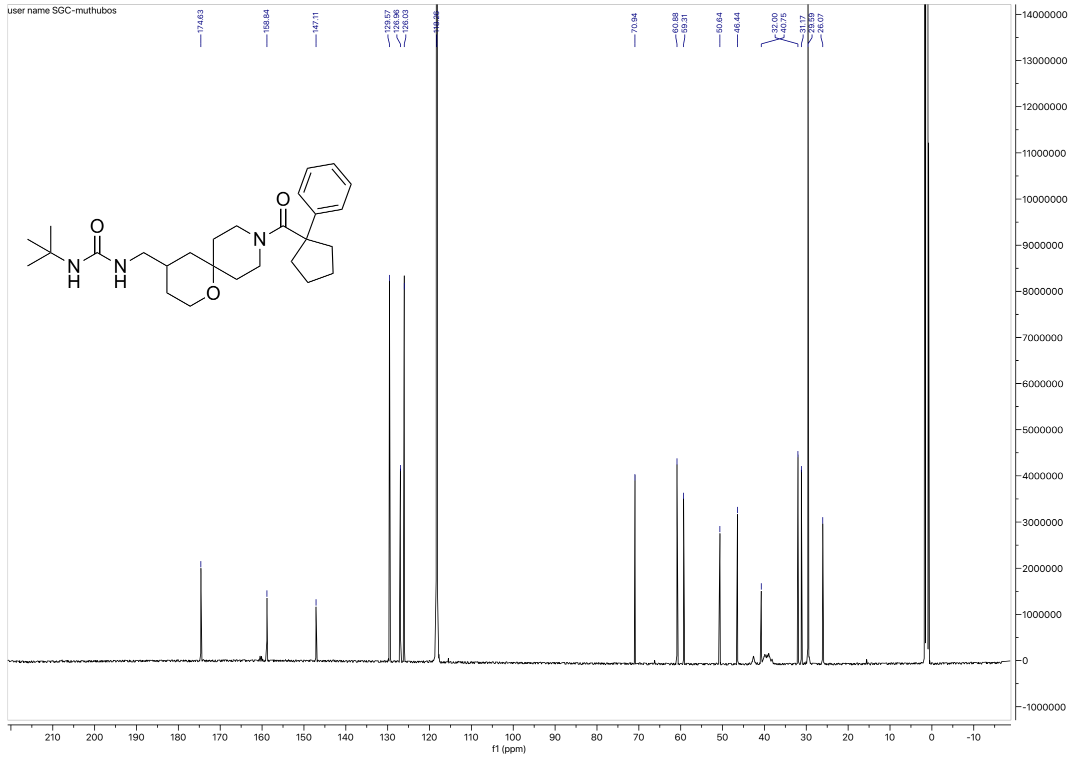

HRMS for **3r**

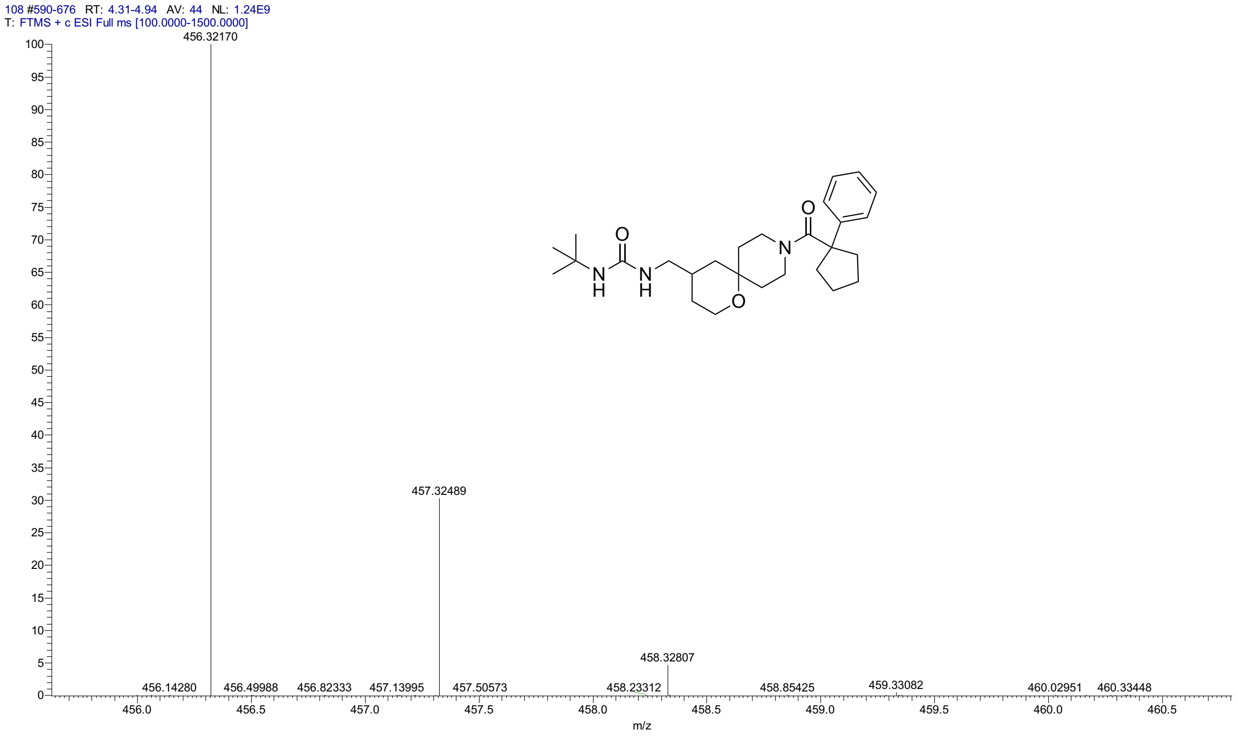


HPLC for **3r**

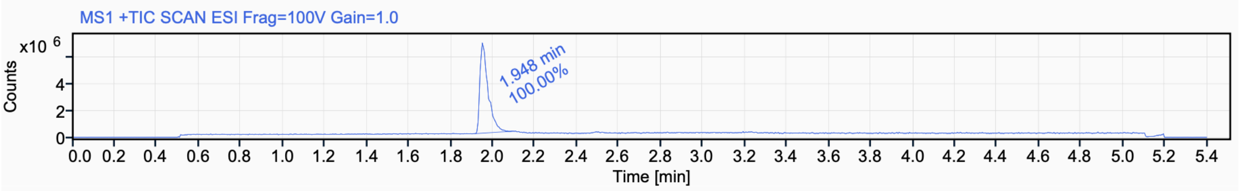


Signal Name MS1 +TIC SCAN ESI Frag=100V Gain=1.0

| RT (min) | Signal description | Area | Area% | Peak Resolution USP |
|---|---|---|---|---|
| 1.948 | MS1 +TIC SCAN ESI Frag=100V Gain=1.0 | 18237574.0 | 100.00 | |

$^{1}$H NMR (500 MHz, $CD_3CN$) for **3s**

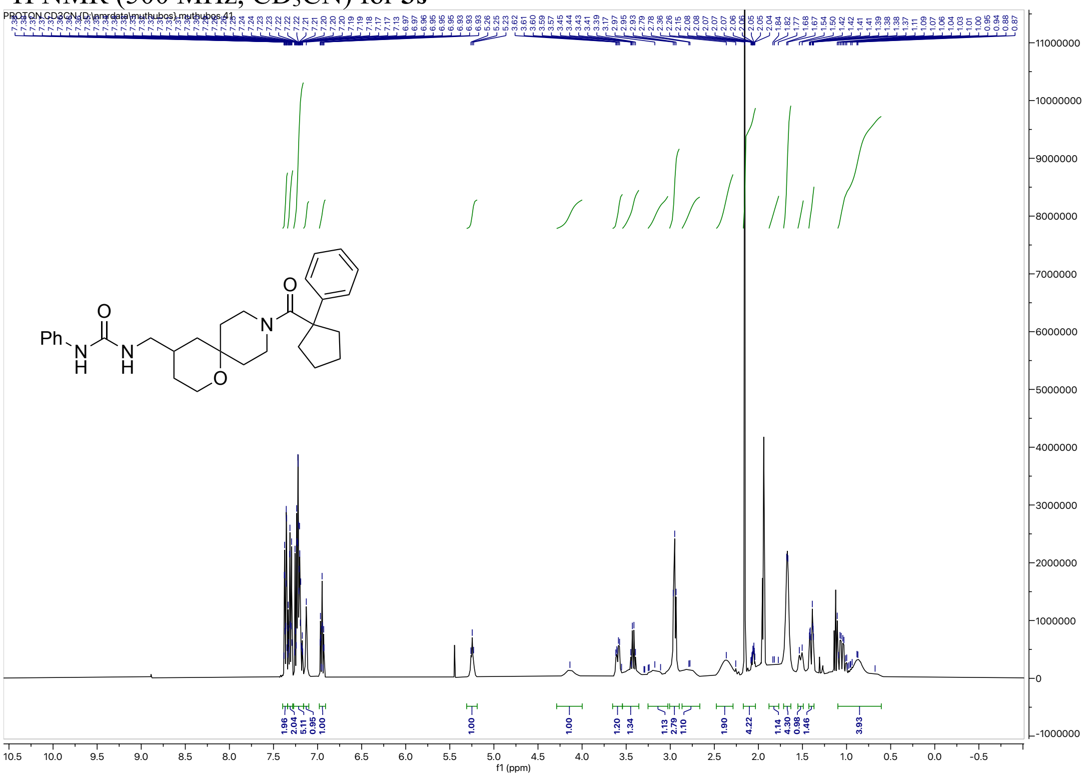

$^{13}$C NMR (126 MHz, $CD_3CN$) for **3s**

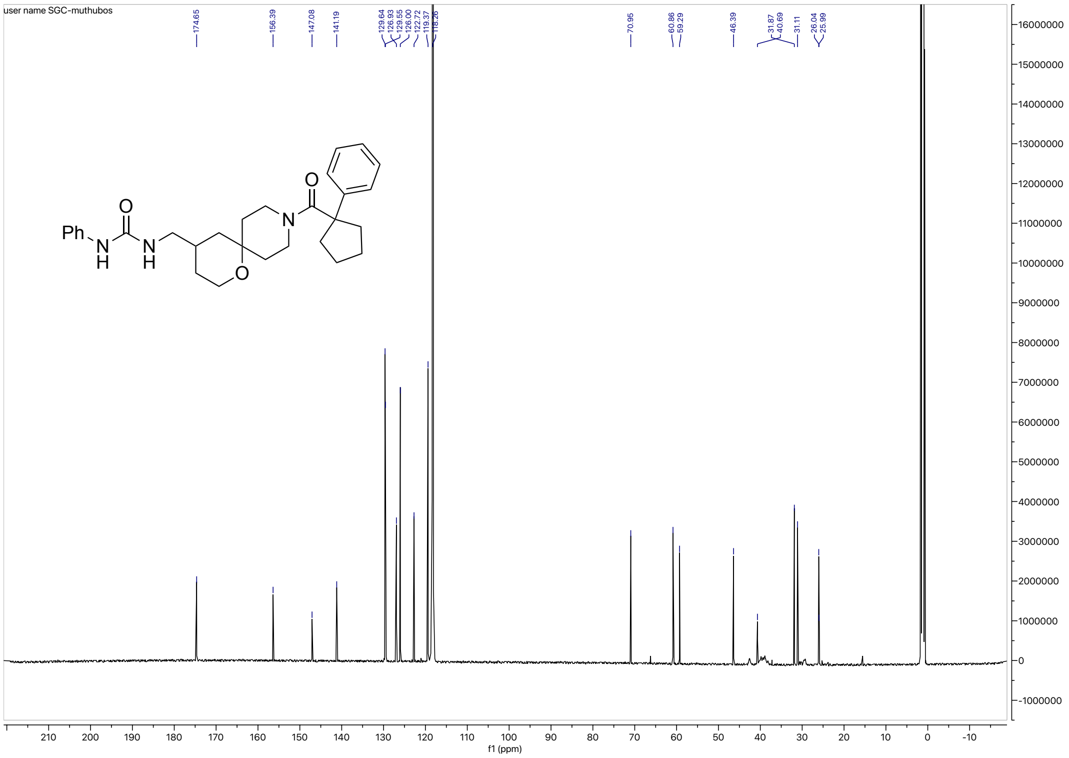

HRMS for **3s**

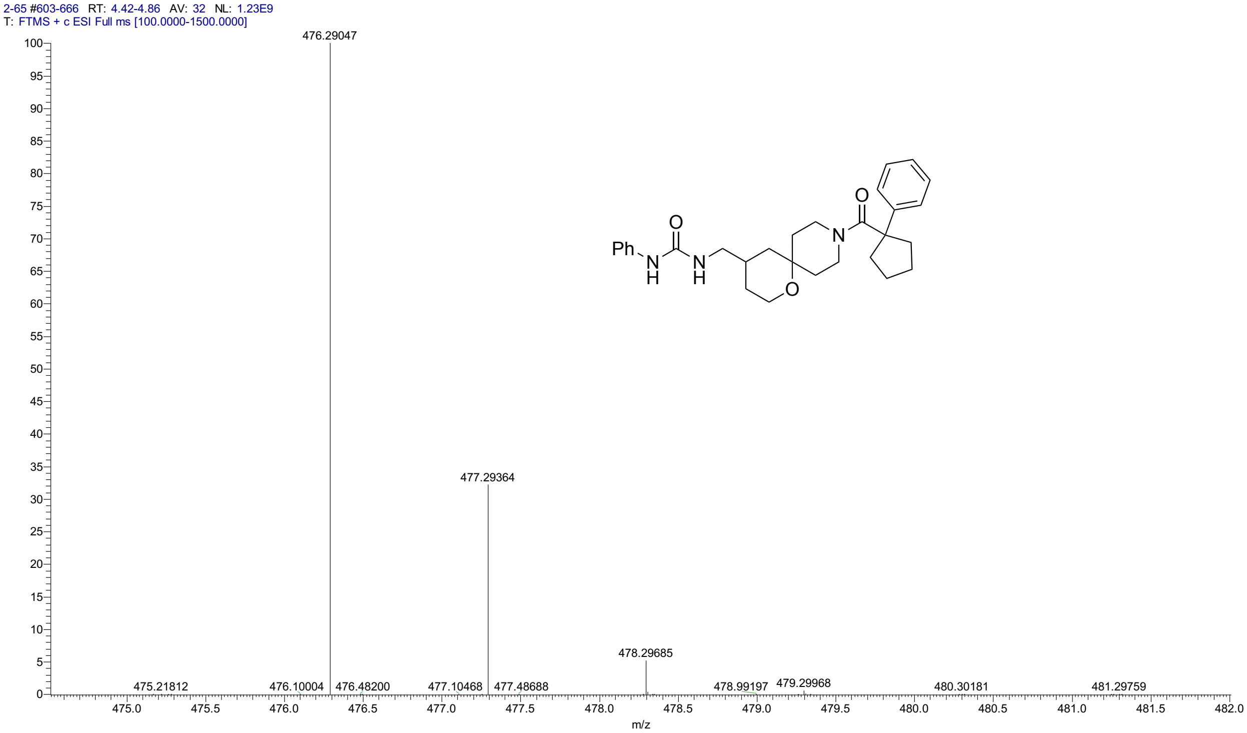


HPLC for **3s**

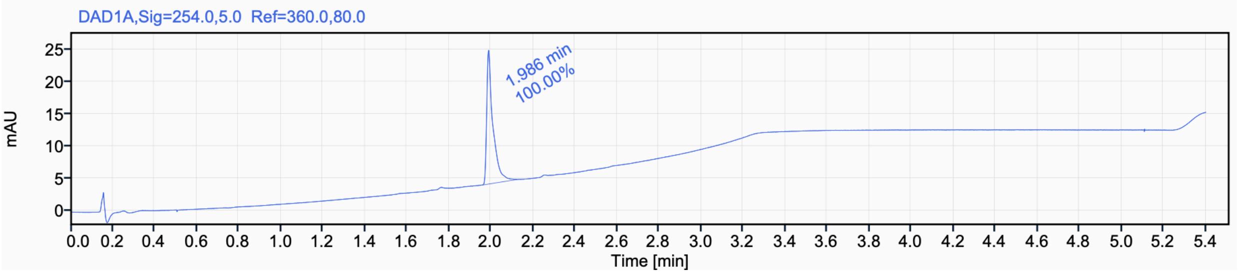


Signal Name DAD1A

| RT (min) | Signal description | Area | Area% | Peak Resolution USP |
|---|---|---|---|---|
| 1.986 | DAD1A,Sig=254.0,5.0 Ref=360.0,80.0 | 42.3 | 100.00 | |

[1]H NMR (500 MHz, $CD_3CN$) for **3t**

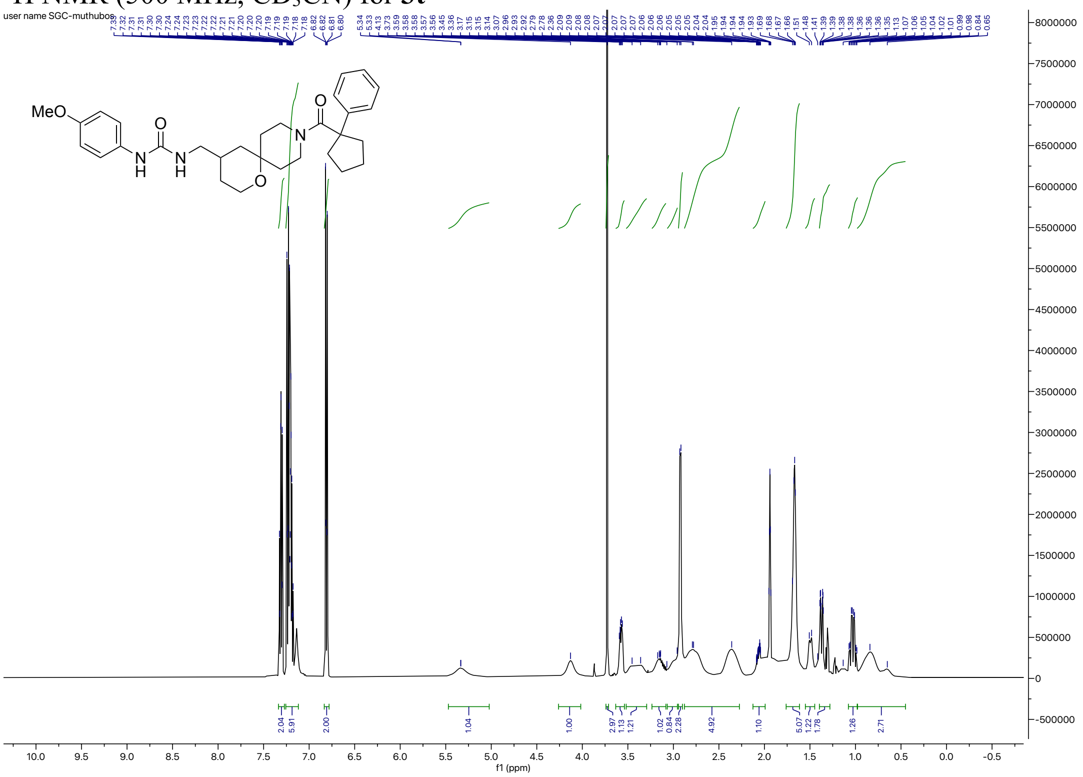

[13]C NMR (126 MHz, $CD_3CN$) for **3t**

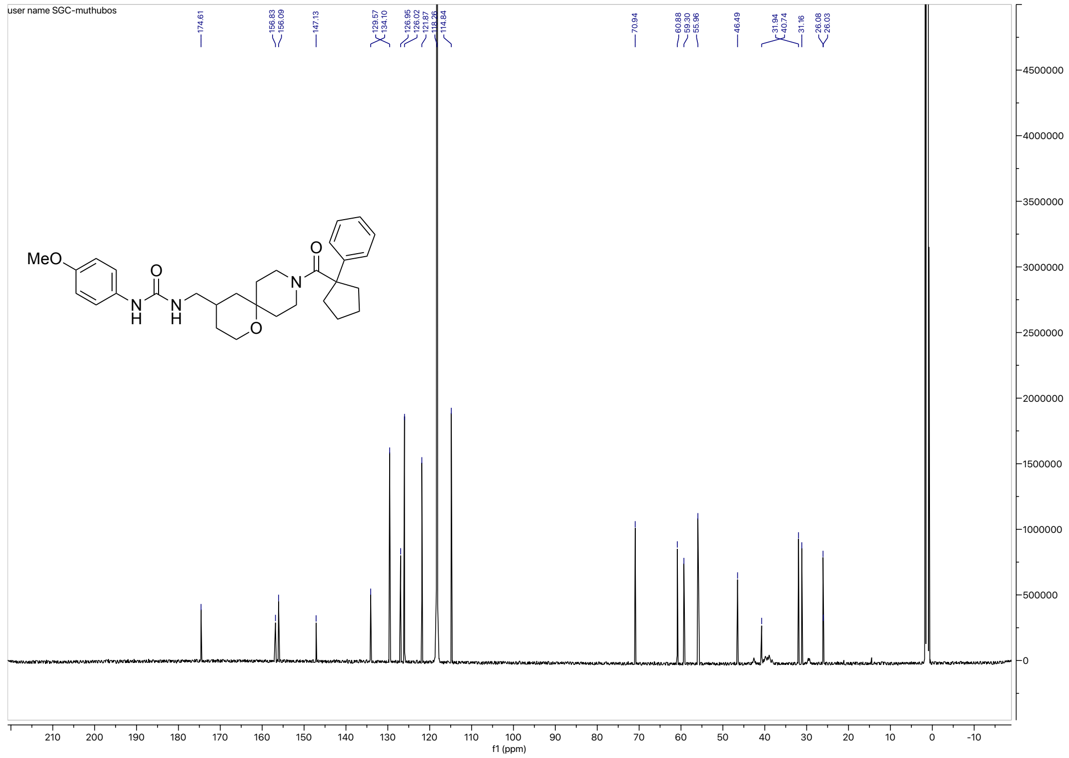

#### HRMS for **3t**

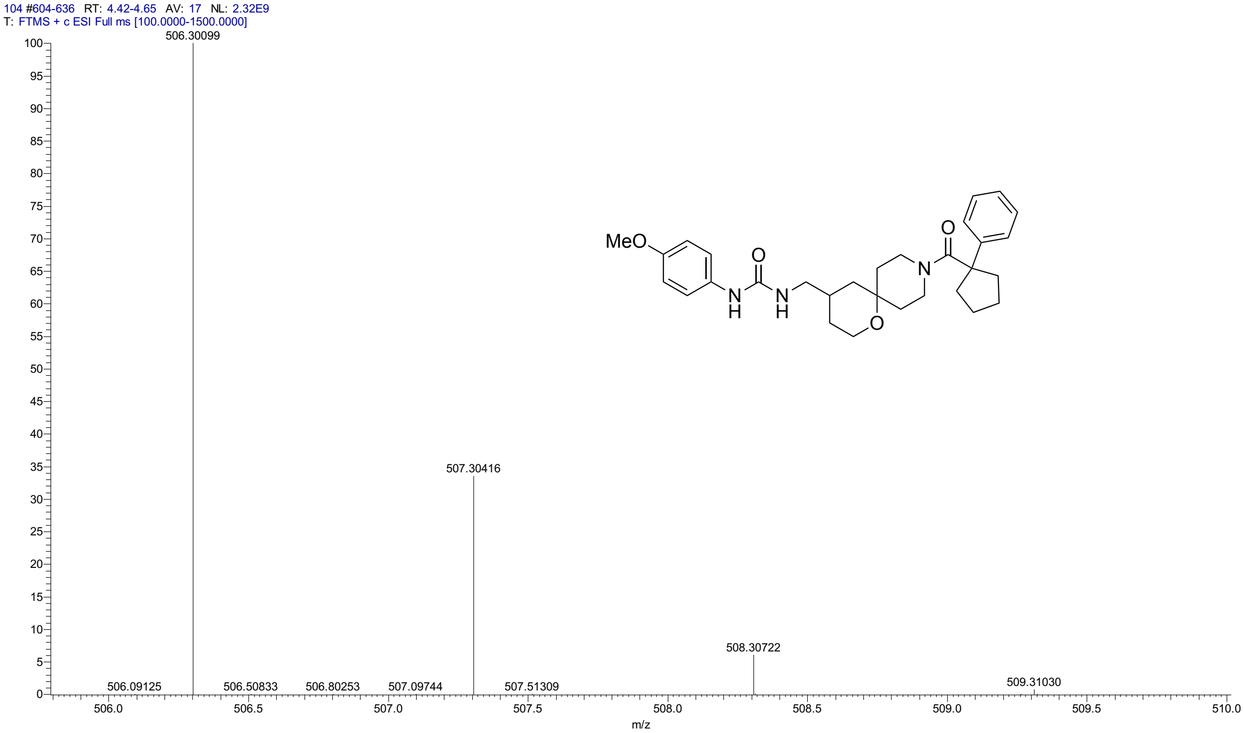


#### HPLC for **3t**

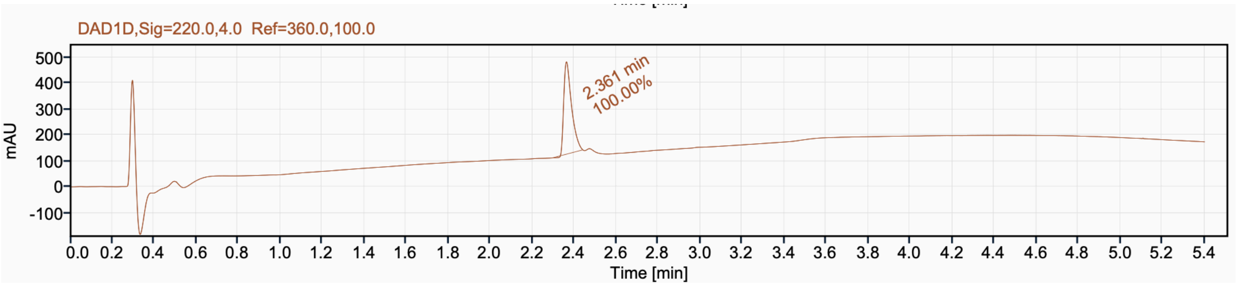


Signal Name DAD1D

| RT (min) | Signal description | Area | Area% | Peak Resolution USP |
|---|---|---|---|---|
| 2.361 | DAD1D,Sig=220.0,4.0 Ref=360.0,100.0 | 916.0 | 100.00 | |

$^{1}$H NMR (500 MHz, $CD_3CN$) for **3u**

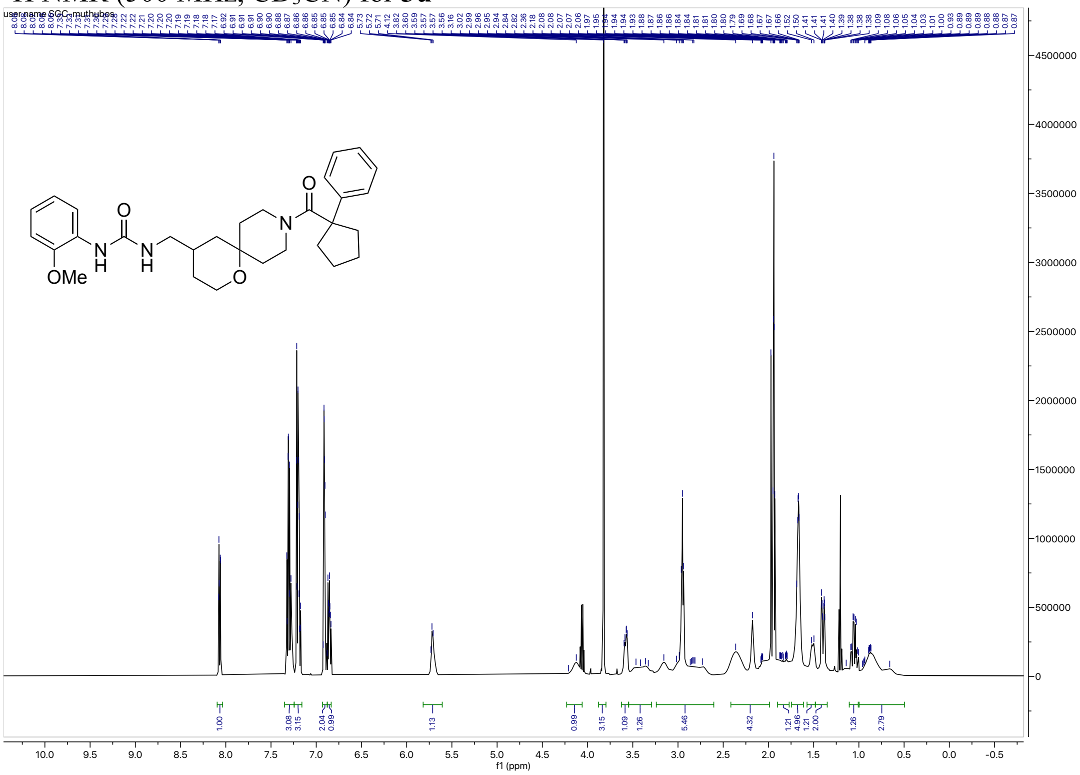


$^{13}$C NMR (126 MHz, $CD_3CN$) for **3u**

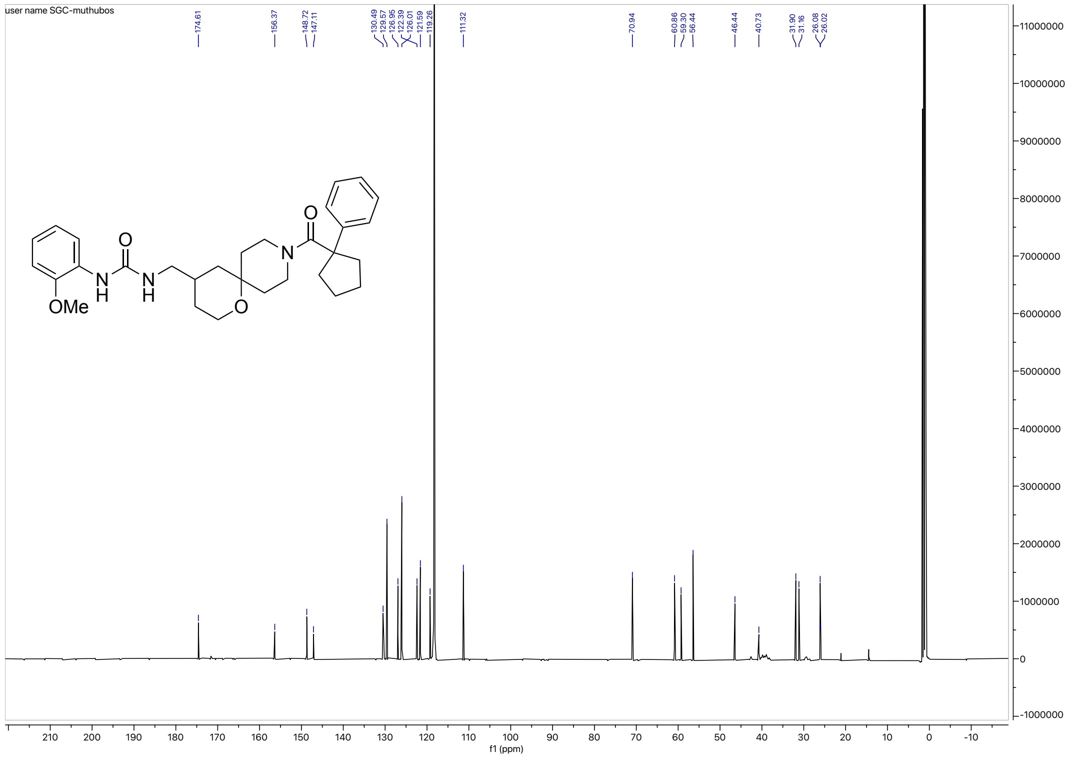

#### HRMS for **3u**

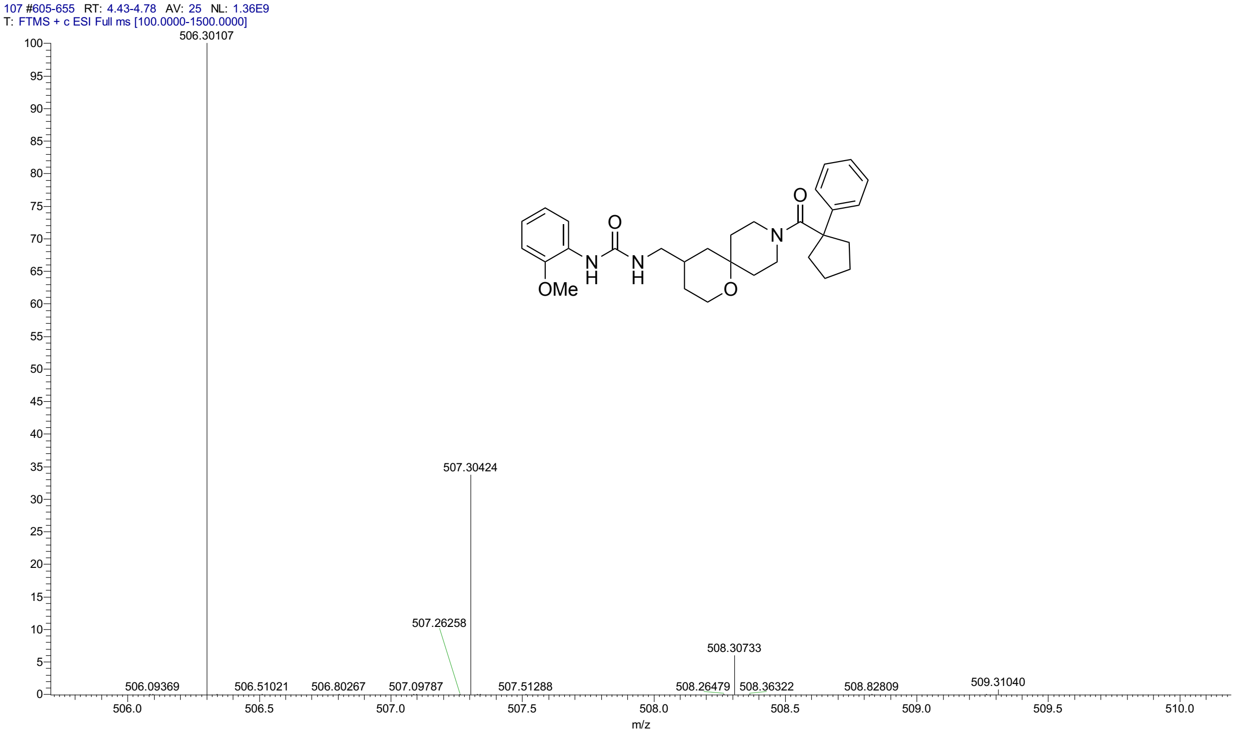


#### HPLC for **3u**

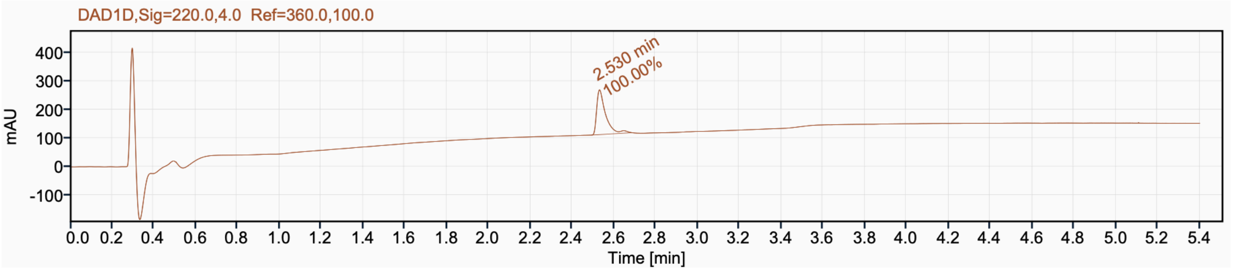


Signal Name DAD1D

| RT (min) | Signal description | Area | Area% | Peak Resolution USP |
|---|---|---|---|---|
| 2.530 | DAD1D,Sig=220.0,4.0 Ref=360.0,100.0 | 464.0 | 100.00 | |

#### $^{1}$H NMR (400 MHz, $CD_3OD$) for **4a**

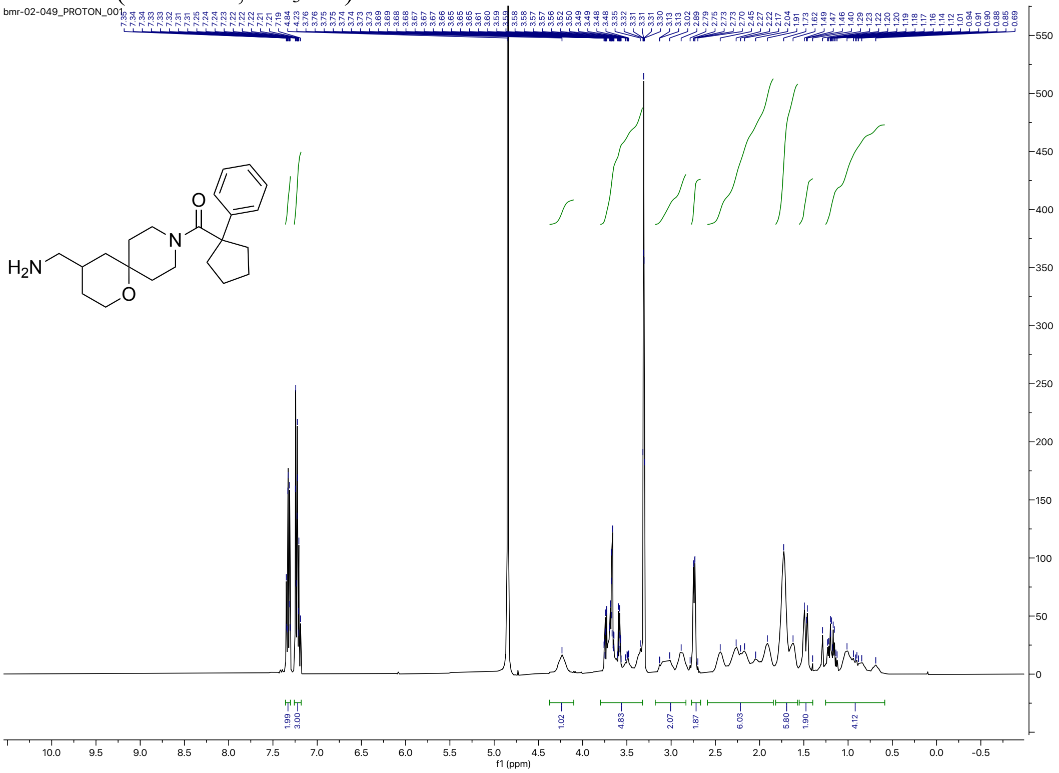


#### $^{13}$C NMR (100 MHz, $CD_3OD$) for **4a**

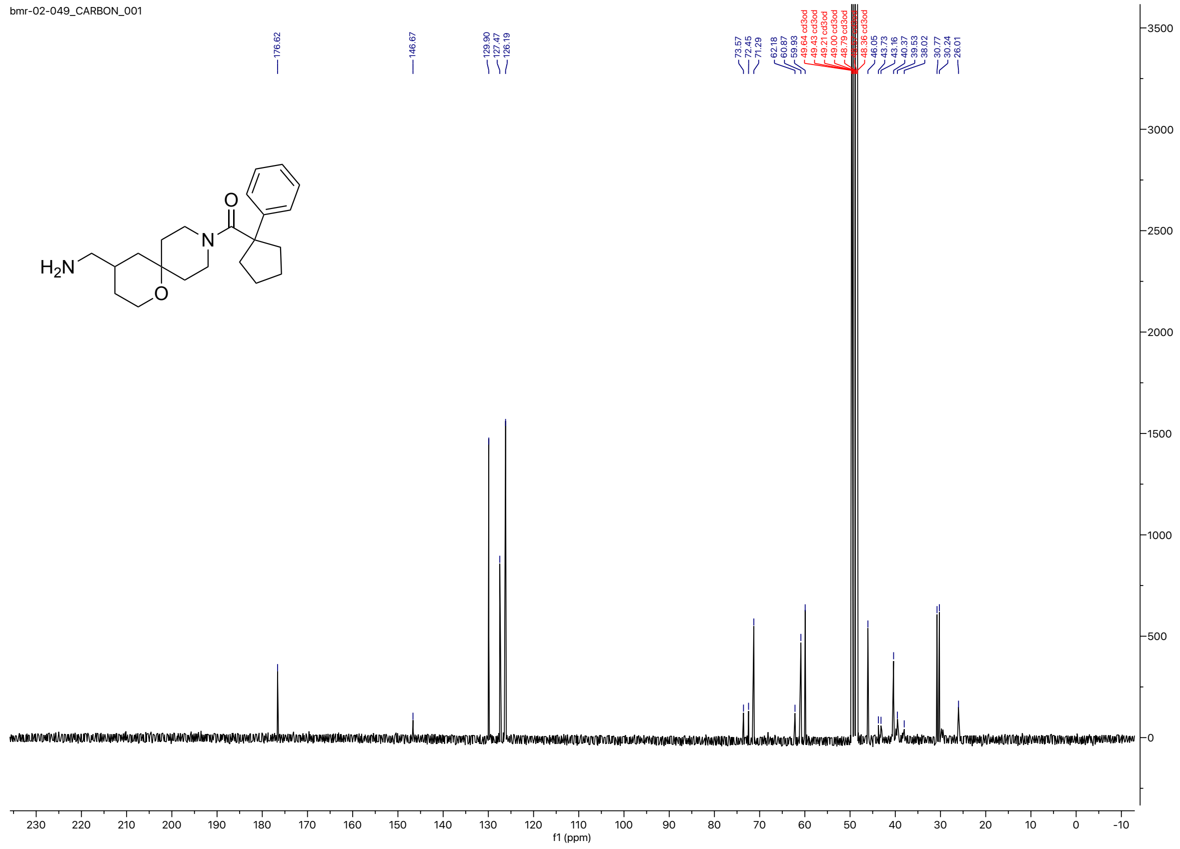

#### HRMS for **4a**

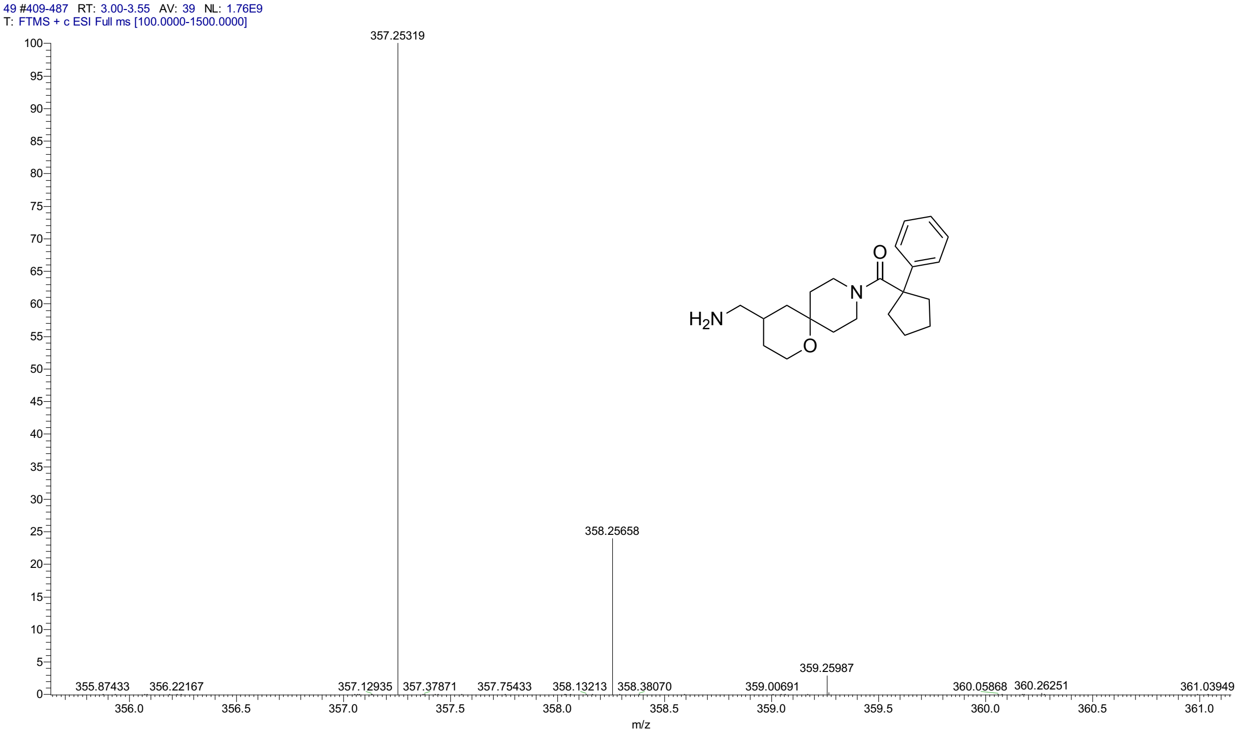


#### HPLC for **4a**

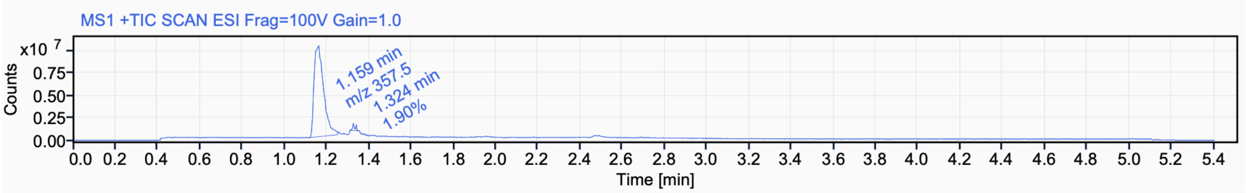


Signal Name MS1 +TIC SCAN ESI Frag=100V Gain=1.0

| RT (min) | Signal description | m/z | Purity UV(‰) / MS(%) | Area | Area% |
|---|---|---|---|---|---|
| 1.159 | MS1 +TIC SCAN ESI Frag=100V Gain=1.0 | 357.5 | | 32209391.5 | 98.10 |
| 1.324 | MS1 +TIC SCAN ESI Frag=100V Gain=1.0 | | | 624767.1 | 1.90 |

$^{1}H$ NMR (500 MHz, $CD_3OD$) for **4b**

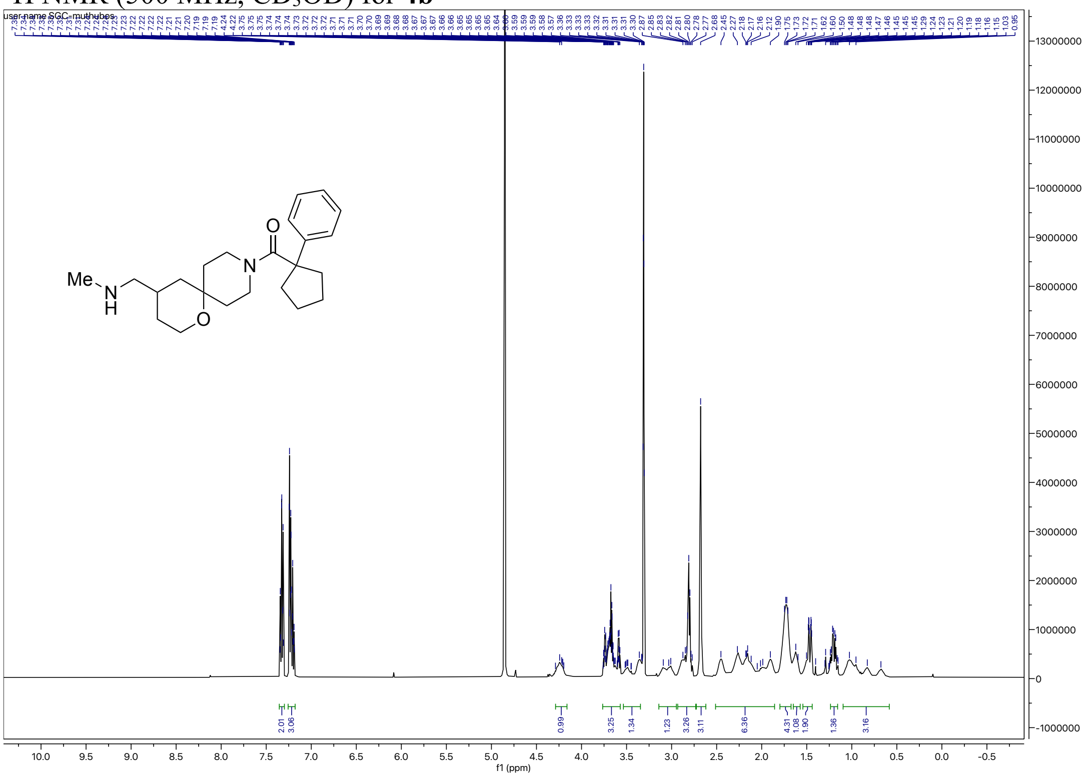

$^{13}C$ NMR (126 MHz, $CD_3OD$) for **4b**

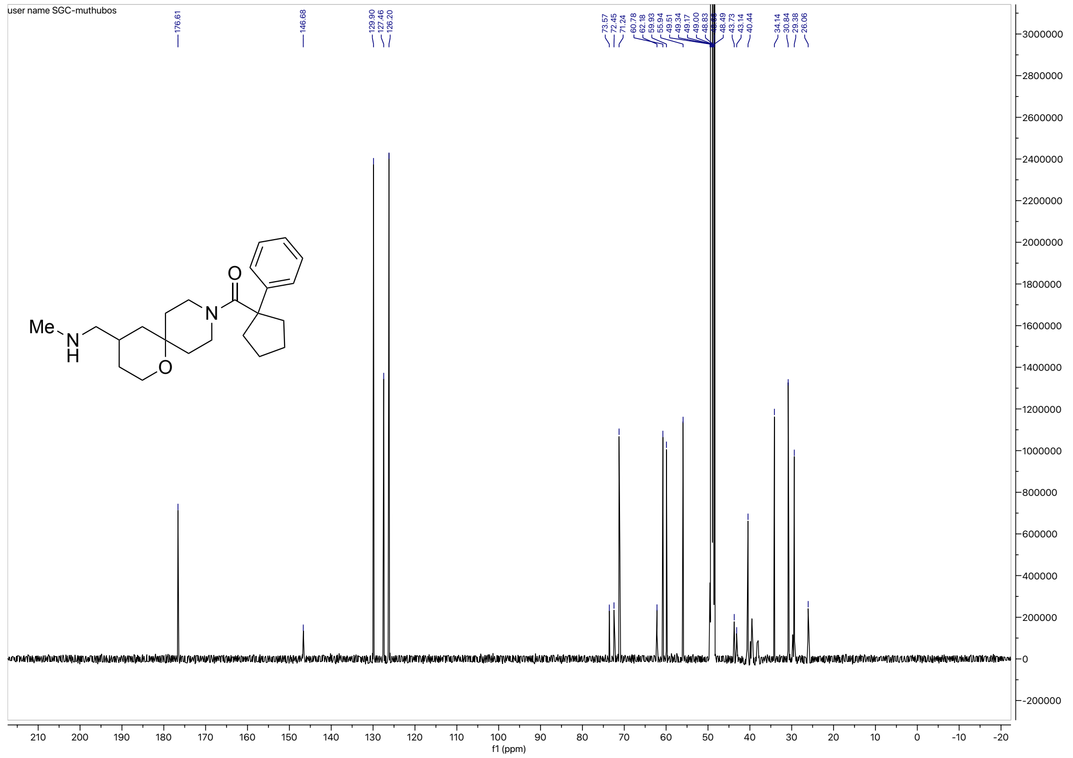

HRMS for **4b**

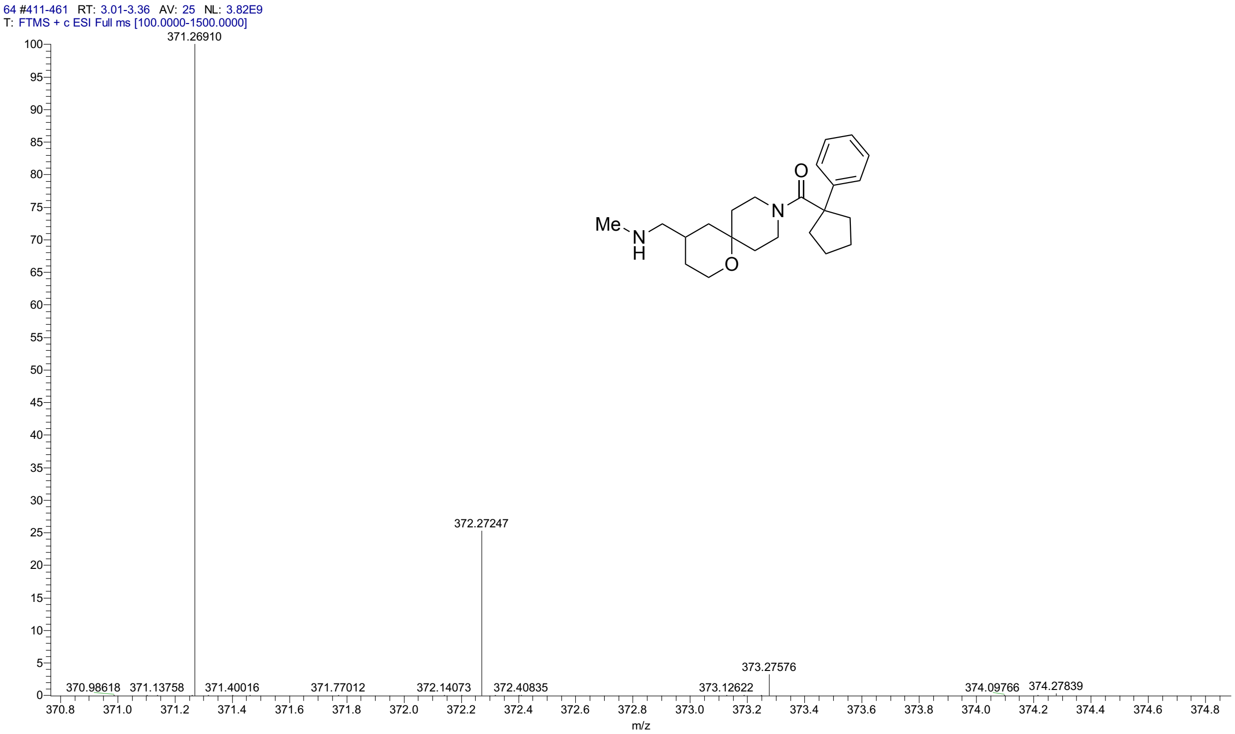


HPLC for **4b**

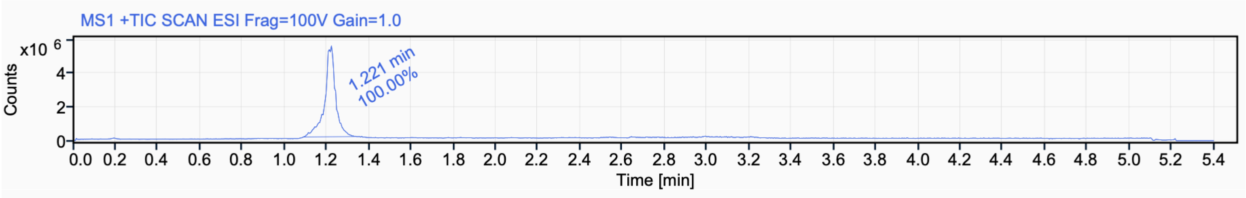


Signal Name MS1 +TIC SCAN ESI Frag=100V Gain=1.0

| RT (min) | Signal description | m/z | Purity UV(‰) / MS(%) | Area | Area% |
|---|---|---|---|---|---|
| 1.221 | MS1 +TIC SCAN ESI Frag=100V Gain=1.0 | | | 19184845.5 | 100.00 |

$^{1}$H NMR (500 MHz, $CD_3OD$) for **4c**

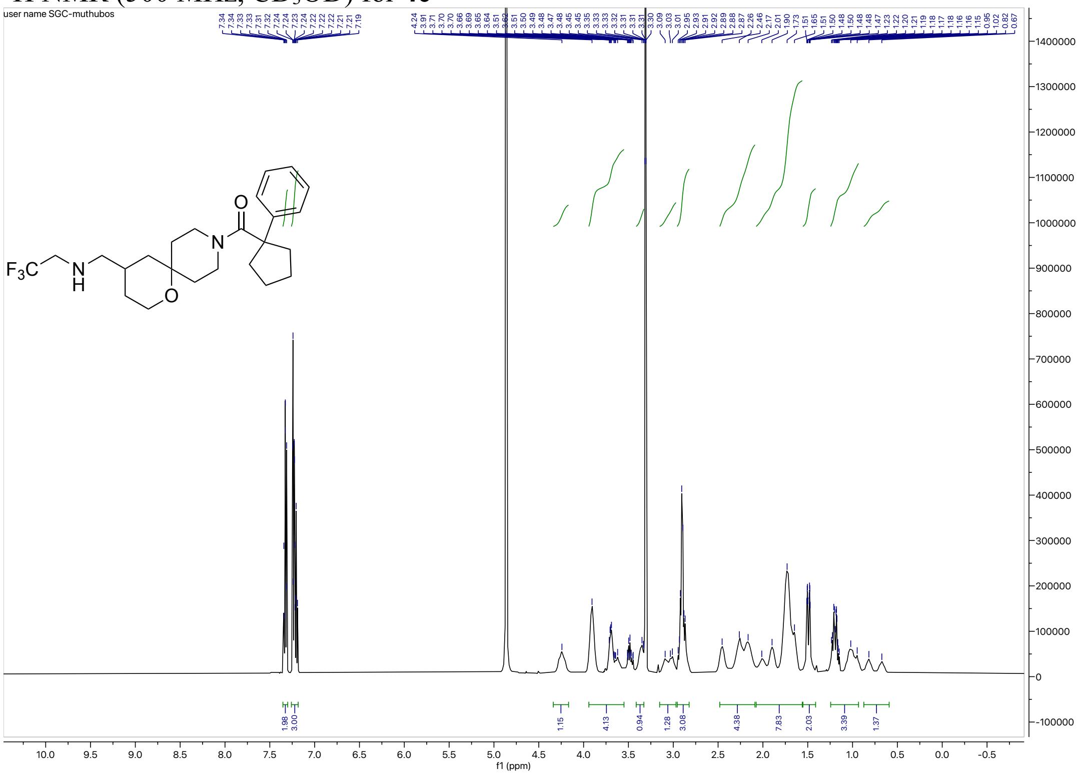

$^{13}$C NMR (126 MHz, $CD_3OD$) for **4c**

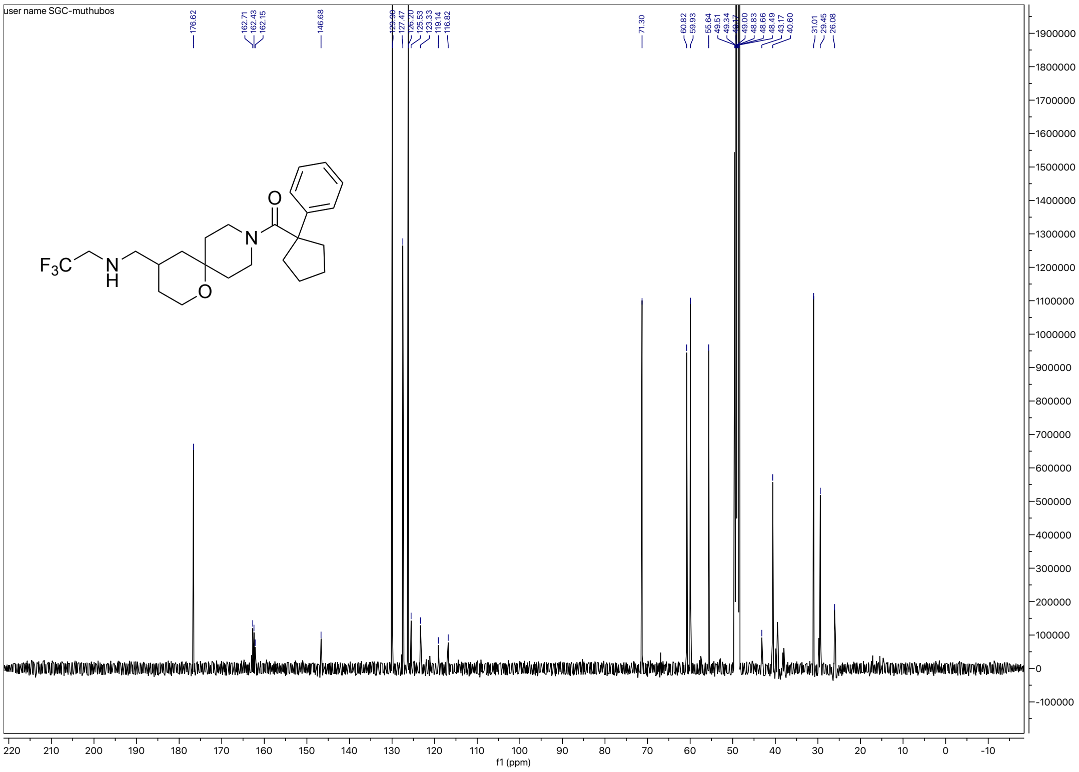

### HRMS for **4c**

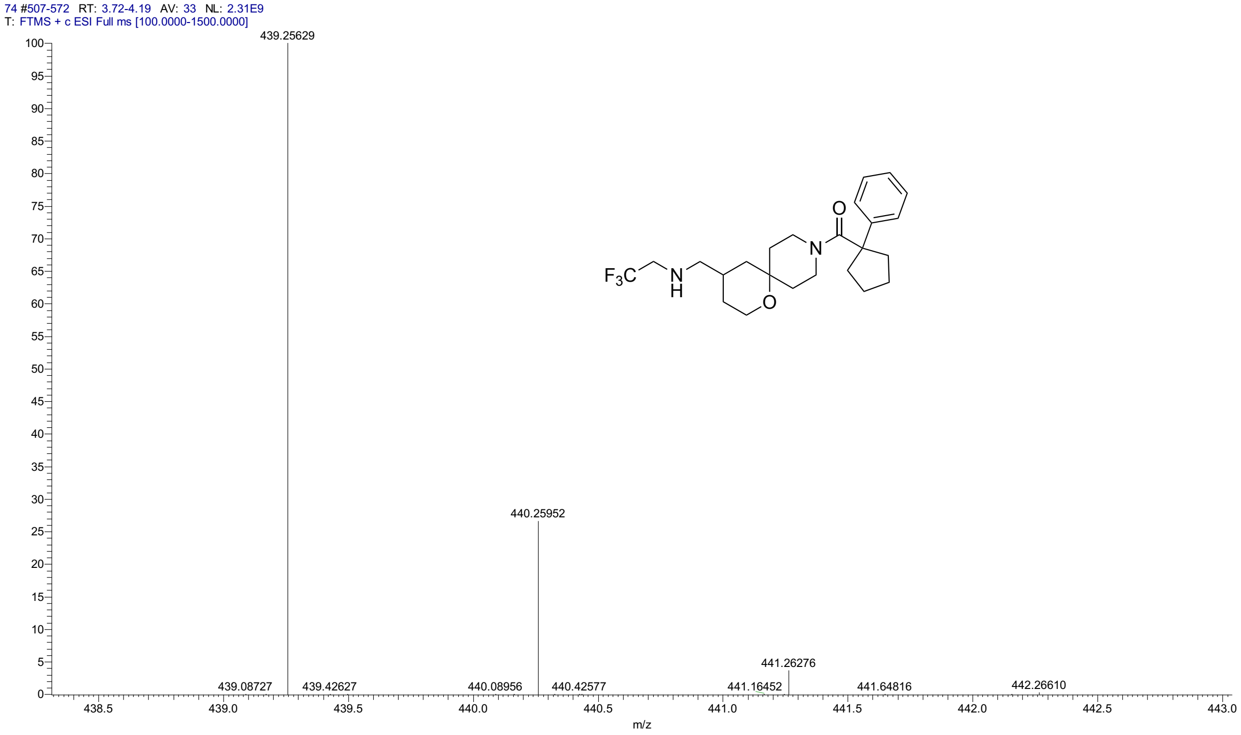


### HPLC for **4c**

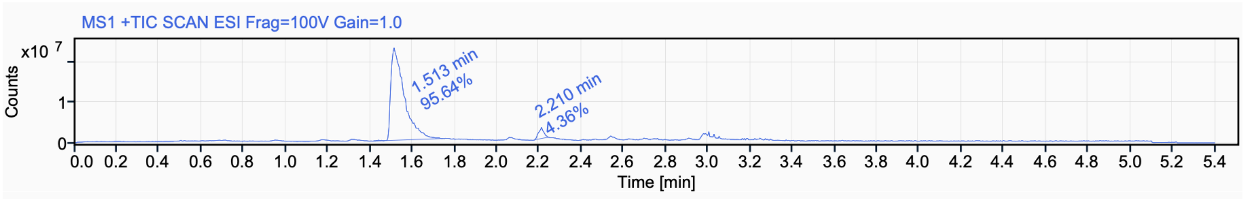


Signal Name MS1 +TIC SCAN ESI Frag=100V Gain=1.0

| RT (min) | Signal description | m/z | Purity UV(‰) / MS(%) | Area | Area% |
|---|---|---|---|---|---|
| 1.513 | MS1 +TIC SCAN ESI Frag=100V Gain=1.0 | | | 101341790.2 | 95.64 |
| 2.210 | MS1 +TIC SCAN ESI Frag=100V Gain=1.0 | | | 4622229.8 | 4.36 |

[1]H NMR (500 MHz, $CD_3CN$) for **4d**

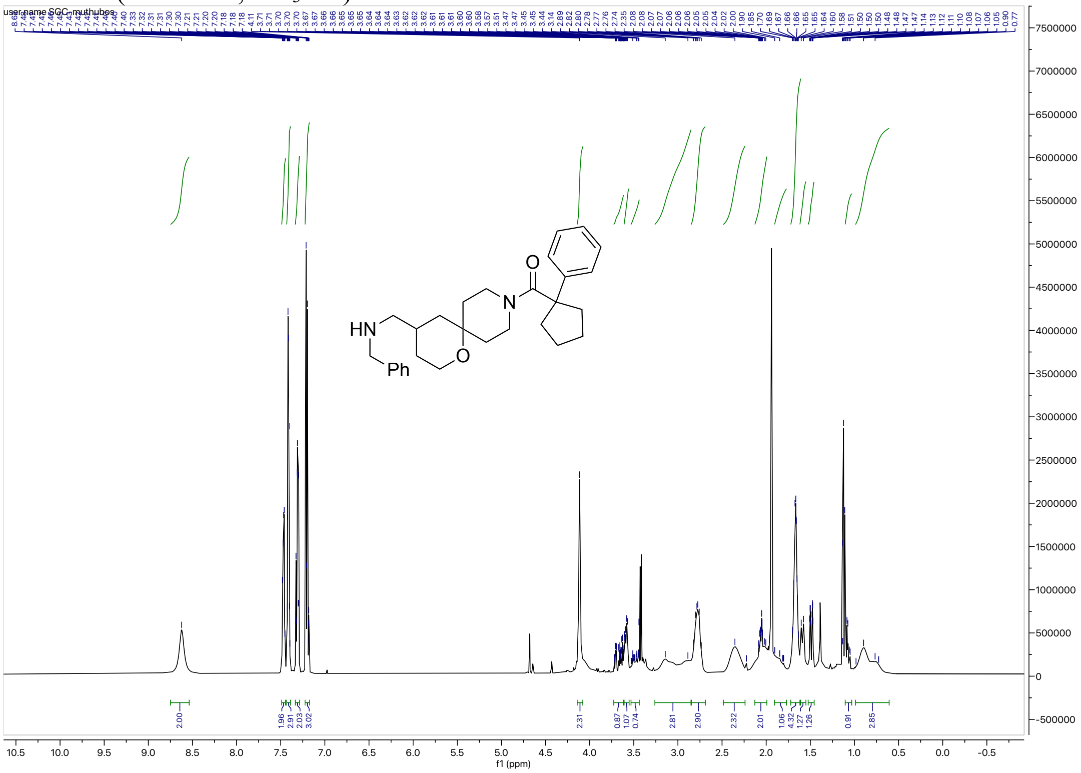

[13]C NMR (126 MHz, $CD_3CN$) for **4d**

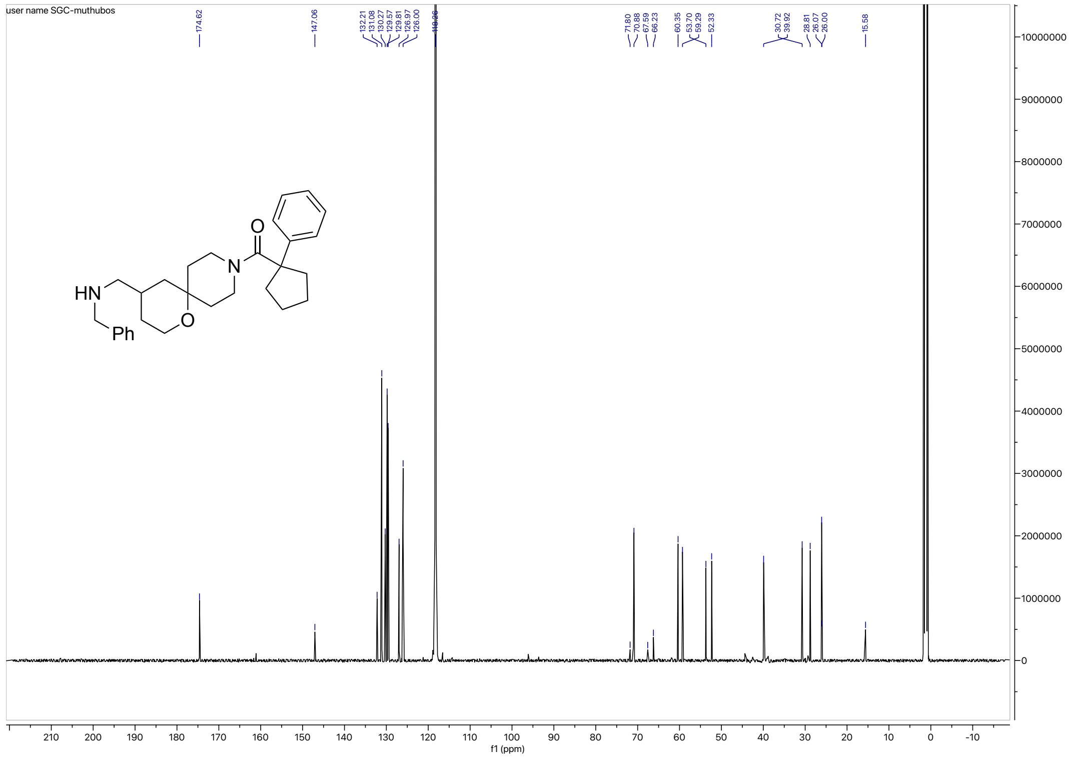

HRMS for **4d**

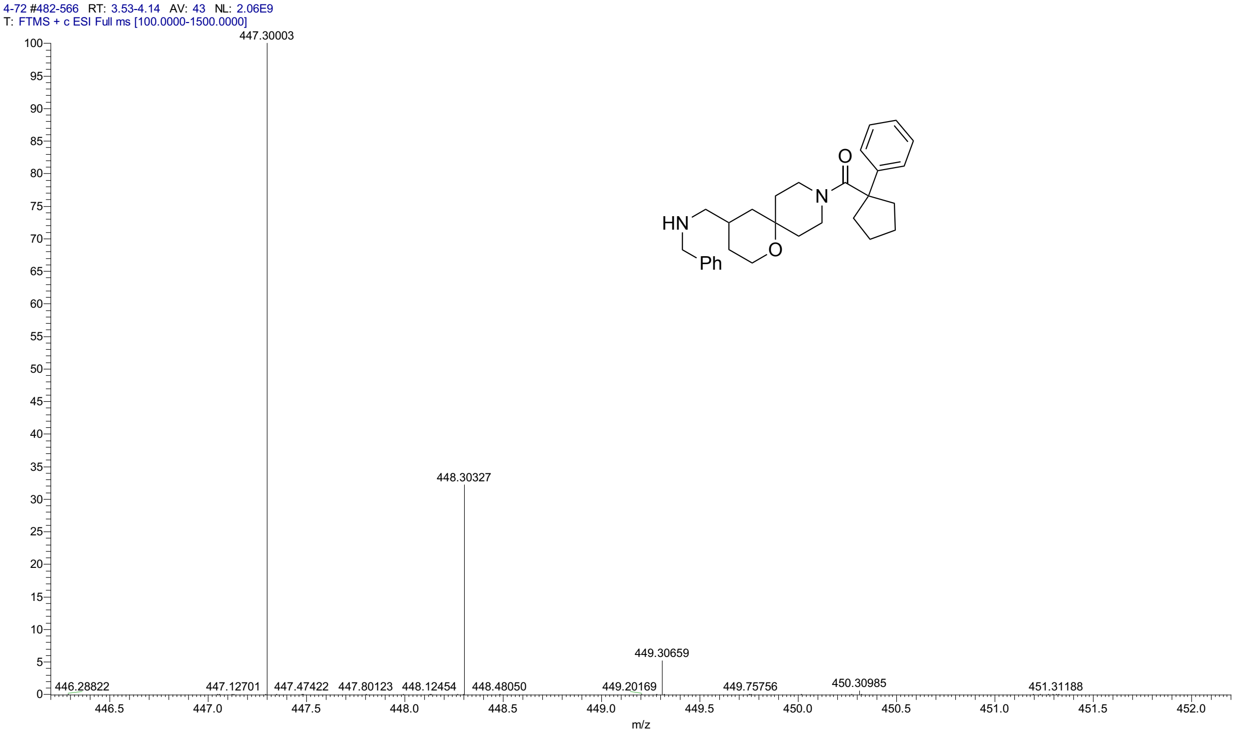


HPLC for **4d**

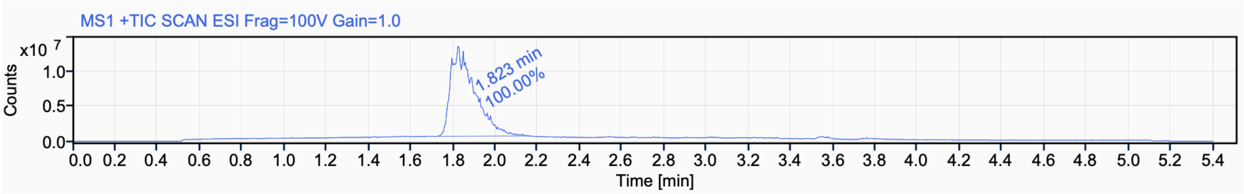


Signal Name MS1 +TIC SCAN ESI Frag=100V Gain=1.0

| **RT (min)** | **Signal description** | **Area** | **Area%** | **Peak Resolution USP** |
|---|---|---|---|---|
| 1.823 | MS1 +TIC SCAN ESI Frag=100V Gain=1.0 | 102452112.6 | 100.00 | |

$^{1}H$ NMR (500 MHz, $CD_3CN$) for **4e**

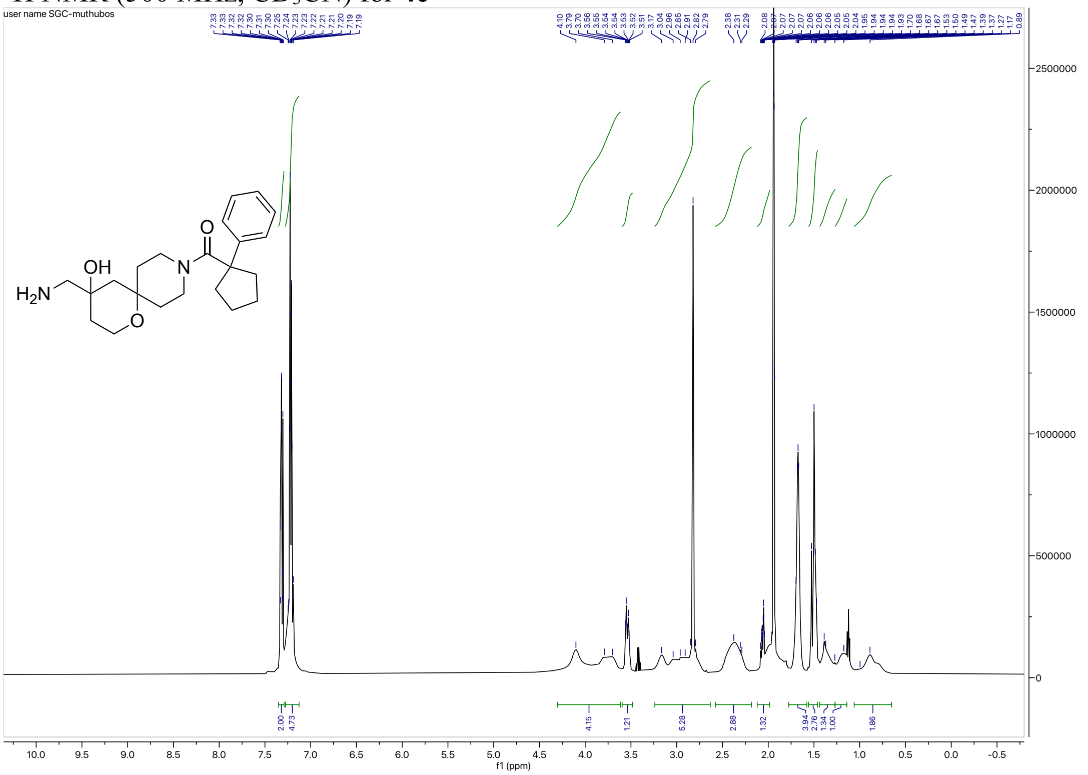


$^{13}C$ NMR (126 MHz, $CD_3CN$) for **4e**

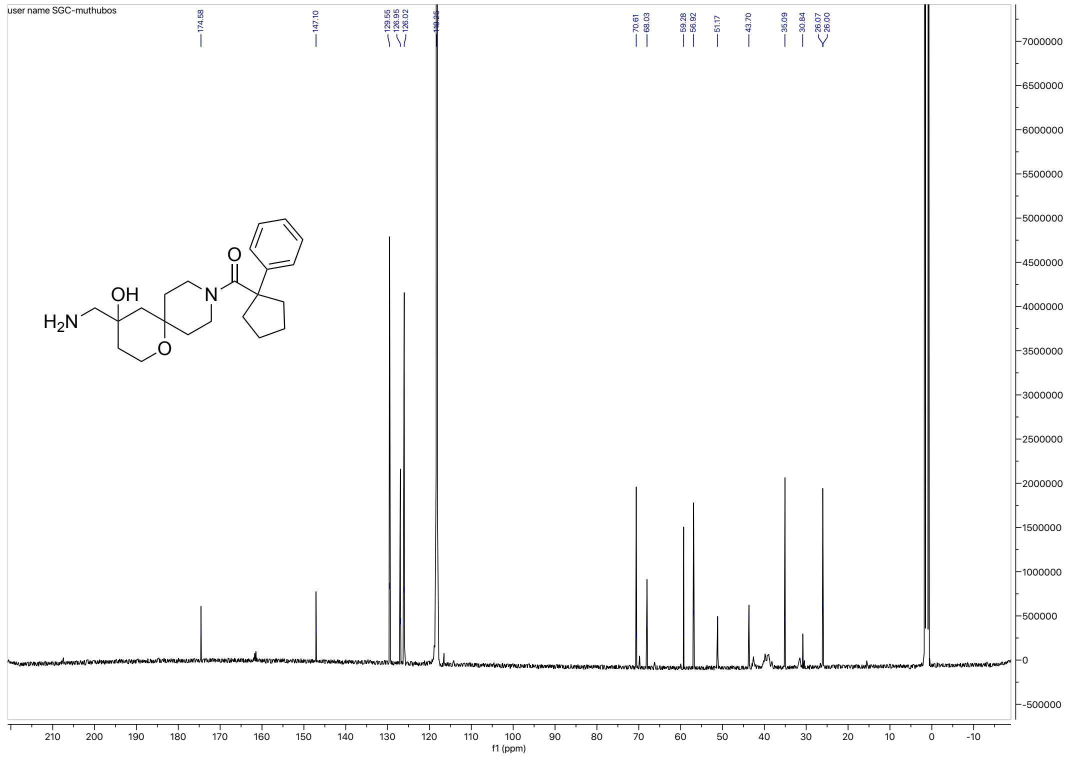

HRMS for **4e**

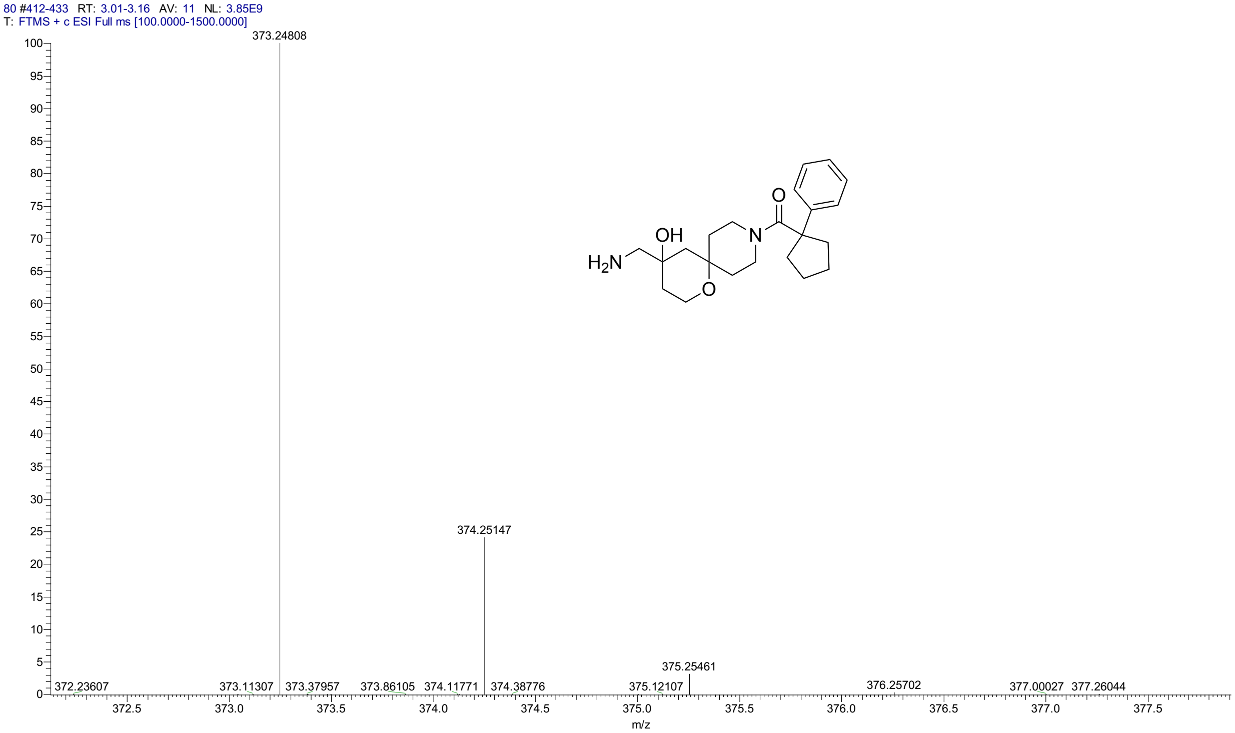


HPLC for **4e**

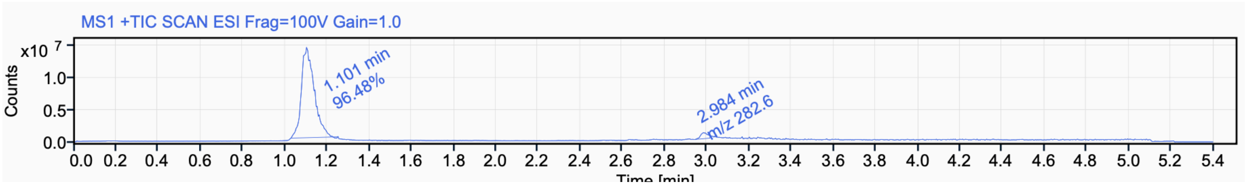

$^{1}$H NMR (500 MHz, $CD_3CN$) for **5a**

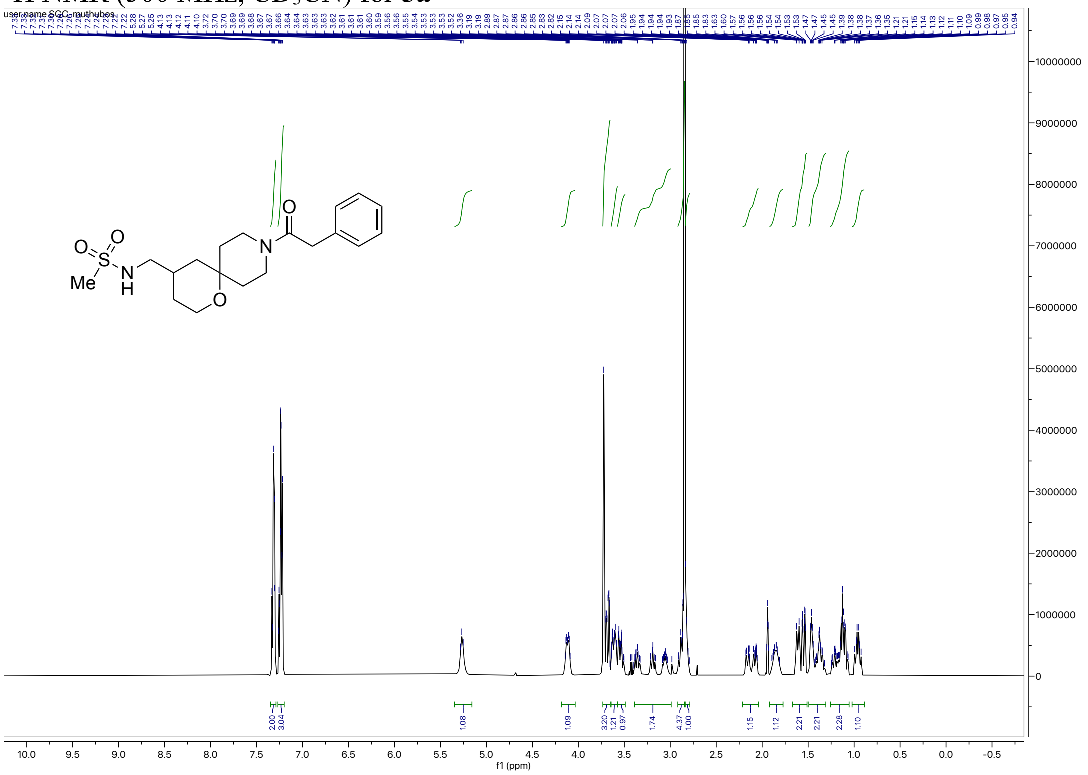

$^{13}$C NMR (126 MHz, $CD_3CN$) for **5a**

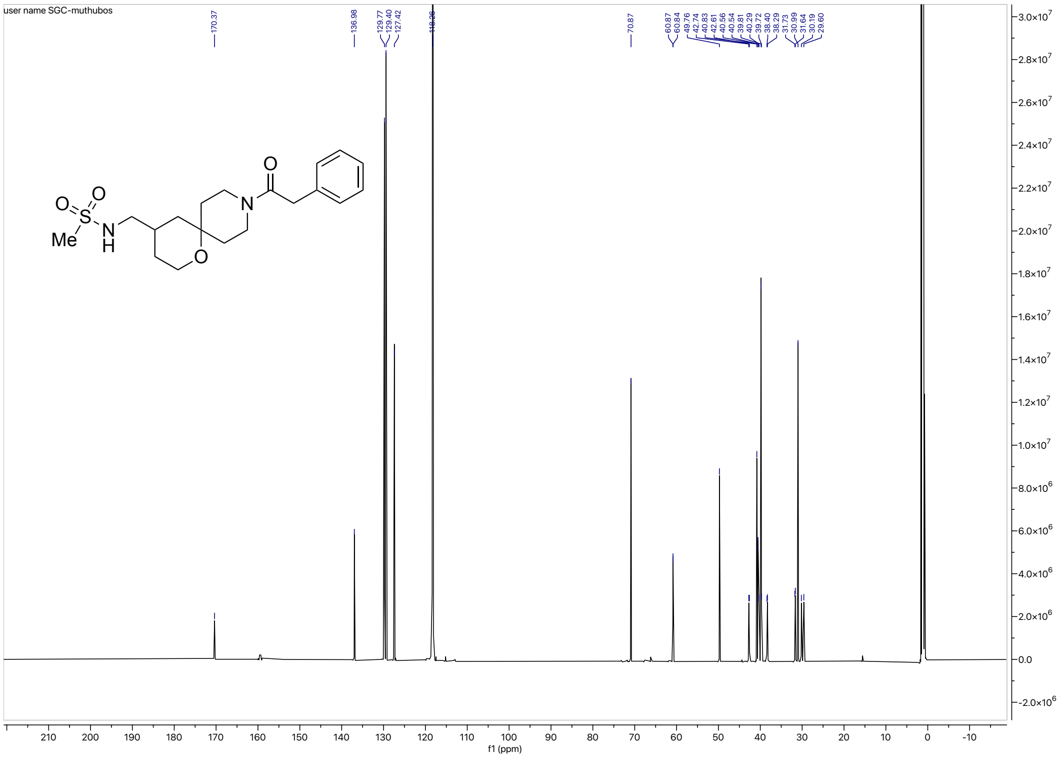

HRMS for **5a**

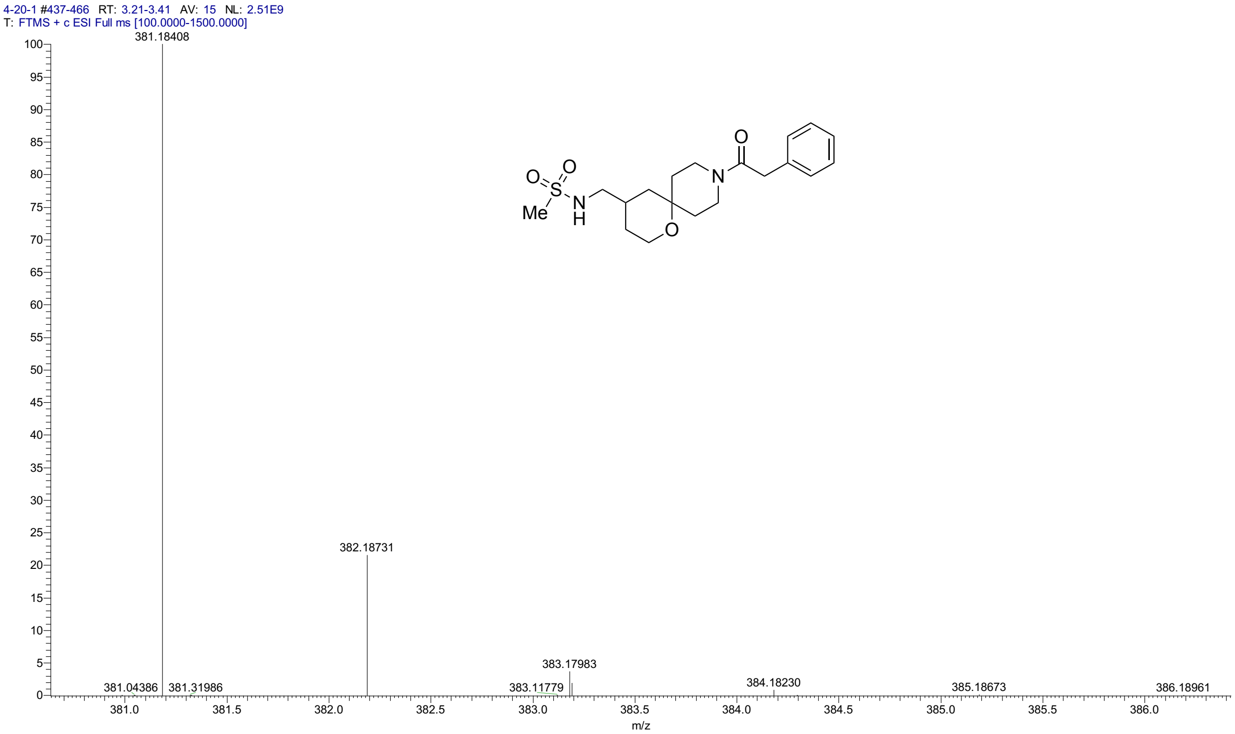


HPLC for **5a**

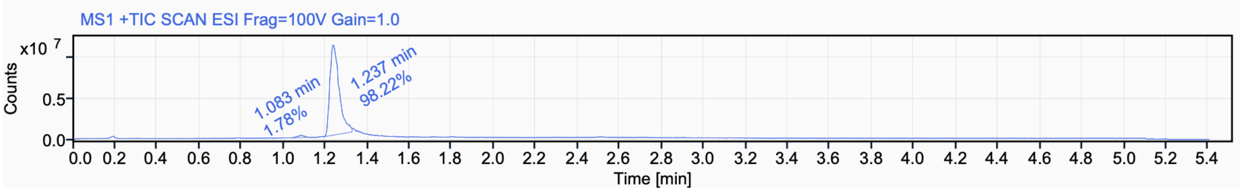


Signal Name MS1 +TIC SCAN ESI Frag=100V Gain=1.0

| RT (min) | Signal description | m/z | Purity UV(‰) / MS(%) | Area | Area% |
|---|---|---|---|---|---|
| 1.083 | MS1 +TIC SCAN ESI Frag=100V Gain=1.0 | | | 608706.0 | 1.78 |
| 1.237 | MS1 +TIC SCAN ESI Frag=100V Gain=1.0 | | | 33661561.3 | 98.22 |

$^{1}H$ NMR (500 MHz, $CD_3CN$) for **5b**

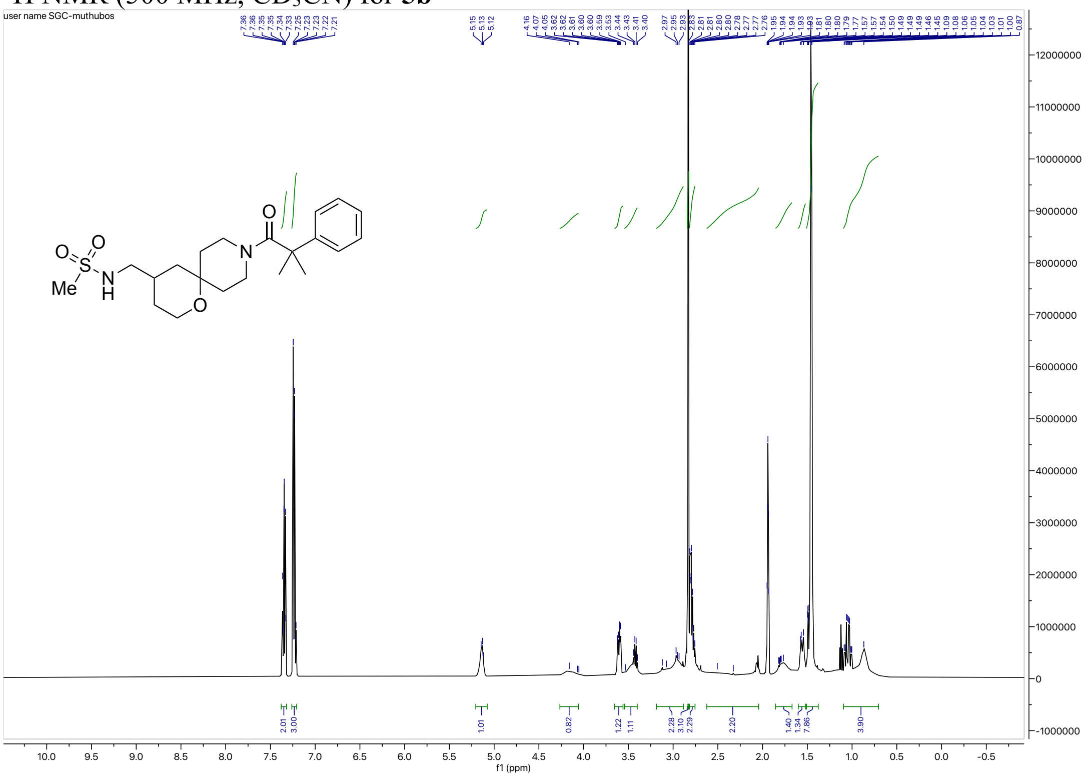

$^{13}C$ NMR (126 MHz, $CD_3CN$) for **5b**

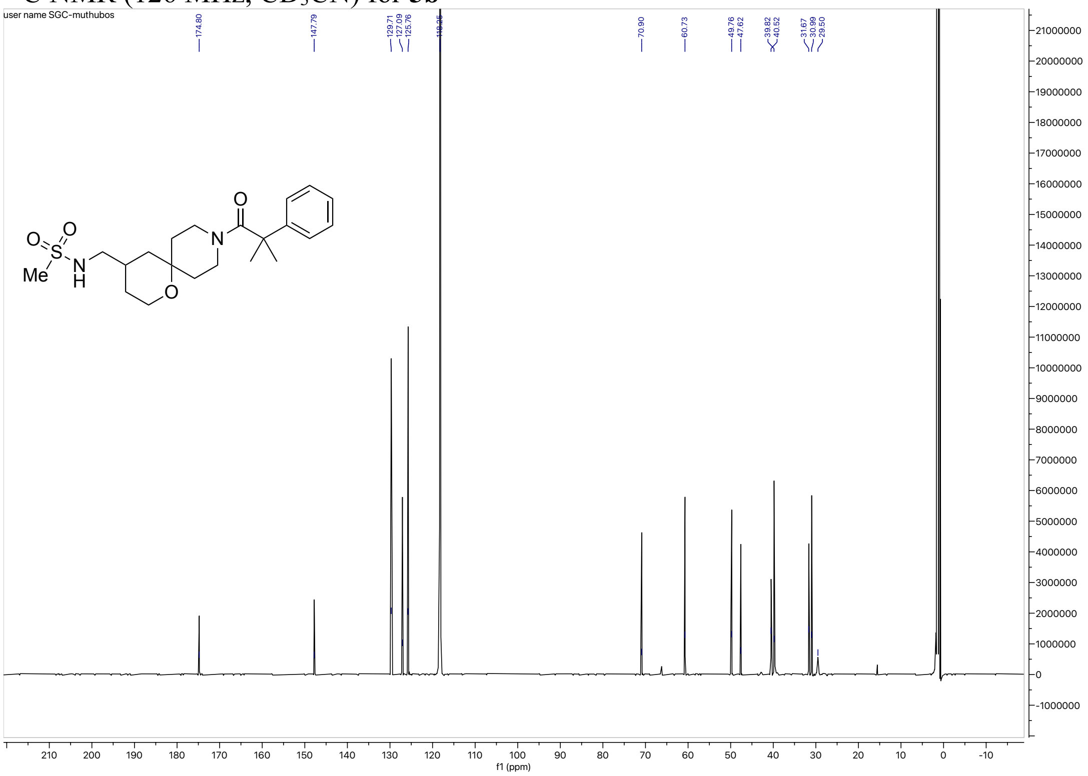

HRMS for **5b**

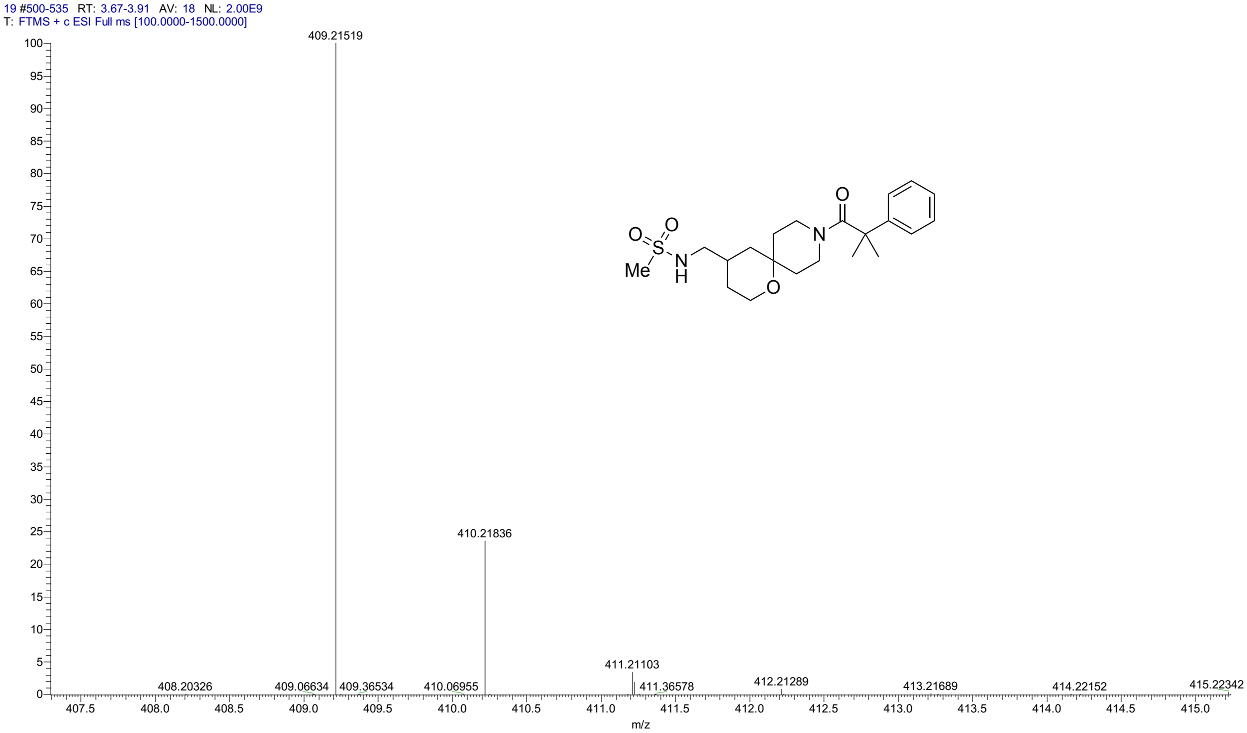


HPLC for **5b**

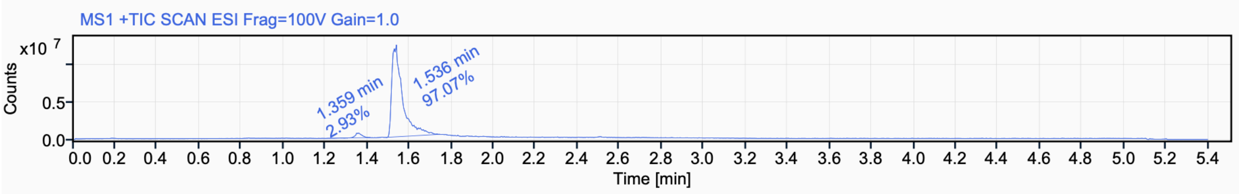


Signal Name MS1 +TIC SCAN ESI Frag=100V Gain=1.0

| RT (min) | Signal description | m/z | Purity UV(‰) / MS(%) | Area | Area% |
|---|---|---|---|---|---|
| 1.359 | MS1 +TIC SCAN ESI Frag=100V Gain=1.0 | | | 1325643.8 | 2.93 |
| 1.536 | MS1 +TIC SCAN ESI Frag=100V Gain=1.0 | | | 43857696.2 | 97.07 |

$^{1}$H NMR (500 MHz, $CD_3CN$) for **5c**

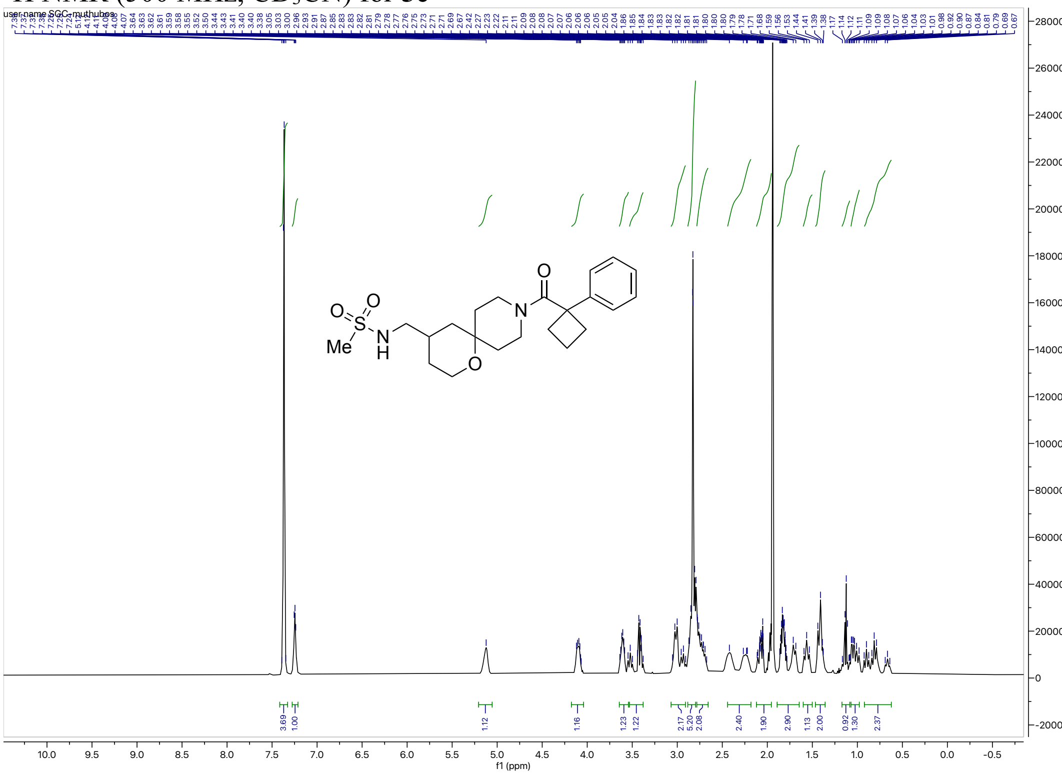


$^{13}$C NMR (126 MHz, $CD_3CN$) for **5c**

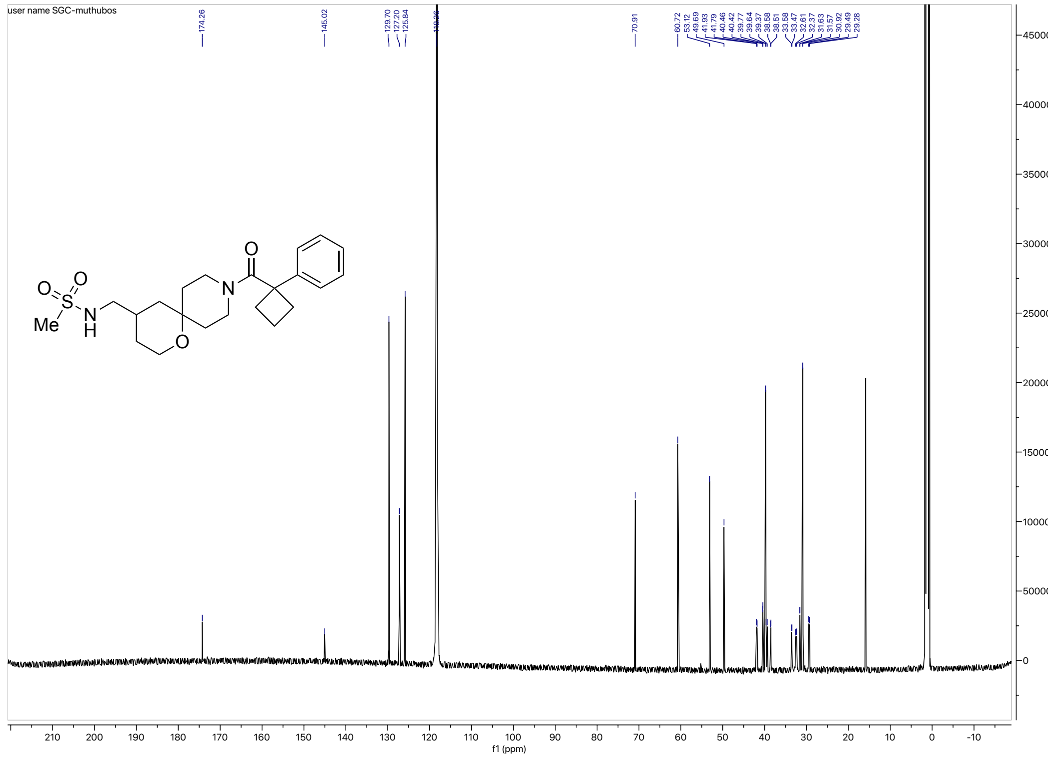

HRMS for **5c**

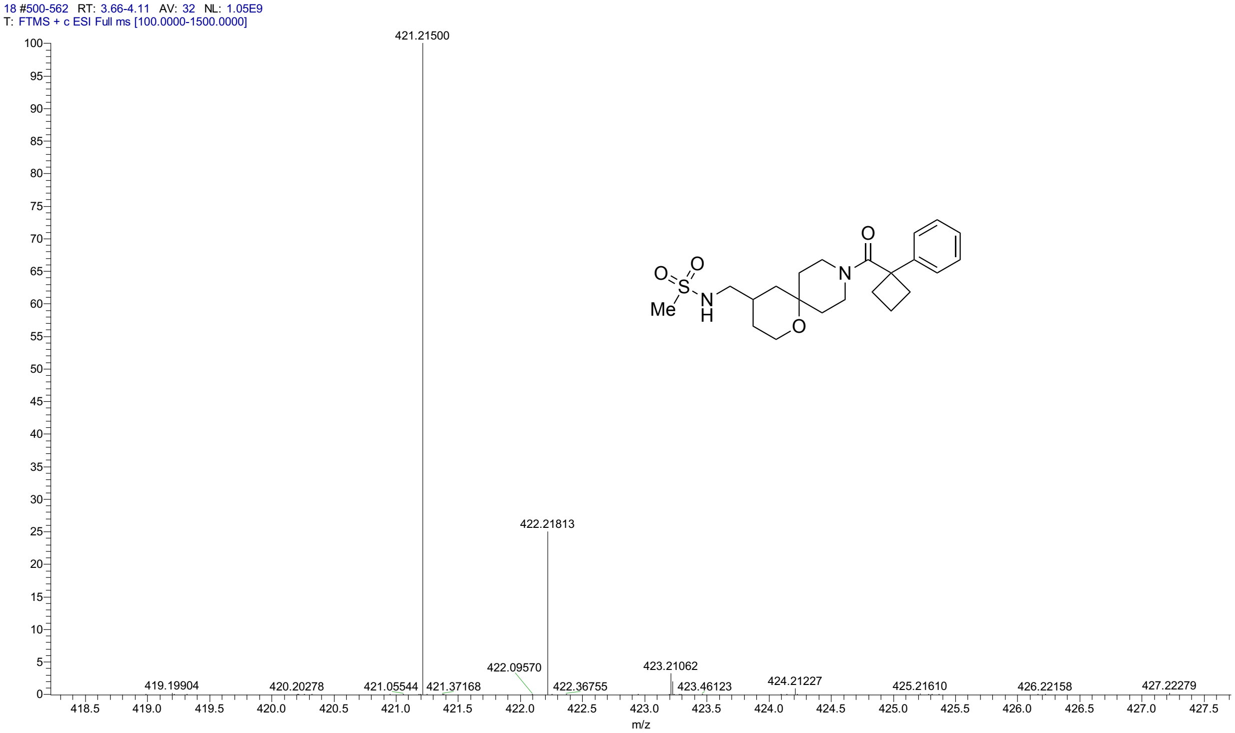


HPLC for **5c**

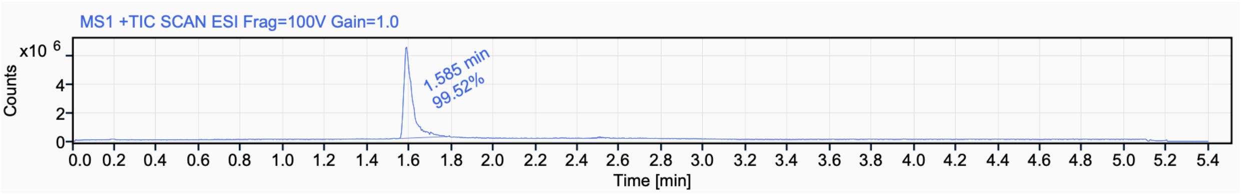


Signal Name MS1 +TIC SCAN ESI Frag=100V Gain=1.0

| RT (min) | Signal description | m/z | Purity UV(‰) / MS(%) | Area | Area% |
|---|---|---|---|---|---|
| 1.585 | MS1 +TIC SCAN ESI Frag=100V Gain=1.0 | | | 18008339.9 | 99.52 |
| 2.506 | MS1 +TIC SCAN ESI Frag=100V Gain=1.0 | | | 87729.3 | 0.48 |

$^{1}$H NMR (500 MHz, $CD_3CN$) for **5d**

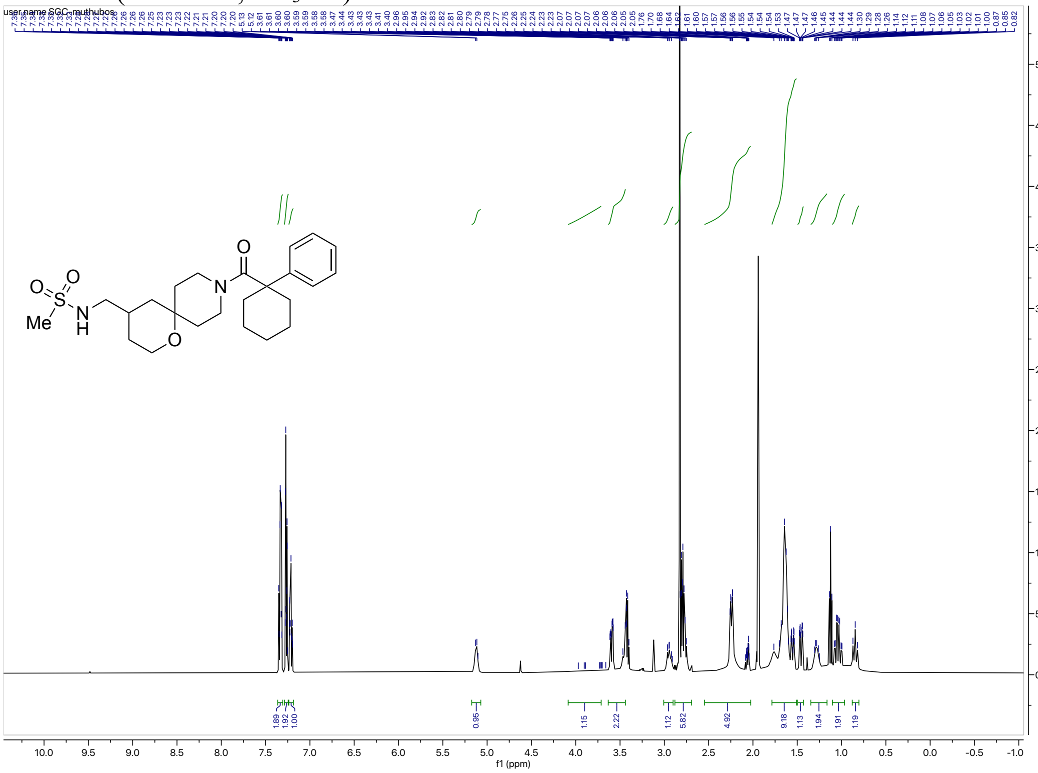

$^{13}$C NMR (126 MHz, $CD_3CN$) for **5d**

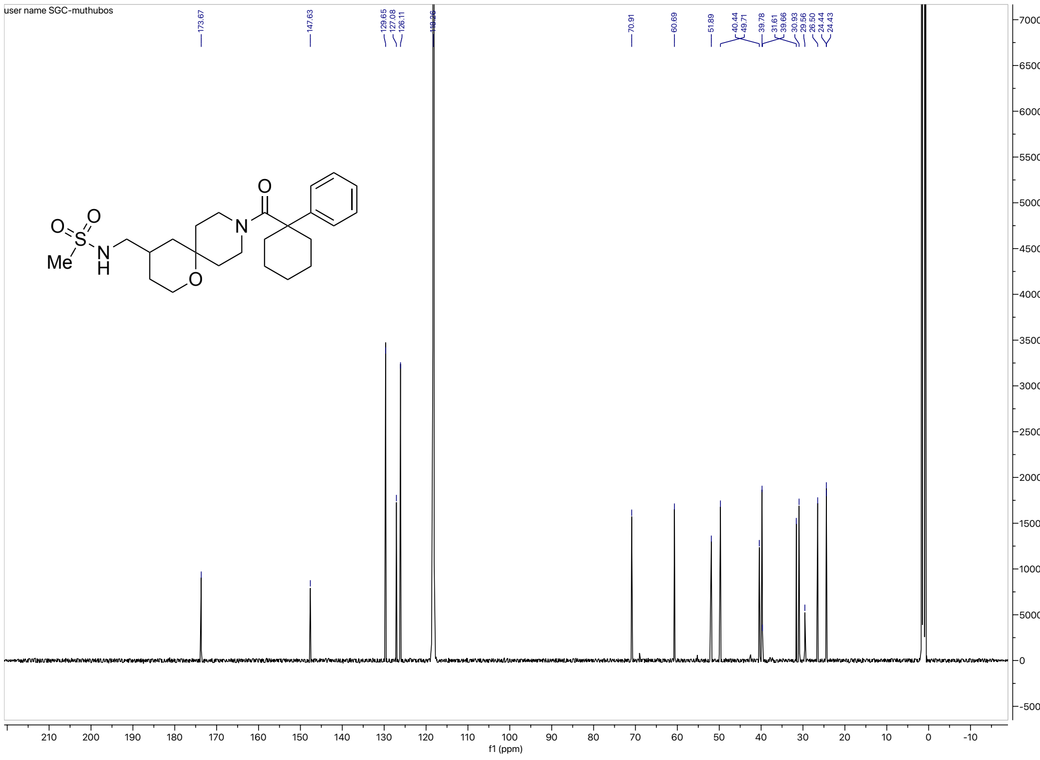

#### HRMS for **5d**

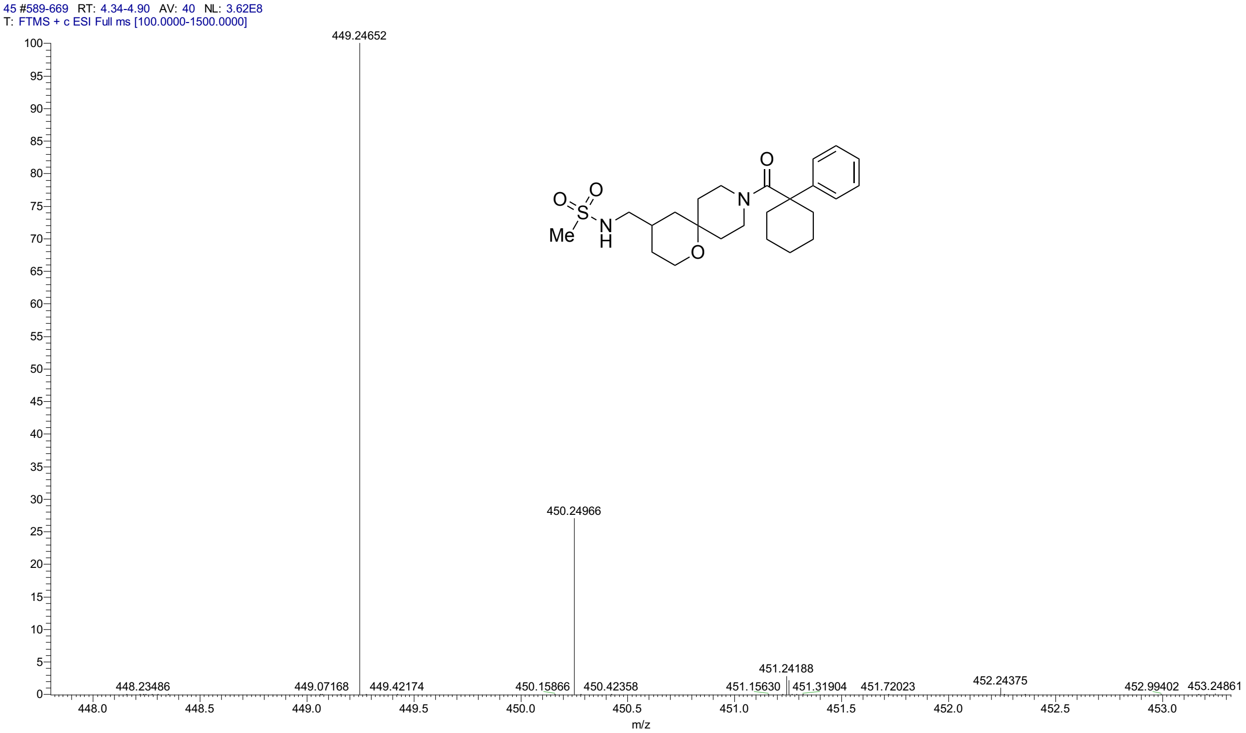


#### HPLC for **5d**

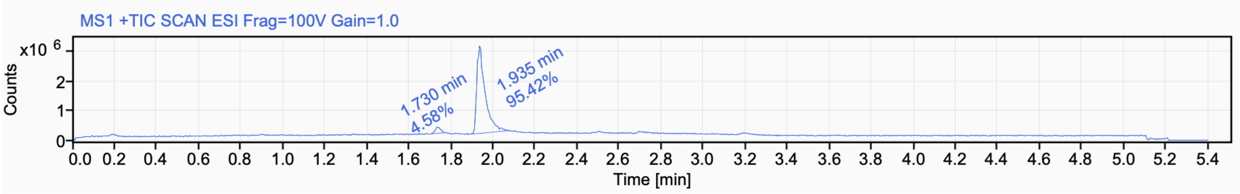


Signal Name MS1 +TIC SCAN ESI Frag=100V Gain=1.0

| RT (min) | Signal description | m/z | Purity UV(‰) / MS(%) | Area | Area% |
|---|---|---|---|---|---|
| 1.730 | MS1 +TIC SCAN ESI Frag=100V Gain=1.0 | | | 389236.6 | 4.58 |
| 1.935 | MS1 +TIC SCAN ESI Frag=100V Gain=1.0 | | | 8104444.9 | 95.42 |

$^{1}$H NMR (500 MHz, DMSO-$d_6$) for **5e**

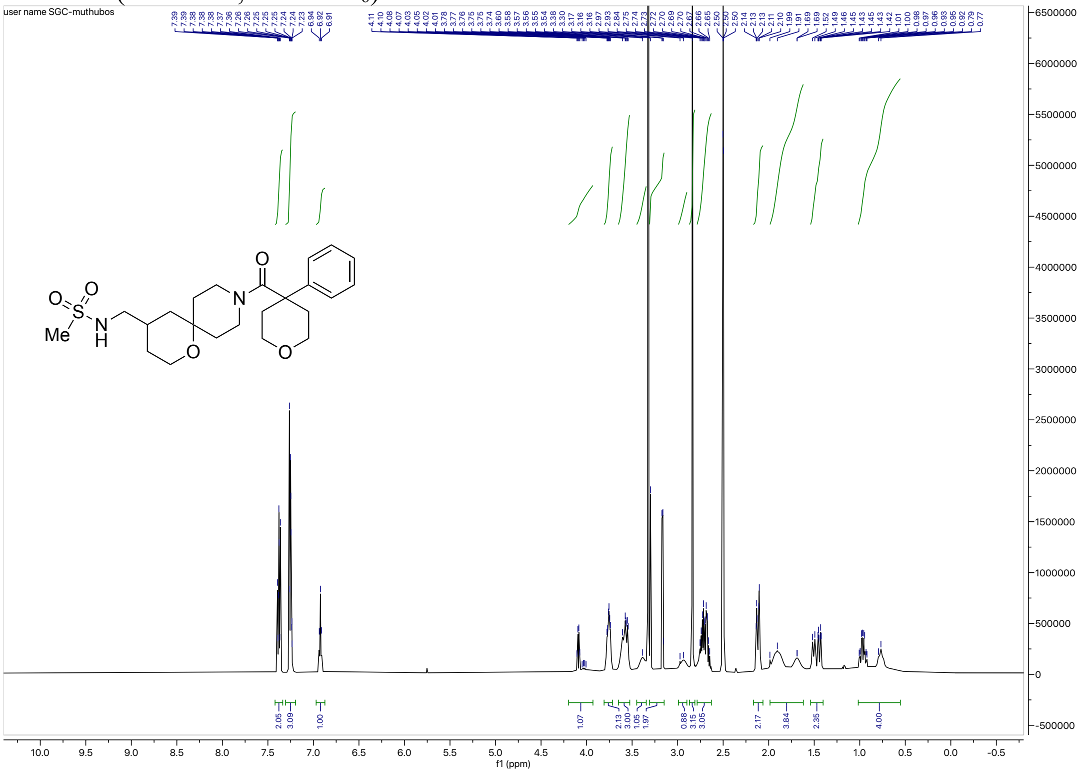

$^{13}$C NMR (126 MHz, DMSO-$d_6$) for **5e**

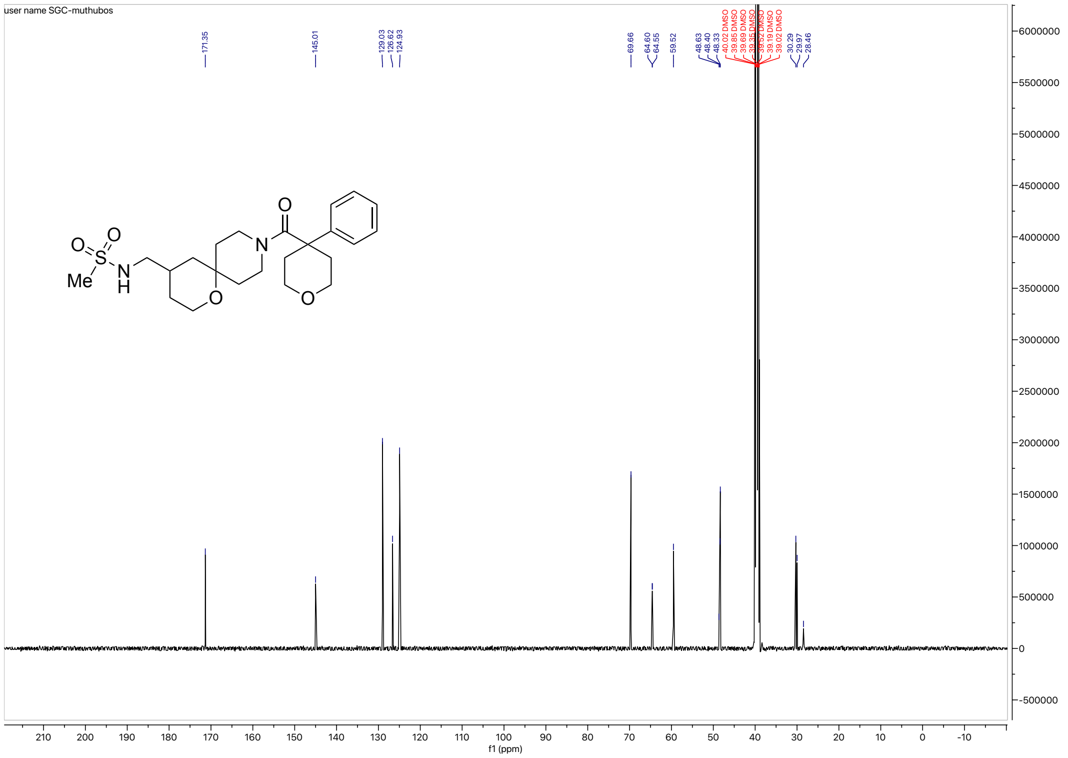

#### HRMS for **5e**

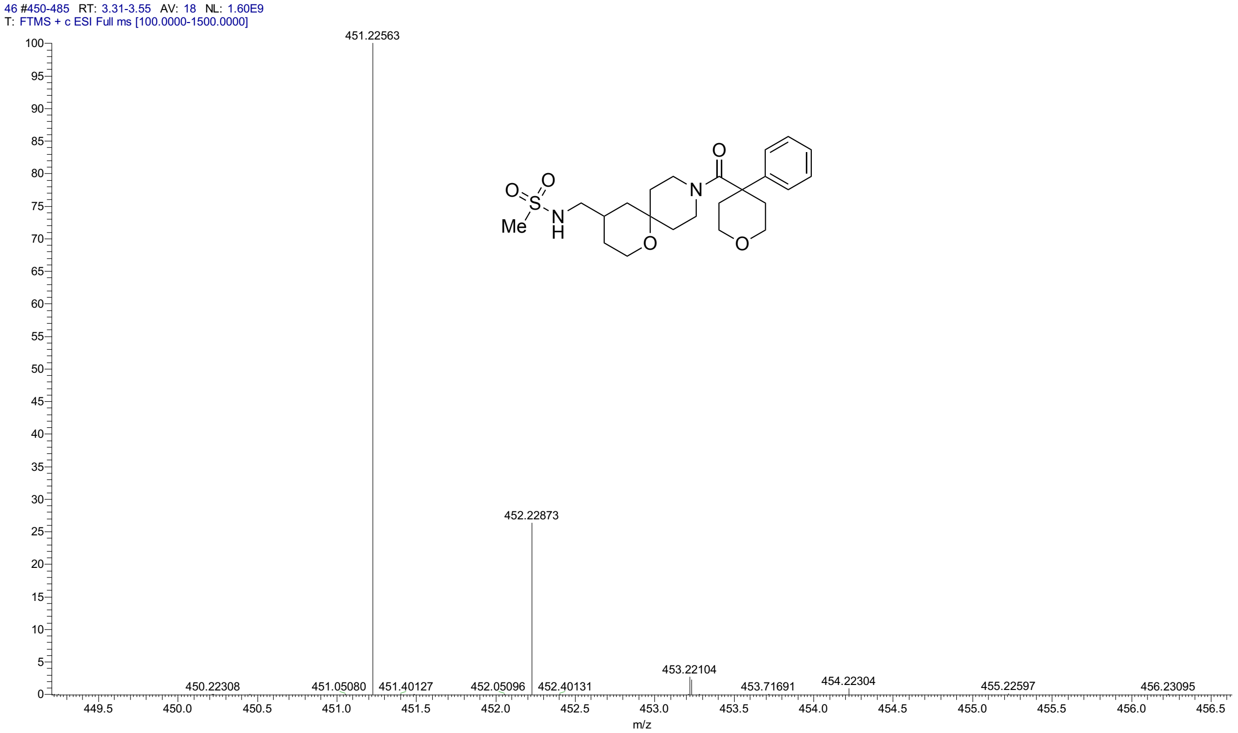


#### HPLC for **5e**

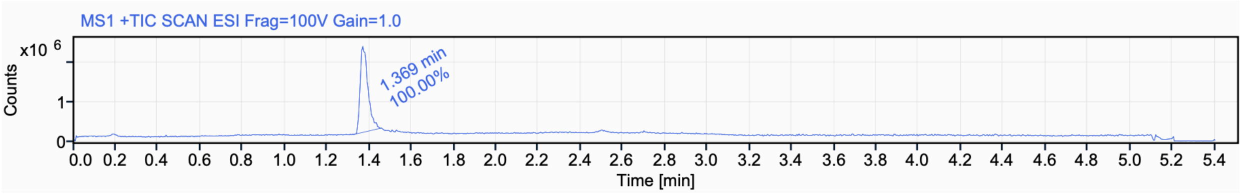


Signal Name MS1 +TIC SCAN ESI Frag=100V Gain=1.0

| RT (min) | Signal description | m/z | Purity UV(‰) / MS(%) | Area | Area% |
|---|---|---|---|---|---|
| 1.369 | MS1 +TIC SCAN ESI Frag=100V Gain=1.0 | | | 5581239.9 | 100.00 |

#### $^{1}$H NMR (400 MHz, DMSO, 80 °C) for **5f**

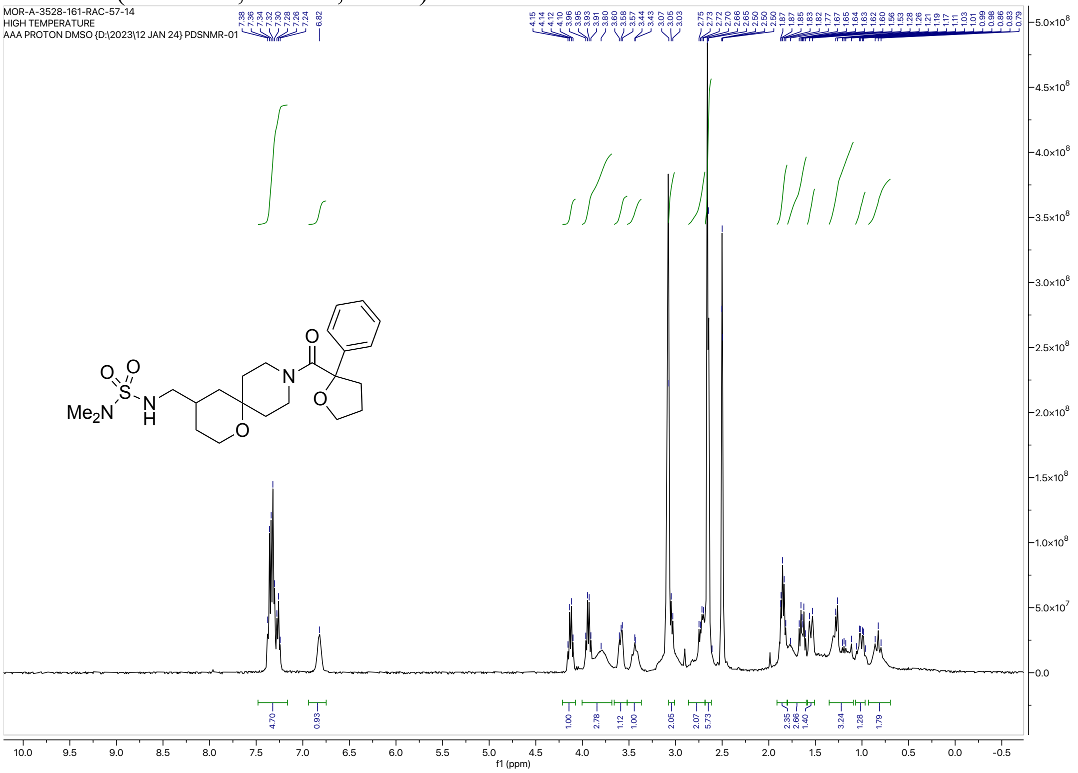


#### $^{13}$C NMR (101 MHz, DMSO-$d_6$) for **5f**

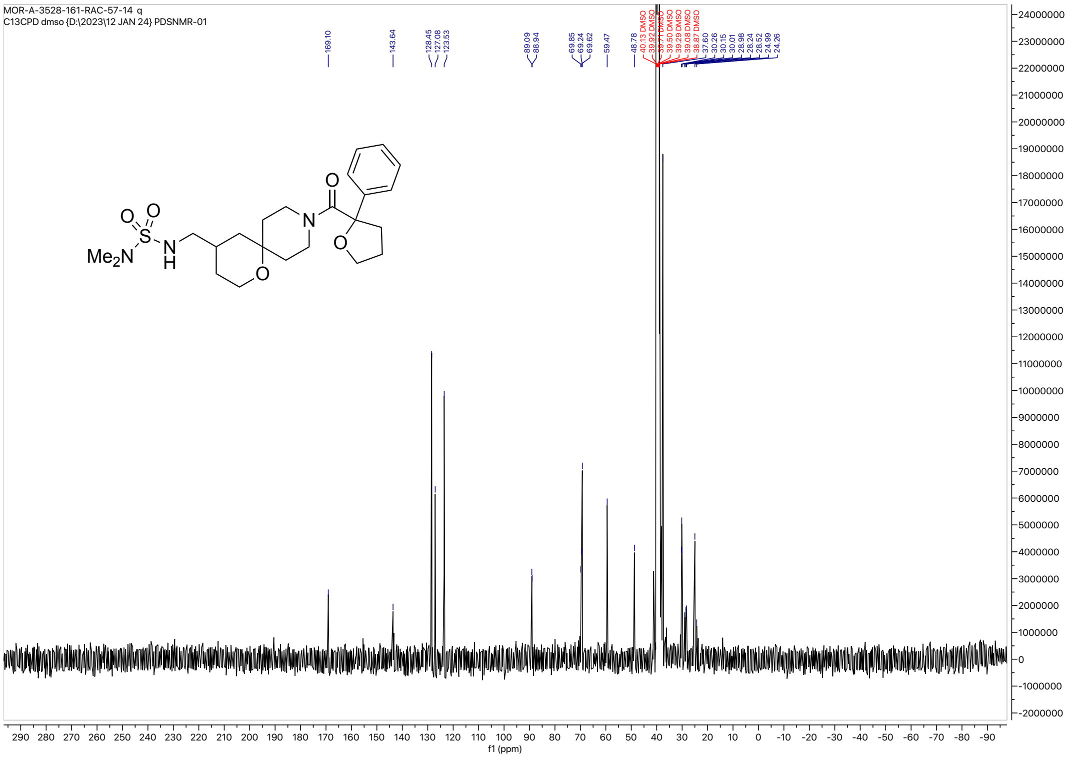

HRMS for **5f**

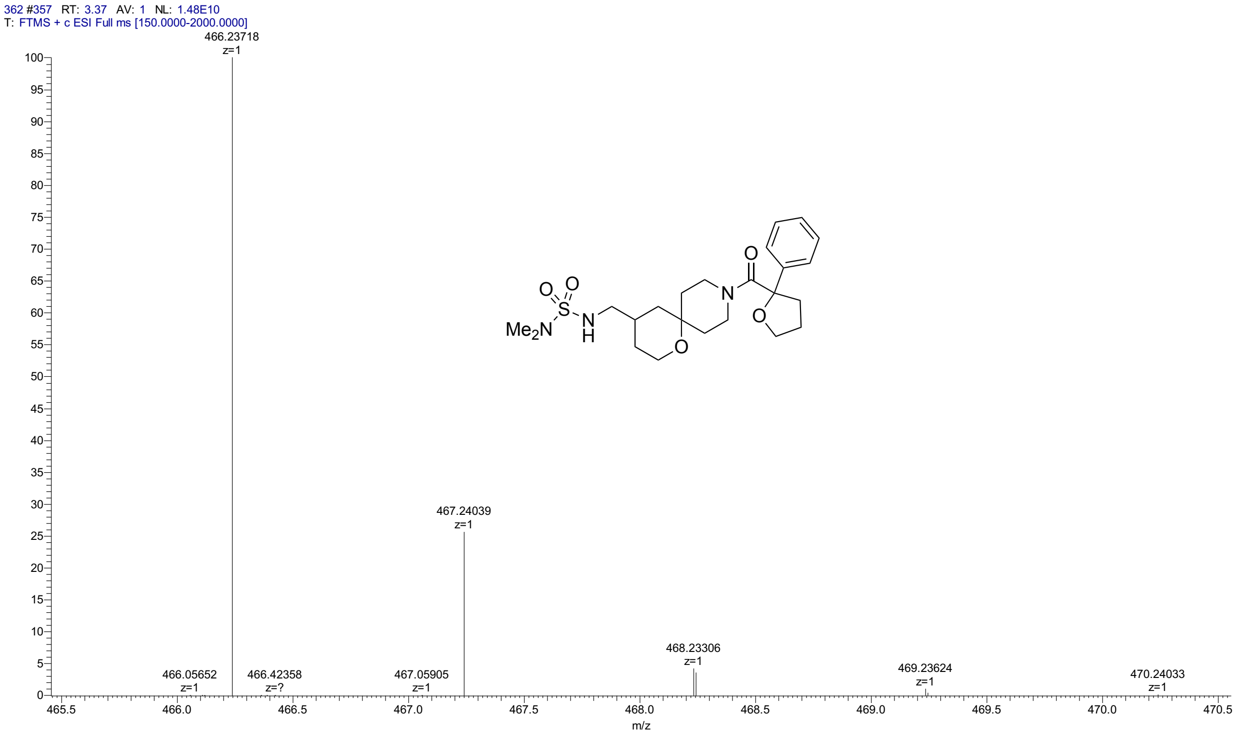


HPLC for **5f**

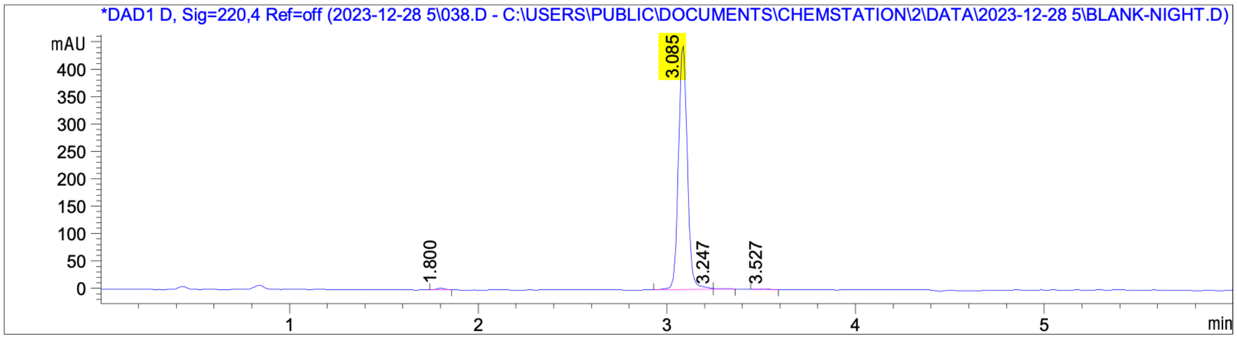


| Peak No | Height | RT (min) | Area | Area % |
|---|---|---|---|---|
| 1 | 3.360 | 1.800 | 10 | 0.64 |
| 2 | 446.661 | 3.085 | 1475 | 98.37 |
| 3 | 1.783 | 3.247 | 8 | 0.51 |
| 4 | 1.957 | 3.527 | 7 | 0.48 |

$^{1}$H NMR (400 MHz, $CDCl_3$, 45 °C) for **5g**

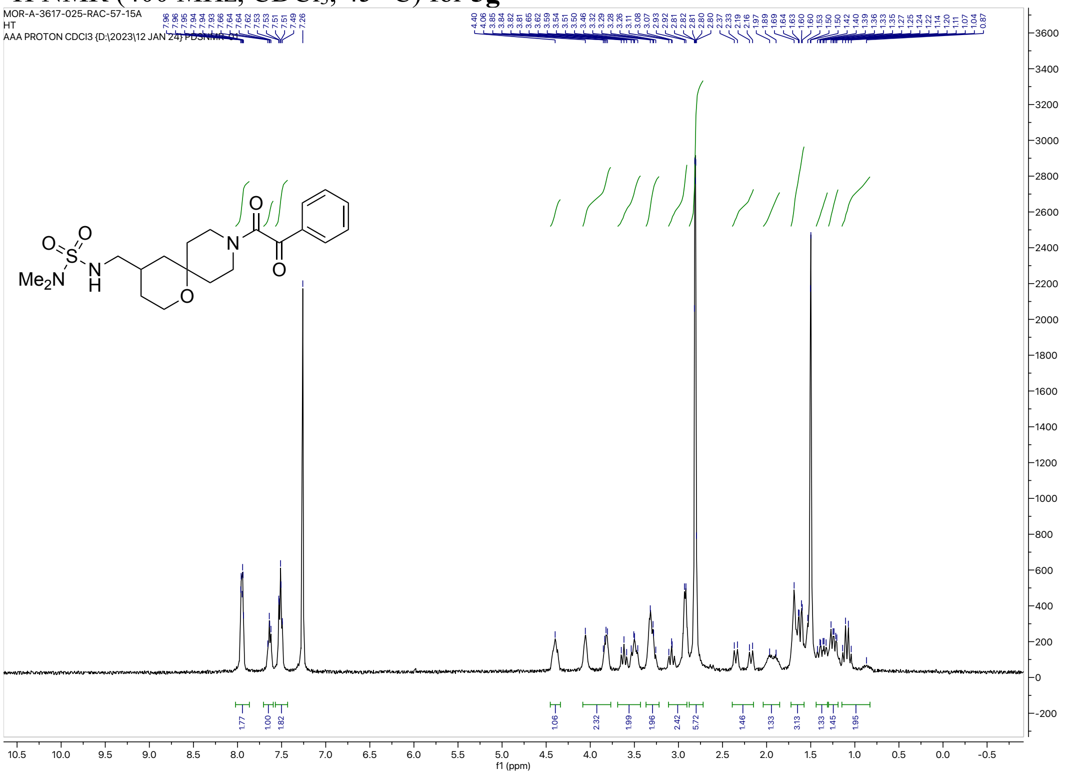

$^{13}$C NMR (126 MHz, DMSO-$d_6$) for **5g**

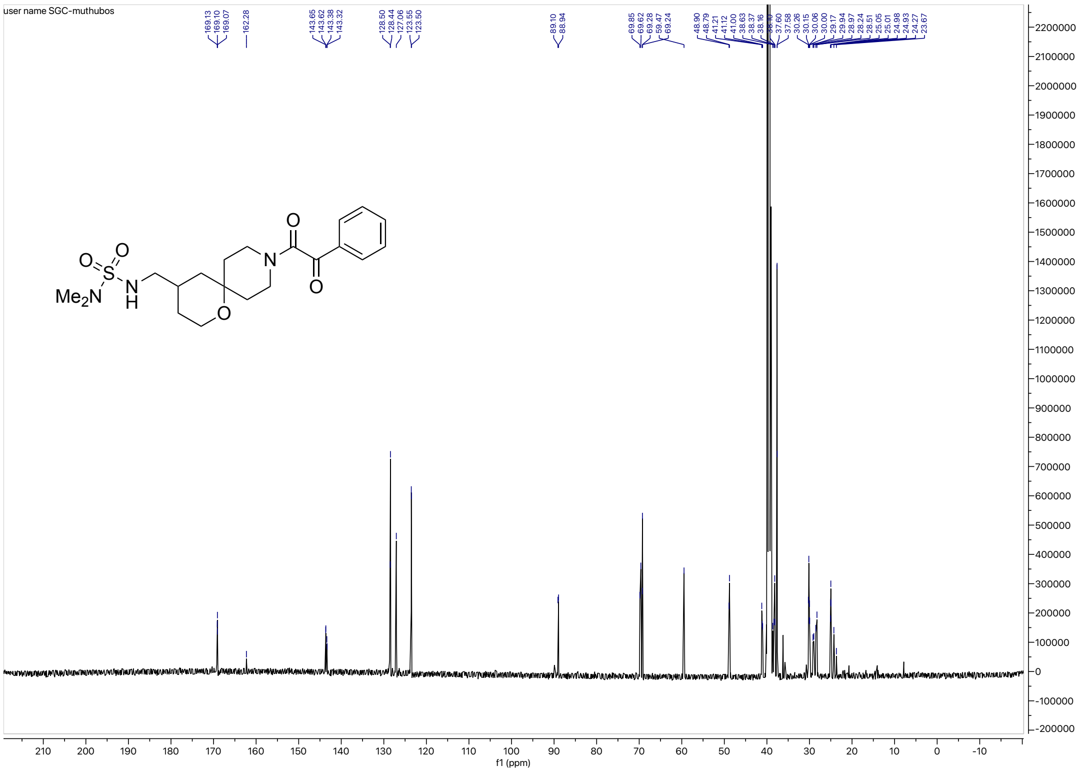

HRMS for **5g**

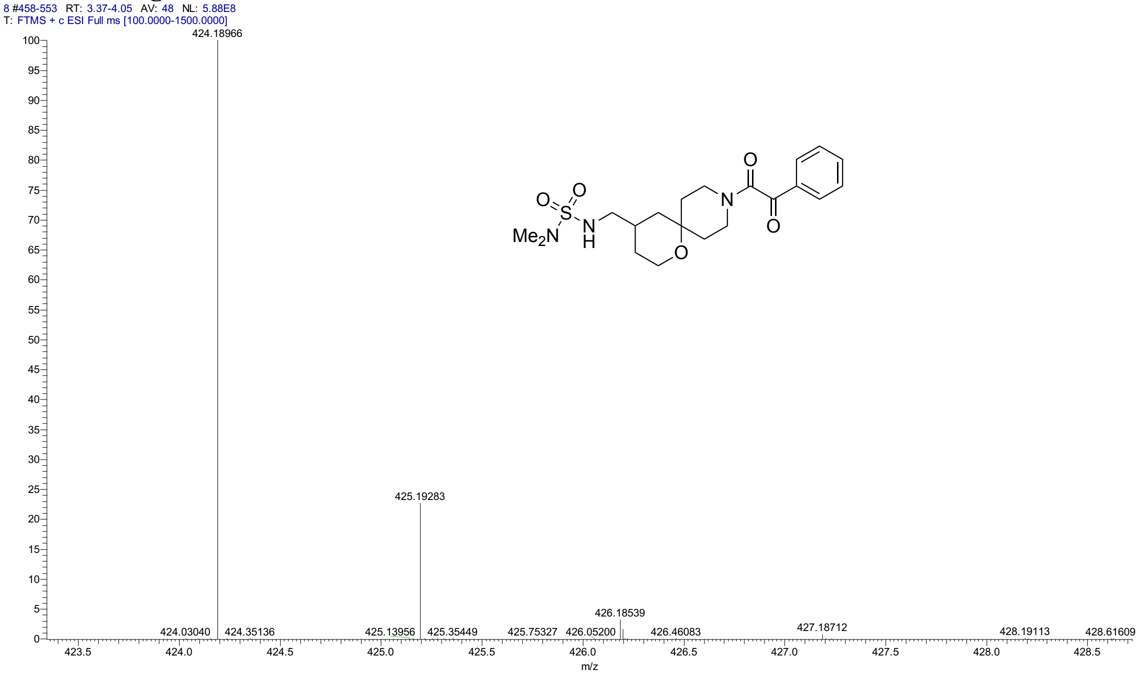


HPLC for **5g**

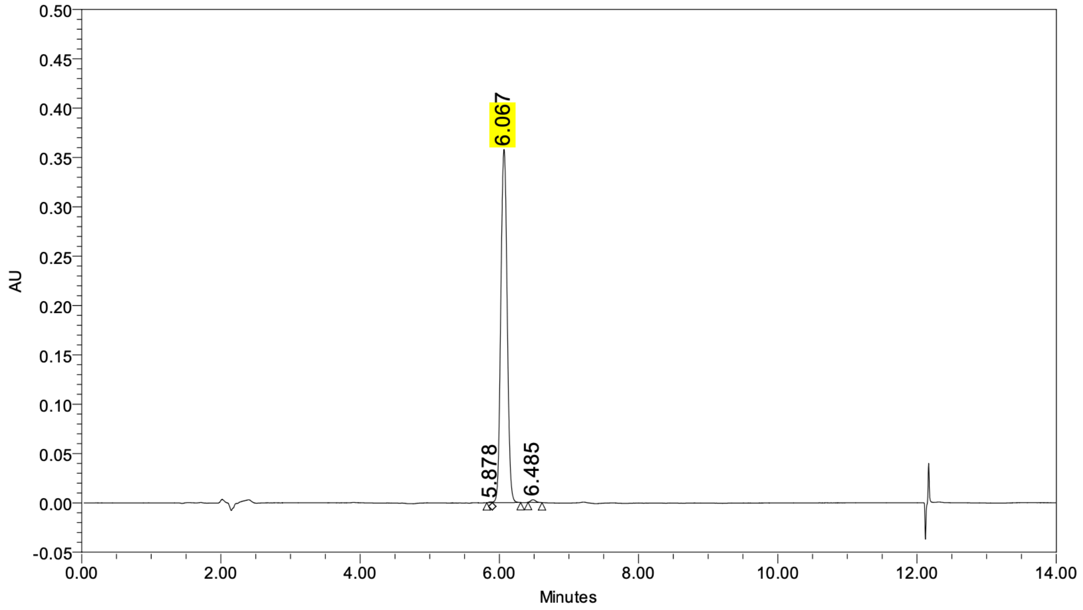


**Peak Results**

| | Name | RT | Height (µV) | Area (µV*sec) | % Area |
|---|---|---|---|---|---|
| 1 | | 5.878 | 859 | 2592 | 0.11 |
| 2 | | 6.067 | 358082 | 2271545 | 99.30 |
| 3 | | 6.485 | 2691 | 13449 | 0.59 |

[1]H NMR (500 MHz, $CD_3CN$) for **6a**

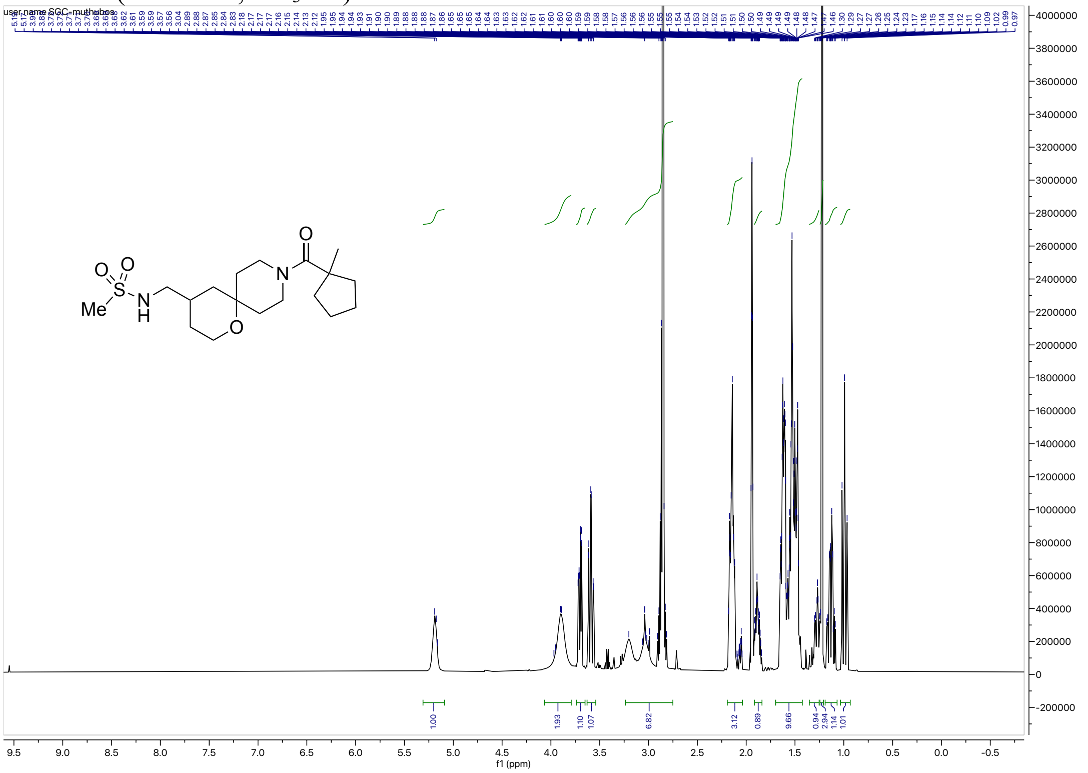


[13]C NMR (126 MHz, $CD_3CN$) for **6a**

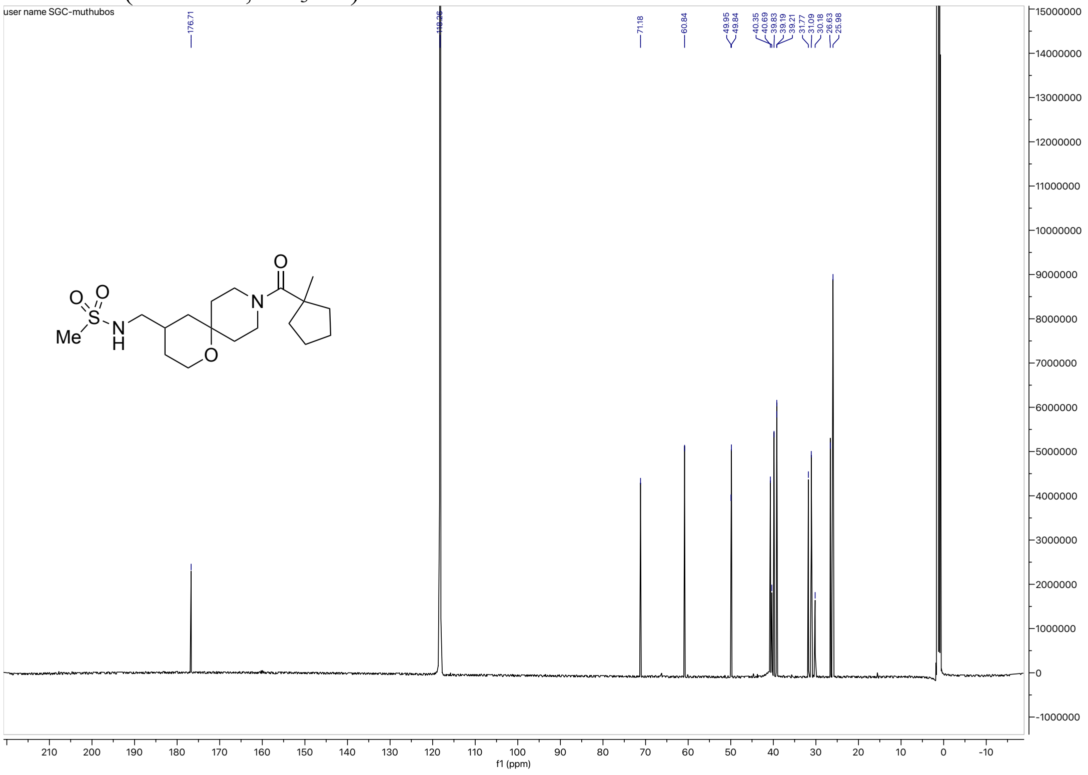

#### HRMS for **6a**

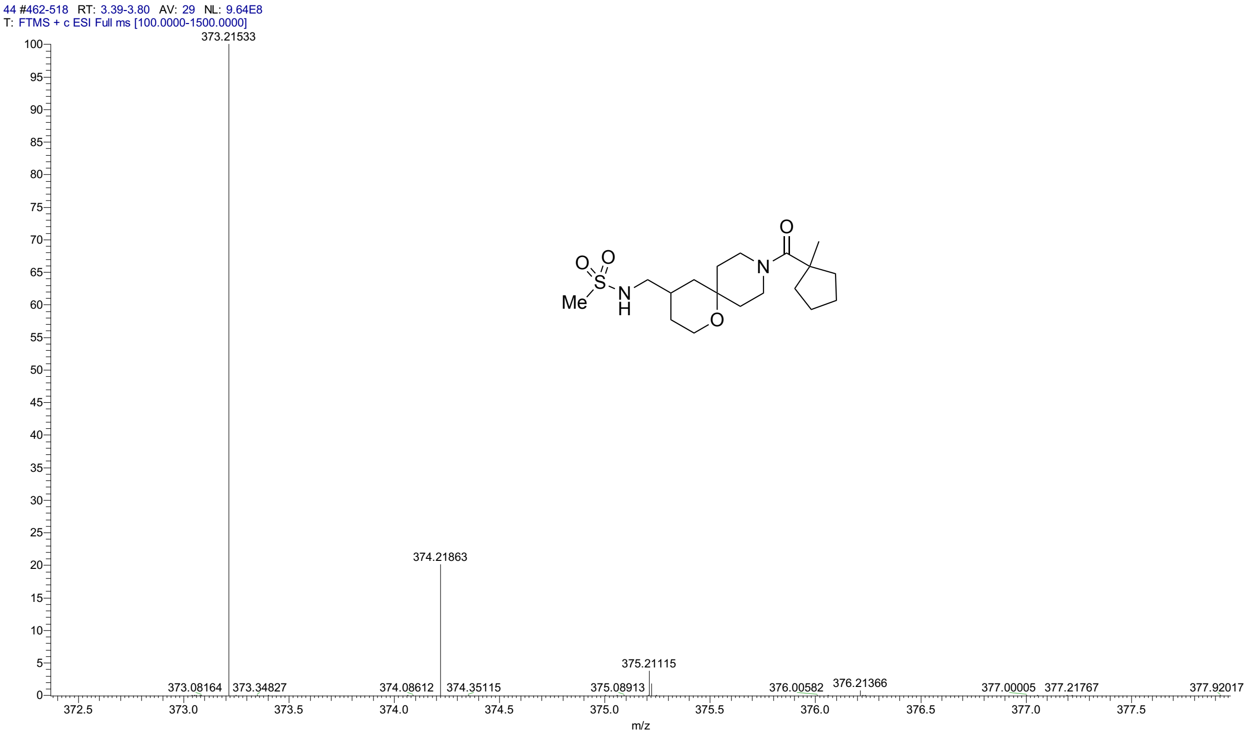


#### HPLC for **6a**

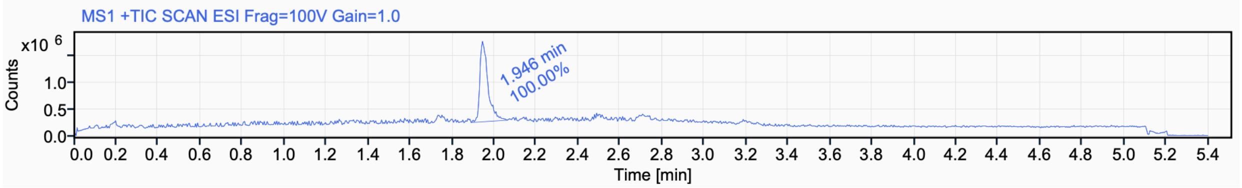


Signal Name MS1 +TIC SCAN ESI Frag=100V Gain=1.0

| RT (min) | Signal description | m/z | Purity UV(‰) / MS(%) | Area | Area% |
|---|---|---|---|---|---|
| 1.946 | MS1 +TIC SCAN ESI Frag=100V Gain=1.0 | | | 4044125.0 | 100.00 |

$^{1}$H NMR (500 MHz, $CD_3CN$) for **6b**

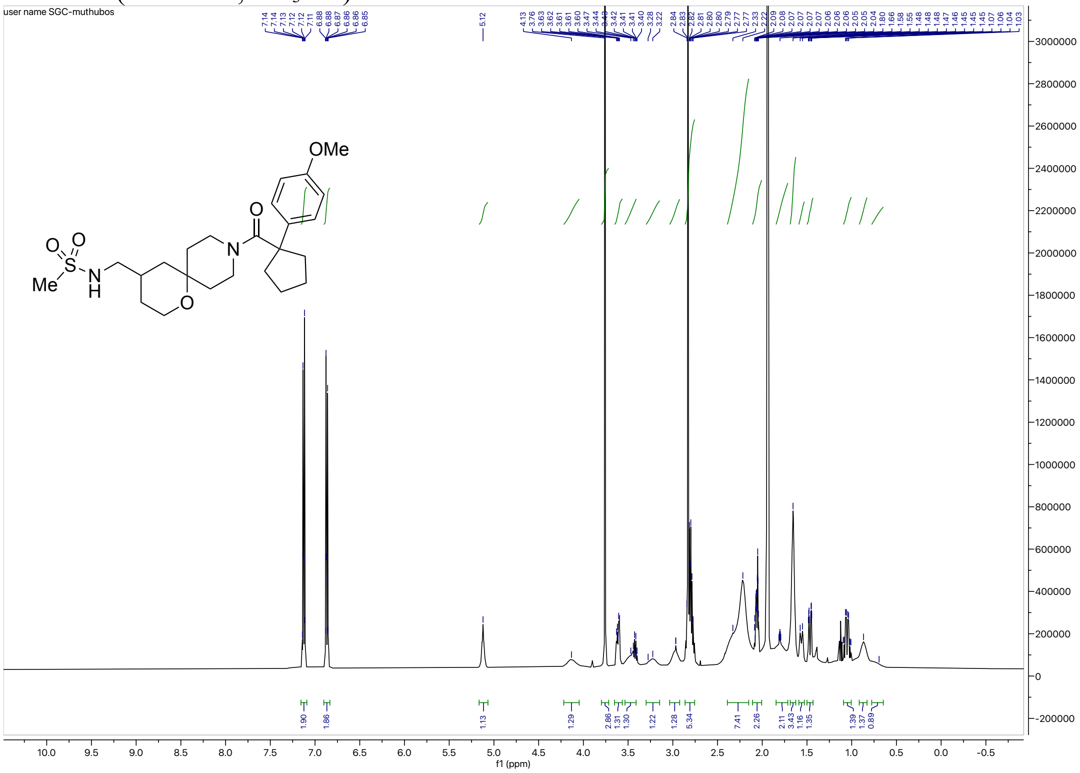


$^{13}$C NMR (126 MHz, $CD_3CN$) for **6b**

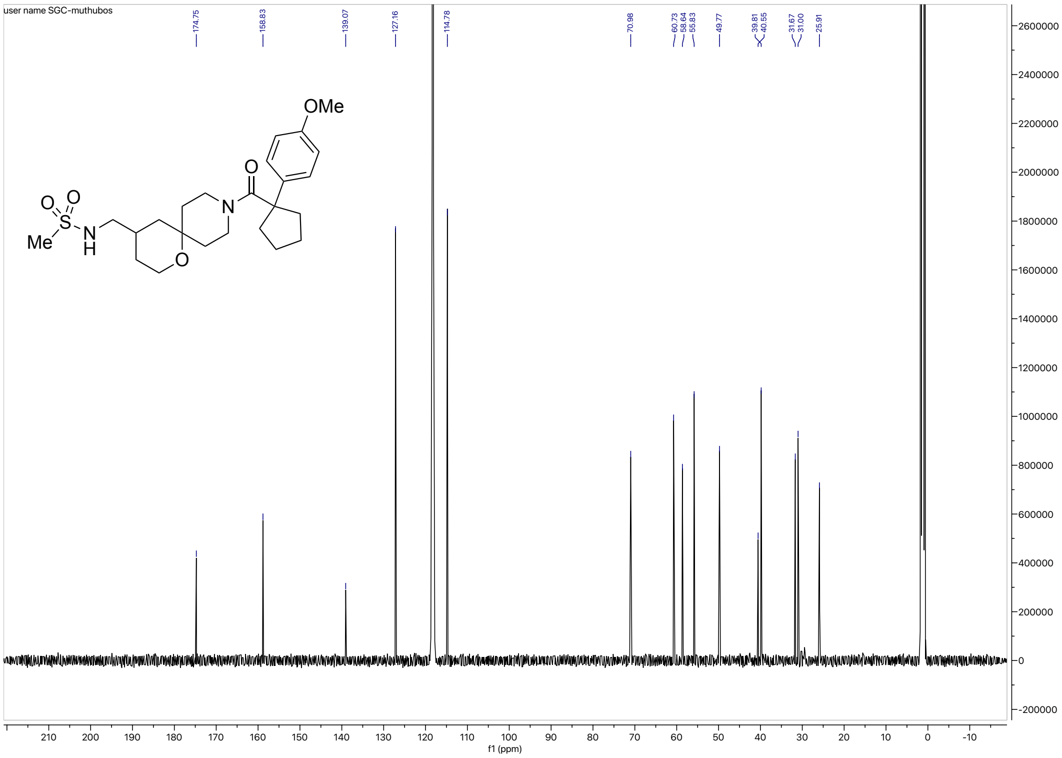

HRMS for **6b**

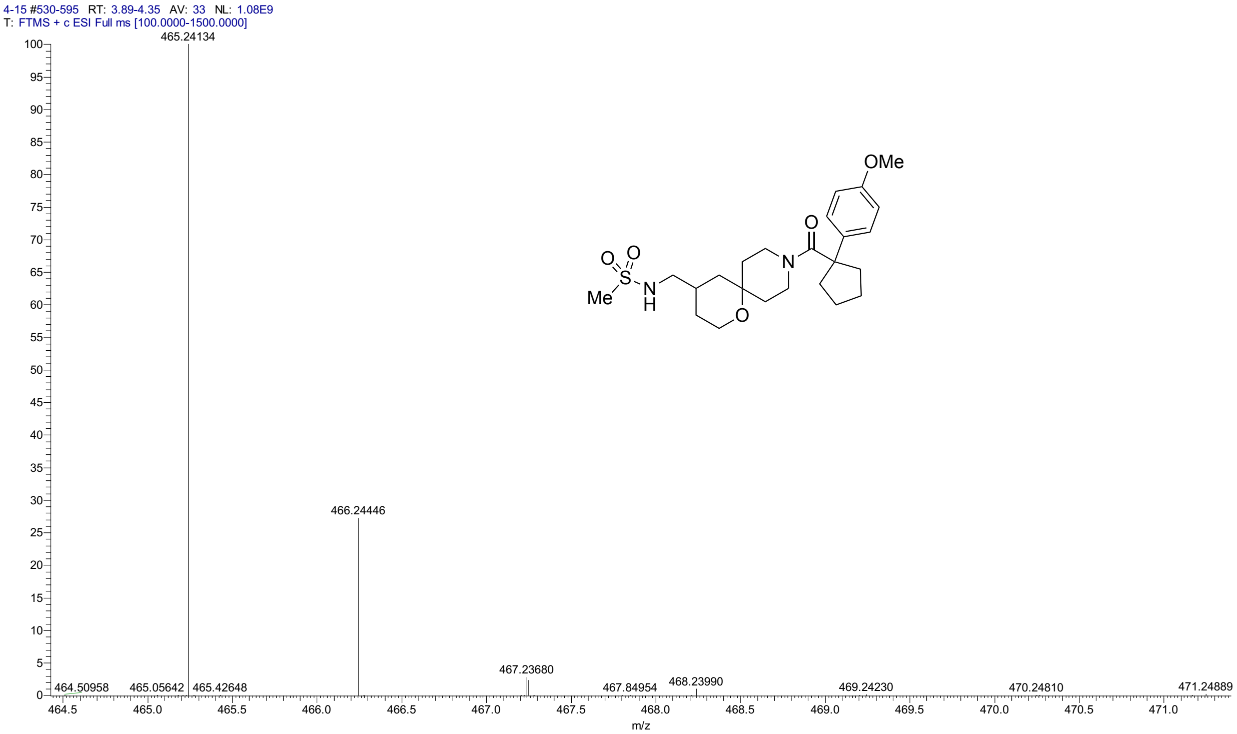


HPLC for **6b**

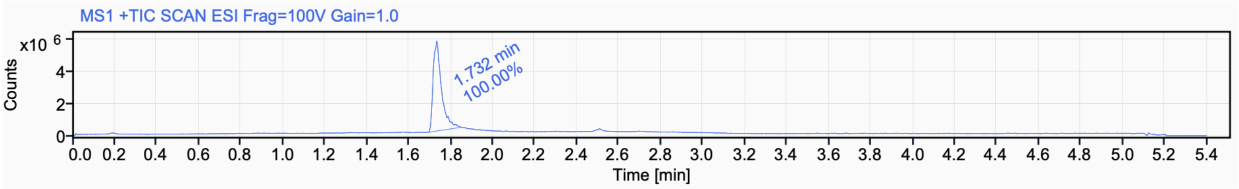


Signal Name MS1 +TIC SCAN ESI Frag=100V Gain=1.0

| RT (min) | Signal description | m/z | Purity UV(‰) / MS(%) | Area | Area% |
|---|---|---|---|---|---|
| 1.732 | MS1 +TIC SCAN ESI Frag=100V Gain=1.0 | | | 15690270.7 | 100.00 |

$^{1}$H NMR (500 MHz, $CD_3CN$) for **6c**

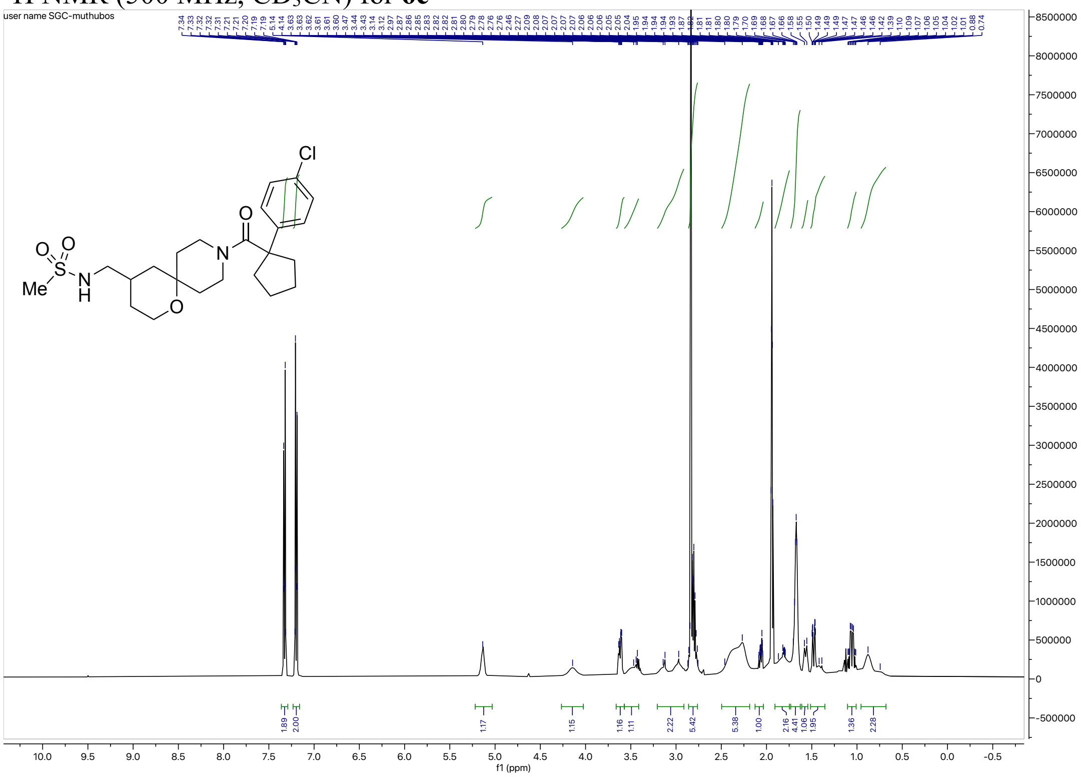

$^{13}$C NMR (126 MHz, $CD_3CN$) for **6c**

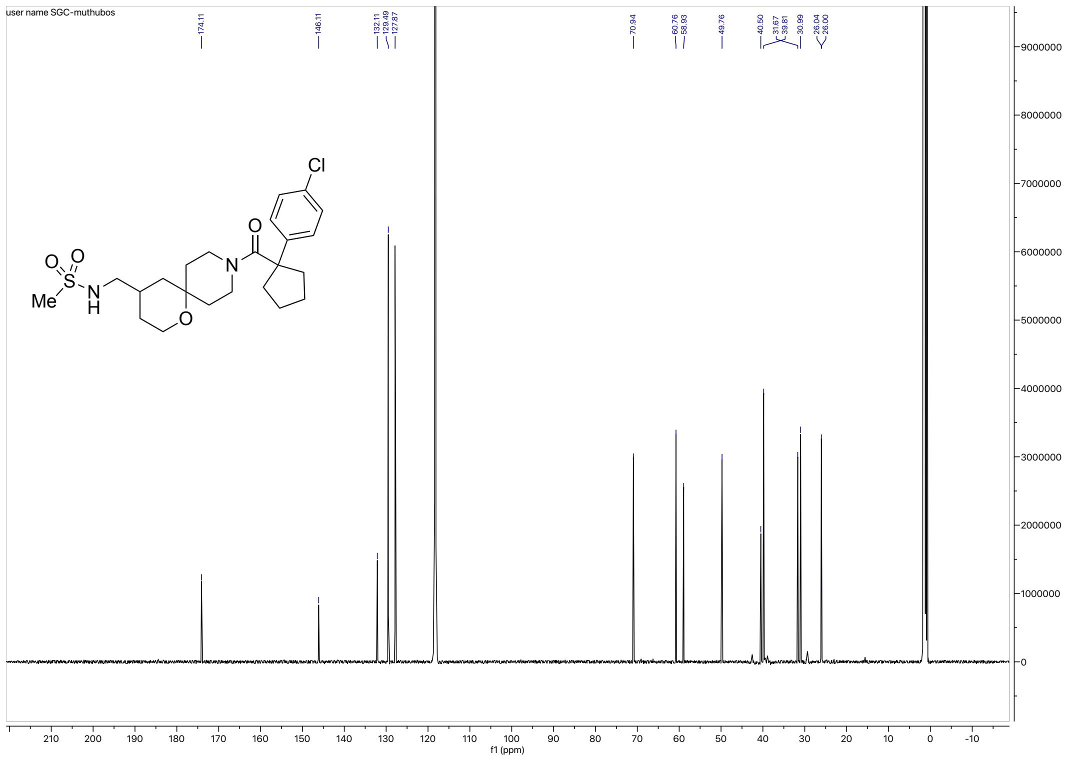

HRMS for **6c**

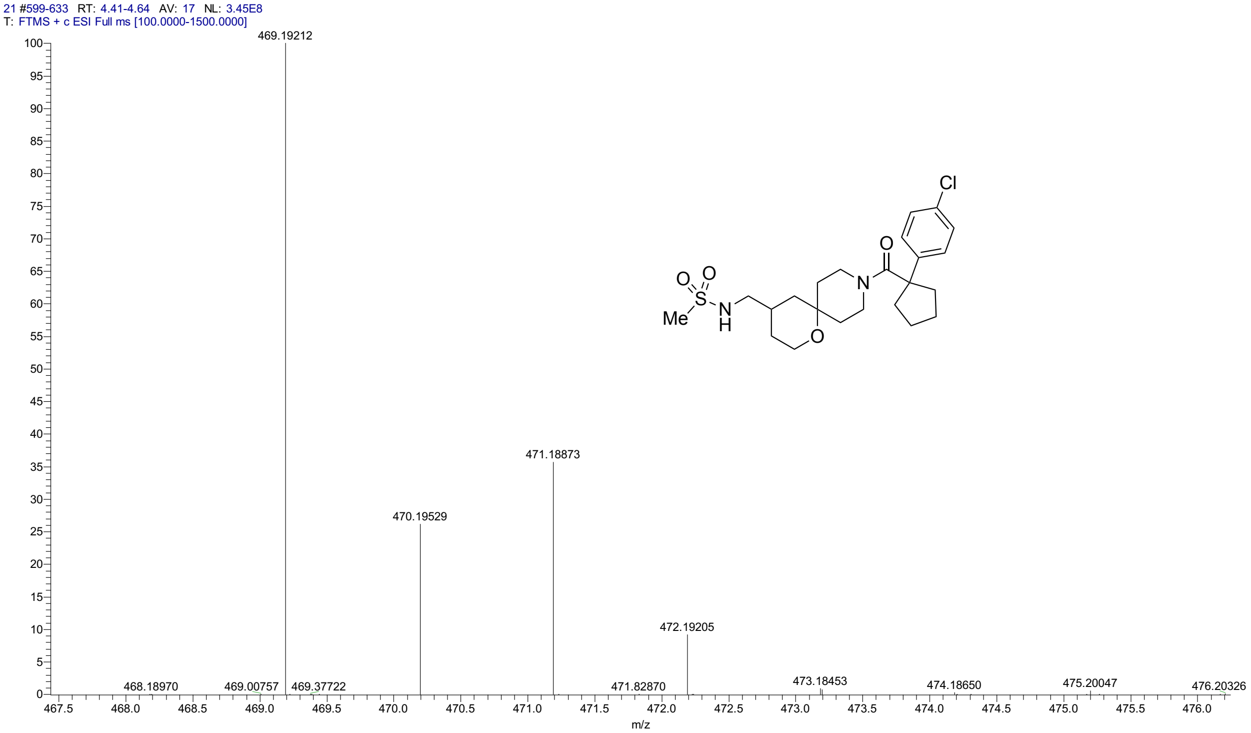


HPLC for **6d**

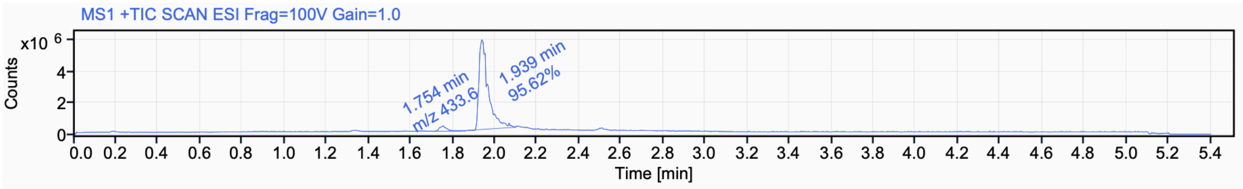


Signal Name MS1 +TIC SCAN ESI Frag=100V Gain=1.0

| RT (min) | Signal description | m/z | Purity UV(‰) / MS(%) | Area | Area% |
|---|---|---|---|---|---|
| 1.754 | MS1 +TIC SCAN ESI Frag=100V Gain=1.0 | 433.6 | | 807013.8 | 4.38 |
| 1.939 | MS1 +TIC SCAN ESI Frag=100V Gain=1.0 | | | 17617918.8 | 95.62 |

$^{1}$H NMR (500 MHz, $CD_3CN$) for **6d**

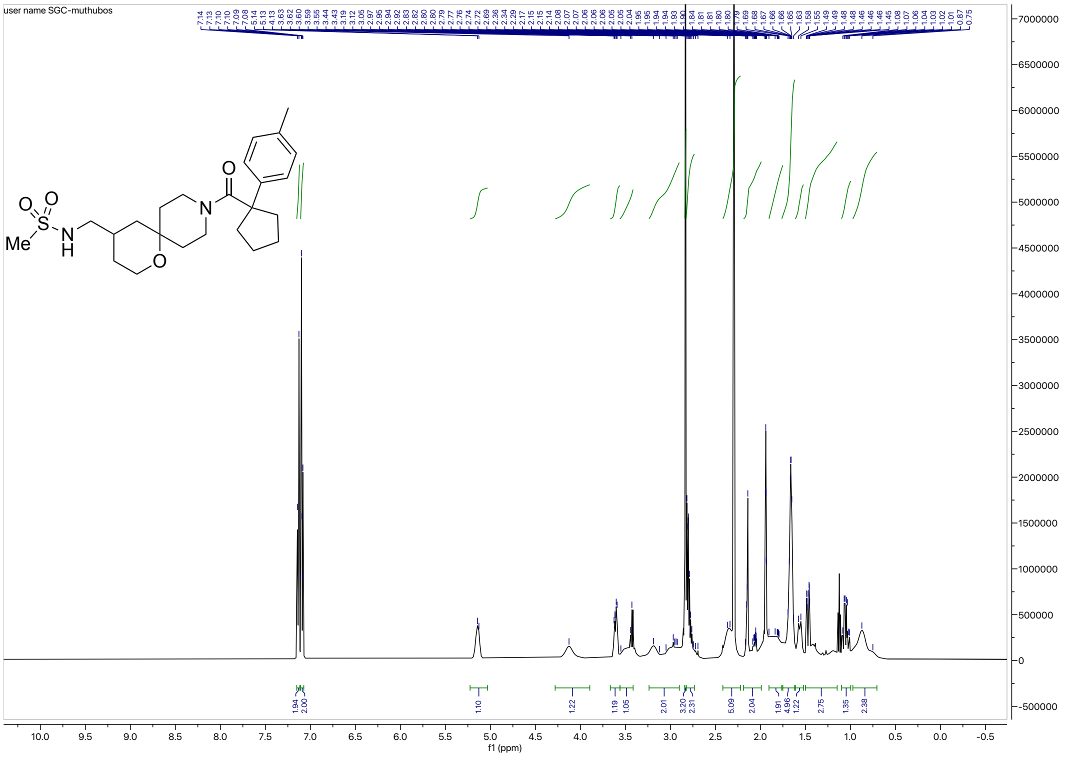

$^{13}$C NMR (126 MHz, $CD_3CN$) for **6d**

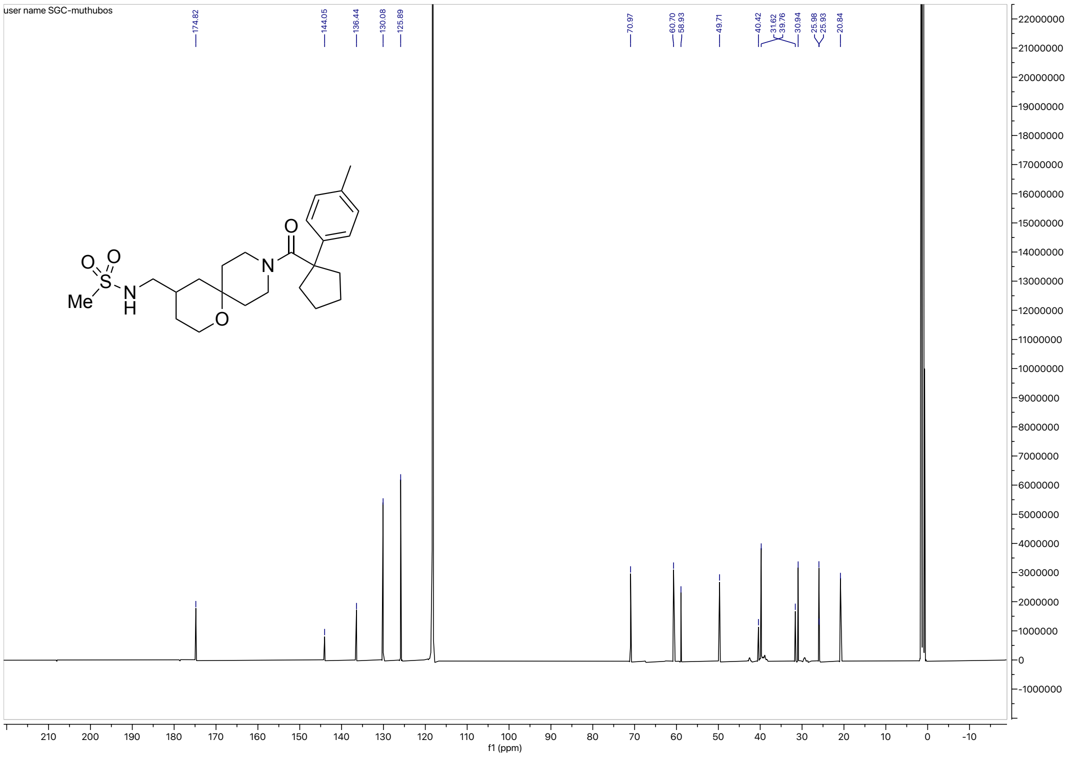

### HRMS for **6d**

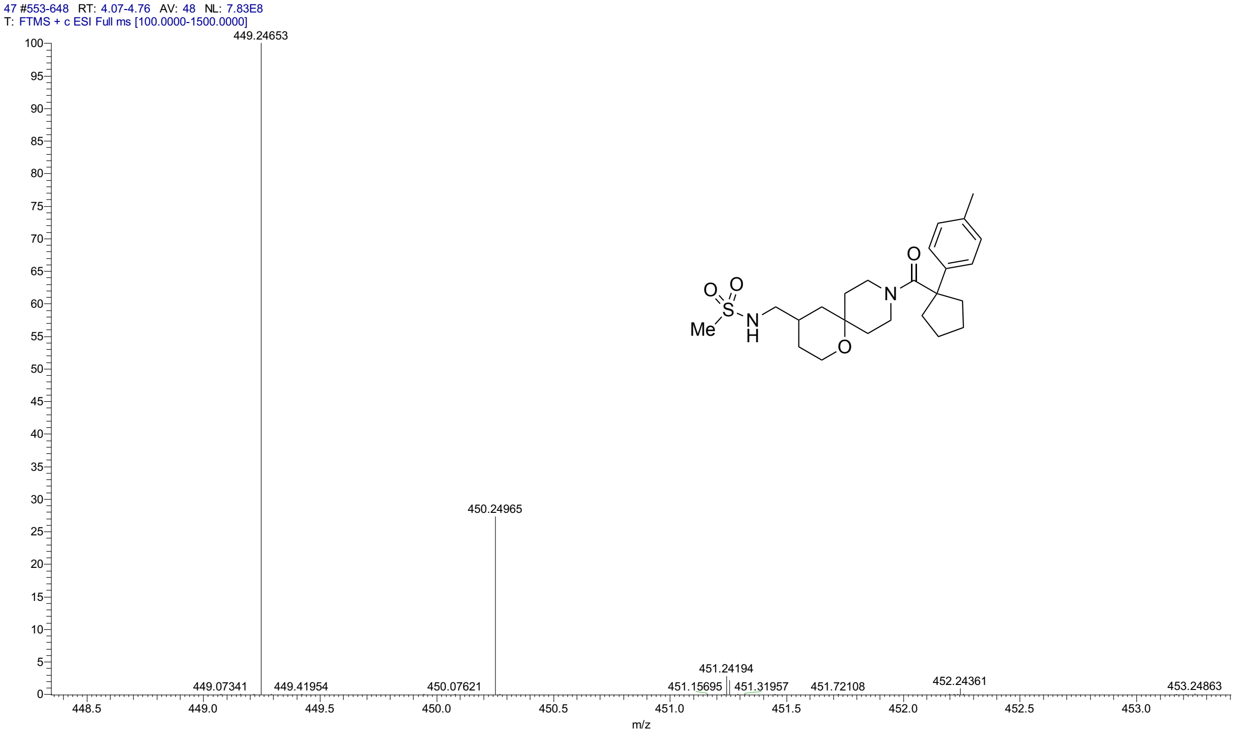


### HPLC for **6d**

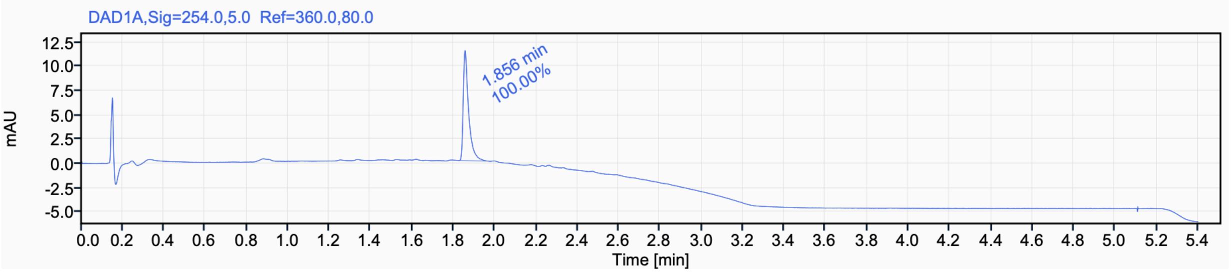


Signal Name DAD1A

| RT (min) | Signal description | Area | Area% | Peak Resolution USP |
|---|---|---|---|---|
| 1.856 | DAD1A,Sig=254.0,5.0 Ref=360.0,80.0 | 20.4 | 100.00 | |

$^{1}H$ NMR (500 MHz, $CD_3CN$) for **6e**

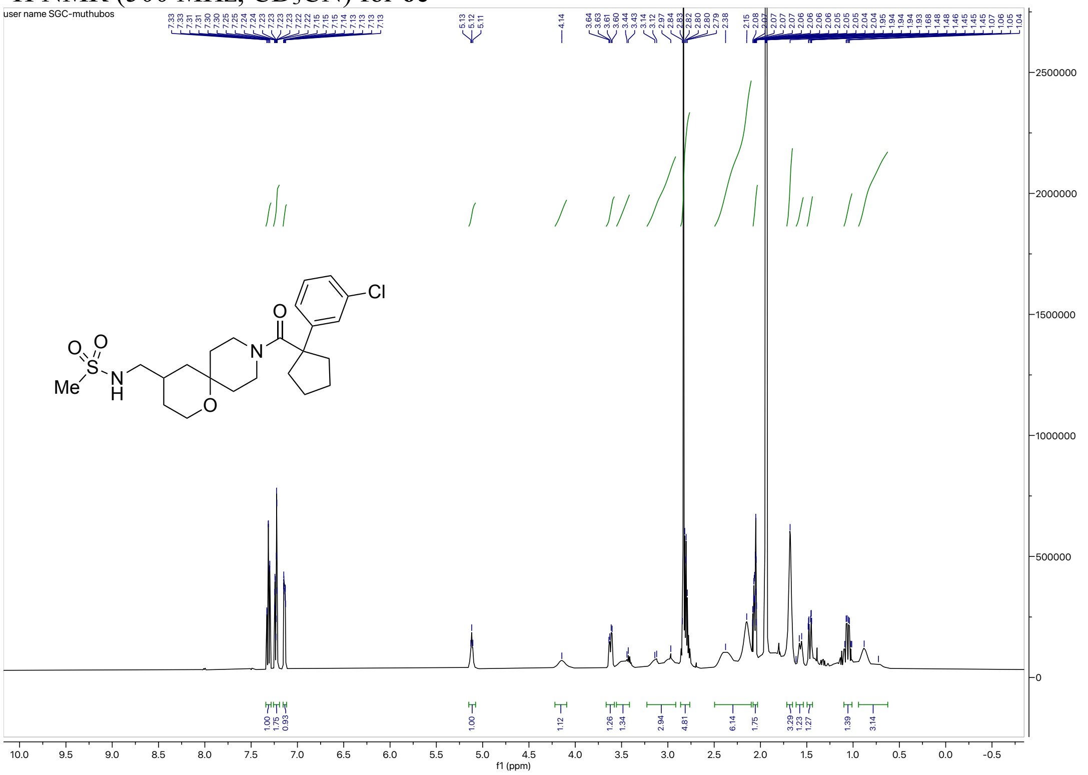


$^{13}C$ NMR (126 MHz, $CD_3CN$) for **6e**

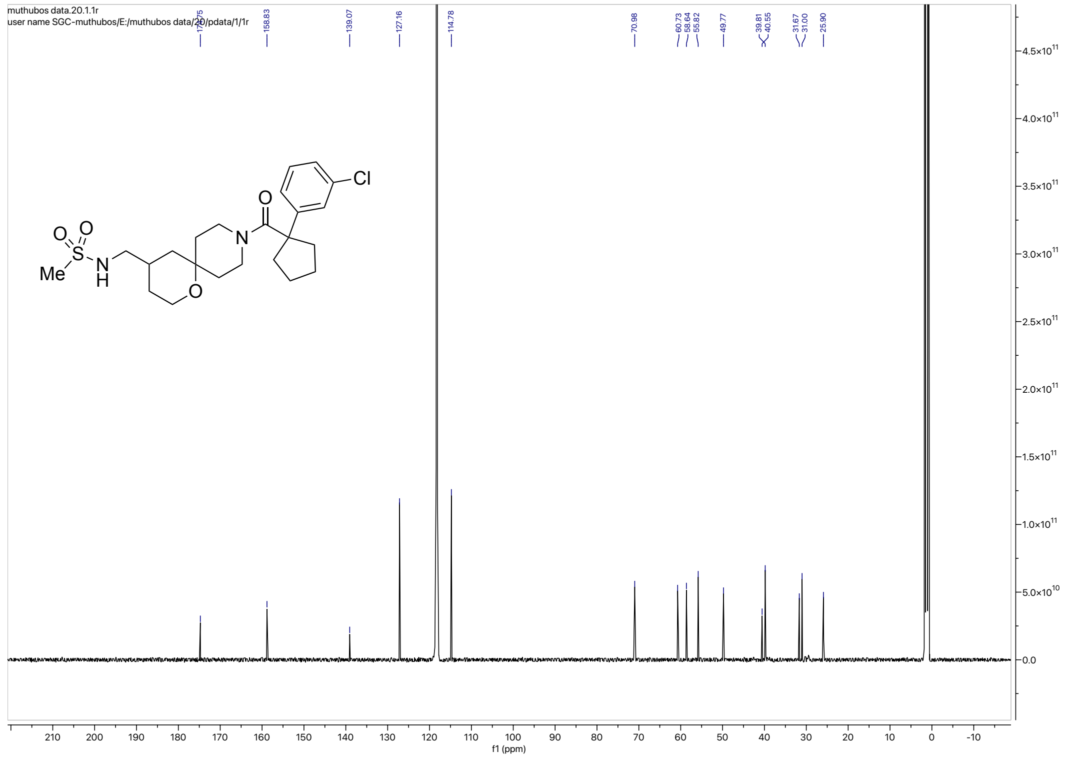

#### HRMS for **6e**

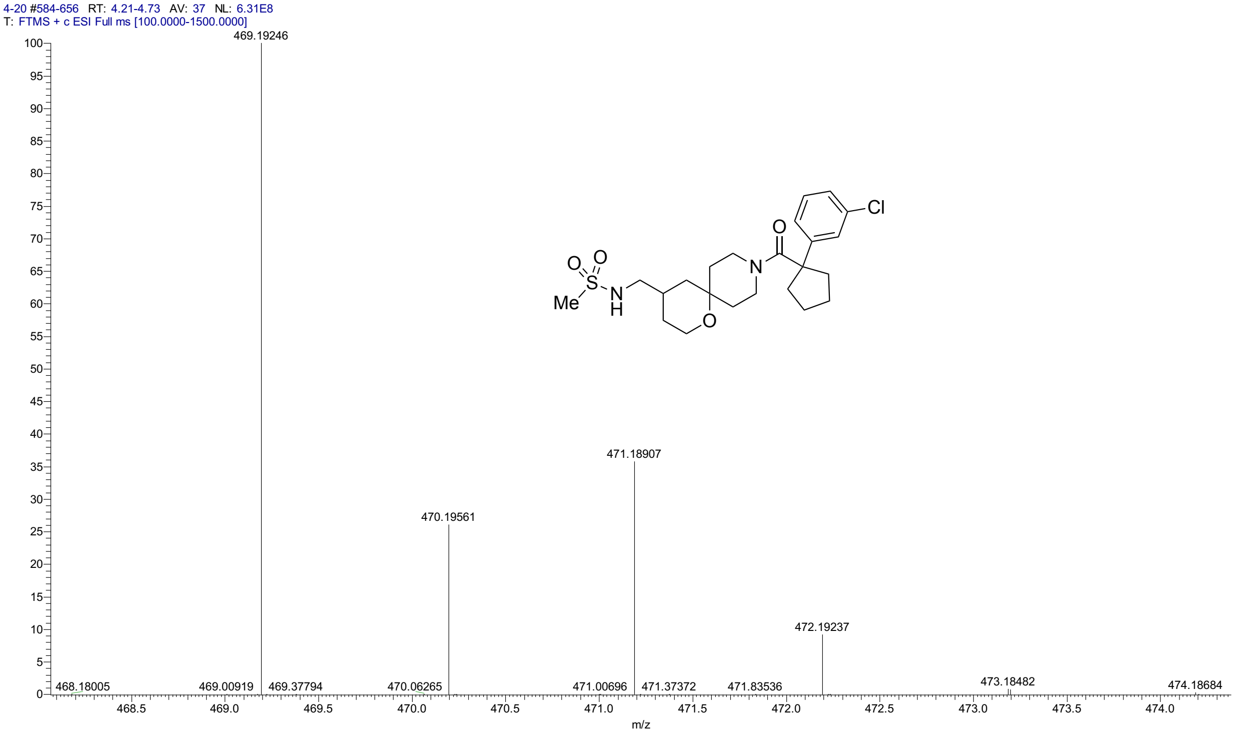


#### HPLC for **6e**

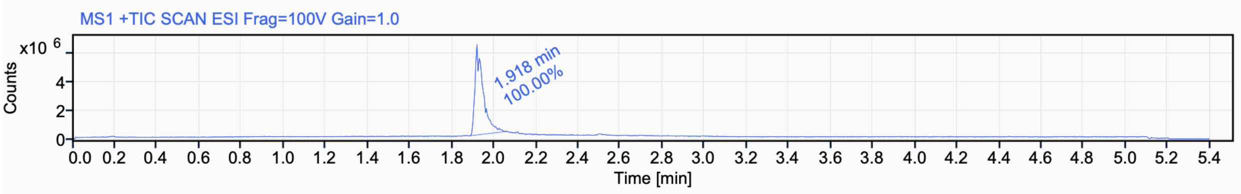


Signal Name MS1 +TIC SCAN ESI Frag=100V Gain=1.0

| RT (min) | Signal description | Area | Area% | Peak Resolution USP |
|---|---|---|---|---|
| 1.918 | MS1 +TIC SCAN ESI Frag=100V Gain=1.0 | 17279929.7 | 100.00 | |

[1]H NMR (500 MHz, $CD_3CN$) for **6f**

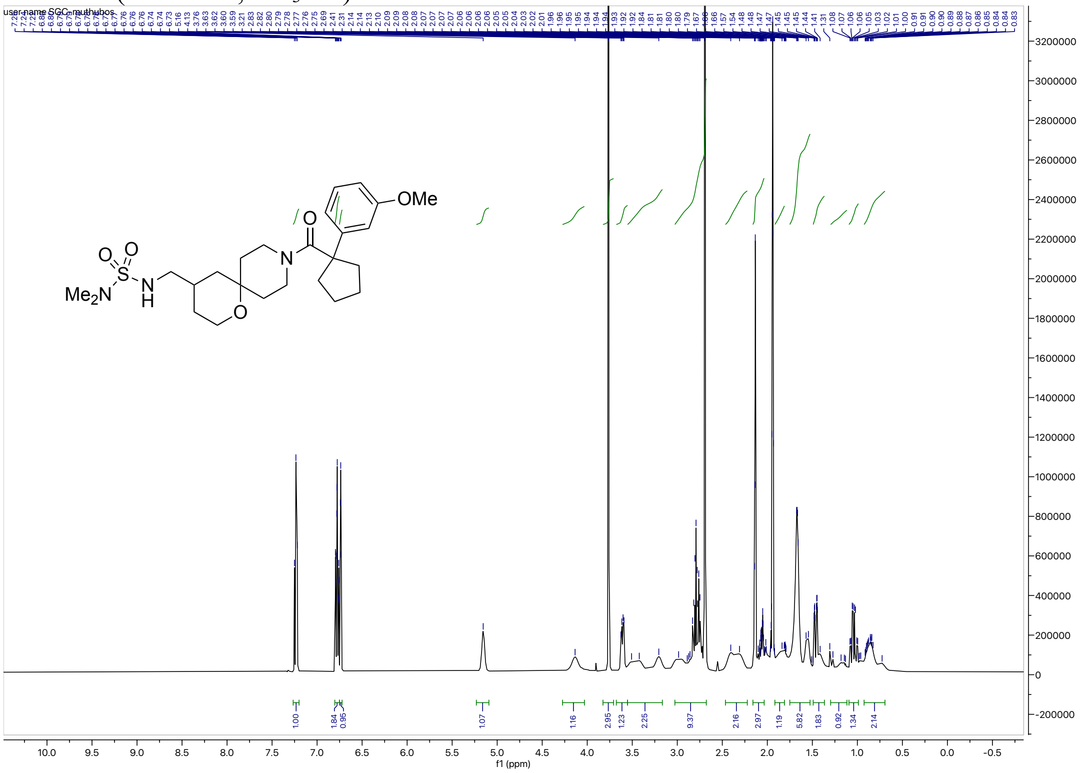

[13]C NMR (126 MHz, $CD_3CN$) for **6f**

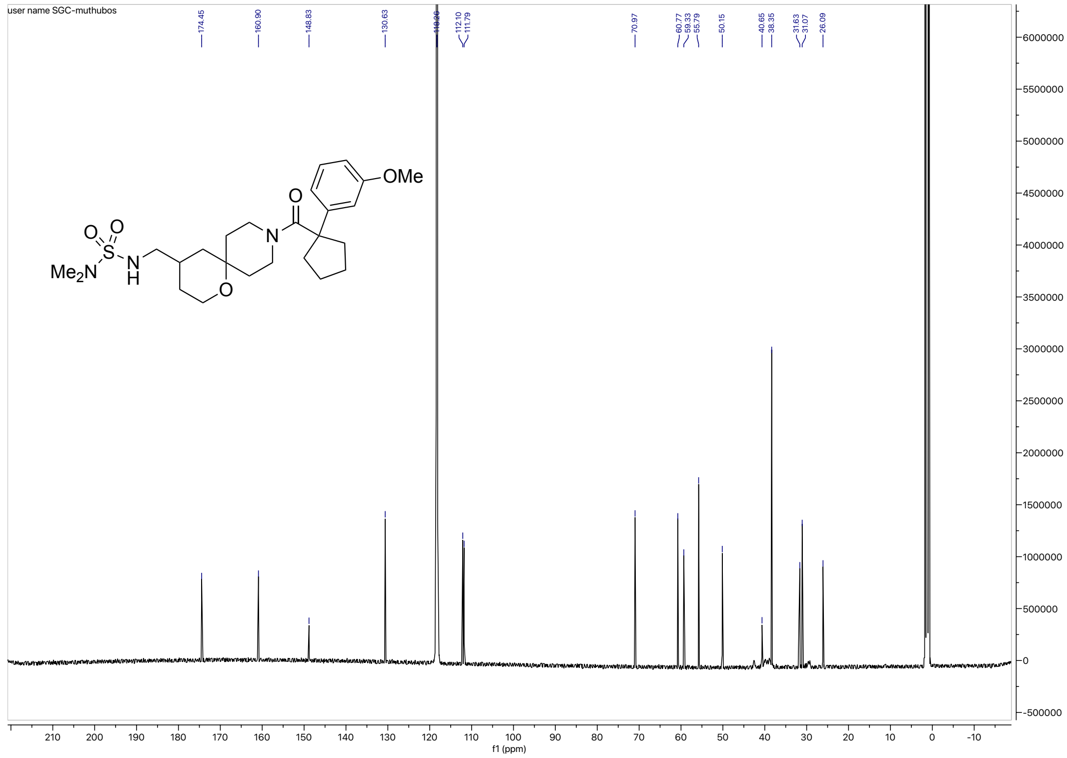

HRMS for **6f**

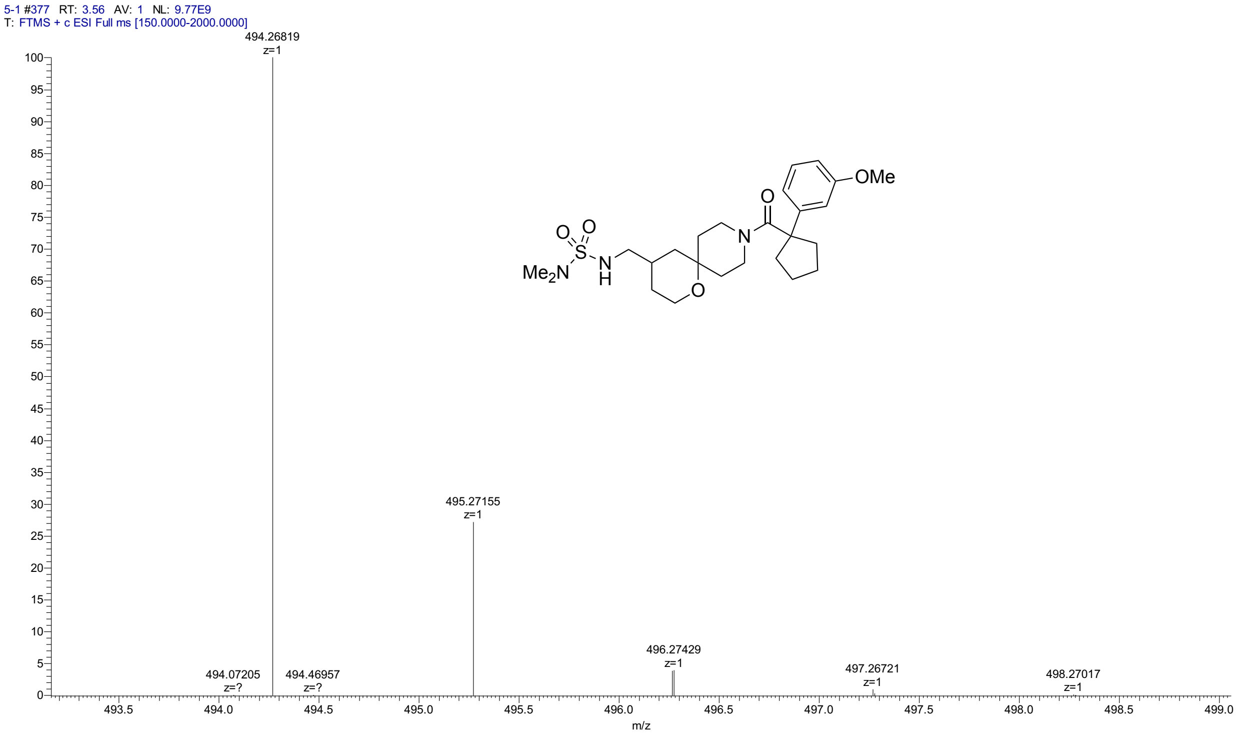


HPLC for **6f**

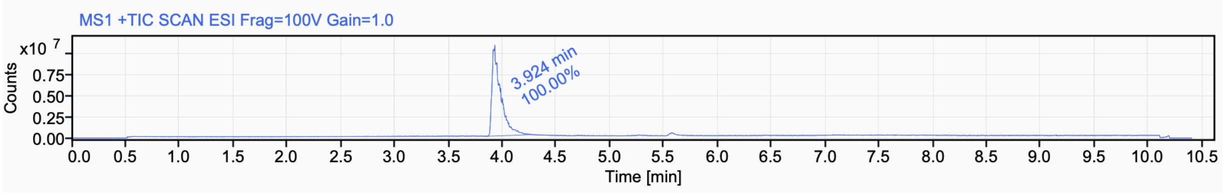


Signal Name MS1 +TIC SCAN ESI Frag=100V Gain=1.0

| RT (min) | Signal description | Area | Area% | Peak Resolution USP |
|---|---|---|---|---|
| 3.924 | MS1 +TIC SCAN ESI Frag=100V Gain=1.0 | 60799978.7 | 100.00 | |

[1]H NMR (400 MHz, DMSO-$d_6$, 80 °C) for **6g**

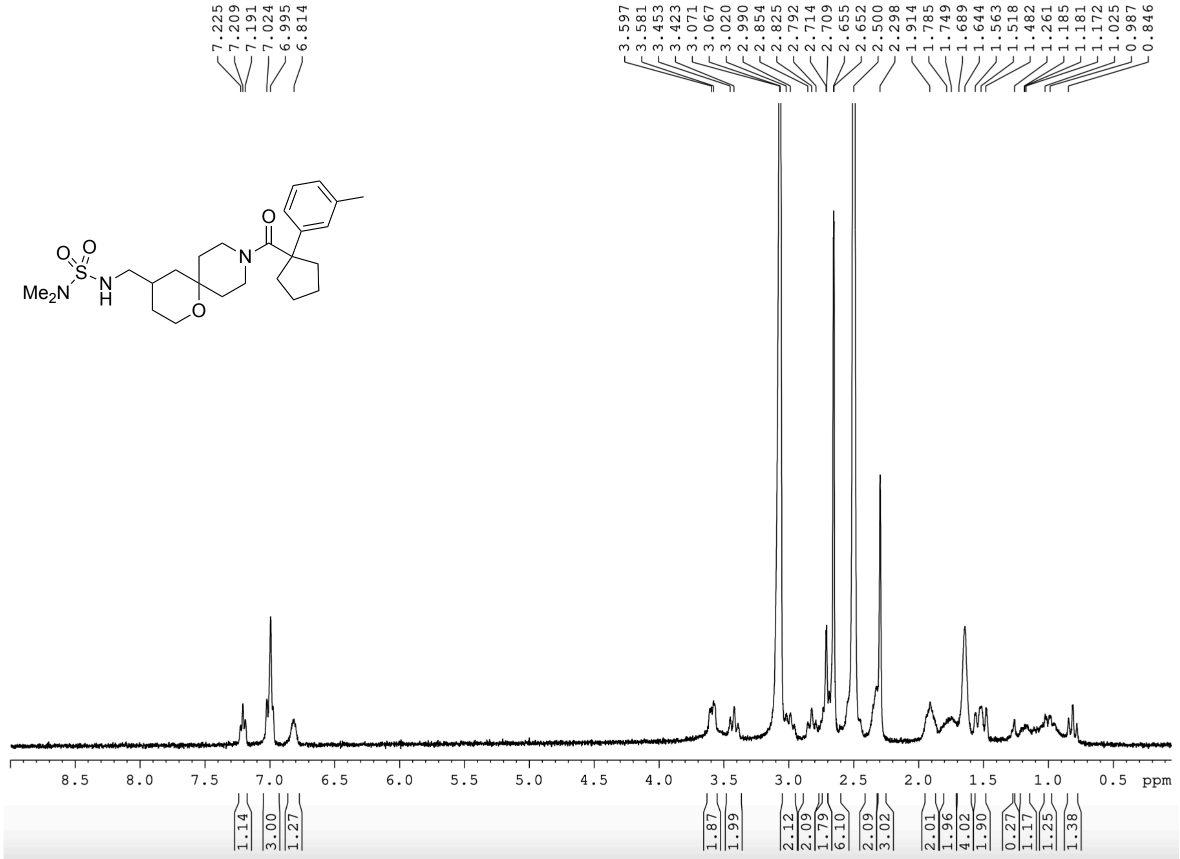


[13]C NMR (101 MHz, DMSO-$d_6$) for **6g**

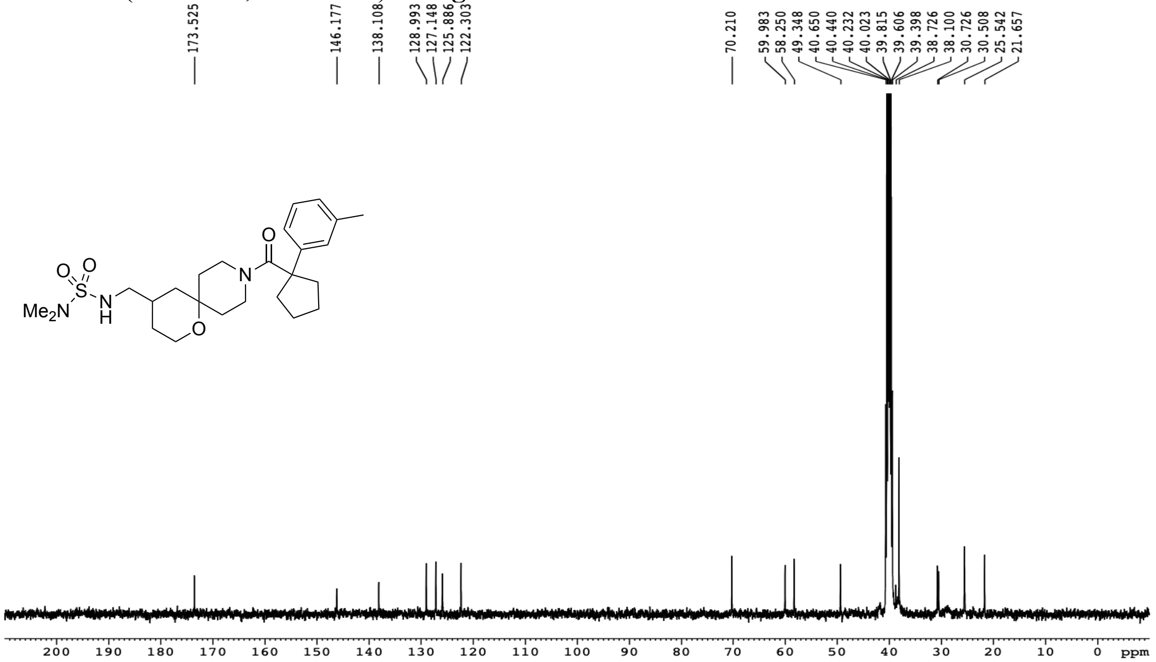

HRMS for **6g**

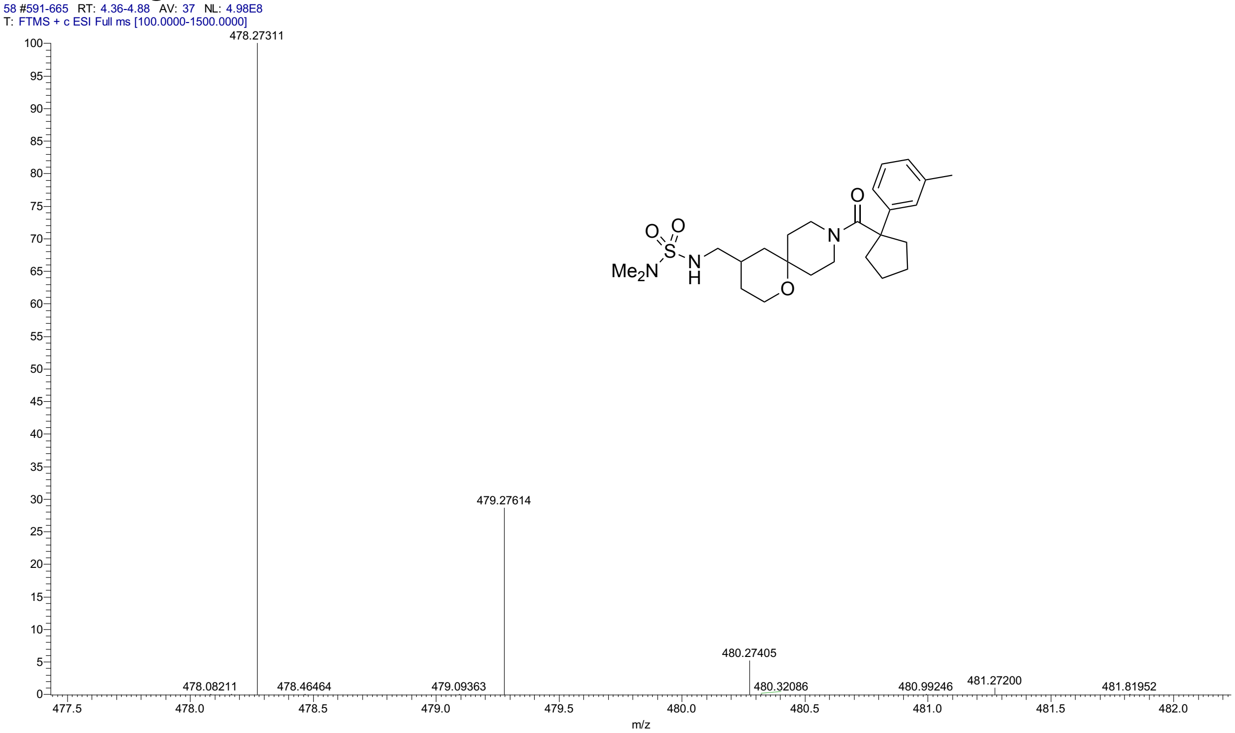


HPLC for **6g**

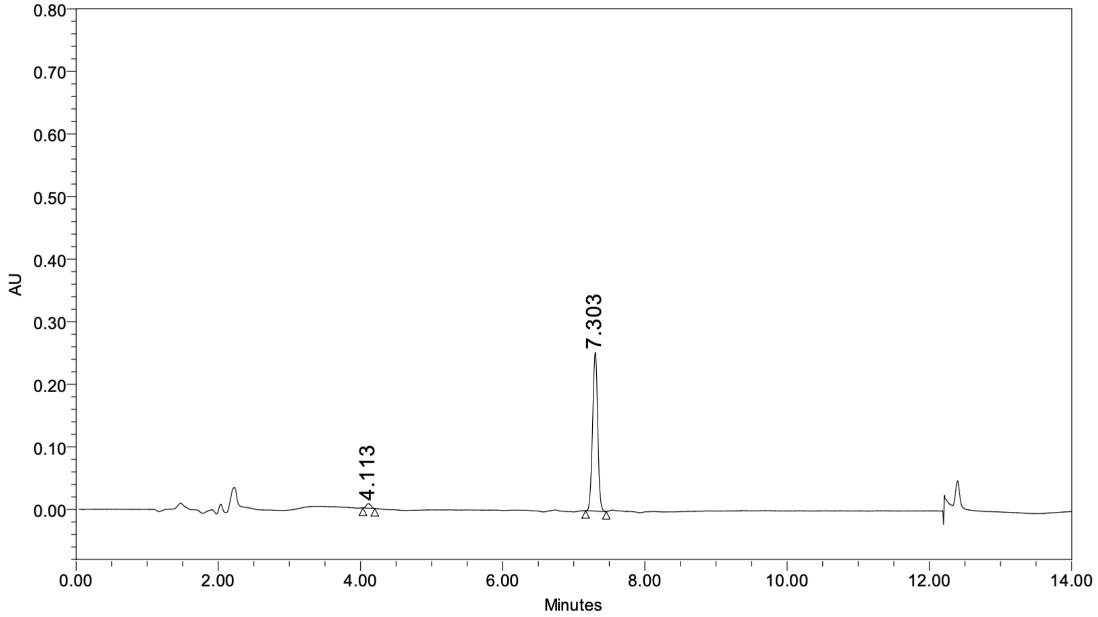


**Peak Results**

| | Name | RT | Height (μV) | Area (μV*sec) | % Area |
|---|---|---|---|---|---|
| 1 | | 4.113 | 7202 | 33098 | 2.53 |
| 2 | | 7.303 | 253587 | 1272970 | 97.47 |

$^{1}H$ NMR (400 MHz, DMSO-$d_6$, 80 °C) for **6h**

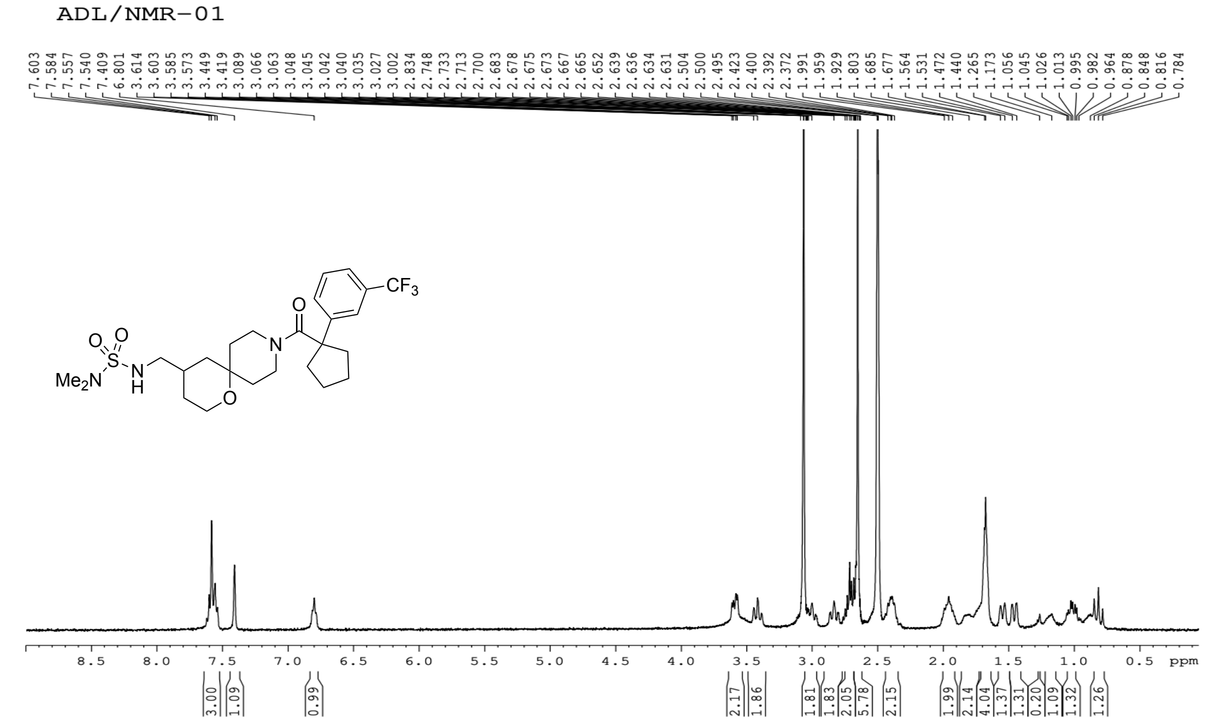


$^{13}C$ NMR (101 MHz, DMSO-$d_6$) for **6h**

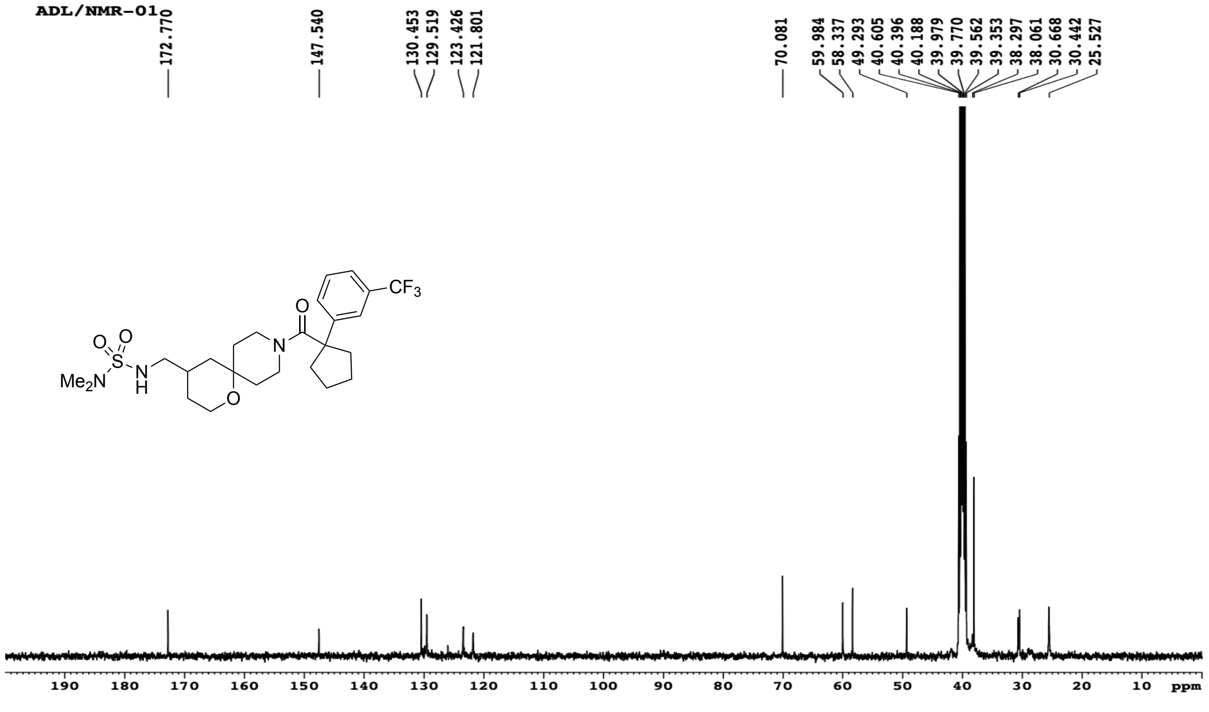

HRMS for **6h**

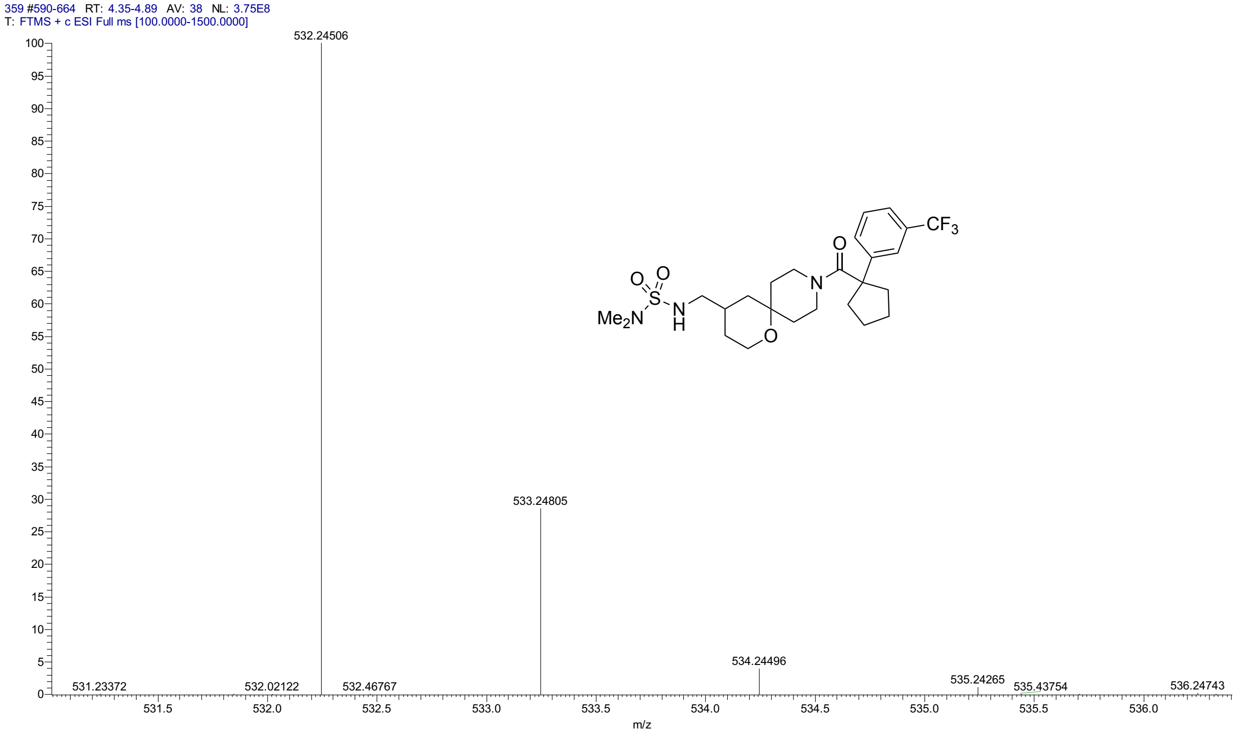


HPLC for **6h**

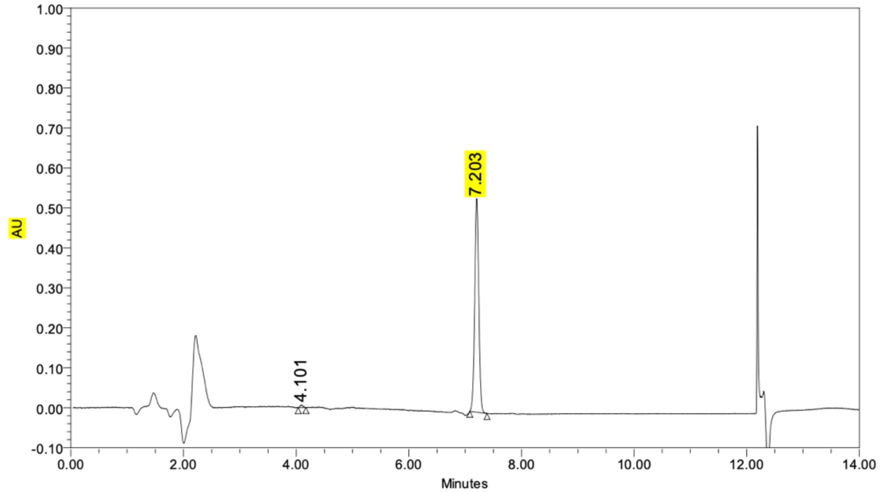


**Peak Results**

| | Name | RT | Height (µV) | Area (µV*sec) | % Area |
|---|---|---|---|---|---|
| 1 | | 4.101 | 6430 | 23650 | 0.87 |
| 2 | | 7.203 | 534666 | 2681779 | 99.13 |

$^{1}$H NMR (500 MHz, $CD_3CN$) for **6i**

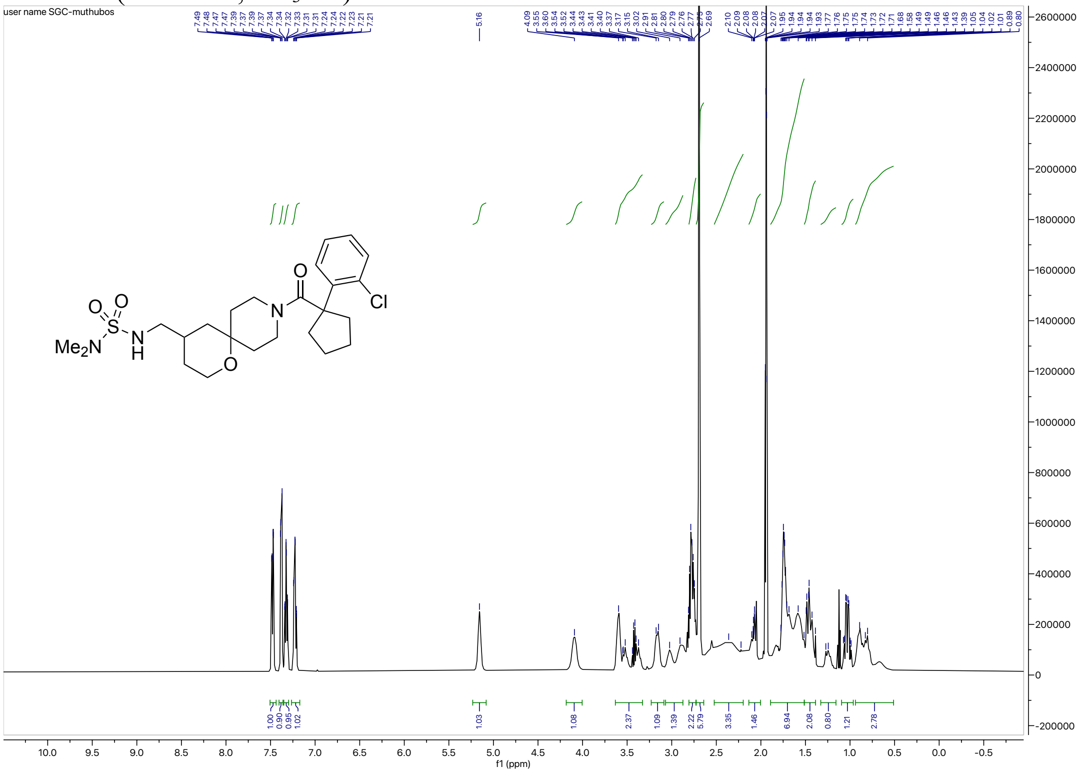

$^{13}$C NMR (126 MHz, $CD_3CN$) for **6i**

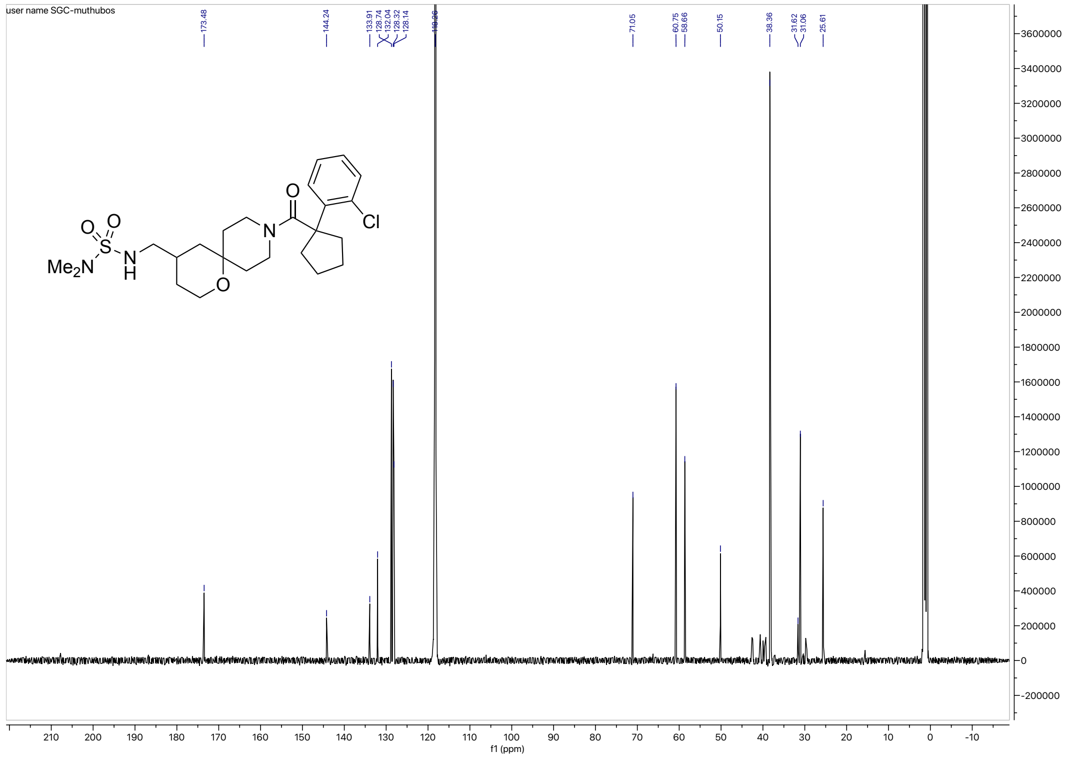

HRMS for **6i**

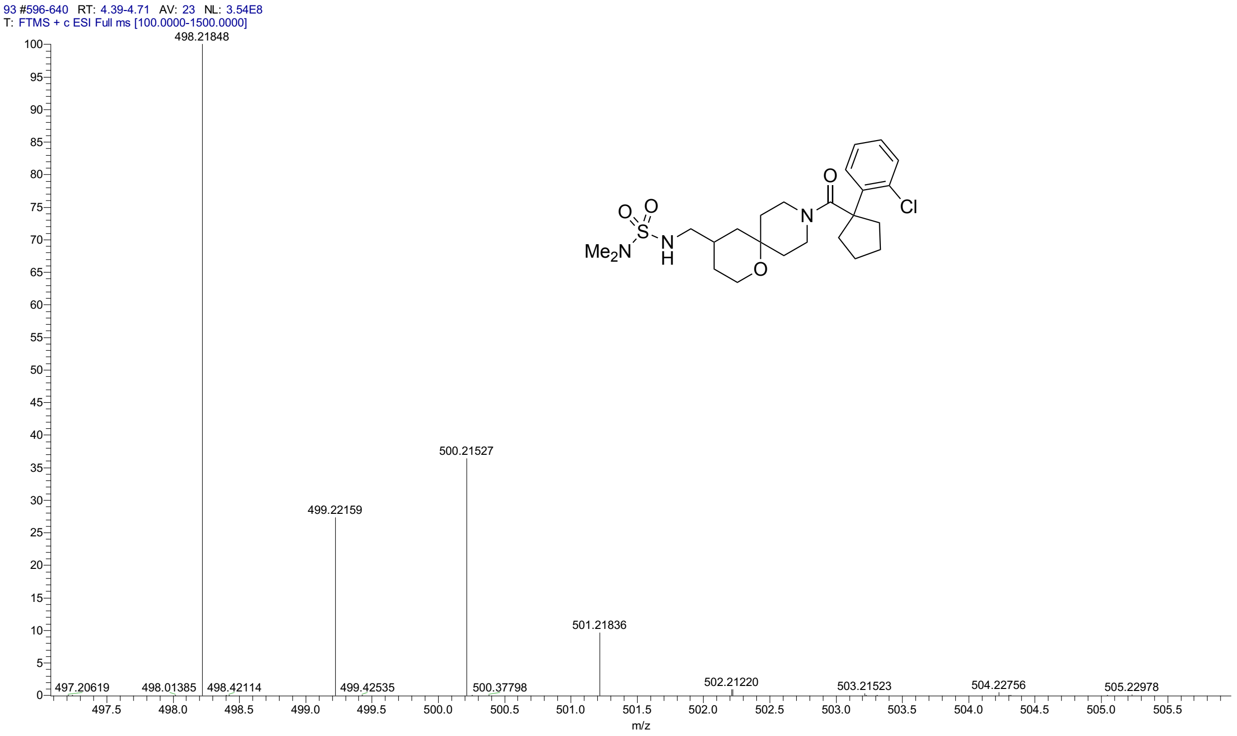


**HPLC for 6i**

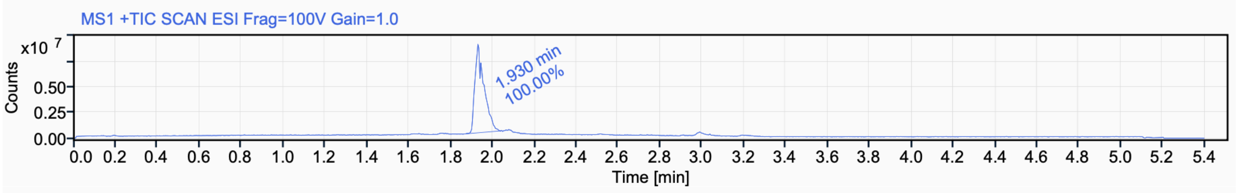


Signal Name MS1 +TIC SCAN ESI Frag=100V Gain=1.0

| RT (min) | Signal description | Area | Area% | Peak Resolution USP |
|---|---|---|---|---|
| 1.930 | MS1 +TIC SCAN ESI Frag=100V Gain=1.0 | 26305054.7 | 100.00 | |

$^{1}$H NMR (500 MHz, $CD_3CN$) for **6j**

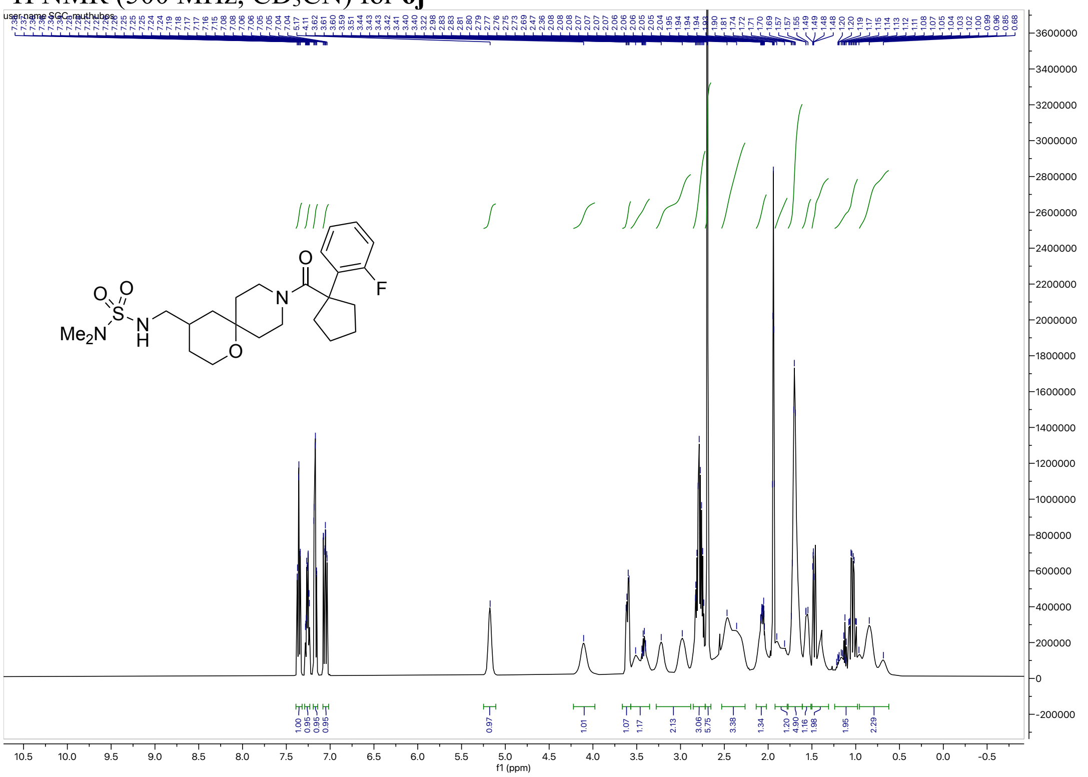

$^{13}$C NMR (126 MHz, $CD_3CN$) for **6j**

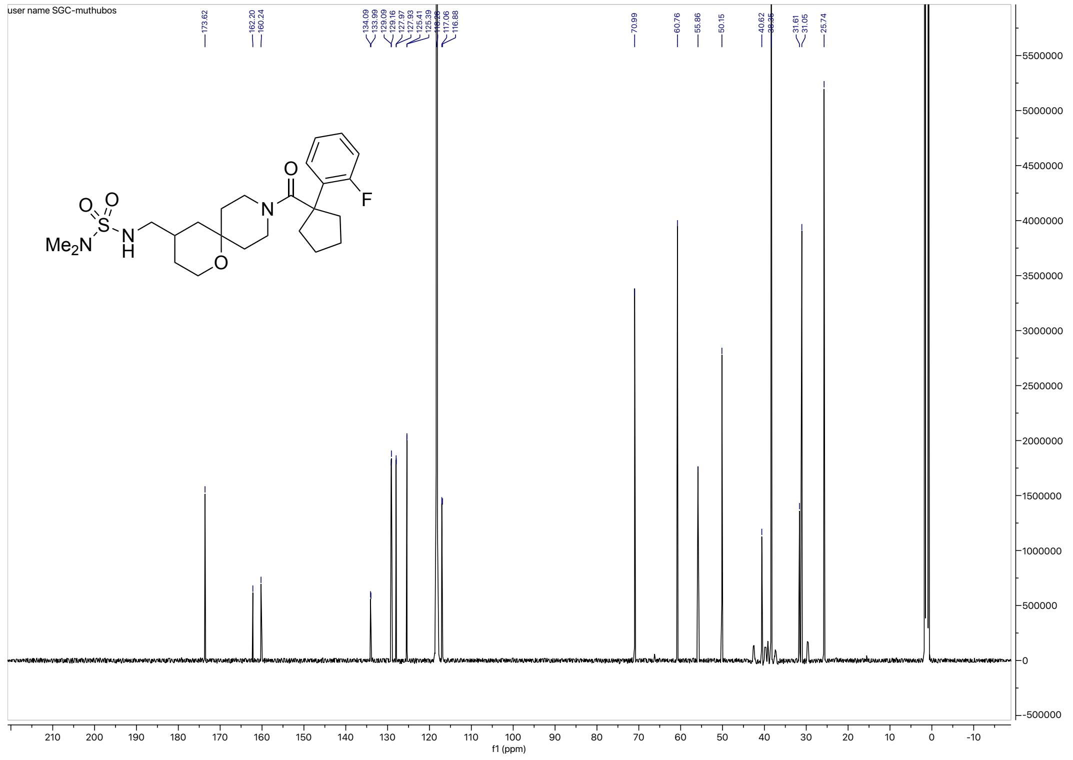

HRMS for **6j**

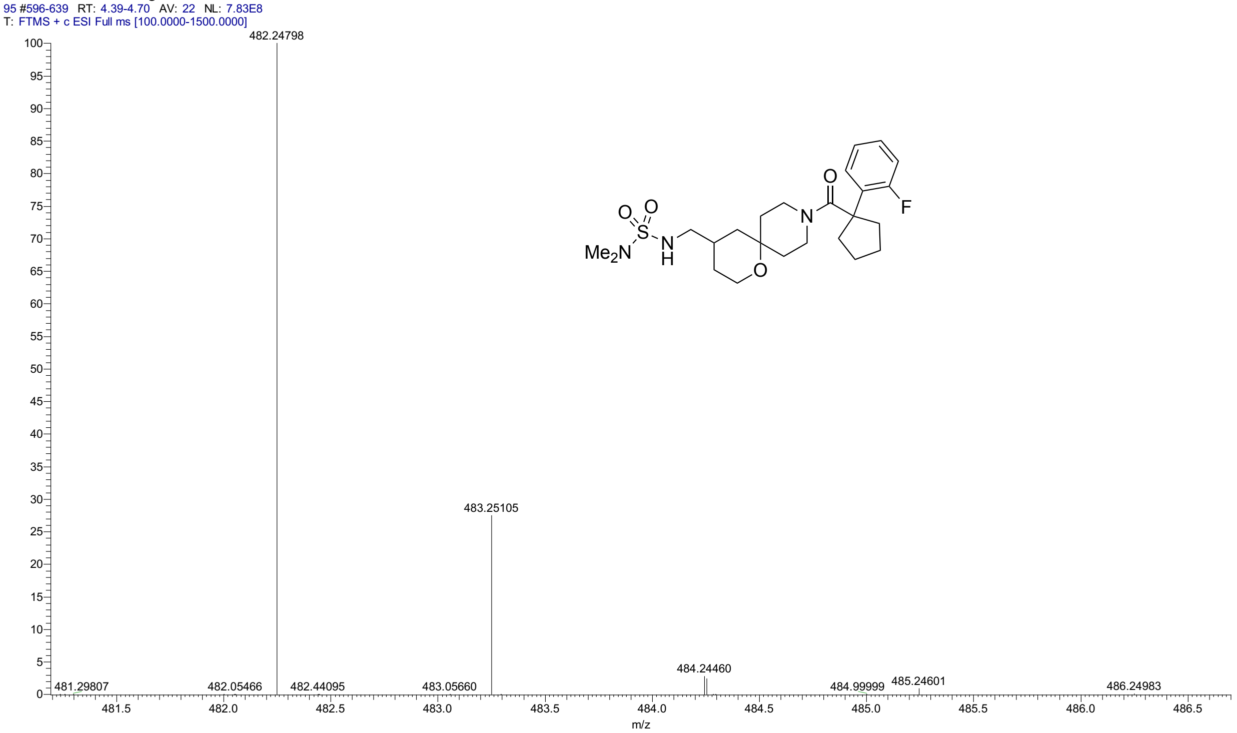


HPLC for **6j**

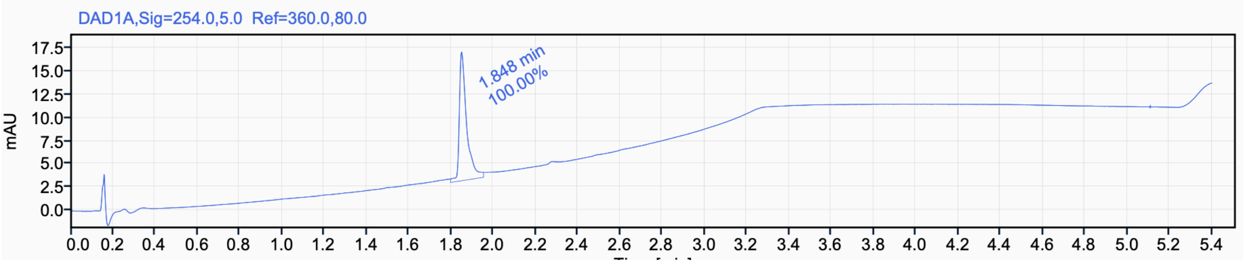


Signal Name DAD1A

| RT (min) | Signal description | m/z | Purity UV(‰) / MS(%) | Area | Area% |
|---|---|---|---|---|---|
| 1.848 | DAD1A,Sig=254.0,5.0 Ref=360.0,80.0 | | | 30.5 | 100.00 |

[1]H NMR (400 MHz, DMSO-$d_6$, 80 °C) for **6k**

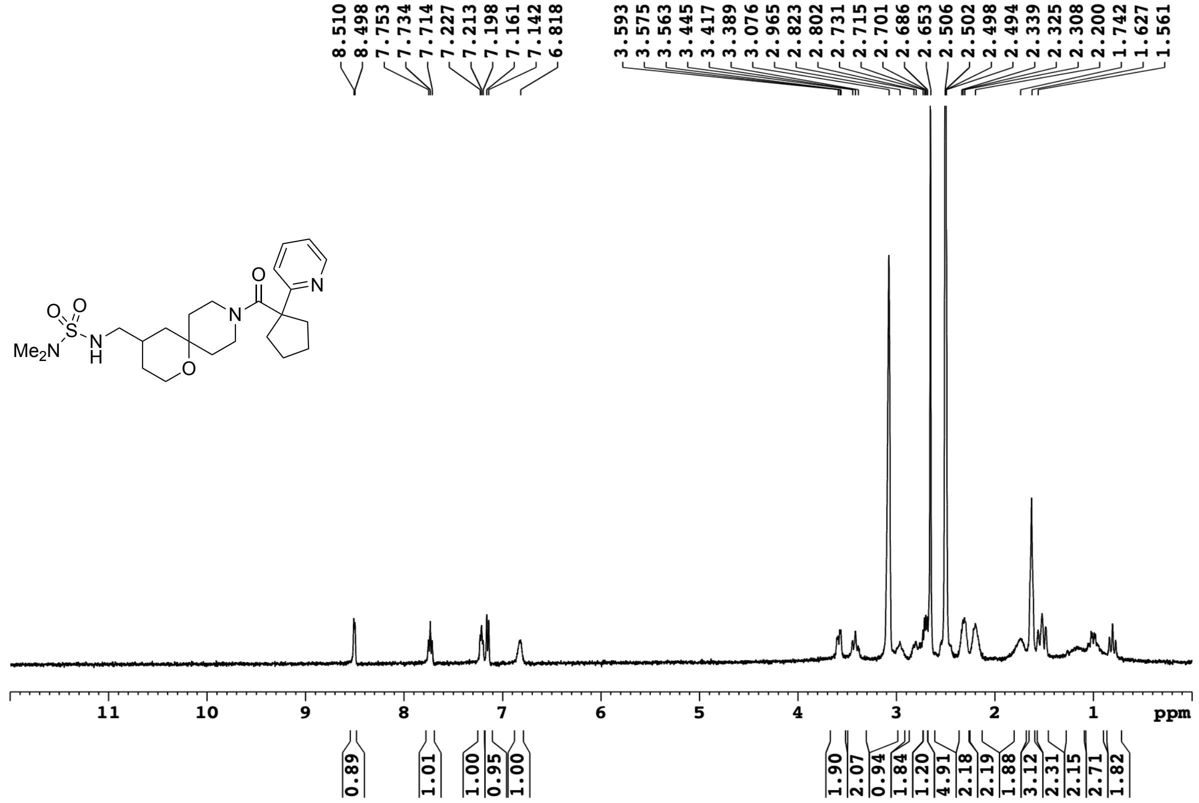


[13]C NMR (101 MHz, DMSO-$d_6$) for **6k**

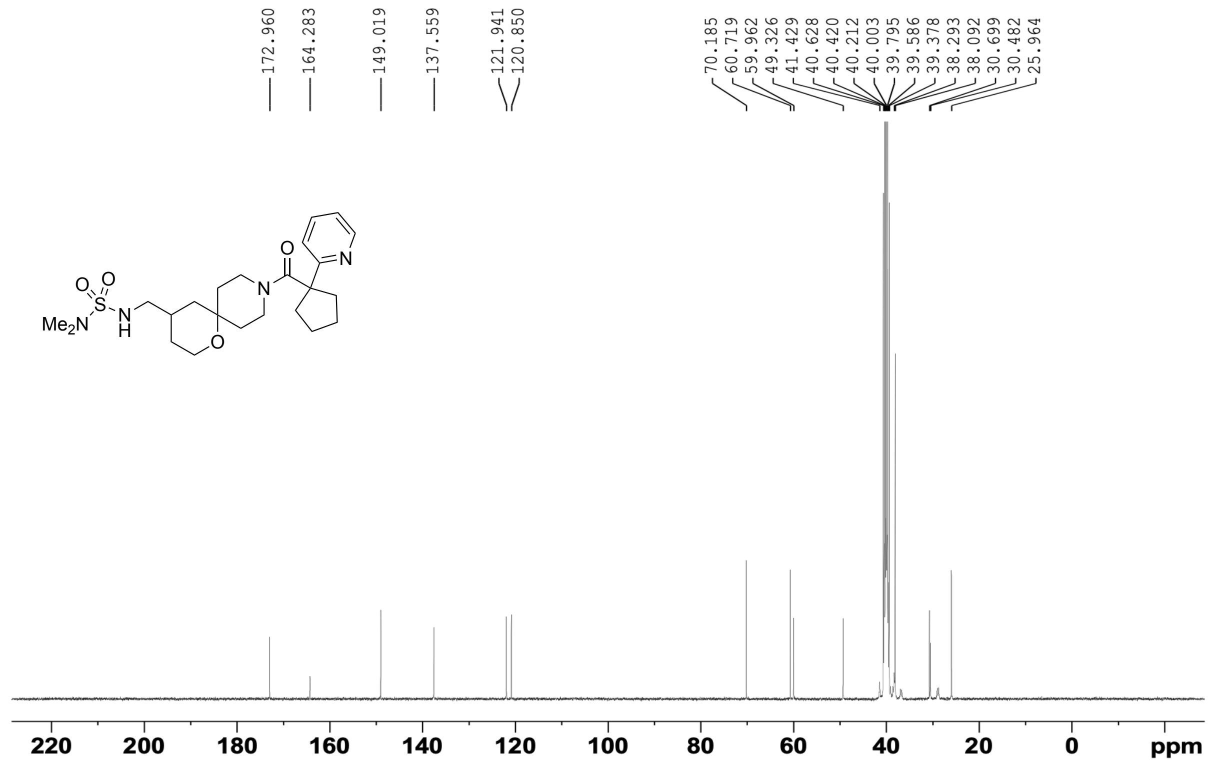

HRMS for **6k**

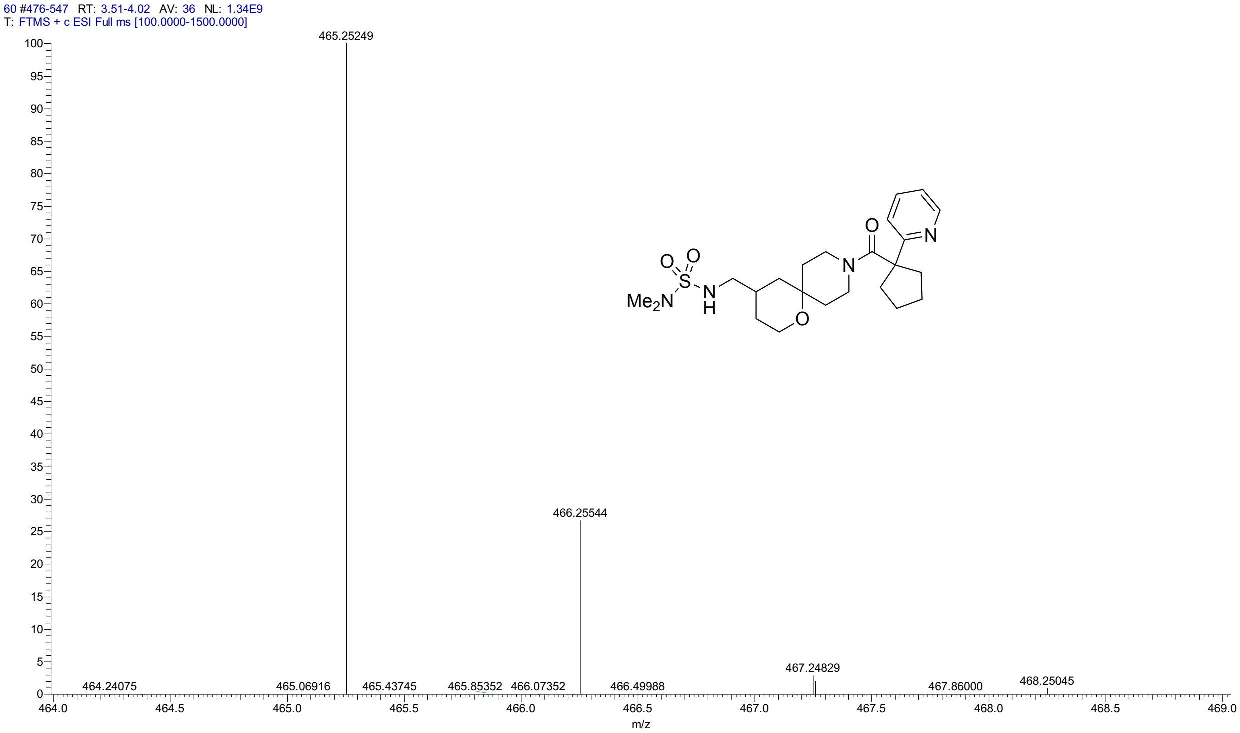


HPLC for **6k**

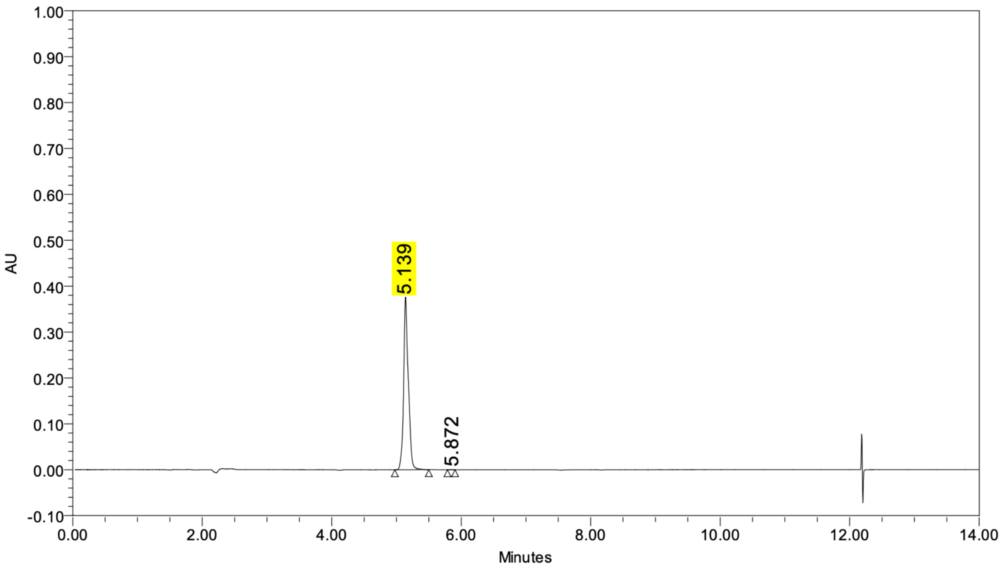


**Peak Results**

| | Name | RT | Height (μV) | Area (μV*sec) | % Area |
|---|---|---|---|---|---|
| 1 | | 5.139 | 376551 | 1900771 | 99.98 |
| 2 | | 5.872 | 129 | 400 | 0.02 |

$^{1}H$ NMR (400 MHz, DMSO-$d_6$, 80 °C) for **6l**

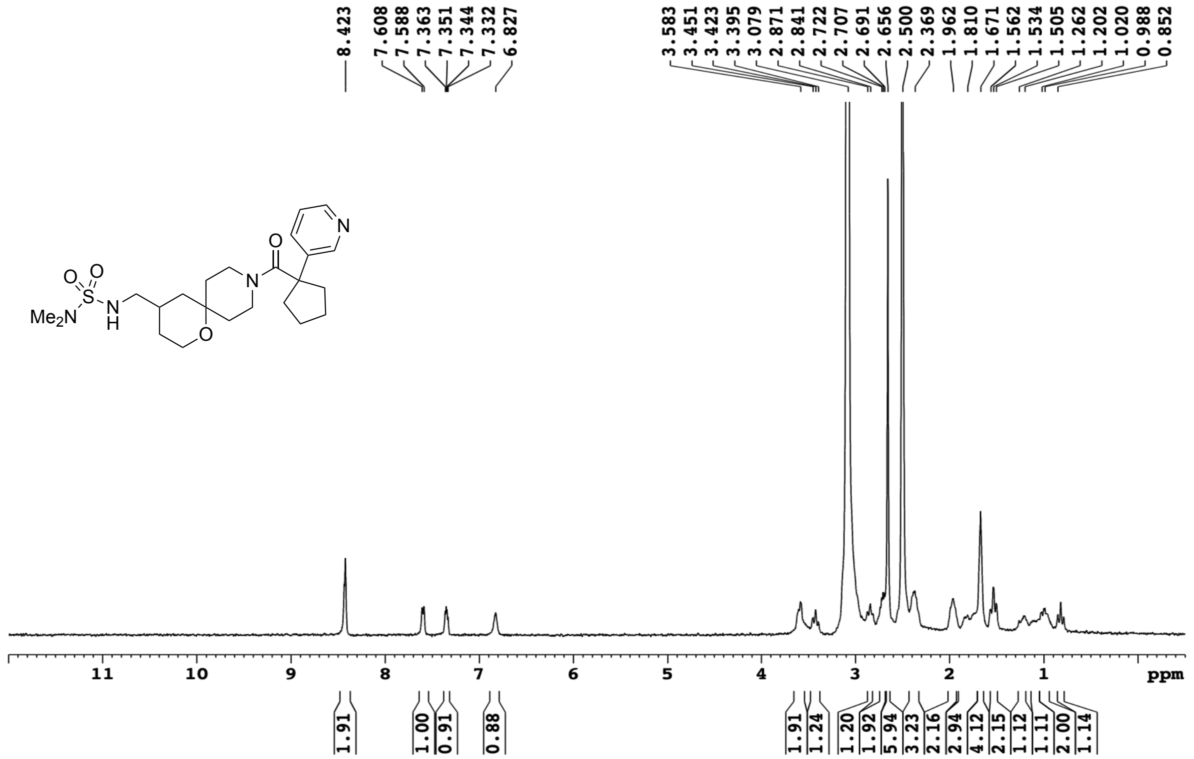


$^{13}C$ NMR (101 MHz, DMSO-$d_6$) for **6l**

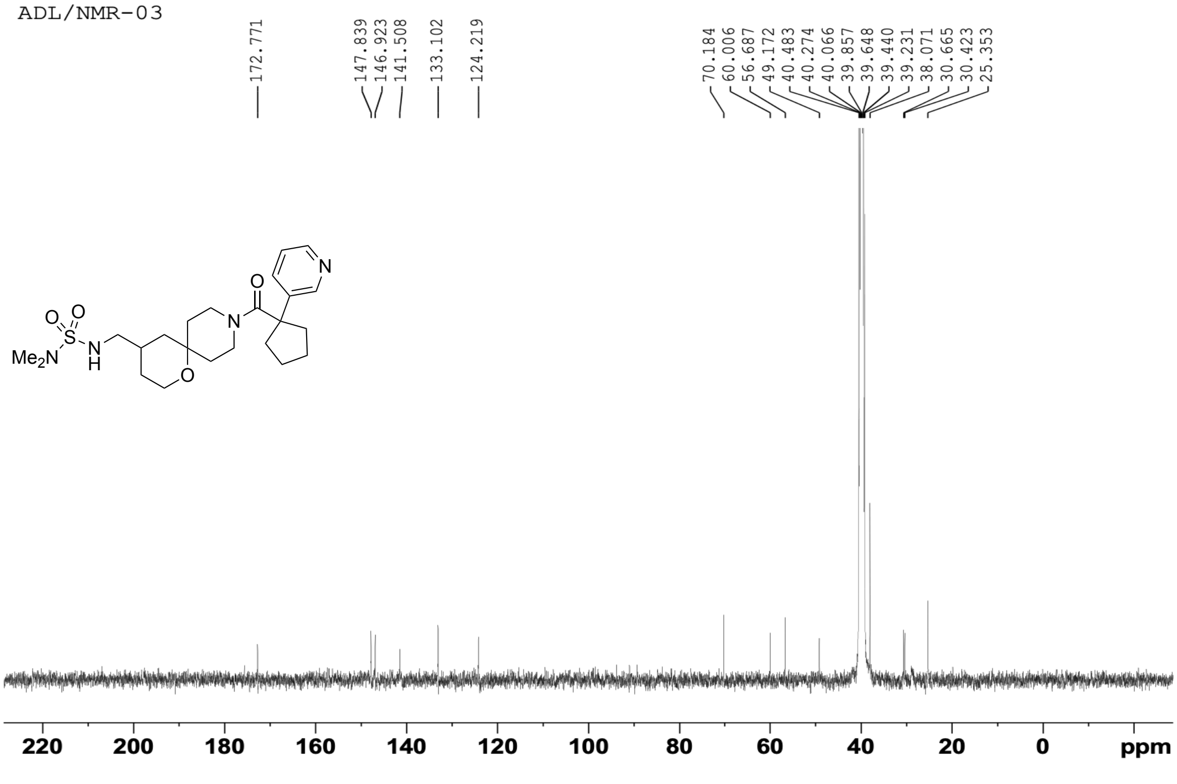

HRMS for **6l**

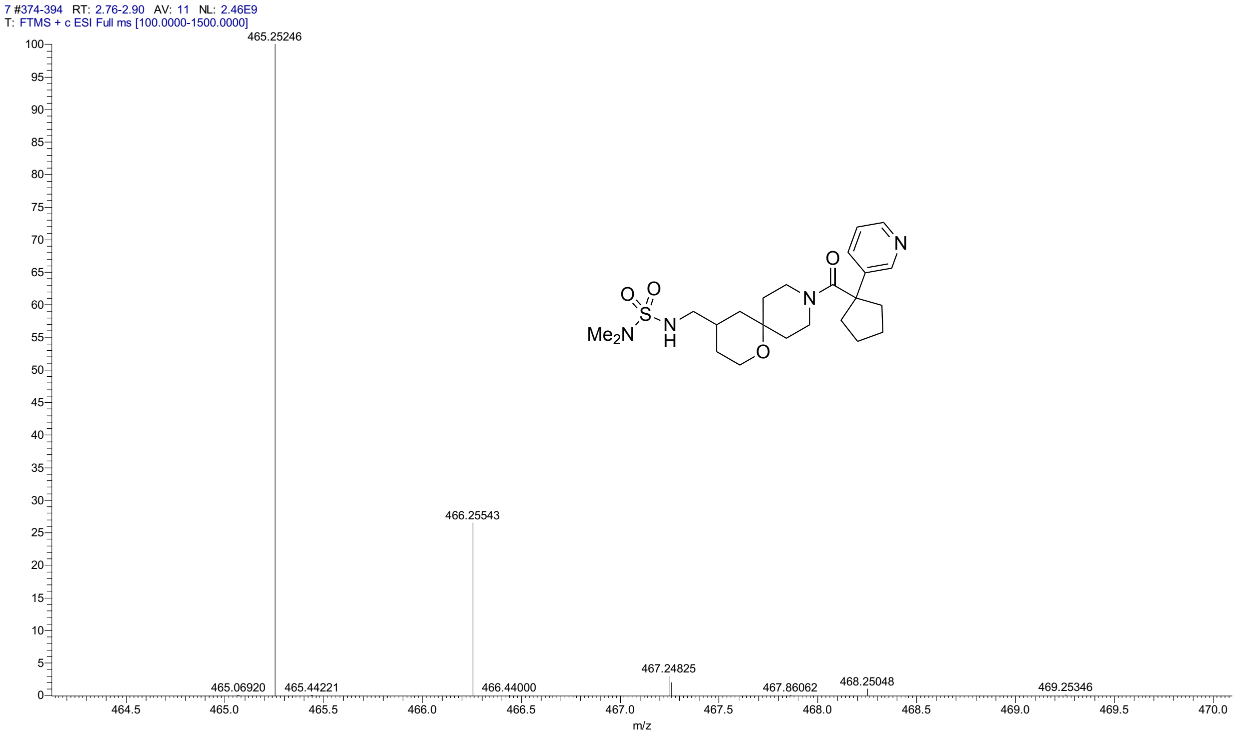


HPLC for **6l**

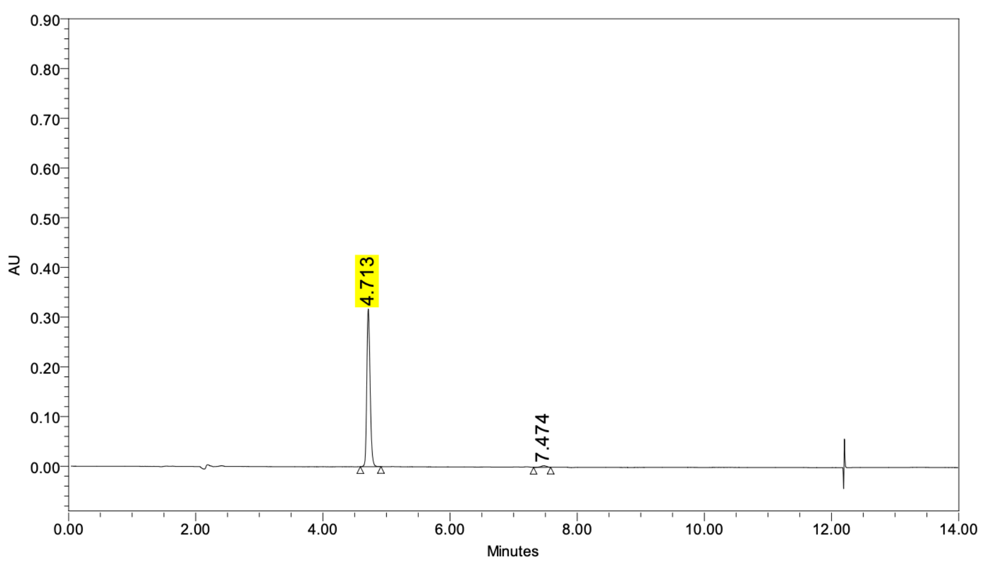


**Peak Results**

| | Name | RT | Height (µV) | Area (µV*sec) | % Area |
|---|---|---|---|---|---|
| 1 | | 4.713 | 317319 | 1146715 | 98.34 |
| 2 | | 7.474 | 3179 | 19403 | 1.66 |

$^{1}H$ NMR (400 MHz, DMSO-$d_6$, 80 °C) for **6m**

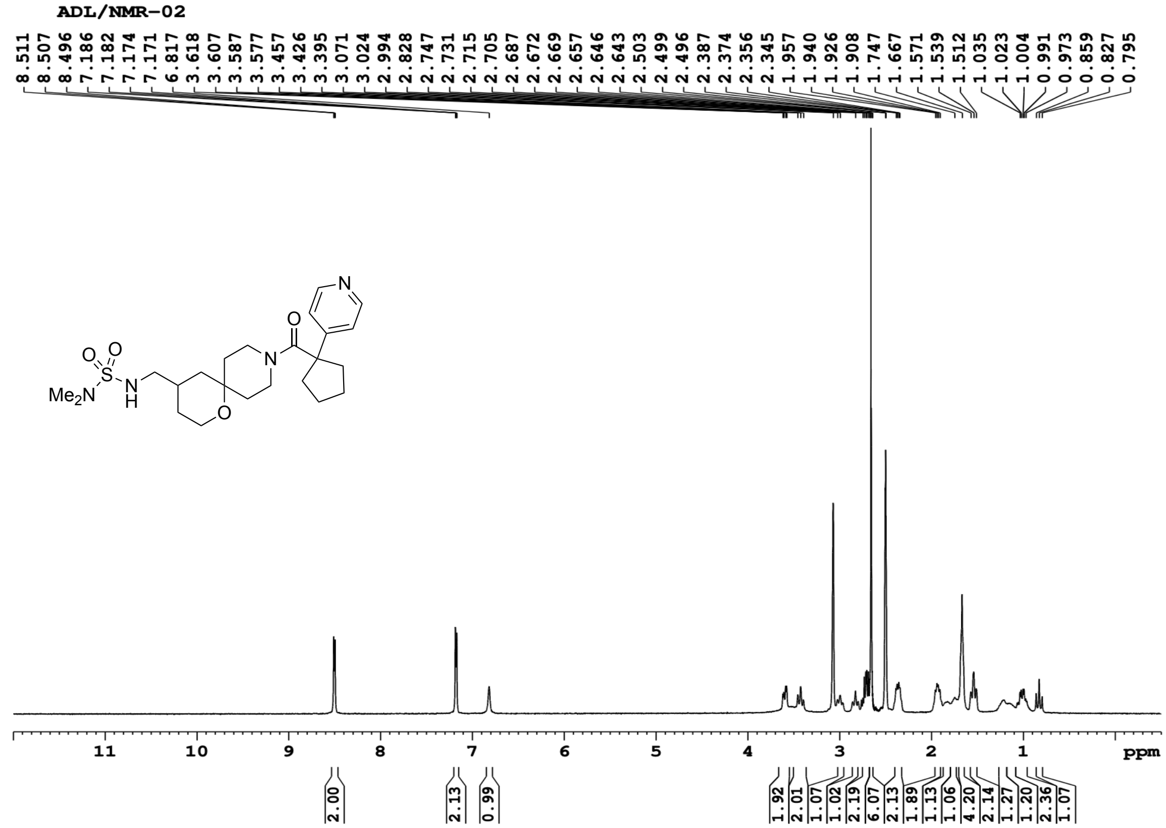


$^{13}C$ NMR (101 MHz, DMSO-$d_6$) for **6m**

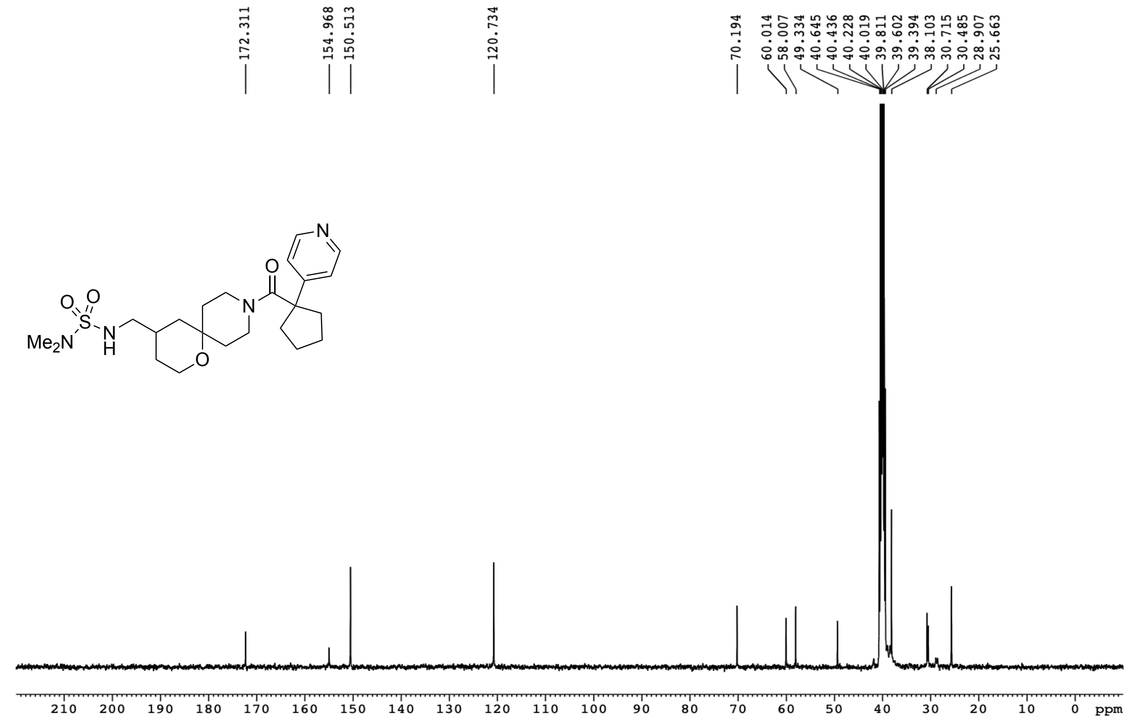

HRMS for **6m**

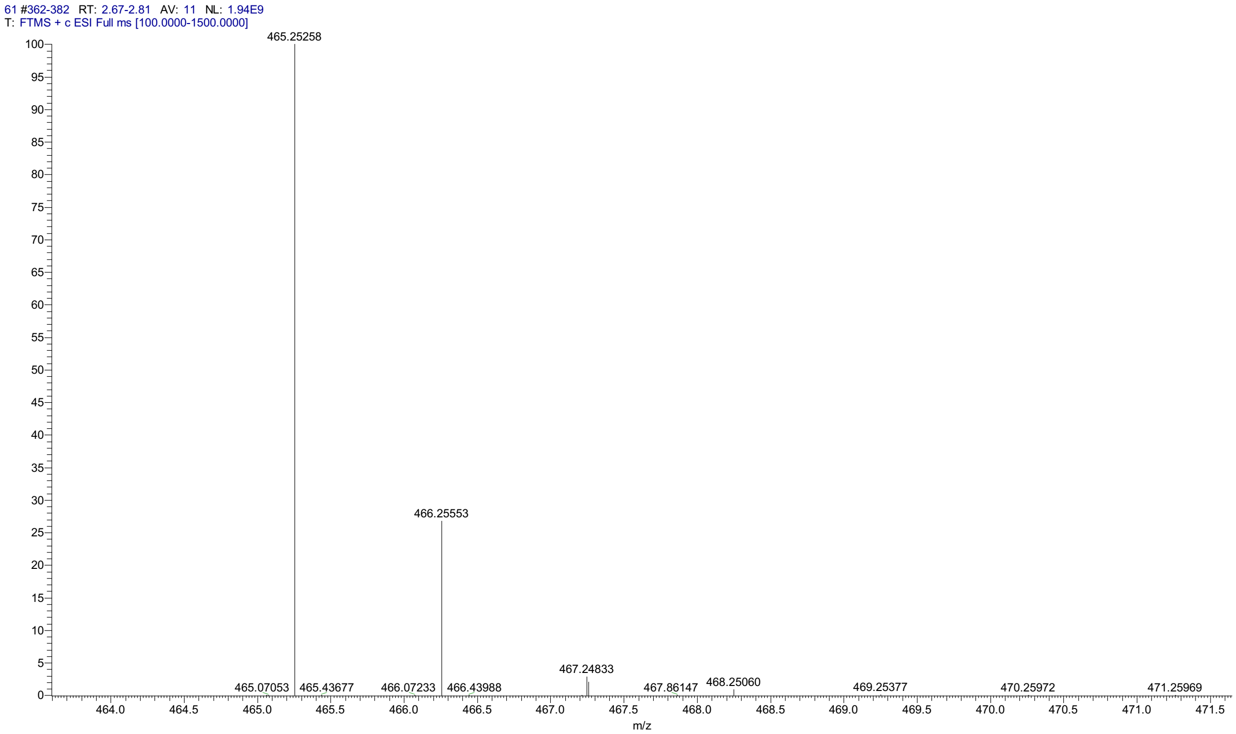


HPLC for **6m**

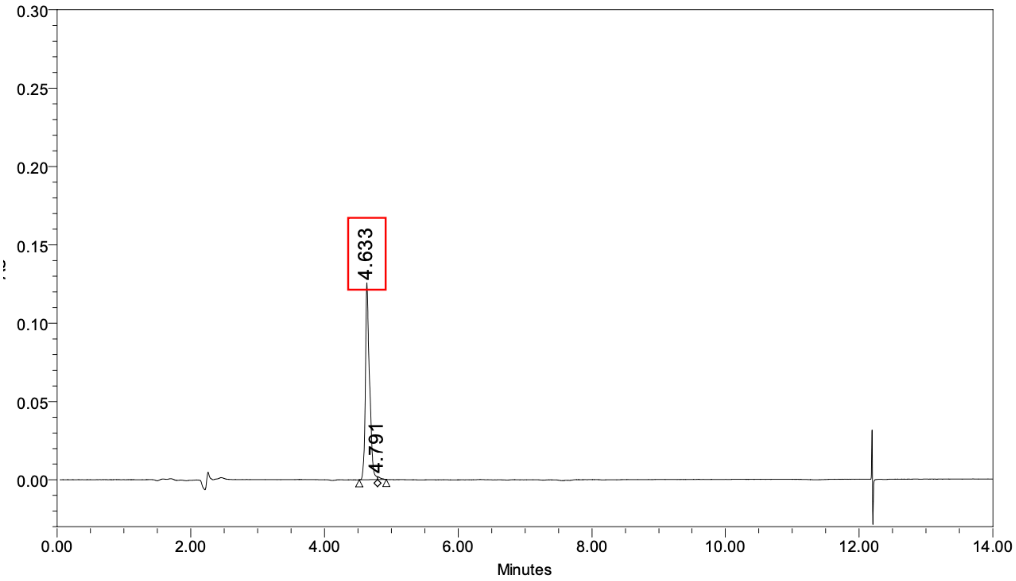


**Peak Results**

| | Name | RT | Height (µV) | Area (µV*sec) | % Area |
|---|---|---|---|---|---|
| 1 | | 4.633 | 125668 | 543018 | 98.96 |
| 2 | | 4.791 | 1804 | 5710 | 1.04 |

[1]H NMR (500 MHz, $CD_3CN$) for **7a**

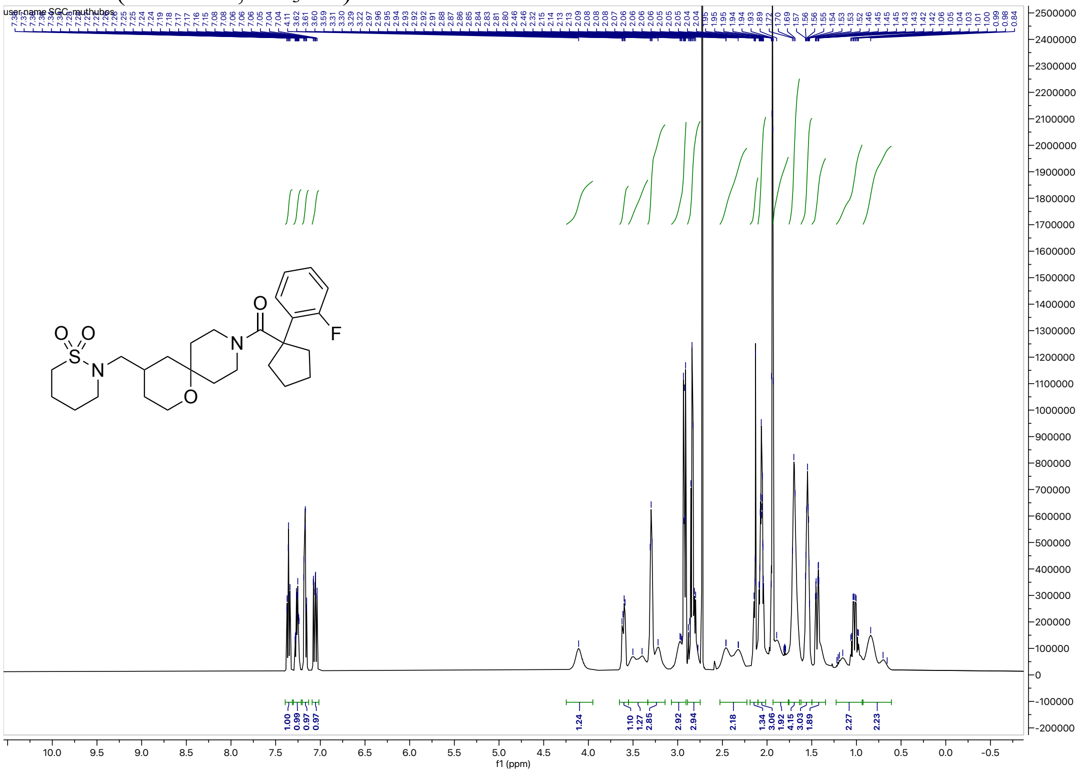


[13]C NMR (126 MHz, $CD_3CN$) for **7a**

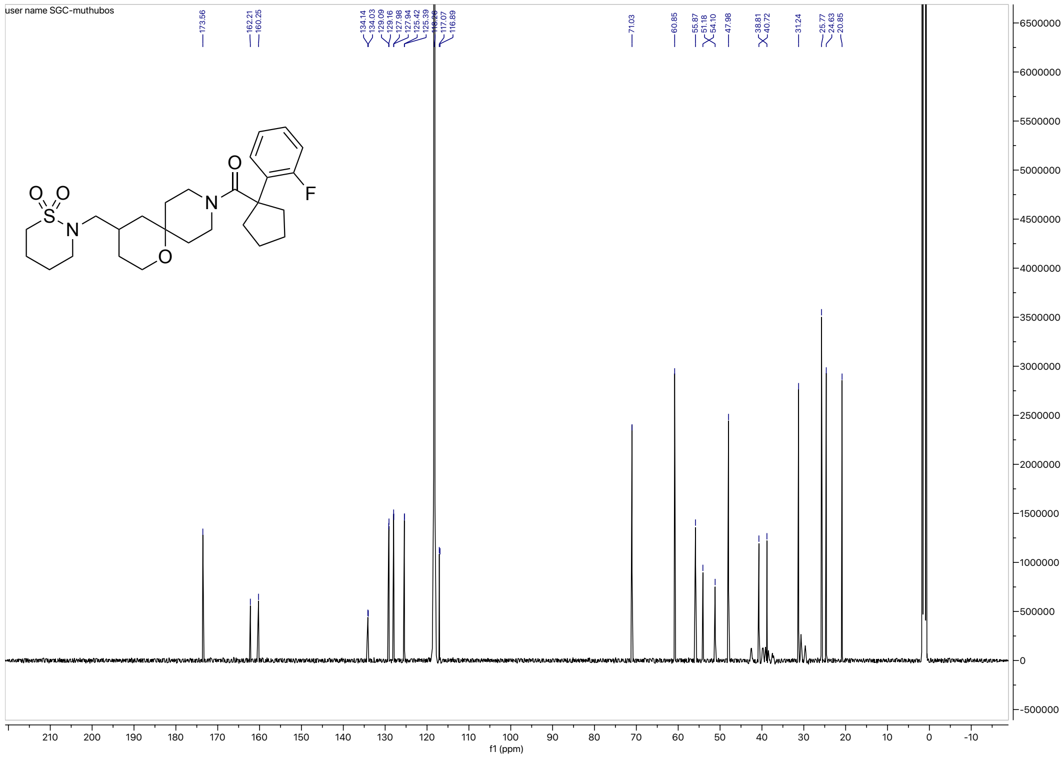

HRMS for **7a**

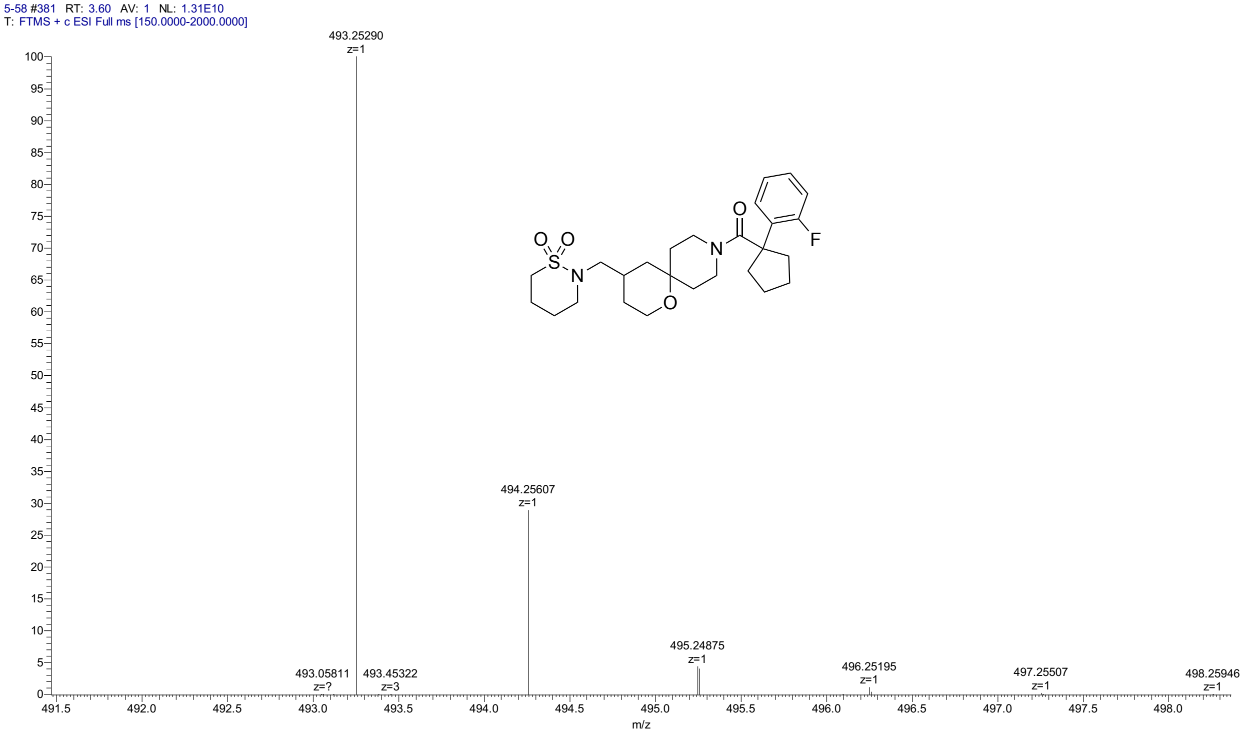


HPLC for **7a**

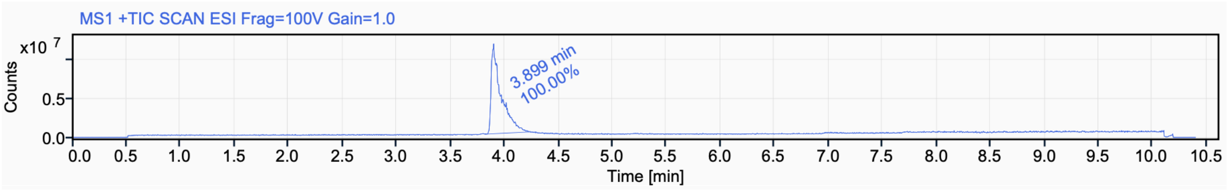


Signal Name MS1 +TIC SCAN ESI Frag=100V Gain=1.0

| RT (min) | Signal description | Area | Area% | Peak Resolution USP |
|---|---|---|---|---|
| 3.899 | MS1 +TIC SCAN ESI Frag=100V Gain=1.0 | 74583278.6 | 100.00 | |

$^{1}$H NMR (500 MHz, DMSO-$d_6$) for **7b**

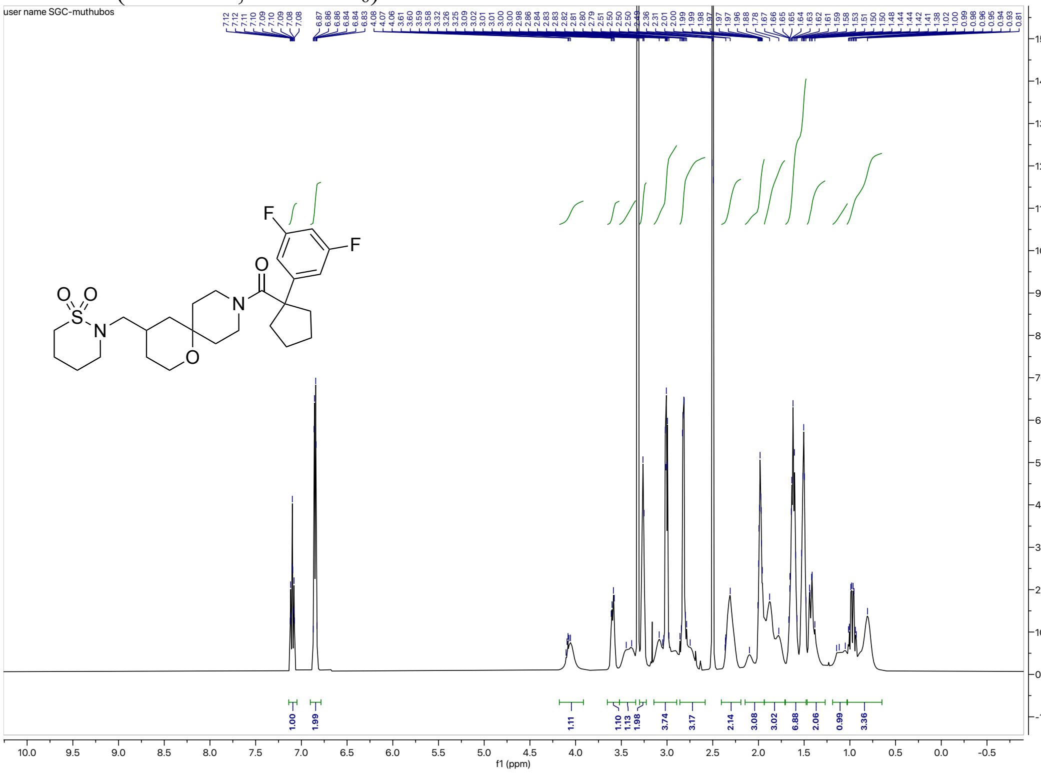


$^{13}$C NMR (126 MHz, DMSO-$d_6$) for **7b**

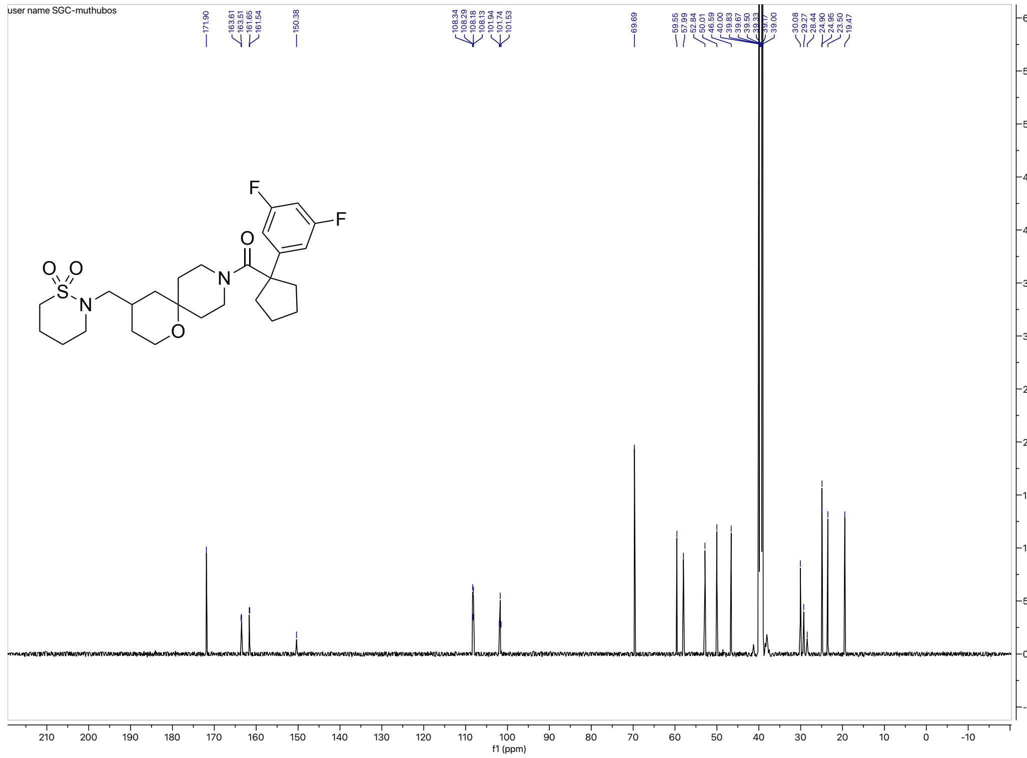

#### HRMS for **7b**

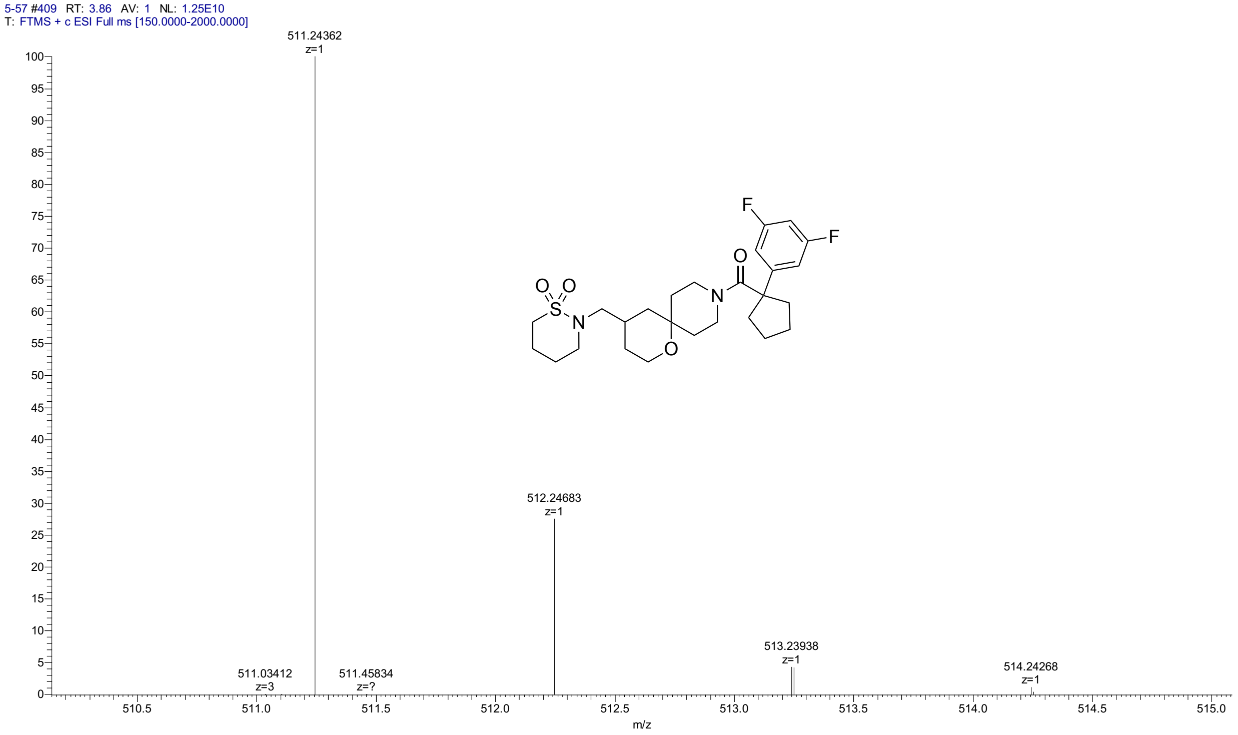


#### HPLC for **7b**

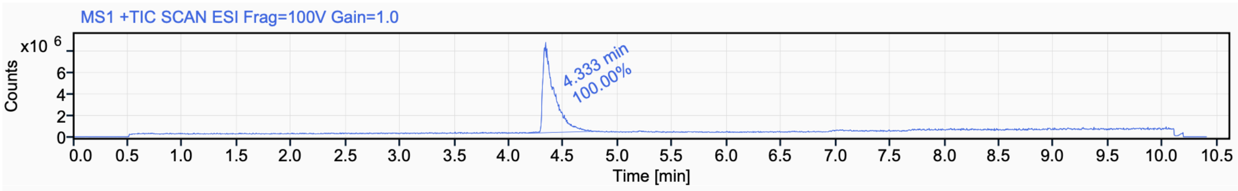


Signal Name MS1 +TIC SCAN ESI Frag=100V Gain=1.0

| RT (min) | Signal description | Area | Area% | Peak Resolution USP |
|---|---|---|---|---|
| 4.333 | MS1 +TIC SCAN ESI Frag=100V Gain=1.0 | 61165362.5 | 100.00 | |

$^{1}$H NMR (500 MHz, $CD_3CN$) for **7c**

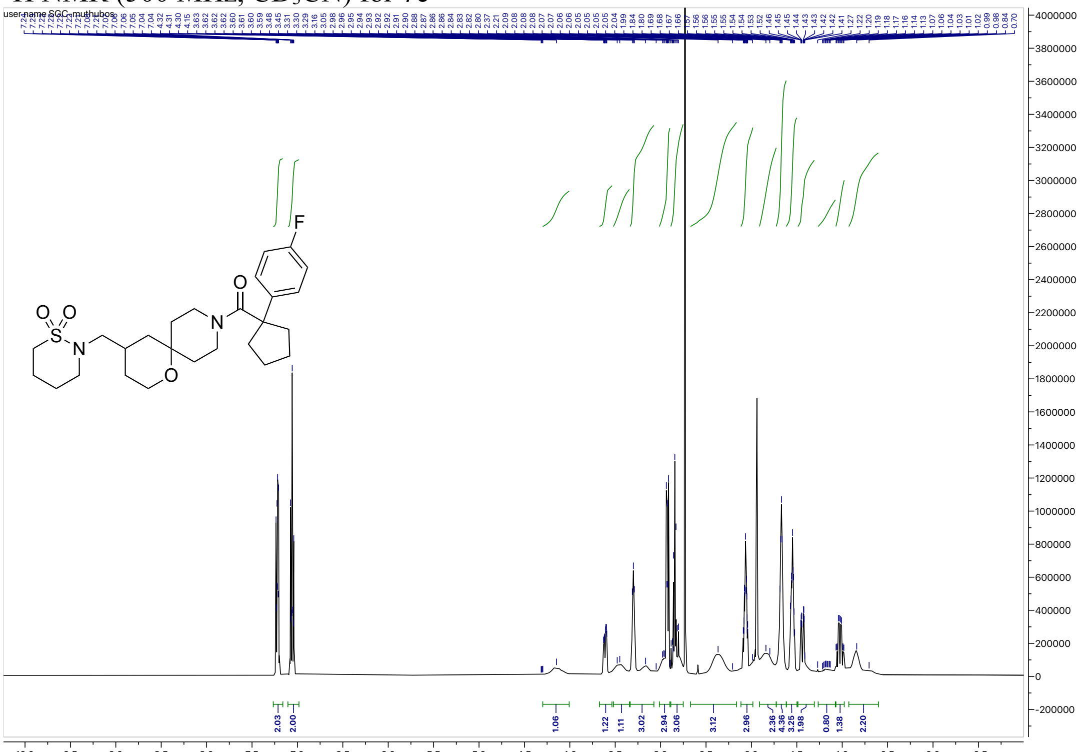

$^{13}$C NMR (126 MHz, $CD_3CN$) for **7c**

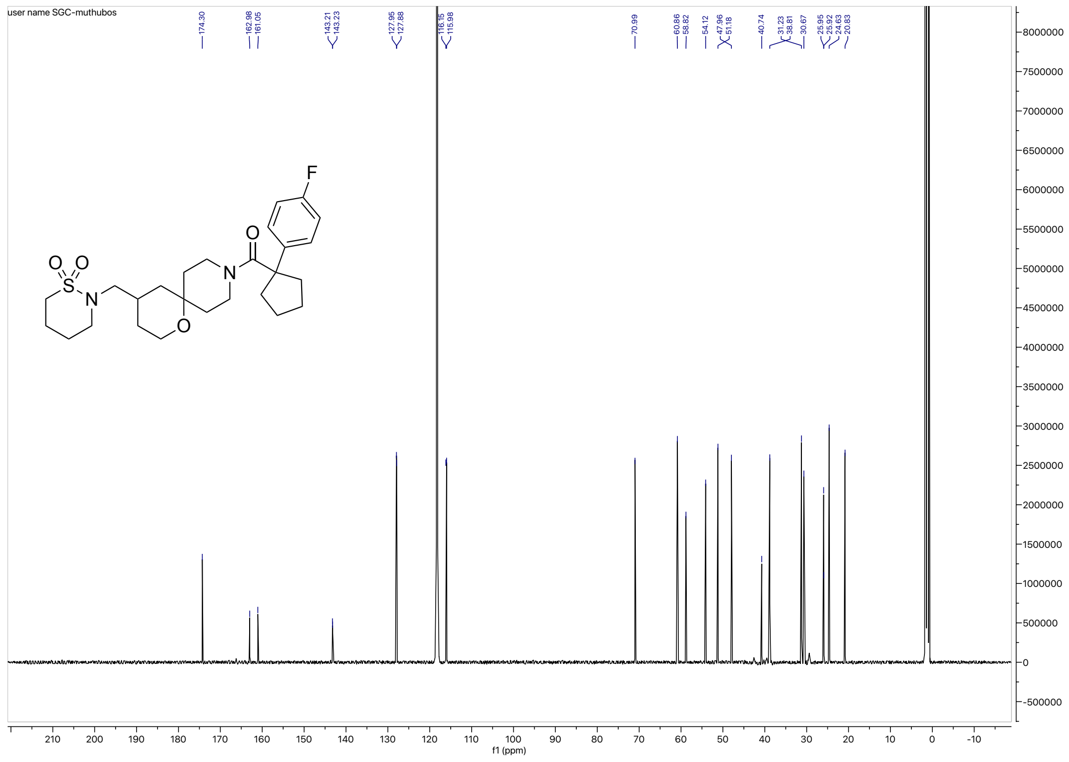

#### HRMS for **7c**

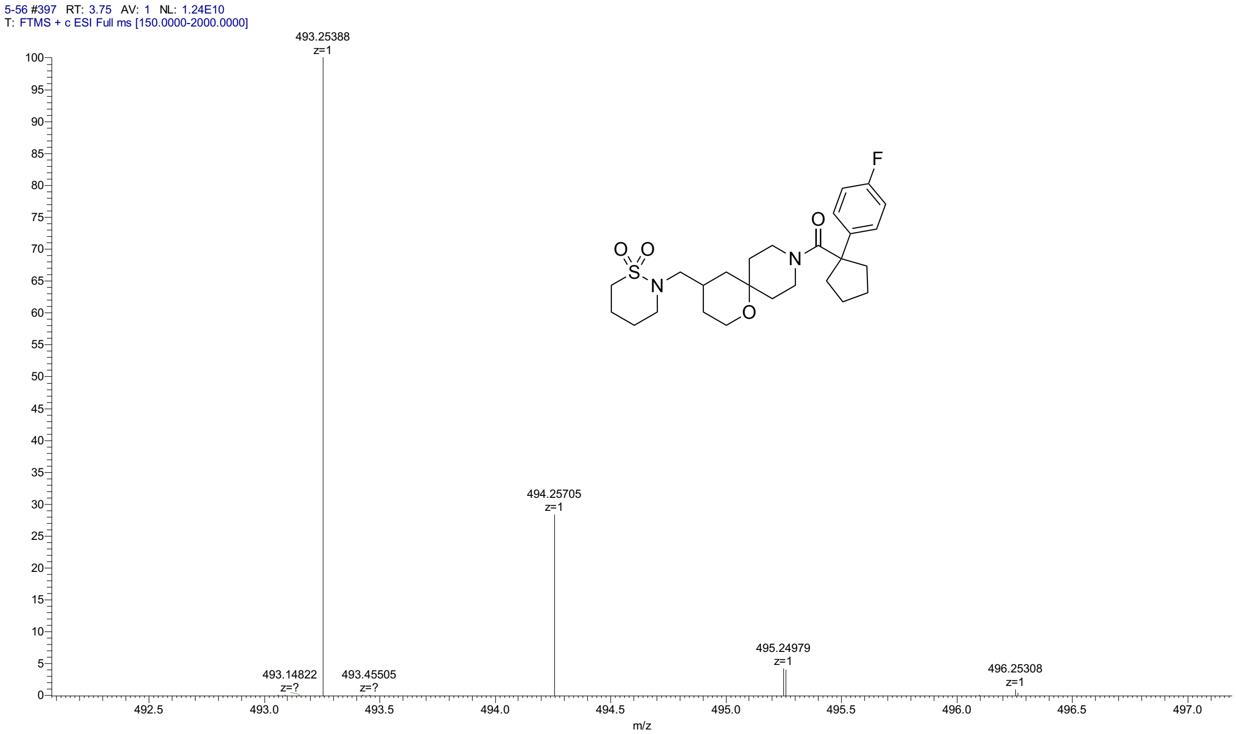


#### HPLC for **7c**

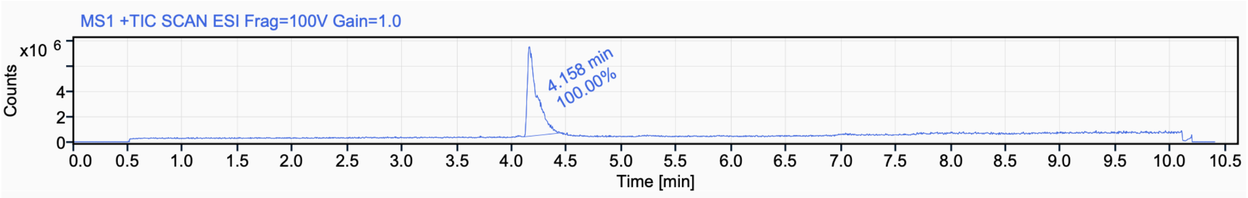


Signal Name MS1 +TIC SCAN ESI Frag=100V Gain=1.0

| RT (min) | Signal description | Area | Area% | Peak Resolution USP |
|---|---|---|---|---|
| 4.158 | MS1 +TIC SCAN ESI Frag=100V Gain=1.0 | 44242672.1 | 100.00 | |

### [1]H NMR (700 MHz, $CDCl_3$) for **IIa**

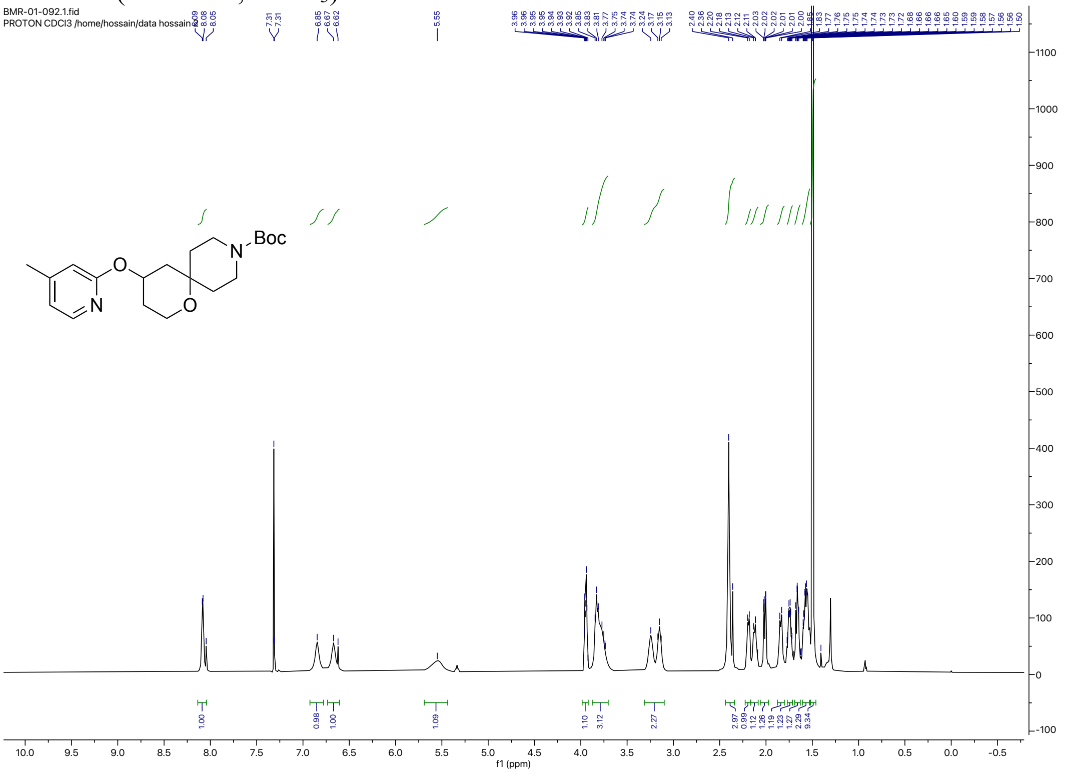


### [1]H NMR (400 MHz, $CDCl_3$) for **IIb**

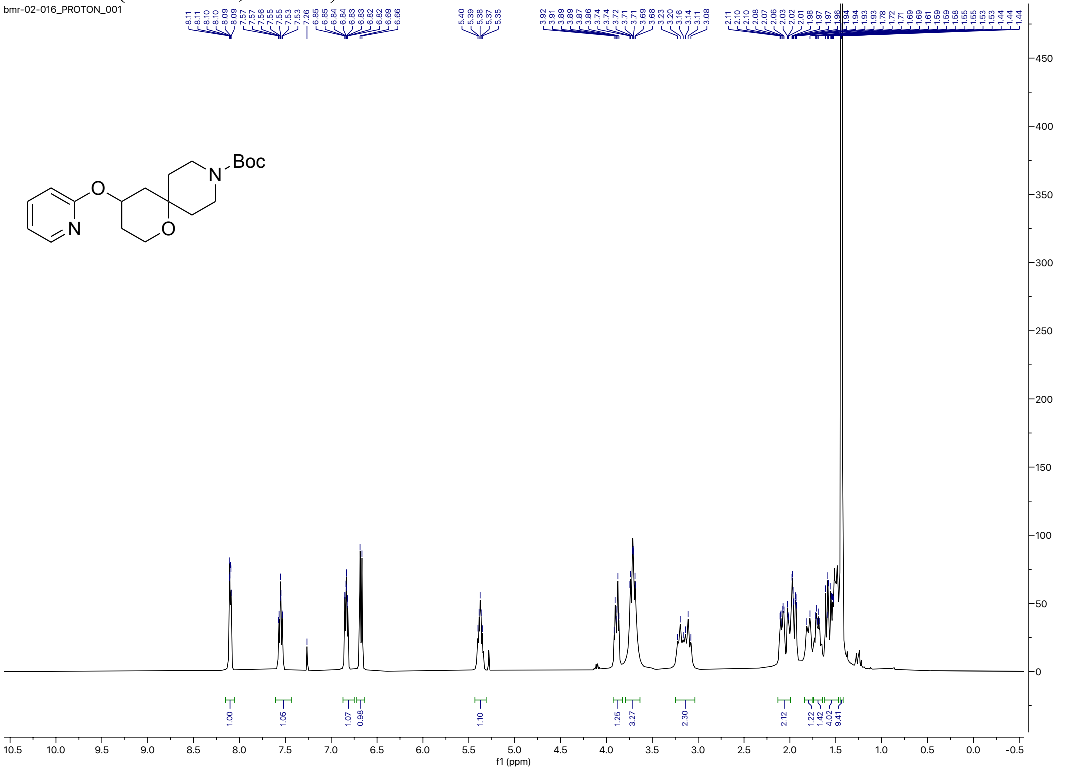

$^{1}$H NMR (400 MHz, $CDCl_3$) for **VI**

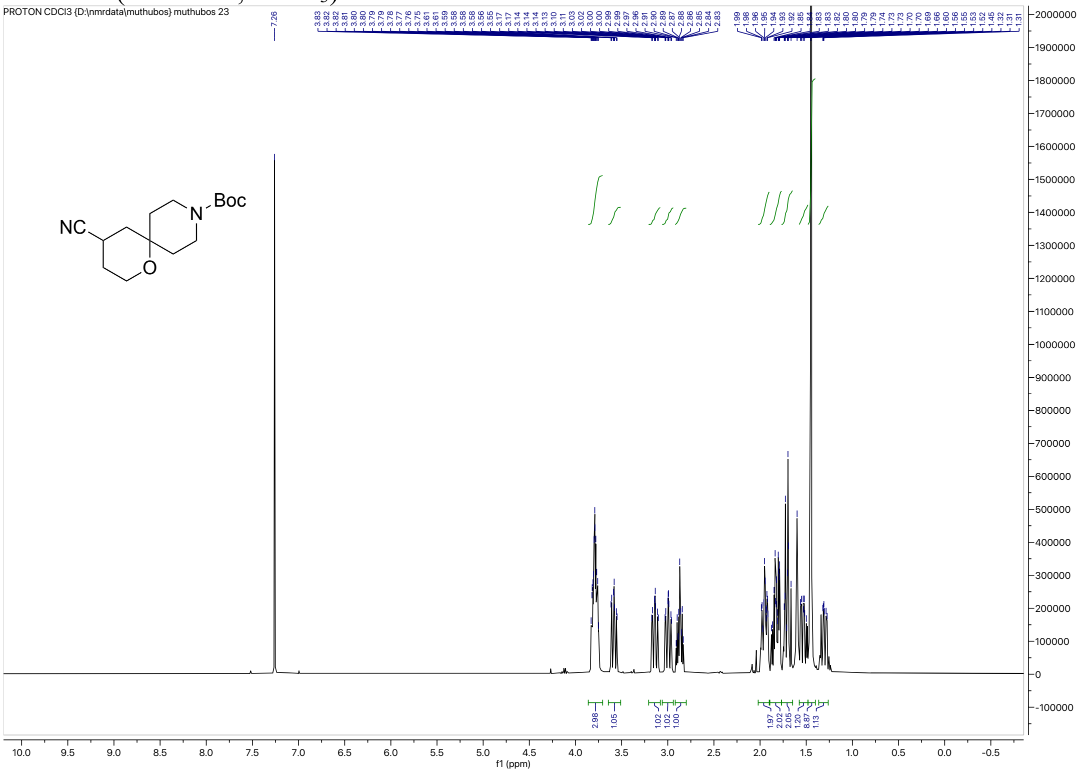


$^{1}$H NMR (500 MHz, $CDCl_3$)) for **VII**

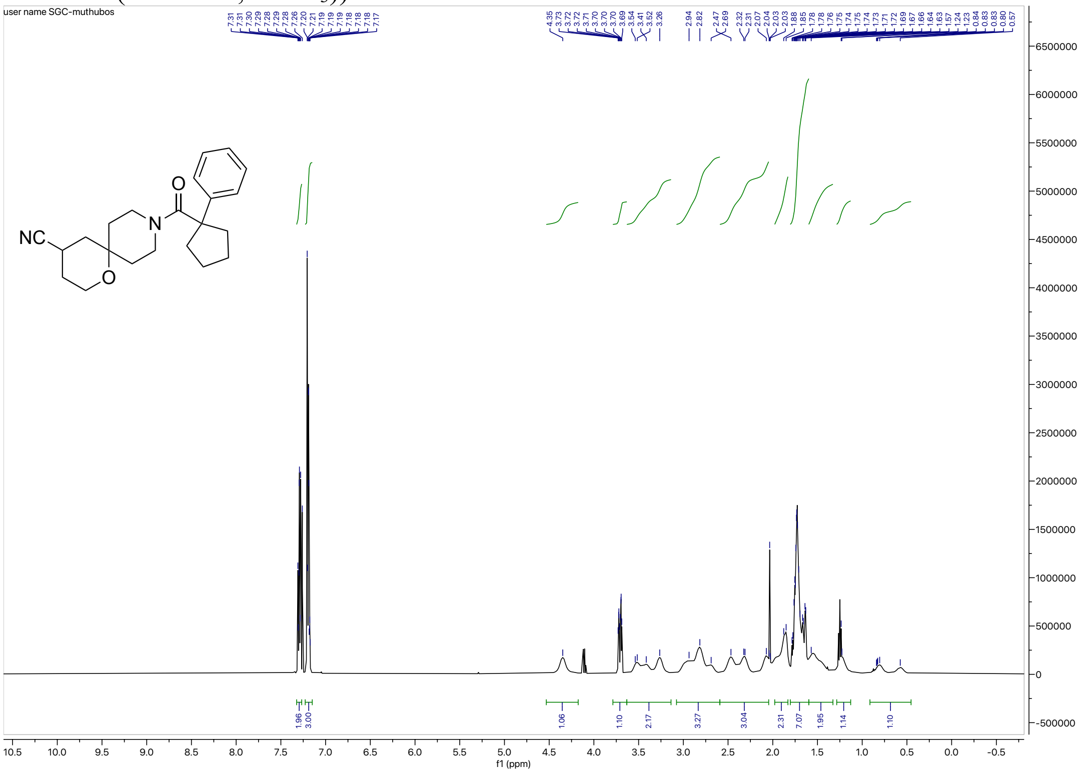

#### $^{1}$H NMR (700 MHz, $CDCl_3$) for **VIII**

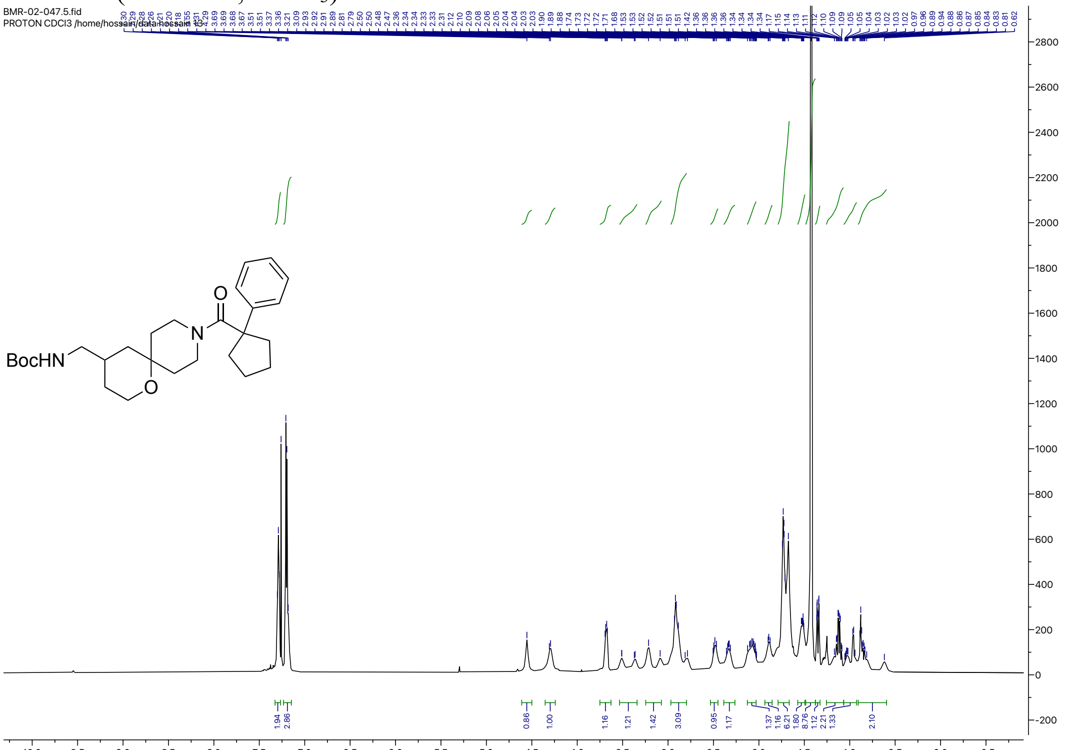


#### $^{1}$H NMR (500 MHz, $CDCl_3$) for **IXa**

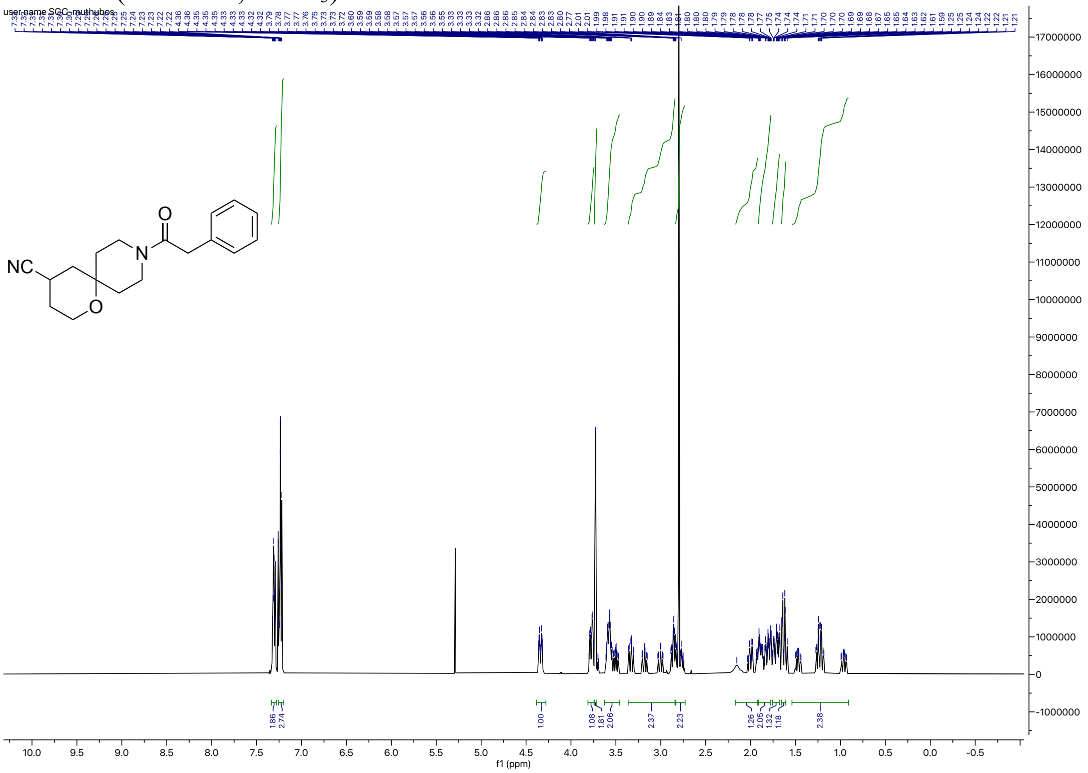

#### $^{1}$H NMR (500 MHz, $CDCl_3$) for **IXb**

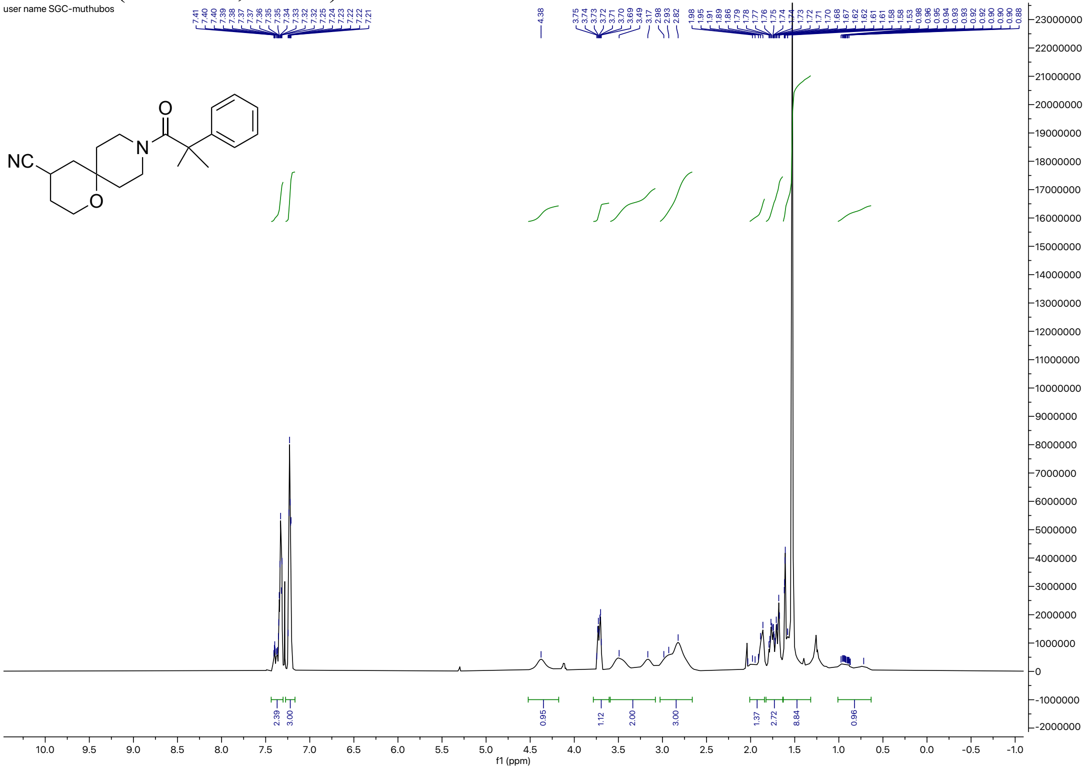


#### $^{1}$H NMR (500 MHz, $CDCl_3$) for **IXc**

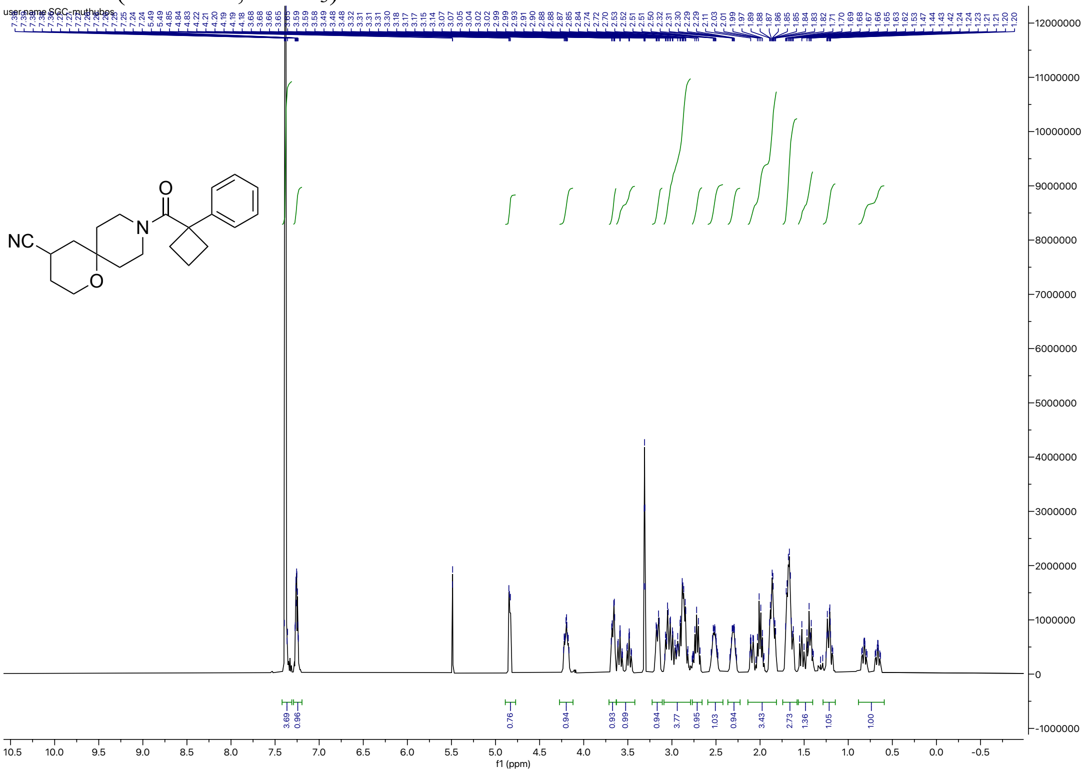

$^{1}$H NMR (500 MHz, DMSO-$d_6$) for **IXe**

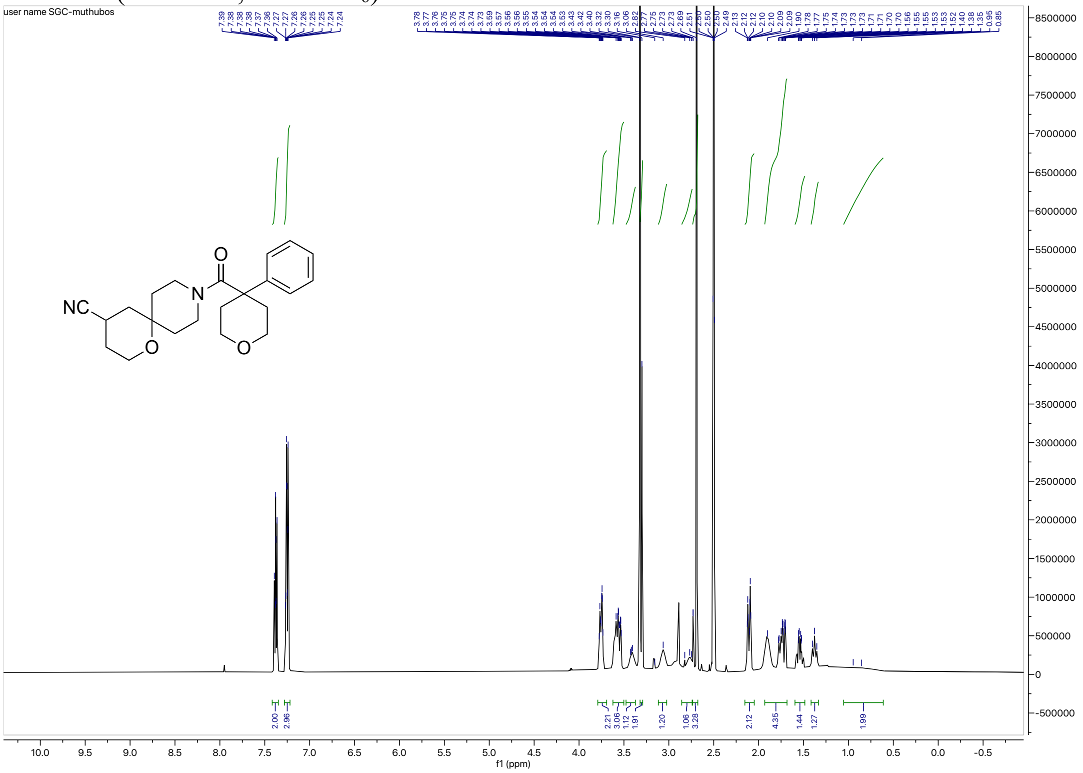

$^{1}$H NMR (500 MHz, DMSO-$d_6$) for **IXf**

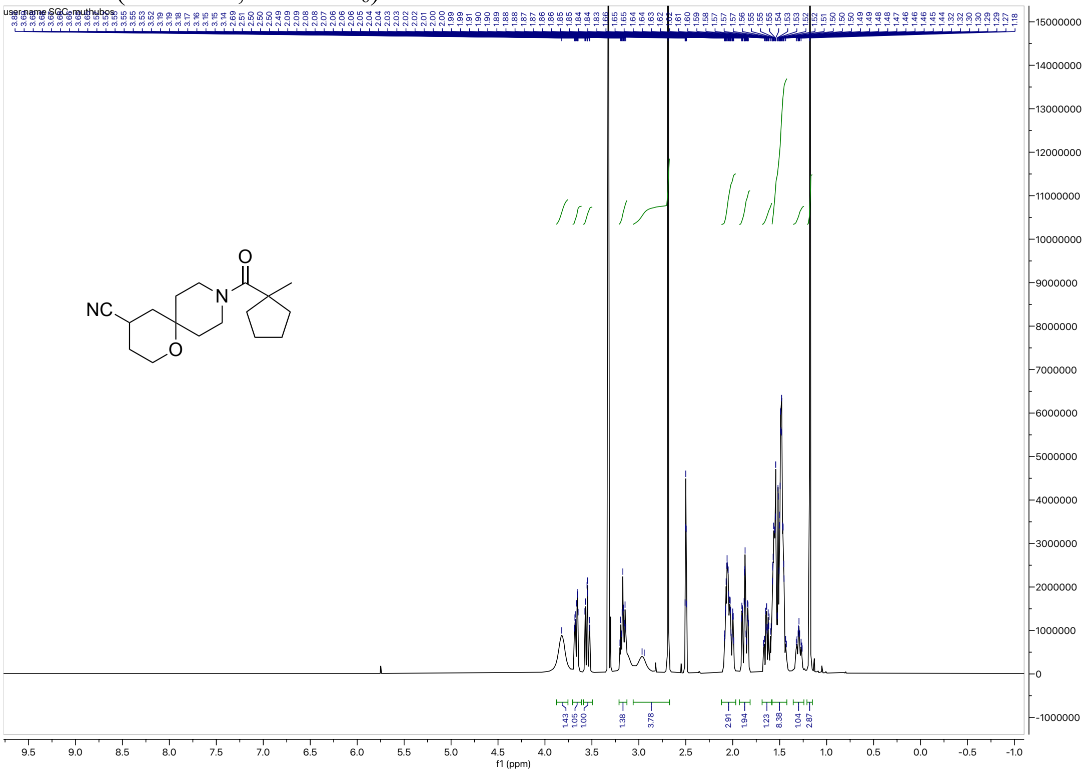

$^{1}H$ NMR (500 MHz, $CDCl_3$) for **IXh**

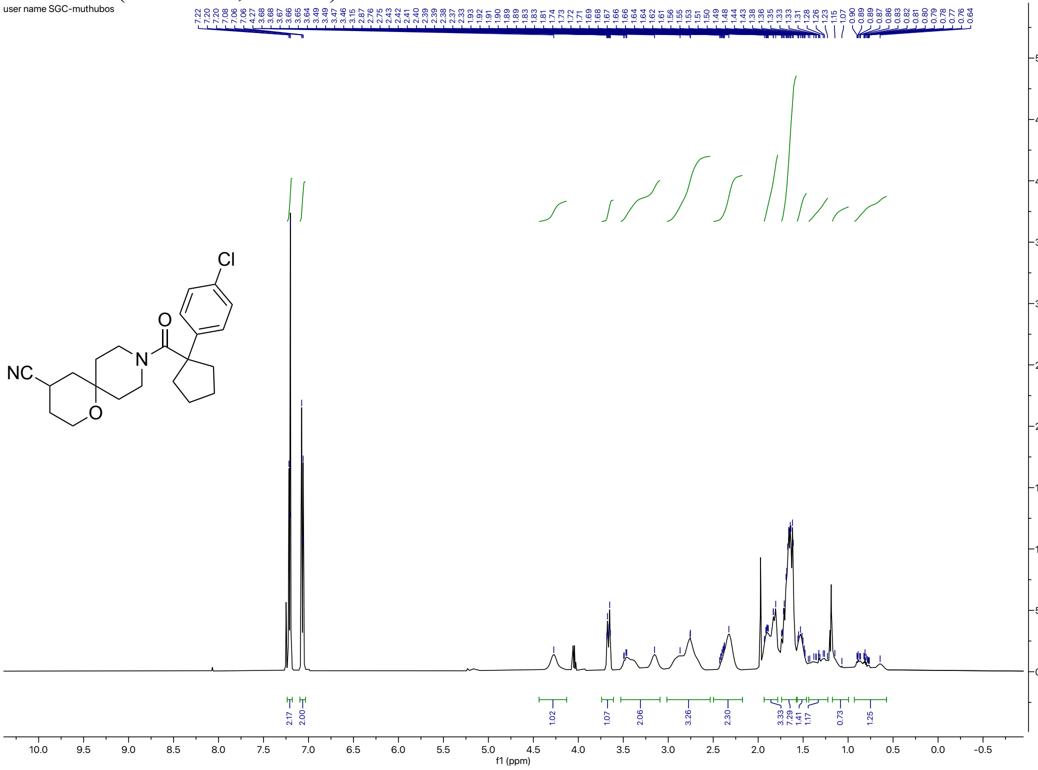


$^{1}H$ NMR (500 MHz, $CDCl_3$) for **IXi**

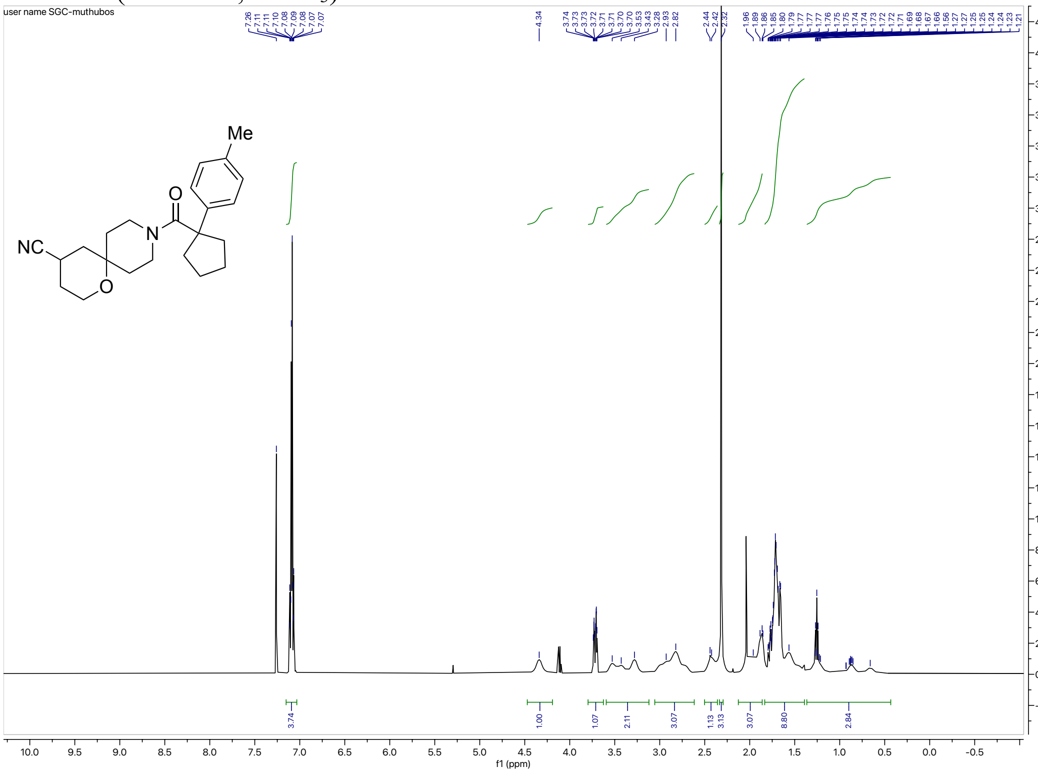

$^{1}$H NMR (500 MHz, $CDCl_3$) for **IXj**

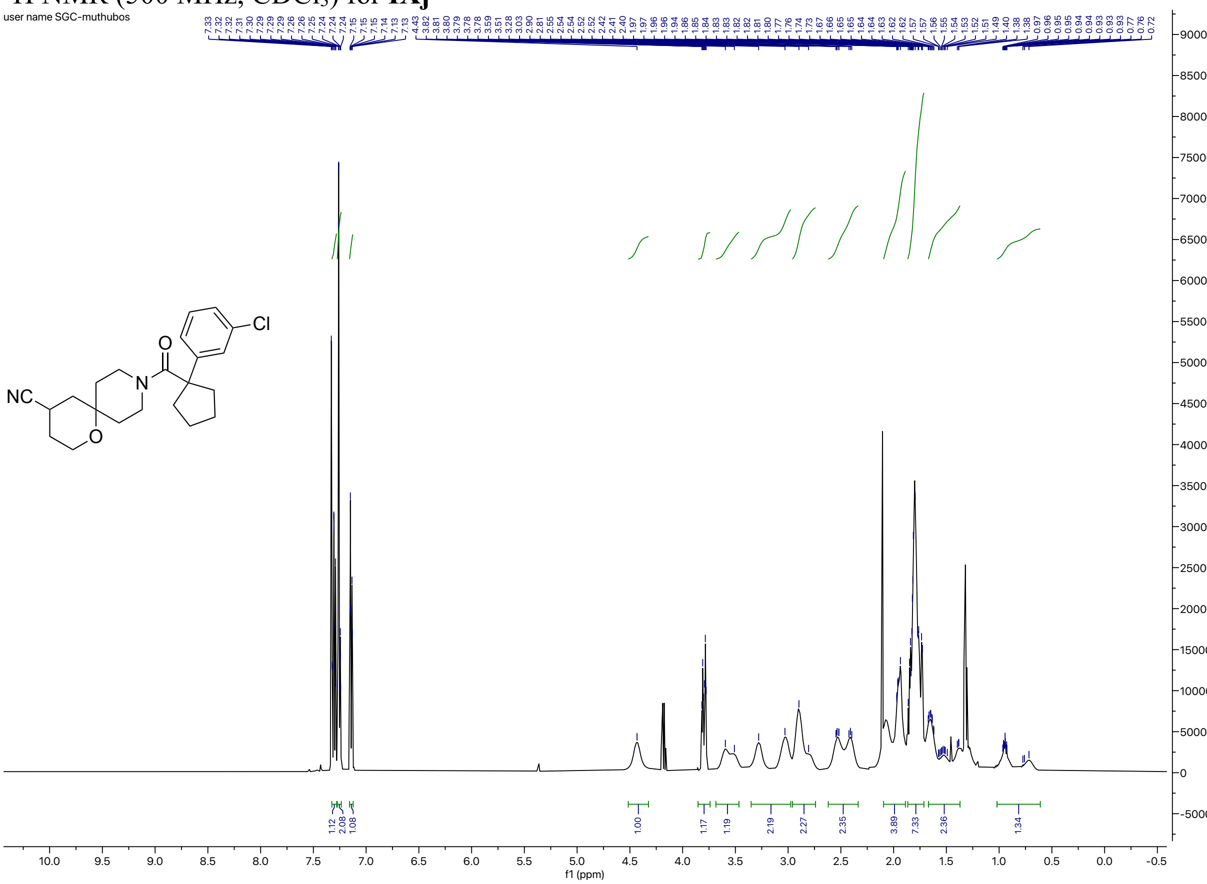


$^{1}$H NMR (500 MHz, $CDCl_3$) for **Xa**

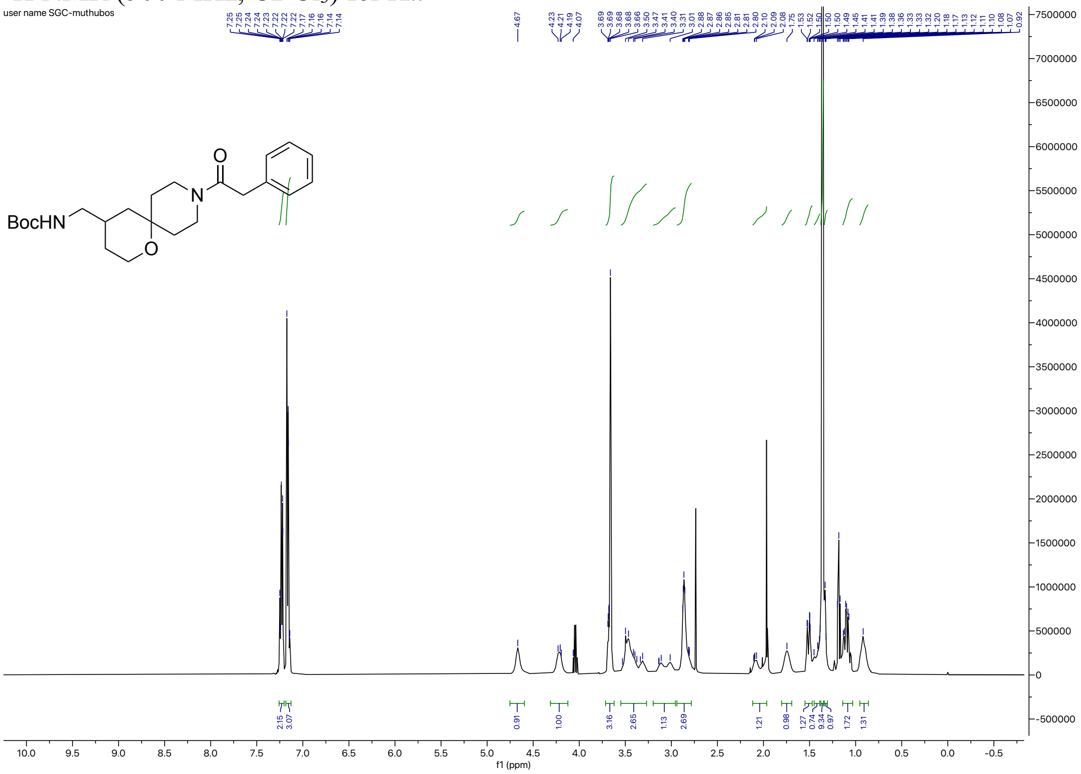

#### $^1$H NMR (500 MHz, $CDCl_3$) for **Xb**

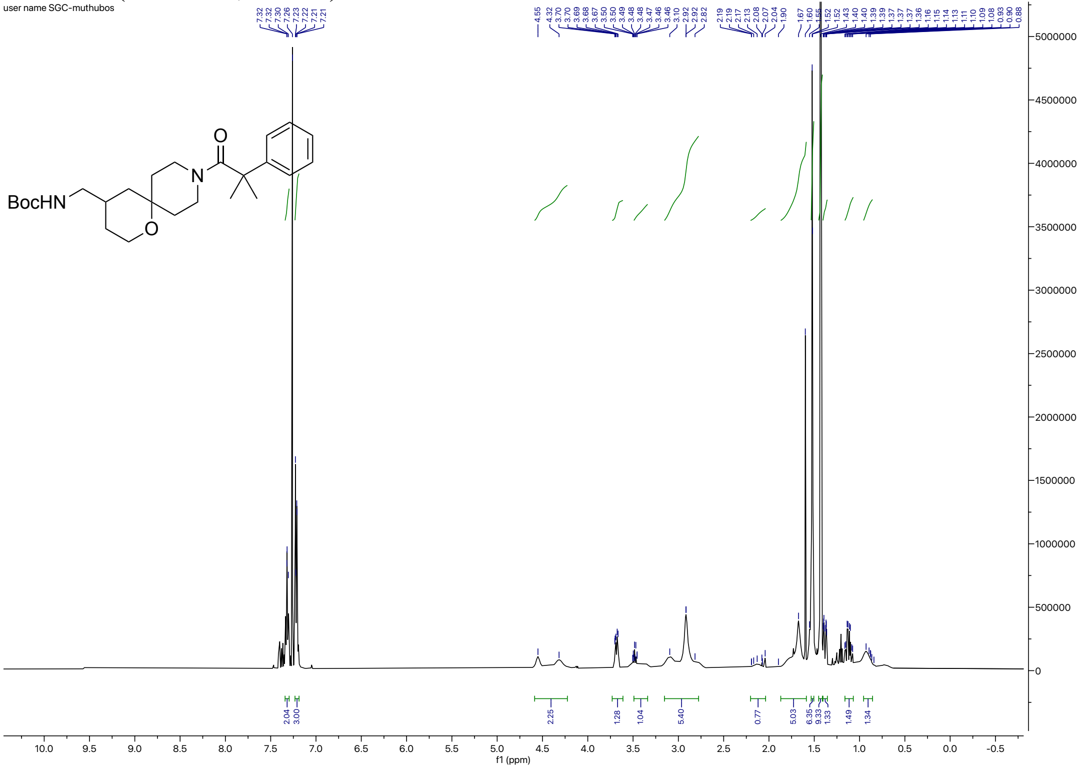


#### $^1$H NMR (500 MHz, $CDCl_3$) for **Xc**

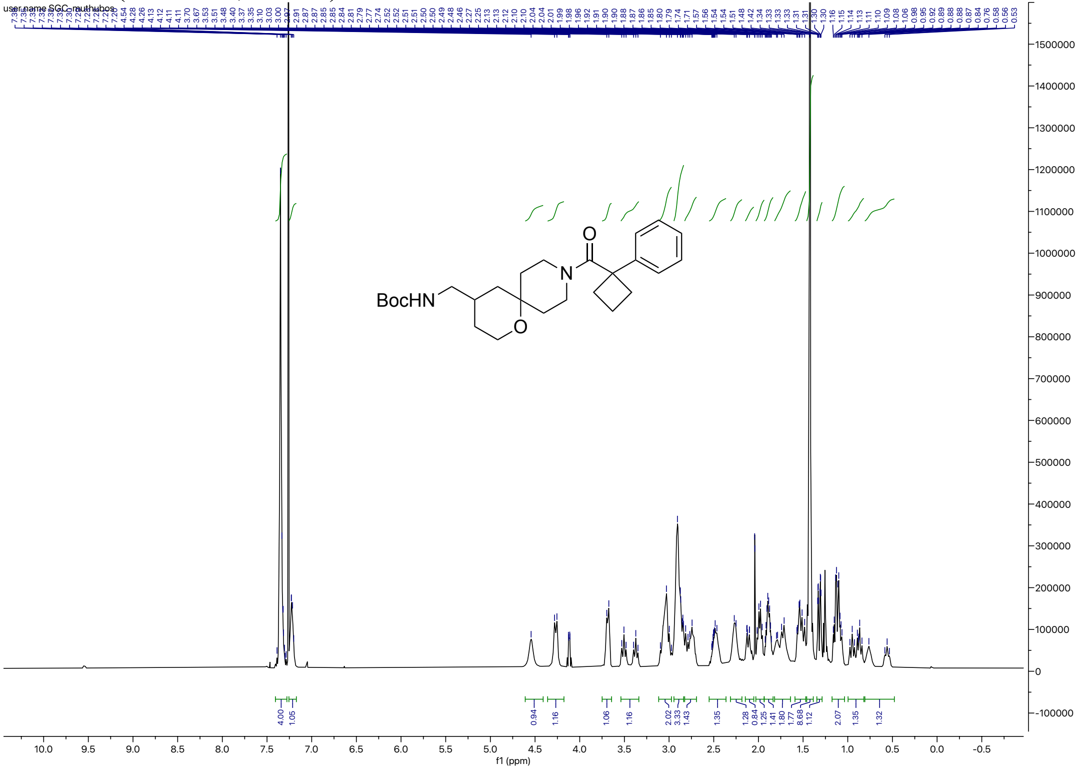

[1]H NMR (500 MHz, $CDCl_3$) for **Xd**

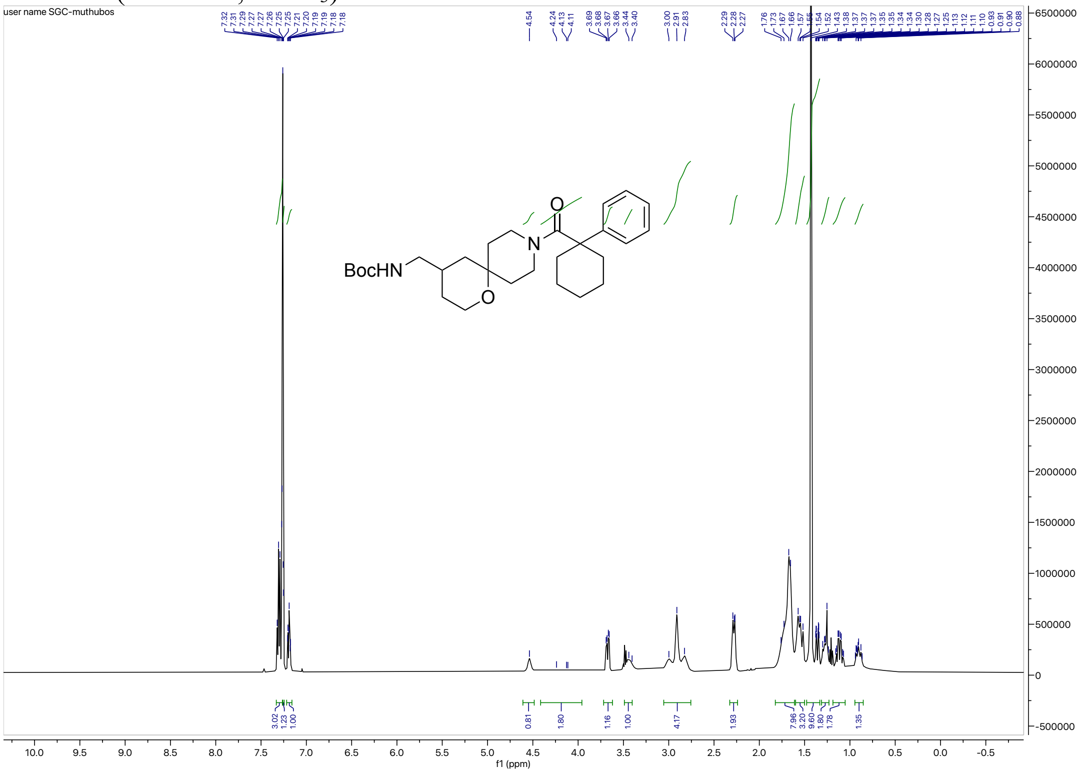

[1]H NMR (500 MHz, $CDCl_3$) for **Xe**

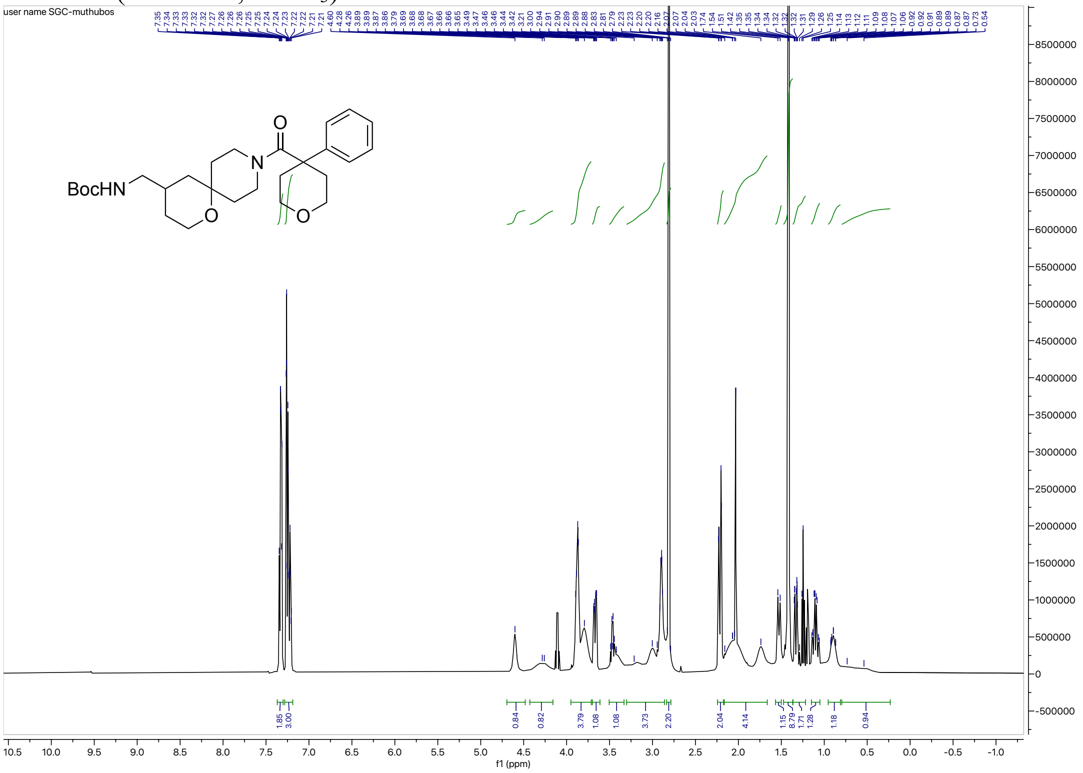

#### $^1$H NMR (500 MHz, $CDCl_3$) for **Xf**

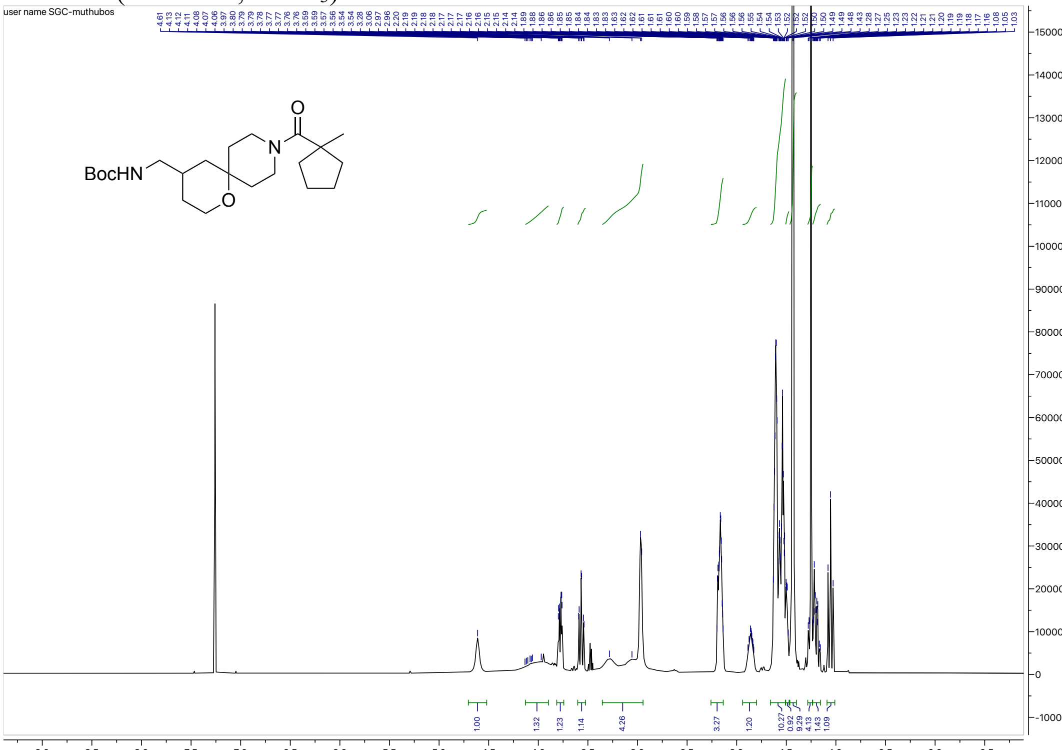


#### $^1$H NMR (500 MHz, $CDCl_3$) for **Xg**

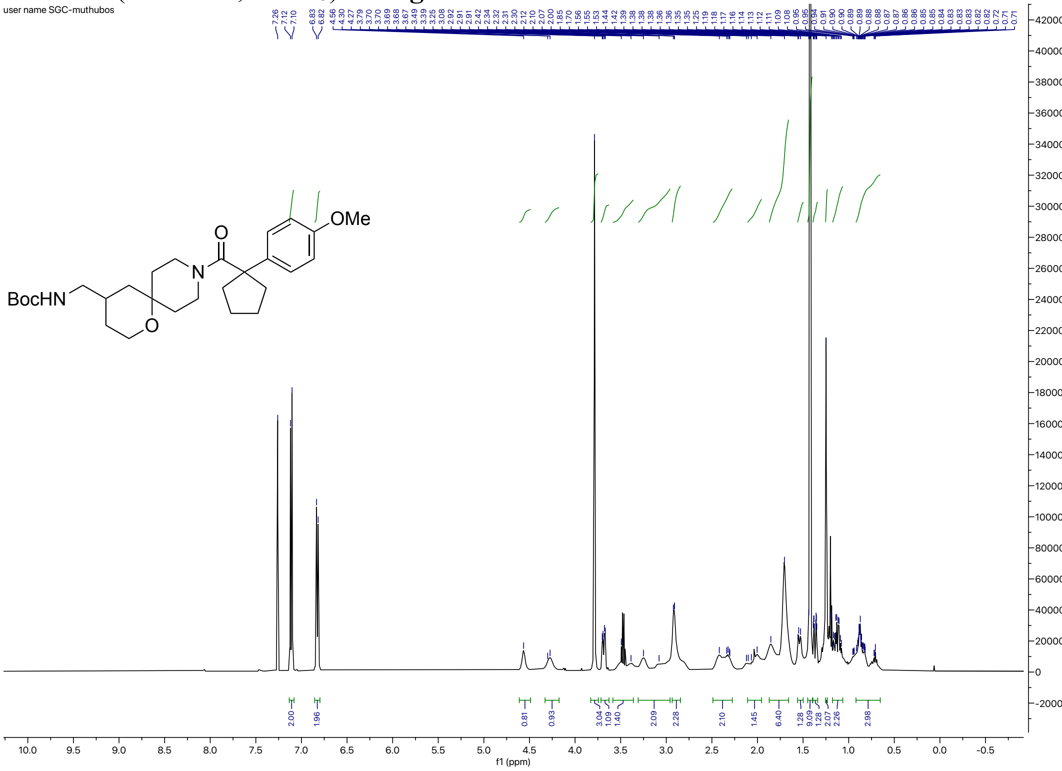

#### $^1H$ NMR (500 MHz, $CDCl_3$) for **Xh**

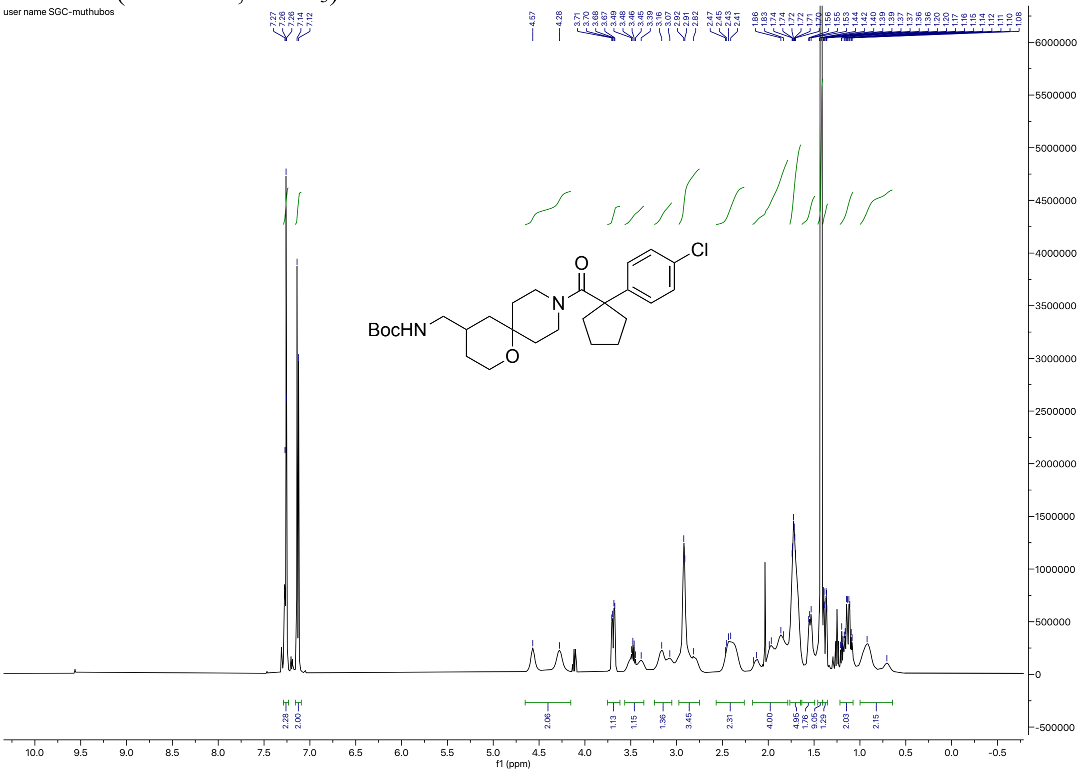


#### $^1H$ NMR (500 MHz, $CDCl_3$) for **Xi**

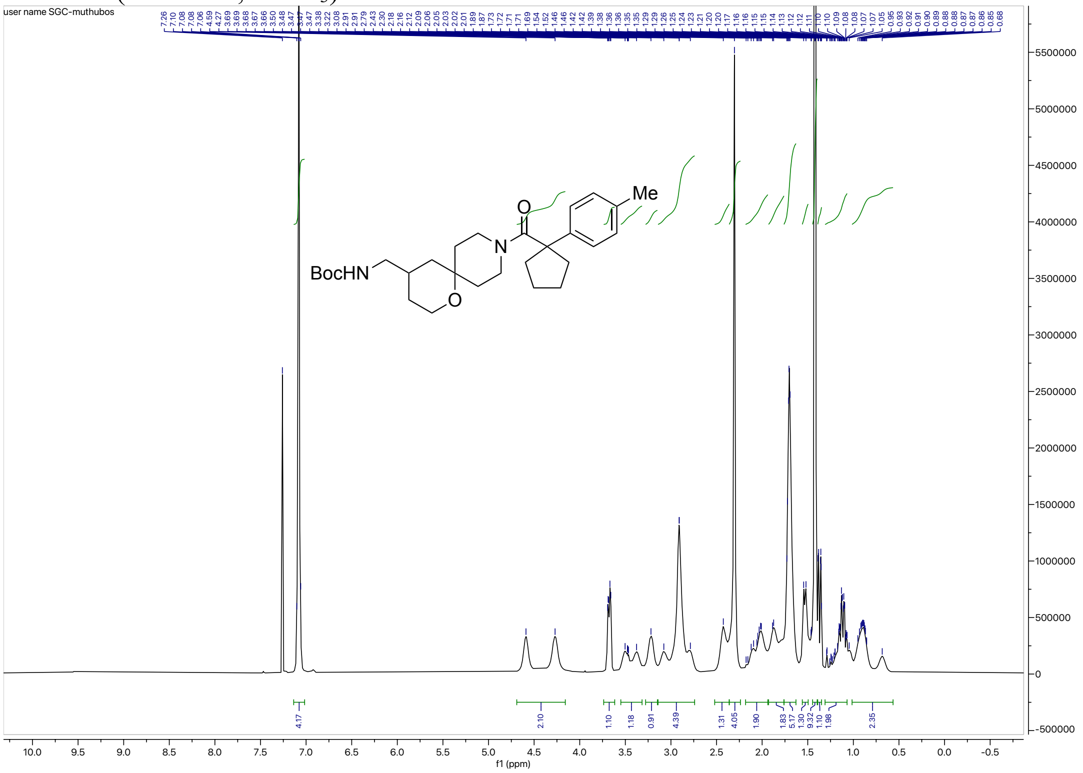

#### $^{1}$H NMR (500 MHz, $CDCl_3$) for **Xj**

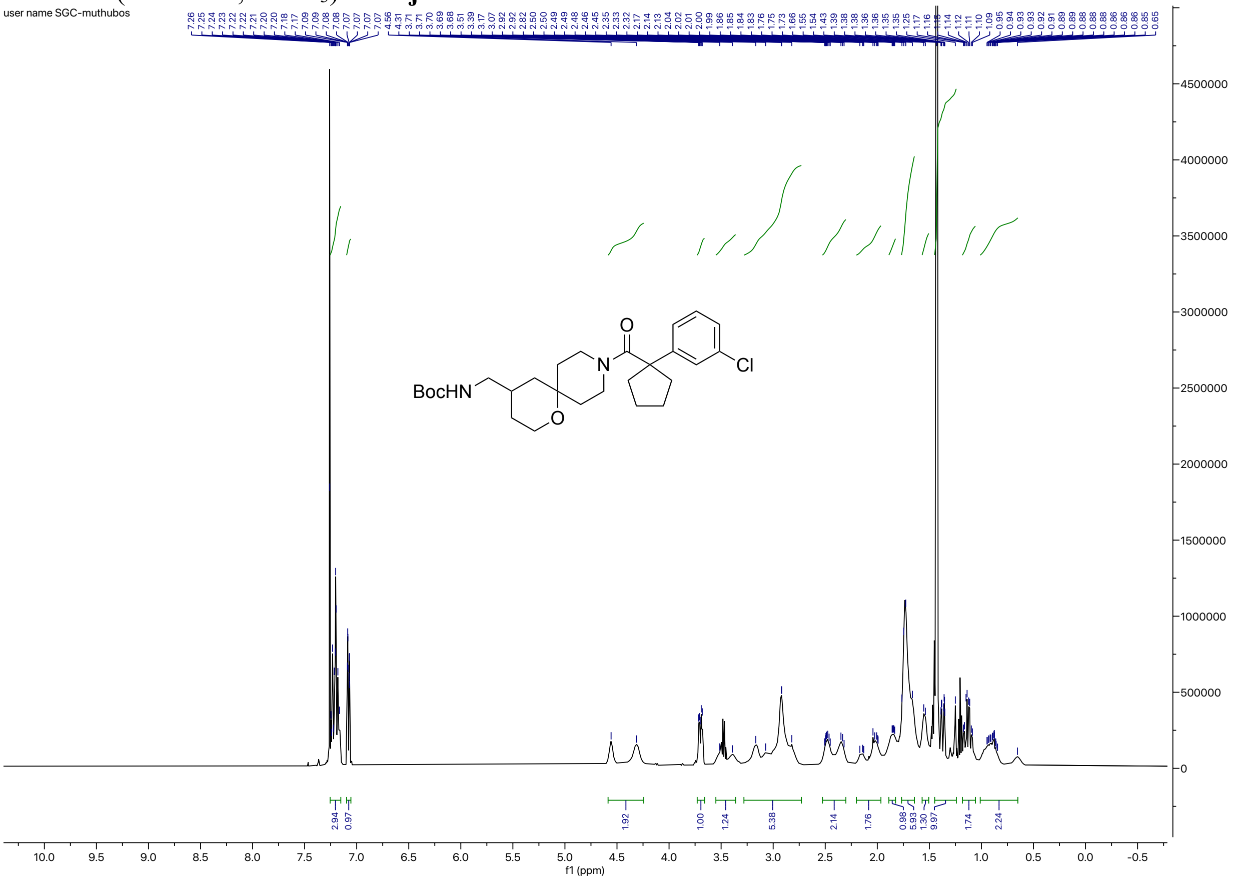


#### $^{1}$H NMR (400 MHz, $CDCl_3$) for **XI**

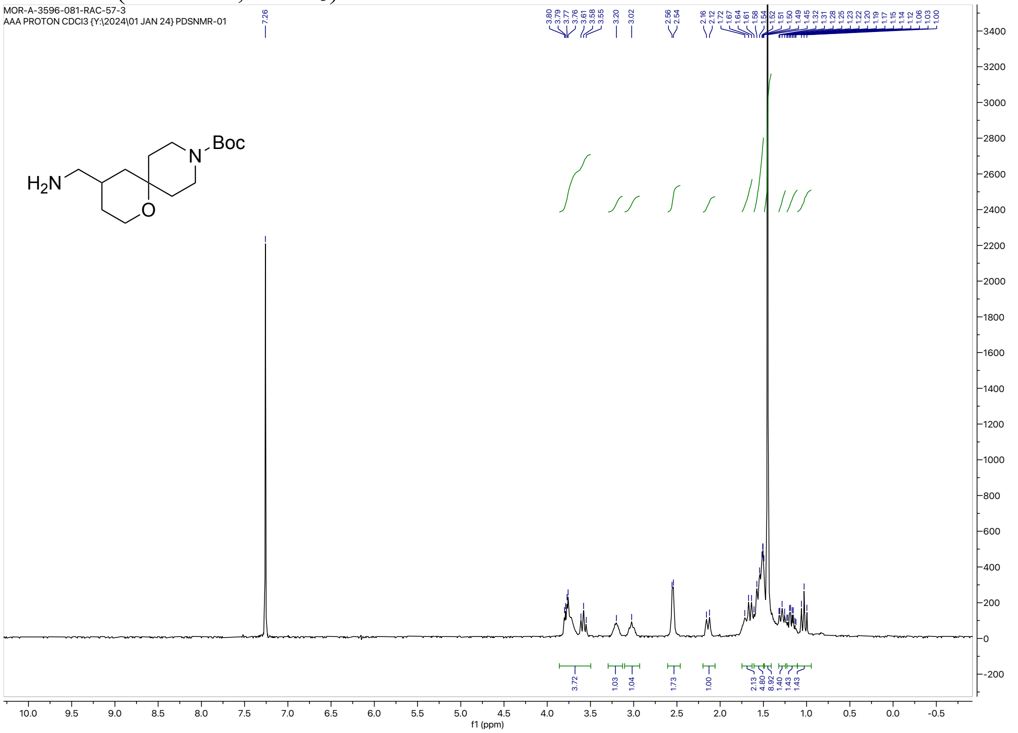

$^{1}$H NMR (500 MHz, $CDCl_3$) for **XIII**

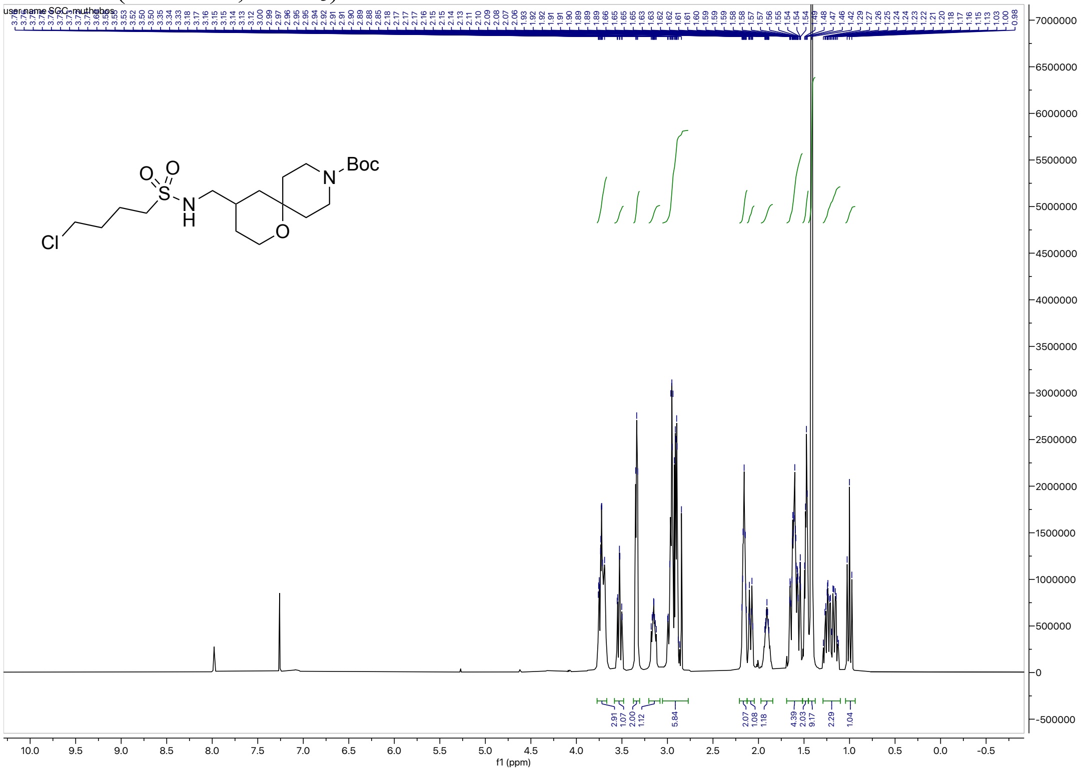

$^{1}$H NMR (500 MHz, $CDCl_3$) for **XIV**

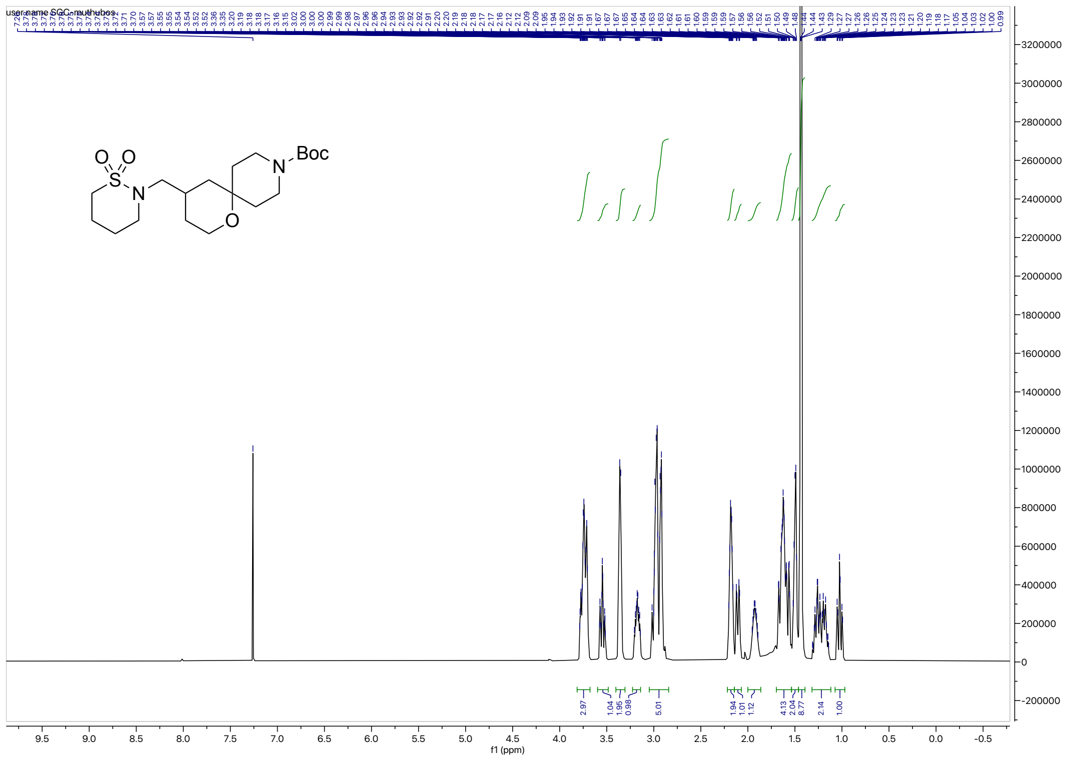

#### $^{1}H$ NMR (500 MHz, $CDCl_3$) for **XVa**

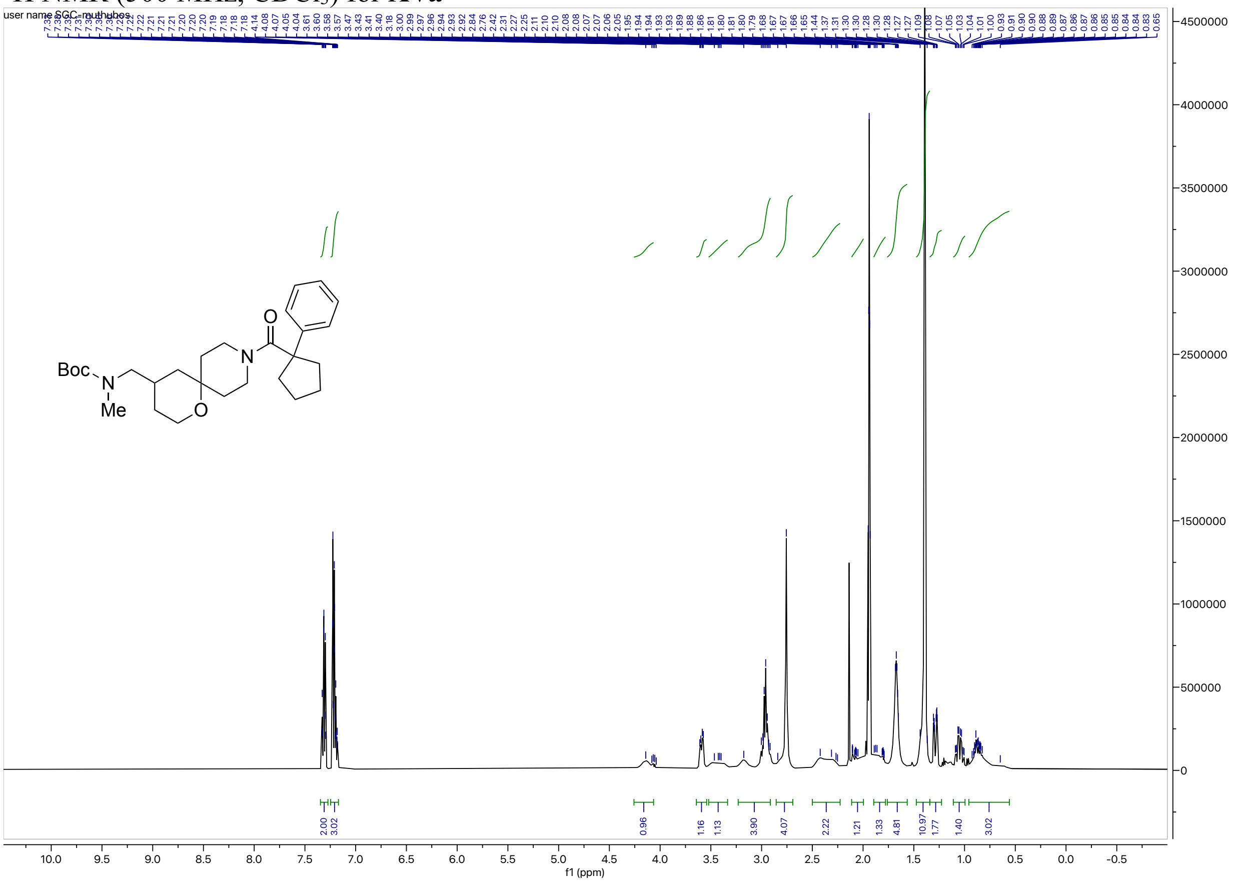

#### $^{1}H$ NMR (500 MHz, $CDCl_3$) for **XVb**

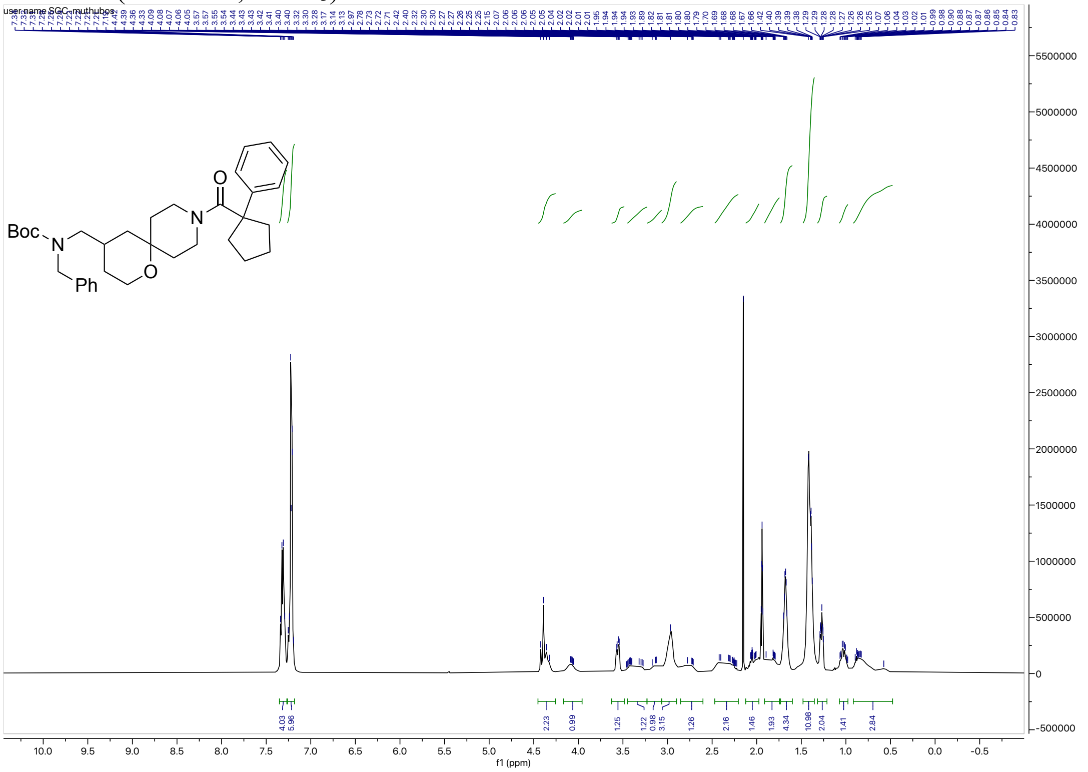
